# GEMOT: Towards Mechanistic World Models for Biology

**DOI:** 10.64898/2026.09.26.754619

**Authors:** Theodore de Pomereu, Branwen Snelling, Fabian Fröhlich

## Abstract

Scientific discovery seeks mechanisms that explain observations and predict beyond the measurements that produced them. Whereas large language models (LLMs) encode knowledge implicitly, mechanistic world models organise it as a parsimonious set of explicit, modular, reusable mechanisms whose predictions can be scored against data. In biology, where measurements are sparse and noisy, mechanistic world modelling must discover latent states and governing equations jointly, yet the prior knowledge that could constrain this search is largely unstructured. We introduce `gemot`, an agentic framework for mechanistic world modelling, and evaluate it on 18 published biological problems spanning molecular biology, epidemiology, and immune-cell differentiation, with up to 1,755 training measurements, 65 observables, 170 experimental conditions, and 3 data modalities per problem. `gemot` is auditable by construction: a semantic layer records each hypothesis, and a Model Context Protocol layer separates hypothesis formulation from numerical evaluation, so hypothesis scoring cannot be fabricated. On every problem, autonomously constructed models match or exceed the reference models in fit and parsimony, and generalise where held-outs exist. We demonstrate that `gemot` can formulate novel, biologically plausible mechanistic hypotheses. This work will enable decoding of interventional biological data into competing mechanistic hypotheses and design experiments that distinguish between them, and may serve as a blueprint for mechanistic world modelling in other domains.

## 1 Introduction

### Mechanisms and models

A mechanism consists of entities structured by causal interactions, working in concert to generate a system’s behaviour. A mechanistic model instantiates these mechanisms in simulatable form, enabling quantitative comparison against experimental and counterfactual predictions under interventions. This naturally connects to concepts developed in the philosophy of science such as *causal models* (Woodward, 2003; Pearl, 2009) and *real patterns* (Dennett, 1991; Ladyman, 2026). Among many formalisms for mechanistic models, Ordinary Differential Equations (ODE) models enjoy wide popularity across fields, and with novel symbolic regression (Cranmer, 2023; Mangan et al., 2016) and distillation (Cranmer et al., 2020) approaches it is now becoming possible to directly discover corresponding governing equations *de novo* from data.

### Biological underdetermination

In biology, mechanistic models link hypotheses to measurements across scales, from molecular networks and interacting cells to tissues and populations. Models have helped relate local regulatory and cell–cell interactions to tissue organisation and generate testable predictions for developmental experiments (Tomlin & Axelrod, 2007). At the population scale, models of transmission and immunity connect individual processes to epidemic dynamics and inform disease control (Keeling & Rohani, 2008). Causal explanation, however, requires experimental and intervention evidence beyond reproducing an observed outcome (DiFrisco & Jaeger, 2020; Briscoe & Kicheva, 2017). In periodic patterning, for example, different molecular, cellular, and mechanical processes can yield similar patterns, so agreement with a reaction–diffusion model does not establish that mechanism (Hiscock & Megason, 2015). The same underdetermination holds at the level of the equations themselves, where distinct model topologies can be indistinguishable (Szederkényi et al., 2011). This makes *de novo* mechanism discovery in biology inherently prone to underspecification.

### Triangulation and parsimony

The practical response of experimental biology is triangulation: combining complementary data modalities to constrain a shared mechanistic hypothesis (Ladyman, 2025, Section 4). ODE models recapitulate triangulation, as governing equations describe the temporal evolution of latent variables that are shared across experiments and contextualised by the interventions applied in each. Although triangulation narrows the space of admissible hypotheses, several hypotheses may survive, where the canonical statistical recourse is Occam’s razor, favouring the simplest model consistent with the evidence (Forster & Sober, 1994), where the Akaike and Bayesian Information Criteria (AIC, BIC) are common complexity criteria. Crick (1988, p. 138), however, cautioned that evolution adds mechanisms on top of existing mechanisms, so whether arguments about simplicity are useful for establishing mechanistic hypothesis remains debated.

### Mechanistic world modelling

Mechanistic world modelling, proposed by Posner, Lei, and Schölkopf (2026), formalises the process of mechanistic model formulation and casts it as knowledge organisation: mechanistic models explain a particular system via a structure of *bindings* that instantiate *mechanisms* over latent *variables*. Scientific discovery is then the joint recovery of these three elements, with parsimony and compositionality as inductive pressures, and where experimental verification and knowledge recombination implement active inquiry. As we outlined above, systems biology has already established counterparts to most of these components. State variables of ODE models correspond to latent variables, and the functional forms of interactions between them to mechanisms. Moreover, frameworks such as BioPAX and INDRA represent biological statements semantically and assemble them into executable models (Demir et al., 2010; Gyori et al., 2017). The binding structure can be encoded in machine-readable formats like SBML (Hucka et al., 2003) and human-readable front-ends such as Antimony (Smith et al., 2009), with PEtab (Schmiester et al., 2021) binding latent states to measurements collected under different interventions and assays, enabling the quantitative scoring of agreement of models with data. What systems biology lacks is the operationalisation of tacit judgement required to assemble these components: which latent states to posit, which mechanisms to bind to them, and what to simplify or omit. This is the variable, mechanism and structure discovery of a mechanistic world model. While some operationalisation has been attempted in textbooks (Klipp et al., 2016; Alon, 2019), most practitioners regard modelling as an art that draws on unstructured literature, expert intuition and tribal folklore.

### Our Contribution

Here, we introduce `gemot`, an agentic system that operationalises the variable, mechanism and structure discovery of mechanistic world models, using large language models (LLMs) to turn biological knowledge and experimental data into scorable hypotheses. Agents jointly construct and revise biological mechanisms and readout mappings, choosing latent states and linking them to measurements, so that one hypothesis can be triangulated against many assays and modalities. An agentic judgement layer retrieves knowledge and assembles hypotheses into models and a model context protocol (MCP) layer insulates numerical scoring, ensuring auditability and preventing fabrication. `gemot` does not resolve underdetermination; it makes alternatives cheap, producing sets of admissible, auditable mechanistic hypotheses that a scientist can adjudicate and discriminate between with new experiments. We scale evaluation to 18 published biological problems from the PEtab Benchmark Models collection (Hass et al., 2019; Grein et al., 2026), comprising 6,638 training measurements across 11 measurement modalities. Agents receive experimental measurements, curated assay and experimental information and knowledge priors, while direct access to the published model and narrative study context is withheld.

### Relation to prior work

Relative to semantic regularisation approaches (Kacprzyk & van der Schaar, 2025; Kacprzyk et al., 2025), our semantic representation describes the biology encoded in mechanistic hypothesis rather than qualitative behaviour of the dynamic systems, enabling validation of predictions on held-out interventional experimental data. Within LLM-based symbolic regression and model discovery (Shojaee et al., 2025a; Ziaei Bideh & Gryak, 2026; Song et al., 2026; Krongauz et al., 2026; Abhyankar et al., 2026), our focus is the joint inference of mechanism and measurement interpretation, which enables consideration of multi-modal data and triangulation based hypothesis filtering. Building on empirical discovery precedents (Holt et al., 2024), we scale from individual case studies to heterogeneous biological datasets requiring shared latent mechanisms to reconcile many observables, interventions and measurement modalities. Compared with synthetic equation- and model-discovery evaluations (d’Ascoli et al., 2024; Ziaei Bideh et al., 2026; Shojaee et al., 2025b; Murphy, 2026) and interactive-discovery benchmarks (Wiemann et al., 2026; Kabra et al., 2026; Gandhi et al., 2025; Zheng et al., 2026; Cerrato et al., 2026; Duan et al., 2025), we evaluate `gemot` on real data, enabling us to assess the relevance of prior knowledge.

## 2 Results

### 2.1 GEMOT: A layered architecture for trustworthy agentic discovery

As LLMs become more capable, auditability by humans is likely going to be one of the main measures of success for any agentic scientific discovery framework. We address this need architecturally (Figure 1) by separating the agentic *judgement layer*, which retrieves knowledge, analyses model fit, and proposes new hypotheses, from the numerical *evaluation layer*, which trains the model on experimental data, evaluates parsimony and compares simulations with data. The two layers are decoupled by a *semantic layer*, which instantiates hypotheses as human readable models, and an MCP *insulation layer*, which passes the formulated model from *judgement* to *evaluation* layer and makes data and model diagnostics query-able in the *judgement layer*. Together, these layers implement two separable design decisions: a trust boundary, across which the agent can submit models and query data and diagnostics but cannot alter data, fits or scores (insulation and evaluation layers); and an intermediate representation, in which each hypothesis is a versioned, human-readable model that can be inspected apart from the agent’s narration of it (semantic layer). Hypotheses and corresponding model instantiations are constructed through iterative loops between *judgement* and *evaluation layers*, where model–data mismatch and other diagnostics inform modelling decisions while BIC serves as the quantitative objective that the LLM is instructed to optimise.

**Figure 1:**
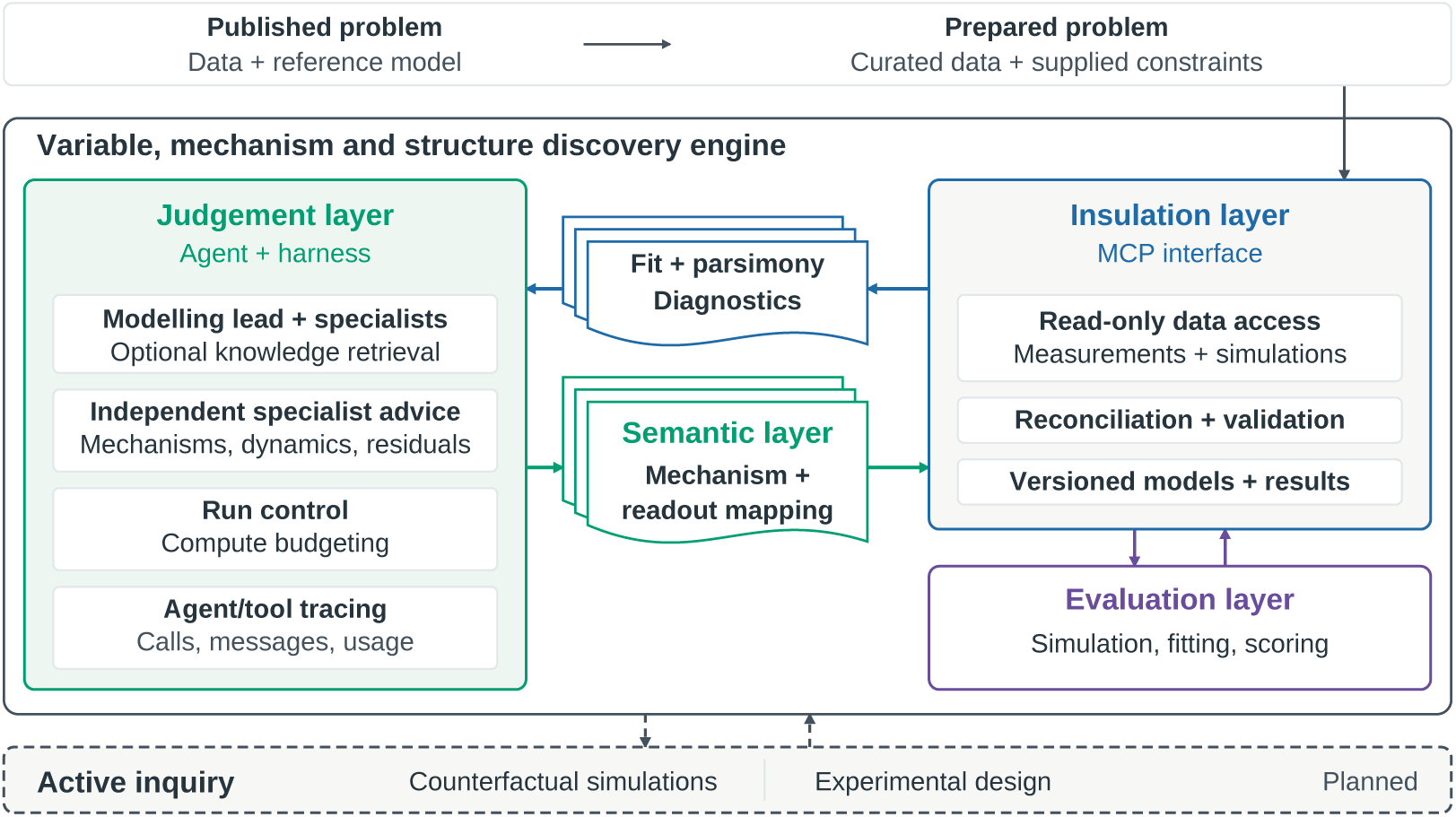
The variable, mechanism and structure discovery engine. Reference models and heldout data are withheld during model construction; feedback uses training data only. The MCP interface protects supplied constraints and preserves model versions and numerical results independently of agent traces. Dashed elements denote planned capabilities. Implementation details are in Appendices A–C.

The ***judgement layer*** is an agent harness built with Deep Agents (LangChain, Inc., 2025) that supplies tools and computational budgets; a modelling lead coordinates mathematical, biological, and signal-analysis specialist subagents. Harness-level tracing records agent interactions and tool calls. Prompts specify a build–fit–diagnose loop with independent specialist advice (Listings 114–119), using benchmark-specific tasks, supplied constraints and entity lists (Appendix I.5). The optional biocurator subagent has access to UniProt (The UniProt Consortium, 2025), Reactome (Milacic et al., 2024), and OmniPath (Türei et al., 2026).

The ***evaluation layer*** is a functional layer controlled by the MCP server that implements model training through established systems biology tools: repeated local, gradient based optimisation from different initial parameter vectors (Grein et al., 2026). Simulation and gradient computation use `AMICI` (Fröhlich et al., 2021), likelihood-based parameter estimation uses `pyPESTO` (Schälte et al., 2023) with trust-region optimisation in `fides` (Fröhlich & Sorger, 2022).

The ***semantic layer*** instantiates sets of hypotheses as human readable models in the Antimony (Smith et al., 2009) format that defines proposed states, reactions, rate laws, and readout mappings linking latent states to measured quantities (Appendix A.3).

The ***insulation layer*** is implemented as MCP server providing read-only access through DuckDB (Raasveldt & Mühleisen, 2019) to training measurements, simulations, raw and noise-normalised residuals, and noise scales by observable, condition, time, and model version. Separately from agent traces, the server preserves model versions, fitted parameters, diagnostics, and justifications. The layer also computes diagnostic information like residual bias, temporal structure, worst-fitting points, parameters at bounds, and local parameter uncertainties and correlations from the Fisher information matrix. Lastly the layer translates human readable models to SBML (Hucka et al., 2003), reconciles and validates them against PEtab’s experimental and observation specification while protecting supplied constraints, and passes the full problem to the evaluation layer.

The insulation layer is also responsible for imposing deterministic safeguards. For example, despite explicit instructions, agents hardcoded numeric literals (unnamed fixed constants) in 12/90 GLM models across 11/18 benchmarks (Figure 2; Section 2.2), and in 0/5 Astra models with `gemot` versus 5/5 without (Figure 3; Section 2.4). Although constants can encode unit normalisation or cooperativity, unjustified fixed values can escape BIC penalties and understate uncertainty. During reconciliation, `gemot` therefore counts literals towards complexity penalties and reports them to the agent; automatic promotion to fitted parameters remains under consideration. This illustrates how deterministic safeguards support reliable model construction and reduce the need for human audit.

**Figure 2:**
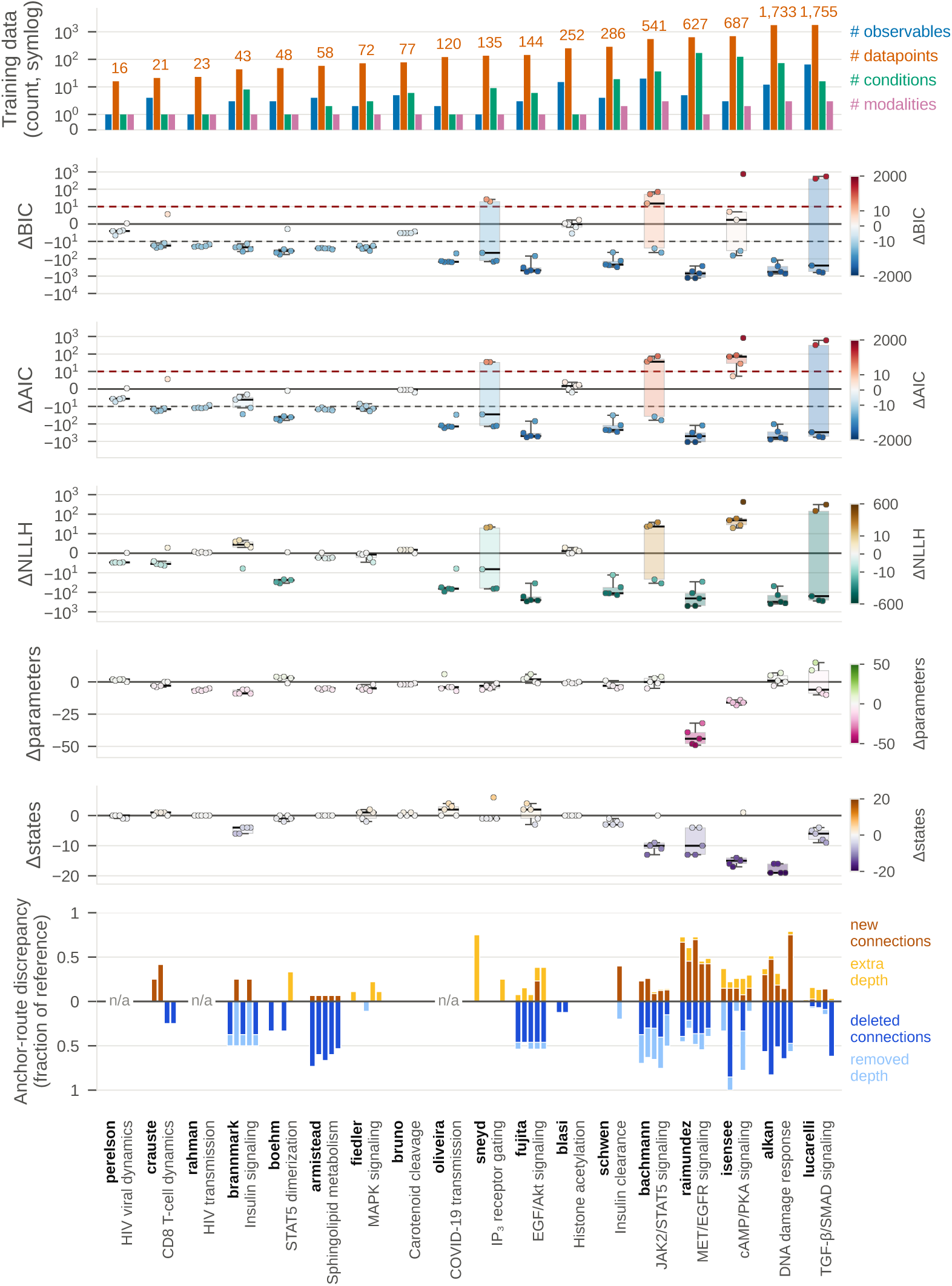
Training performance, model size, and structural differences across 18 published problems. Columns are ordered by training measurement count. Points show individual runs; boxplots summarise available values. ΔBIC and ΔAIC are candidate minus reference; ΔNLLH denotes the training negative-log-likelihood difference. Bottom-row stacked bars show each candidate’s normalised anchor-route discrepancy: new connections (brown) and extra depth on shared routes (yellow) above zero; deleted connections (dark blue) and removed depth on shared routes (light blue) below. Normalisation uses the reference route summary, not the full network’s edge count (Appendix D.3).

**Figure 3:**
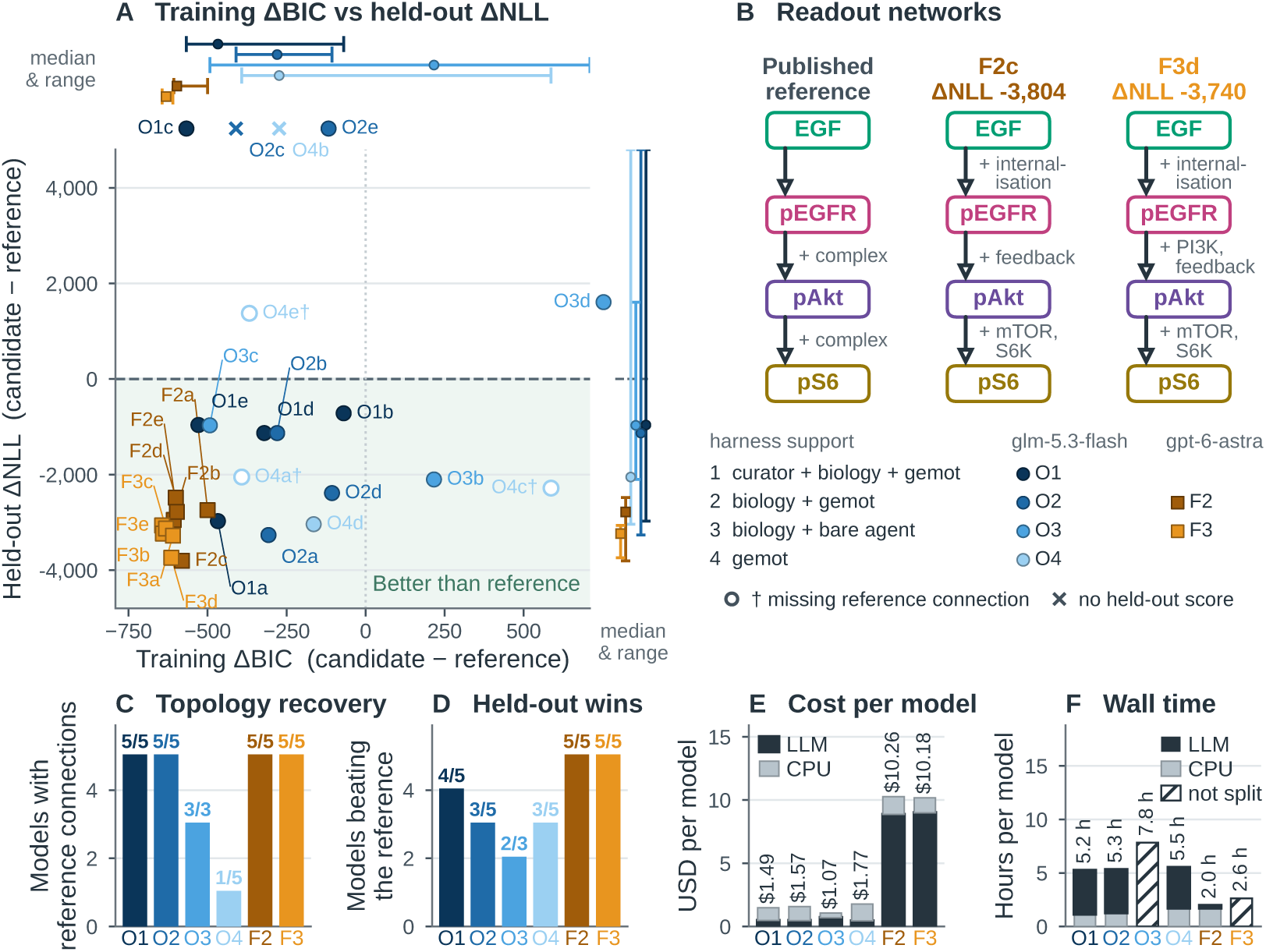
Agent configurations, model structure, prediction, and cost. **A,** The 28 Fujita runs that built a model (the two that built none are omitted), with linear training ΔBIC and held-out ΔNLL relative to the reference. Marginals show medians and ranges, a failed held-out integration ranking worst; off-scale values and missing scores are indicated separately; daggers in A mark models lacking a required reference connection. **B,** Readout networks of the reference and the two best held-out candidates: the EGF input and the three readouts, linked by the forward readout connections counted in C; side notes name what each model places on a connection (full maps in Figure 29). **C,** Recovery of unsigned reference connections among built models; Table 9 gives the models each configuration built. **D,** Candidates with held-out ΔNLL *<* 0, by configuration, over every delivered model; a model whose held-out integration failed counts as not beating the reference. **E,** Cost per delivered model charges all launched runs: adjusted LLM usage plus a CPU list-price estimate. **F,** Wall time per model, split between LLM and numerical work where available; hatched bars show unsplit totals. Configurations, evaluation protocols, controls, full run mappings, and cost assumptions are in Appendix D.5.

### 2.2 GEMOT MATCHES OR EXCEEDS PUBLISHED MODELS IN FIT AND PARSIMONY

We first tested whether `gemot` could construct parsimonious mechanistic models competitive with human-built references. We selected 18 problems from the PEtab Benchmark Models collection (Hass et al., 2019; Grein et al., 2026): 13 smaller problems, followed by five larger problems with more training measurements and generally more distinct interventions (conditions), observables (measured signals), and data modalities. The selection spans molecular biology, interacting cell populations, and epidemiology. The training datasets span 1–65 observables, 16–1,755 datapoints, 1–170 conditions, and 1–3 measurement modalities per problem (Figure 2, row 1; Table 1). Modalities comprise immunoblotting, mass spectrometry, qRT-PCR, ELISA, flow cytometry, HPLC, patchclamp recordings, cell imaging, immunofluorescence microscopy, and viral RNA assays, alongside epidemiological surveillance. Agents receive measurements, experimental inputs, assay annotations, and curated biological constraints, but start from an empty model without the published mechanism or narrative study context (Appendix A.2).

We applied `gemot` using GLM-5.3-Flash as the underlying LLM, with 5 runs per problem (90 in total; model listings in Appendices G and H). Median usage per run was 2.0 million LLM tokens and 2.0 CPU-hours for smaller problems, versus 7.9 million tokens and 514 CPU-hours for larger problems (Appendix D.2), for a total of 400.3 million tokens and 17,230 CPU-hours. Overall, 78/90 models matched or improved on the reference by BIC (Schwarz, 1978), using the conventional ΔBIC *<* 2 threshold (candidate minus reference; (Kass & Raftery, 1995)). At least two runs achieved lower BIC than the reference in every problem, and lower AIC in every problem except Isensee (Figure 2, row 2/3). In 63/90 models (15/18 problems; exception Perelson, Bruno, and Blasi), runs scored ΔBIC *<* −10, conventionally indicating very strong evidence favouring the candidate (Kass & Raftery, 1995). 55/90 runs achieved a better fit to the data, indicated by a negative candidate-minus-reference training negative log-likelihood difference (ΔNLLH *<* 0) (Figure 2, row 4). In sum, `gemot` constructs models competitive with human references in fit and parsimony.

### 2.3 GEMOT PROPOSES NOVEL MECHANISTIC HYPOTHESIS

Published benchmarks may occur in LLM training data. We therefore compared candidate and reference structures to assess whether `gemot` proposes mechanisms beyond the published models.

Per-problem median differences ranged from −9 to +3 parameters (Figure 2, 5th row) and −4 to +2 states (Figure 2, 6th row) on the 13 smaller problems, versus −44 to +1 parameters and −19 to 6 states on the five larger problems, suggesting that parsimony gains are likely achieved through coarse-graining. To identify novel mechanisms, we computed model topology in terms of weighted graphs that quantify the latent depth—the shortest path along directly interacting state variables that are not part of the readout mapping— between interventions and readouts (anchors; Appendix D.3). Comparing these graphs to published models (SBGN Activity Flow maps in Appendix F), we found differences in 56/75 defined comparisons: 49 changed connectivity (brown/dark blue bars) and seven only route lengths (yellow/light blue bars; Figure 2, bottom row). All 25 larger-problem runs added connections, despite generally using fewer states. As extensive connection deletions and route shortening make their new connections and extra depth difficult to interpret, we concentrated on models with smaller anchor-route discrepancies:

#### Added connections

Two Crauste runs (Listings 35 and 36) added an early-effector-to-memory influence (brown) through pathogen clearance relieving suppression of memory differentiation, consistent with antigen-withdrawal experiments (Sarkar et al., 2008). One of these runs (Listing 36) also adds a direct differentiation step from naive to late effectors cells, for which we found neither strong supporting nor refuting evidence in the literature.

#### Latent intermediates

Longer routes (yellow) separated IP_3_-dependent priming from channel opening in Sneyd (Listing 78), introduced an unnamed dynamic mediator of MAPK feedback in Fiedler (Listing 63), and inserted mTORC1–S6K signalling between Akt and S6 in Fujita (Listing 4), consistent with literature (Rossi et al., 2009; Dougherty et al., 2005; Inoki et al., 2002) (Appendix D.3).

We discuss more findings in Appendix D.3. In summary, `gemot` thus deviates substantially from published models, proposing both novel hypotheses that are consistent with, and, therefore, likely informed by, prior literature, as well as likely data-driven hypotheses with little literature support.

### 2.4 Held-out interventions partly discriminate mechanisms

To evaluate whether `gemot` generated mechanistic hypotheses generalise beyond training data, we evaluate predictive performance on the Fujita EGF–Akt–S6 benchmark (Fujita et al., 2010), where held-out data was available from the PEtab Benchmark Collection. Models trained on 144 constant-EGF-dose measurements predicted 276 held-out EGF-pulse and EGF-ramp measurements with all fitted parameters fixed (Figures 7–12). We report ΔNLL as candidate minus reference where negative values favour the candidate. To identify what drives performance, we evaluated `gemot` in six configurations, crossing two LLM drivers—O (open weight, blue) for GLM-5.3-Flash and F (frontier, brown) for GPT-6-Astra—with an ablation ladder (shading) over capability: 1, the full `gemot`; 2, the `gemot` without the biocurator; 3, a barebones harness with sandboxed file access and packages pre-installed (prompts in Listings 120–123); and 4 `gemot` with biology blinded and biocurator removed (prompt in Listing 124; Figure 3). We evaluated O1–O4 and F2–F3 with five runs each, labelled a–e (protocol and model listings in Appendices D.5 and G). F1 and F4 were cost cut.

#### Parsimony and prediction

We generally observed poor correlation between BIC and held-out performance (GLM: -0.1, Astra: 0.3 Spearman’s *ρ*). For example, O1c achieved lower training ΔBIC than O1a (−567 versus −467), yet far worse held-out ΔNLL (+44 625 versus −2974). The two best-performing models (F2c ΔNLL −3804; F3d ΔNLL −3740), recapitulated the forward activation structure of the reference model, but gained predictive power in face of added complexity such as EGFR internalisation (F2c, F3d), integral negative feedback from pAKT onto activation by pEGFR (F2c, F3d), and latent intermediates such as PI3K (F3d), MTOR (F2c, F3d) and S6K (F2c, F3d) (Figure 3B; Figure 29). This complexity is affordable in training BIC through lower NLL and coarse-graining elsewhere, e.g. ommiting EGFR/AKT and AKT/S6 complexes or merging dephosphorylation rate constants. This suggests that BIC, while likely helpful in guiding model construction, cannot discriminate mechanistic hypotheses, echoing Crick’s caution about parsimony.

#### Implicit biological knowledge

To test whether model construction requires curated knowledge, we removed the biocurator and its external knowledge tools from the GLM panel (O2). All five models retained the reference’s forward readout connections, and 3/5 models outperformed the reference on held-out data, compared with 4/5 for the full panel (Figure 3A, shaded region; C, O1/2). Thus, GLM could construct predictive mechanisms without curated retrieval, consistent with sufficient implicit biological knowledge in this LLM.

#### Biological context

To test how important implicit biological knowledge is for generalisation, we blinded `gemot` by removing data annotation, removing the biocurator and obfuscating identifiers, including those of the stimulus protocol (O4). Blinding barely changed median training ΔBIC (280 versus 273; Figure 3A, top errorbars) and reduced recovery of the forward readout connections from 5/5 to 1/5 (Figure 3C; Figure 3A, daggers mark runs that did not recover topology), but did not impair generalisation: median held-out ΔNLL improved from −1132 to −2052 (Figure 3A, right errorbars), and 3/5 blinded models beat the reference (Figure 3A, pale-blue circles). Biological context thus shaped which mechanisms were built rather than how well they predicted.

#### Cost and reliability

Practical model construction requires reliability at an affordable cost. Both biology-visible `gemot` GLM arms (O1/2) delivered 5/5 models, compared to 3/5 for the bare agent (O3), while Astra maintained 5/5 models for the barebones harness (F3). However, GLM delivered models in 5-8 hours at a cost of less than $2 per model versus more than $10 in 2-3 hours for Astra (Figure 3E,F). For larger models, where model training time exceeds Astra’s 1h cache timeout, cost differences were more pronounced (data not shown). Astra nevertheless produced stronger predictions more consistently: all ten candidates (F2/3) beat the reference, versus 9/13 biology-visible GLM (O1–3) candidates (Figure 3A, brown/orange squares versus blue circles). The best GLM model achieved 86% of the best Astra model’s improvement over the reference (held-out ΔNLL −3265 versus −3804). Bare Astra also had lower median held-out ΔNLL than its panel (−3235 versus −2781). Thus, `gemot` supports reliable construction with cheaper LLMs, while the frontier model’s more consistent predictions came at higher cost.

As the forward topology implemented by Fujita implements textbook biology, we conclude that biological context is necessary to build plausible models, and that ambiguity between competing mechanistic hypotheses persists even with additional experimental validation.

### 2.5 Robustness to biology blinding extends beyond Fujita

We tested whether Fujita’s robustness to blinding extends to four further systems, comparing biology-visible (O1) and biology-blinded (O4) models (Figure 4). We reuse selected published validation tests for Raimundez and Crauste, but not Crauste’s forecasting prediction tests. Perelson and Lucarelli use adapted held-out protocols (Raimúndez et al., 2020; Crauste et al., 2017; Perelson et al., 1996; Lucarelli et al., 2018). For Lucarelli, we refit all estimated parameters in both candidates and reference to pooled calibration and validation measurements; the original study kept gene-regulatory parameters fixed during transfer, which is not applicable to with blinded biology (Appendix D.6).

**Figure 4:**
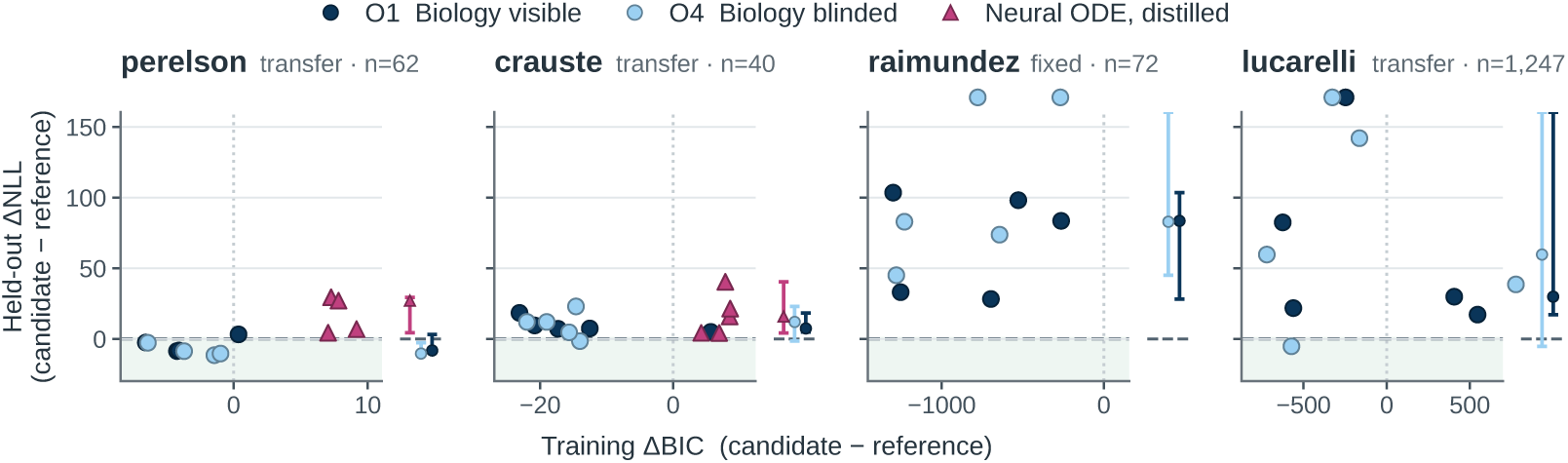
Training fit against held-out prediction on four further systems. Each panel plots training ΔBIC against held-out ΔNLL, both candidate minus reference, so the shaded region below zero is better than the reference on held-out data. Panels are ordered by training measurement count; *n* is the number of held-out measurements each ΔNLL is computed on. Crauste, Lucarelli, and Perelson are transfer tests, with candidate and reference parameters refitted per held-out unit; Raimundez holds dynamical parameters fixed while calibrating new per-blot observation scales. Circles are the Figure 2 runs (biology visible) and their biology-blinded counterparts; triangles are neural-ODE models distilled into rate laws (Crauste and Perelson only). Marginals show median and range per arm; runs beyond a panel’s scale are drawn outside its frame, above it for held-out ΔNLL and past its right edge for training ΔBIC.

#### Blinding and neural baselines

Blinding did not consistently worsen held-out ΔNLL: O1 and O4 largely overlapped, although blinded runs had larger errors in some Raimundez and Lucarelli cases (dark and light blue circles; Appendix D.6). Distilled neural ODEs (Appendix D.7) performed comparably on Crauste and Perelson, with slightly higher median ΔBIC in both systems (magenta triangles). On Fujita, neural ODE integration produced unphysical simulations and failed numerically, preventing evaluation; training and distillation was not successful on Raimundez and Lucarelli.

#### Implications for model discovery

Competitive performance under blinding supports `gemot`’s ability to construct useful models with little to no cues for potential recalling of published models. The transfer examples assess model reuse across contexts rather than prediction under novel interventions, which may partly explain why these held-outs appear less discriminating than Fujita’s fixed-parameter pulse/ramp evaluation. Biological context was therefore not needed for competitive generalisation on any system tested, pointing to a need for better experimental design.

## 3 Discussion

Across 18 biological problems, `gemot` constructed models competitive with human references in fit and parsimony, and in doing so generated novel hypotheses. Variable, mechanism and structure discovery, the part of mechanistic world modelling that systems biology has left informal, can thus be operationalised with LLM agents. Across five benchmarks with held-out data, blinded runs generalised competitively; on Fujita, blinding changed biological plausibility but not how well models predicted. `gemot` thus leaves mechanistic underdetermination to experiment and human judgement, but makes alternative hypotheses explicit and cheap to generate.

### Towards mechanistic world models

A mechanistic world model is maintained over successive experiments. After each, variables and bindings are revised, new mechanisms are added to a library, and the next intervention is chosen to discriminate between the remaining hypotheses (Posner et al., 2026). Our benchmarks provide fixed data and therefore only test the first step, which `gemot` implements as variable, mechanism and structure discovery. Furthermore, `gemot` draws mechanisms from the implicit knowledge of the LLM and does not represent them separately from their bindings, so they cannot enter a library. Structured prior knowledge could make these mechanisms explicit, as BioPAX (Demir et al., 2010) and INDRA (Gyori et al., 2017) already do for molecular systems. The models `gemot` constructs often also use coarser mechanisms than the references, which brings a general difficulty to light. Biology is multiscale, and while Posner et al. (2026) call for hierarchically organised mechanisms, they do not specify how mechanisms at different levels relate. INDRA partly addresses this (Bachman et al., 2023), but only with predefined levels for individual interactions, fixed in advance rather than discovered. Symbolic regression could complement it by discovering missing coarse-grained mechanisms from data (de Pomereu & Fröhlich, 2026). `gemot` also does not yet implement experimental design, which could be implemented by proposing interventions or measurements of latent states that maximally distinguish model candidates.

### Trustworthiness by design

We expect future LLM generations to increase in capability and costeffectiveness, rendering some of `gemot`’s components and observed capability augmentations obsolete. We therefore see `gemot`’s contribution mainly in its trustworthy architecture that make numerical claims easy to audit. Through the configured interface, the MCP layer prevents agents from replacing protected evaluation data or scores, and the judge uses these records to assess success (Appendix B.4). For mathematical modellers, this separation between judgement, semantics, and scoring enables empirical evaluation of modelling strategies.

### Limits of published benchmarks

Evaluation on public data allows us to compare candidate models against a human reference, but withholding published reference models and validation data cannot exclude exposure during LLM training. As we only found current-generation LLMs capable enough (data not shown), training cutoff was not a viable mitigation strategy. Structural departures from the references demonstrate that `gemot` synthesises novel mechanistic hypotheses, which we, based on results for biology-blinded problems, expect to generalise to unseen problems (Section 2.3). The held-out evaluation is moreover unannounced to the LLM and supplies no feedback during construction. Even if a LLM could anticipate this evaluation and recall the held-out measurements, it would still have to translate them into an executable mechanism and readout mapping that can predict recalled measurements from training data. Verbatim reproductions of the reference model or recall of data cannot explain better performance on training data alone, and improvement through knowledge recombination would fulfill a stated design goal of mechanistic world models.

### Complexity measures

Our study employed BIC as the only complexity criterion to guide model construction, with a parameter-count penalty that grows with sample size (Schwarz, 1978). `gemot` also supports AIC (Akaike, 1974), AICc (Hurvich & Tsai, 1989), CAIC (Bozdogan, 1987), HQC (Hannan & Quinn, 1979), and a Fisher-information approximation to TIC (Takeuchi, 1976) for online scoring (Appendix B.7). Our preliminary evaluation suggests that the choice among these criteria has no major impact on constructed models (data not shown). Predictive criteria such as WAIC (Watanabe, 2010) and Bayesian leave-one-out cross-validation (Vehtari et al., 2017) may better reflect generalisation, although their value for predicting new interventions remains to be established. Obtaining posterior samples for each candidate, with additional refitting where needed, would add substantial cost to our optimisation-based workflow and was beyond the scope of this evaluation. We anticipate that `gemot` will enable the development and empirical evaluation of better complexity measures in the future.

## Reproducibility Statement

Code is not released with this preprint; we will release all code upon publication. In its place, the Appendix provides a human-curated, LLM-narrated summary of the code together with the artefacts it generated. We document code and prompt versions throughout, but exact reproduction remains out of reach: the required compute and token budgets are high, and responses from external language models cannot be reproduced exactly. Because we observed large run-to-run variability, we performed five repeat runs for every configuration and refrained from interpreting small-to-medium performance or pass rate differences.

## AI Use Statement

Generative AI was used both in the evaluated system and in preparing this work.

### Research ideation and execution

In the reported experiments, GLM-5.3-Flash and GPT-6-Astra agents developed mechanistic models, proposed and refined biological hypotheses, implemented these hypotheses as executable models, and interpreted fit diagnostics. Their configurations and evaluation protocols are described in Appendices C.6 and D.

During preparation of the research, OpenAI Codex and Anthropic Claude helped develop and refine conceptual frameworks, implement methods and analysis code, and interpret results. AI assistance also supported qualitative and thematic analysis and critique of research methods and experiments, including ablations, statistical tests, and held-out evaluation protocols. Anthropic Claude was used for code writing, spot-checks of agent traces, and diagnosing failed runs from them, and was the sole AI assistant used for data curation: extracting measurements, cleaning and reformatting datasets, and checking units and metadata. Claude also implemented, ran, and checked analysis and figure code on the computing cluster, including the held-out and transfer rescoring and the refits of the rebuilt distilled models, and generated the manuscript’s reported values from the frozen inputs as LaTeX macros checked against the data. Codex and Claude assisted with preparing scientific figures and research artifacts.

### Retrieval and discovery

Codex and Claude assisted with searching for, identifying, and analysing relevant literature.

### Drafting

Codex helped draft sections of the paper. Claude drafted figure captions and appendix passages describing its analyses.

### Writing and polishing

Codex helped edit and polish the manuscript for clarity, concision, and flow. Claude audited the manuscript for outdated values, statements the data no longer supported, and inconsistent notation, and applied the corrections the authors directed.

AI was not used for benchmark selection, synthetic dataset generation, formulation or proof of mathematical claims, or translation.

AI assistance identified several issues in human-curated benchmarks. Code was checked by at least one author, and held-out data curation was checked by two authors. Numerical claims are supported by archived computational outputs and statistical analyses. The human authors directed the work and retain responsibility for its scientific claims, references, code, and final text, including all AI-assisted contributions.

## Acknowledgements

We thank the whole Dynamics of Living Systems lab for helpful discussions and feedback on the manuscript. We also thank James Briscoe, James DiFrisco and Mihaela van der Schaar for helpful discussions and feedback on the manuscript, and the Hasenauer lab for helpful discussions.

## A Benchmark construction and reference problems

The reported experiments span multiple software revisions due to time and computational cost constraints. Unless specified otherwise, differences between the revisions used for these experiments did not materially affect model construction, evaluation, or the conclusions reported here. We will rerun all experiments using a single software revision upon publication.

This appendix documents benchmark preparation in the `petab-mcp` source repository at revision `e0d6921`, with input-protocol blinding documented at `7d22712`. It describes the available implementation, rather than asserting that every historical run used that revision. The current generated package, imported at manuscript revision `b4209e6`, carries producer stamp `09af5ef+dirty`; the Figure 3 score export retains `604449e+dirty`, while the refreshed training sweep records `d958571+dirty` and its constituent run revisions. Appendix D.1 separates these evidence sources. These identifiers distinguish implementation documentation, generated summaries, and analysis provenance.

### A.1 Collection and materialisation

The empirical suite comprises 18 problems selected from the Benchmark-Models-PEtab collection (Hass et al., 2019). We initially selected 13 smaller problems, then added five larger problems with more training measurements and generally more conditions and observables: Isensee, Alkan, Bachmann, Lucarelli, and Raimundez. The authoritative roster is defined in `src/petab_harnes s/paths.py`. It spans biochemical networks, cell-population dynamics, within-host infection, and epidemic transmission. The biological problems include dynamic and steady-state measurements, so constructing an ODE model does not imply that every dataset directly observes a trajectory.

Preparation is performed by `examples/setup.py`, which reads each example’s `run.py` configuration and calls the materialisation functions in `src/petab_harness/benchmark.py`. The default collection revision is `1763dfcd66d25bfd97492cfedf6a730472c383d0`. An existing collection can instead be supplied through `BENCHMARK_MODELS_PETAB`; this override uses that checkout as provided, so its revision must be recorded separately. Ordinary materialisation performs no parameter fitting and leaves the committed reference summary unchanged.

Each problem is written as PEtab version 1 tables and a YAML specification (Schmiester et al., 2021), accompanied by an empty SBML model (Hucka et al., 2003) and an empty Antimony representation. PEtab version 2 and PEtab SciML extend the broader modelling ecosystem (Pathirana et al., 2026; Persson et al., 2026); the materialisation path described here uses version 1. The model identifier is a neutral placeholder. Measurements are retained, while identifiers and table representations undergo configured relabelling, observable merging, and observable-formula blinding (below; distinct from the biology blinding of O4). This is consequently a curated transformation of a published estimation problem. Compact observable tables are retained during preparation; timepoint-specific observable and noise parameter overrides are expanded only in a simulation-facing copy when required by AMICI.

### A.2 Information supplied to the agent

Agents receive empirical measurements, their observable and condition identifiers, assay annotations, prescribed experimental inputs, and curated parameter information. The published dynamical model and narrative study context are withheld. Source-bearing identifiers are relabelled where configured, and free-text condition names are removed. The agent must construct a candidate mechanism and the mappings connecting its states to measured quantities. The measurement, condition, and observable tables provide the fixed experimental specification within which that construction takes place. With assistance from Anthropic Claude, we spot-checked agent traces to verify that the configured information-access restrictions operated as intended (Appendix B.4).

The retained information can include substantive biological priors. Per-example setup notes (Appendix I.5) specify known totals, stimulus rules, initial populations, or qualitative constraints that cannot be recovered from the supplied measurement rows alone. For example, the Crauste setup (prompt in Listing 148) identifies an unobserved pathogen and describes its growth, clearance, and role in T-cell activation. Such supplied constraints delimit the model-construction task; they must be considered when interpreting which components an agent independently proposes. Withholding the published model does not establish that its source is unrecognisable from the remaining biological entities or observations.

These statements describe the configuration retaining biological annotations. The current biology-blinding implementation removes biological names, descriptive metadata and mechanistic coupling guidance, while preserving prescribed time-dependent inputs as anonymised numerical rules. The agent is instructed to implement these rules and use their defined stimuli in its equations (Appendix D.5).

Where a held-out measurement table is configured, preparation stores it outside the agent-facing problem directory. The implementation checks compatibility of the observable, condition, and parameter tables and rejects overlapping observable–condition–time tuples between training and heldout measurements. This allows later timepoints of an existing condition to be withheld. The documented preparation path also rejects held-out problems requiring incompatible timepoint-specific override expansion.

### A.3 Observation mappings, parameters, and initial conditions

For a candidate model *m*, PEtab couples a dynamical system to an experimental and measurement specification (Schmiester et al., 2021). Let *c* identify a simulation condition together with its optional pre-equilibration condition. The model states satisfy

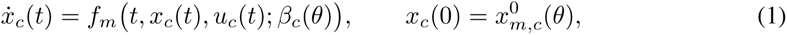

where *u_c_*(*t*) contains prescribed inputs and *θ* ∈ Θ*_m_* contains estimated parameters. The map *β_c_* combines these parameters with fixed values and condition-specific overrides; an override can be a supplied number or a reference to an estimated parameter. Each measurement row *i* specifies a value *y_i_*, time *t_i_*, observable *o_i_*, and condition pair *c_i_*. Replicates remain separate rows. A steady-state measurement uses the simulated equilibrium in place of 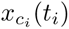.

Published observable formulas can reveal species names, complex composition, stoichiometry, or combinations of latent states. Preparation replaces this mechanistic content with `readout_*` placeholders that the agent must define through readout rules (model assignment rules); a place-holder is named `readout_<observableId>` unless the problem shares one readout between several observables. Assay descriptions retain selectivity, such as whether a measurement detects total protein or a phosphorylated form. This information constrains the interpretation of the readout without providing the reference model’s state decomposition.

The separation is between a biological readout and its experimental calibration. An agent may need to define a sum or weighted combination of its own states, while the supplied observable template retains assay-specific scales, offsets, and noise parameters. Writing *h_m_* for the readout mapping, the vector of agent-defined biological readouts (one readout rule each; the prompts call these rules measurement mappings), the prediction and noise scale for row *i* are

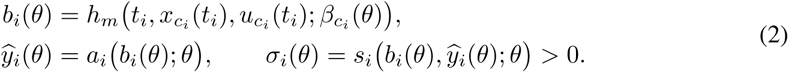

Here *a_i_* and *s_i_* are the supplied observable and noise templates for *o_i_*, including measurement-row parameter overrides; known time and input arguments are suppressed. For example, *a_i_*(*b*; *θ*) = *α_i_b_j_* + *γ_i_* calibrates readout *j* by an observation scale *α_i_* and offset *γ_i_*. Several observables can read the same readout with different calibrations, so a problem can have fewer readouts than observables. The composition of *h_m_* and *a_i_* gives the observation mapping *g_m_* in Appendix B.3.

Let *T_i_* be the declared linear, natural-log, or base-10-log observable transformation, and *q_i_* the declared standard normal or unit-scale Laplace error density. PEtab assumes independent measurement errors, giving the measurement NLLH

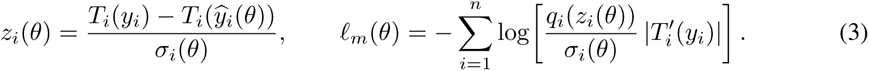

The noise scale is a standard deviation for normal errors and a scale parameter for Laplace errors, both in the transformed observation space. The Jacobian accounts for the density of measurements supplied in linear space; its contribution cancels in ΔNLL when measurements and transformations are shared. Fitting searches Θ*_m_* to minimise *ℓ_m_*(*θ*) + *P_m_*(*θ*), where *P_m_* contains retained objectiveprior terms and is zero when none are present (Appendix A.4). Signal-dependent noise expressions are rewritten in terms of the same `readout_*` symbol, preserving their statistical form. Keeping this noise model specification fixed was required to make NLLH values directly comparable between models constructed by agents and the published references; estimated noise parameter values can still differ between fits.

The parameter table is a curated subset of the published problem. It can retain observation parameters, fixed setup quantities, and selected initial-condition parameters, together with their estimation flags, bounds, estimation scales, and available prior annotations. Estimated kinetic parameters are introduced through the candidate model. Bounds constrain *θ*^min^ ≤ *θ* ≤ *θ*^max^ componentwise in linear parameter units; the declared linear or logarithmic optimisation coordinates are distinct from the observable transformations *T_i_*. A quantity fixed in a publication is not automatically treated as given knowledge: a value encoding an absent reaction or a previous fitting decision would itself constrain the mechanism being sought.

Initial conditions require corresponding care. Known baselines can be supplied as fixed parameters; estimated initial populations and compound initial expressions must preserve their intended dependence on model parameters. Condition-specific initial values remain experimental inputs. When pre-equilibration is prescribed, 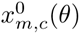 is constructed from the simulated steady state under the pre-equilibration condition, followed by any initial-state overrides in the simulation condition. For Fiedler, preparation adds a basal pre-equilibration condition and stimulus gate to preserve the prestimulus steady-state constraint after removing the published model. These choices influence both the admissible trajectories and the parameter count, and belong to the benchmark specification.

### A.4 Reference construction and interpretation

Reference summaries are regenerated only through the explicit `-refit` preparation option. The reference fitting path loads the collection’s published SBML model and parameter table directly. It uses AMICI, pyPESTO, and Fides, with per-example fitting budgets. The reported objective is the better of the multistart result and evaluation at the published nominal parameter vector. Objective-prior columns are removed for reference fitting and scoring, so the reference objective is the measurement NLLH. At the documented revision, candidate fits retain objective priors on supplied parameters, where present, and their contributions remain in the reported training objective; newly introduced kinetic parameters do not inherit the published model’s priors. These differences in prior treatment have only small effects on scores and do not materially influence the results of this study.

Documented exceptions adjust the reference problem when necessary for comparison. Selected published-fixed parameters are freed within their published bounds; both constrained and freed fits are evaluated, the better attainable likelihood is retained, and the larger parameter count is charged. Oliveira applies matching noise-parameter changes to candidate and reference specifications. Curated fitted literals embedded in the published equations can also contribute to the reference count, as in Fujita. These adjustments mean that the reference is derived from the publication rather than universally identical to its original estimation specification. Negative ΔBIC or ΔNLL favours the candidate over this numerical reference without establishing a uniquely correct mechanism. Appendix F visualises all 18 reference models using SBGN Activity Flow notation.

### A.5 Roster and table counts

Table 1 includes the generated roster without modification. Its reference values come from committed summaries; its observable and condition counts are the training counts of Figure 2, which include only observables and conditions that carry training measurements. The materialised tables can declare more: Fujita supplies sixteen condition rows but training responses for six, and Fiedler adds a basal pre-equilibration row. Neither table-row counts nor parameter counts measure the number of independently identifiable biological mechanisms.

**Table 1:** Published benchmark problems in the generated manuscript snapshot. *k*_ref_ is the reference parameter count used for scoring, *n* is the number of training measurement rows, and nll_ref_ is the stored reference offset used to compute training ΔNLL. Obs. and Cond. count the observables and simulation conditions that carry training measurements, which is what the reference fit used; a problem may ship more of either, and the surplus is not part of the estimation task. Fujita declares sixteen conditions and is trained on six, the other ten being the held-out arms of Figure 3, and Perelson’s second condition is its pre-equilibration. The offsets depend on each problem’s measurements and noise model and should not be compared directly across rows.

| Example | $k_{\text{ref}}$ | $n$ | $\text{nll}_{\text{ref}}$ | Obs. | Cond. |
| --- | --- | --- | --- | --- | --- |
| perelson | 3 | 16 | 223.49 | 1 | 1 |
| blasi | 9 | 252 | -1090.56 | 15 | 1 |
| rahman | 9 | 23 | 21.15 | 1 | 1 |
| crauste | 12 | 21 | 190.97 | 4 | 1 |
| armistead | 14 | 58 | -301.92 | 4 | 2 |
| sneyd | 15 | 135 | -319.79 | 1 | 9 |
| fiedler | 22 | 72 | -58.58 | 2 | 3 |
| boehm | 9 | 48 | 138.22 | 3 | 1 |
| brannmark | 22 | 43 | 141.82 | 3 | 8 |
| schwen | 30 | 286 | 939.88 | 4 | 19 |
| bruno | 13 | 77 | -46.69 | 5 | 6 |
| oliveira | 14 | 120 | 379.37 | 2 | 1 |
| fujita | 20 | 144 | -66.58 | 3 | 6 |
| isensee | 46 | 687 | 3941.46 | 3 | 123 |
| alkan | 65 | 1733 | 3171.38 | 12 | 73 |
| bachmann | 115 | 541 | -418.41 | 20 | 36 |
| lucarelli | 84 | 1755 | 1681.61 | 65 | 16 |
| raimundez | 142 | 627 | 539.19 | 5 | 170 |

## B Implementation and execution model

This appendix describes the implementation at revision `e0d6921`. Paths are relative to the software repository. The description explains how a proposed biological model becomes a numerical estimation problem and an inspectable record. It is distinct from the protocols of the archived evaluations: the reported runs span earlier revisions and different fitting budgets. In particular, current validation, stopping, parameter-counting, and checkpoint selection rules must not be assumed to have operated identically throughout those cohorts. The run exports (Appendix C.6) and cohort-specific methods (Appendix D) determine which settings support each result.

### B.1 Preparation, orchestration, and numerical state

The implementation separates problem preparation, agent orchestration, and numerical evaluation (Table 2). Preparation converts a published estimation problem into an agent-facing construction task. The harness then creates a working copy and launches a separate MCP server process. The server owns the mutable PEtab problem; the harness owns the agent interaction, run limits, and final comparison with the withheld reference. This boundary lets different agent configurations use the same editing and estimation interface. It does not make their information access or computational budgets identical; those remain experimental settings.

**Table 2:** Principal implementation boundaries. Modules are grouped by their responsibility in a run rather than by their individual helper functions.

| Responsibility | Main entry points and role |
| --- | --- |
| Preparation | `examples/setup.py` and `src/petab_harness/benchmark.py`: materialise the construction problem and manage separate reference preparation. |
| Orchestration | `src/petab_harness/runner/` and `src/petab_harness/agent/`: launch services, configure agents, enforce run limits, and archive outcomes. |
| Model revision | `src/petab_mcp/server.py`, `src/petab_mcp/session.py`, and `src/petab_mcp/operations.py`: apply a proposed change through reconciliation, validation, fitting, and version creation. |
| Numerical evaluation | `src/petab_mcp/engine.py` and `src/petab_mcp/fitting.py`: construct the AMICI/pyPESTO objective, estimate parameters, and produce simulation and optimization diagnostics. |
| Inspection and storage | `src/petab_mcp/data_access.py` and `src/petab_mcp/provenance.py`: expose diagnostic tables and preserve model versions and events. |

Materialisation starts from a pinned source collection and applies the example’s declared curation: identifier replacement, supplied parameter definitions, and observable-formula blinding, where required. It retains empirical measurements and experimental information while replacing the reference mechanism with an empty SBML model. Reference fitting is a separate operation, so preparing an agent problem does not silently regenerate its comparison score. The server receives the working problem and numerical configuration, while the benchmark judge retains the published-reference information needed for final assessment.

Communication uses MCP over Streamable HTTP. This also separates tool messages from compiler output: AMICI compilation writes to the server’s output stream, which would interfere with a standard-input/output JSON-RPC channel. The harness captures server logs and manages the subprocess lifecycle, including termination when its parent disappears.

### B.2 Mcp tools and resources

The following overview covers the `petab_mcp` interfaces relevant to the reported study. The harness assigns tools by role and evaluated ablation configuration (Appendix B.9); an available tool need not have been invoked in every run. The held-out evaluation tool is not registered with LLM agents.

#### Problem definition and measurements

`describe_problem` returns the problem dimensions, whether a model has been authored, the current version, and the default number of optimisation starts. `list_observables` exposes the supplied observable descriptions (`observableName`), observable formulas, and noise models; `list_conditions` exposes the experimental conditions and their prescribed model inputs. `get_measurements` filters measurements by condition and observable. Its default response summarises counts, time ranges, value ranges, means, and preview points for each observable–condition pair; setting `summarize=false` returns paginated individual measurements, with a default of 200 rows per page. These tools inspect the supplied experimental specification.

#### Model and parameter inspection

`get_model` returns the current model as Antimony or SBML, or a requested version’s stored Antimony source. `get_model_structure` returns a structured account of species, reactions, rate laws, and parameters, including whether each species’ initial value is fixed, estimated, or supplied by a condition. `get_parameters` reports parameter values, bounds, scales, priors, estimated/fixed status, and roles such as `kinetic`, `observable_scaling`, `noise_sigma`, and `initial_condition`. Filters select parameter identifiers or roles. It also identifies supplied parameters not yet declared in the current model; the SQL parameter table, in contrast, contains only the live estimation problem.

#### Model revision and estimation

`make_changes` accepts a complete Antimony model and/or changes to parameter bounds and estimation scales. The alternative `model_edits` argument accepts a batch of structured operations: add or remove a reaction, replace a rate law, set an initial value, set or remove an assignment rule, or rename a symbol. `base_version` applies such a batch to a stored model and its search space. A non-empty justification is required; references and an override of the multistart count are optional. The call reconciles (Appendix B.6) and validates the proposal, fits it, and records a version, returning fit scores, diagnostics, warnings, and flags indicating whether the change was applied and committed. Fitted parameter values come from estimation; they cannot be supplied through the parameter-edit argument. There is no separate agent-facing fit call. The treatment of rejected changes, failed fits, and duplicates is described in Appendix B.5.

#### SQL access to data and numerical records

`list_data_tables reports` available table names, columns, and row counts. query_data runs DuckDB SQL over measurements, `conditions`, `observables`, and `parameters`, together with `simulations`, `simulations_history`, `optimization_starts`, and `optimization_trace`. Simulation records associate each measurement with its prediction, residual, noise-normalised residual, and noise scale. The history table adds version identifiers for comparisons between candidates; optimisation tables describe individual starts and their iteration traces. Queries support filtering, joins, and aggregation, with 50 returned rows by default and pagination through `offset`. Raw measurement and SQL pages are capped at 1,000 rows. Explicit ordering is needed for stable SQL pagination. SQL runs on disposable copies with external file and network access disabled, so it cannot overwrite the problem or its stored scores (Appendix B.8).

#### Fit diagnostics

`get_fit_summary` returns the current likelihood and complexity scores, active selection criterion, parameter count including charged literals, fitted values, parameters near their bounds, and residual/noise flags. `get_fit_diagnosis` provides detailed residual or parameter reports, selected using `sections=residuals`, `parameters`, or `all`. These include the worst-fitting points, observable-wise residual tests, local parameter uncertainties and correlations, and variation across near-optimal starts (Appendix B.7). `get_fitting_log` returns a requested tail of the optimisation log, defaulting to 200 lines.

#### Version history and audit records

`list_versions` returns stored versions and their scores; `get_version` retrieves a selected model, parameter table, metadata, and available simulation and optimisation records. `diff_versions` compares two versions through a model-text diff, parameter additions/removals/changes, and fit-score differences. `restore_version` reinstates a stored model and fitted parameters without refitting, while appending a restoration record rather than erasing history. `get_provenance` and `get_errors` return recent events and rejected or failed operations, respectively, with a configurable limit of 50 records by default.

#### Progress and computational cost

`get_progress` reports the best stored version under the configured criterion, the number of fitted operations, recent improvement, the non-improving streak, and whether a stopping condition has been reached. `get_run_timing` reports accumulated serverside compilation and fitting times and counts. These are timings of numerical work, distinct from total agent elapsed time, token usage, or CPU consumption.

#### Modelling guidance and resources

When guidance files are present, `list_skills` lists their names, descriptions, and supporting files; `get_skill` retrieves a selected document or supporting file. The server also exposes read-only MCP resources under `petab://` for model text, PEtab tables, versions, provenance, errors, simulations, the data schema, and the fitting log, with additional resources for available guidance. These resources provide an alternative retrieval interface, rather than additional model-editing operations. External biological databases and delegation to specialist agents are supplied by the harness, not by `petab_mcp` itself.

#### Held-out evaluation for the harness

When validation data are configured, `evaluate_validation_data` scores the current model using the dataset’s declared protocol. Fixed-parameter evaluation retains the fitted parameter vector; transfer evaluation refits the estimated parameters for the target system, optionally overriding the multistart count. The target fit is not written back to the working model. The response records the evaluation status, mode, score, parameter and measurement counts, conditions, and observables. This tool is excluded from all modelling-agent allowlists: the harness invokes it for evaluation. That separation is implemented by tool selection in the harness, not by authenticated server roles (Appendix B.4).

### B.3 Semantic layer and mechanistic world model notation

We adopt the mechanistic world model notation of Posner et al. (2026) to describe the relation between our semantic layer and variable, mechanism, and structure discovery:

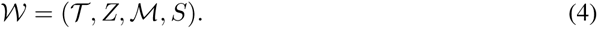

Here *T* denotes semantic variable types, *Z* the latent variables, including mechanistic parameters, *M* a library of mechanisms, and *S* their bindings to particular variables. Each mechanism combines a typed input/output-role signature with a transformation; a binding assigns those roles to variables of the corresponding types.

We use *semantic layer* for a model representation together with the interpretation that connects its entities and processes to executable mathematics. For the numerical implementation, let *m ∈ L_S_* denote a complete model proposal in the language of a semantic layer *S*, and let *E* denote the supplied experimental and observation specification. We denote its translation to dynamics, an observation mapping, and a parameter space by *C_S_*:

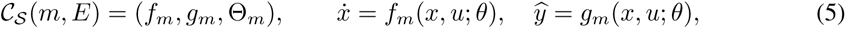

where *x* contains model states, *u* denotes prescribed experimental inputs, and *θ* ∈ Θ*_m_* contains the estimated parameters. Time dependence, initial conditions, and observation noise are specified by the model together with *E*; the displayed equations suppress these additional arguments (the full PEtab specification is given in Appendix A.3). With *V* (*m, E*) denoting the implemented acceptance checks, the candidate class is

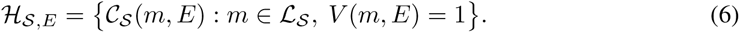

In the mechanistic world model notation, the states *x* and mechanistic parameters in *θ* instantiate *Z*. Reaction kinetics together with their stoichiometric effects, and model rules, specify transformations; references to particular species and parameters encode their bindings *S*. `gemot` stores these transformations together with their bindings in each authored model. It does not yet maintain a separate reusable mechanism library *M*, and semantic types *T* are not explicitly learned or enforced (Section 3). The notation therefore makes explicit both the variable, mechanism, and structure discovery implemented here and the remaining steps towards a mechanistic world model. Semantic regularisation can operate through restrictions on *H_S,E_* and through biological priors guiding proposals within it. It is distinct from the parameter-count penalty used in scoring; no separate semantic loss is optimised in the reported runs.

In our implementation, the model-language component is Antimony, implemented by libAntimony (Smith et al., 2009). It represents named species, compartments, reactions, kinetic laws, and rules. In `src/petab_mcp/operations.py`, the authoring path invokes PEtab’s `SbmlModel.from_antimony` to translate that description into SBML. Reconciliation connects it to the supplied parameter table and observable and noise templates (Appendix B.6); in particular, the agent’s assignment rules define the biological readouts required by PEtab. The resulting problem is compiled for numerical evaluation. Thus libAntimony supplies the mechanism’s formal interpretation, while the complete mechanism-to-measurement contract also depends on PEtab and the server’s checks. Merely giving a variable a biological name does not validate that interpretation. The current validation settings do not establish unit or thermodynamic consistency, and no automatic biological-ontology verification is claimed.

The inspected conversion path uses `getSBMLString`, which flattens Antimony submodules into a single executable model. This is a property of the present pathway, not a limitation of Antimony’s modular syntax or SBML’s `comp` package (Smith et al., 2015). The implementation check used PEtab 0.8.2 and Antimony 3.1.3; these local dependency versions do not identify every historical run environment. Structured authoring operations address reactions, rules, initial values, and symbols. They do not currently expose a curated library of modules with validated biological interfaces and applicability conditions.

Changing this layer would change the units in which an agent proposes and revises mechanisms. For example, a BNGL rule can specify a transformation of molecular sites across many possible complexes, while PySB can package rule generation into reusable Python components (Harris et al., 2016; Lopez et al., 2013). These alternatives would need their own translation, validation, complexity accounting, and readout-mapping interfaces before being evaluated under the same benchmark. A Python-based authoring layer would also need an execution policy appropriate to code submission; the current MCP access restrictions should not be assumed to carry over unchanged. Comparing such layers is prospective work, not a completed ablation.

### B.4 Trust boundary and resistance to score manipulation

The server separates the agent’s authority to propose a model from the authority to evaluate it. The exposed authoring operations accept model definitions and parameter-search settings; they provide no operation for rewriting measurement, condition, or observable tables, changing the scoring implementation, or submitting a replacement likelihood. Fitted parameter values, likelihoods, complexity counts, and version records are produced by the numerical pipeline. The benchmark judge reads these records when assessing a run, rather than treating the agent’s final narrative as evidence of a successful fit. Consequently, compliance with the evaluation contract does not depend solely on instructions asking the agent to respect it.

The boundary is enforced at several points. Parameter-edit schemas restrict the editable fields, and server-side checks reject unsupported changes. Model proposals are reconciled and validated before replacing the working problem. Checks cover the required model structure, readout rules, use of experimental inputs, and preservation of protected supplied constants. They also reject specified forms of observation-parameter reuse, including the use of a noise parameter inside the mechanism. The current complexity calculation charges non-exempt numerical literals embedded in model equations, reducing the opportunity to hide tunable choices outside the estimated-parameter table. These are executable checks with defined scopes and exceptions, rather than assessments made by another language model. Structural checks establish the presence and use of model elements, without proving their causal meaning; some checks are best-effort when their required metadata cannot be read.

Inspection preserves the same separation. DuckDB queries use disposable copies with external access disabled, so SQL updates cannot modify the underlying problem or stored scores. Restoring an earlier model selects a recorded version without deleting the intervening history. Accepted revisions, numerical outcomes, and rejected operations remain available for auditing. Together, these controls make reported improvements traceable to an evaluated candidate and prevent direct alteration of the benchmark records through the supplied tools.

This claim is scoped to the configured harness and MCP interface. The default agents receive curated tool sets with direct access to the host filesystem disabled. The local server is not an operatingsystem sandbox, and withholding a tool from an agent is not equivalent to authenticating separate server roles. Broader host or endpoint access would require additional isolation. Evaluation integrity also differs from scientific adequacy: a model can improve a correctly computed objective through overfitting or poorly constrained assumptions. The checks do not prove that every possible strategy for exploiting an objective has been excluded.

### B.5 From a proposed revision to a stored version

The principal authoring operation, `make_changes`, requires a non-empty justification. It accepts either a complete Antimony model or a batch of structured operations addressed to model identifiers. Parameter bounds and estimation scales can be changed in the same call. The two model-authoring routes are mutually exclusive. Structured edits may target an earlier version; the implementation retrieves that version’s submitted source and parameter search space and records the resulting branch relationship. Bibliographic references can accompany an edit but are not mandatory.

Reconciliation (Appendix B.6) aligns the proposed SBML model with the PEtab parameter and observation interface. Parameter edits run after reconciliation so that they can address newly introduced parameters. These operations act on a candidate copy. Validation checks the resulting problem, including the presence of dynamics and preservation of protected supplied constants, before the candidate replaces the live state. A rejected candidate therefore leaves the previous working problem intact and returns an error explaining the rejected operation.

An accepted edit proceeds immediately to fitting. When evaluation creates a version, its model, fitted parameters, scores, and diagnostics are retained together. The submitted source is stored separately from the reconciled Antimony representation, allowing subsequent edits to refer to the agent’s model text. If fitting or compilation fails before creating a version, the server restores the previous working state. The edit and its failure remain in the event record. If a later diagnostic or response-construction error occurs after a snapshot was created, that version is retained; an error response alone is therefore insufficient to infer whether the model state changed.

Duplicate detection further distinguishes tool calls from fitted versions. An unchanged candidate can return its existing score without creating another version, whereas an increased multistart budget can justify a new fit. Structure-level checks also identify repeated visits and can stop repeated fitting of the same structure. Consequently, tool-call counts, optimization counts, and version counts measure different activities.

### B.6 Reconciliation of model and PEtab specification

#### Purpose and scope

Reconciliation makes an authored model consistent with the supplied PEtab specification (Appendix A.3). After Antimony conversion or structured model edits, `src/petab_ mcp/reconcile.py` updates the candidate SBML model and parameter table before validation and fitting. Measurement, condition, and observable tables remain unchanged. The agent supplies the mechanism and its readout rules; reconciliation infers neither missing interactions nor readout rules.

#### Removing unused symbols

The pass first removes species and parameters that are not referenced by reactions, model equations, initial assignments, or the fixed PEtab tables. Rule targets are retained, and initial assignments belonging to removed symbols are removed with them. This is a syntactic cleanup, not a test of dynamical relevance or identifiability: a referenced but redundant state can remain. If no species survives, an internal placeholder keeps the model safe for downstream handling; it does not replace the separate requirement for an authored model with dynamics.

#### Making initial conditions explicit

Existing parameterised initial conditions, including compound expressions such as *x*_2_(0) = *θ*_1_*θ*_2_, are preserved. Other nonzero initial values generally receive a named initial-condition parameter and become estimable. Zero or unspecified initial pools normally receive no new parameter, and an initial value of one is treated as a fixed unit normalisation; supplied parameter definitions take precedence over these defaults. Species initialised through condition-table overrides receive no additional initial-condition parameter. Assignment-rule targets also receive none, and conflicting initial assignments are removed. These conventions prevent the introduction of free parameters that are immediately overridden, while retaining the parameter dependence of the agent’s initial state assumptions.

#### Synchronising the parameter table

New model parameters receive PEtab rows, and obsolete rows are removed. Parameters needed only by observable or noise formulas and measurement-row overrides are retained. Symbols that are species, computed rule targets, or condition-table override targets are excluded where PEtab forbids them as parameter-table entries. If a reintroduced parameter was supplied with the problem, its original row is restored, including estimation status, bounds, scale, nominal value, and available prior annotations (Appendix A.4). Other new kinetic parameters are estimated, with default bounds that include the submitted nominal value; positive values normally use logarithmic coordinates. Parameters used as exponents receive linear coordinates and narrower default bounds. Existing search settings are retained, apart from the initial-condition conventions above. Explicit edits to bounds or scales run after reconciliation, so they can address newly introduced parameters.

#### Validation and feedback

The reconciled candidate then undergoes PEtab and SBML checks and the additional safeguards in Appendix B.4. These check required readout rules, use of prescribed inputs, protected supplied constants, and prohibited reuse of observation parameters. Missing readouts are reported to the agent for correction, rather than filled automatically. The edit pipeline records added and removed parameters and pruned species, and reports how each species’ initial condition is fixed, estimated, or set by a condition. Rejection leaves the working problem unchanged; accepted proposals proceed to fitting under the versioning rules in Appendix B.5.

#### Relation to complexity accounting

Reconciliation determines which named quantities enter the estimated-parameter count. The scorer separately adds non-exempt numerical literals in model equations to that count (Appendix B.7); these literals are not automatically promoted to fitted parameters. Consequently, the submitted model and its reconciled representation are both retained for audit. Reconciliation and validation check consistency with the experimental specification; they do not establish biological correctness or uniqueness of the mechanism.

### B.7 Parameter estimation and numerical diagnostics

AMICI supplies compiled simulation and sensitivity calculations, pyPESTO constructs and coordinates parameter estimation, and Fides performs local trust-region optimization (Fröhlich et al., 2021; Schälte et al., 2023; Fröhlich & Sorger, 2022). Multistart optimization explores the supplied parameter bounds; the best returned solution is written back to PEtab nominal values. Reported fitted values are in linear units, with their estimation scales recorded separately. Neither multiple starts nor convergence of an individual start establishes a global optimum.

#### Warm starts across model iterations

To reuse numerical work after a model revision, the implementation reserves ⌊*N*_starts_*/*2⌋ starts for warm initialisations when usable parameter values are available; the remaining starts are sampled afresh. The current nominal point, normally the preceding fit’s best solution, provides the reference point. Distinct finite solutions from the preceding multistart are also reused, matching shared parameters by identifier and converting between estimation scales through linear values; parameters absent from that fit retain their current nominal values. For newly added parameters, additional candidates test unit values and lower or upper bounds, jointly and, when several parameters are added, one at a time against unit values for the others. These candidates receive priority within a limited part of the warm-start budget, followed by the nominal point and previous solutions ranked by objective. Candidates are clipped to the current bounds and deduplicated in the free parameter coordinates.

Any remaining warm-start slots are filled by Gaussian jitter around the clipped nominal point, with standard deviation 0.05 times each parameter’s bound width in its estimation coordinates (linear, log, or log_10_), followed by clipping. Unbounded coordinates receive no jitter. A separate random-number generator with seed zero makes these perturbations reproducible without changing the fresh-start sampling stream. If warm initialisation is unavailable, all starts are sampled afresh. This combines reuse of earlier fits with exploration after structural changes; the candidate values do not guarantee recovery of the preceding model’s likelihood.

For eligible observation parameters, the current implementation attempts hierarchical estimation through pyPESTO (Loos et al., 2018; Schmiester et al., 2020), with a joint-estimation fallback when that construction is unavailable. Parameters eliminated from the outer optimization remain estimated parameters in the PEtab problem and its complexity accounting. The current scorer additionally charges non-exempt numerical literals embedded in model equations. Archived parameter counts must be interpreted using each run’s scoring convention; the availability of hierarchical estimation does not imply that every archived fit used it.

#### Diagnostic interfaces and scope

Fit results and `get_fit_summary` expose scores and compact warnings; `get_fit_diagnosis` supplies residual or parameter reports on request. DuckDB tables provide individual simulated measurements, residuals, and optimisation records for further analysis, while the fitting log exposes numerical messages (Appendix B.2). The diagnostics below describe the implementation at the documented revision for the study’s tool configuration, rather than the exact feedback available in every archived run. They concern the supplied training data. Held-out evaluation is reserved for the harness and excluded from modelling-agent tool sets. Numerical agreement, diagnostic passes, and successful integration do not establish biological adequacy or uniqueness of a mechanism.

#### Fit scores and numerical consistency

The summary reports fit status, the likelihood objective, the number of measurements, and the complexity count *k*, separating estimated parameters from charged numerical literals. It returns BIC, AIC, AICc, CAIC, HQC, and TIC where defined, together with the configured criterion used for ranking versions (configuration in Appendix C.4). Undefined scores are returned as missing, for example AICc when the sample is too small relative to *k*, or TIC when its effective parameter count cannot be computed. Compact reports omit that effective count, while archived records retain it. Improvement reports compare the active criterion with the previous best checkpoint; a decrease of at least two units updates that checkpoint, allowing smaller successive improvements to accumulate towards the threshold.

The fit archive retains both the best optimiser objective and the likelihood obtained by independently re-simulating its fitted parameters. A finite optimiser objective is used as the authoritative fitting score. An `objective_note` identifies failed re-simulation or a discrepancy exceeding 10*^−^*^3^ max(1*, |ℓ*_opt_), where *ℓ*_opt_ is the optimiser’s negative log-likelihood. The summary also counts missing simulated measurements. Thus a finite score does not imply that the independent simulation reached every measurement. The reported *χ*^2^ requires a complete residual evaluation, rather than summing only successfully simulated points. The server also returns *χ*^2^*/*(*n* − *k*) and an upper-tail 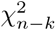 probability when defined; these should not be interpreted as calibrated adequacy tests when noise scales are fitted.

#### Individual residuals and observable summaries

Each simulation-table row associates an observable, condition, and measurement time with the measurement *y_i_*, prediction 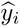, evaluated noise scale *σ_i_*, raw residual, and noise-normalised residual:

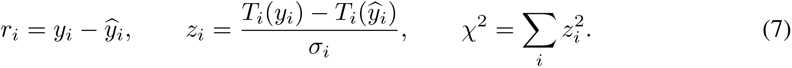

Here *T_i_* is the declared linear, natural-log, or base-10-log observable transformation; the noise formula can give a different *σ_i_* at each point. The field `residual_weighted` contains *z_i_*, not *z*^2^. Missing predictions, unevaluable noise, or zero noise leave the normalised residual missing. The history table adds a version identifier, enabling comparisons at the same measured point across models.

The detailed residual report gives each observable’s usable point count, contribution to *χ*^2^, mean *z_i_*, largest *z_i_*, and residual-test summaries. It also lists the eight worst individual discrepancies by default, including their measurements, predictions, conditions, and times. To bound the response, the default observable table combines the 12 largest *χ*^2^ contributions, three largest absolute mean residuals, and six observables flagged for residual structure: at most 21 distinct entries. Omitted counts, omitted *χ*^2^, and the number of omitted flagged observables remain visible. Setting `max_observables=0` removes this truncation; `worst_n` controls the point list separately. The aggregate residual-structure verdict is returned automatically with fit results and score summaries.

#### Bias and temporal structure

Sign and runs tests use points with available weighted residuals, average replicate residuals at each condition–time pair, and discard zero or non-finite residuals. Within each condition, an exact two-sided binomial sign test checks the balance of positive and negative residuals. An exact two-sided Wald–Wolfowitz runs test checks their ordering in time, conditional on the sign counts. For each observable and test, condition-specific probabilities are combined using Fisher’s statistic 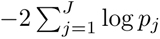, compared with 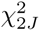 under the independence assumption. Either combined *p <* 0.05 flags the observable. There is no adjustment across observables or between the two test families, so these flags are screening diagnostics. Reports include combined probabilities, contributing-series counts, sign counts, and low-power flags. Fewer than eight nonzero design-point residuals, or no computable runs test, triggers a heuristic low-power warning; this is not a formal power calculation. Runs tests cannot assess a series containing only one residual sign.

#### Replicate lack of fit

A separate check compares predictions with replicate scatter without relying on the fitted noise scale, so it can remain available when that scale is zero or unevaluable. At a design point *i* with *n_i_* replicates, sample standard deviation sd, and common prediction 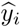, it computes

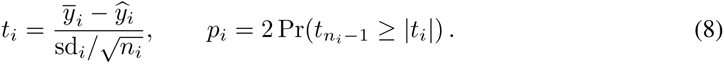

Measurements and predictions are expressed in the noise-additive, scale-normalised coordinate; proportional noise uses an approximate logarithmic coordinate. A usable point requires at least three finite replicates, positive sample variance, and a finite prediction shared by its replicate rows. At least three usable design points must share a noise-parameter family. Probabilities are Holm-adjusted within each such family and flagged below 0.01, without additional correction across families. The compact report gives availability, usable and flagged counts, the largest standardised discrepancy, the smallest adjusted probability, and up to five worst flagged points.

An additional guardrail subtracts the number of estimated non-noise parameters from the pooled number of usable design points. Parameters that also affect observation scaling remain in this count. A non-positive difference makes the report unavailable and discards its flags; a difference of one or two marks an unflagged result as weak; larger differences receive the implementation’s powered designation. These are heuristic evidence-strength categories, not formal power estimates. Unknown parameter counts prevent a power claim. Reports distinguish absent replicates, insufficient replication, too few usable points, and unresolved inputs, and identify partially untested noise families. A flagged fitted prediction indicates remaining discrepancy; it does not prove that no parameter vector could fit that point.

#### Estimated noise scales

For estimated noise-only parameters, the noise report gives the fitted value, effective observation standard deviation, supporting point count, empirical data spread, and their ratio. It also reports the data-derived upper-bound cap, a data-derived noise estimate where available, the method used to obtain it, and the ratio of fitted to estimated noise. Comparisons use a common noise-additive, scale-normalised coordinate rather than comparing a raw parameter directly with measurements in incompatible units. An effective scale above 0.75 of the empirical data standard deviation is labelled inflated, and below 0.01 is labelled collapsed. These are heuristics; fewer than five supporting points or unevaluable quantities yields an unknown status. Fixed noise parameters are not included in this report.

The preferred data-derived noise estimate pools within-design-point replicate variance and requires at least six residual degrees of freedom. Otherwise, the implementation uses successive measurement differences within time series, 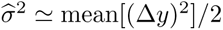. These estimates do not use fitted residuals, although their scale normalisation can depend on fitted observation scales. The difference-based fallback includes true temporal change and is not a reliable lower bound on measurement noise. These additional ratios provide advice rather than formal tests. A `scale_coupled` flag identifies noise formulas in which the noise parameter is combined with another estimated parameter, such as an observation scale or an additive offset, cautioning against direct cross-model comparisons of noise parameters. Reporting these diagnostics does not itself change the parameter bounds.

#### Parameter values, bounds, and local uncertainty

Fitted values are reported on their original scale alongside the estimation scale. After the first fit, compact responses may list only added, removed, or changed parameters (more than a 1% relative change from the parent version), with an unchanged-parameter count; complete values remain available through parameter and version inspection. A bound warning identifies estimates within 1% of the allowed bound span of either endpoint, measured in the estimation coordinate. It reports the parameter, value, bound, and side. A bound warning does not by itself demonstrate non-identifiability.

Parameter diagnosis uses local objective geometry at the stored fitted optimum. The preferred sensitivity-based information approximation includes measurement-residual and noise contributions,

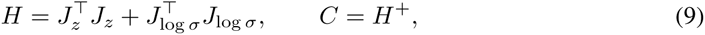

with an objective-Hessian or residual-sensitivity fallback when required. Here *J_z_* is the Jacobian of the noise-normalised residuals *z_i_* defined above, and *J*_log_ *_σ_* is the Jacobian of the natural logarithms of the evaluated measurement noise scales *σ_i_*. The subscripts identify the quantities being differentiated. Writing *η_j_* for free parameter *j* in its estimation coordinate (linear, natural-log, or log_10_), the entries are (*J_z_*)*_ij_* = *∂z_i_/∂η_j_* and (*J*_log_ *_σ_*)*_ij_* = *∂* ln *σ_i_/∂η_j_*, evaluated at the stored optimum. Rows therefore index measurements and columns index free outer parameters. Residual derivatives include dependence of both predictions and noise scales on these parameters. The second term captures parameter dependence of the noise scales and vanishes when those scales are parameter-independent. Here *H*^+^ is the Moore–Penrose pseudoinverse; *H* is a local approximation, not necessarily the exact expected Fisher information. The per-parameter report gives 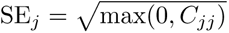 in the estimation coordinate. For log and log10 parameters it also expresses this as a one-standarderror multiplicative factor, exp(SE*_j_*) or 10^SE^*^j^*, rather than a confidence interval. For hierarchical fits, these sensitivities hold the fitted inner parameters fixed; those parameters are excluded from the outer-parameter geometry, and the report explicitly notes their omission. Pseudoinversion discards null directions, so even small finite standard errors must be read together with the rank diagnostics.

#### Correlations, flat directions, and multistart variation

Marginal correlations are 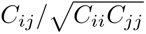; partial correlations use 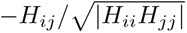 . Pairs with absolute correlation at least 0.9 are reported in descending order. The report gives matrix rank, a rank-deficiency flag, and a condition number based on the largest and smallest strictly positive eigenvalues; the latter can remain finite for a rank-deficient matrix. Eigenvectors with *λ/λ*_max_ *<* 10*^−^*^3^ identify flat directions, with up to four dominant parameter loadings above 0.15 in absolute value per direction. Pair and direction lists default to eight entries each; the per-parameter table is not subject to that truncation.

The complementary multistart report gives each parameter’s standard deviation and range in estimation coordinates across solutions within ΔNLL ≤ 1 of the best objective. If fewer than three starts qualify, it instead uses up to five best recorded starts, which can lie outside this band. At least two values are needed for a parameter’s dispersion estimate. Up to 12 simplification candidates are ranked by the number of warning signs: a bound flag, scaled standard error at least one, high marginal correlation, participation in a flat direction, or multistart standard deviation at least 0.5. Reasons accompany each candidate. These signals motivate refitting or sensitivity analysis; they neither justify automatic removal nor establish structural identifiability.

#### Suggestions for sharing parameters

The parameter report also estimates the local cost of tying two parameters in estimation coordinates. For *ω* = *e_i_ − e_j_* and fitted vector 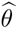, the quadratic approximation is

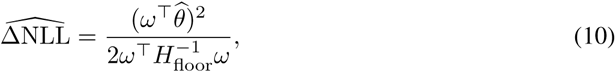

where *H*_floor_ is a numerically regularised version of the local information matrix *H* in Equation (9). For the eigendecomposition *H* = *Q* diag(*λ*_1_*, . . ., λ_d_*)*Q^⊤^*, it retains the eigenvectors but replaces eigenvalues below a positive threshold *τ* by that threshold:

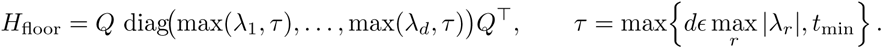

Here *d* is the number of free parameters represented in *H*, *ɛ* is machine precision, and *t*_min_ is the smallest positive normalised floating-point number. Unlike the pseudoinverse used above, this inverse retains large contributions from nearly flat directions, allowing a parameter tie along such a direction to have a small estimated cost. The calculation is unavailable for material negative curvature, 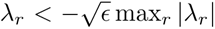. By default, candidate pairs must share an inferred parameter role and estimation scale, and neither may be near a bound; metadata limitations and disabled filters are disclosed. Reports include the estimated fit cost, largest required movement in scaled coordinates, and predicted change in BIC: tying one parameter saves log *n*, before accounting for the estimated fit cost. The detailed report defaults to the eight cheapest pairs, ordered by estimated cost and then movement, and gives total priced and score-improving pair counts and reasons for skipped pairs. Up to three score-improving suggestions accompany fit summaries automatically. These are proposals for an actual constrained refit: approximate costs are not additive across multiple ties, and an empty list does not certify that the model is parsimonious.

#### Optimisation, revision, and stopping diagnostics

The fit result gives the actual multistart count and a histogram of termination flags, distinguishing convergence criteria from budget or numerical exits. The per-start SQL table contains objective-ranked starts, objective values, elapsed times, and function, gradient, and Hessian evaluation counts; the trace table contains available iteration-wise objectives and times. Termination flags are not columns of the per-start table. `get_fitting_log` returns the last 200 lines by default, or all lines when requested, including convergence, gradient, and conditioning messages. Server timing separately accumulates compilation and fitting counts and wall times; these are not summed CPU time or total agent elapsed time.

Revision feedback includes validation errors, reconciliation warnings, and a nesting advisory when a child gains estimated parameters but worsens the fitting objective by more than 10*^−^*^3^. Where detectable, it names a parent parameter no longer estimated or referenced less often. This is a prompt to check whether the edit preserved the parent as a special case and whether optimisation was adequate, not a symbolic test of nesting. Non-nested alternatives remain legitimate proposals. The outstanding advisory persists across rollback until a qualifying parameter-addition comparison clears it. Progress reports additionally expose the best version, recent improvement, non-improving streak, and stopping status; their availability to agents depends on role.

Residual flags influence stopping as well as diagnosis. At this revision, any sign, runs, or replicate lack-of-fit flag prevents the server’s early adequate-plateau stop. The harness’s explicit finish gate refuses replicate lack-of-fit flags for up to three consecutive attempts before relenting; sign and runs flags alone do not cause that refusal. Estimated-noise inflation or collapse and a current, non-stale rank-deficient information matrix each trigger similarly bounded refusals. Low-power and weak-pass qualifications are advisory, and unavailable tests do not block completion. Consequently, reaching a stopping condition is not a statistical certificate that the mechanism is adequate.

#### Unavailable, stored, and stale reports

Diagnostics expose availability and reasons for missing evidence. Parameter geometry uses the frozen fitted optimum and an already compiled objective; reading the report does not compile a new model. If local-information evaluation fails, multistart-only information can remain available. Stored reports survive restoration and restart, with a stale flag for a changed structure; a live report also warns when current parameter settings differ from those used for its fitted geometry. Replaying a stored report does not expand lists that were truncated when it was generated. Residual inspection may simulate an already compiled model when needed. Missing simulation rows, unavailable uncertainty estimates, and empty candidate lists must therefore be distinguished from successful checks with negative findings. Compilation and integration failures require numerical investigation before being interpreted as evidence against a mechanism.

### B.8 Data access and durable provenance

The `query_data` interface uses DuckDB to query PEtab tables, simulated measurements, residuals, and optimization diagnostics (Raasveldt & Mühleisen, 2019). A version-history table supports comparisons across stored simulations. Each query receives newly created in-memory tables copied from the session data. SQL may modify these disposable tables, but changes are not written back to the PEtab problem. External file and network access are disabled and the database configuration is locked before executing the query. Results are bounded and paged; stable pagination requires an explicit ordering. A version without readable simulation records contributes no trajectory rows and must not be interpreted as a successful fit.

The provenance store keeps the append-only event stream in `provenance.jsonl`, separate from snapshots in `versions/vN/`. A snapshot contains model source and reconciled model text, `parameters.tsv`, metadata, and available simulation and optimization records. A stored pyPESTO result can accompany the structured summaries. This distinction preserves attempted changes as well as numerical outcomes and supports restoration without discarding earlier candidates. Run-level records additionally identify configuration, code revision, timing, and available token-use measurements. A dirty-tree digest can distinguish uncommitted source states, but cannot reconstruct an unavailable patch. These records make the process auditable; they do not guarantee exact replay of external language-model or knowledge-service responses.

### B.9 Agent roles and access to biological knowledge

In the panel configuration, the modelling lead (prompts in Listings 114 and 115) authors revisions and delegates bounded analyses. The mathematician (Listing 116) examines structure and estimation behavior; the systems scientist (Listing 117) assesses the biological mechanism; and the signal engineer (Listing 118) queries measurements and residuals. Tool sets are assigned by role. In particular, detailed measurement and SQL access is delegated to analysts, whose summaries return to the lead, limiting repeated inclusion of large tool outputs in its context. The biological curator (Listing 119) receives curated entity information and observable descriptions, rather than the fitting tools or authority to edit the model.

Knowledge tools are discovered from configured services and filtered through a named allowlist. The default set covers identities, protein domains, interactions, pathways, and ontologies, including UniProt, Reactome, and OmniPath (The UniProt Consortium, 2025; Milacic et al., 2024; Türei et al., 2026). These resources are useful only for models that describe molecular biology. Literature retrieval was implemented but disabled in the reported experiments. This separates structured retrieval from free-text searches that may reveal a benchmark’s source study. It does not establish the absence of leakage from other knowledge sources. Unavailable services are skipped rather than replaced by unrestricted tools, and the resolved tool set is recorded. Access to a tool therefore establishes a capability, not evidence that it was used in every run.

### B.10 Budgets, stopping, and final candidate selection

Run limits distinguish agent steps, model-output limits, total elapsed time, and numerical work per fit. Graph-step limits do not directly bound calls made within delegated analyses. The multistart count is a run setting that individual permitted calls can override. Per-start time limits and the number of parallel workers determine the expected fit duration; transport and stall timeouts include additional compilation and execution allowances. Budget accounting must therefore distinguish one long fit from an unresponsive agent step. Stopping and finish checks also distinguish an agent’s declared completion from the existence of a scoreable model. Their thresholds and enabled controls are configuration- and revision-dependent.

In the reported `gemot` runs, the judge ranks stored candidates by BIC. It scans finite scores in order and prefers the first candidate whose available noise diagnostics do not trigger its plausibility check. Missing reports are unassessable rather than evidence of plausible noise. If none passes, the best-scoring candidate remains available for reporting with the failed check. Restoration failure is reported and leaves the current model to be assessed. Held-out predictions are reported separately from the construction-success decision. These distinctions are essential when comparing archived summaries or reselecting checkpoints for a new analysis.

## C Software setup, execution, and run records

This section describes the implementation at code revision `e0d6921` and gives commands for installing the software and exercising the modelling workflow. This is the version documented here, rather than a claim that every reported experiment used this revision. Historical cohorts must instead be associated with their recorded code, problem, prompt, and configuration versions. The examples below are starting points for new runs; reproducing a reported cohort additionally requires its resolved configuration and retained inputs and outputs.

### C.1 Installation and dependencies

The distribution contains both `petab_mcp`, which implements the server, and `petab_harness`, which coordinates experiments. It requires Python 3.12 or newer. The numerical stack is a core dependency: model evaluation uses AMICI, pyPESTO, and Fides (Fröhlich et al., 2021; Schälte et al., 2023; Fröhlich & Sorger, 2022). A working C++ compiler, CMake, Ninja, and SWIG are needed because AMICI compiles generated model code. Starting from the root of a source checkout with the compiler available, a local environment can be prepared as follows:

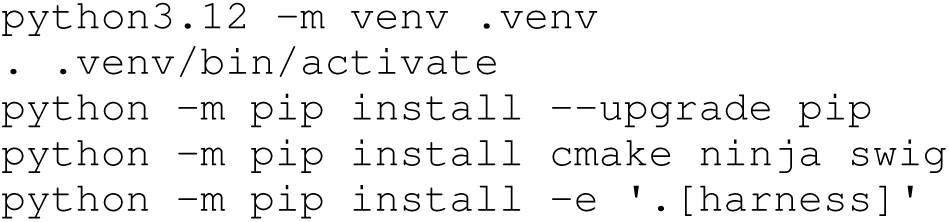

The `agent` extra contains the Deep Agents (LangChain, Inc., 2025) and LangChain dependencies; the `harness` extra additionally installs Phoenix tracing and Weights & Biases logging. The optional `explorer` extra supports the run browser, and `dev` supplies testing and linting dependencies. An editable installation does not freeze dependency resolution: the environment’s resolved package versions and compiler information must be retained with an experiment. The repository also supplies a Linux Conda environment with the compiler and HDF5 development dependencies for cluster use.

### C.2 A local agent run

For a local agent run on the Fujita benchmark, configure OPENROUTER_API_KEY in the execution environment, prepare the empirical problem (Appendix C.3), and select the model explicitly:

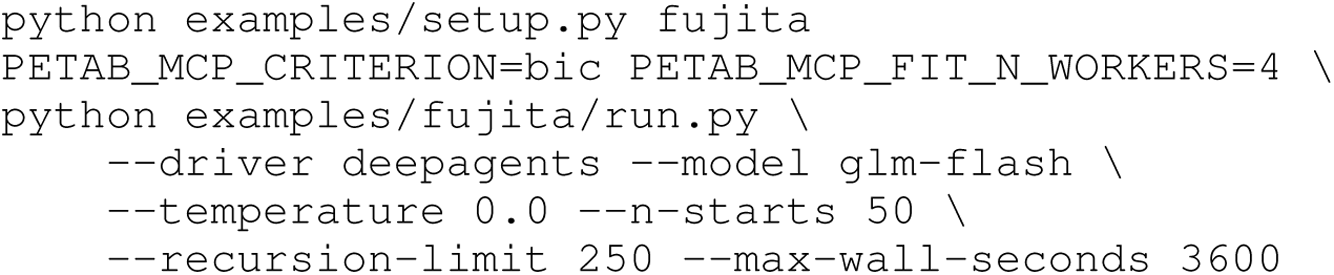

At the documented revision, `glm-flash` resolves to `z-ai/glm-5.3-flash`. The model registry also specifies provider routing, so the alias alone is insufficient to identify the service configuration across revisions. Phoenix tracing is enabled for every agent run; without a configured collector endpoint (`PHOENIX_COLLECTOR_ENDPOINT` or `OTEL_EXPORTER_OTLP_ENDPOINT`), the runner starts a local collector. Weights & Biases logging is separately enabled with `-wandb`.

Each invocation recreates the example’s `work/` directory from its prepared `problem/` directory, launches a stateful MCP server, runs the selected driver, and writes `work/result.json`. The server uses HTTP transport to keep compiler output separate from protocol messages. Model compilation and server logs are written to `amici_build.log` and `server.log`. Concurrent repetitions of one example require separate example directories because the local working path is reused.

### C.3 Preparing empirical problems and reference fits

Published problems are materialised from the Benchmark-Models-PEtab collection at revision `1763dfcd66d25bfd97492cfedf6a730472c383d0`. The setup script obtains this pinned collection and applies each example’s declared configuration:

~~~
python examples/setup.py
python examples/setup.py fujita alkan raimundez
~~~

The first command prepares the complete empirical suite; the second prepares a subset. This step constructs the agent-facing problem and leaves the committed reference scores unchanged. After materialisation, an empirical problem is run through its own wrapper, for example `examples/f ujita/run.py`, with the same runner options as above.

Reference fitting is an explicit additional operation:

~~~
python examples/setup.py --refit fujita
~~~

This regenerates the reference record using the published model and the example’s declared adjustments. These include, where applicable, freeing parameters on both sides of the comparison, parameter overrides, and charges for fitted values represented by model literals. Reference multistart counts and per-start limits are declared in each example. A regenerated reference should be accompanied by convergence information; increasing the number of starts cannot compensate for truncating nearly every start before convergence. The cluster utilities `slurm/refit_references.sh` and `slurm/verify_refit.py` support reference submission and inspection of per-start outcomes. Reference regeneration is therefore a separate numerical experiment, not a routine consequence of downloading the benchmark.

### C.4 Selection criteria and computational budgets

The reported experiments used BIC to guide search and final selection. The standalone server selects it with `-criterion bic`; the example runner inherits `PETAB_MCP_CRITERION=bic`. Other information criteria reported in the results are retrospective comparisons, not separate search arms.

The local runner defaults to 50 multistarts and a recursion limit of 250 graph steps. `-n-starts` sets the default fit budget; an individual model-edit call can request a different count. `-max-wallseconds` requests a graceful stop at a subsequent step boundary, allowing a committed candidate to be judged. It is not a hard process deadline and does not replace a scheduler allocation.

Numerical budgets have separate controls. `PETAB_MCP_FIT_N_WORKERS` sets the worker limit, and `PETAB_MCP_FIT_PER_START_TIMEOUT` bounds individual optimisation starts, with a default of 180 seconds. Transport and stall timeouts account for the requested starts, parallel worker waves, and compilation allowance. The documented implementation sets AMICI’s absolute tolerance to 10*^−^*^10^ unless `PETAB_MCP_FIT_AMICI_ATOL` overrides it. These implementation defaults should not be substituted for the settings recorded for historical fits or held-out rescoring.

### C.5 Cluster execution and retained records

The supplied Slurm workflow uses cluster-specific partition names and resource allocations that require adaptation to the target system. `slurm/setup_env.sh` builds or updates `envs/pet ab-mcp/` inside a compute allocation. It installs the development, harness, and explorer extras. A shared Phoenix collector is started using `slurm/phoenix_ctl.sh` with the `start` argument before submitting agent sweeps.

`slurm/make_sweep.py` expands problems, models, temperatures, and repetitions into a uniquely named configuration table and submits one array task per combination. Its resource options include `-cpus-per-task`, `-mem`, `-time`, and `-throttle`. Unlike the local runner, it defaults to 100 multistarts and 500 graph steps. The resolved per-example count is raised to AGENT_N_STARTS, or to REFIT_N_STARTS when no agent count is declared, if this exceeds the requested count. Per-start limits similarly prefer `AGENT_PER_START_TIMEOUT` and otherwise use the reference limit. Reference and agent budgets can therefore differ, especially for large models; the resolved configuration table records these allocations. The default agent deadline leaves 1,200 seconds within the allocation for judging and archiving. The evaluated configurations are listed in Appendix C.6.

Array tasks copy example files into private working directories, align fit parallelism with the allocation unless explicitly overridden, and collect result and log files with the combination identifier and exit status. At normal completion, the harness attempts to archive model versions, parameters, simulations, accepted-change provenance, and rejected-tool-call records under `examples/_runs_a rchive/`. The result records code and prompt versions, resolved ablations, and the trace identifier linking it to Phoenix. Archives omit large optimisation files such as `pypesto_result.hdf5`; these require separate retention for subsequent optimisation audits. A working-tree digest identifies differences but cannot reconstruct the patch. Consequently, reproducibility requires preserving the actual source changes, environment, prepared problems, reference records, configuration tables, and traces alongside the compact run archive.

### C.6 Configurations of the scored runs

Tables 3–5 give the configuration of every agent run that Figures 2–4 score, read from each run’s archived record and scheduler log rather than from the implementation defaults above; `paper/ge nerated/run_configurations.csv` lists every run with every field. The harnesses carry the names used in Figure 3. The support a run was given is reported as it ran: the biology-blinded panel records the curator as enabled in its ablation state but seats none. The null-arms experiment of Figure 3 kept no scheduler logs, so its budget is the one its submission manifest records. No run set a reasoning effort, so every call ran at its model’s default as the provider serves it: medium effort for GPT-6-Astra and max, the highest of its three levels, for GLM-5.3-Flash, with reasoning on for both (Table 5). The exception is a call whose response spent its whole completion budget on reasoning and returned nothing; the harness re-issued it at low effort.

**Table 3:** **Configuration of the scored agent runs**, by figure and arm. *Given*, the support available to the run as in Figure 3; *prompt set*, a hash over the run’s complete prompt registry; *commit*, the code revision the run executed. A count after a value gives the number of runs with it.

| Arm | Language model | Given | Prompt set | Commit | Runs |
| --- | --- | --- | --- | --- | --- |
| *Figure 2: 90 runs over 18 benchmarks* |  |  |  |  |  |
| O1 | GLM-5.3-Flash | curator + biology + gemot | 637a341 | 604449e (80),<br>c1a851f (5), 59cde21 (5) | 90 |
| *Figure 3: 30 runs* |  |  |  |  |  |
| F2 | GPT-6-Astra | biology + gemot | 971aaa6 | 5969dd5 | 5 |
| F3 | GPT-6-Astra | biology + bare agent | 637a341 | 5969dd5 | 5 |
| O1 | GLM-5.3-Flash | curator + biology + gemot | 637a341 | 59cde21 | 5 |
| O2 | GLM-5.3-Flash | biology + gemot | 971aaa6 | 59cde21 | 5 |
| O3 | GLM-5.3-Flash | biology + bare agent | 637a341 | 5969dd5 | 5 |
| O4 | GLM-5.3-Flash | gemot | 9faaf29 (4), ecfb848 (1) | 7d22712 | 5 |
| *Figure 4: 20 runs over 4 benchmarks* |  |  |  |  |  |
| O4 | GLM-5.3-Flash | gemot | ecfb848 | a56d77b | 20 |

## D Evaluation cohorts, scoring, and numerical protocols

### D.1 Evidence sources and version scope

The software description in Appendices A–C is anchored to source revision `e0d6921`. That revision is a description of the available implementation, not a claim that every reported run used the same code. The manuscript combines the run-level training export accompanying Figure 2, the expanded Fujita evaluation in Figure 3, and the additional validation systems in Figure 4. Their candidate selection, numerical settings, and denominators must be interpreted within each cohort.

As noted in Appendix A, the use of multiple software revisions reflects time and computational cost constraints. Unless specified otherwise, revision differences did not materially affect model construction, evaluation, or the reported conclusions. We will rerun all experiments using a single software revision upon publication.

### D.2 Cross-problem sweep summaries

The sweep summaries use the frozen run-level export `paper/generated/sweep_data.jso n`, with selection and exclusions described in Table 6. Runs started between 10 and 21 September 2026; the export retains job and archive identifiers, code revisions, fitting budgets, and candidate metrics. We used the recorded candidate summaries without reselecting model versions. The Fujita tasks and selected versions coincide with configuration O1 in Figure 3. The model-source annexes additionally contain the Fujita ablations (Appendix G). The biology-visible validation cohorts of Figure 4 are those systems’ Figure 2 runs and are listed in Appendix H; the 20 biology-blinded validation models are not listed.

Per-problem medians use available finite values independently for each metric; these marginal medians need not describe the same candidate. The generated L^A^T_E_X macros used within this manuscript and the unrounded `sweep_medians.csv` use the same observations. Training condition counts include conditions represented in the measurement table and can differ from the number of supplied experimental-design rows. Likelihood differences are not pooled across problems with different measurement and noise models. Per-run differences satisfy ΔBIC = 2ΔNLL + Δ*k* log *N*, where *N* is the number of training measurements and *k* is the parameter count used by the scoring implementation.

Recorded multistart budgets (Table 4) are 1,000 for Crauste, 200 for Bachmann, and 100 for the remaining problems. Token and fit counts come from run summaries; token counts include cached inputs. Wall time is full-task elapsed time. CPU time is measured consumption summed over workers in Slurm accounting, including local compilation and fitting but excluding remote LLM inference. All 90 selected runs have performance scores, token usage, and CPU accounting. The smaller/larger usage summaries in Section 2.2 pool all 65 and 25 runs, respectively, rather than problem-level medians. Missing fit counts for two Lucarelli runs are omitted from fit-count summaries only. Export provenance and numerical consistency checks are recorded in `paper/drafted/sweep_provenance.md`.

**Table 4:** Fit and wall budgets of the scored agent runs. , per benchmark as each run’s scheduler log states them (for the null-arms experiment, which kept no logs, as its submission manifest records them); the Figure 3 arms share one budget. *Starts*, multistarts per fit; *per-start cap*, the optimiser’s time limit per start; *wall budget*, the agent’s deadline; *fit workers*, parallel optimisation workers.

| Benchmark | Starts | Per-start cap (s) | Wall budget (h) | Fit workers |
| --- | --- | --- | --- | --- |
| *Figure 2* |  |  |  |  |
| perelson | 100 | 60 | 3.67 | 100 |
| crauste | 1000 | 60 | 3.67 | 100 |
| rahman | 100 | 60 | 3.67 | 100 |
| brannmark | 100 | 240 | 3.67 | 100 |
| boehm | 100 | 60 | 3.67 | 100 |
| armistead | 100 | 60 | 3.67 | 100 |
| fiedler | 100 | 90 | 7.33 | 100 |
| bruno | 100 | 60 | 3.67 | 100 |
| oliveira | 100 | 60 | 7.33 | 100 |
| sneyd | 100 | 60 | 7.33 | 100 |
| fujita | 100 | 90 | 6.67 | 32 |
| blasi | 100 | 60 | 3.67 | 100 |
| schwen | 100 | 120 | 7.67 | 100 |
| bachmann | 200 | 2400 | 33.3 | 100 |
| raimundez | 100 | 2400 | 33.3 | 100 |
| isensee | 100 | 12000 | 167 | 100 |
| alkan | 100 | 2400 | 33.3 | 100 |
| lucarelli | 100 | 1200 | 33.3 | 100 |
| *Figure 3* |  |  |  |  |
| all arms | 100 | 90 | 6.67 | 32 |
| *Figure 4* |  |  |  |  |
| perelson | 100 | 60 | 3.67 | 100 |
| crauste | 1000 | 60 | 3.67 | 100 |
| raimundez | 100 | 2400 | 33.3 | 100 |
| lucarelli | 100 | 1200 | 33.3 | 100 |

**Table 5:** Settings shared by the scored agent runs. A count after a value gives the number of runs with it; — marks a setting a run does not record. *Reasoning effort requested*, the setting each run’s requests carried; with none, the provider’s default applies, as OpenRouter’s model metadata listed it on the date shown. *Calls re-issued at low effort*, responses that spent the whole completion budget on reasoning and were retried.

| Setting | Value |
| --- | --- |
| Sampling temperature | 0.0 |
| Agent step limit | 500 |
| Output tokens per call | 131072 |
| Reasoning effort requested | none (provider default) |
| Calls re-issued at low effort | 0 (128), 1 (9), 2 (1), 3 (1), 4 (1) |
| Selection criterion | bic (138), — (2) |
| ΔBIC threshold | 10.0 (138), — (2) |
| OmniPath snapshot | 2026-08-11 (105), — (35) |
| Code-execution timeout (s) | — (130), 1344.0 (10) |
| Provider, GLM-5.3-Flash | z-ai (served by Z.AI) |
| Provider, GPT-6-Astra | openai (served by OpenAI) |
| Default reasoning, GLM-5.3-Flash | on, max effort (of low, high, max) |
| Default reasoning, GPT-6-Astra | on, medium effort (of low, medium, high, xhigh, max) |
| Defaults as listed by | OpenRouter model metadata, 2026-09-26 |

### D.3 Topological comparisons with the published models

We compared the selected candidate from each of the 90 sweep runs with its published reference. The scores in `paper/generated/sweep_data.json` were imported at manuscript revision `b4209e6`, with export stamp `d958571+dirty`. The definitions below follow the method description accompanying the earlier `51bd3d2` package; implementation inspection and the matched source examples refer to producer revision `5375241`. Current cohort selection, including the replacement Isensee run, is described in Table 6. These comparisons concern model structure, not the correctness of its biological interpretation.

**Table 6:** Evidence provenance and run selection. Infrastructure-failed attempts were excluded when they prevented a usable run or result; interrupted runs with a scored model were retained. Failures to construct a model were retained in the run denominators. Source stamps describe export environments; per-run code revisions describe the executions.

| Evidence | Scope | Provenance and selection |
| --- | --- | --- |
| Training comparison | 90 runs over 18 problems | Latest-started 5 runs per problem from the listed jobs, excluding infrastructure failures. An Isensee rerun replaces the earlier out-of-memory attempt; all selected runs have CPU accounting. Export source d958571+dirty, imported at manuscript revision b4209e6. |
| Fujita configurations | 30 runs, 28 models | Initial environment-activation failures before agent execution and associated cancellations were excluded; replacement batches entered the analysis. All 30 audited runs remain in the denominator, including two without a model. Final panel head or delivered bare-agent problem; run revisions 59cde21 and 5969dd5, export stamp 604449e+dirty. |
| Additional validation, O1 | 20 runs over four systems | Separate transfer and fixed-parameter evaluations; compact records in fujita_validation_deltas.csv. |

#### Weighted graphs of interventions and readouts

The topology analysis reported in Section 2.3 uses weighted graphs to compare connectivity and latent depth. Their nodes represent interventions and readouts, termed intervention and readout anchors. We first extract a directed, unsigned graph of the underlying state-variable influences. A reactant, modifier, or species read by a reaction’s rate law points to the species that the reaction changes. Rate rules contribute influences from their righthand-side dependencies to the integrated variable. Assignment dependencies are resolved except at retained anchors; self-edges and product-only influences are omitted. Signs, parameter values and stoichiometric magnitudes are not compared, and conservation laws are not reduced.

Each shipped observable formula names its readout, so observables that read the same readout form one readout anchor, and the species underlying each readout are aligned between models; the shipped condition table identifies interventions. A declared intervention enters the graph as the species whose initial value it sets, as the assignment-rule variable it feeds, or as the column itself where it appears only in a rate law. In the latter two cases the symbol is retained as a graph node rather than resolved into its own definition, so that its downstream influences are retained. Routes *into* exogenous interventions are not scored. Fifty-nine of the 90 candidates share at least one intervention anchor with their published reference. One further candidate has a declared intervention without a shared anchor; matching its remaining routes does not establish recovery of the intervention response.

#### Connectivity and latent depth

An edge in the weighted graph records a connection between an ordered pair of anchors whose underlying route does not pass through a third anchor’s species. Its weight is the shortest such route’s hop count, which quantifies latent depth between the anchors. Let *d_c_*(*A, B*) and *d_r_*(*A, B*) be the candidate and reference hop counts where a route exists, and let *R_c_* and *R_r_* denote the sets of reachable ordered pairs.

Pairs present in only one model are grouped by their initial route segment to reduce repeated charges for a shared branch reaching multiple readouts. Write Γ*_m_*(*P*) for the partition of a pair set *P* induced by the source anchor and the first node of model *m*’s selected shortest route; directly adjacent endpoint pairs form their own groups. These groups summarise structural differences, not independent biological hypotheses. The discrepancy is

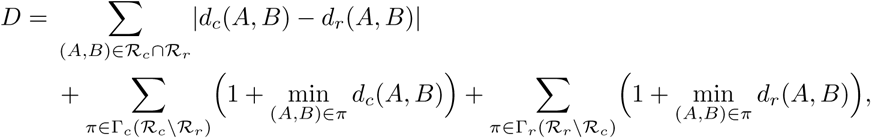

normalised by Σ*_π∈_*_Γ_*_r_* _(_*_Rr_* _)_ 1 + min_(*A,B*)_*_∈π_ d_r_*(*A, B*) . This is an exact comparison of the aligned anchor-route summaries, not an exact edit distance between the full reaction networks. Shared species can give zero-hop routes; the leading unit ensures that adding or removing such a route still has a cost.

We separate the score into added connections, increased latent depth on shared routes, deleted connections, and reduced latent depth on shared routes. Depth denotes the shortest route’s hop count, rather than a count of uniquely added or removed latent species: an intermediate can serve multiple routes, and readout mappings affect which species delimit them. The sum of these four normalised components equals the reported total, up to rounding.

A normalised comparison is available for 75 of the 90 candidates, across 15 problems. Rahman has only one readout anchor and no intervention anchors. Oliveira has two readout anchors, cumulative cases and deaths, but both are terminal readouts with no directed route between them and no intervention anchors. Their reference route summaries therefore have zero normalising denominators; these comparisons are undefined and excluded rather than counted as matches.

Perelson also has a single readout, viral load, and its published reference encodes treatment implicitly rather than through an explicit drug-input variable. The current export has no intervention anchor for Perelson, so all five comparisons have zero normalising denominators and are undefined.

Of the 75 defined comparisons, 49 differ in connectivity and seven only in latent depth; 19 match both summaries. Added connections occur in 38 candidates and deleted connections in 41, with overlap. All 25 candidates for the larger problems add connections, although 23 use fewer states, one the same number, and one more than its reference. In contrast, the nonzero Fiedler and Sneyd comparisons change only latent depth. These distinctions are concealed by the total. Figure 2 and Table 7 therefore report the components separately. The normalised total is not bounded by one: six candidates exceed one (maximum 1.36), and longer shared routes alone can produce such a score without adding any connection.

**Table 7:**
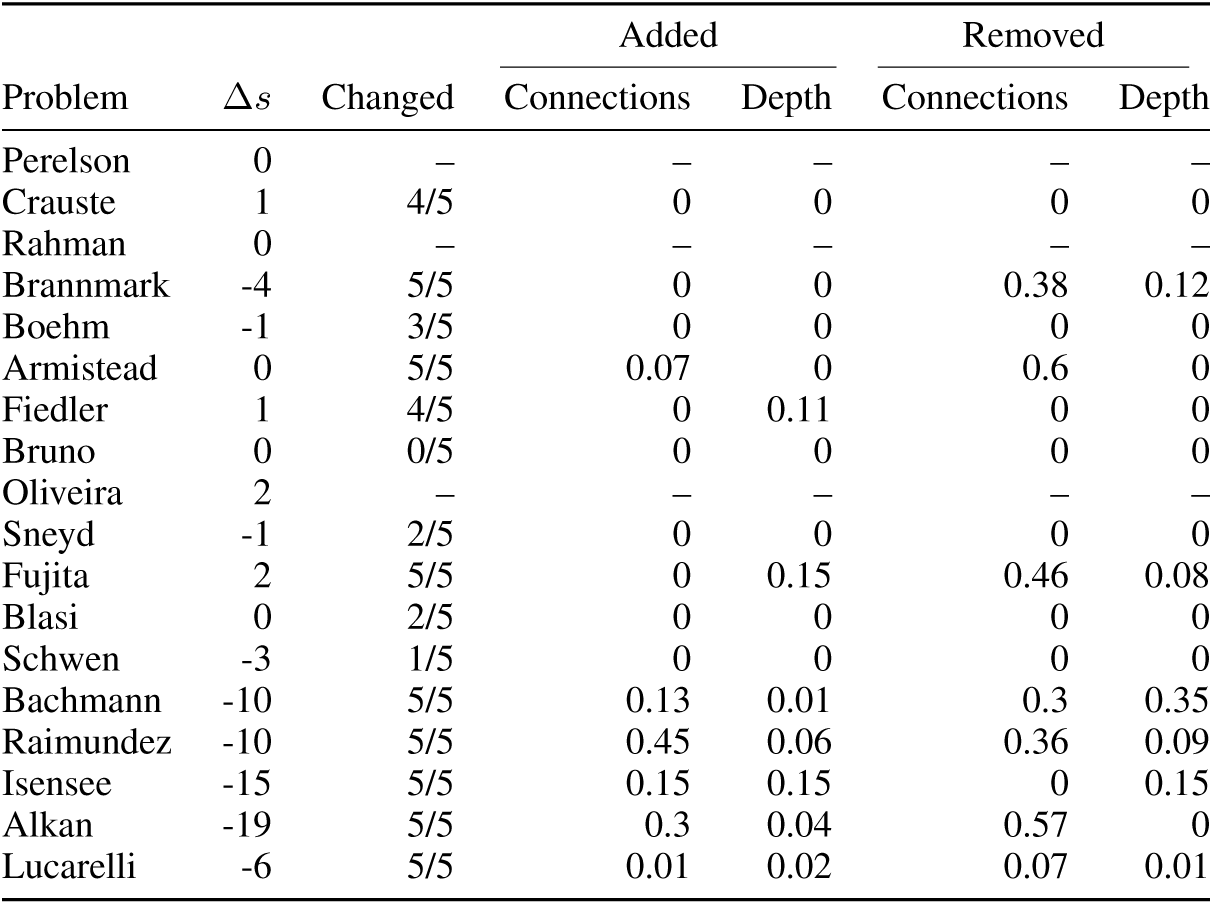
Connectivity and latent depth in the training sweep. Each problem has five selected candidates. Δ*s* is the median candidate-minus-reference state count; “Changed” counts candidates whose connectivity or latent depth differs from the reference among defined comparisons. The last four columns give median weighted contributions from added connections, increased latent depth, deleted connections and reduced latent depth, using the reference-summary denominator defined above. They are not fractions of recovered biological interactions. Separate component medians need not sum to the median total. A dash denotes an undefined normalised comparison.

#### Further structural examples

Matching connectivity and latent depth does not imply that the underlying models are identical. For example, Schwen candidate 3 (Listing 86) uses eight states instead of the reference’s eleven while retaining both connectivity and latent depth, with training ΔBIC = −131.78. Four Schwen candidates match both connectivity and latent depth; this includes candidate 5 (Listing 88), whose declared insulin-dose intervention enters through `E = init_Ins` and binding reactions `bind1` and `bind2`, although `init_Ins` has no shared anchor in the exported comparison.

Raimundez candidate 1 (Listing 94) adds MAPK-dependent inhibition of EGFR activation. Reaction `egfr_actL` converts `E_b` to `E_p` at rate `a_E*E_b/(1 + f_eEb*M_p)`, with `M_p` identified by `readout_pMAPK := M_p`. The coefficient `f_eEb` is unset in the listing but equals 39.91 in the saved parameter table, `paper/model_sources/benchmark/raimundez_1.parame ters.tsv`. This positive coefficient makes the activation rate decrease with MAPK; the reference influence graph has no MAPK-to-EGFR route. This candidate has four fewer states and training ΔBIC = −696.68. The signed interpretation comes from the rate law and fitted coefficient, not from the unsigned topology score.

#### Relation to Fujita topology recovery

The recovery counts in Figure 3C (Appendix D.5) test whether each model contains the three forward connections EGF→pEGFR→pAkt→pS6, with each route excluding other intervention or readout anchors. Additional forward connections and different latent depths are permitted; backward connections are not assessed. The sweep instead scores added and removed connections and changes in latent depth across all ordered anchor pairs, including feedback between readouts but excluding routes into interventions. Both analyses use EGF as the intervention anchor for the five shared Fujita candidates.

The numerical path extraction also differs from the sweep and the equation-based SBGN maps in Appendix F. The Fujita extractor treats products as sources of influence on other reaction participants even when the rate law does not read them. For example, reference reaction `v3_reaction_3`, `pEGFR_Akt`→pEGFR+pAkt, creates a pAkt→pEGFR link solely because the two species are coproducts. The sweep omits this link unless supported by a reactant, modifier or rate-law dependency. This rule can alter connectivity and shortest path lengths. Consequently, recovery in Figure 3C does not imply zero discrepancy in Figure 2; neither establishes biological causality or dynamical equivalence.

#### Mechanistic interpretation of added connections and latent intermediates

We inspected the equations and saved parameter tables for six systems, including the examples discussed in Section 2.3. Table 8 identifies representative candidates with added connections or increased latent depth. These examples illustrate different interpretations of the scores, rather than estimating how often agents recover supported mechanisms. Signs below follow the fitted equations, not the unsigned graphs. The literature assessment concerns each specific proposal, not validation of the whole candidate. Candidate reaction and assignment-rule names below identify the corresponding entries in the linked listings. Sources are listed in Appendices H and G; the Fujita example belongs to the shared training-sweep cohort, without using its held-out performance for selection.

**Table 8:**
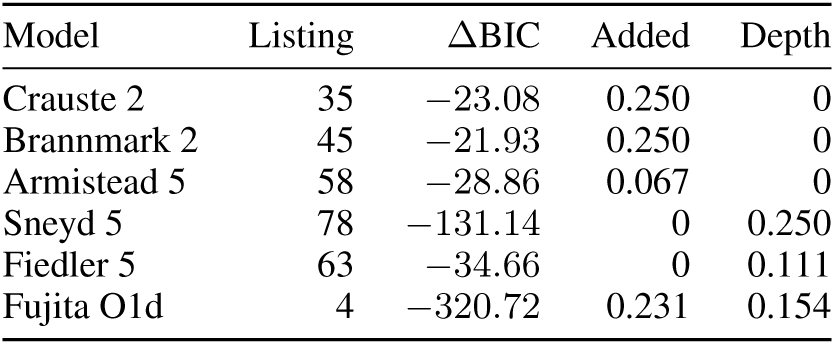
Inspected examples from the training sweep. Added and Depth are the normalised contributions from added connections and increased latent depth, rounded to three decimals; other components may also be nonzero. All selected examples improve training BIC. Listings identify the archived models and retain their task and version provenance.

**Table 9:** Fujita configurations and denominators. All configurations have five audited runs. “Topology” counts models containing every required forward readout connection among delivered models; “Improved” counts complete held-out scores below the stored reference.

| Code | Configuration | Models | Complete | Improved | Topology |
| --- | --- | --- | --- | --- | --- |
| O1 | GLM panel, curator on | 5 | 5 | 4 | 5/5 |
| O2 | GLM panel, curator off | 5 | 4 | 3 | 5/5 |
| O3 | GLM bare agent | 3 | 3 | 2 | 3/3 |
| O4 | GLM panel, biology blinded | 5 | 4 | 3 | 1/5 |
| F2 | Astra panel, curator off | 5 | 5 | 5 | 5/5 |
| F3 | Astra bare agent | 5 | 5 | 5 | 5/5 |

#### Crauste (reference: **Figure 34**): pathogen-dependent memory differentiation

Candidate 2 (Listing 35) makes late-effector-to-memory conversion pathogen-dependent through reaction e2_mem, with flux *v_L→M_* = *c*_fate_(1 −*a_P_*)*L* = *c*_fate_*α*_act_*L/*(*α*_act_ + *P*) instead of the reference’s constant multiple of *L*. Assignment rule `a_P` defines *a_P_* = *P/*(*α*_act_ + *P*), with *P* = `Pathogen`, *L* = `LateEffector`, *α*_act_ = `alpha_act` and *c*_fate_ = `c_fate`. The saved fit gives *α*_act_ = 1.2332 and *c*_fate_ = 0.09673. Early effectors clear pathogen through `p_clear`, so this creates an early-effector–pathogen–memory route that avoids the measured late-effector anchor. The hypothesis is release of suppression as pathogen declines, not a direct early-effector-to-memory lineage. Candidate 3 (Listing 36) uses the same qualitative gate in `l_to_m`, but also sends 9.6% of modelled naive-cell activation directly to late effectors through `n_to_l`, alongside the earlyeffector route `n_act`; that shortcut has less support from the reference study’s predominantly linear ontogeny (Crauste et al., 2017). Vaccinia experiments in Sarkar et al. (2008) separated cognate antigen from infection-associated stimuli and found that shortening late antigenic stimulation favoured memory-associated phenotype and function. This supports the gate’s sign, not its exact kinetics or a larger final memory population. Moreover, antigen can persist after infectious-virus clearance (Tamburini et al., 2014); the single pathogen variable is an approximation to antigenic stimulation.

#### Brannmark (reference: **Figure 39**): a shared response approximation

Candidate 2 (Listing 45) assigns *b*_IR_ = *R_p_* and *b*_IRS1_ = 1 + *K_d_R_p_* in `readout_IR_P` and `readout_IRS1_P`, with fitted *K_d_* = 16.0742, before assay-specific observation scaling. Candidate 4 (Listing 47) uses the same readout names, with *R_p_* and *R_p_* + *r*_0_ via `IRS1p_lin` (*R_p_* is `Rp` and `IRp`, respectively). Both therefore force the two biological signals to share a temporal shape, rather than supplying separate IRS1 dynamics. Insulin reaches their shared state directly through `receptor_activation` and `ir_activation`, respectively, creating an additional insulin–IRS1 anchor route; in the reference, that route passes through the IR readout anchor. This is a testable reduction of the readout mapping, not receptor-independent IRS1 activation. The primary adipocyte study supports receptor trafficking together with downstream feedback (Brännmark et al., 2010), but does not establish affine equivalence of the two signals. Candidate 2 retains trafficking (`receptor_internalization` and `receptor_recycling`) while omitting downstream feedback; candidate 4 retains a delayed inhibitory variable (`x2_prod`/`x2_deg`, acting through `ir_dephos`) but no explicit internalisation. Such differences make the published trafficking interventions relevant discriminating evidence.

#### Armistead (reference: **Figure 35**): genotype-dependent ceramide supply or clearance

All five candidates introduce a direct genotype effect on a ceramide rate. The reference already connects genotype to ceramide through measured sphinganine; the added anchor route bypasses this intermediate. In candidate 5 (Listing 58), the reaction named `s_salvage` adds *k_cs_*(1 + *g*_on_*S*_on_) independently of sphinganine, with *k_cs_* = 0.60754 and *g*_on_ = −0.48799 (parameter `g`). Because *S*_on_ is one in wild type and zero in the Hai1a-deletion mutant, this source is about 49% lower in wild type. Candidate 1 (Listing 54) instead changes clearance through `cer_exit`; together with the source `csl_to_cer`, this gives *Ċ* = 0.03145−0.22143*S*_on_*C* for *C* = `Cer`. These are distinct kinetic hypotheses despite sharing the added unsigned connection. In the original zebrafish study, serine palmitoyltransferase inhibition and ceramide-synthase knockdown supported de novo synthesis, whereas sphingomyelinase inhibition did not alleviate apoptosis (Armistead et al., 2024). This evidence favours de novo supply and does not validate a sphinganine-independent source or altered clearance. The unassigned source may instead absorb omitted processes in the restricted training data.

#### Sneyd (reference: **Figure 36**): priming before pore opening

Candidate 5 (Listing 78) contains the route IP_3_ → *g*_primed_ → *g*_open_ through J_prime and J_gate, with states gprimed and gopen. Their fluxes are *k_pr_f*_IP3_(1 − *g*_primed_ − *g*_open_) and *k_pr_g*_primed_, respectively, with fitted 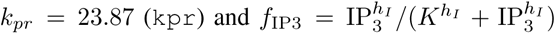 (assignment rule `fip3`). The primed state is absent from `readout_open_probability`, giving an explicit nonconducting intermediate. The reference input directly influences a readout-contributing gating state, so this adds one hop while using fewer declared states overall. Partial-agonist and mutational experiments support conformational coupling between ligand binding and pore opening (Rossi et al., 2009), although principally in IP_3_R1 rather than the benchmark’s type-2 receptor. They do not identify this fitted gate. Ligand-step experiments also constrain strict binding-order interpretations (Mak et al., 2007). The reference already describes dynamic activation and eliminates fast intermediates (Sneyd & Dufour, 2002); added depth resolves these dynamics differently, rather than establishing a previously unknown receptor state.

#### Fiedler (reference: **Figure 37**): feedback with a finite response time

Candidate 5 (Listing 63) replaces direct ERK inhibition of RAF activation by an inhibitory state *w*: fb_charge and `fb_decay` yield *ẇ* = *k_a_*(pERK^2^ − *w*), while `raf_act` makes RAF activation proportional to 1*/w*. The fitted *k_a_* = 0.32935 (`k_adapt`) gives a relaxation time of approximately 3.04 model-time units. Together with `mek_phos`, the return route becomes pERK → *w* → pRAF → pMEK, one hop longer than the reference. The hypothesis is that RAF sensitivity depends on recent ERK activity, rather than only its instantaneous abundance. ERK-dependent inhibitory RAF phosphorylation and subsequent recovery provide an experimental precedent (Dougherty et al., 2005). They do not identify *w* as a distinct protein, validate reciprocal inhibition, or establish its timescale in the benchmark’s HeLa experiments (Fiedler et al., 2016). Positive ERK–RAF feedback has also been reported under different cellular and stimulation conditions (Balan et al., 2006), further limiting identification of *w* with a specific RAF phosphoform. Candidate 4 (Listing 62) also introduces a feedback mediator `Xinh` through `feedback_recruitment` and `feedback_decay`, with inhibition encoded by assignment rule `Ffb`. Part of its additional depth comes from moving Sorafenib’s action from reference MEK phosphorylation (`v3_v_2`) to `raf_activation` through assignment rule `I_sor`, which does not enter `mek_phosphorylation`, showing how pharmacological abstraction can affect the score.

#### Fujita (reference: **Figure 43**): resolving mTORC1 and S6K

Candidate O1d (Listing 4) inserts the three-hop route pAkt → *M* → pS6K → pS6 through mtorc1_act, s6k_act and s6_phos; the first reaction identifies *M* as mTORC1 activity. The first two activation rates are *V_M_* pAkt(1 − *M*)*/*(*K_M_* + pAkt) and *k_M_ M* (1 − pS6K); the saved coefficients are positive (*V_M_* = 1.17 10*^−^*^7^, *K_M_* = 0.04119 and *k_M_* = 1.3212; `Vmax_M`, `Km_M` and `k_M`). Akt regulation of TSC2 and mTOR-dependent S6K activation support this pathway ordering (Inoki et al., 2002; Brown et al., 1995), although the candidate still lumps upstream regulation and does not establish these kinetics in EGF-stimulated PC12 cells. The reference Akt readout includes the Akt–S6 complex, which reaches pS6 in one hop through `v6_reaction_6`: the two-hop increment therefore reflects both an explicit cascade and a changed readout mapping. O1c (Listing 3) goes further, adding serial `S6K2`/`S6K3` stages through `s6k2_act`/`s6k3_act`, without established molecular identities. Such stages can implement filtering consistent with the original study’s functional observation (Fujita et al., 2010), but their names and depth do not demonstrate recovery of additional biological actors.

### D.4 Training fit per model version

Figures 5 and 6 follow every scored run through its model versions. At each version a run contributes the best training ΔBIC it had fitted so far, carried forward once it stops; a panel shows the median over its runs as a line and the minimum–maximum range as a band. These curves include all fitted versions, without applying the final noise-plausibility check. The judge instead scans versions in BIC order and can skip those whose fitted noise is flagged as implausibly inflated or collapsed (Appendix B.10). It therefore can select a higher-BIC model; if every version is flagged, it falls back to the lowest-BIC version. The delivered model differs from the raw minimum in 9 of the 90 Figure 2 runs (four of the five on Alkan). The trajectory export does not retain individual selection reasons, so it does not establish that this filter explains all nine discrepancies. Each run’s delivered ΔBIC, the value its figure plots, is therefore marked where the run ended. The two bare-agent arms of Figure 3 (O3, F3) archive only the model they delivered, so they are shown as their delivered end state.

### D.5 Fujita configurations and fixed-parameter prediction

Figure 3 combines three audited Fujita experiments, comprising 30 runs across 6 configurations. The first experiment used revision `59cde21`; the later bare-agent and Astra experiment is associated with `5969dd5` in its submission manifest, and the biology-blinded re-run used `7d22712`, which appends the stimulus protocol to the blinded task (described below). We verified that the other differences between these three run-code revisions had no impact on agent behaviour. These run revisions differ from the export stamp `604449e+dirty`, preserved in manuscript commit `16a2651`. Construction used 144 measurements from six constant-dose conditions; fixed-parameter evaluation used 276 withheld measurements from six pulse and four ramp conditions (Fujita et al., 2010). The published dynamical model was not supplied to the agents. The experiment records specify 32 CPUs, 100 fitting starts, and a 90-second per-start limit.

We ran these experiments with `z-ai/glm-5.3-flash` and `openai/gpt-6-astra`. The figure uses C and F for these models, followed by a configuration number and a run letter. Display names differ from archive labels: O1a–O1e map to B1–B5, O2a–O2e to O1–C5, O3a–O3e to G1– G5, O4a–O4e to X1–X5, F2a–F2e to P1–P5, and F3a–F3e to N1–N5. O3a and O3e delivered no fittable model. Panel runs contribute their recorded final `head_version`; bare-agent runs contribute the delivered PEtab problem, represented by the sentinel version −1 in the CSV. Neither selection uses held-out performance.

O1 includes the biocurator and its external knowledge tools. O2 and F2 remove that role, its tools, and the lead’s curator guidance, while retaining biological identifiers, assay descriptions, and the three modelling analysts. O3 and F3 use a stock agent with file tools and a sandboxed shell to author and fit a PEtab problem directly. They retain curated biological knowledge tools and scenario information, but lack the panel’s MCP modelling operations, version store, specialist analysts, and finish gate. The comparison therefore changes a collection of modelling and coordination facilities rather than isolating knowledge retrieval. The configurations are not a complete factorial design: the Astra curator-on (F1) and biology-blinded (F4) arms were not run because of cost.

Biology-visible configurations receive the time-dependent EGF rule and guidance to use EGF in receptor binding. Biology blinding anonymises identifiers, removes descriptive metadata and replaces the biological task with numerical instructions. The current implementation (revision `7d22712`) preserves the experimental input protocol: it reads the rules from `inputs.tsv`, translates stimulus and condition-input identifiers to anonymous aliases, and appends them to the blinded task. For Fujita, the rule specifies

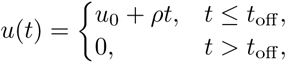

where *u*_0_, *ρ* and *t*_off_ are fixed condition inputs. The prompt requires the agent to implement the assignment rule exactly and use *u*(*t*) wherever the stimulus enters the model, rather than substituting its parameter columns or estimating them. This supplies the prescribed step, pulse and ramp waveform definitions without revealing the stimulus’s biological identity or receptor-binding role. Rules may reference only their declared condition-input columns, time and mathematical functions; both preparation and prompt assembly validate the allowed symbols. The driver constructs the task from this validated input, discarding caller-supplied scenario prose. A private identifier mapping supports evaluation. Blinding does not remove pretrained biological knowledge, and enabling external tools does not establish which sources a particular run consulted.

We designed both pipelines to avoid an explicit incentive to optimise held-out performance. The supplied objectives concern training fit and BIC (Listings 153, 120 and 124); held-out evaluation is neither announced as an objective nor returned as construction feedback. The panel tool allowlist excludes the held-out evaluator. The bare agent sees a staged problem in a sandbox, with reference files withheld; its scoring server is started only after delivery. In both pipelines the judge evaluates held-out data after construction. We cannot exclude that an agent inferred later evaluation from the supplied experimental design.

#### Scoring and numerical availability

The archived parameters, including observation scales, remain fixed for Fujita evaluation (Appendix E.1). We rescored all delivered Fujita models with a script that varied solver tolerances and integration step sizes; the results were consistent across these settings. O2c and O4b lack a complete score over the ten conditions. Together with the two runs that delivered no model, they leave 26 complete held-out scores among 30 audited runs. An incomplete integration receives no joint likelihood over a reduced subset of measurements. It counts as predicting worse than the reference: it ranks worst in the held-out medians and ranges, and counts against its configuration in Figure 3D. A model without a held-out score still contributes its training ΔBIC to the training summary, whereas a no-model run contributes to neither.

All differences are candidate minus reference. The reference offsets for ΔNLL are 66.5807726637496 for training and 4156.3460971713175 for held-out evaluation. Training ΔBIC uses the committed reference with *k* = 20 and *N* = 144; the export corrects only its reference offset, with every recorded adjustment equal to zero. The held-out reference uses a stored score.

#### Readout topology and structural features

For each forward intervention/readout pair, the topology audit searches for a directed influence-graph path that does not pass through the species of another intervention or readout. The reference contains three adjacent connections under this rule. Inclusion requires all three, permitting additional connections and different latent-path lengths. Of 28 models, 24 satisfy this criterion; the two runs without a model are reported separately. This unsigned comparison does not test edge signs, reaction identities, rate laws, backward feedback, or the complete latent mechanism, and does not reduce conservation laws. Panel B illustrates the reference and two candidates selected for having the lowest held-out ΔNLL. That illustrative selection is retrospective. Its arcs all use one activity-flow terminator and are not sign-tested; they are separate from the unsigned recovery criterion used in panel C.

Depths count shortest directed influence-graph hops between implemented intervention/readout anchors. Assignment dependencies are expanded and readout rules add no hop. The blinded intervention anchor is the anonymous ligand/dose species mapped from `EGF_0`; other models use their implemented EGF stimulus. These are propagation proxies, not exclusively counts of unobserved intermediates. Reaction count includes all SBML reactions. State count includes dynamical species and rate-rule variables before conservation reduction, excluding constant, boundary, and assignment-defined species. Charged parameter count includes estimated observation scales and charged hardcoded literals.

#### Predictive success and rank association

Panel D counts models with held-out ΔNLL *<* 0 among those with complete scores, irrespective of topology. The same complete-score population underlies the Spearman correlations between training ΔBIC and held-out ΔNLL reported in Section 2.4: 16 GLM models across O1–O4 and ten Astra models across F2–F3. O3a and O3e produced no model; O2c and O4b lack complete held-out scores. We correlate the ranks of the two quantities within each language-model sample, using average ranks for ties, without covariate adjustment or topology filtering. The resulting coefficients are *ρ* = −0.14 for GLM and *ρ* = 0.30 for Astra. These pooled-configuration associations are descriptive and do not establish causal effects of model complexity.

#### Cost accounting

Panel E divides each configuration’s total expenditure estimate across all five audited runs by its number of fittable delivered models. Runs with no model are charged, and models with incomplete held-out integration remain in the denominator. Both models use their recorded usage amount, so the language-model component is the provider’s invoiced spend with no list-price correction applied.

The CPU component values Slurm `TotalCPU`, measured in core-seconds, at the exported `c7i.8xlarge` Linux on-demand rate in `us-east-1`: $1.428 per 32-vCPU hour, quoted 12 September 2026. It is computed as

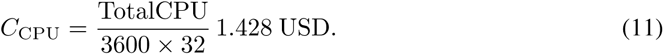

This prorates actual CPU use; it does not price an allocated instance for the full elapsed time or measure a cloud invoice. Storage, data transfer, and reservation or spot discounts are excluded. Under this convention the 30 Fujita runs total $130. The separate experiments in Figure 4 are outside this cost total.

#### Held-out predictions per arm

Figures 7–12 draw every model of Figure 3 on the two heldout protocols, the one-minute EGF pulse at six doses and the EGF ramp at four rates, at its stored parameters and the `stricter` solver setting. The first column of each figure is the published model at the collection’s nominal parameters; every other column carries the training ΔBIC and held-out ΔNLL that Figure 3 plots for that model. O4b integrates on no held-out condition, and O4e, which reads the condition inputs directly instead of implementing the supplied stimulus rule, predicts no response to the ramps; the other three biology-blinded models respond to both protocols.

**Figure 5:**
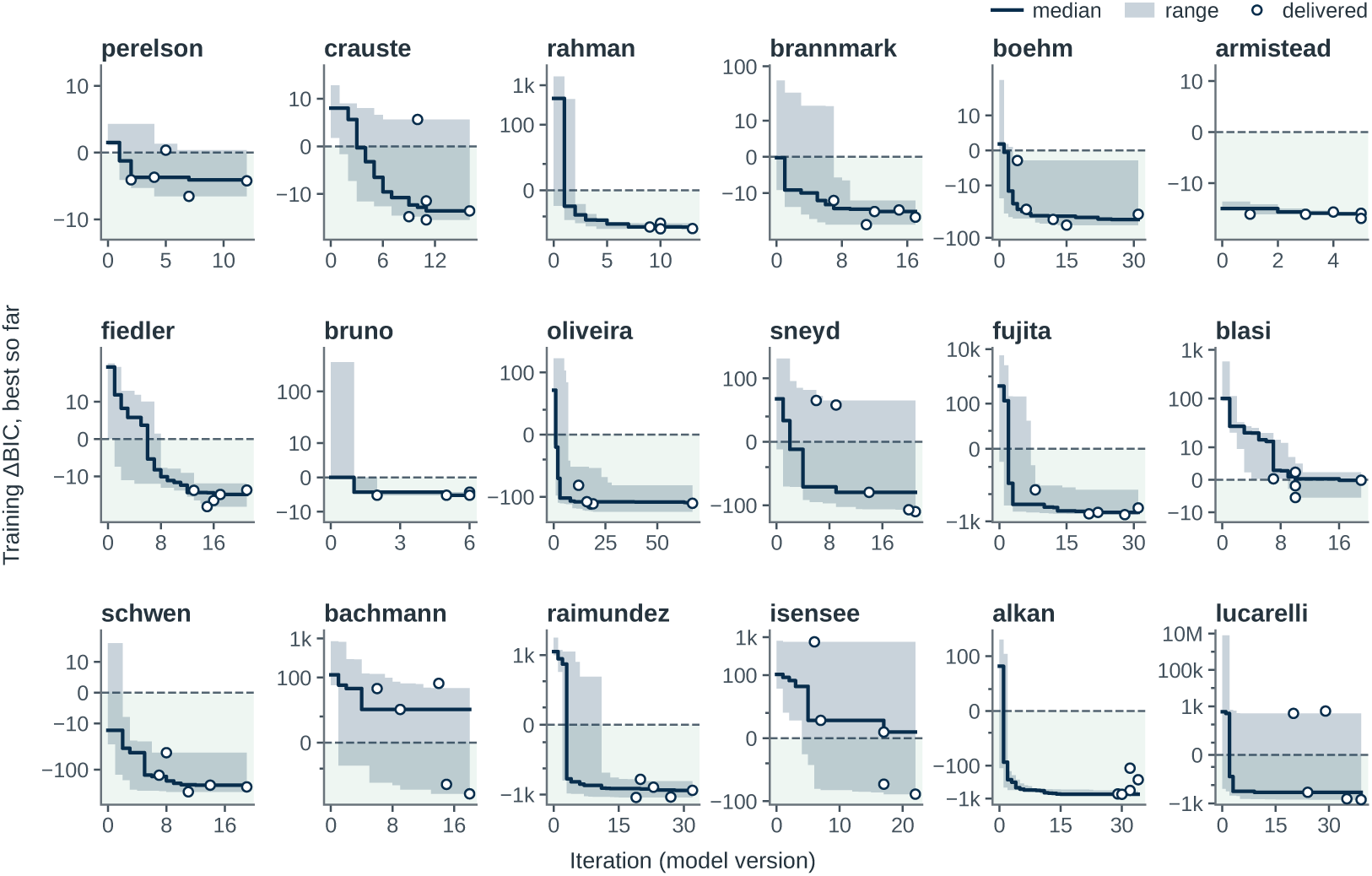
Training ΔBIC per model version, Figure 2 runs. One panel per benchmark, five runs each, on its own scale. Line, median best-so-far training ΔBIC against the committed reference; band, range over runs; open marks, each run’s delivered model at the version it ended on. Symmetric-log axis, linear within ±10; the shaded region is better than the reference.

**Figure 6:**
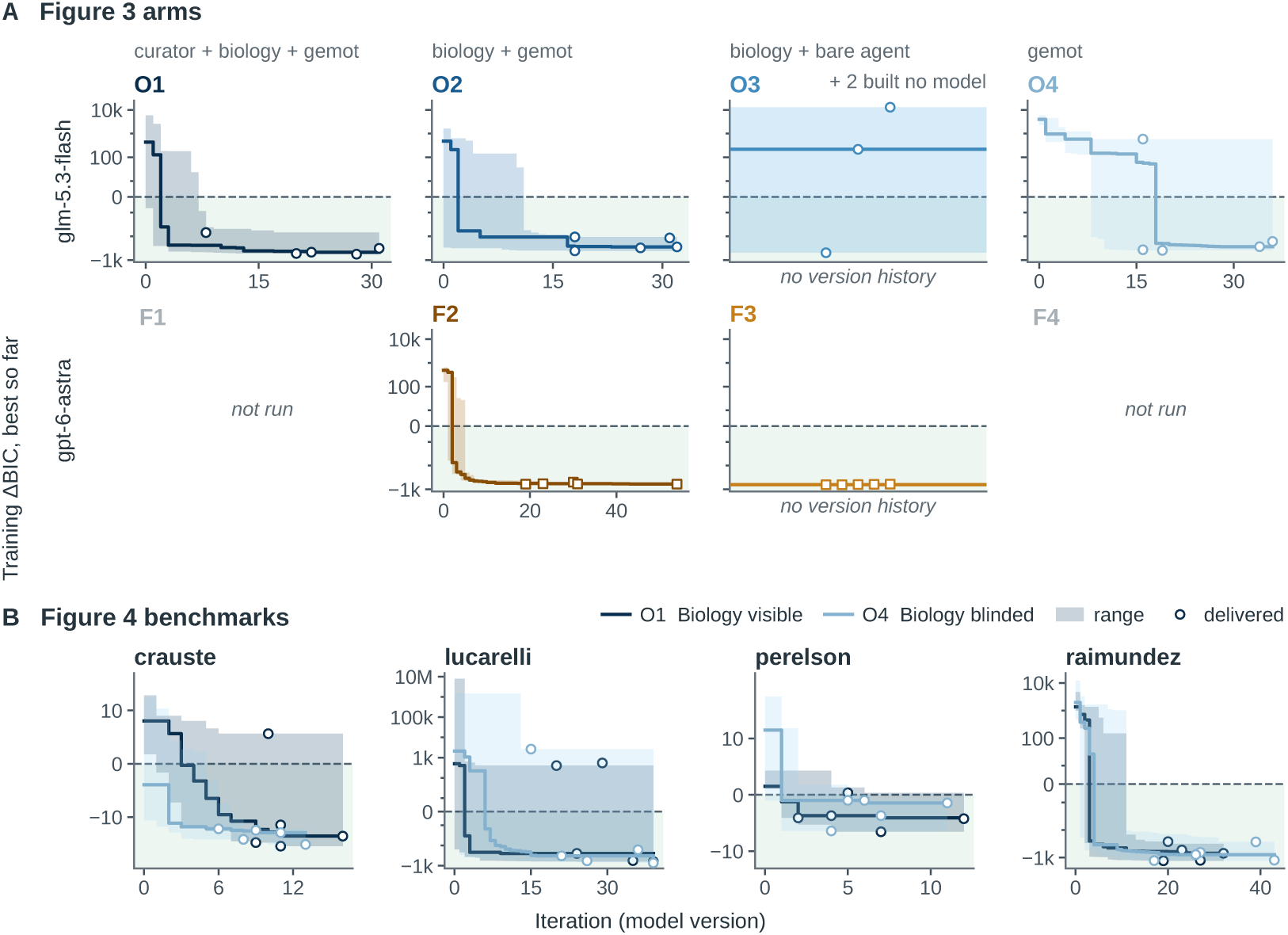
Training ΔBIC per model version, Figures 3 and 4. (A) The Fujita arms by language model (rows) and support rung (columns, in Figure 3’s colours); O3 and F3 keep no version history and show their delivered models only; F1 and F4 were not run. (B) The four Figure 4 systems, biology-visible (O1) and biology-blinded (O4) runs overlaid. Encoding as in Figure 5.

**Figure 7:**
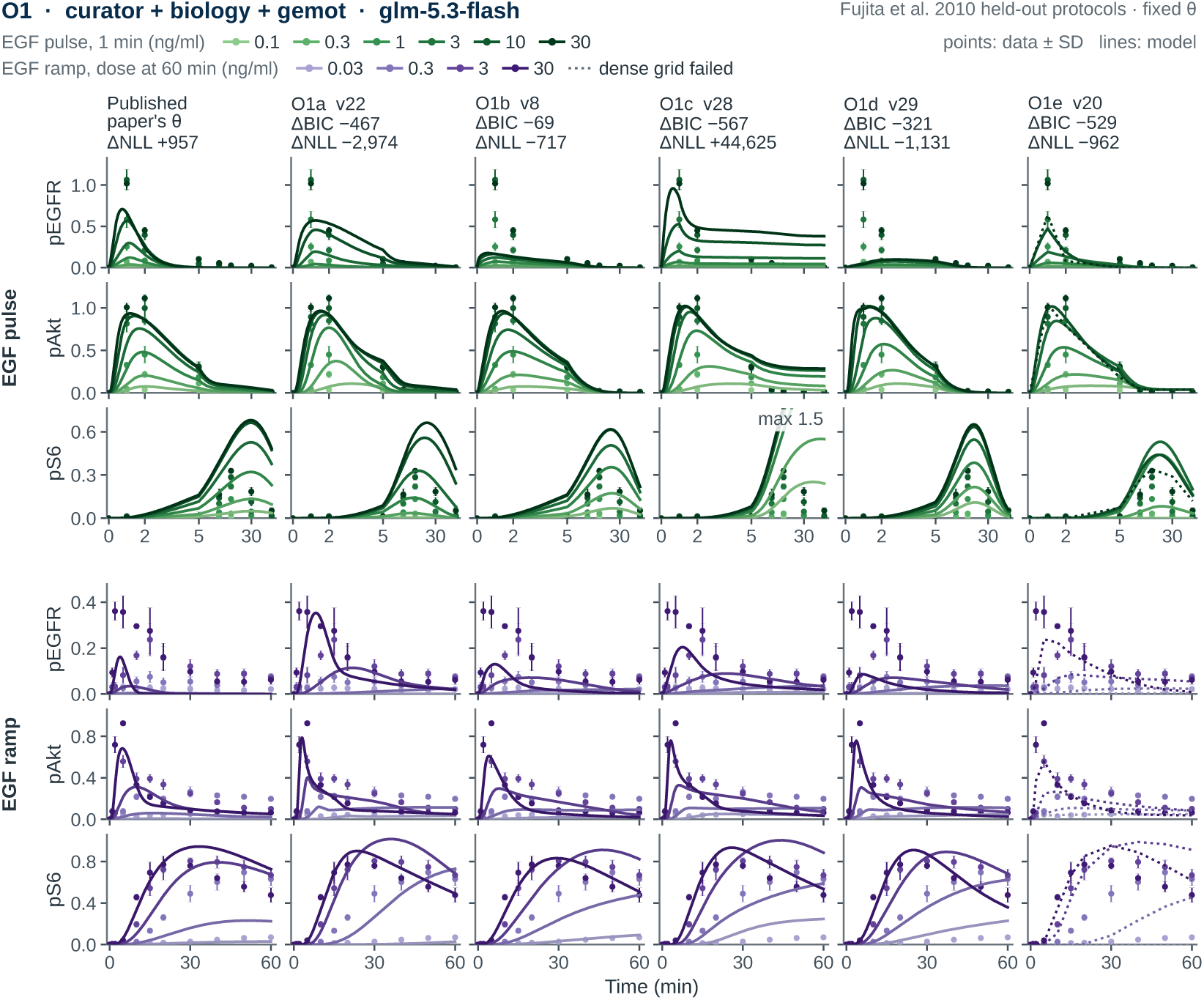
Held-out Fujita predictions, O1. (curator, biology and gemot; GLM-5.3-Flash). Rows, the pulse block then the ramp block, three observables each; columns, the published model at its nominal parameters, then O1a–O1e. Points, data ± SD; lines, each model at its stored parameters; one sequential scale per protocol, lightest for the weakest stimulus. Rows share a scale; a prediction leaving it is clipped with its maximum printed. Dotted, a condition that integrated at the measured times but not on the drawing grid.

**Figure 8:**
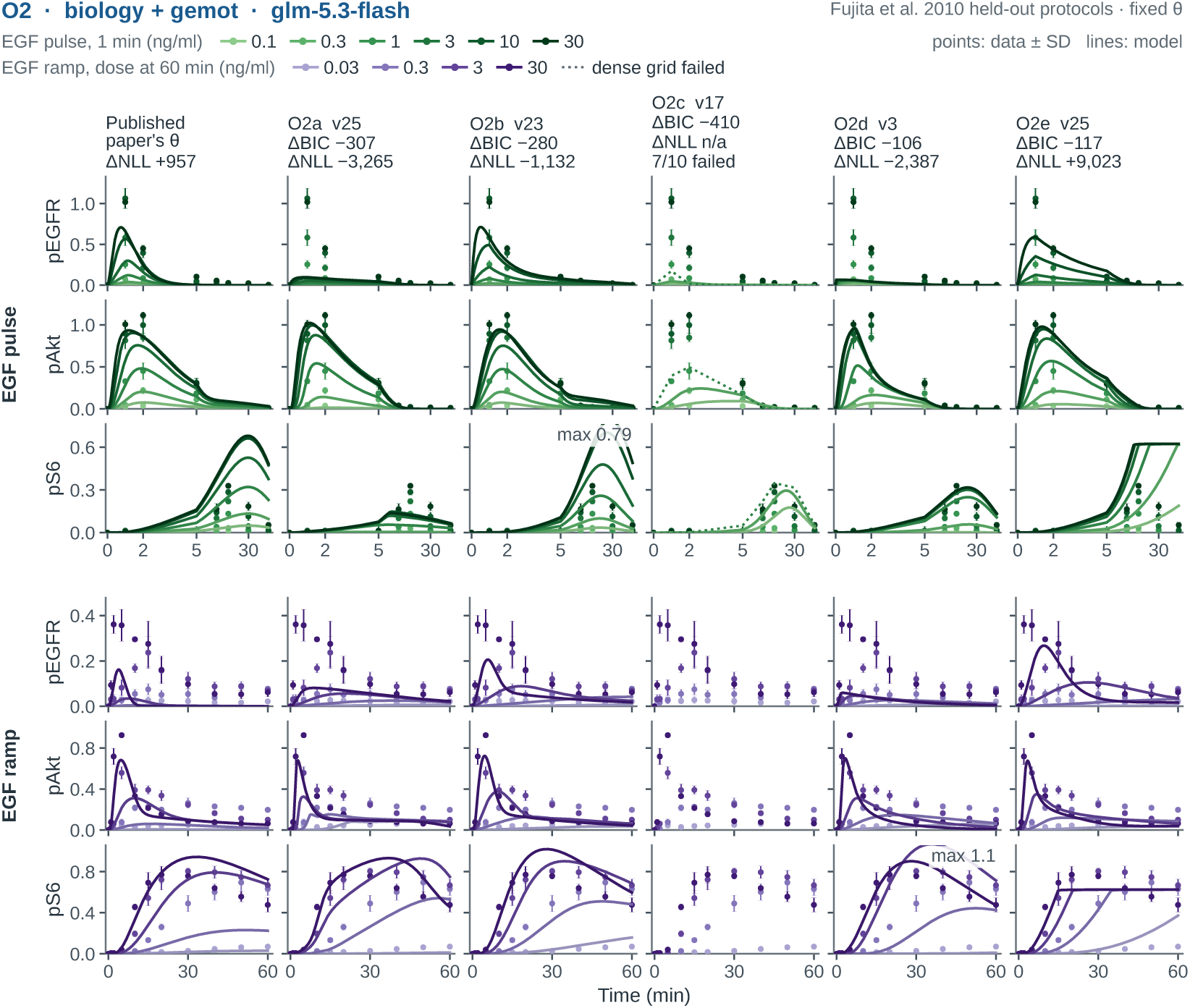
Held-out Fujita predictions, O2. (biology and gemot; GLM-5.3-Flash). As Figure 7; columns, the published model, then O2a–O2e.

**Figure 9:**
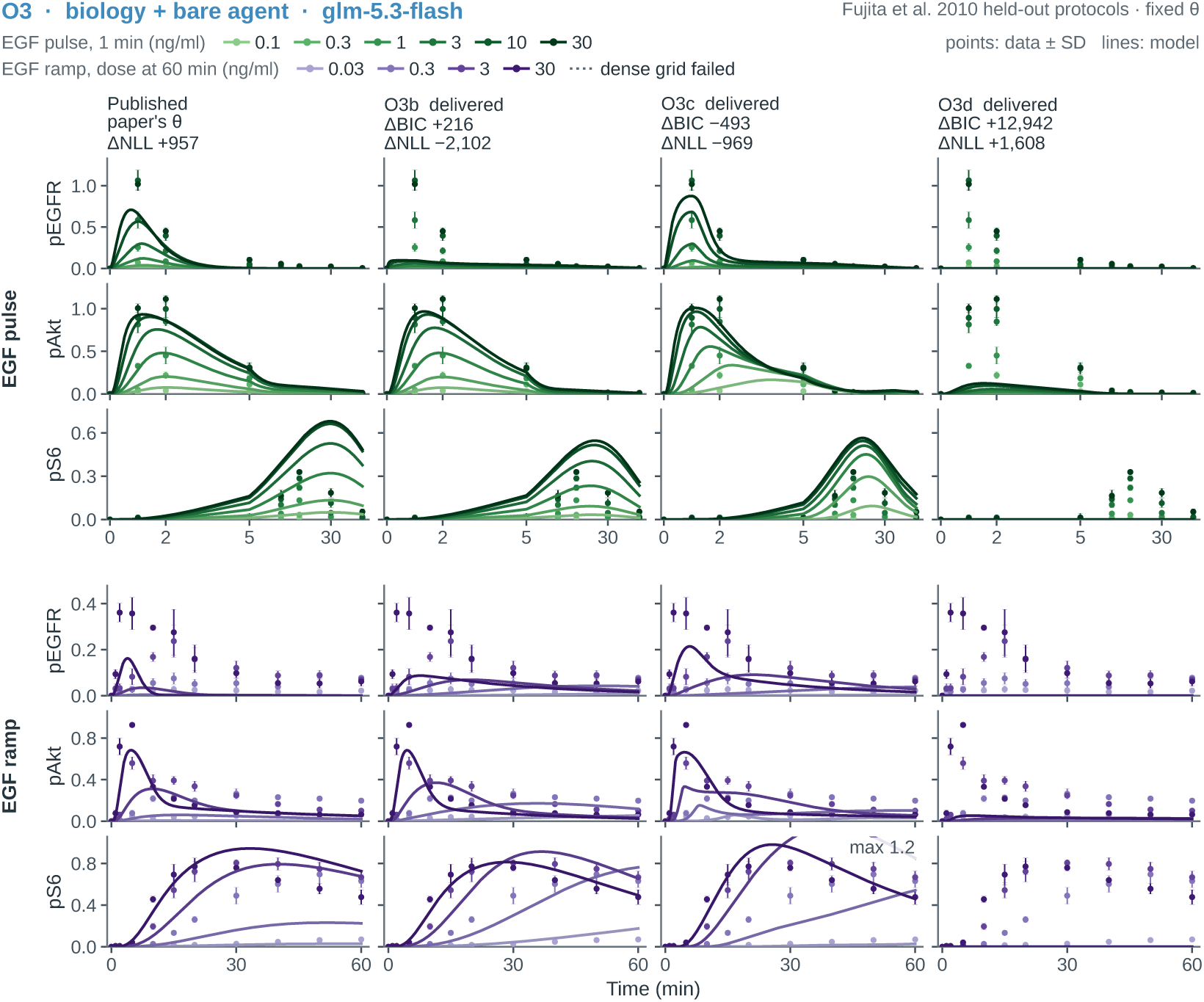
Held-out Fujita predictions, O3. (biology and bare agent; GLM-5.3-Flash). As Figure 7; columns, the published model, then O3b–O3d; two O3 runs delivered no model.

**Figure 10:**
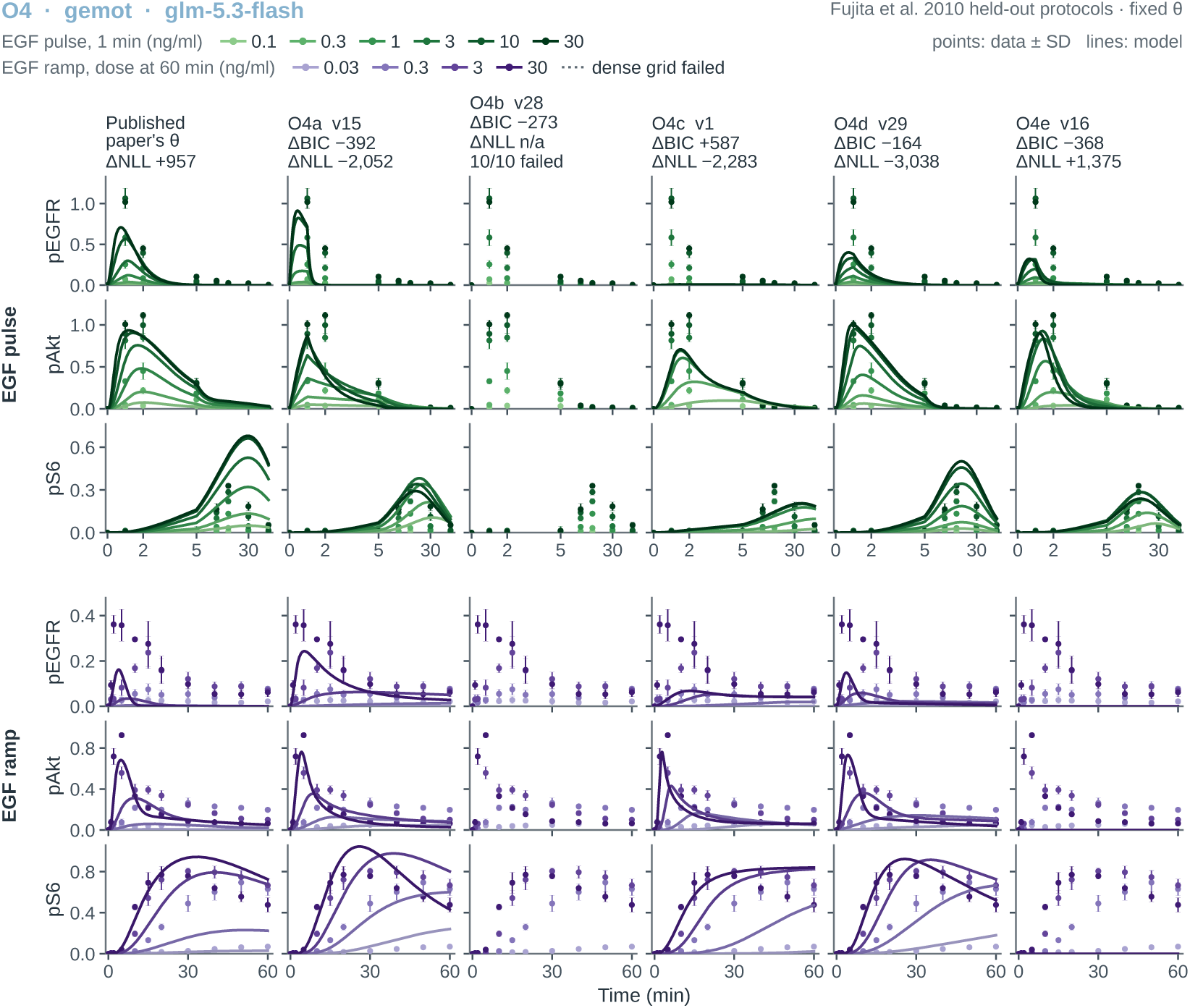
Held-out Fujita predictions, O4. (gemot, biology blinded; GLM-5.3-Flash). As Figure 7; columns, the published model, then O4a–O4e.

**Figure 11:**
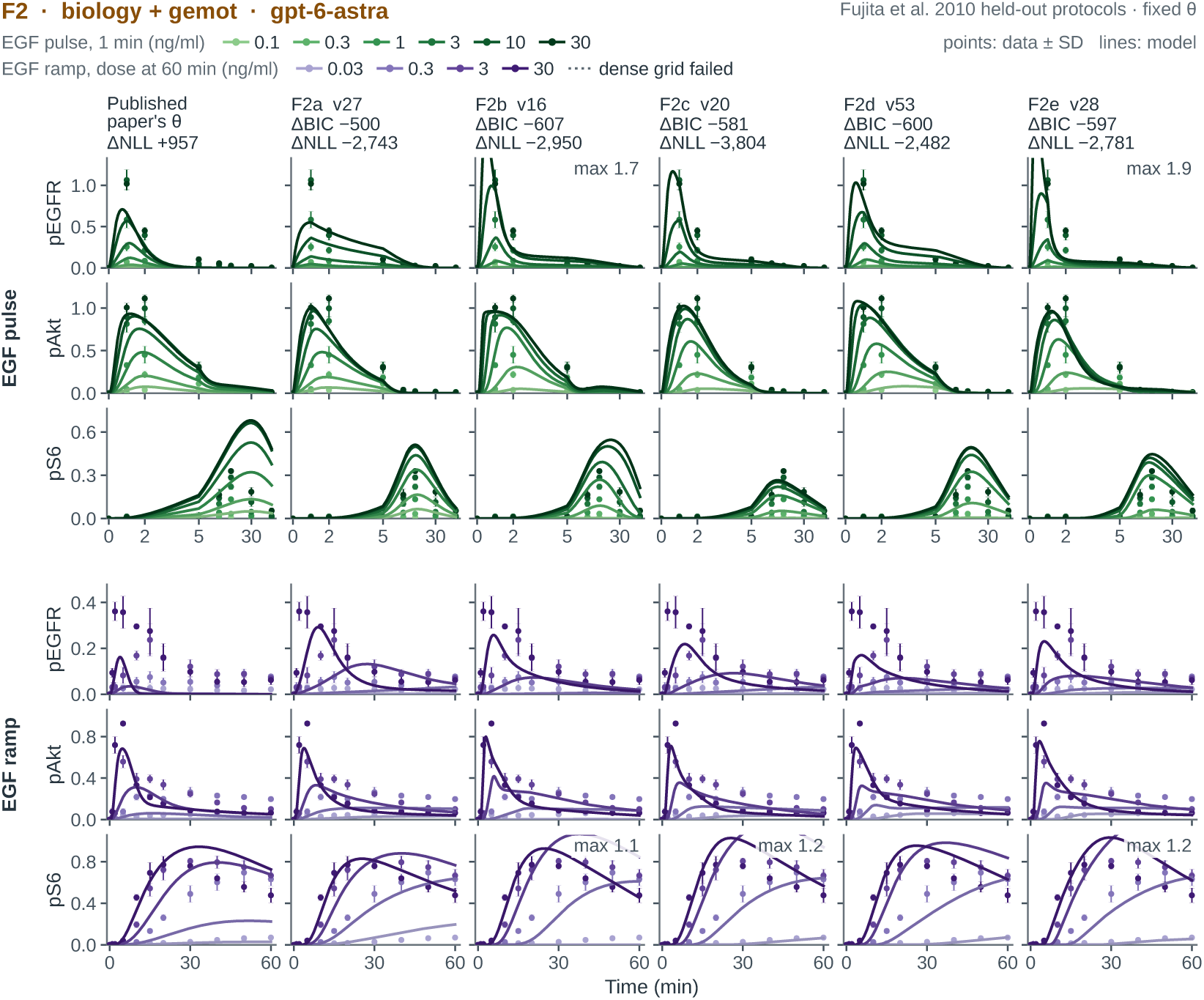
Held-out Fujita predictions, F2. (biology and gemot; GPT-6-Astra). As Figure 7; columns, the published model, then F2a–F2e.

**Figure 12:**
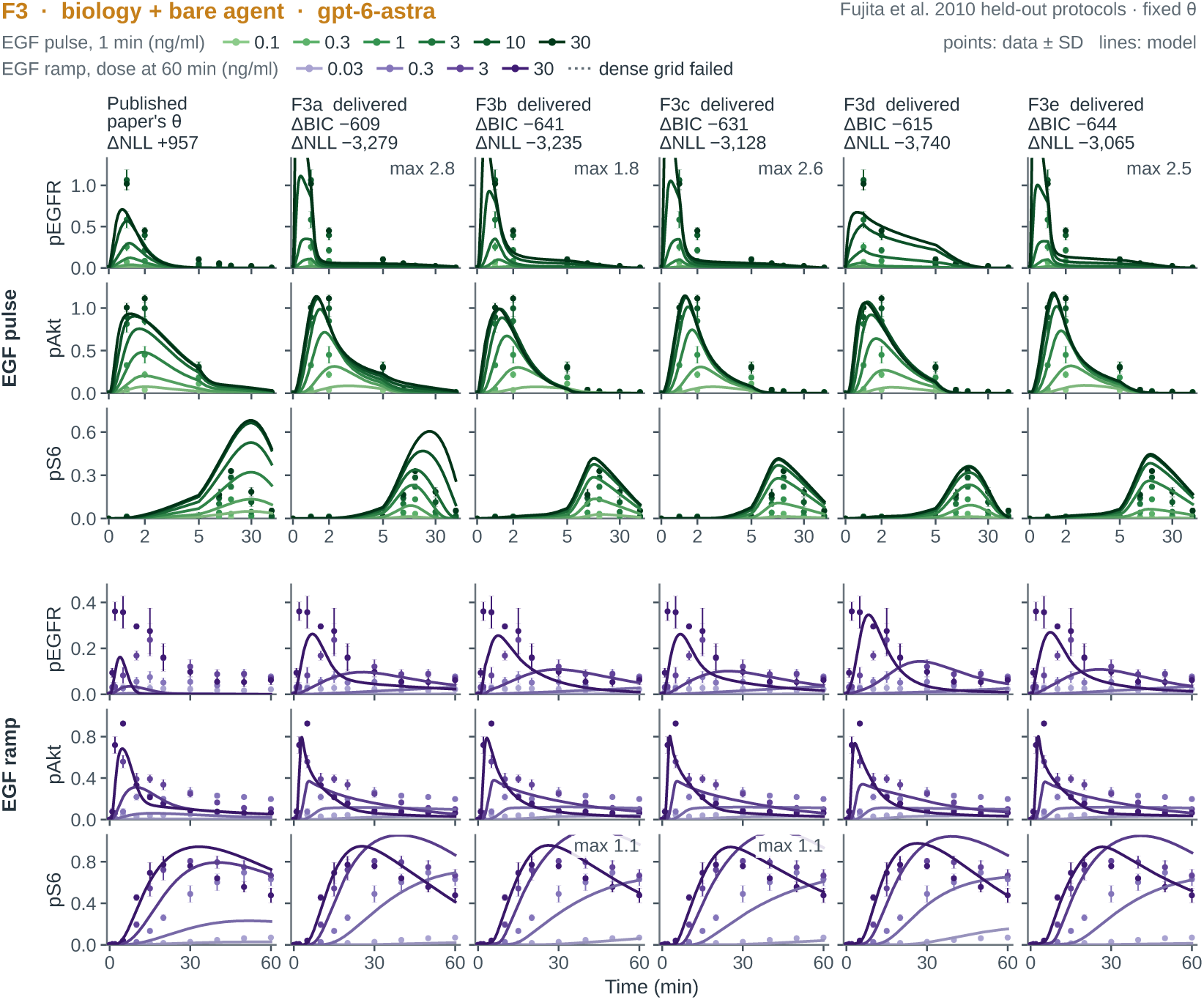
Held-out Fujita predictions, F3. (biology and bare agent; GPT-6-Astra). As Figure 7; columns, the published model, then F3a–F3e.

### D.6 Additional validation systems in Figure 4

The compact score export covers 20 biology-visible O1 GLM models in Figure 4: five each for Crauste, Lucarelli, Perelson and Raimundez. It does not cover the biology-blinded or neural-model comparisons in that figure.

The evaluation protocols only partly reproduce the original studies’ validation designs. Raimundez keeps dynamical parameters fixed, using published nominal values for the reference, and analytically fits new per-blot observation scales for the subset of validation experiments in the original Figures 5D and 6E (Raimúndez et al., 2020). Crauste refits each structure to the tumour and vaccinia responses in original Figures 2D and 2B; the former was a published transfer test, whereas the original early-to-late prediction tests in Figures 6 and S6 are not reproduced here (Crauste et al., 2017). Perelson uses four additional patients withheld from model construction, retaining patient-specific refitting as in the original study (Perelson et al., 1996). Lucarelli transfers to three further cell types, pooling genes from original Figure 6B (used for fitting) and Figure S6 (originally withheld), and refits all estimated parameters for candidate and reference rather than only signalling parameters as in the original study (Lucarelli et al., 2018). This broader refit treats candidates and reference symmetrically because the harness cannot reliably separate signalling from gene-regulatory parameters in arbitrary agent-built models. It tests whether model structures can be refitted across cell types, without testing whether the learned Smad-complex-to-gene-expression mapping transfers unchanged.

Median held-out ΔNLL values were +7.4 for Crauste, +29.8 for Lucarelli, −8.2 for Perelson, and +83.5 for Raimundez; all favour the references except Perelson, whose median favours the candidates. Four of five Perelson candidates improved on the reference; no Crauste, Lucarelli or Raimundez candidate did.

#### Held-out and transfer predictions

Figures 13–26 draw the runs of Figure 4, biology visible (O1) and biology blinded (O4), on their held-out data; Figures 27 and 28 draw its distilled neural-ODE models the same way. Raimundez is predicted at fixed dynamical parameters with each heldout blot’s observation scale solved at the fitted point; Crauste, Lucarelli, and Perelson are drawn at the optimum each model reached when refitted to each held-out unit from its training optimum. Column titles carry the training ΔBIC and held-out ΔNLL plotted in Figure 4, except that a distilled model’s title gives the held-out ΔNLL of its drawn refit: these are refitted with the neural judge’s 50 starts, as Figure 4 scored them, and need not settle in the optimum the judge’s refit found. The Raimundez figures and the biology-visible Perelson runs are drawn at a tighter tolerance than their scores were computed at (10*^−^*^16^ absolute, 10*^−^*^14^ relative); one blinded Raimundez run (7fd274ec), whose held-out objective is not finite there, is drawn instead at the fit tolerance its score used, where its drawn likelihood reproduces the score. The other transfer figures are drawn at the fit tolerance their scores used (10*^−^*^10^ absolute, AMICI’s default relative tolerance). Four of the five blinded Raimundez models do not respond to the EGFR knockdown, where every biology-visible model does: the knockdown input is constant in the training data and, under blinding, was not wired into EGFR synthesis.

**Figure 13:**
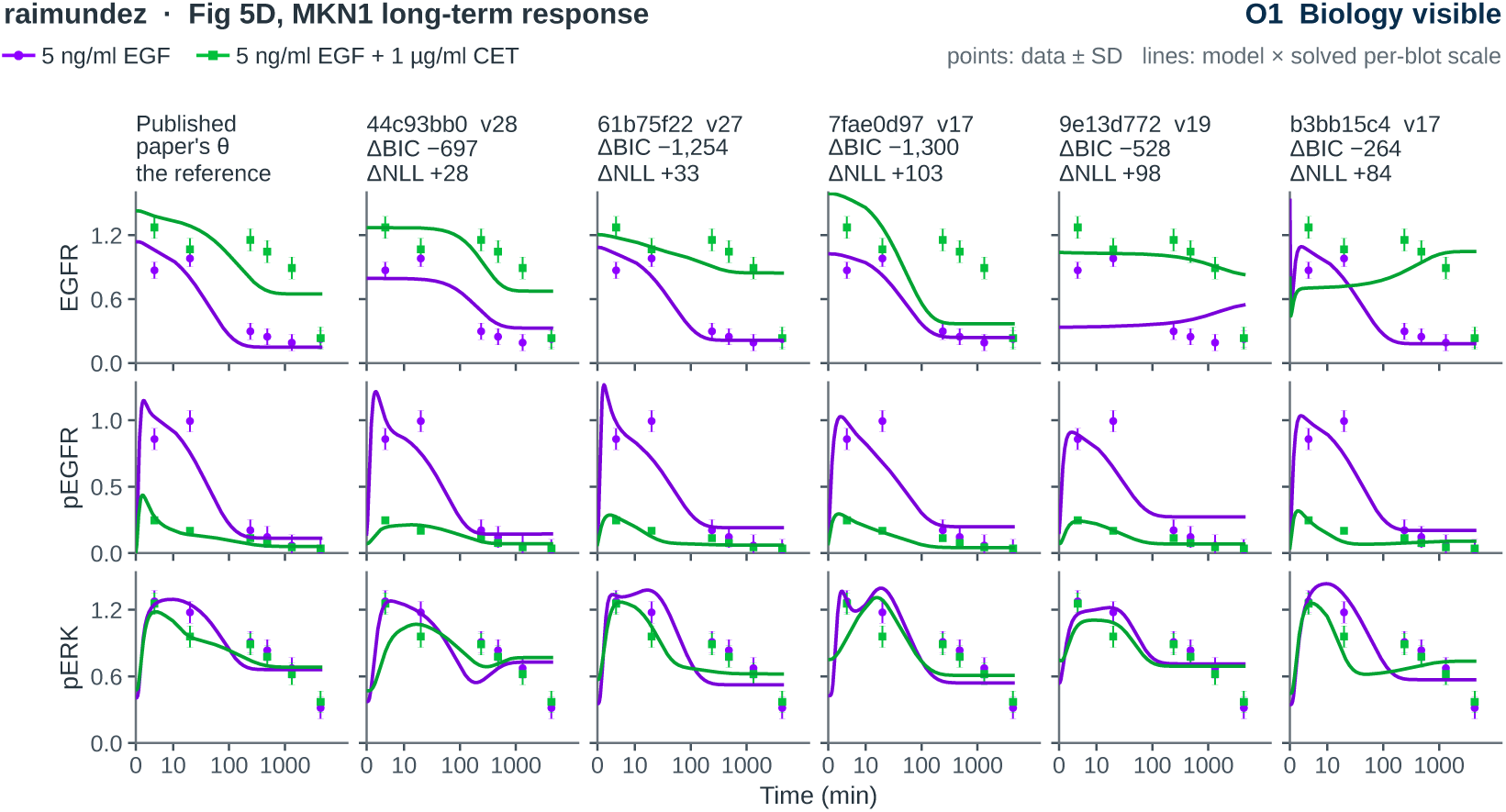
Held-out Raimundez predictions, Fig. 5D, biology visible (O1). MKN1 long-term response to 5 ng/ml EGF (purple) and EGF with 1 *µ*g/ml cetuximab (green). Columns, the published model at its nominal parameters (the reference), then the five Figure 2 runs. Points, data ± SD; lines, each model’s readout times the per-blot observation scale solved at the fitted point, the quantities the held-out likelihood scores.

**Figure 14:**
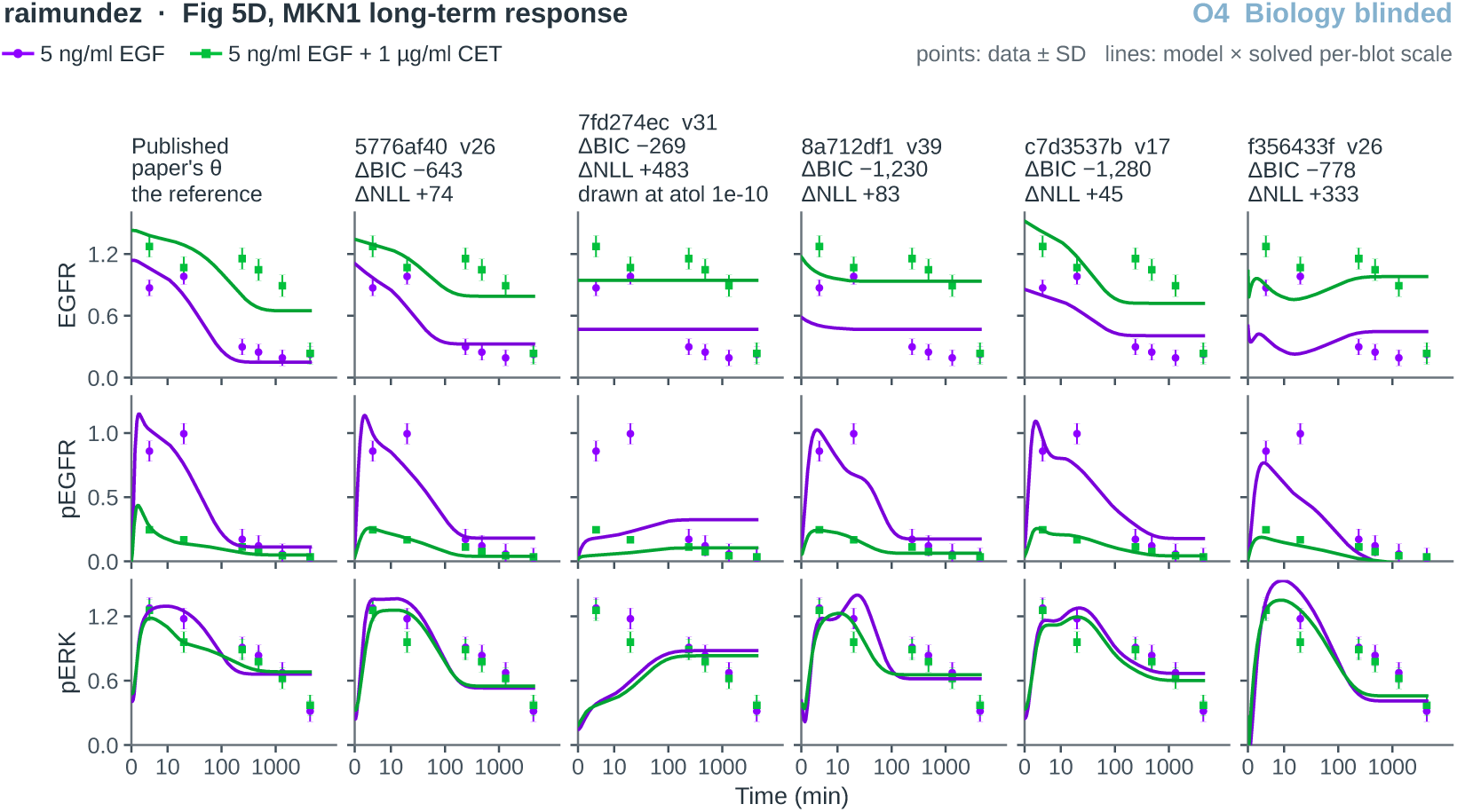
Held-out Raimundez predictions, Fig. 5D, biology blinded (O4). As Figure 13, for the five biology-blinded runs.

**Figure 15:**
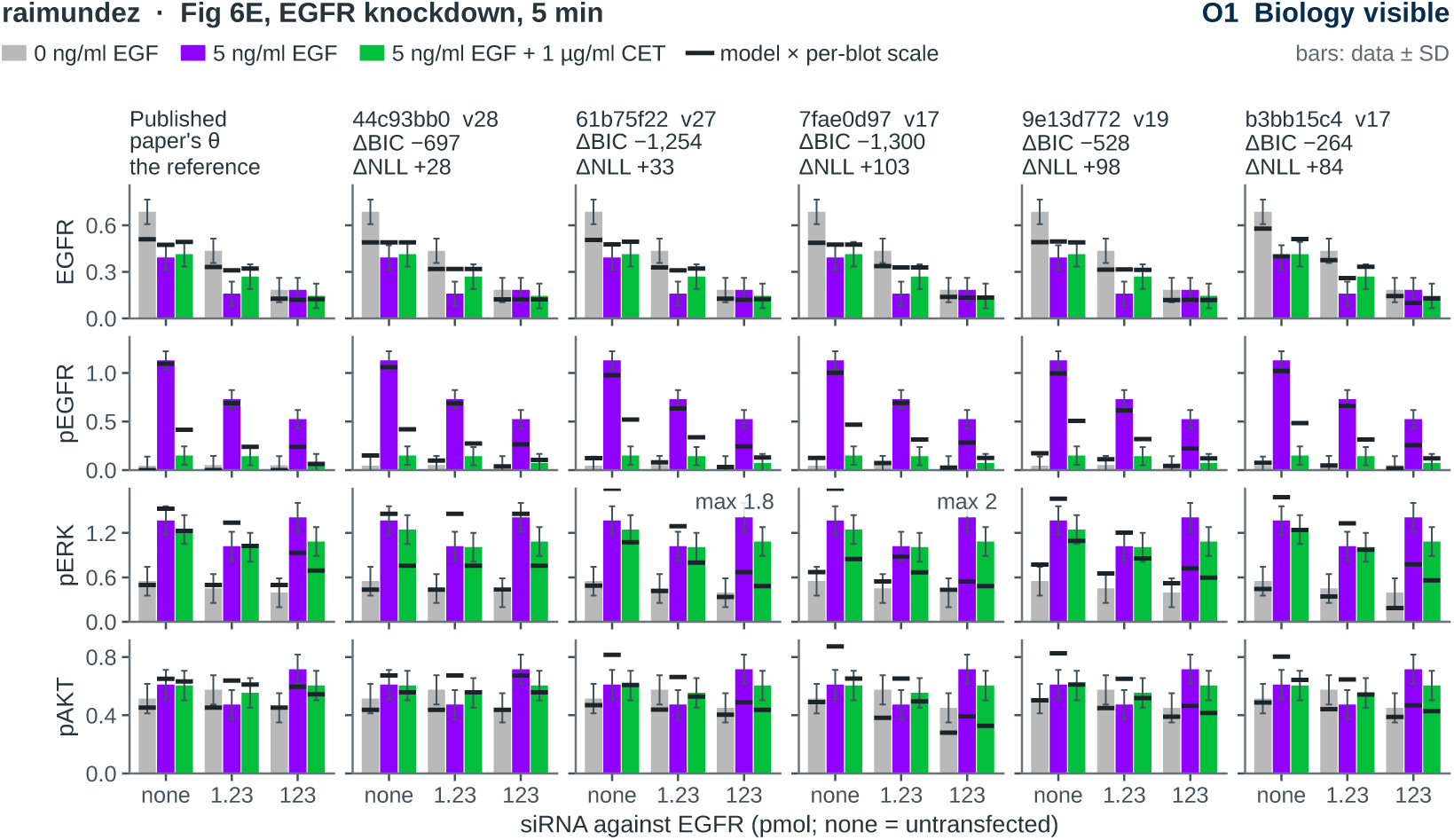
Held-out Raimundez predictions, Fig. 6E, biology visible (O1). EGFR knockdown, read 5 min after no stimulus (grey), EGF (purple), or EGF with cetuximab (green). Bars, data ± SD; dark levels, each model’s scaled prediction.

**Figure 16:**
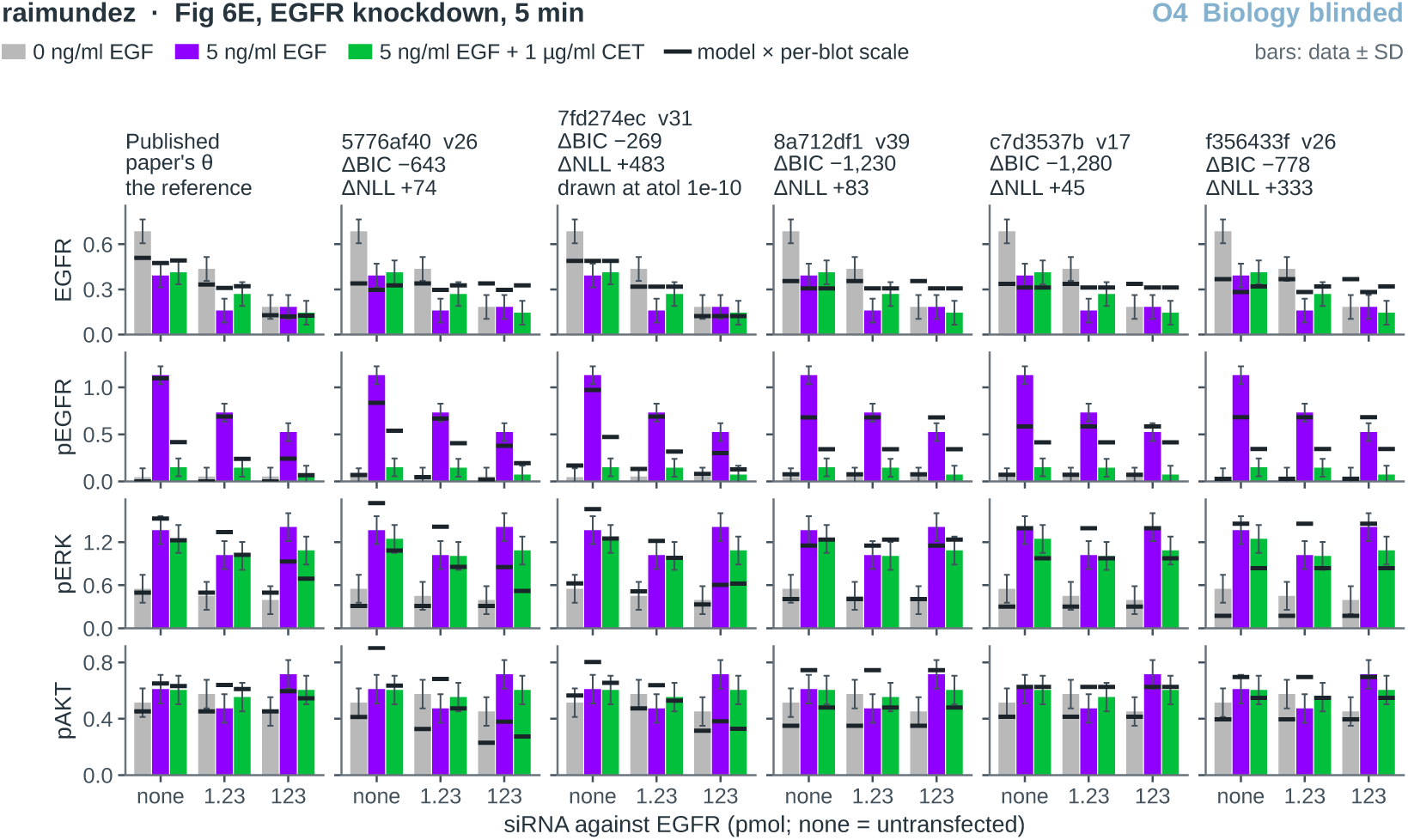
Held-out Raimundez predictions, Fig. 6E, biology blinded (O4). As Figure 15, for the five biology-blinded runs.

**Figure 17:**
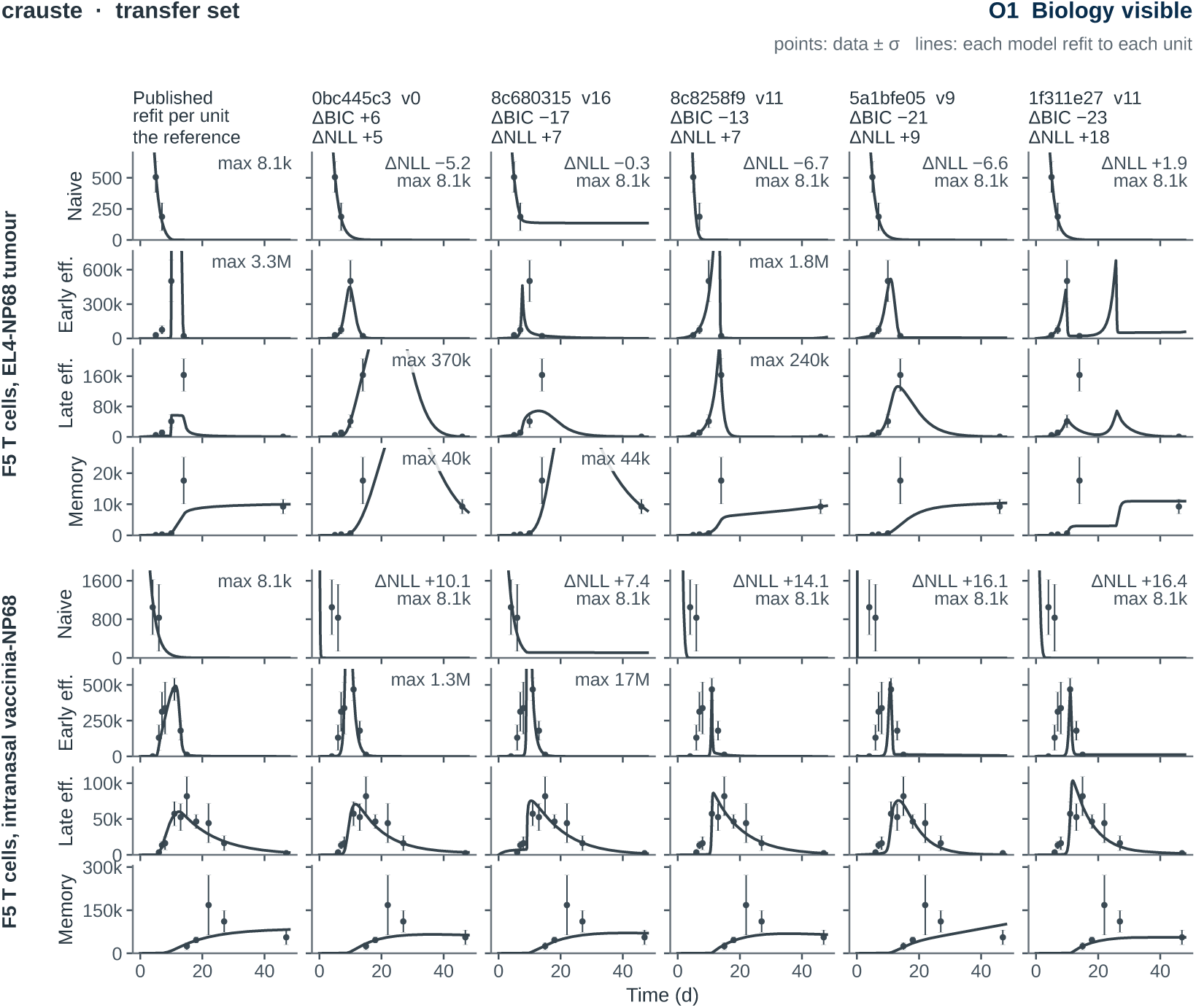
Crauste transfer refits, biology visible (O1). Two F5 T-cell experiments as units, four populations each. Columns, the published structure refitted per unit (the reference), then the five Figure 2 runs, each refitted per unit. Points, data ± *σ* (the refit’s own where estimated); lines, the refitted model; the first panel of each unit gives that unit’s ΔNLL.

**Figure 18:**
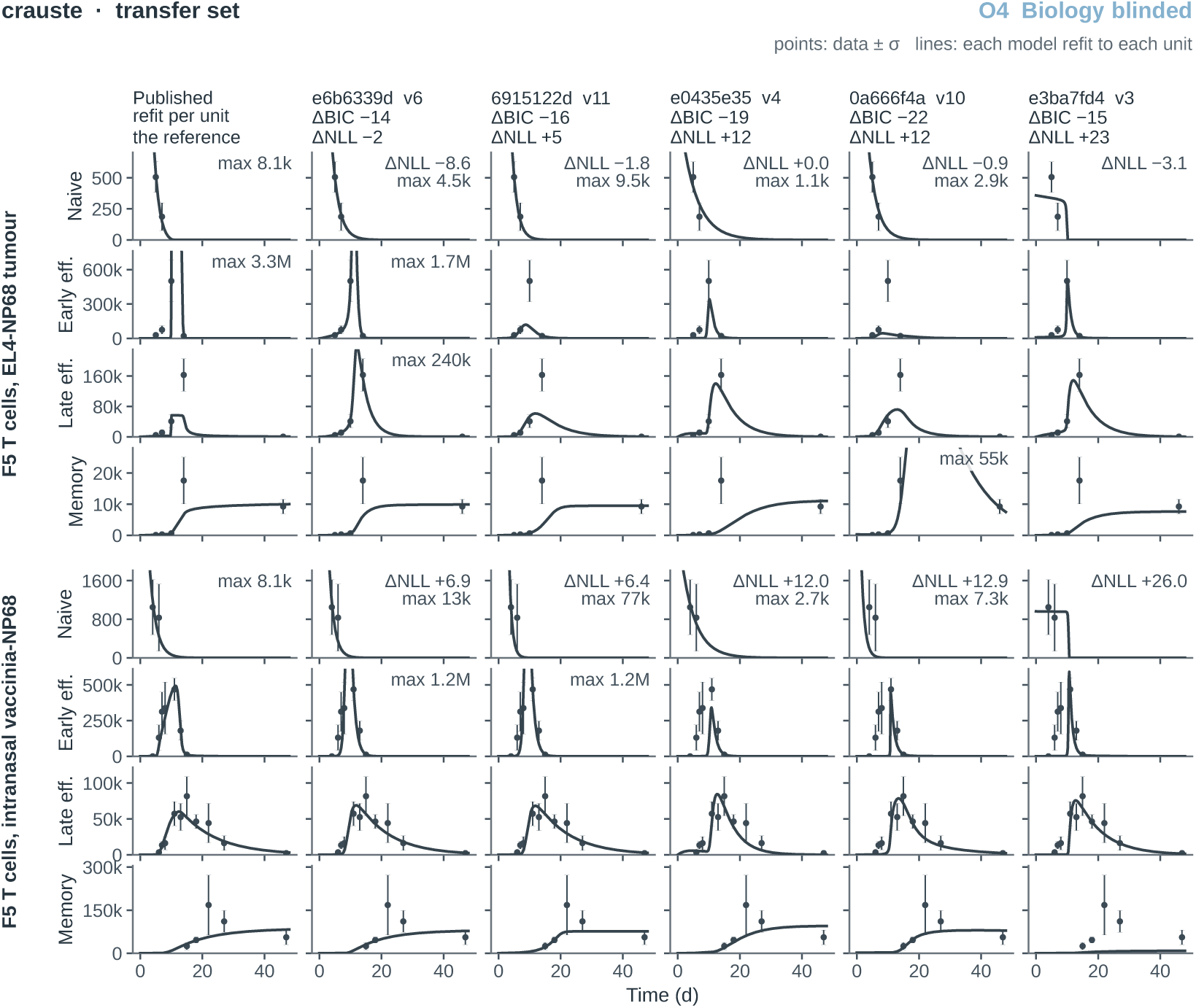
Crauste transfer refits, biology blinded (O4). As Figure 17, for the five biologyblinded runs.

**Figure 19:**
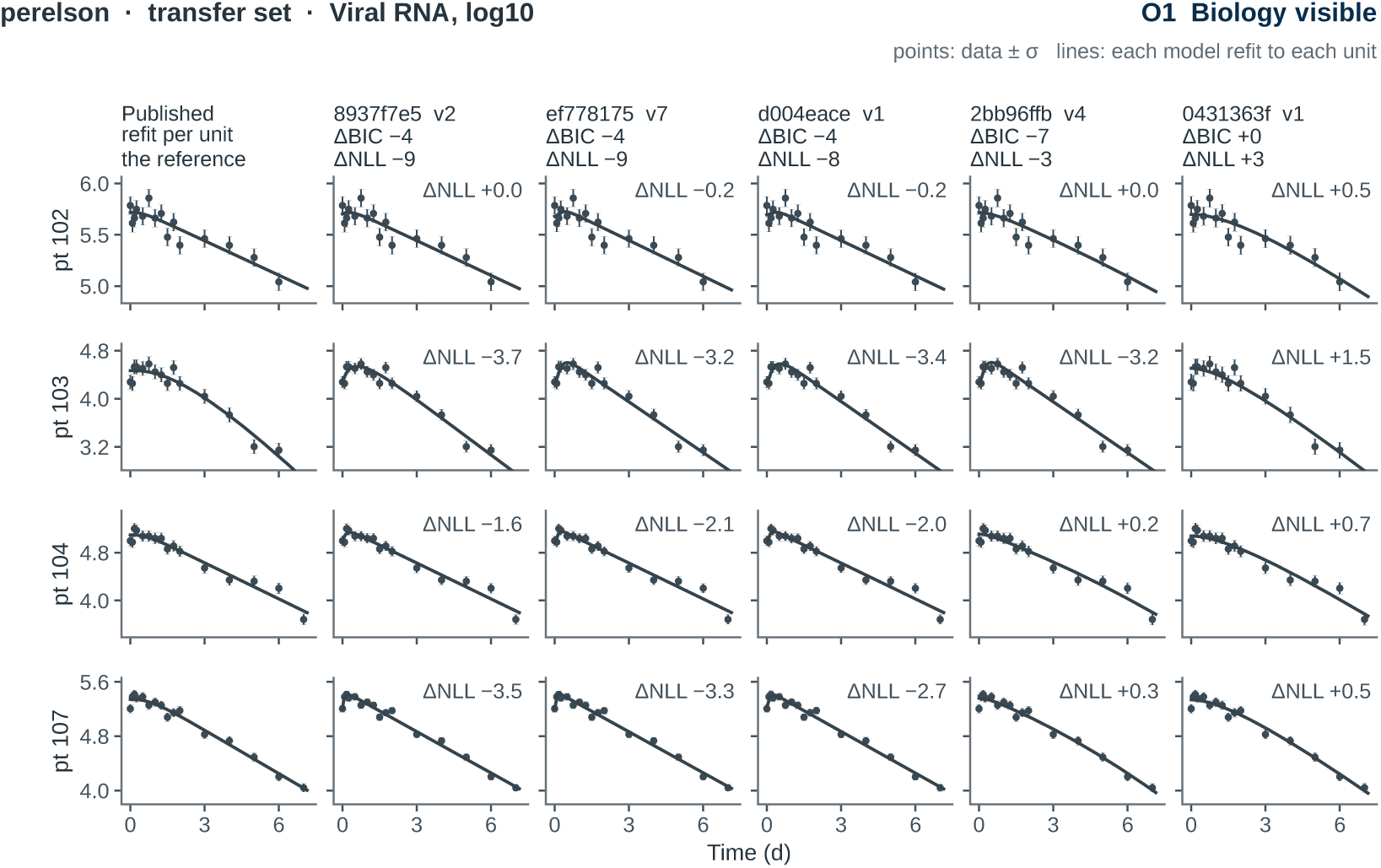
Perelson transfer refits, biology visible (O1). Viral RNA (log_10_) for four patients as units. Encoding as in Figure 17.

**Figure 20:**
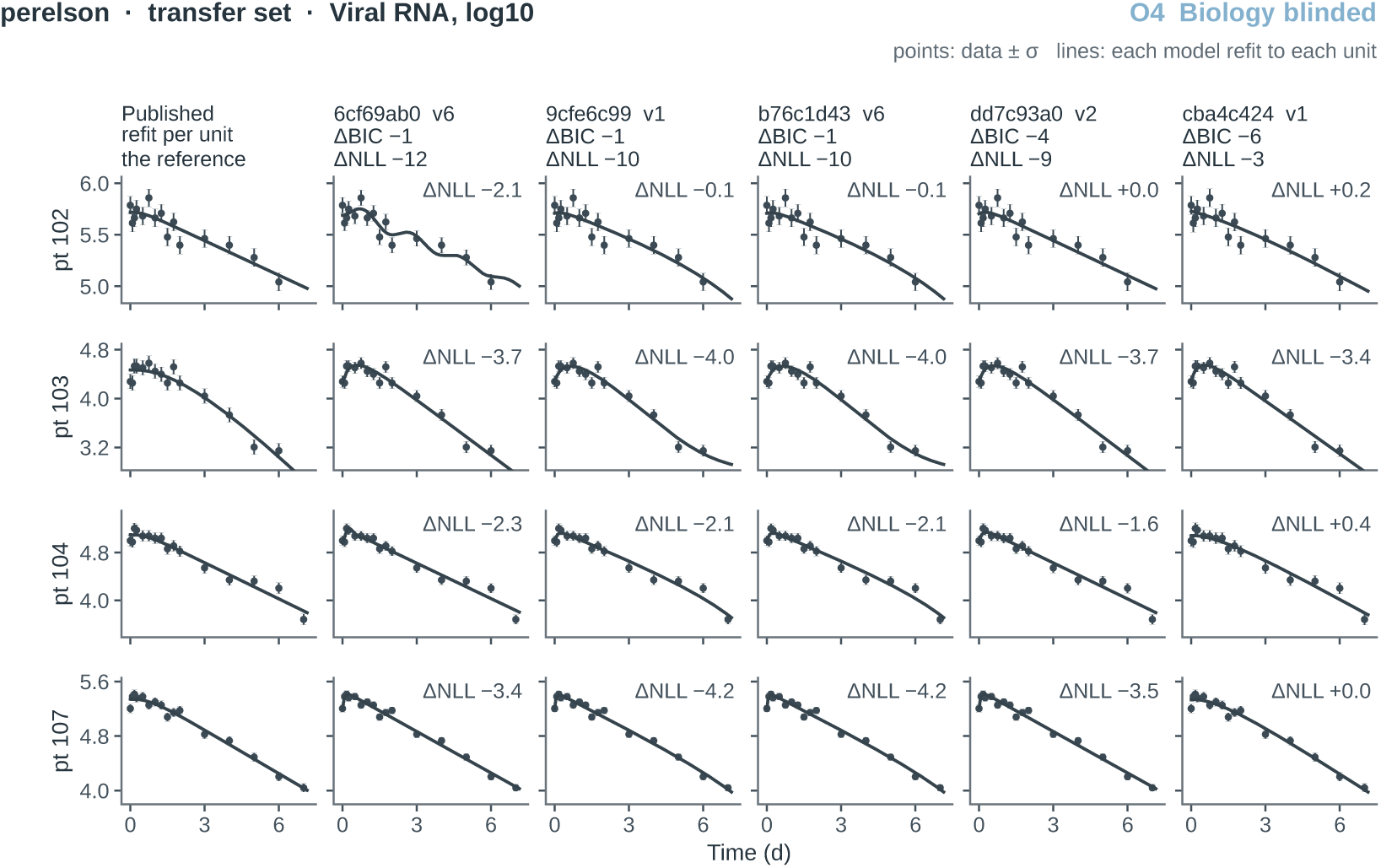
Perelson transfer refits, biology blinded (O4). As Figure 19, for the five biology-blinded runs.

**Figure 21:**
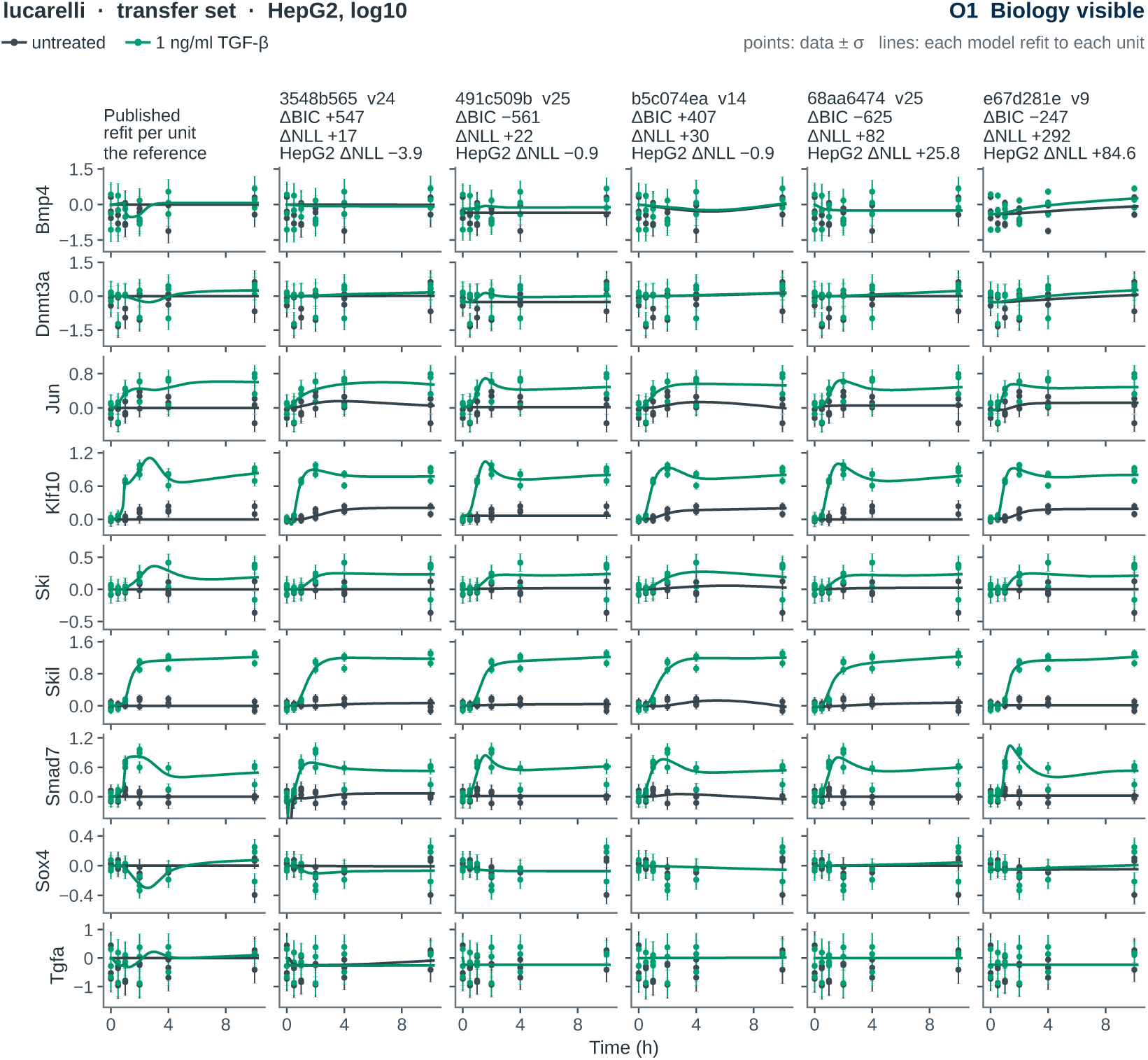
Lucarelli transfer refits, HepG2, biology visible (O1). Gene expression (log_10_) in HepG2 cells, untreated (slate) and with 1 ng/ml TGF-*β* (green). Encoding as in Figure 17; the fourth title line gives the cell type’s share of each run’s ΔNLL.

**Figure 22:**
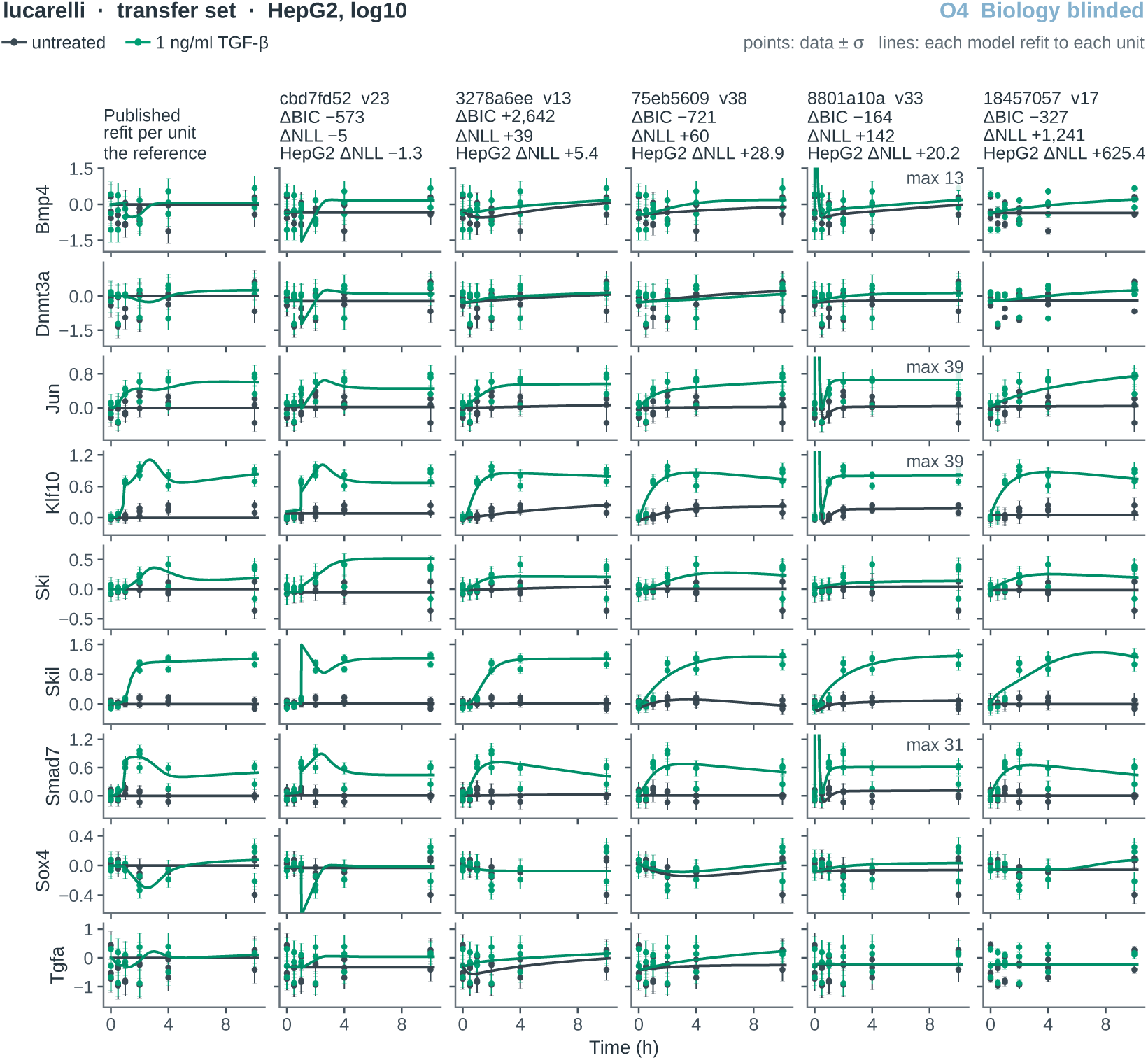
Lucarelli transfer refits, HepG2, biology blinded (O4). As Figure 21, for the five biology-blinded runs.

**Figure 23:**
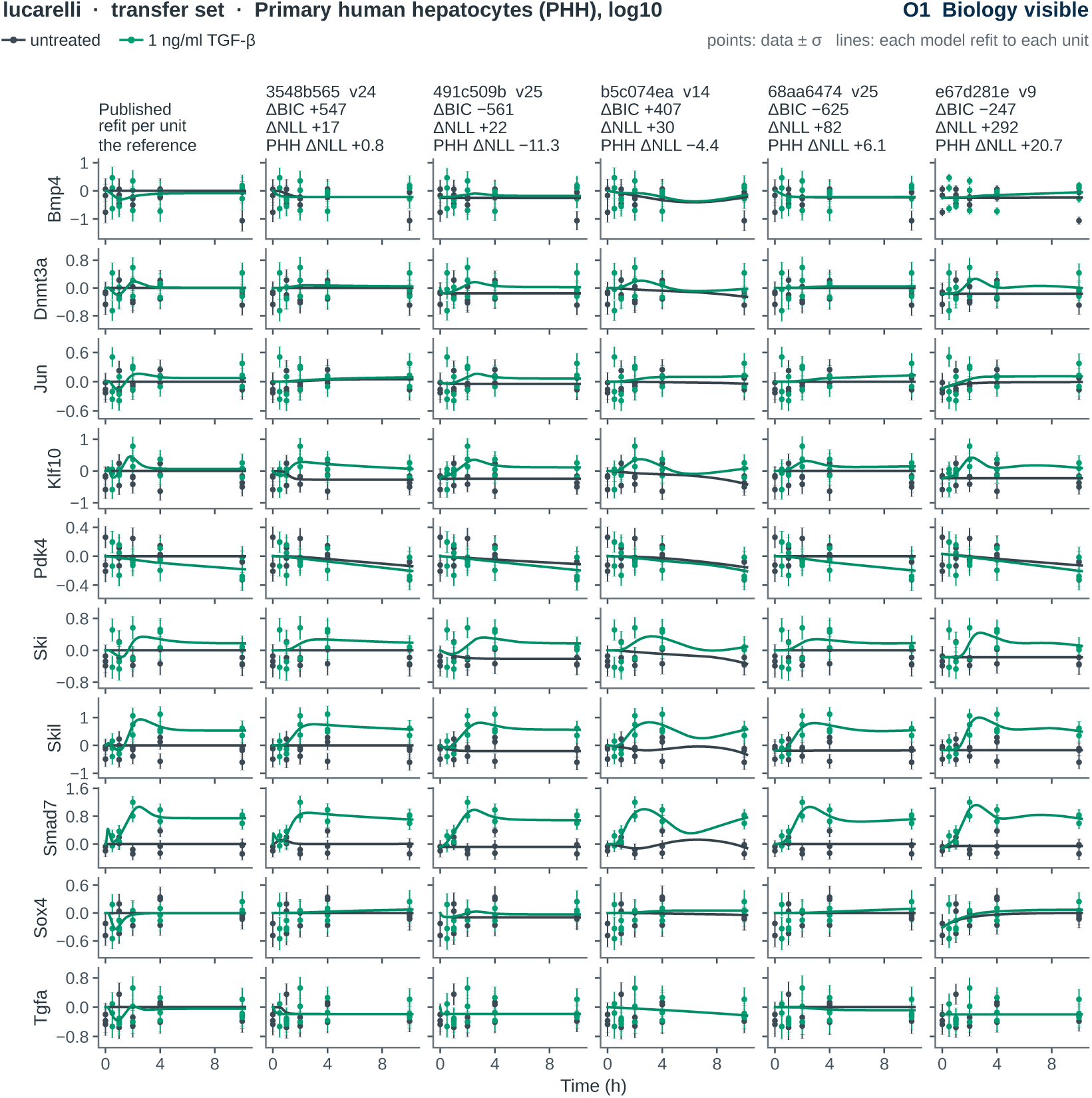
Lucarelli transfer refits, PHH, biology visible (O1). Gene expression (log_10_) in primary human hepatocytes, untreated (slate) and with 1 ng/ml TGF-*β* (green). Encoding as in Figure 17; the fourth title line gives the cell type’s share of each run’s ΔNLL.

**Figure 24:**
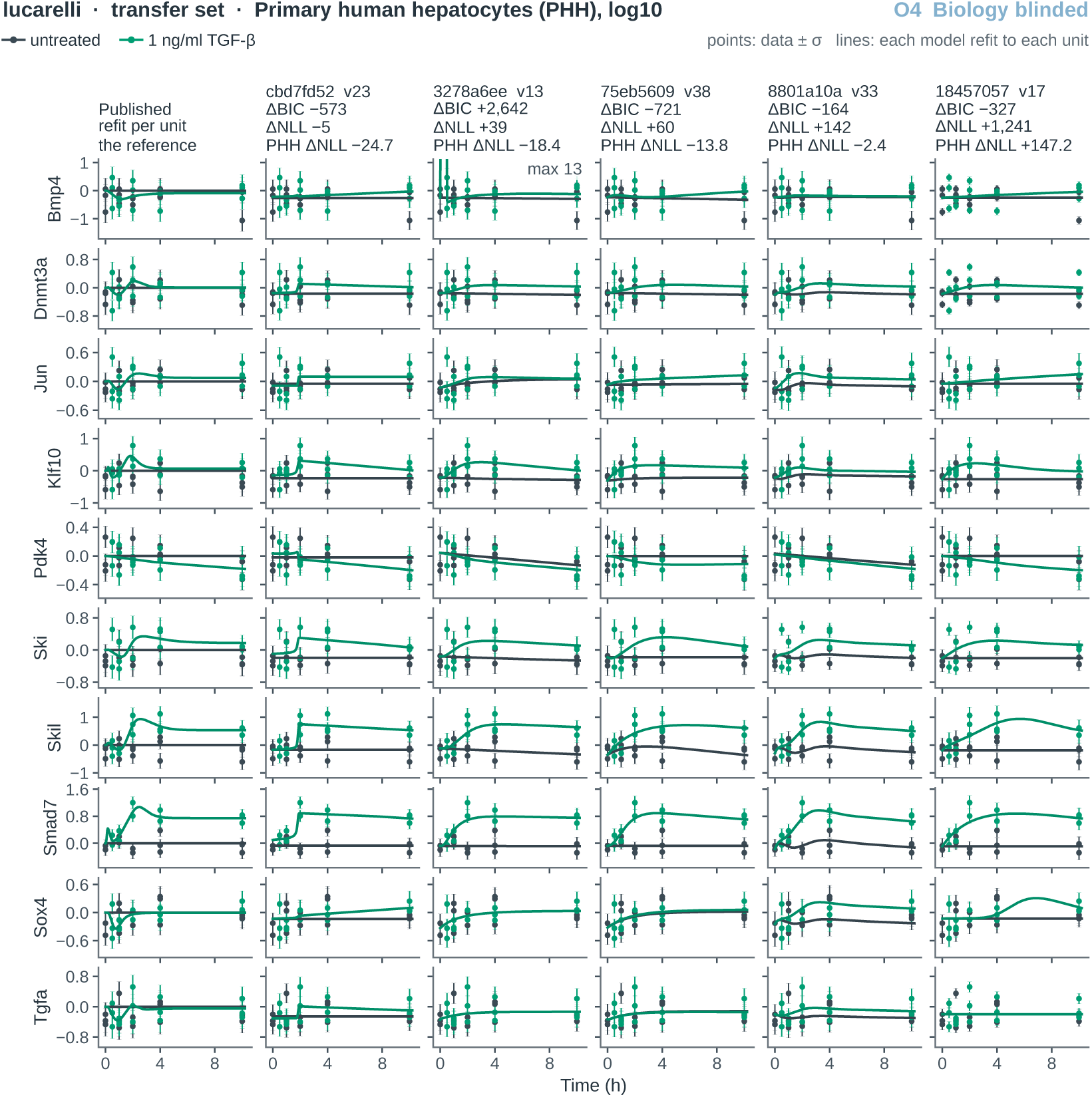
Lucarelli transfer refits, PHH, biology blinded (O4). As Figure 23, for the five biology-blinded runs.

**Figure 25:**
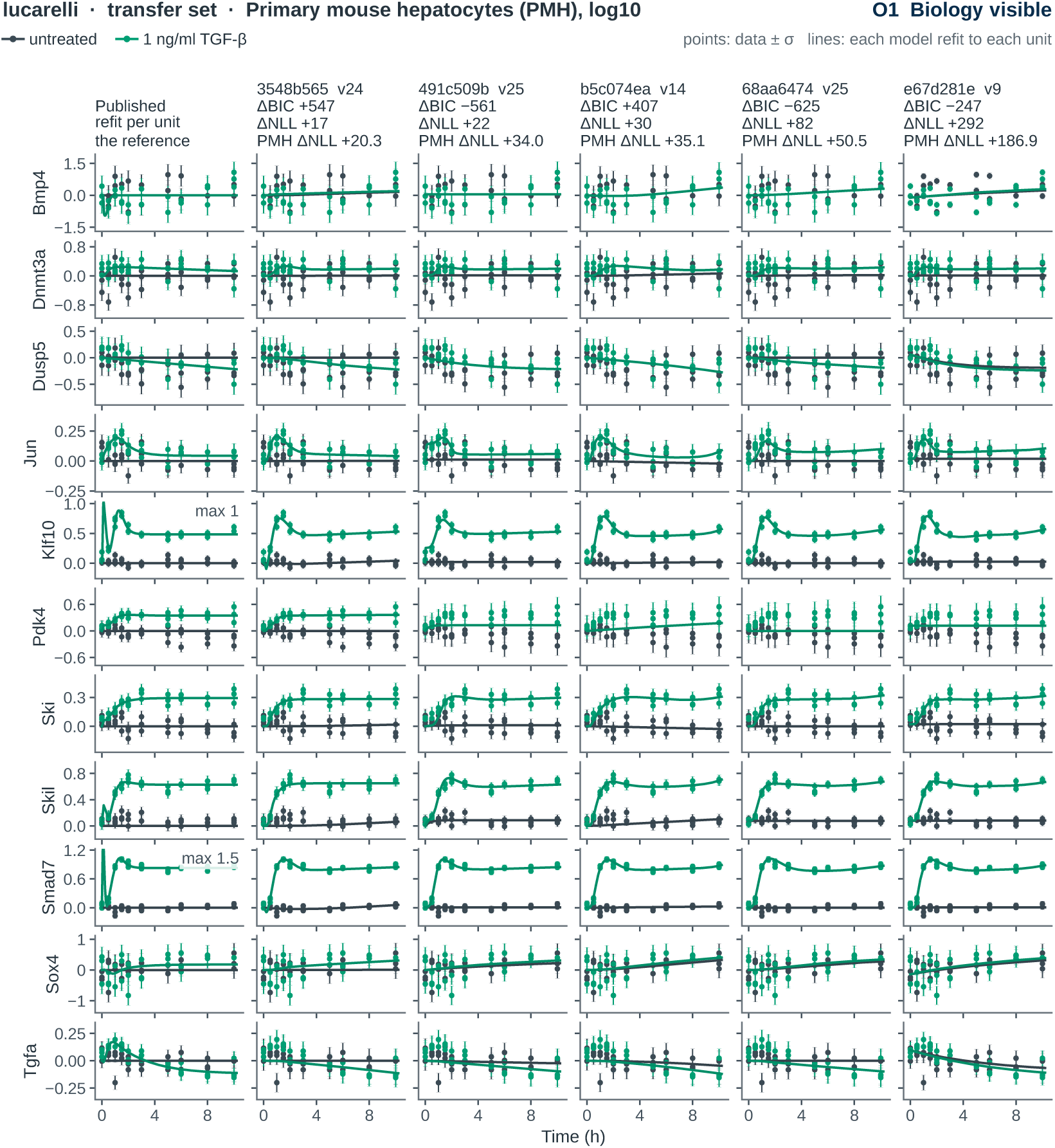
Lucarelli transfer refits, PMH, biology visible (O1). Gene expression (log_10_) in primary mouse hepatocytes, untreated (slate) and with 1 ng/ml TGF-*β* (green). Encoding as in Figure 17; the fourth title line gives the cell type’s share of each run’s ΔNLL.

**Figure 26:**
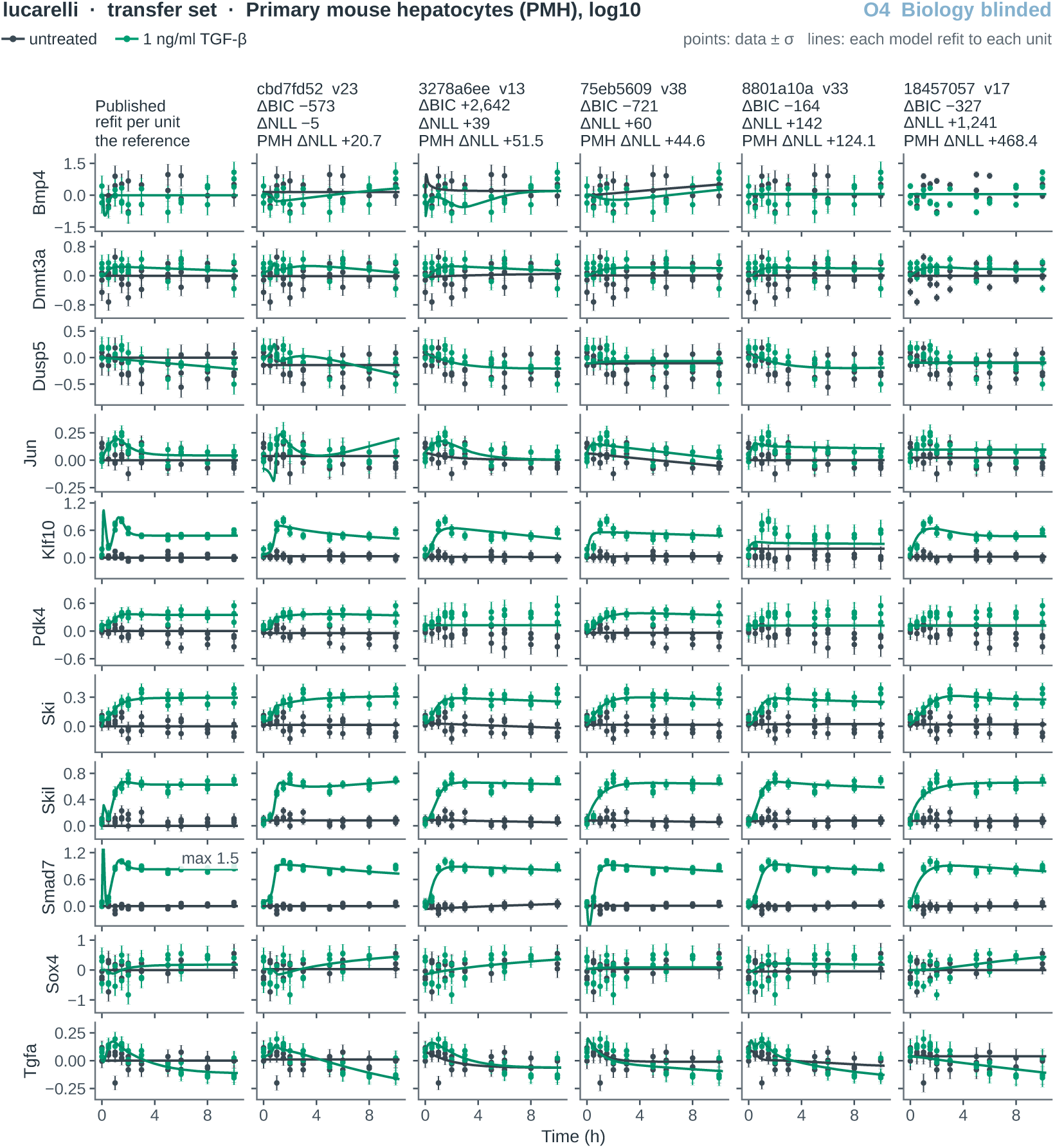
Lucarelli transfer refits, PMH, biology blinded (O4). As Figure 25, for the five biology-blinded runs.

**Figure 27:**
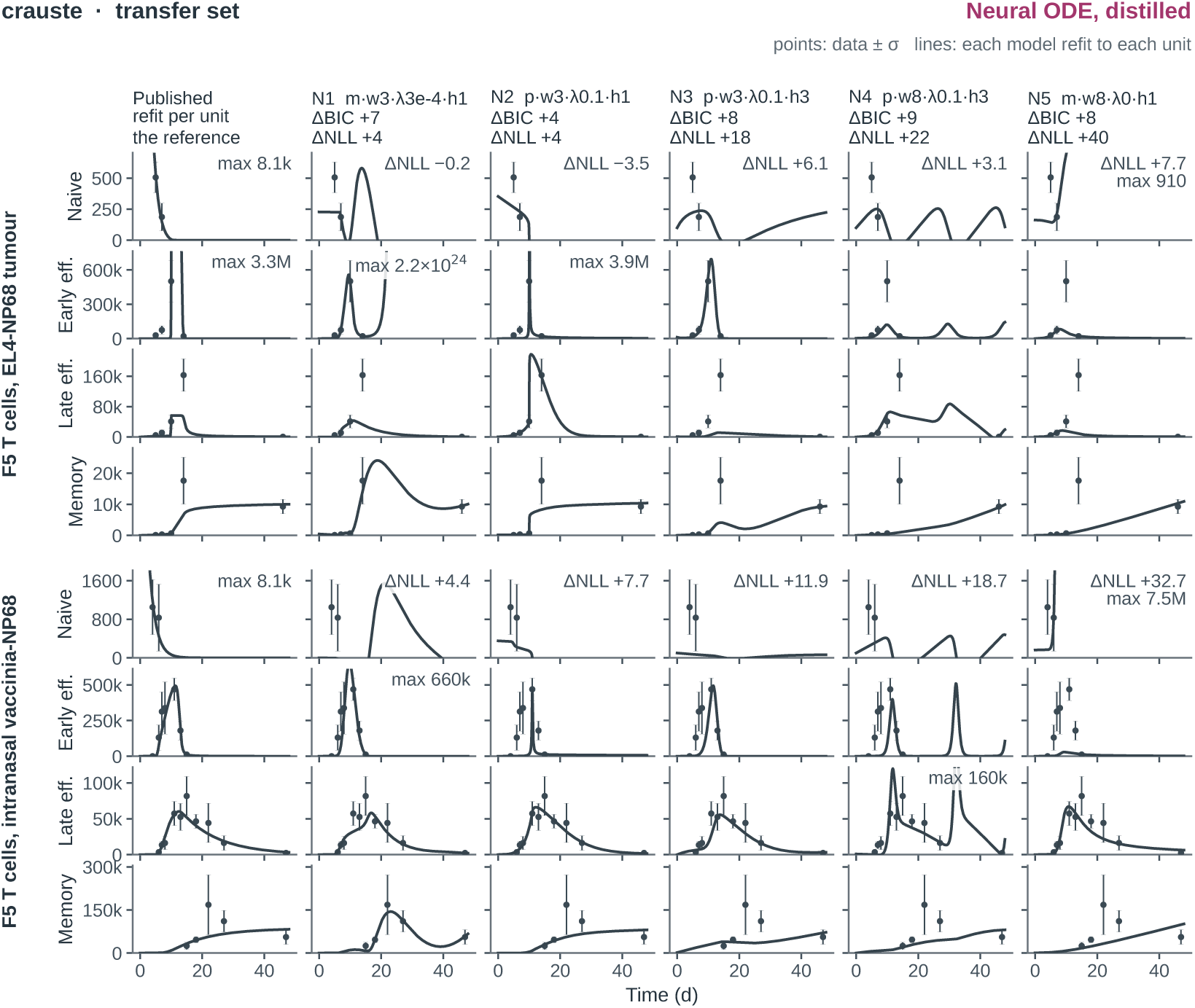
Crauste transfer refits, distilled neural ODEs. As Figure 17, for the five distilled neural-ODE models of Figure 4 (Appendix D.7), each refitted per unit with the neural judge’s 50 starts. Columns N1–N5 are titled by grid cell: plain (p) or multiplicative (m) field, hidden width w, *ℓ*_2,1_ weight *λ*, and horizon stages h. The held-out ΔNLL in each title is that of the drawn refit.

**Figure 28:**
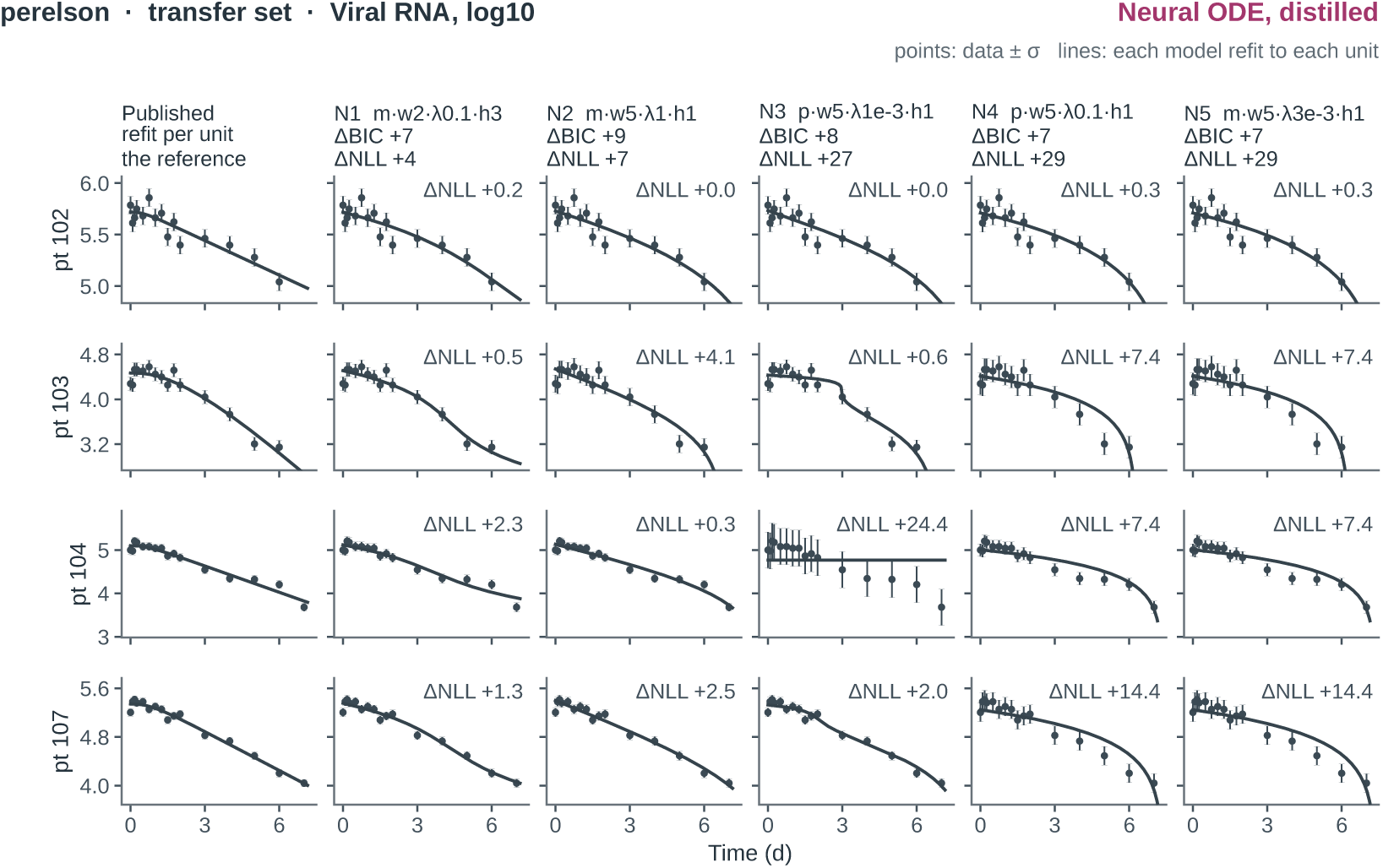
Perelson transfer refits, distilled neural ODEs. As Figure 19, for the five distilled neural-ODE models of Figure 4, each refitted per unit with the neural judge’s 50 starts; columns as in Figure 27. N4 and N5 come from different grid cells but distil to the same equation, viral RNA falling at a constant rate under therapy, and differ only in fitted constants.

### D.7 Neural ODE distillation protocol

The neural baseline constructs models without an LLM: a neural ODE (Chen et al., 2018) is fitted to training measurements, then its vector field is distilled into symbolic rate laws using PySR (Cranmer, 2023). The symbolic model is subsequently refitted through `gemot`’s numerical evaluation layer (Appendix B.7). Figure 4 reports these distilled models; their parameter penalties count the fitted symbolic model’s parameters, including estimated observation parameters, after the neural network has been removed.

#### Data interface and neural training

The implementation reads the supplied measurement, observable, condition and parameter tables, together with time-dependent input specifications where present. It does not read the reference SBML model. The `readout_*` symbols of the observable formulas and explicitly dosed species define the initial state representation, which can be augmented with latent states. The supplied observable and noise formulas, experimental inputs and preequilibration conditions are retained. Known initial amounts remain fixed; unspecified initial amounts are estimated. During neural training, observation scales are fixed at their supplied nominal values, allowing neural states to carry the corresponding amplitudes.

A fully connected network with tanh hidden activations predicts the state derivatives from the states and experimental inputs. States and inputs are normalised using data-derived scales, and time by the last finite training measurement time. Neural training problems are represented using PEtab SciML with PEtab version 2 (Pathirana et al., 2026; Persson et al., 2026) and fitted with AMICI’s JAX backend in double precision, with adaptive Kvaerno5 integration and gradient-based optimisation. Positive scalar parameters, including estimated initial amounts and noise scales, are optimised in log space. The reported grid used multistart Adam, with one or three horizon stages and either no sparsity penalty or an *ℓ*_2,1_ penalty (Equation 13; Table 10). Failed integrations are recorded rather than treated as fitted models.

**Table 10:** The neural-ODE search grid. The axes above the rule define 216 configurations per benchmark; shared recorded settings appear below it. Each cell was allocated two CPUs, 16 GB of memory and a 12 h scheduler limit. These are search configurations, not counts of successful fits.

| Setting | Values |
| --- | --- |
| Hidden width | crauste 3, 5, 8; fujita 4, 7, 12; perelson 2, 3, 5 |
| Augmented states | 0, 1 |
| Multiplicative field | no, yes |
| Horizon stages | 1, 3 |
| $L_{2,1}$ weight (0 = none) | 0, $3 \times 10^{-4}$, $1 \times 10^{-3}$, $3 \times 10^{-3}$, 0.01, 0.03, 0.1, 0.3, 1 |
| Network depth | 2 |
| Optimiser | Adam |
| Learning rate | $3 \times 10^{-3}$ |
| Steps | 5000 |
| Restarts | 4 |
| Patience (steps) | 300 |
| Solver tolerances (rel., abs.) | $1 \times 10^{-6}$, $1 \times 10^{-8}$ |
| Solver step limit | 8192 |
| Shooting windows | 1 |
| Seed | 0 |
| Wall time per cell (s) | 36000 |
| Symbolic regression iterations | 40 |
| Expression size limit | 20 |
| Population size | 33 |
| Parsimony | $3.2 \times 10^{-3}$ |
| Operators | +, −, ×, ÷ |
| Samples of the field | 2000 |
| Proposals per cell | 11 (complexity budgets 1–19, and the distiller’s own selection) |
| Judge fit starts | 50 |

#### Multiplicative field

Let *ξ* ∈ R*^d^* collect the normalised states, including any latent states, and experimental inputs. The plain field feeds *ξ* directly to the network. The multiplicative field augments it with all unordered pairwise products, including squares:

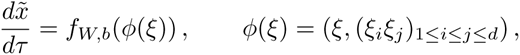

where *x̃* and *τ* denote normalised states and time, and *f_W,b_* is the neural network. This expands the first-layer input dimension from *d* to *d* + *d*(*d* + 1)*/*2, making bilinear interactions, such as massaction terms, and quadratic dependence directly available to the network. The products are algebraic features; the network outputs the state derivatives. Symbolic distillation uses the original variables as regressors, so PySR must recover any products in the symbolic rate laws.

#### Sparsity penalties

Let *W* ^(*ℓ*)^, *ℓ* = 1*, . . ., L*, denote the neural field’s weight matrices, *b* its biases, and *ϑ* the fitted scalar parameters. For training data *D*, the regularised objective is

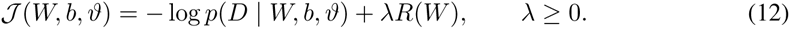

The grid uses *λ* = 0 for unpenalised training and otherwise applies

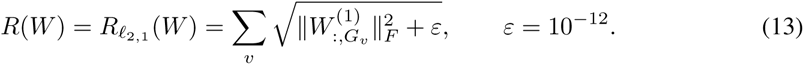

Here *G_v_* indexes first-layer columns carrying input variable *v*. For the plain field, each group is one column; in the multiplicative field, each product column belongs to the groups of its constituent variables, so groups can overlap. Biases and fitted scalar parameters are not penalised. The *ℓ*_2,1_ penalty targets input-variable dependence. Penalised training additionally applies group-wise soft thresholding after each step with strength *ηλ*, where *η* is the learning rate. After penalised training it attempts an unpenalised refit, initialised at the selected solution and holding zero first-layer weights fixed. This penalty guides neural-field construction; the reported BIC is computed from the subsequently fitted symbolic model.

#### Symbolic distillation and refitting

The trained field is sampled along simulated training trajectories and around the settled states reached during preequilibration, using the corresponding experimental or basal inputs and local state perturbations. Each preequilibrated experiment contributes 25 basal-state rows to the sampling pool. PySR regresses each state derivative against the sampled states and inputs. These regression targets are neural-field evaluations, rather than finite differences of the experimental measurements. Candidate expressions are retained along an accuracy–complexity frontier and combined into complete ODE systems. The equations are converted back to data units and exported as Antimony rate rules; reaction stoichiometry is not inferred by this step. Non-integer coefficients become named parameters initialised at their regressed values. Each proposed system is then reconciled, compiled and fitted to the experimental training data through the same MCP evaluation path used for agent-built models. Agreement with the neural field is therefore a proposal criterion, while the reported fit and BIC come from the refitted symbolic model.

#### Selection and transfer evaluation

The neural points in Figure 4 were selected from the `grid3-20260922` campaign. The export collects judged candidates with recorded training BIC and held-out scores, sorts them by training BIC, removes duplicate BIC values after rounding to six decimal places, and retains the first five per problem. Thus, these points summarise selected candidates from a hyperparameter search; their spread does not represent five independent repetitions of the full procedure. Deduplication by BIC is a numerical heuristic, not a test of symbolic model identity. For Crauste and Perelson, the symbolic structures undergo the transfer refitting described in Appendix D.6. Their held-out ΔNLL values use the same refitted references as the O1 and O4 comparisons.

#### Search grid and implementation

The search comprised 648 configurations, with 216 each for Crauste, Fujita and Perelson: three hidden widths, two augmented-state counts, two multiplicative-field settings, two horizon curricula and nine sparsity weights (Table 10). The archived specification, `paper/neural_distillation/neural_ode_grid.json`, records these axes, shared training, solver and distillation settings, evaluation budgets and execution revision `4d7da49`. It was extracted from per-cell configurations, recorded trainer and distiller settings, and launch metadata; `paper/generated/neural_ode_grid.csv` enumerates every configuration. All ten plotted neural candidates map to cells in this grid; their selection procedure and scores are archived alongside the specification. The construction workflow above was documented from `src/petab_ harness/neural_ode/` at revision `249d7c5`.

## E Fujita model diagrams and parameter uncertainty

The expanded Figure 3 compares 28 delivered models from 30 audited runs. Its display names identify the language model, configuration, and run; Appendix D.5 maps them to the original archive labels. The source annex (Appendix G) contains Antimony listings and parameter tables for all 28 delivered models across the six configurations. Twenty listings preserve the archived Antimony; eight bare-agent listings were converted from archived SBML. The manifest `paper/generate d/fujita_models.csv` connects each exported source to its task, version, and hashes. Source initializations need not equal the separately saved fitted parameter values.

### E.1 Parameter uncertainty around the plotted optima

Figure 3A plots one parameter vector per model. To test whether that choice drives the held-out ranking, the vectors from a 1000-start multistart that the training data cannot distinguish from the best optimum found were re-scored on the held-out protocols in Appendix D.5 (Figure 30), every band member of every model. 342 of them, 192 in O1e, 95 in O2c, 52 in O4b, and 3 in O4e, have no held-out value because their held-out simulation did not integrate. Band membership was decided at the tolerance the held-out score is computed with (atol 10*^−^*^16^, rtol 10*^−^*^14^): at the fit tolerance, apparent spreads of optima were integration error rather than non-identifiability. Tightening the tolerance moves an archived optimum’s training NLL by at most 8.5 × 10*^−^*^4^ (median 2.3 × 10*^−^*^6^; more than 10*^−^*^5^ in 4 of 27 models) and its held-out NLL by at most 3.7 × 10*^−^*^8^. All 2704 band members whose simulated values exceed ten times the largest held-out measurement fall outside the band at the tight tolerance.

**Figure 29:**
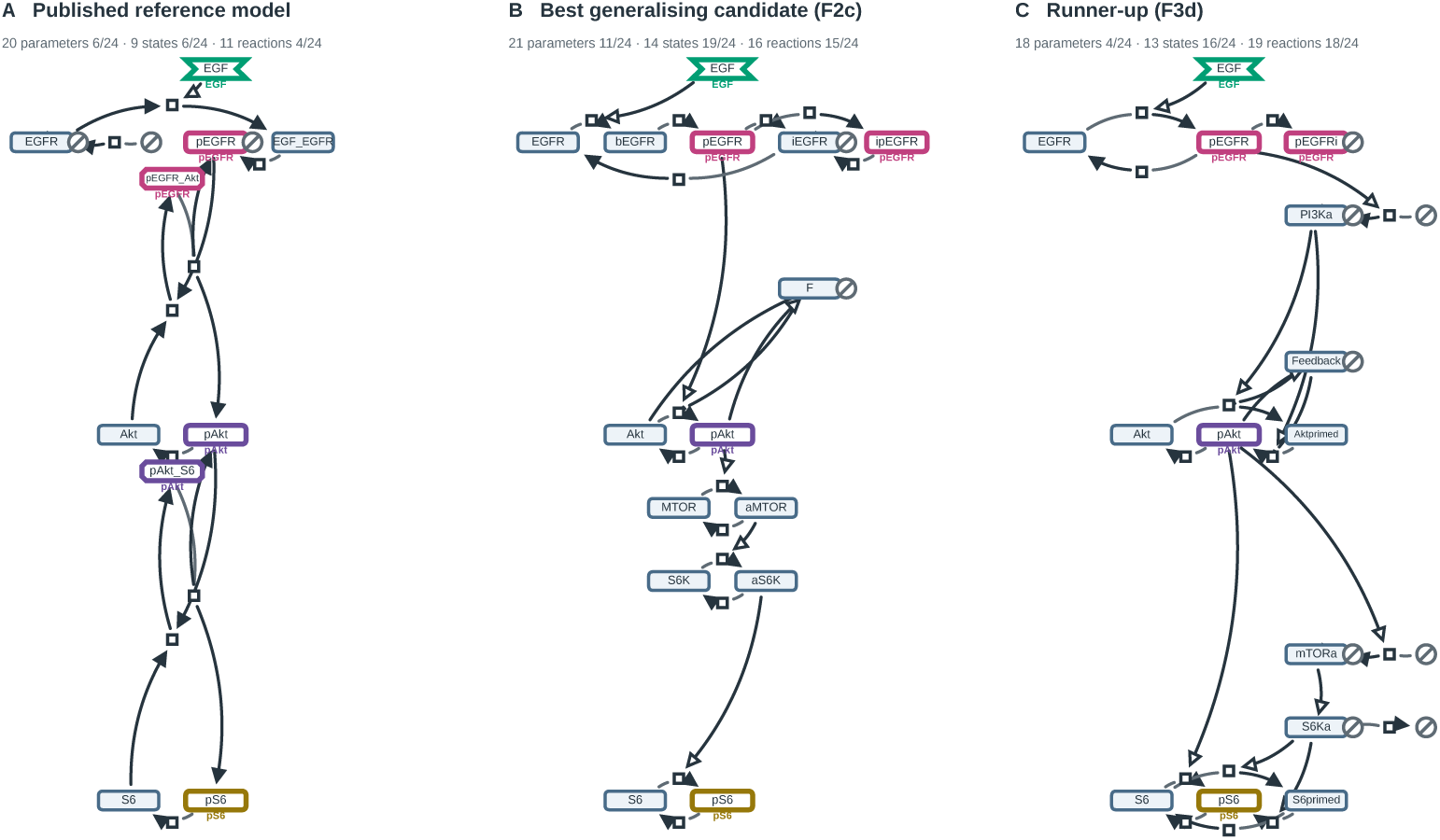
The published reference model and the two best generalising candidates. SBGN process description maps of **A,** the published reference model, **B,** the candidate with the lowest held-out NLL (F2c, held-out NLL 352.09, ΔNLL -3804.26) and **C,** the candidate with the next lowest (F3d, held-out NLL 416.59, ΔNLL -3739.75), drawn from one portable export with one deterministic layout, so no model is loaded or simulated to produce the figure. Glyphs: rounded, macromolecule; cut corners, complex; concave hexagon, perturbation; small square, process; struck circle, source or sink. Arcs: plain line, consumption; filled arrowhead, production; open arrowhead, stimulation; bar, inhibition. Entity pools claim the rows and processes the layers between them. A row is SHARED between the maps wherever something in them is the same: an entity summed into the same readout, or an entity carrying the same name. Those rows sit at identical heights in every map, in the order Figure 3 stacks the readouts, so the mechanisms can be read against one another row by row. Everything else is placed between the shared rows in proportion to how many arcs separate it from each, and every gap is made exactly as wide as the busiest map needs there, so pinning can stretch a map but never squeeze two glyphs together. A pool is outlined and named in the colour of the readout it is summed into (the EGF input likewise), the colours of Figure 3B. Entity classes follow a naming heuristic recorded with the export; modulation arcs are signed from the rate laws. Each heading gives that model’s size and its rank, smallest first, among the 24 candidates that recovered the reference connections; a model that missed one is small or large for reasons unrelated to parsimony. Rows are grouped by conserved moiety, one entity in all the states any map gives it, found from the arcs: a process with one reactant and one product is a state change. Each moiety takes ONE row, the unmodified form on the left and each further state one step to the right, and every row is centred on the same axis. A complex joins neither moiety, so it sits on a half-row under the earlier of its parts. A process sits 30% of the way between the rows it joins, or 30% of a row off a row its pools share, towards the earlier row when it carries material rightwards and the later one when leftwards.

**Figure 30:**
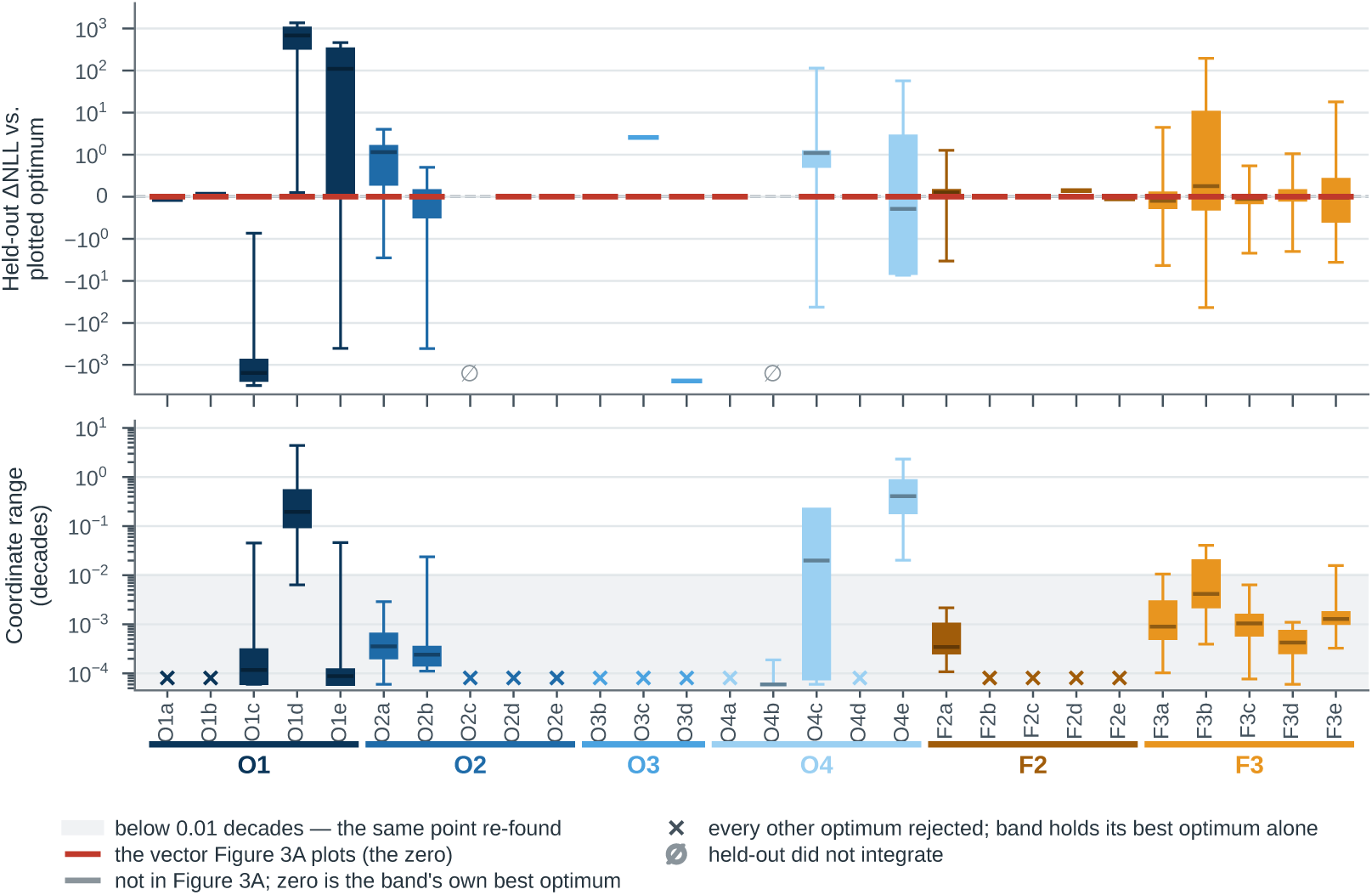
Held-out score across each Fujita model’s likelihood-ratio band. Band members are the multistart optima whose training likelihood, re-evaluated at atol 10*^−^*^16^ / rtol 10*^−^*^14^, lies within the 99% *χ*^2^ threshold (df = *k*) of the best among them; 1000 starts per model, 154–1000 converged, every band member re-scored. **Top,** held-out NLL of each band member relative to the vector Figure 3A plots (red), on a symmetric log scale. **Bottom,** per-coordinate range across the band in decades; the shaded region (below 0.01 decades) is the same point re-found, and a cross marks a band holding only its best optimum.

## F Reference models in SBGN Activity Flow notation

The following diagrams cover all 18 published reference models, using the biological-activity and unknown-influence glyphs of SBGN Activity Flow (Mi et al., 2015). They depict structural dependencies between model variables: each rectangle represents a state or time-dependent input, labelled with its original SBML identifier. Blue rectangles contribute to at least one observable formula; this does not imply that these states are individually measured. A diamond at the target end denotes a dependency whose sign is not assigned.

Arcs are extracted from kinetic laws and assignment or rate rules, with algebraic definitions expanded where necessary. Reaction arcs connect quantities read by a rate law to states with nonzero net stoichiometric changes. Self-loops are retained: they can represent depletion or turnover, rather than biological feedback. These equation-level abstractions omit parameter values, stoichiometric magnitudes, initial conditions and observable formulas; they do not establish biological causality or replace the SBML models. They are distinct from the collapsed anchor-route graphs used in the topology analysis.

Sources are the SBML models and observable tables in Benchmark-Models-PEtab at revision `1763dfc`, the collection pin used for this benchmark suite. The manuscript sources include vector diagrams, the extracted dependencies and file hashes in `paper/reference_activity_flo w/`, together with the extraction and rendering scripts.

**Figure 31:**
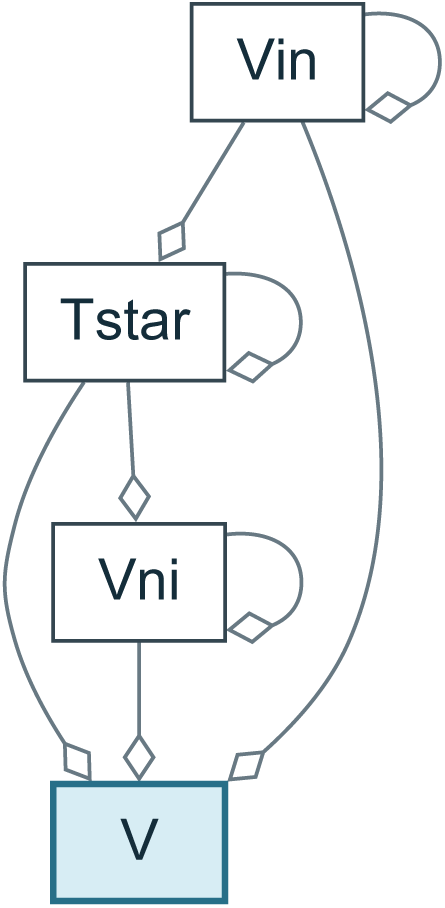
Perelson reference model. SBGN AF abstraction of `Perelson_Science1996`: 4 nodes and 8 dependency arcs. Blue nodes contribute to observable formulas; diamonds leave influence signs unspecified.

**Figure 32:**
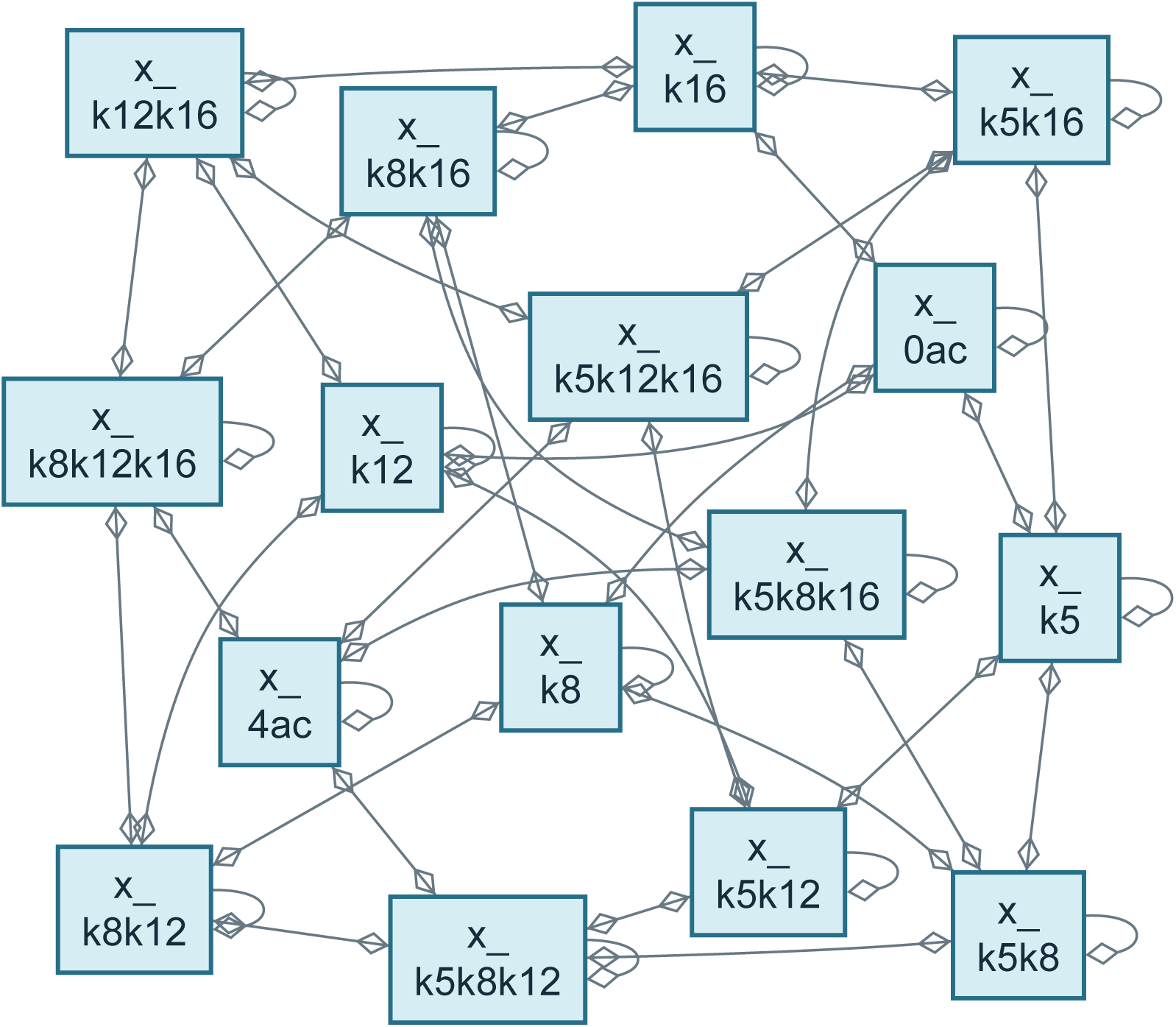
Blasi reference model. SBGN AF abstraction of `Blasi_CellSystems2016`: 16 nodes and 80 dependency arcs. Blue nodes contribute to observable formulas; diamonds leave influence signs unspecified.

**Figure 33:**
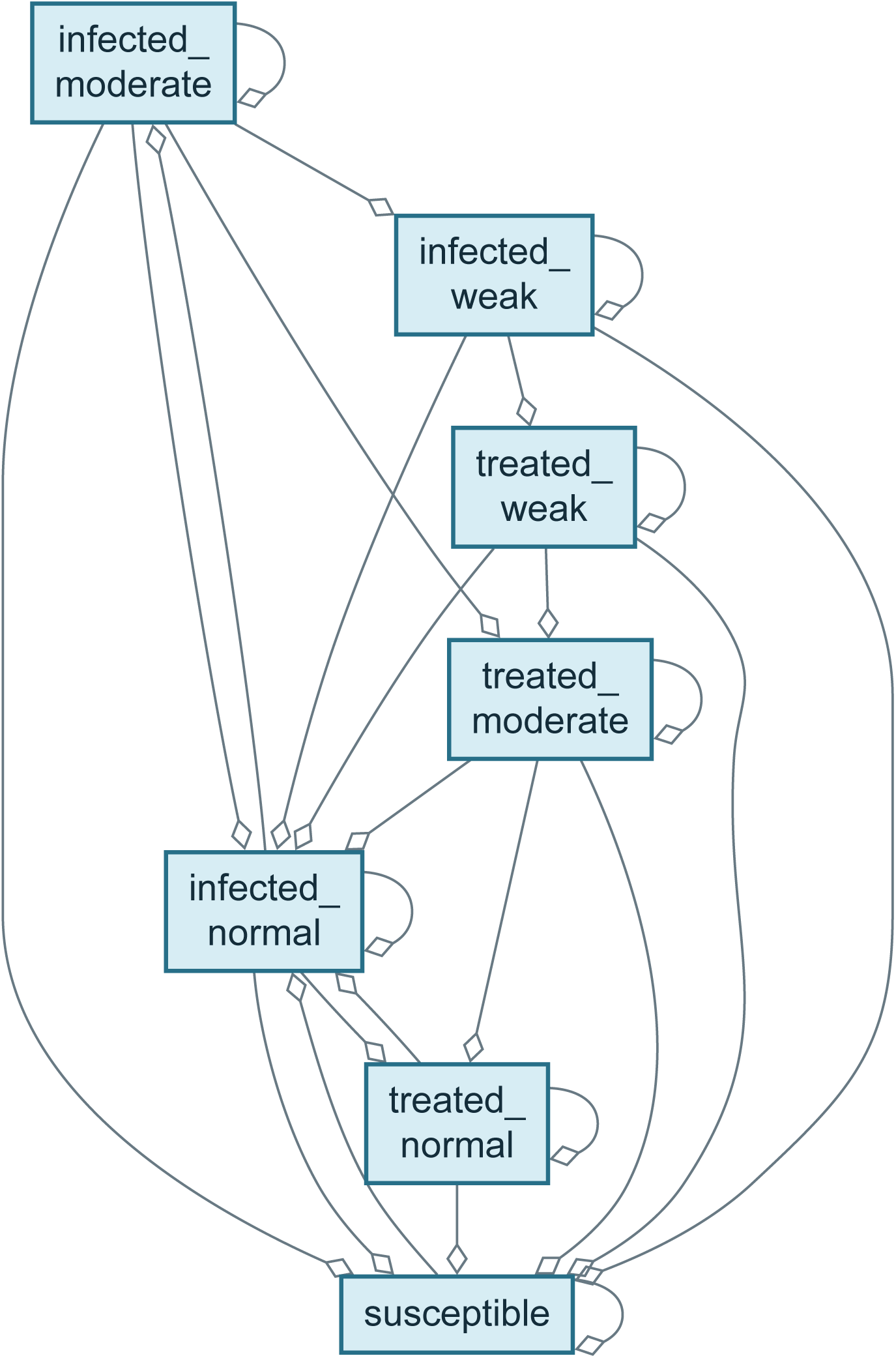
Rahman reference model. SBGN AF abstraction of `Rahman_MBS2016`: 7 nodes and 26 dependency arcs. Blue nodes contribute to observable formulas; diamonds leave influence signs unspecified.

**Figure 34:**
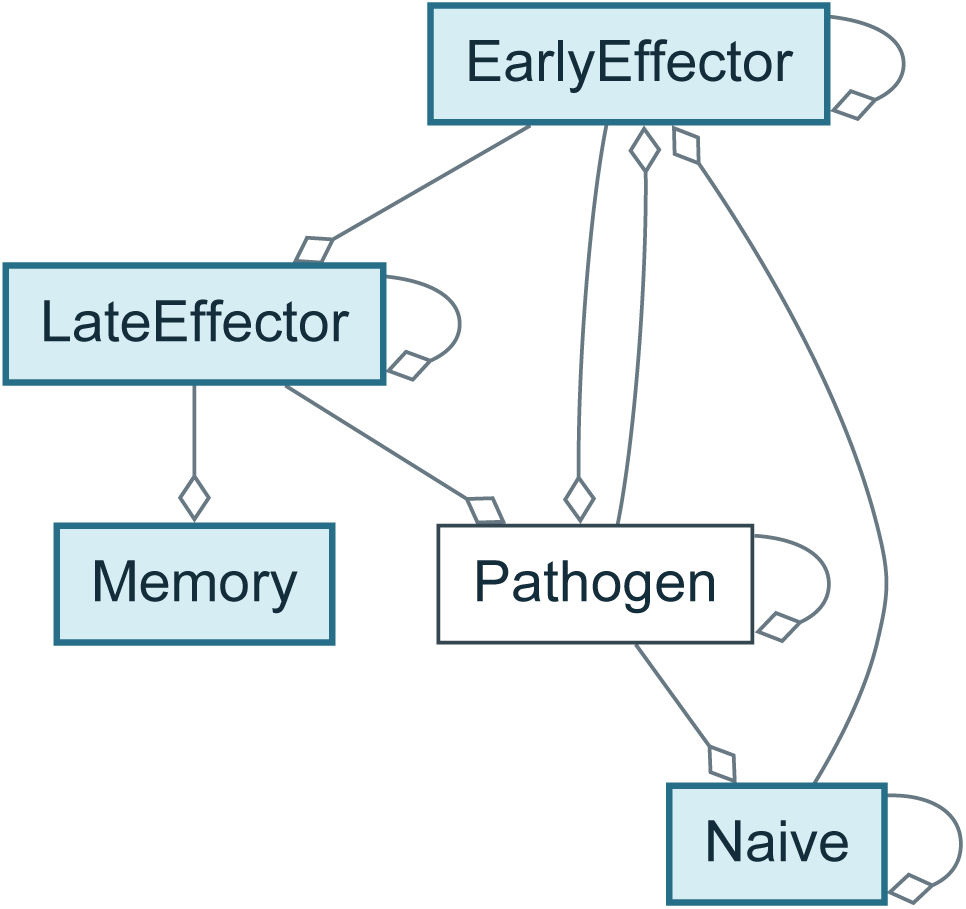
Crauste reference model. SBGN AF abstraction of `Crauste_CellSystems2017`: 5 nodes and 11 dependency arcs. Blue nodes contribute to observable formulas; diamonds leave influence signs unspecified.

**Figure 35:**
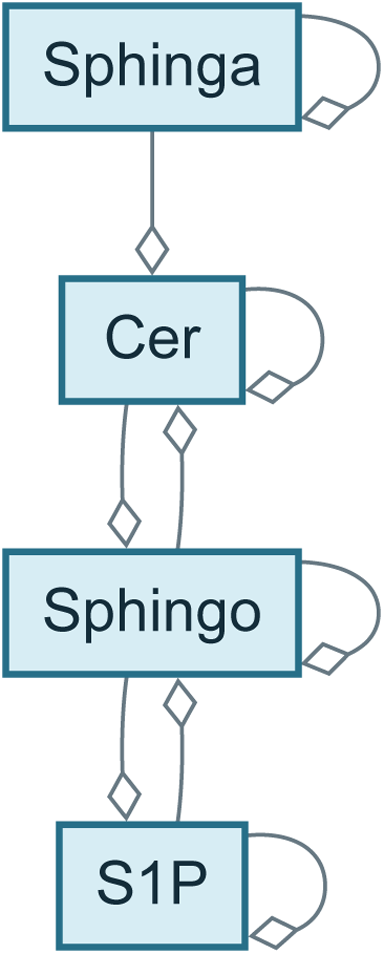
Armistead reference model. SBGN AF abstraction of `Armistead_CellDeathD is2024`: 4 nodes and 9 dependency arcs. Blue nodes contribute to observable formulas; diamonds leave influence signs unspecified.

**Figure 36:**
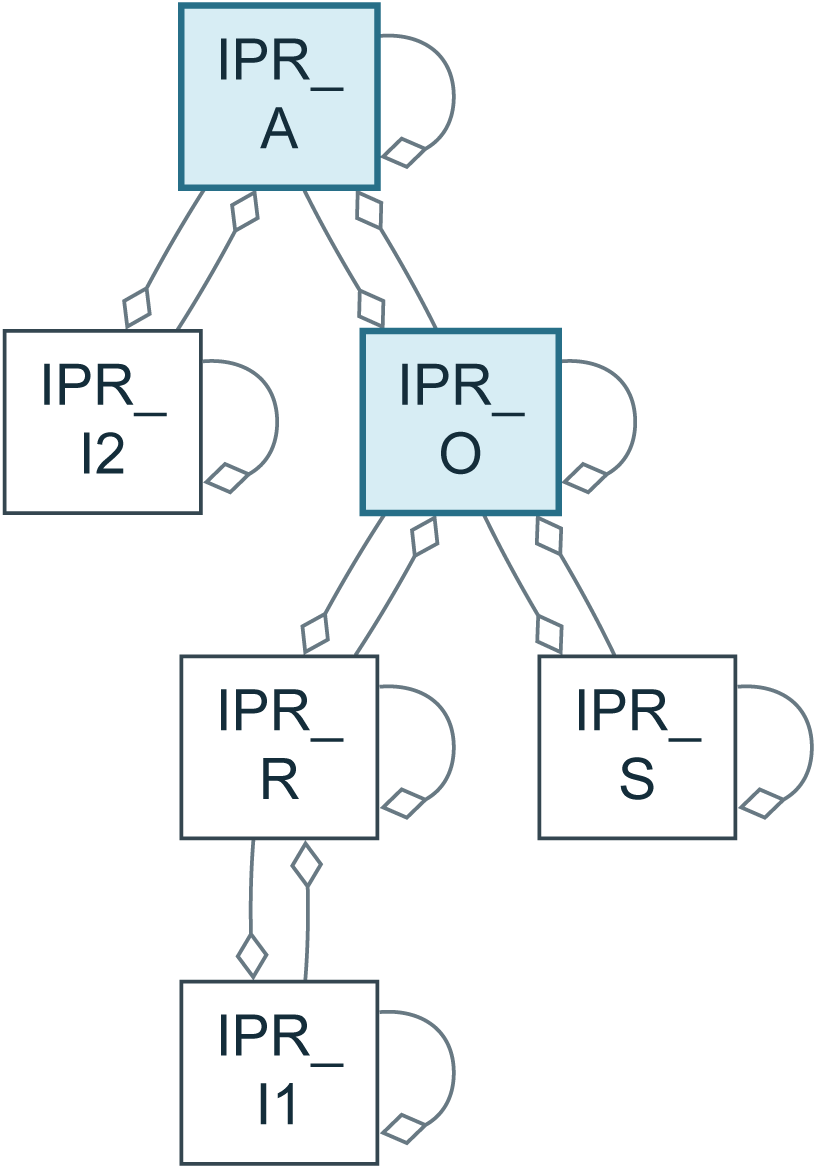
Sneyd reference model. SBGN AF abstraction of `Sneyd_PNAS2002`: 6 nodes and 16 dependency arcs. Blue nodes contribute to observable formulas; diamonds leave influence signs unspecified.

**Figure 37:**
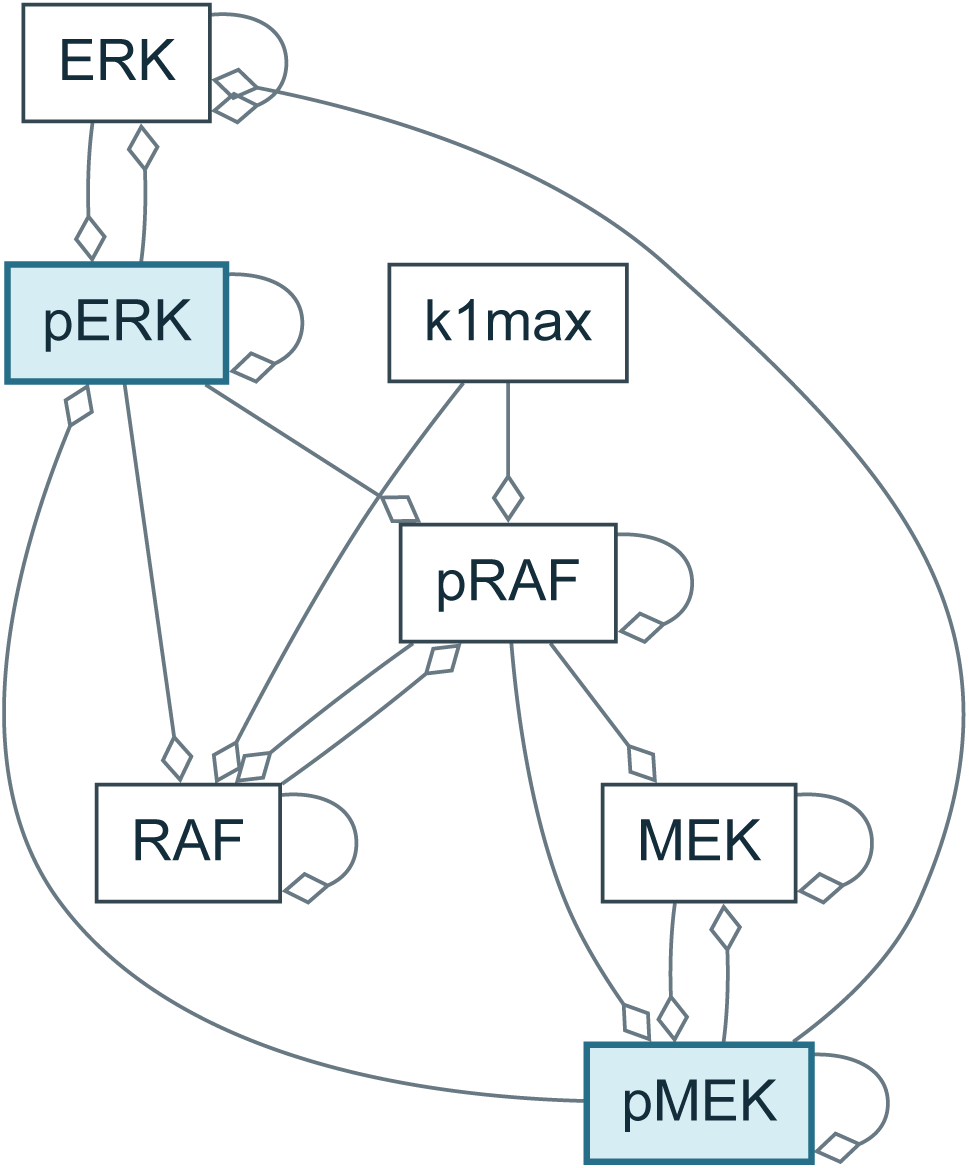
Fiedler reference model. SBGN AF abstraction of `Fiedler_BMCSystBiol2016`: 7 nodes and 20 dependency arcs. Blue nodes contribute to observable formulas; diamonds leave influence signs unspecified.

**Figure 38:**
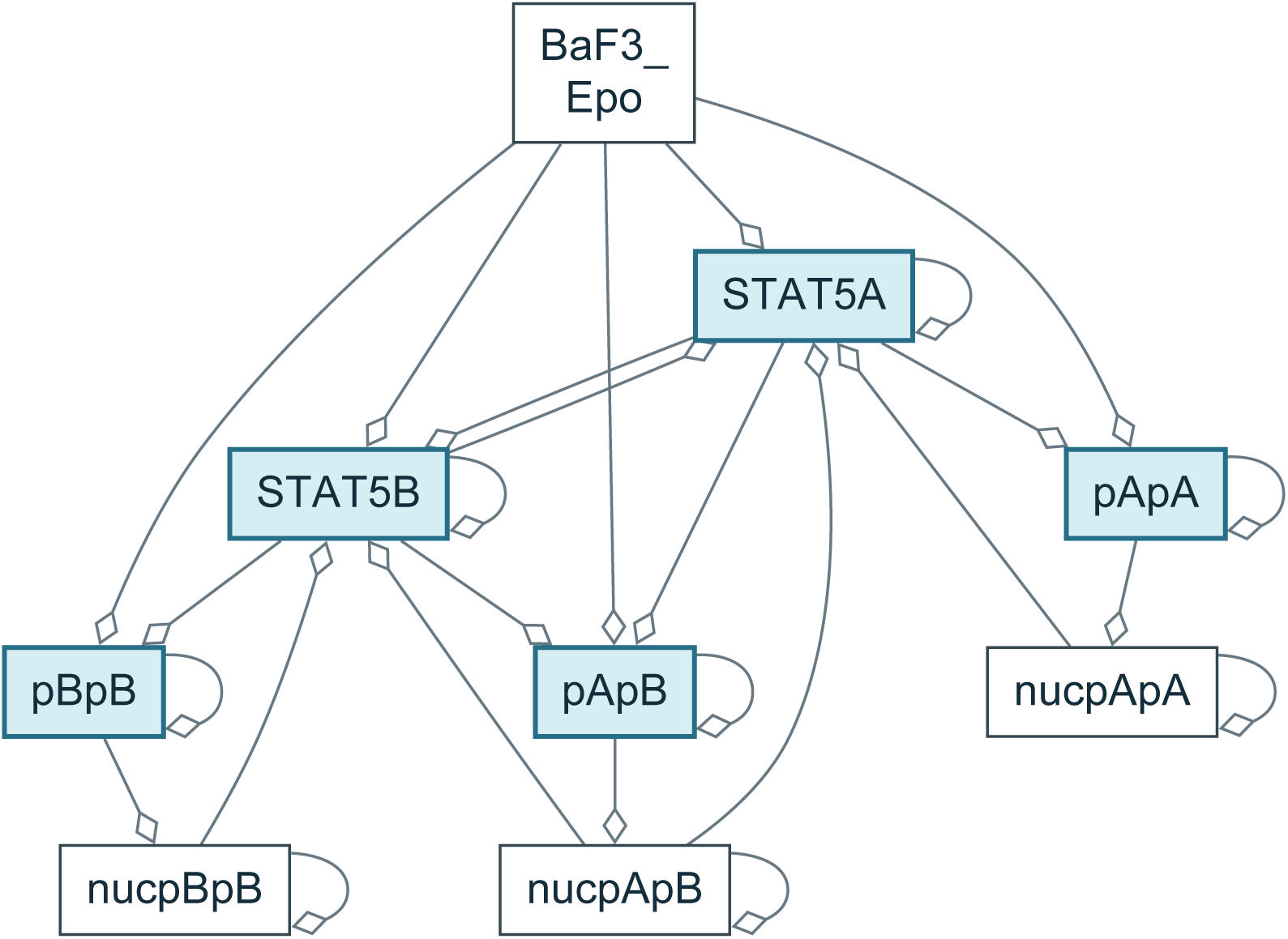
Boehm reference model. SBGN AF abstraction of `Boehm_JProteomeRes2014`: 9 nodes and 26 dependency arcs. Blue nodes contribute to observable formulas; diamonds leave influence signs unspecified.

**Figure 39:**
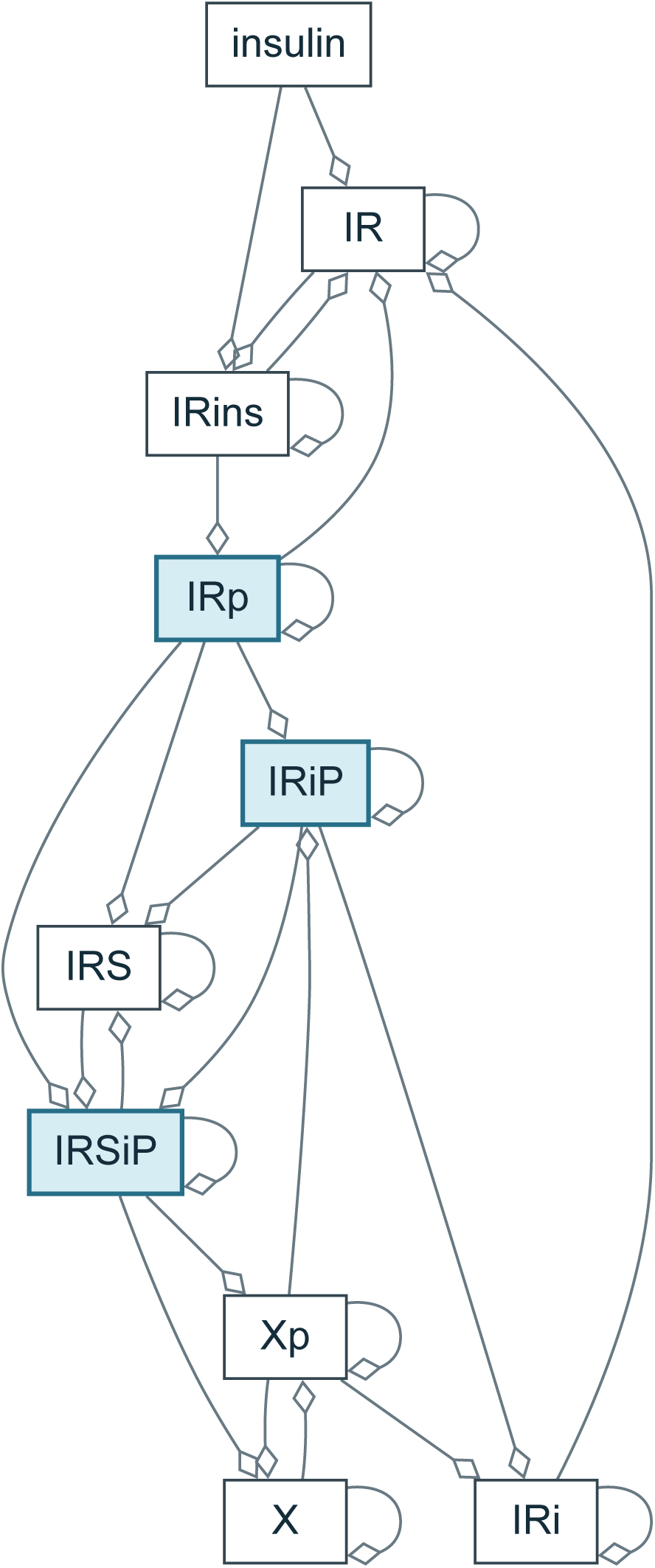
Brännmark reference model. SBGN AF abstraction of `Brannmark_JBC2010`: 10 nodes and 30 dependency arcs. Blue nodes contribute to observable formulas; diamonds leave influence signs unspecified.

**Figure 40:**
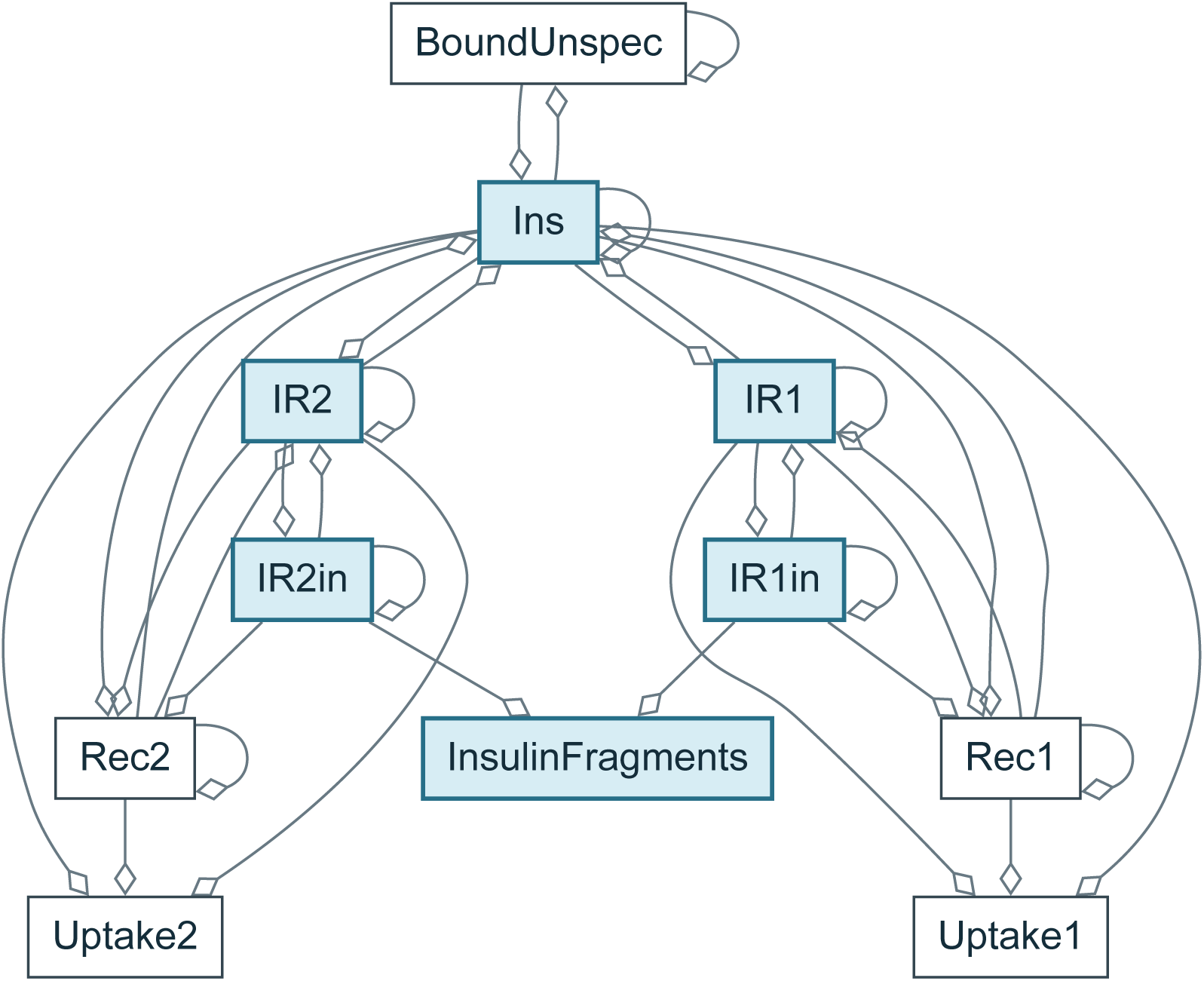
Schwen reference model. SBGN AF abstraction of `Schwen_PONE2014`: 11 nodes and 36 dependency arcs. Blue nodes contribute to observable formulas; diamonds leave influence signs unspecified.

**Figure 41:**
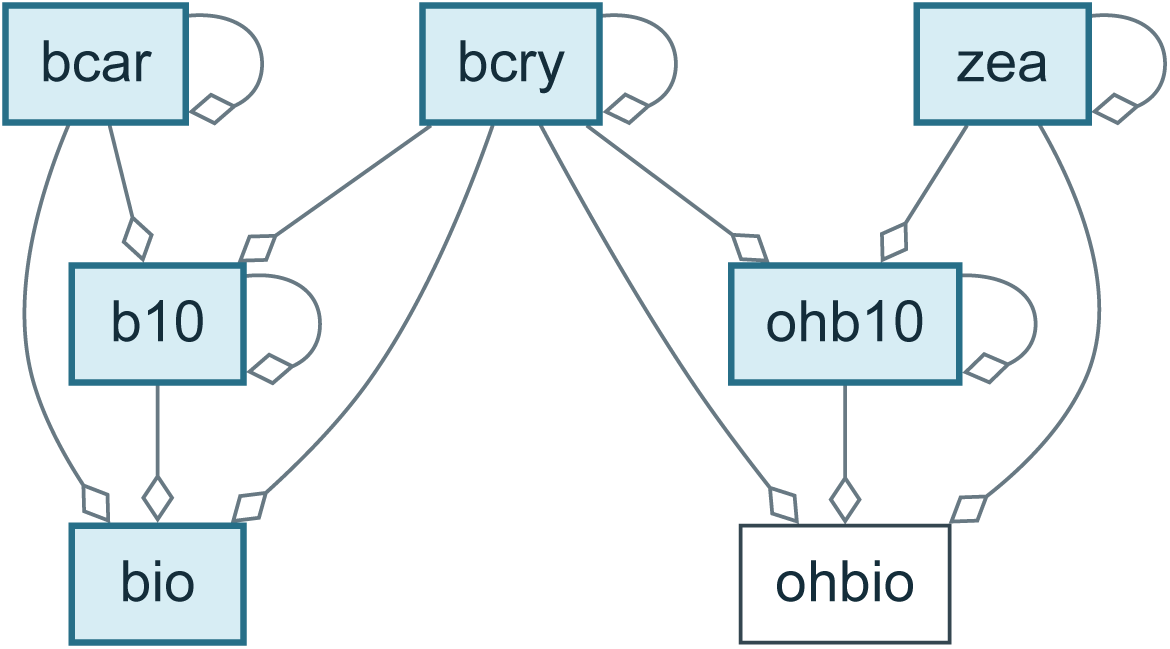
Bruno reference model. SBGN AF abstraction of `Bruno_JExpBot2016`: 7 nodes and 15 dependency arcs. Blue nodes contribute to observable formulas; diamonds leave influence signs unspecified.

**Figure 42:**
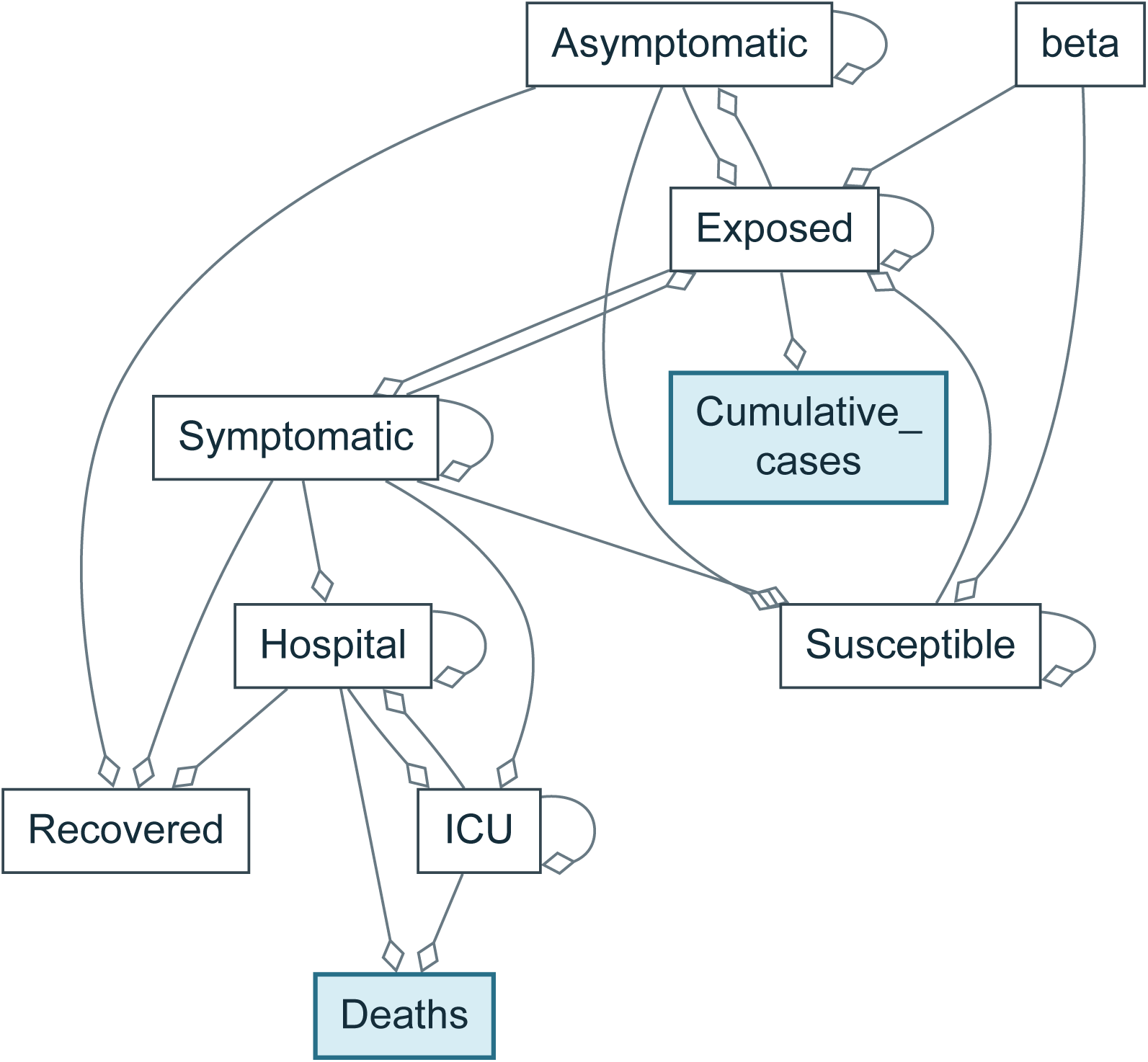
Oliveira reference model. SBGN AF abstraction of `Oliveira_NatCommun2021`: 10 nodes and 25 dependency arcs. Blue nodes contribute to observable formulas; diamonds leave influence signs unspecified.

**Figure 43:**
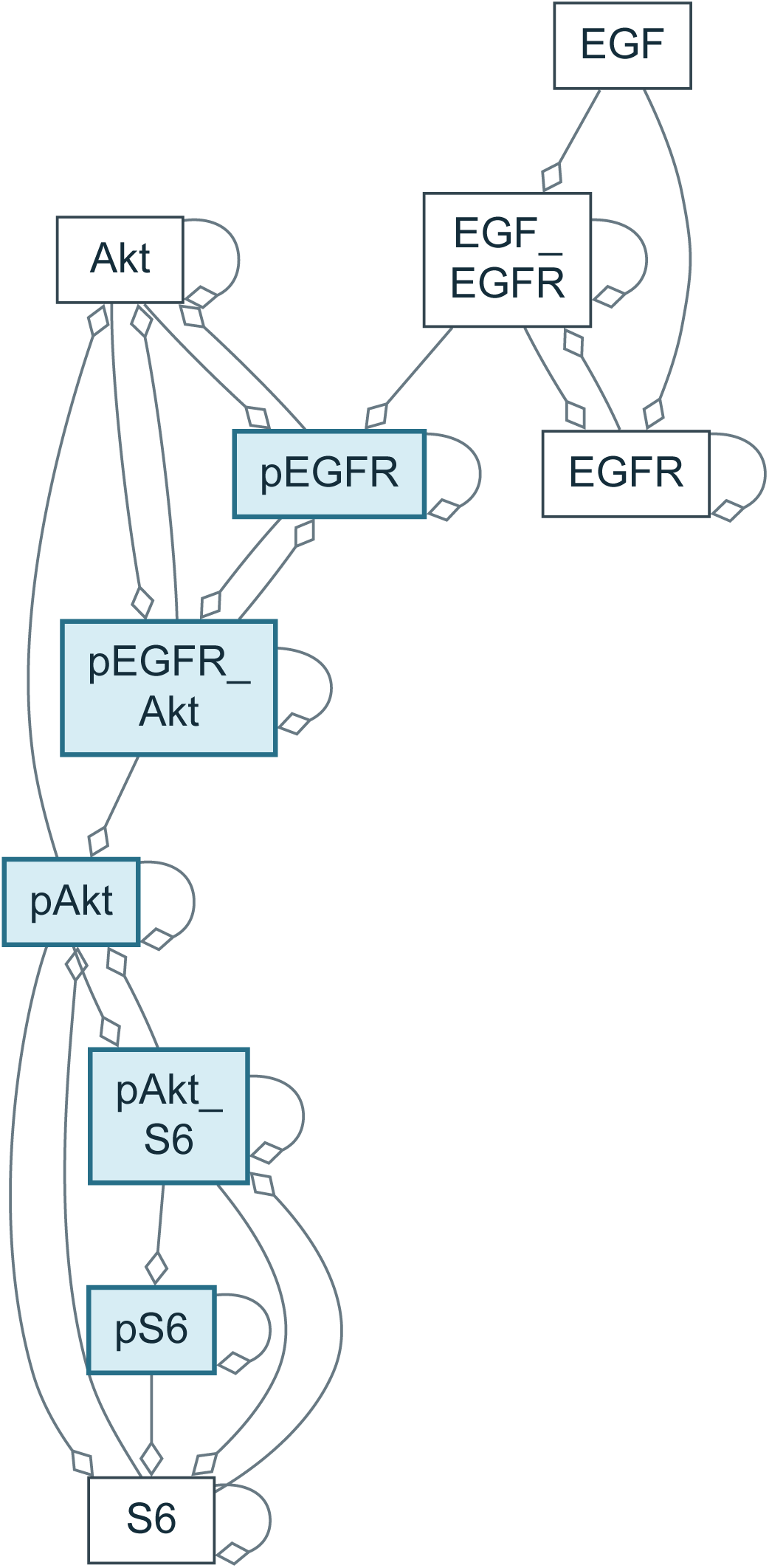
Fujita reference model. SBGN AF abstraction of `Fujita_SciSignal2010`: 10 nodes and 30 dependency arcs. Blue nodes contribute to observable formulas; diamonds leave influence signs unspecified.

**Figure 44:**
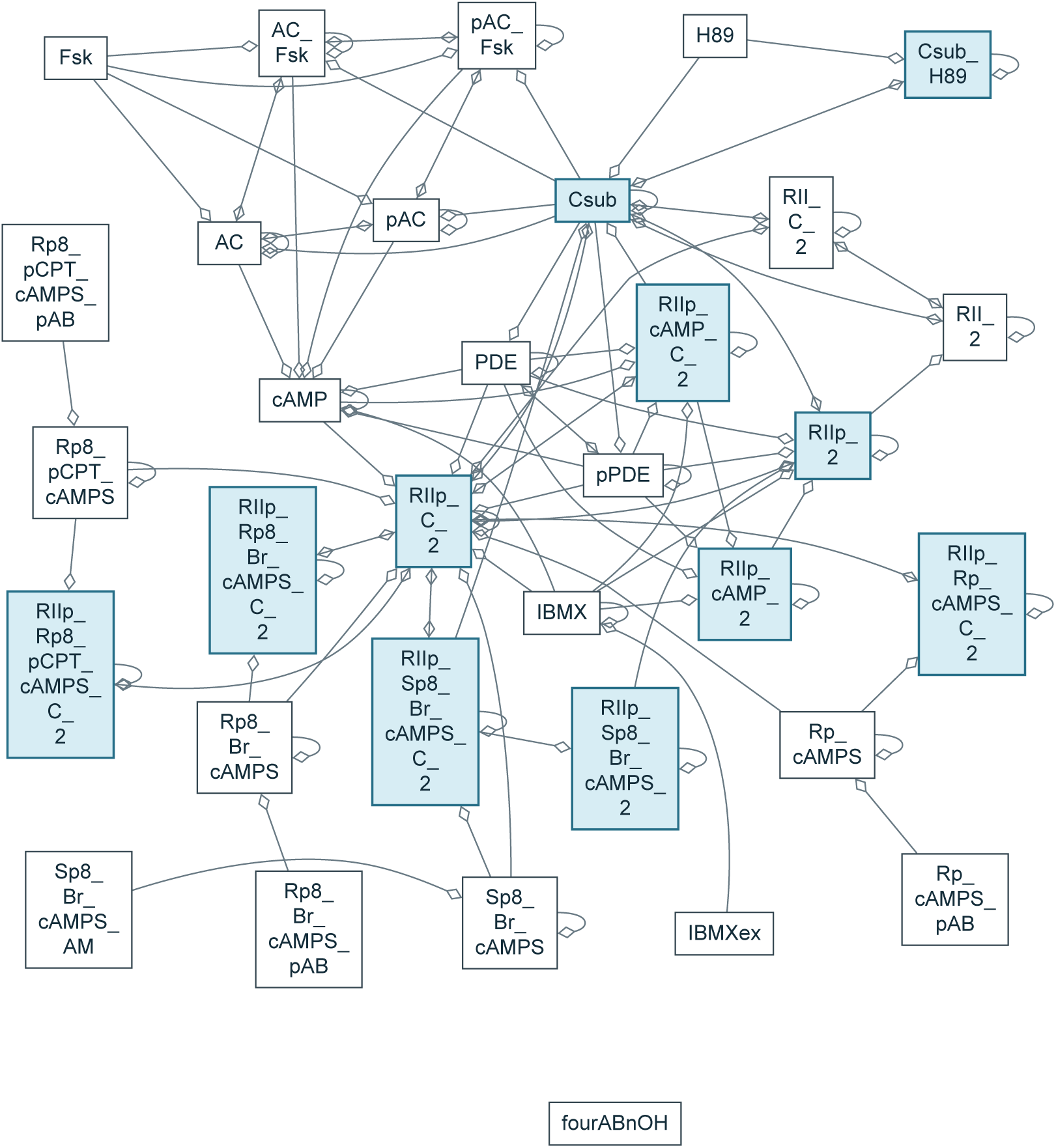
Isensee reference model. SBGN AF abstraction of `Isensee_JCB2018`: 33 nodes and 114 dependency arcs. Blue nodes contribute to observable formulas; diamonds leave influence signs unspecified.

**Figure 45:**
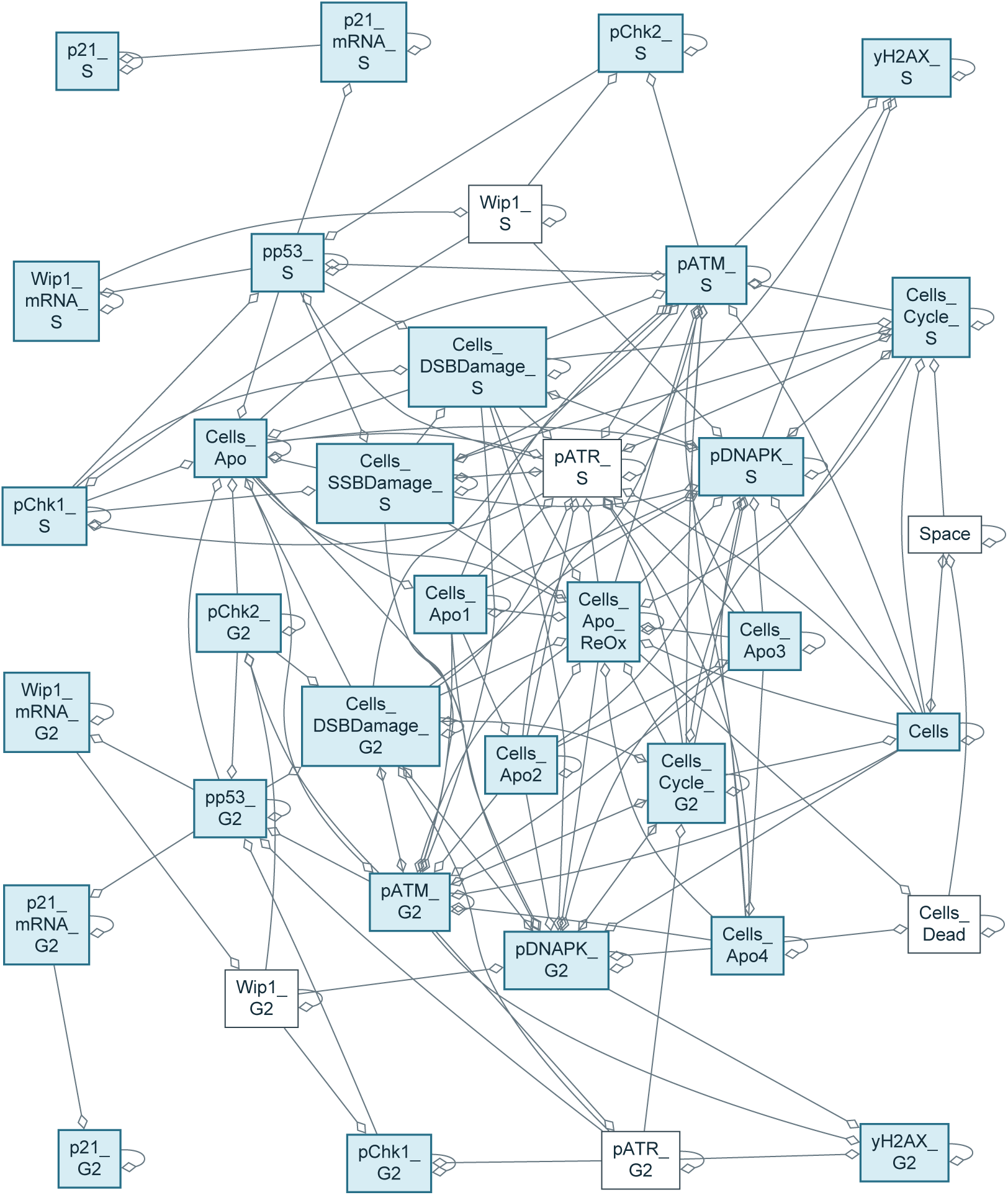
Alkan reference model. SBGN AF abstraction of `Alkan_SciSignal2018`: 36 nodes and 181 dependency arcs. Blue nodes contribute to observable formulas; diamonds leave influence signs unspecified.

**Figure 46:**
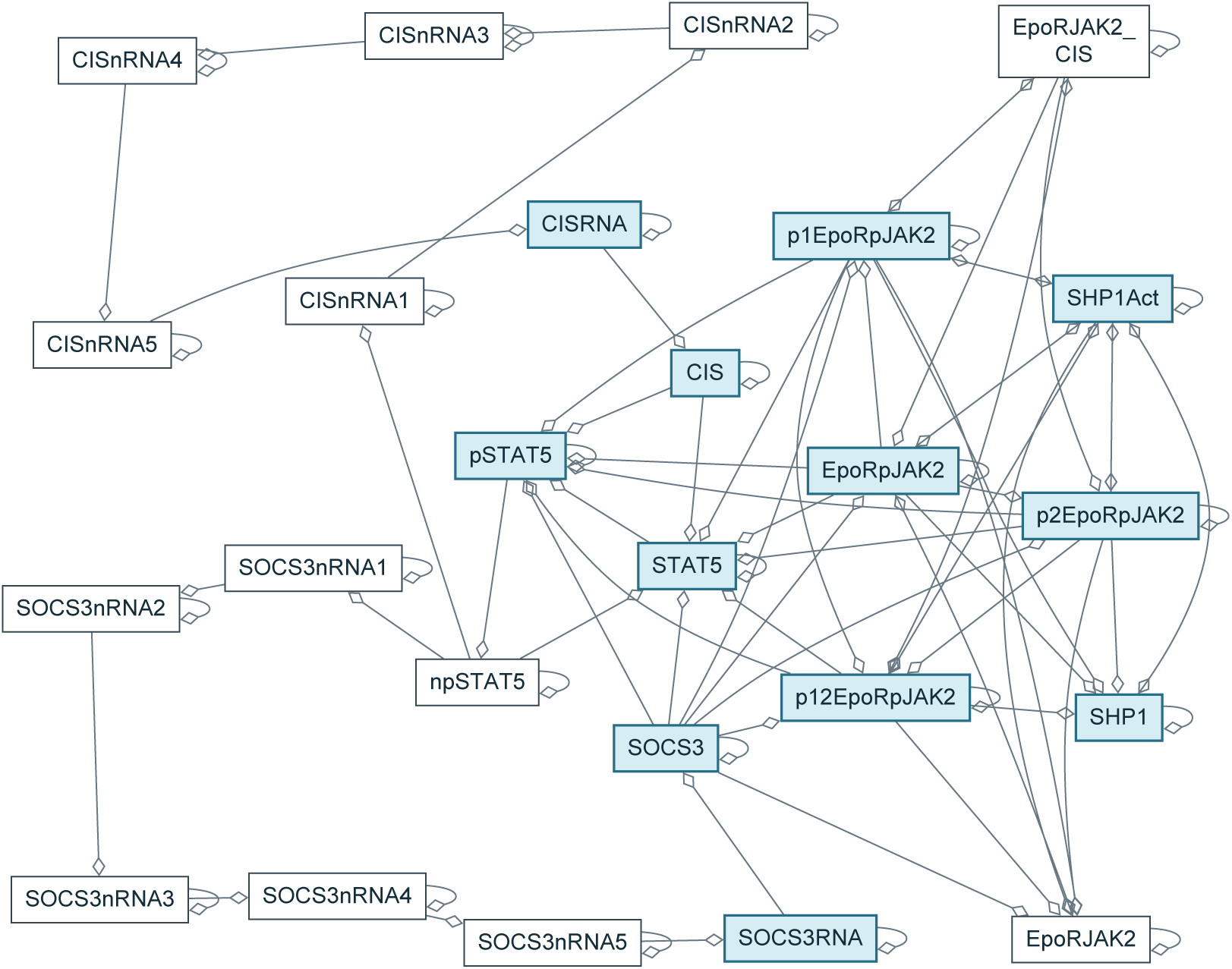
Bachmann reference model. SBGN AF abstraction of `Bachmann_MSB2011`: 25 nodes and 89 dependency arcs. Blue nodes contribute to observable formulas; diamonds leave influence signs unspecified.

**Figure 47:**
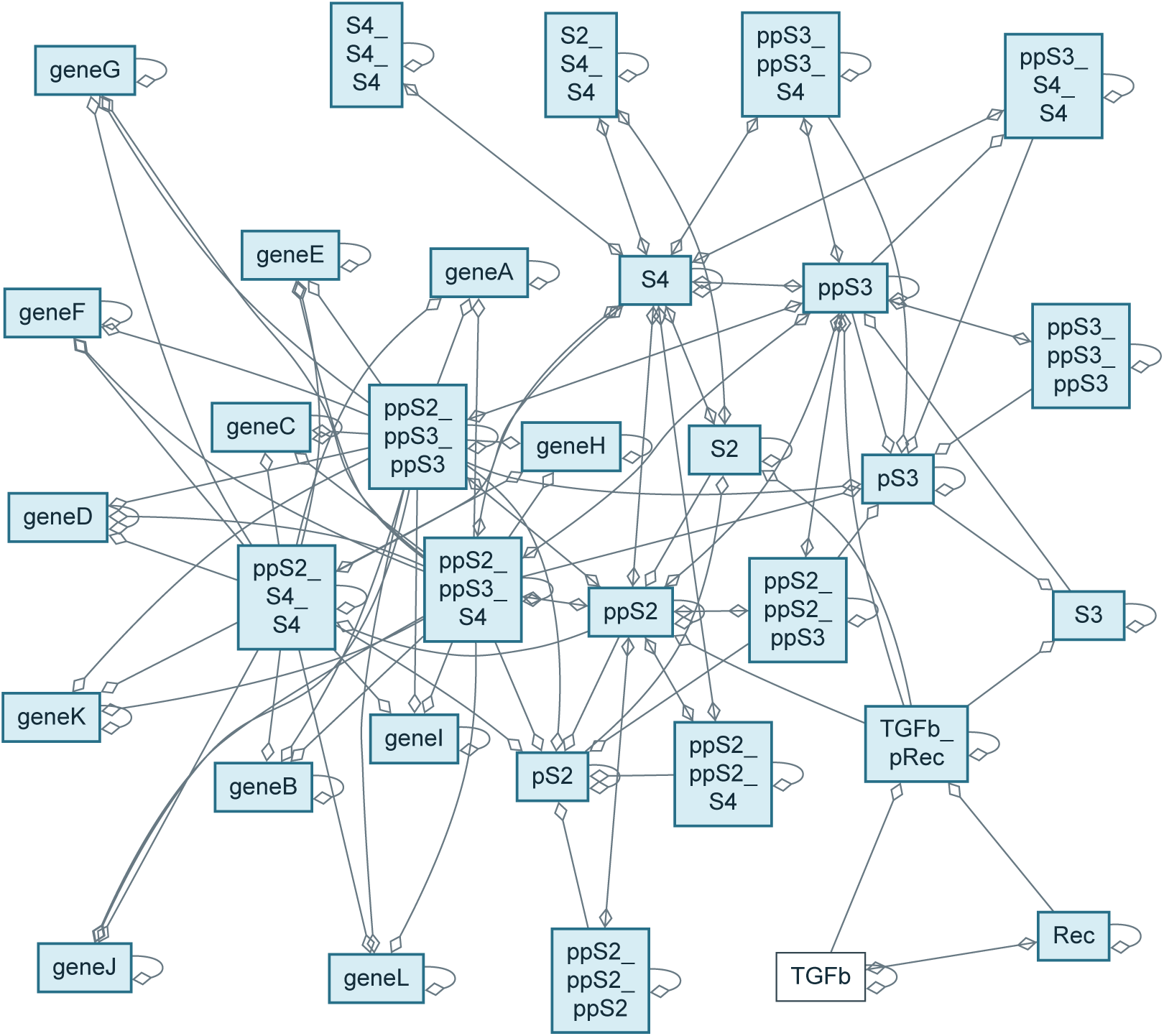
Lucarelli reference model. SBGN AF abstraction of `Lucarelli_CellSystems2 018`: 33 nodes and 141 dependency arcs. Blue nodes contribute to observable formulas; diamonds leave influence signs unspecified.

**Figure 48:**
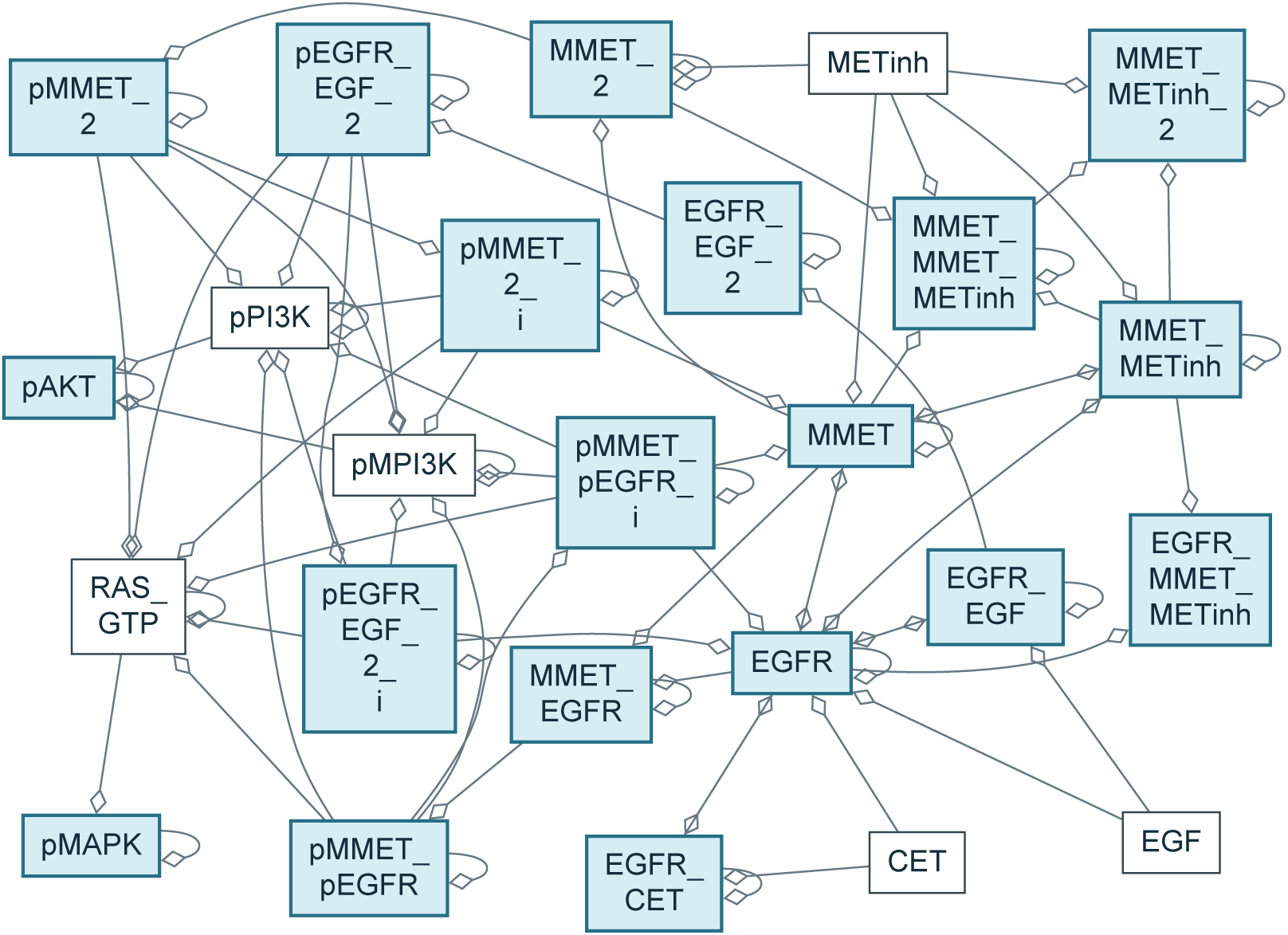
Raimundez reference model. SBGN AF abstraction of `Raimundez_PCB2020`: 25 nodes and 82 dependency arcs. Blue nodes contribute to observable formulas; diamonds leave influence signs unspecified.

## G Agent-generated Fujita model sources

The following listings contain the archived Antimony for the 28 models of Figure 3 whose sources were exported with their hashes. The label, arm, array task and selected version identify each source. Listings marked *converted from the archived SBML* are libantimony renderings of the `model.xml` those runs delivered: the bare-agent arms wrote no Antimony, so none exists to reproduce. The hash recorded for such a row is their `model.xml`, because verifying a derived listing against itself would prove nothing; that SBML is not shipped, so the annex is Antimony throughout. Every other listing is the archived file itself, preserved exactly, including anonymous identifiers and unset initializations. Numeric assignments in these source listings are not necessarily the fitted values. The matching parameter tables (including observation scales and fixed entries) and SHA-256 hashes are exported alongside them; see `fujita_models.csv`. These sources and tables document the selected structures and parameters; reproducing simulations also requires the original PEtab conditions, observables and initialization semantics. No parameters were refitted for this annex.

**Listing 1:**
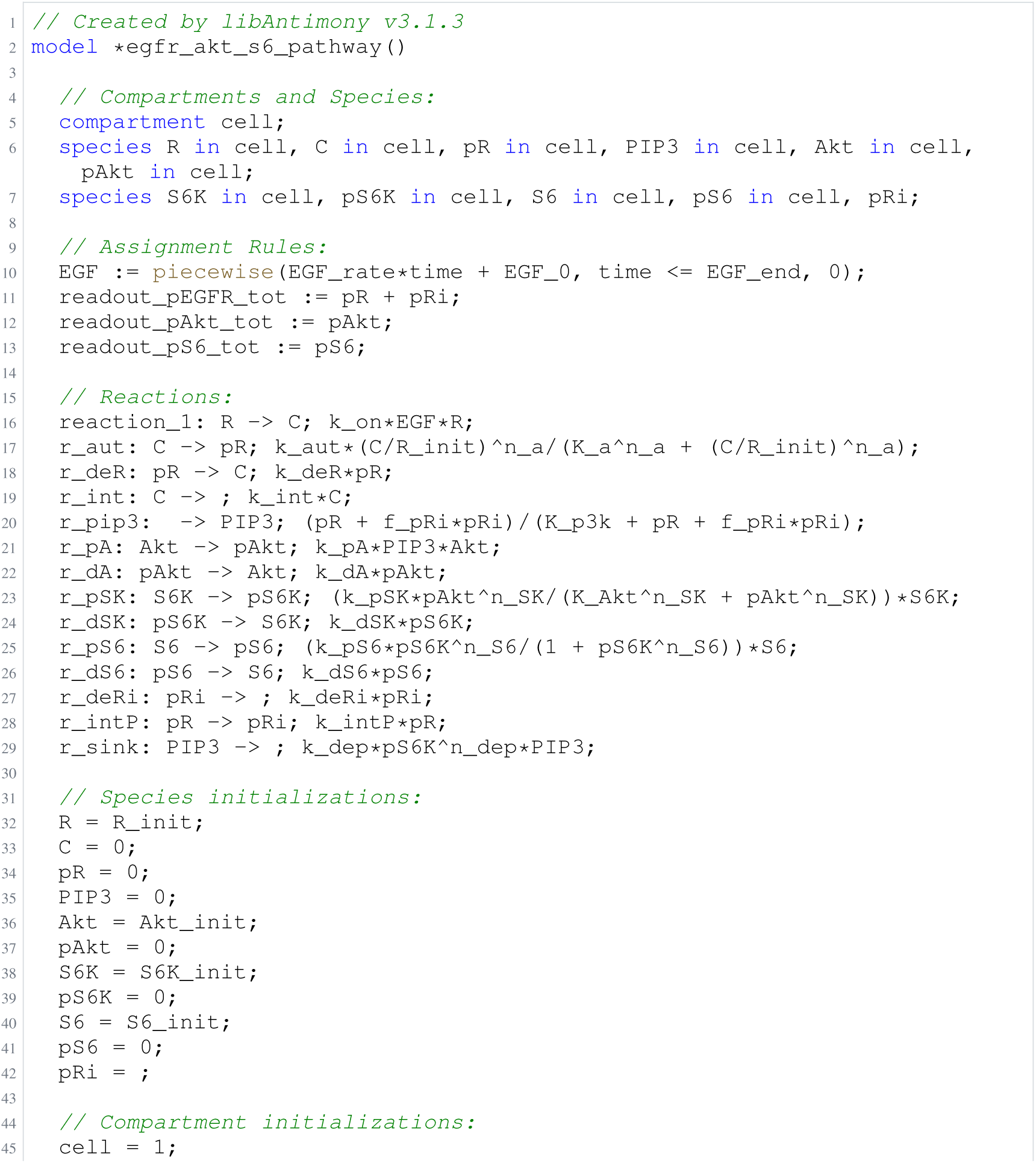
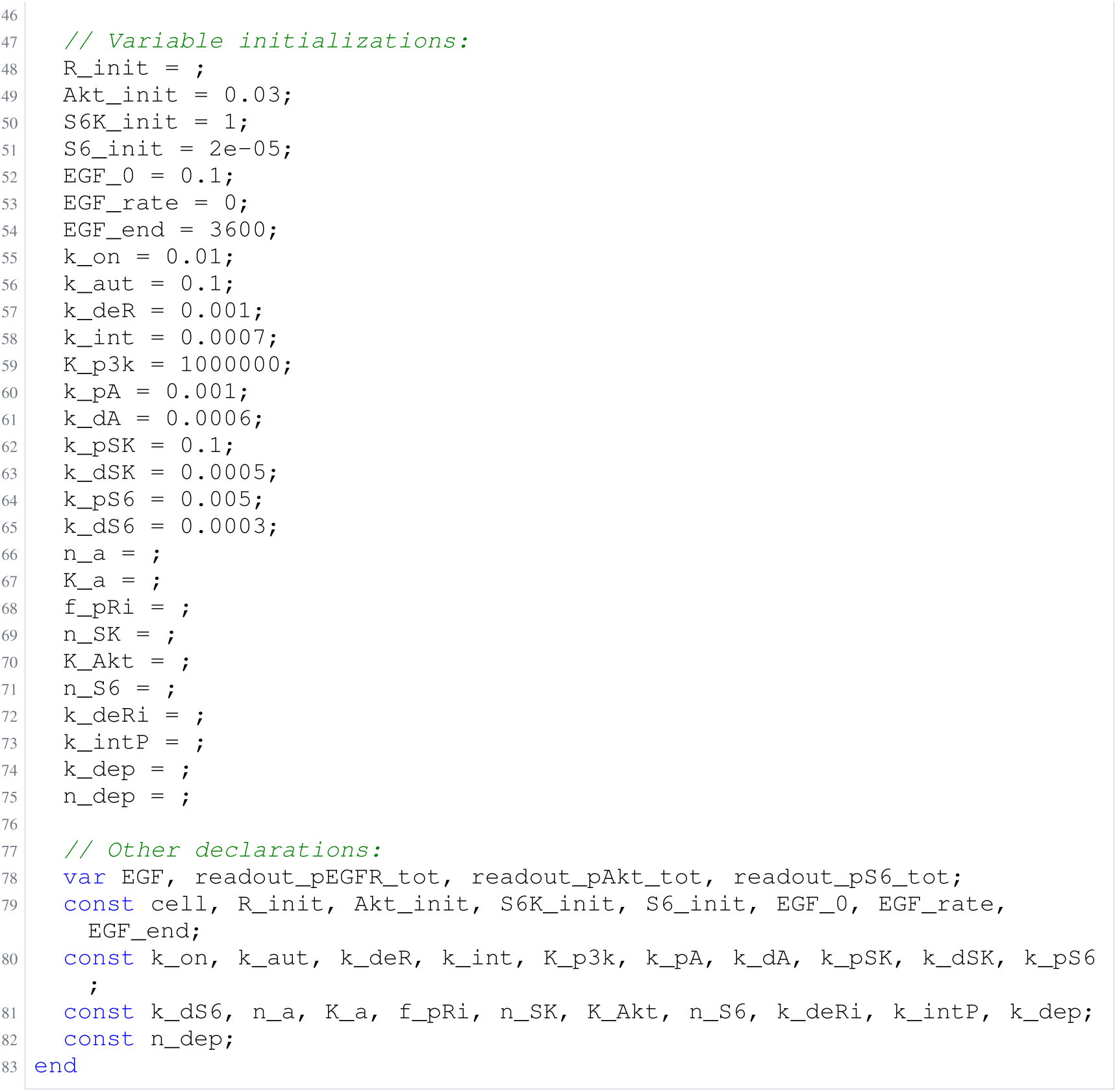
Archived Antimony for O1a: Panel, glm-5.3-flash; task 56476116_0; selected version 22.

**Listing 2:**
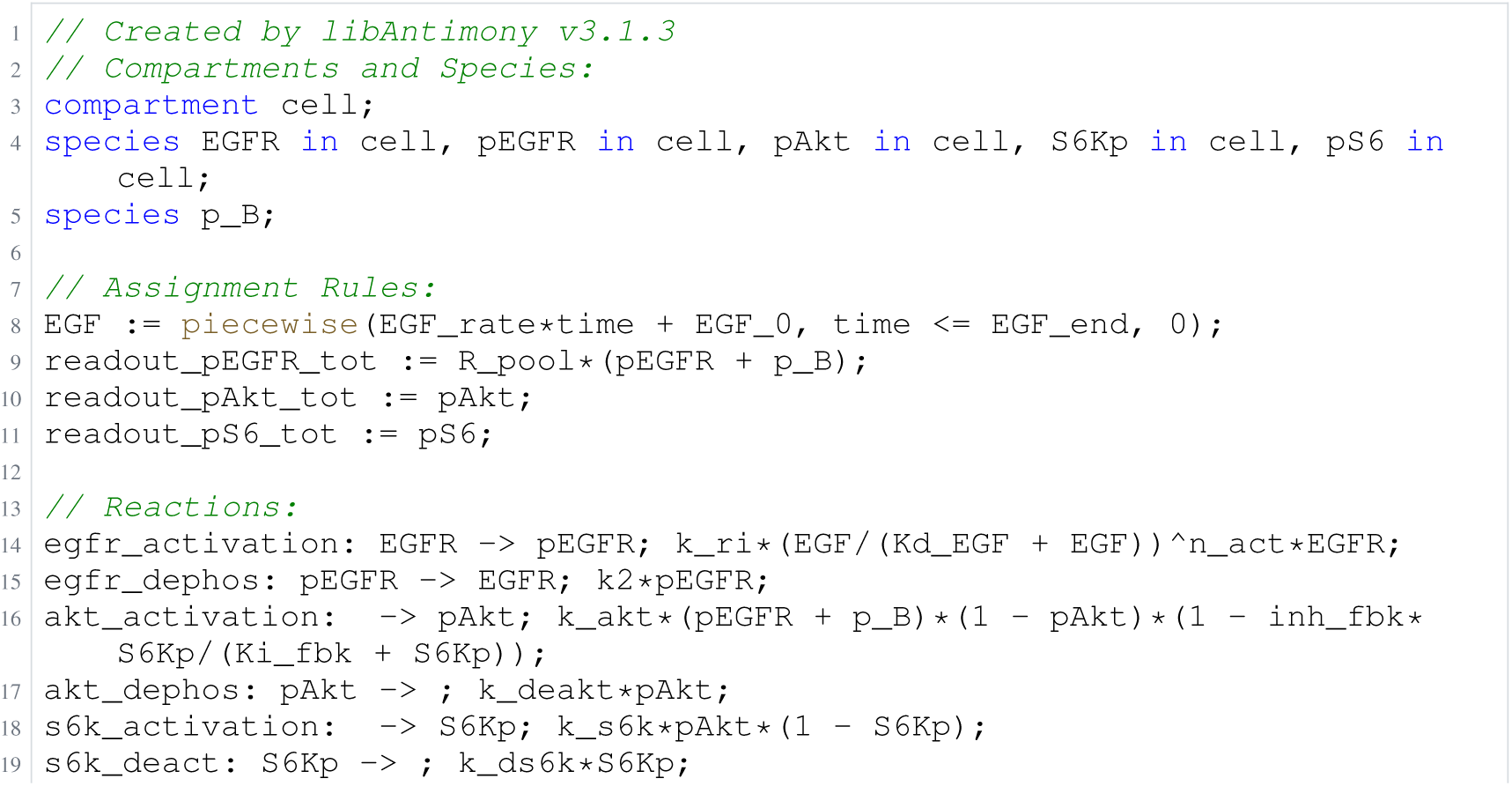
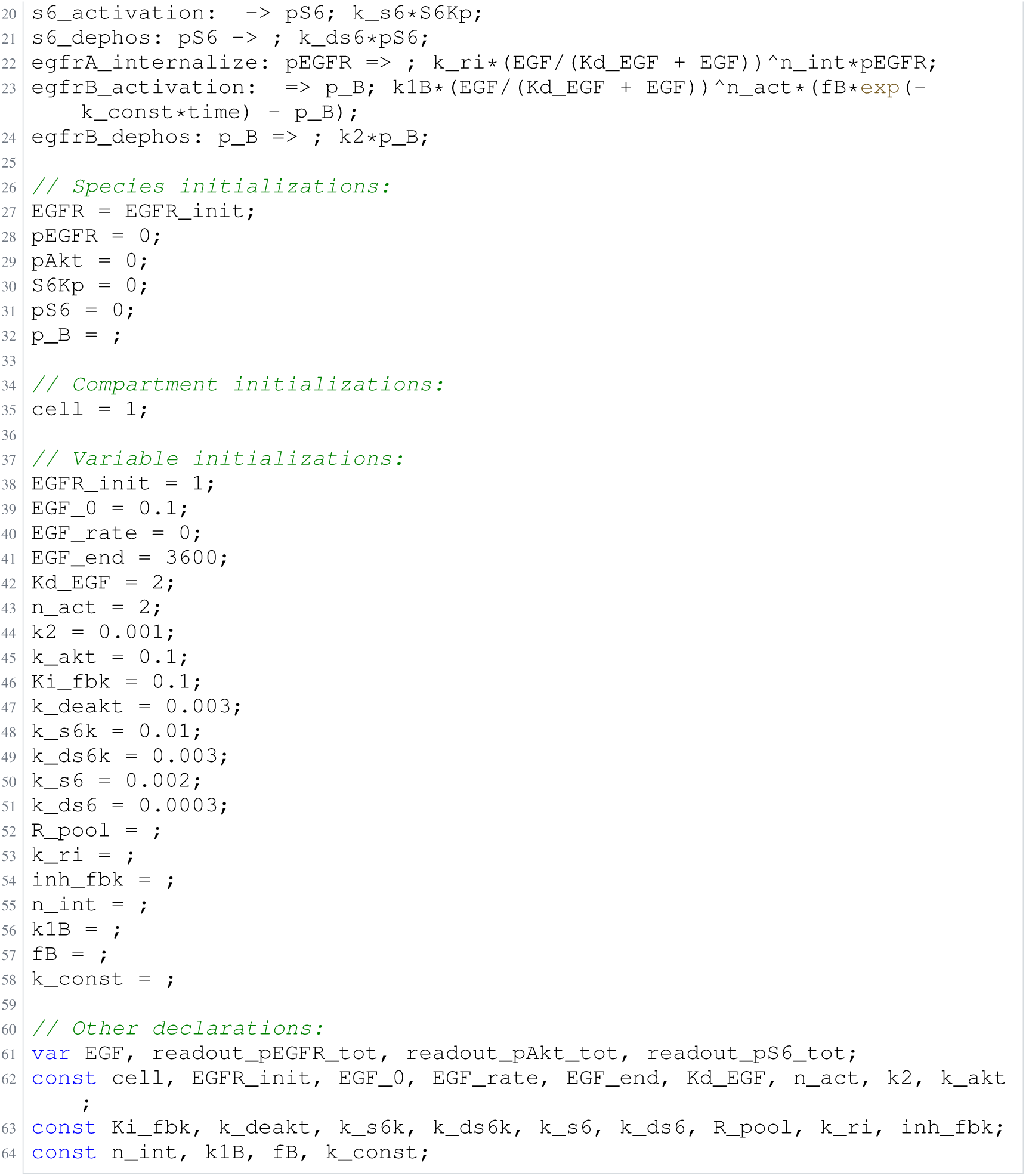
Archived Antimony for O1b: Panel, glm-5.3-flash; task 56476116_1; selected version 8.

**Listing 3:**
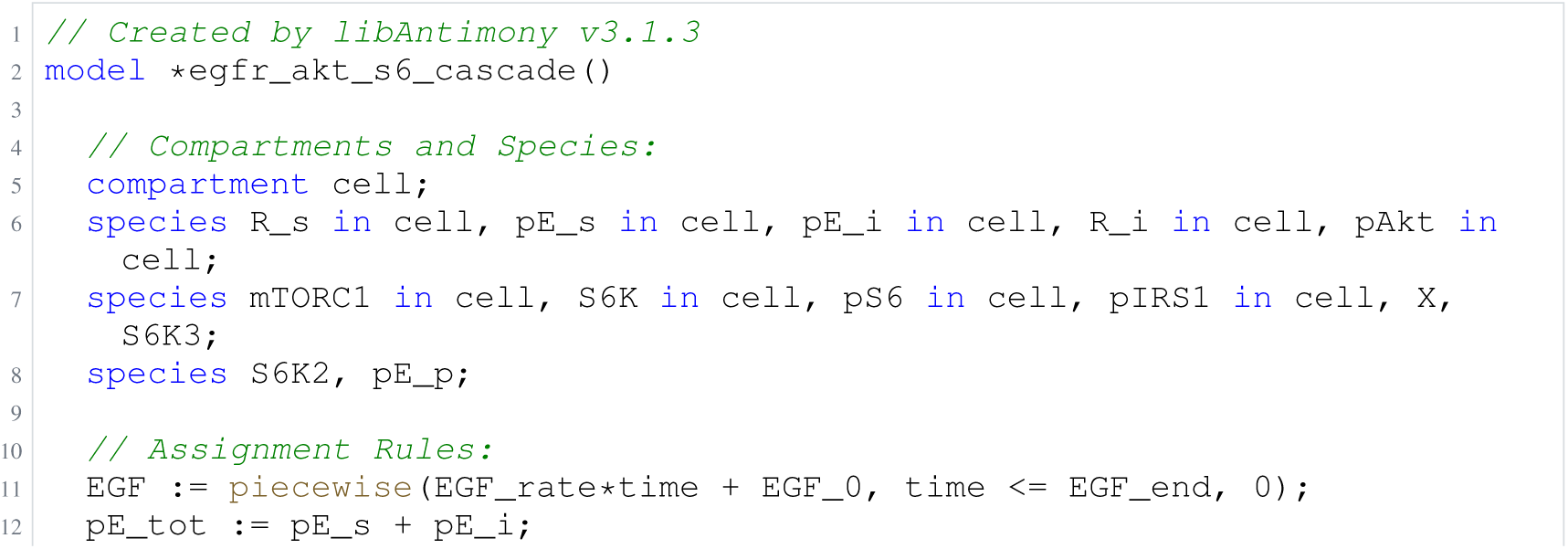
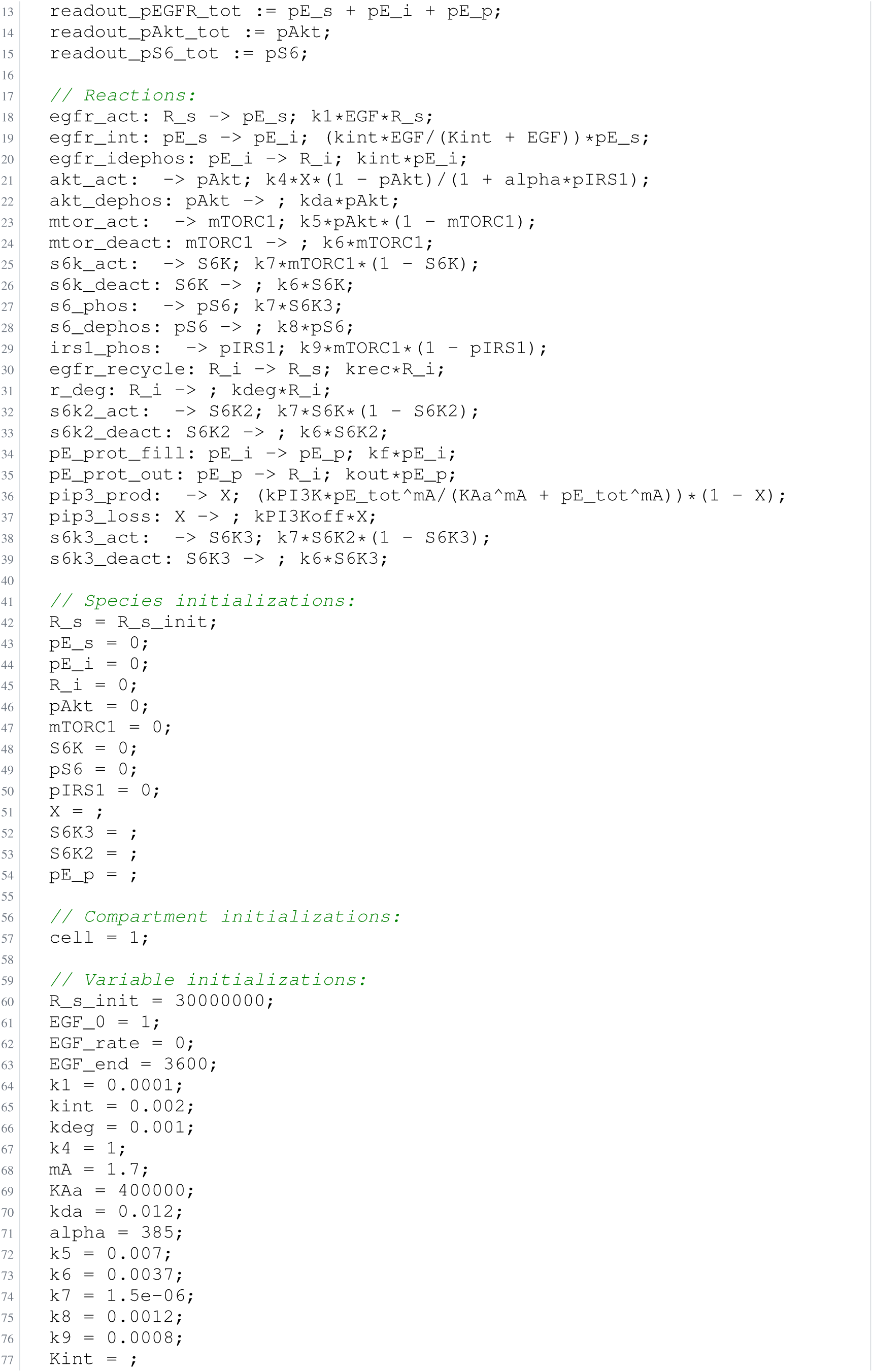
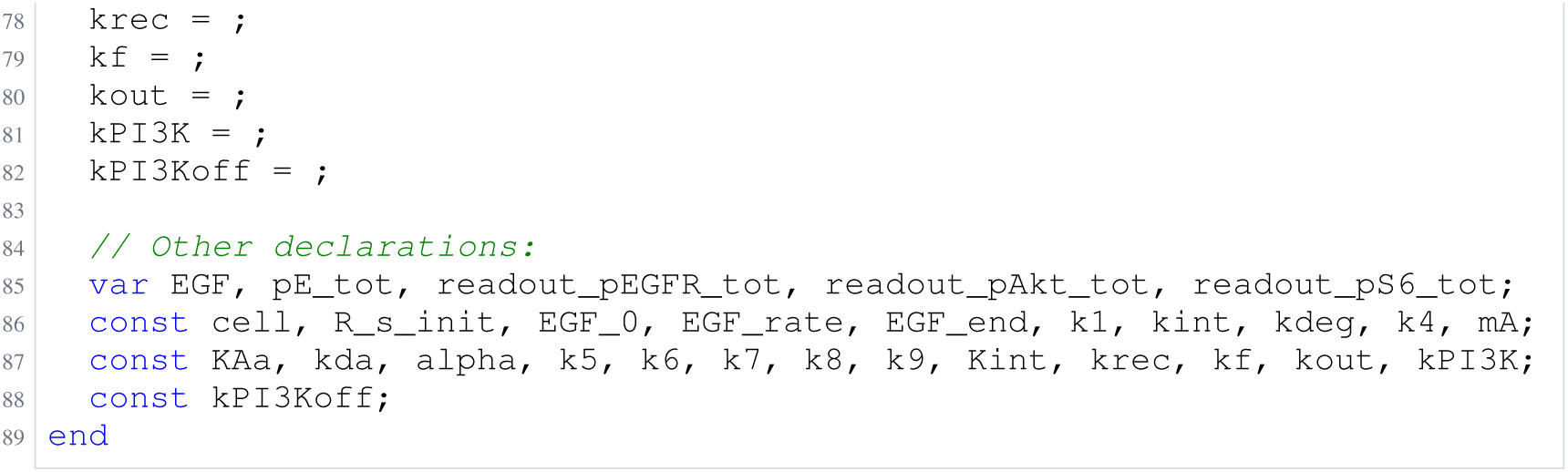
Archived Antimony for O1c: Panel, glm-5.3-flash; task 56476116_2; selected version 28.

**Listing 4:**
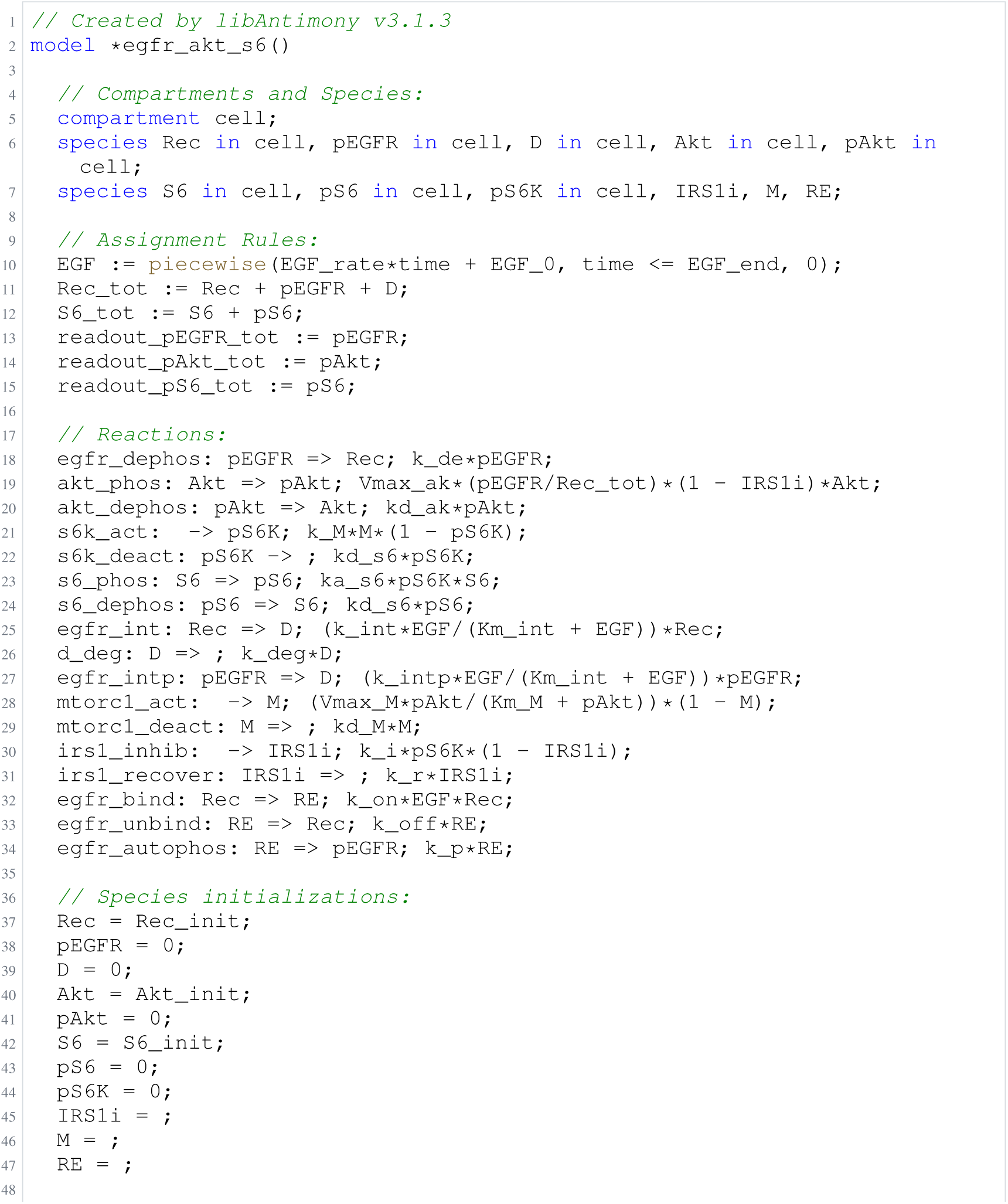
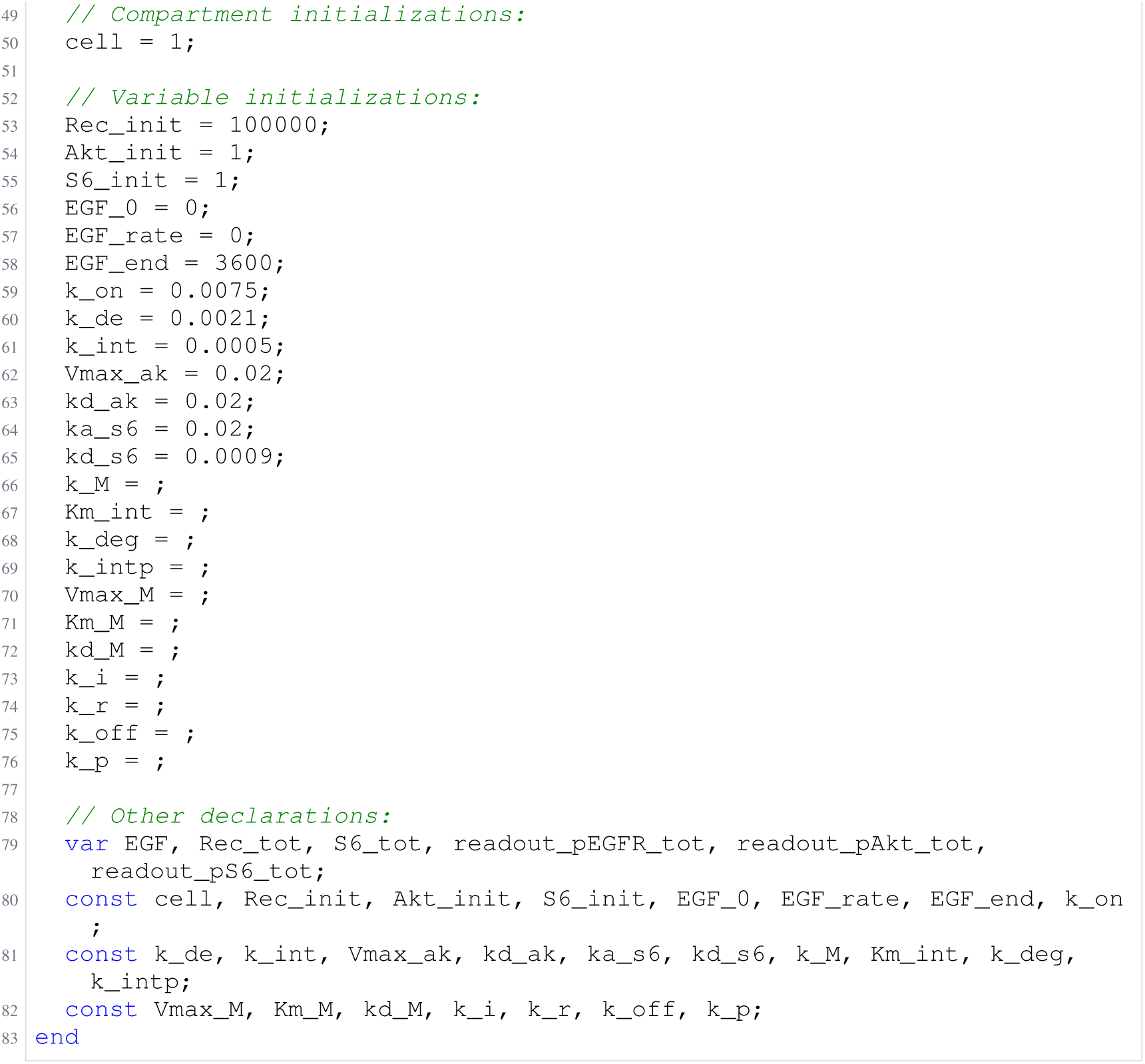
Archived Antimony for O1d: Panel, glm-5.3-flash; task 56476116_3; selected version 29.

**Listing 5:**
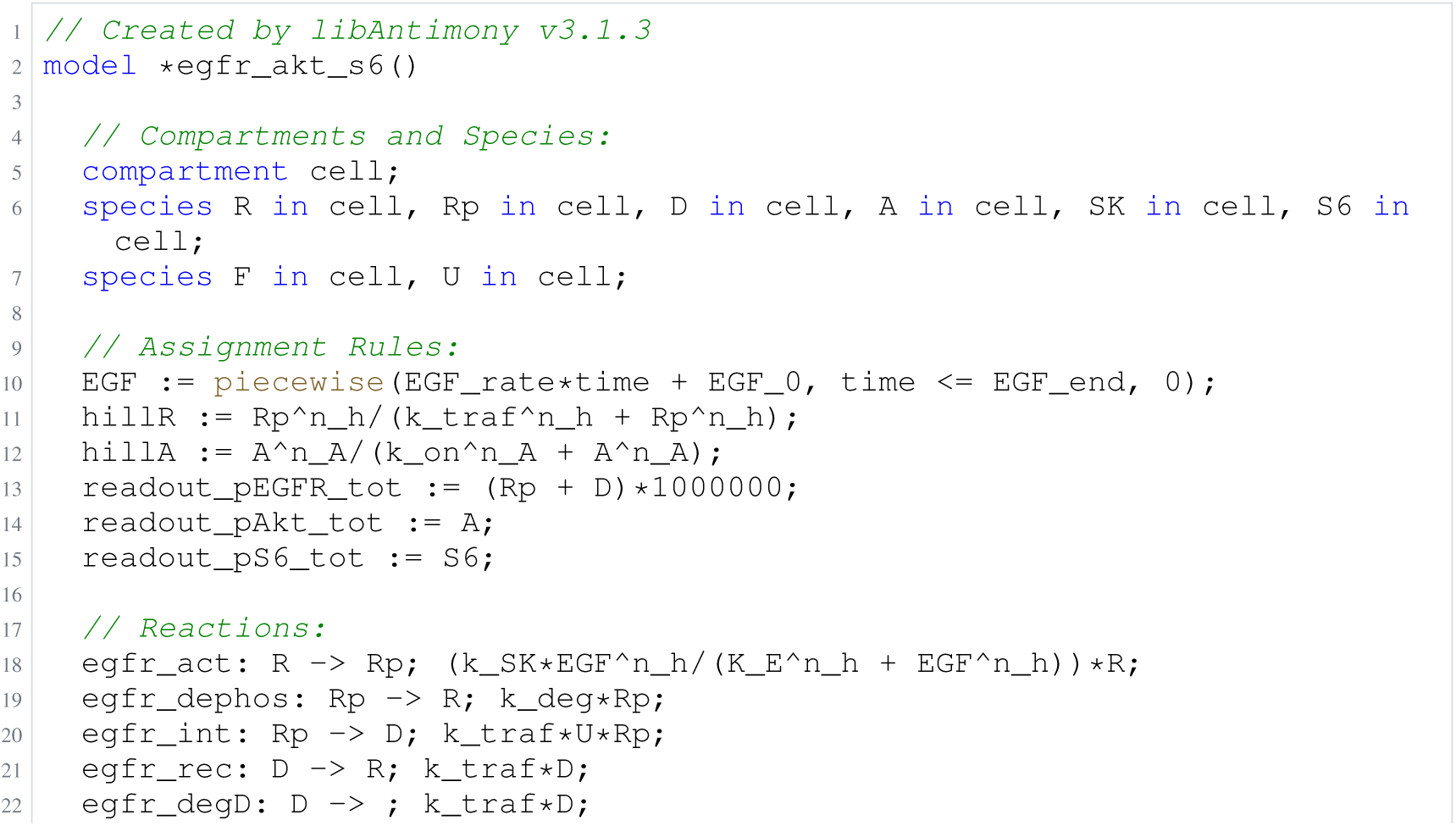
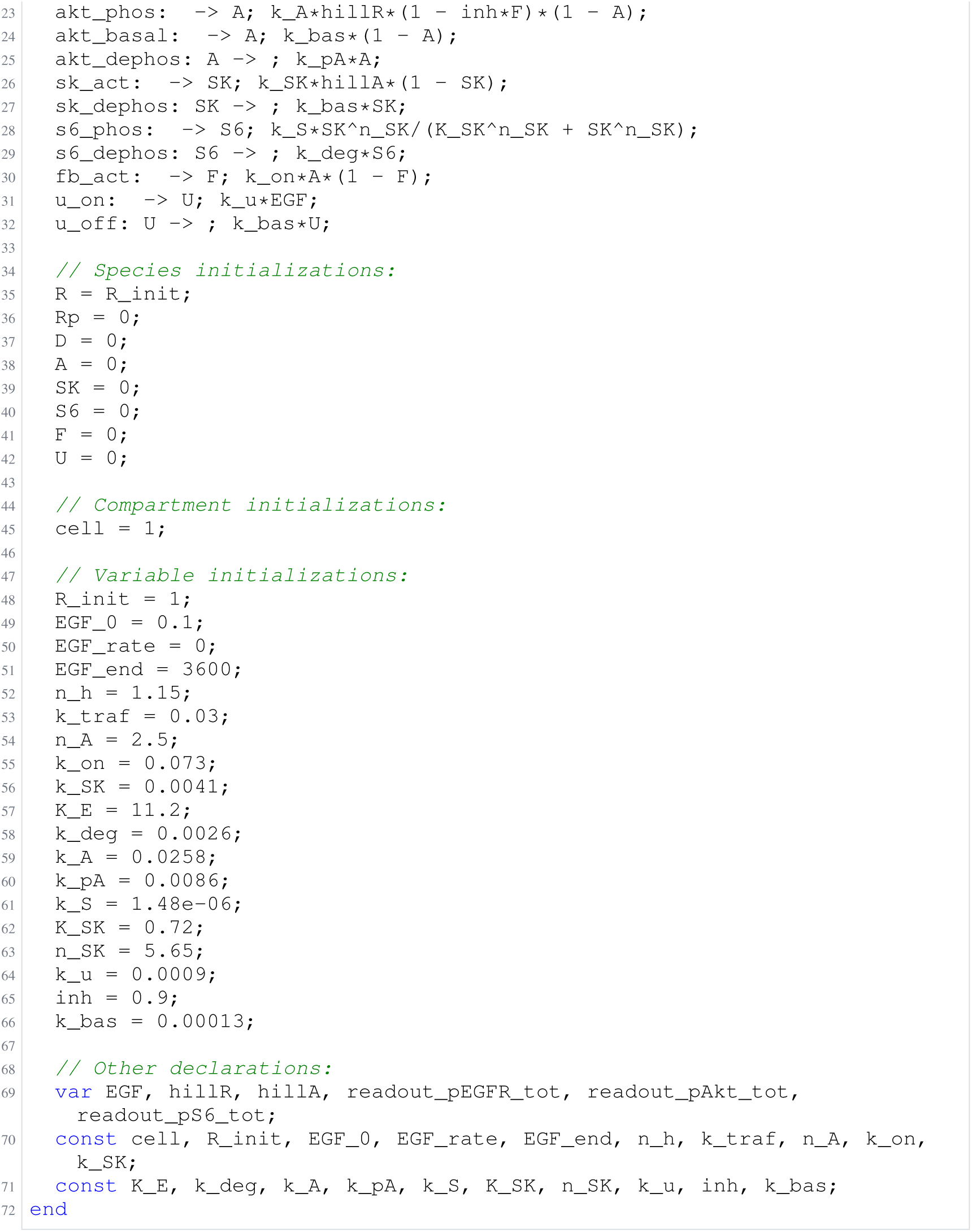
Archived Antimony for O1e: Panel, glm-5.3-flash; task 56476116_4; selected version 20.

**Listing 6:**
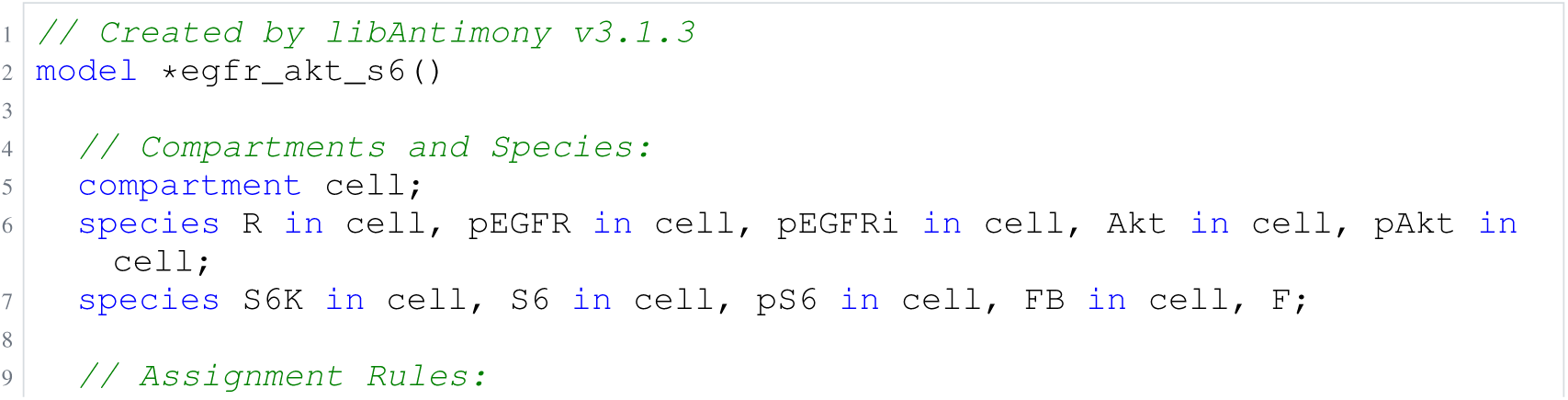
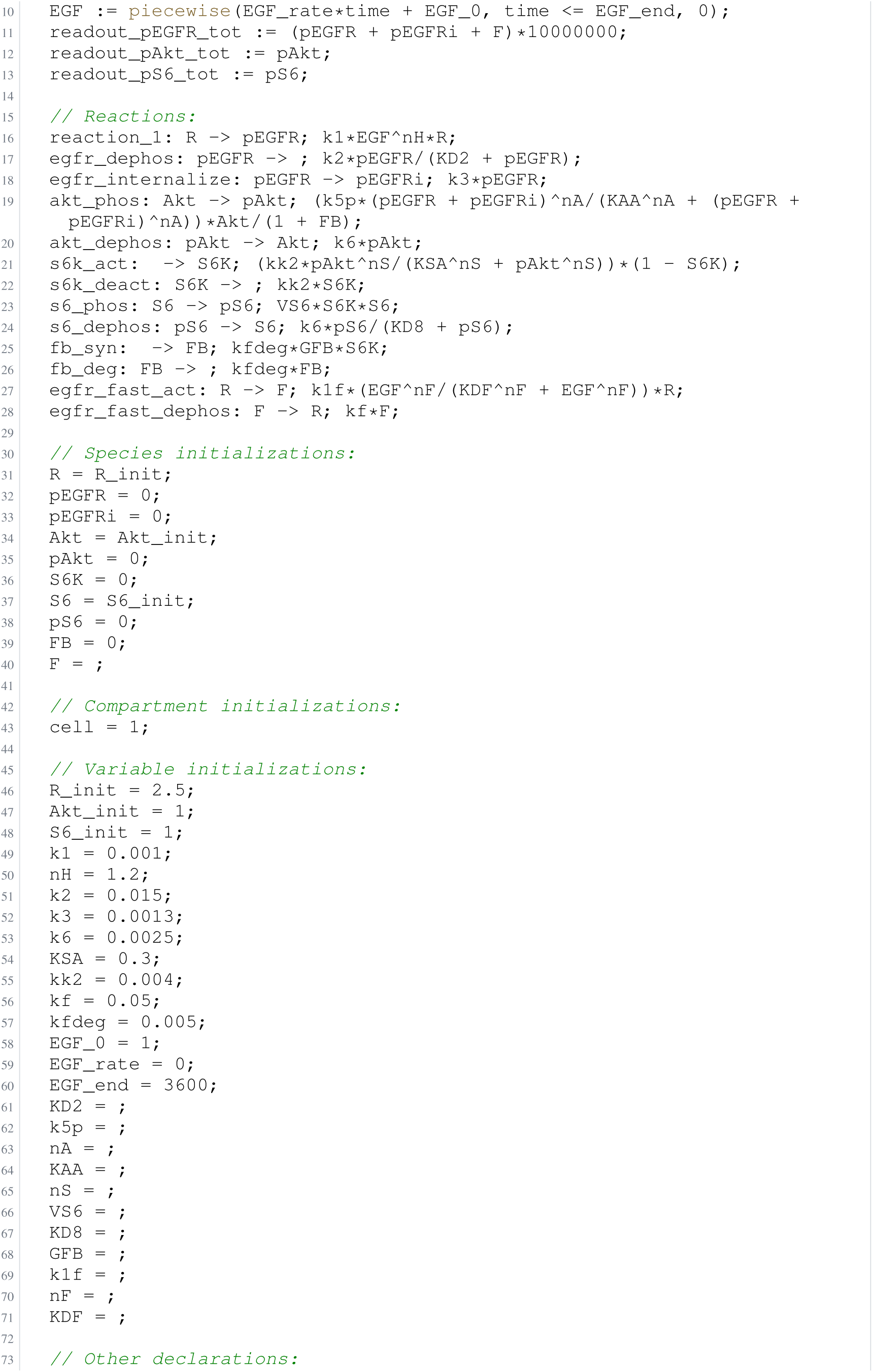
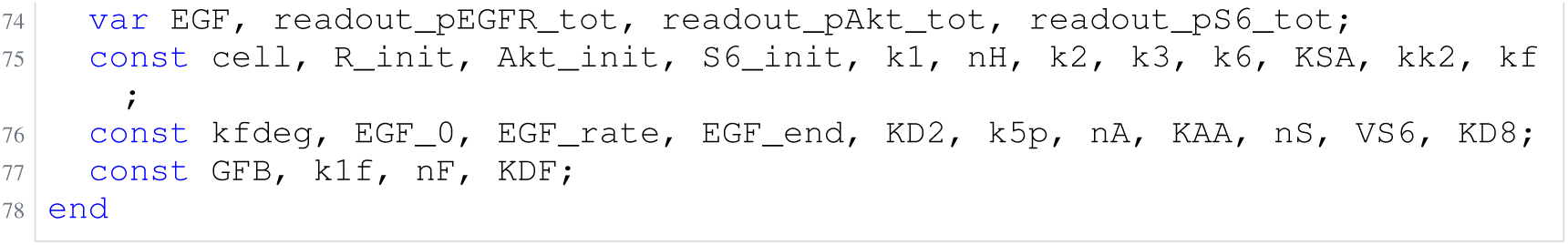
Archived Antimony for O2a: Biocurator off; task 56476121_0; selected version 25.

**Listing 7:**
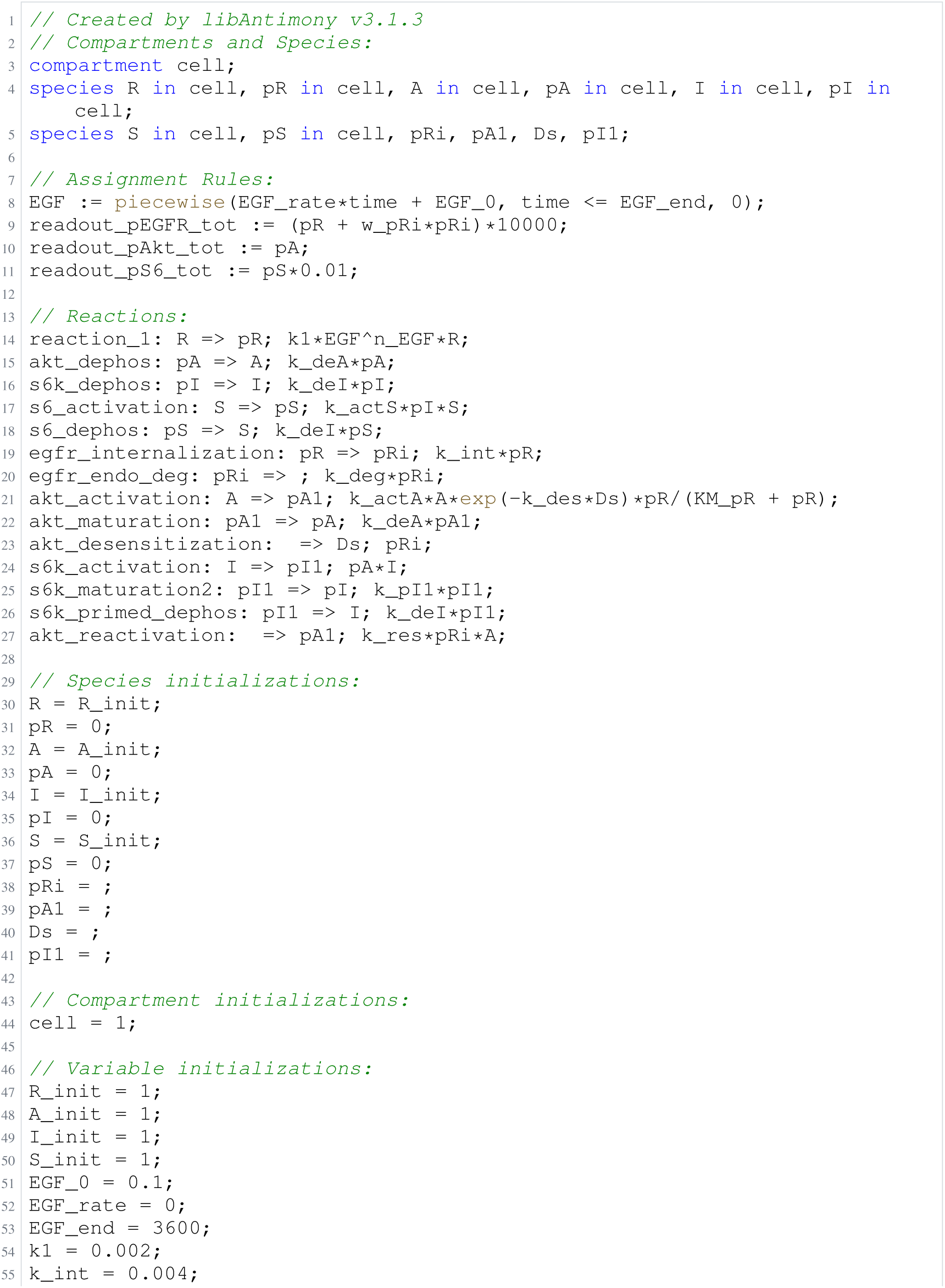
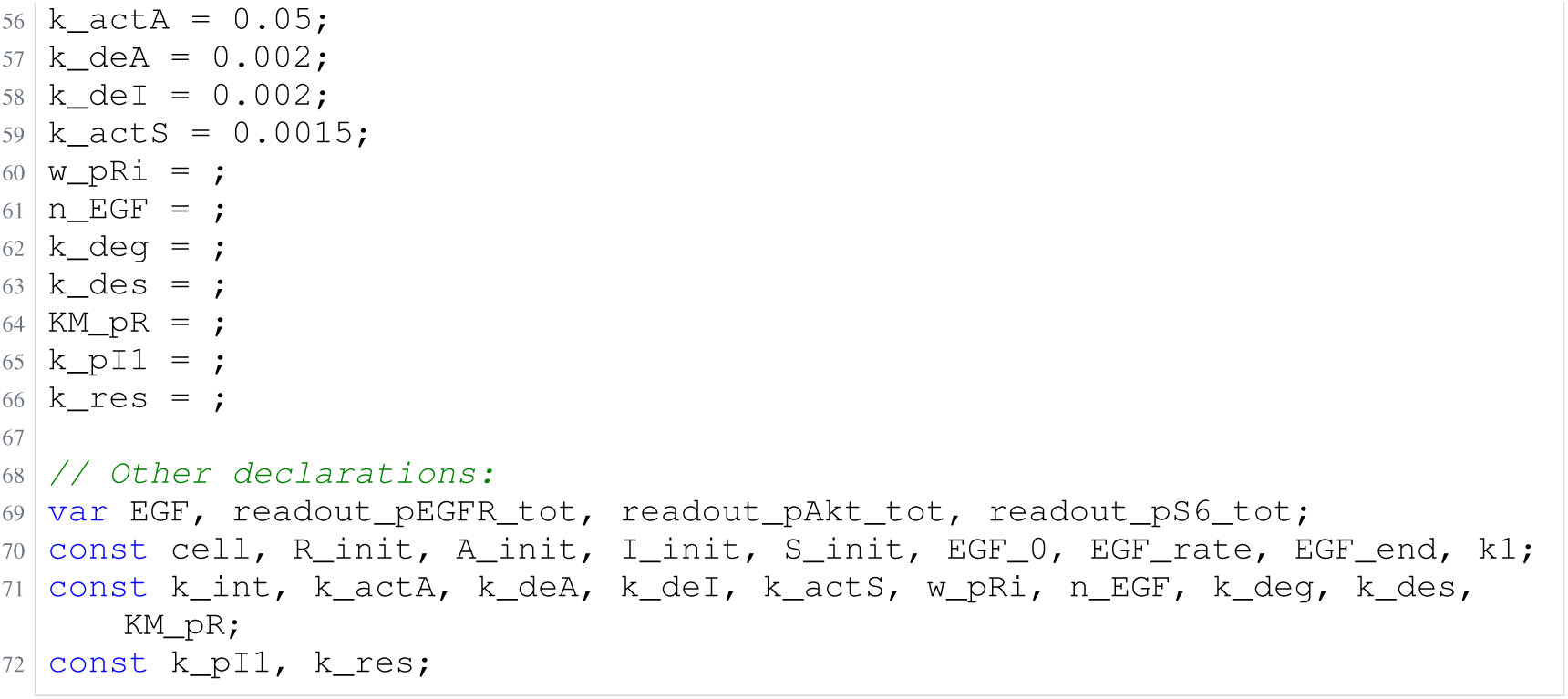
Archived Antimony for O2b: Biocurator off; task 56476121_1; selected version 23.

**Listing 8:**
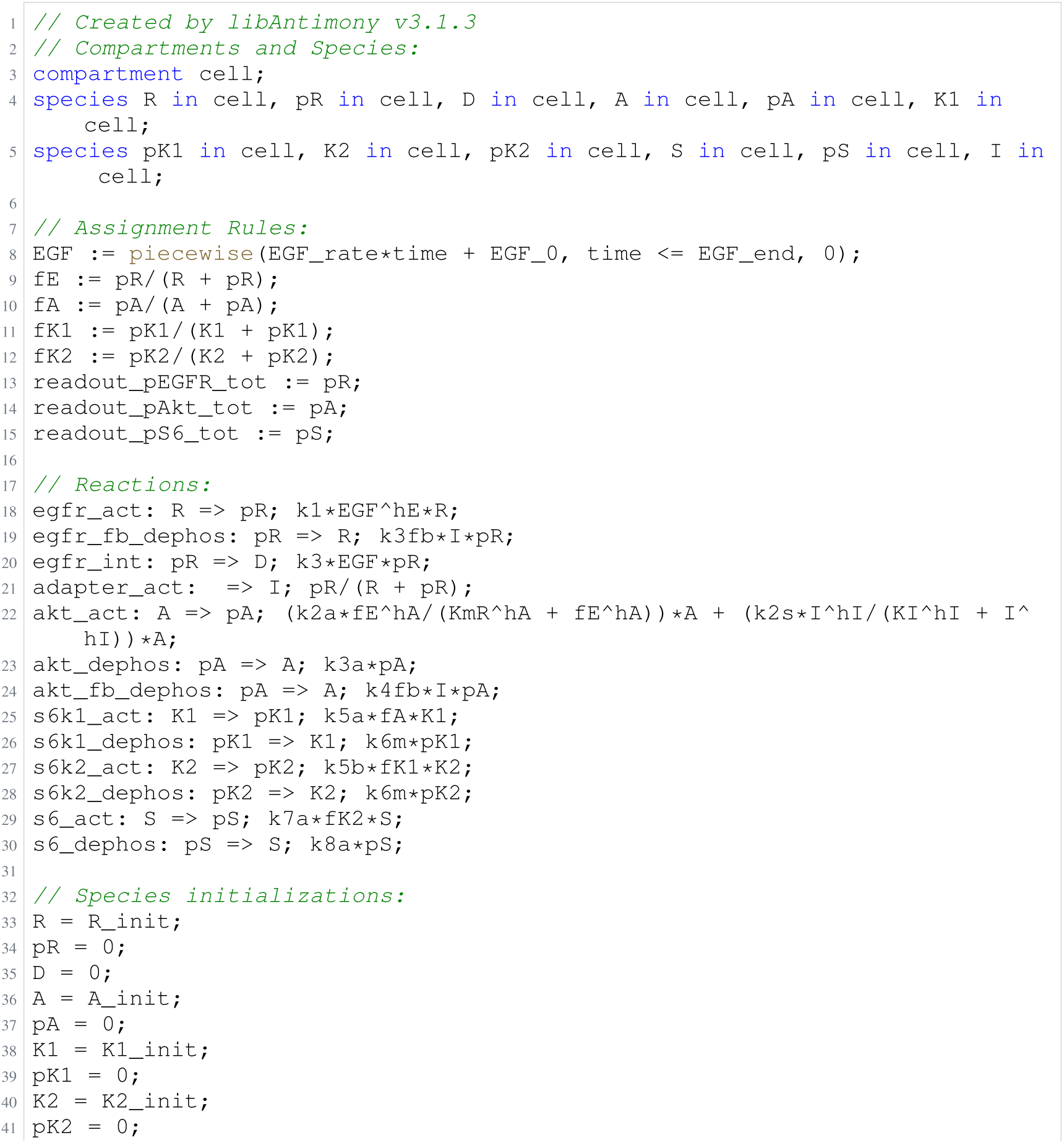
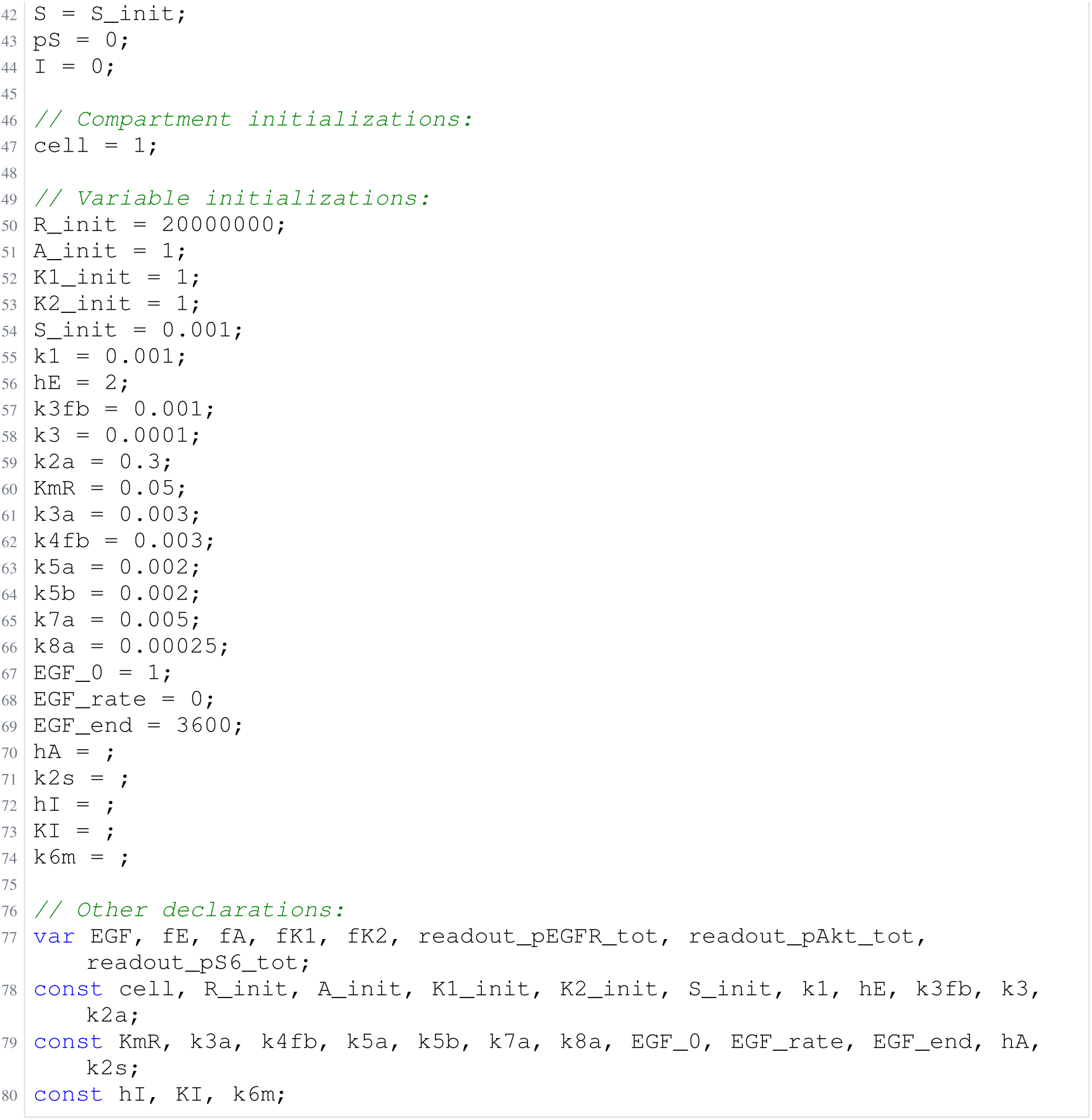
Archived Antimony for O2c: Biocurator off; task 56476121_2; selected version 17.

**Listing 9:**
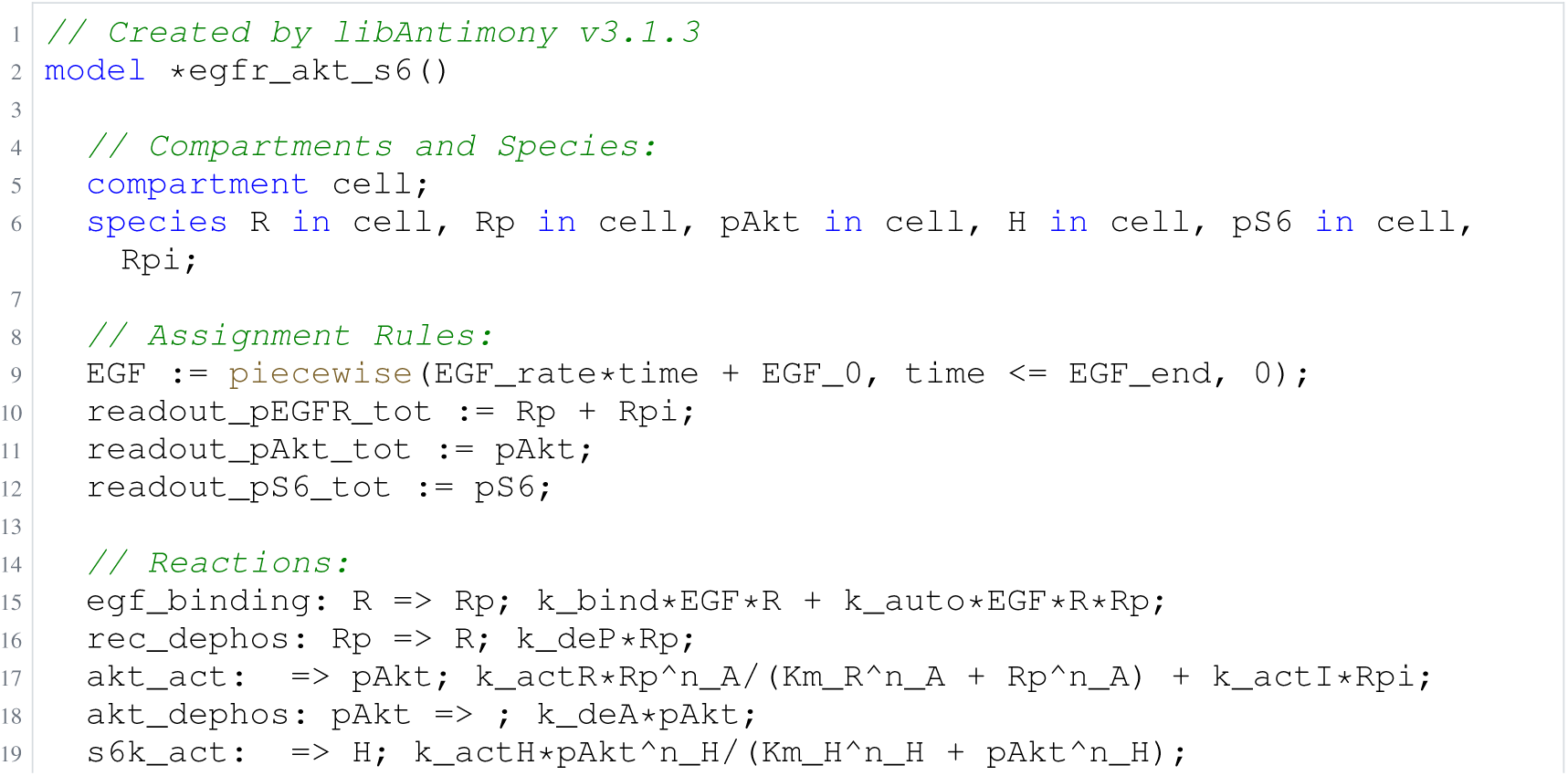
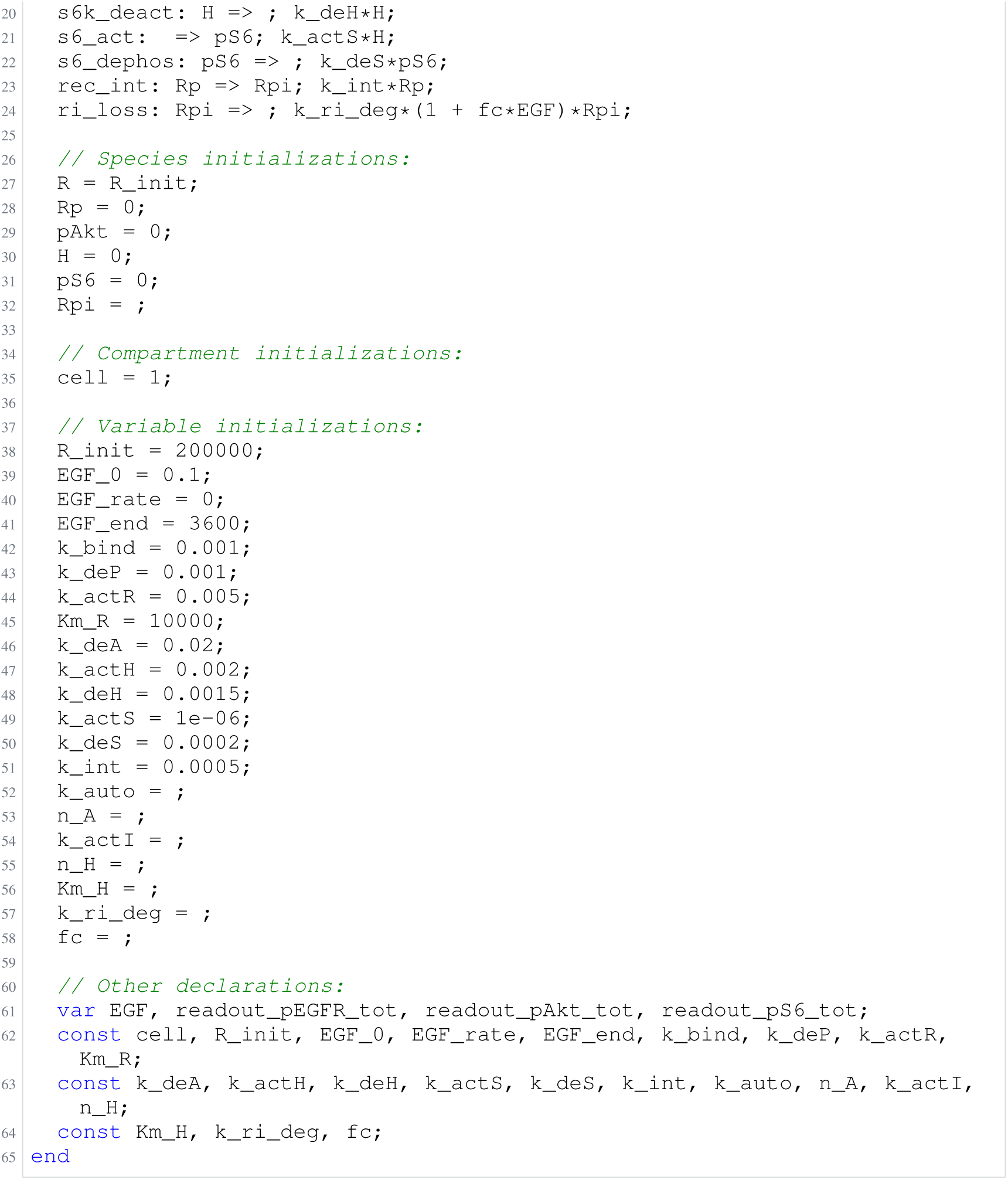
Archived Antimony for O2d: Biocurator off; task 56476121_3; selected version 3.

**Listing 10:**
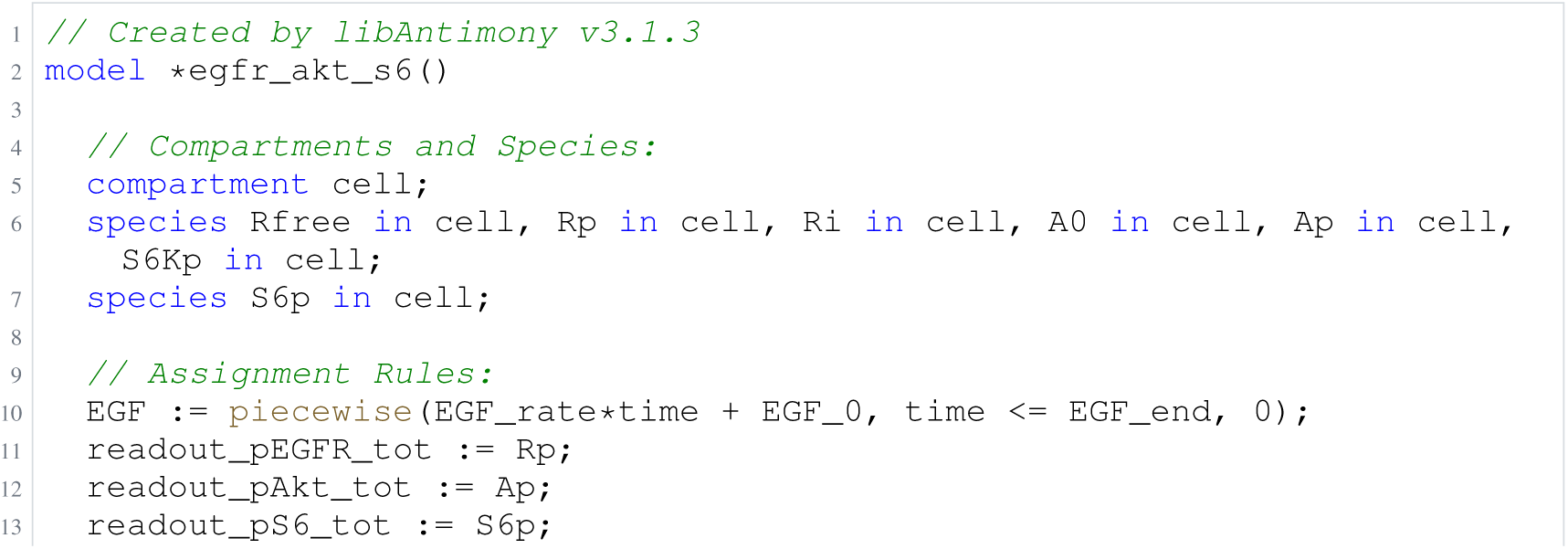
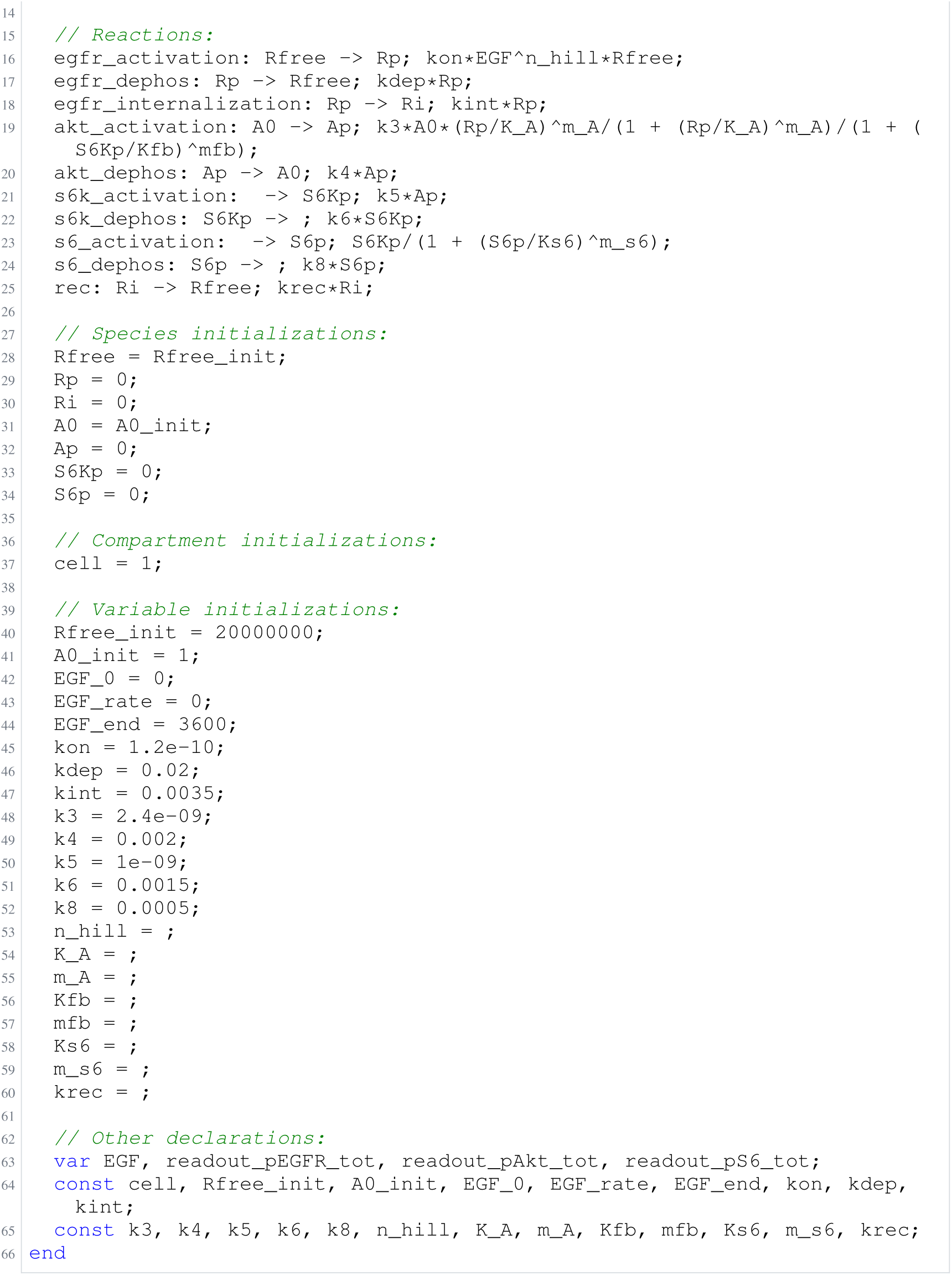
Archived Antimony for O2e: Biocurator off; task 56476121_4; selected version 25.

**Listing 11:**
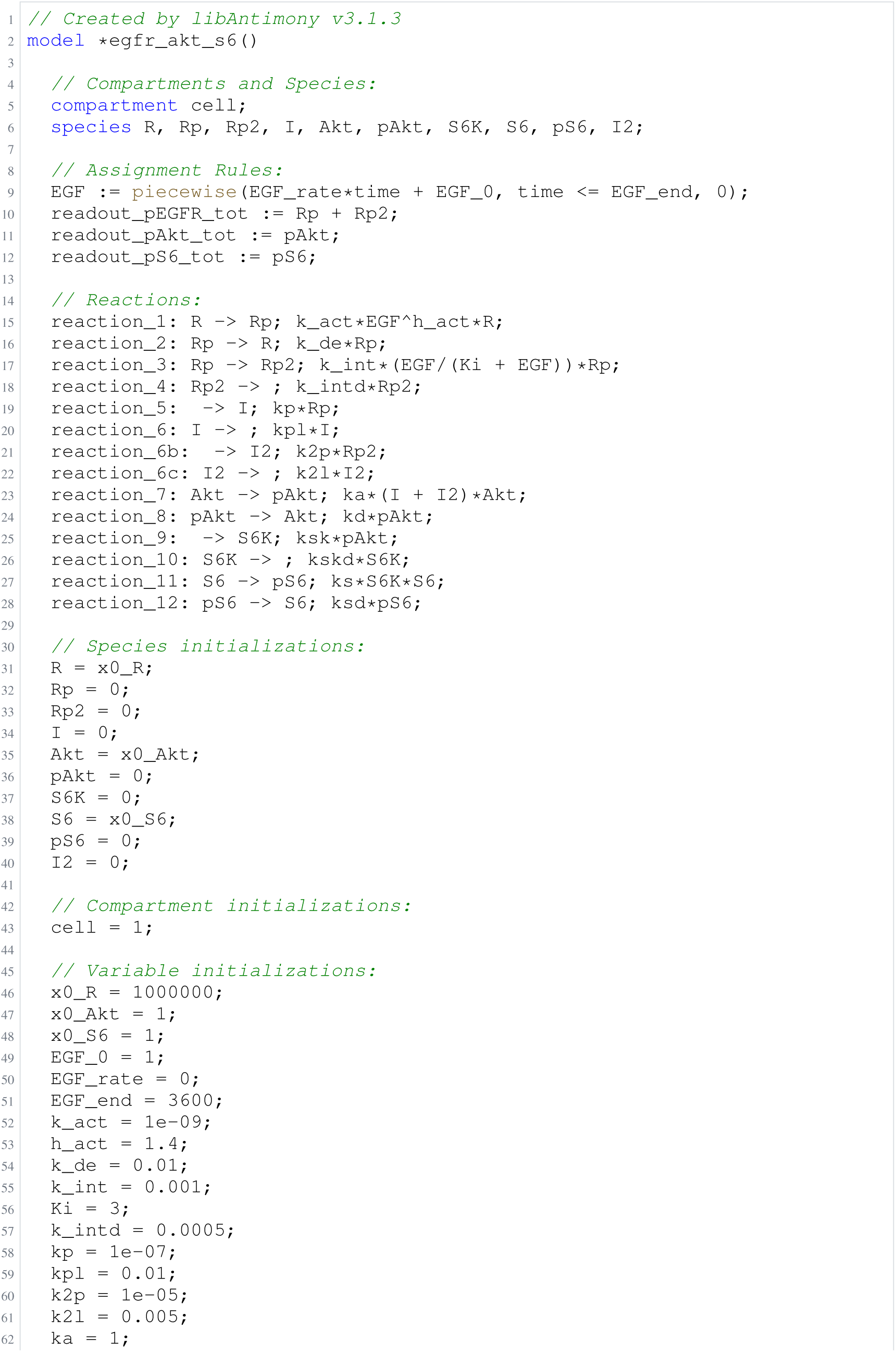
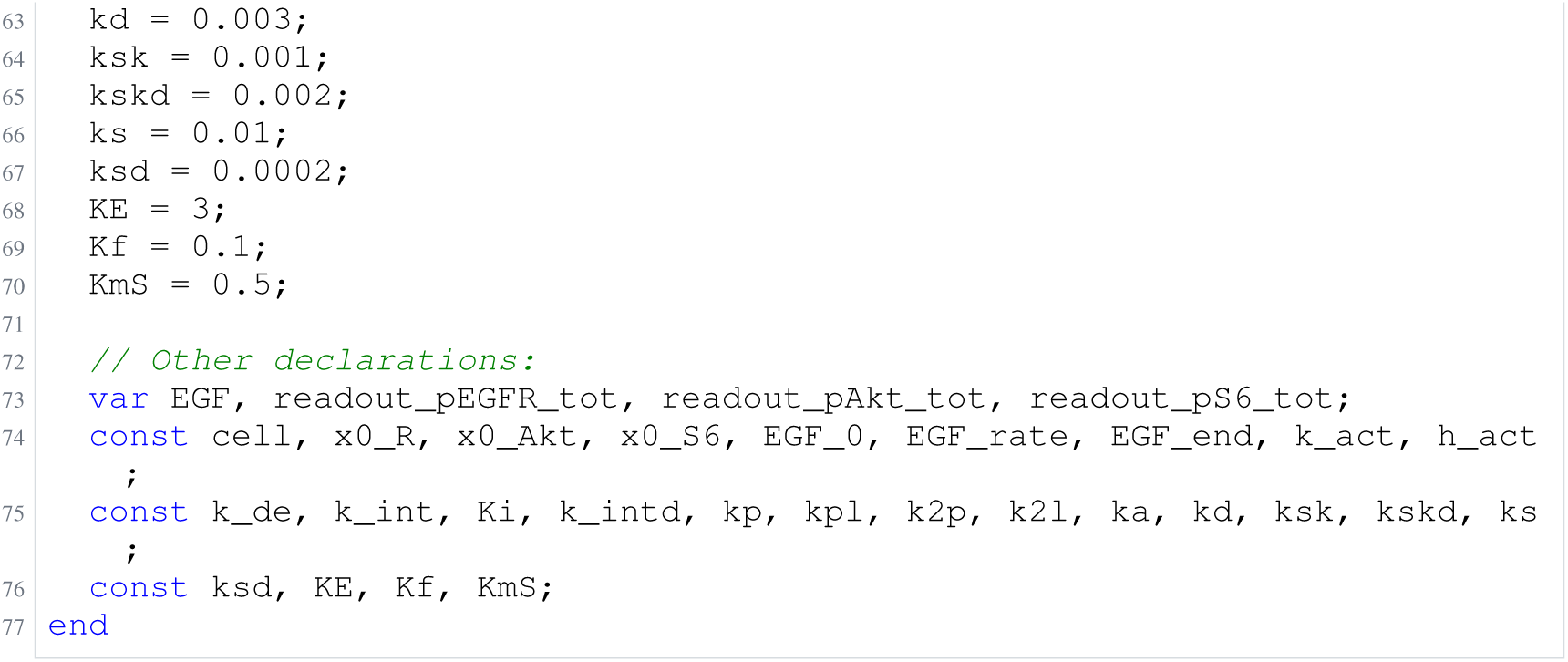
Archived Antimony for O3b: Null arm, glm-5.3-flash; task 56550502_1; delivered problem, no version store; Antimony converted from the archived SBML.

**Listing 12:**
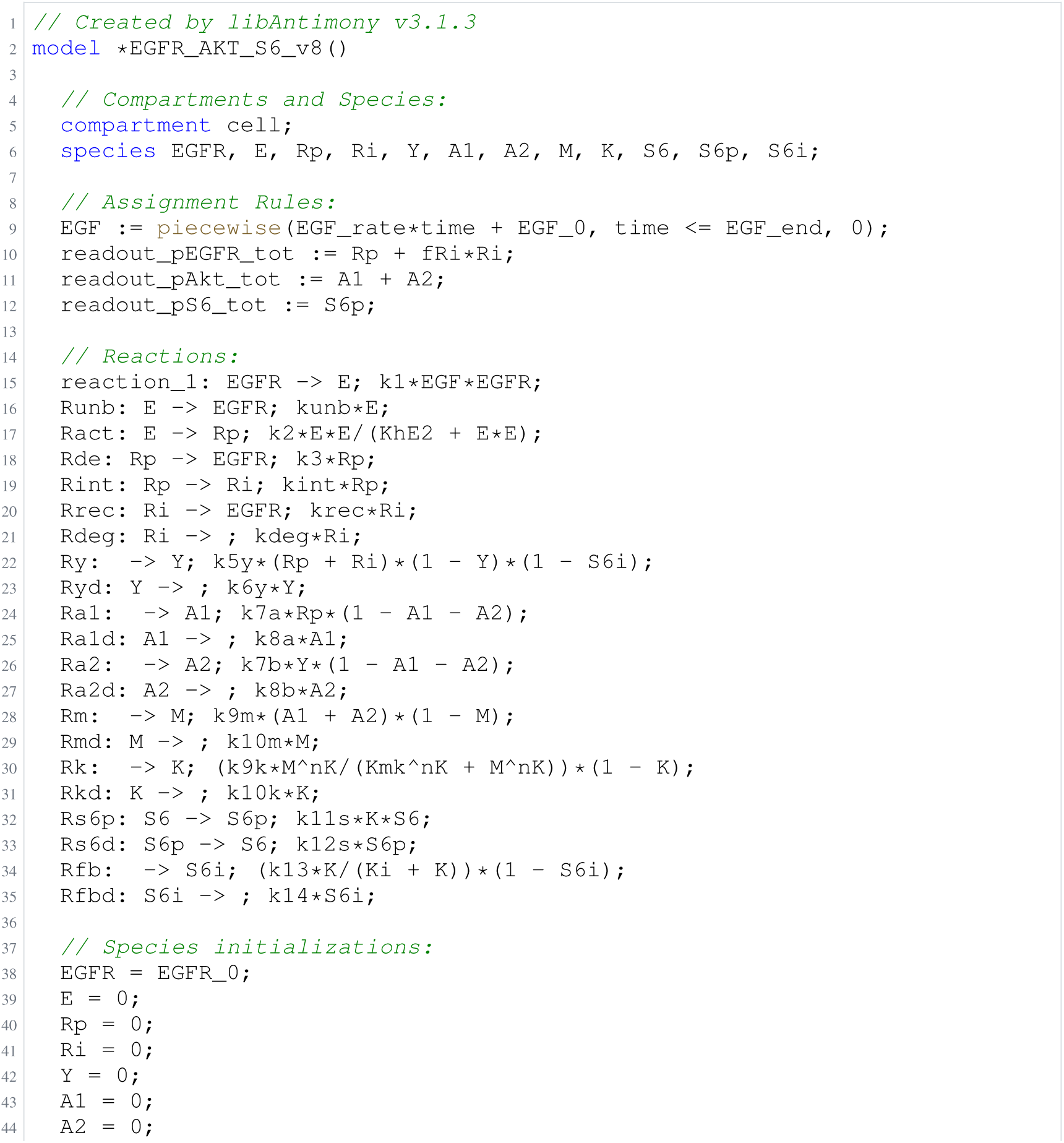
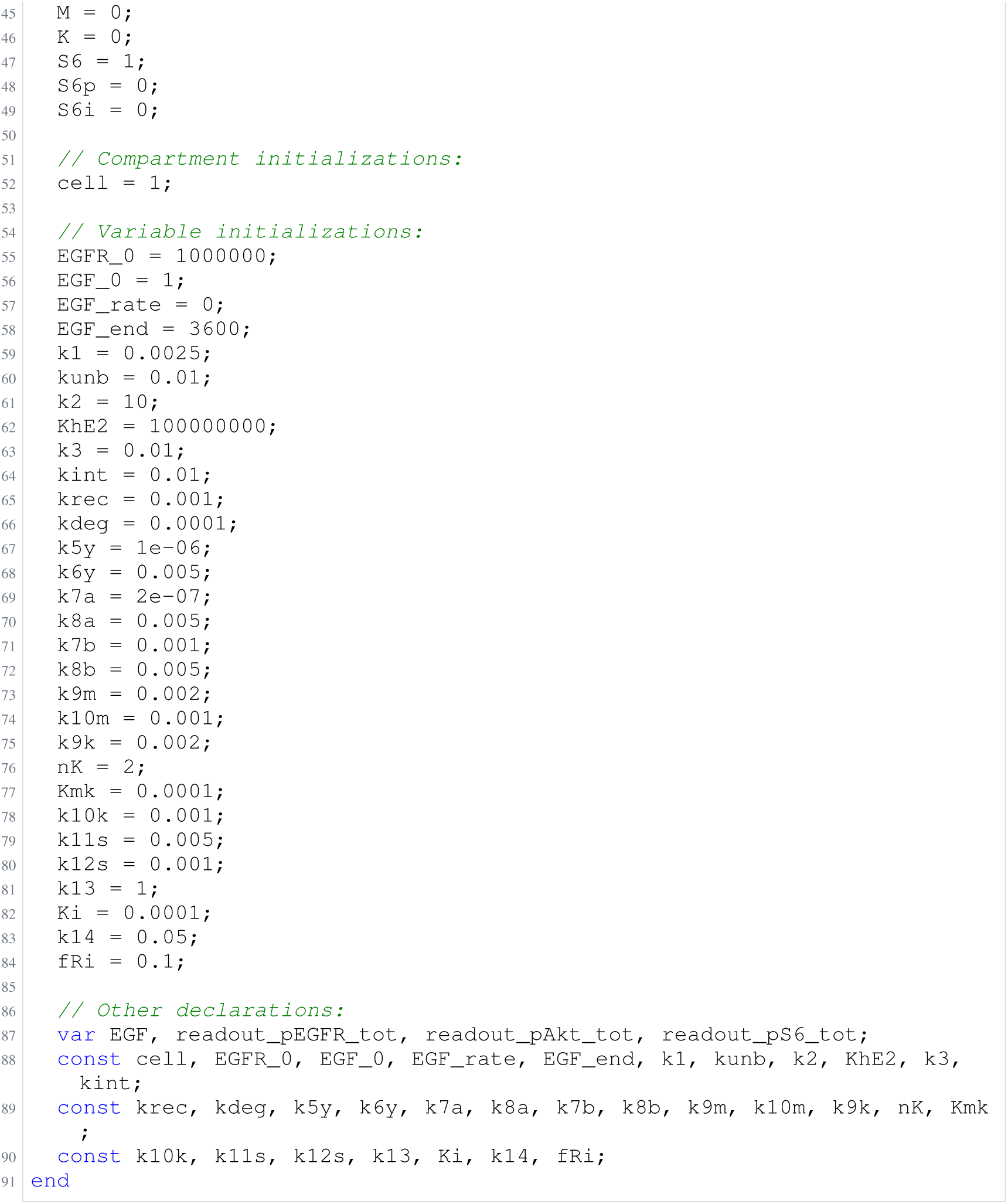
Archived Antimony for O3c: Null arm, glm-5.3-flash; task 56550502_2; delivered problem, no version store; Antimony converted from the archived SBML.

**Listing 13:**
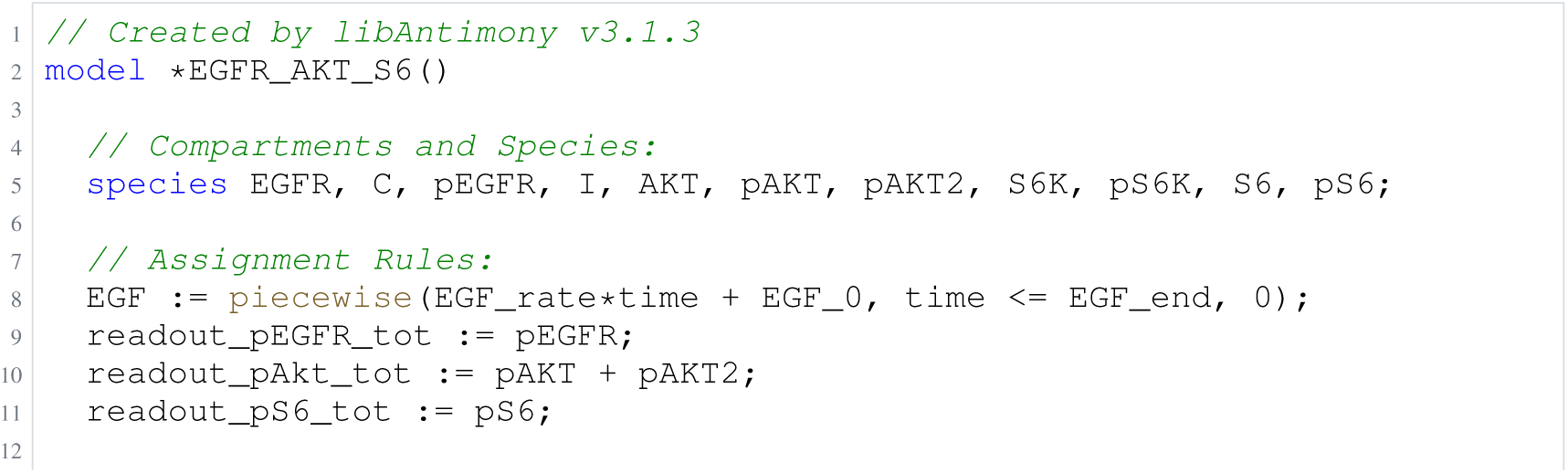
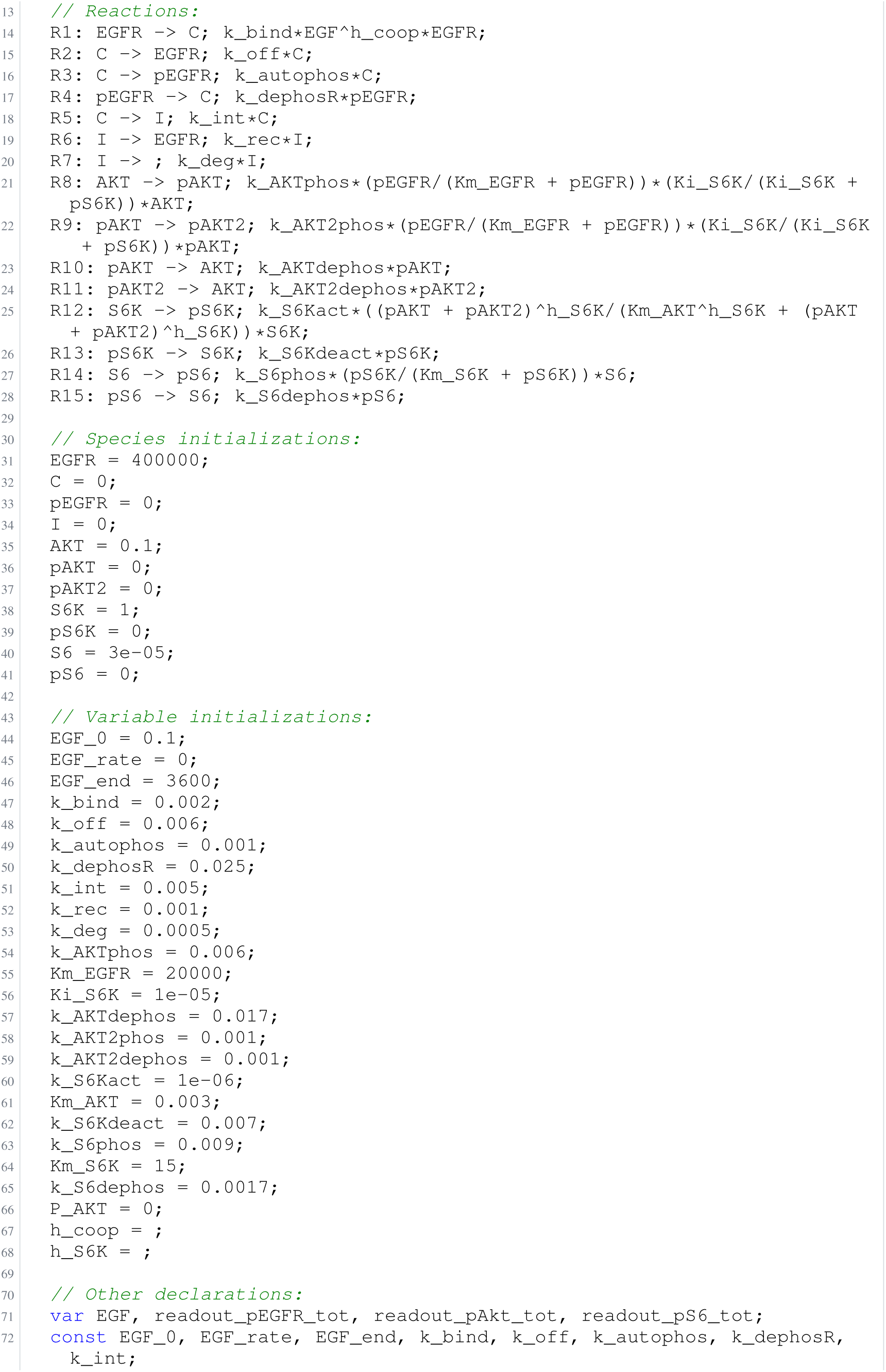
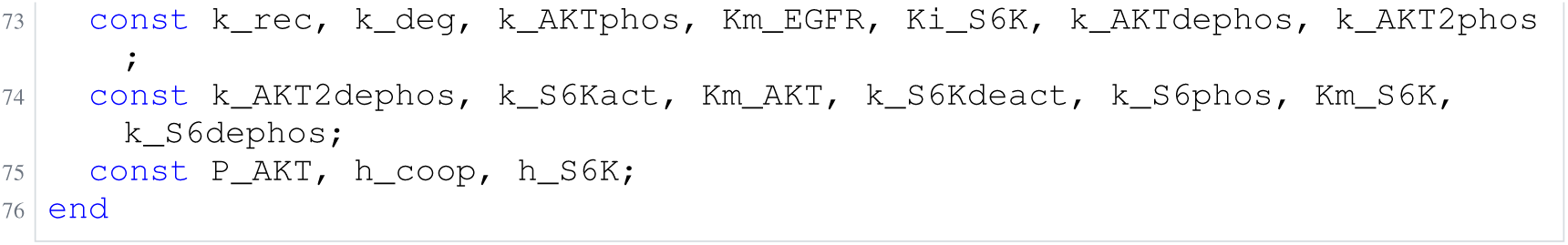
Archived Antimony for O3d: Null arm, glm-5.3-flash; task 56550502_3; delivered problem, no version store; Antimony converted from the archived SBML.

**Listing 14:**
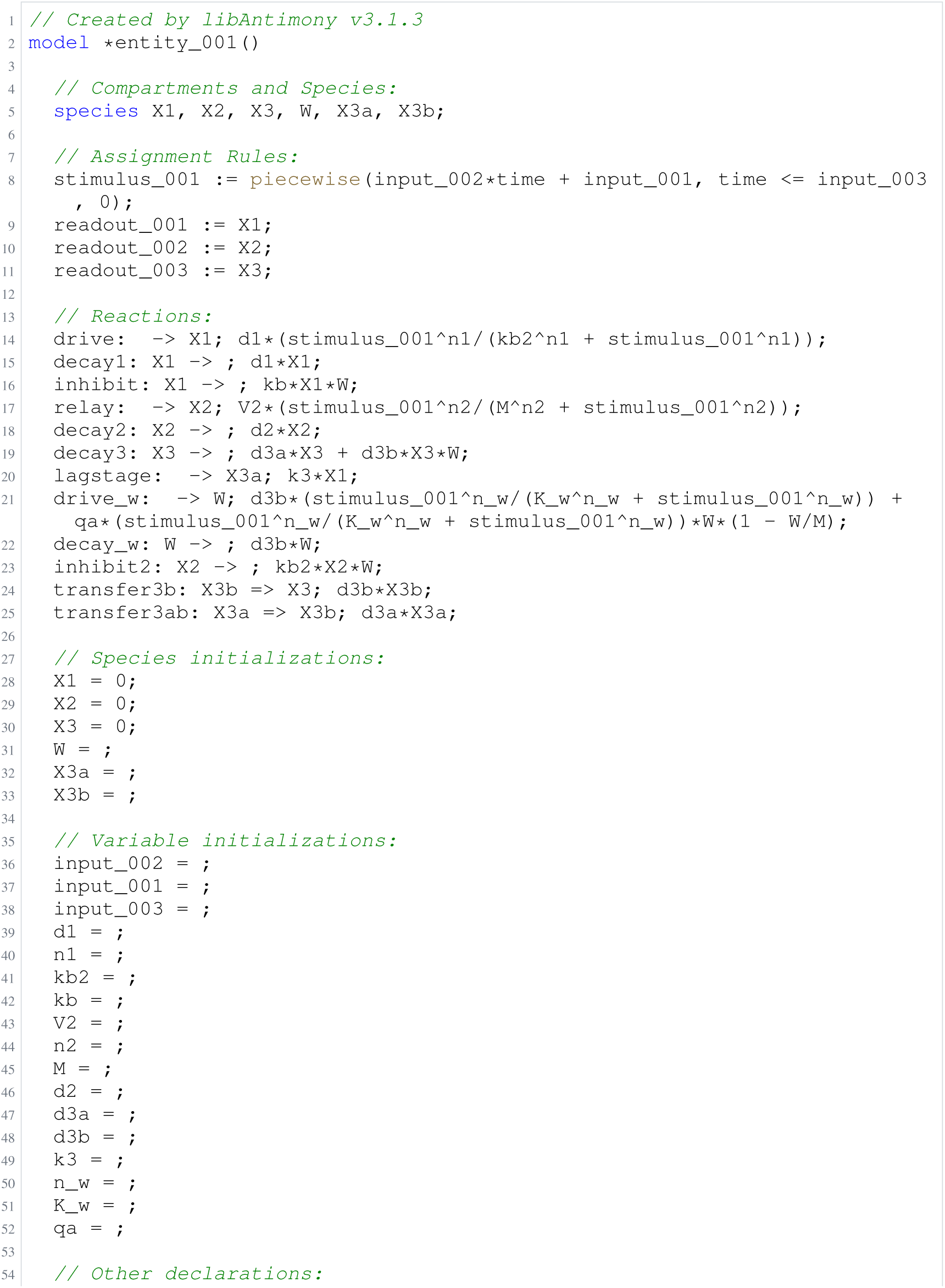
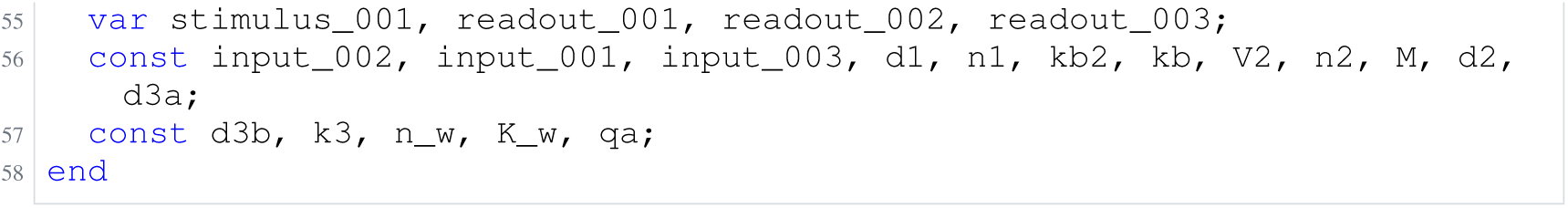
Archived Antimony for O4a: Biology blinded; task 58644985_0; selected version 15.

**Listing 15:**
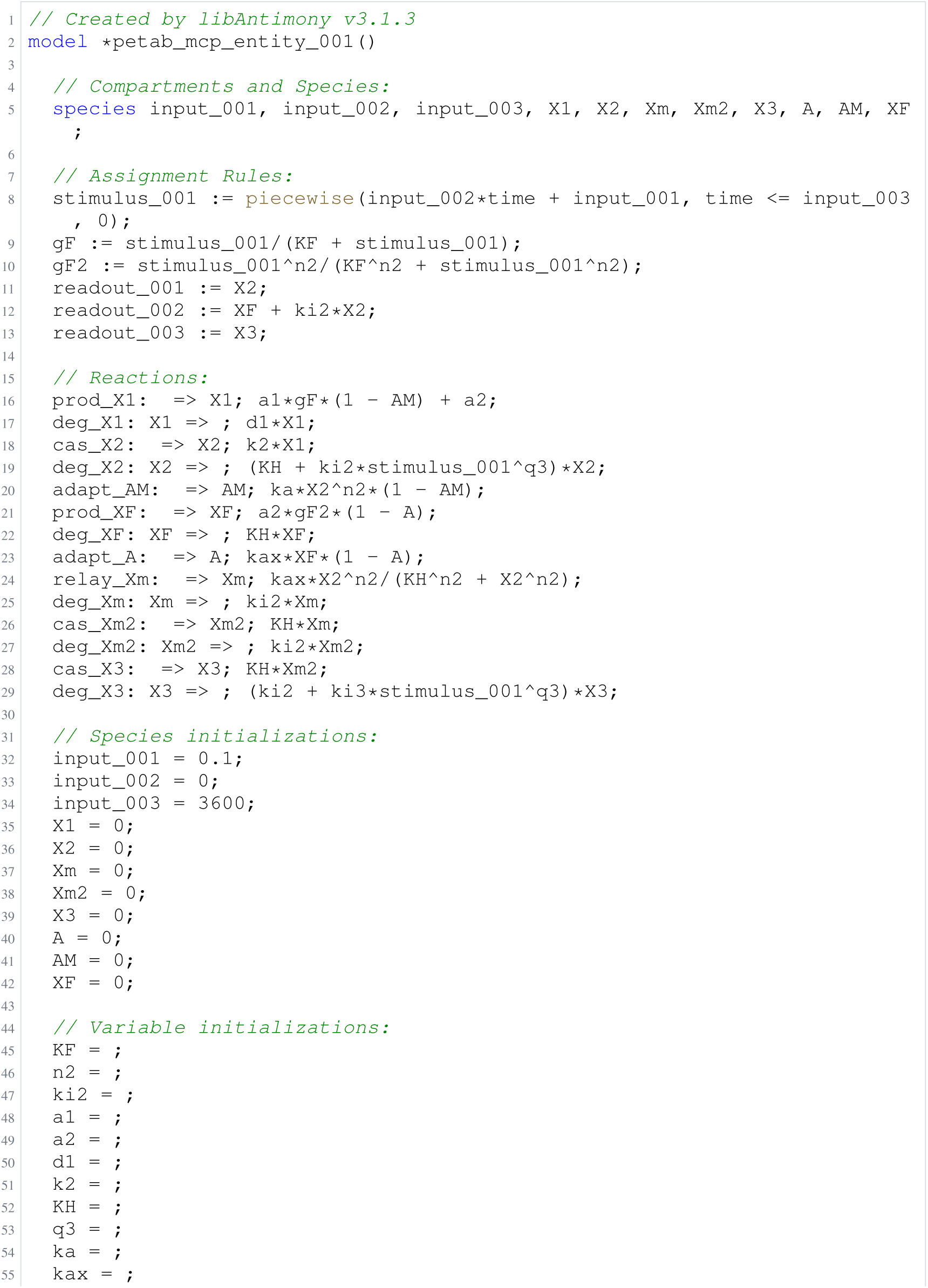
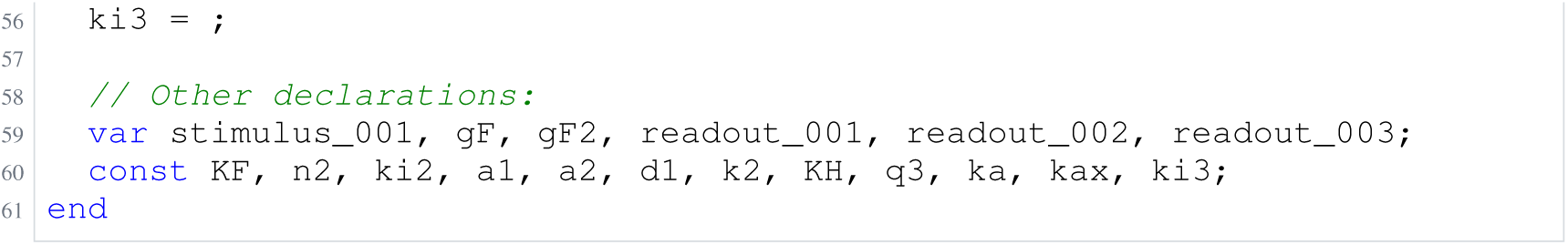
Archived Antimony for O4b: Biology blinded; task 58644985_1; selected version 28.

**Listing 16:**
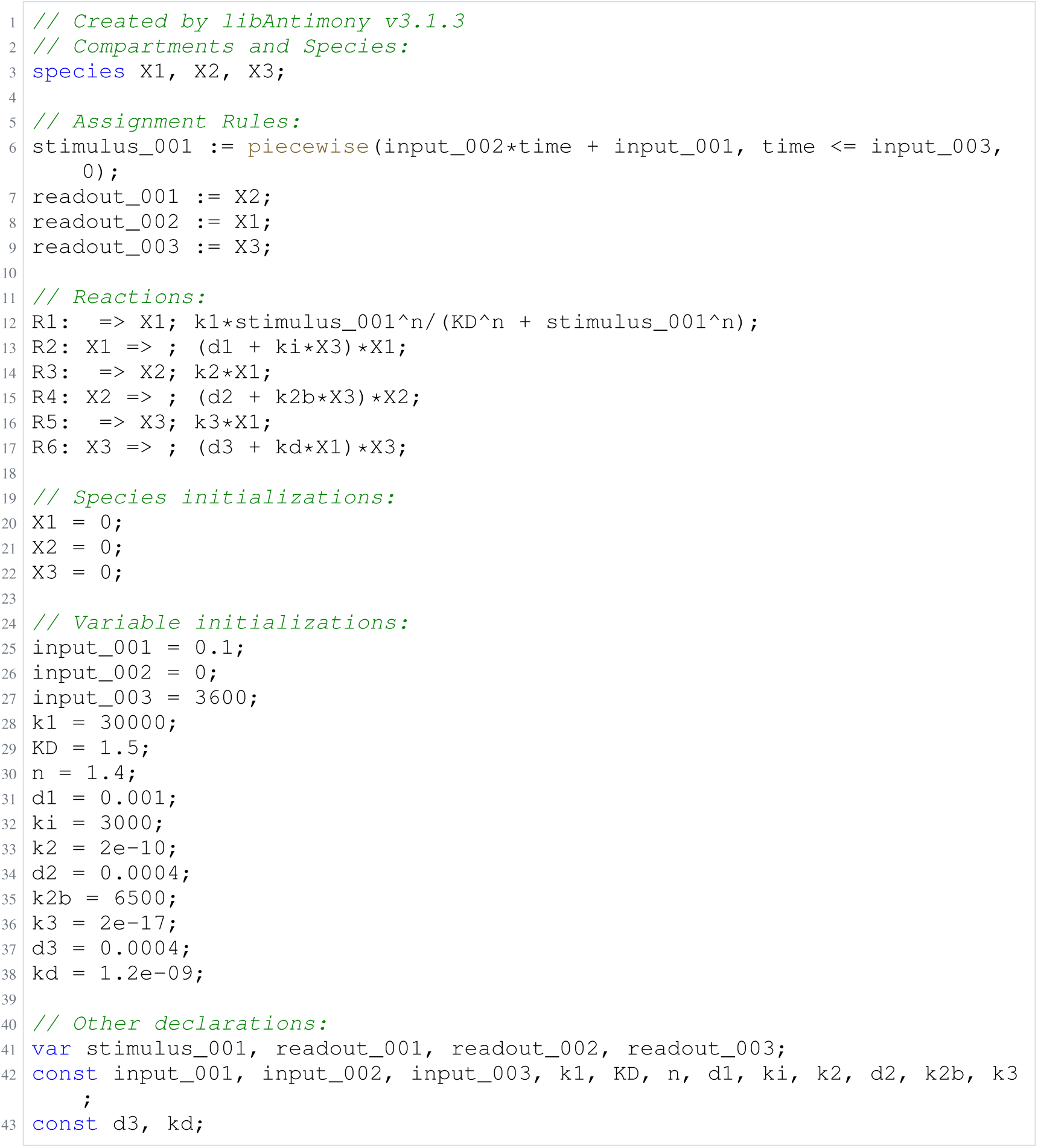
Archived Antimony for O4c: Biology blinded; task 58644985_2; selected version 1.

**Listing 17:**
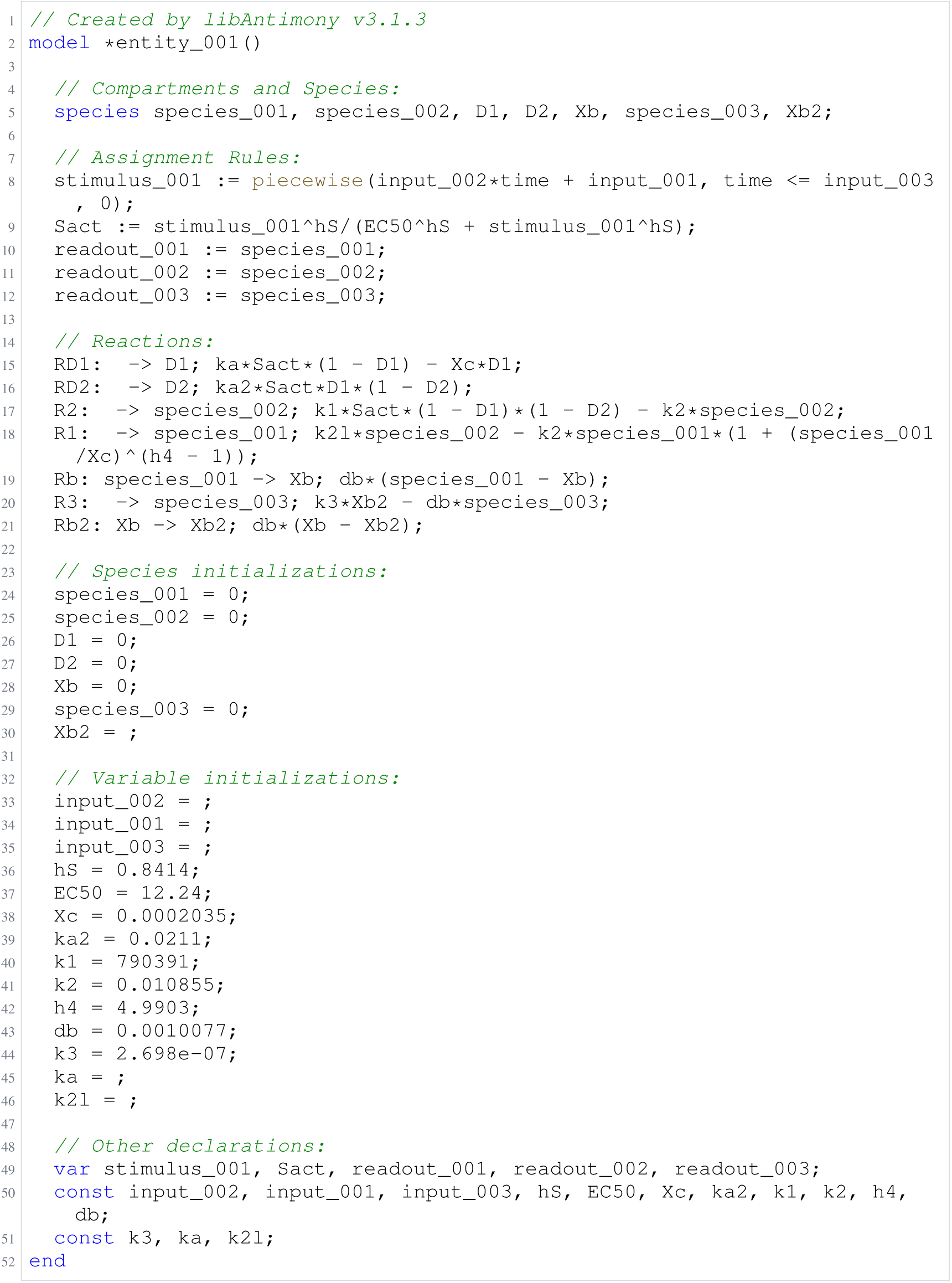
Archived Antimony for O4d: Biology blinded; task 58644985_3; selected version 29.

**Listing 18:**
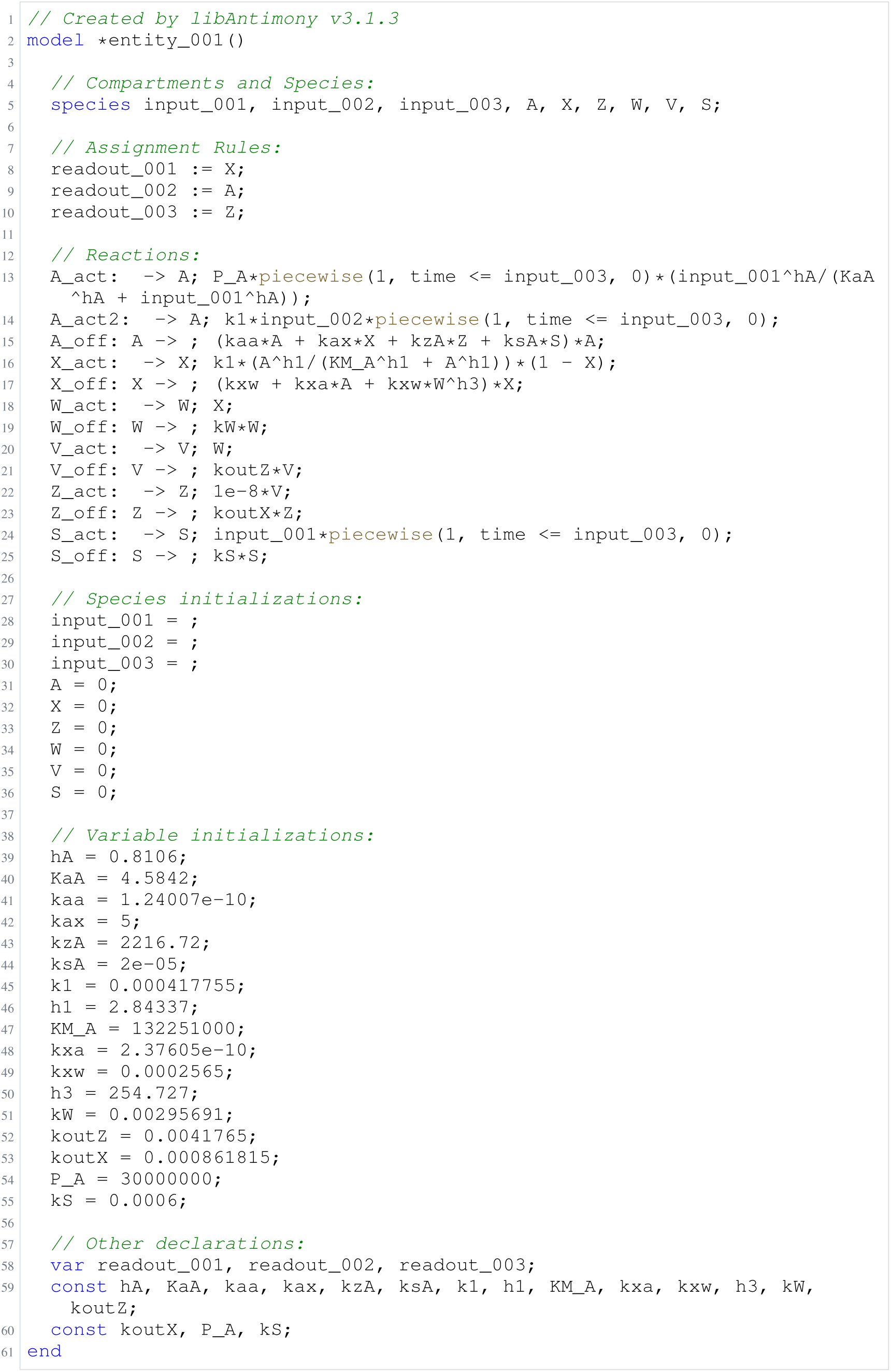
Archived Antimony for O4e: Biology blinded; task 58661494_4; selected version 16.

**Listing 19:**
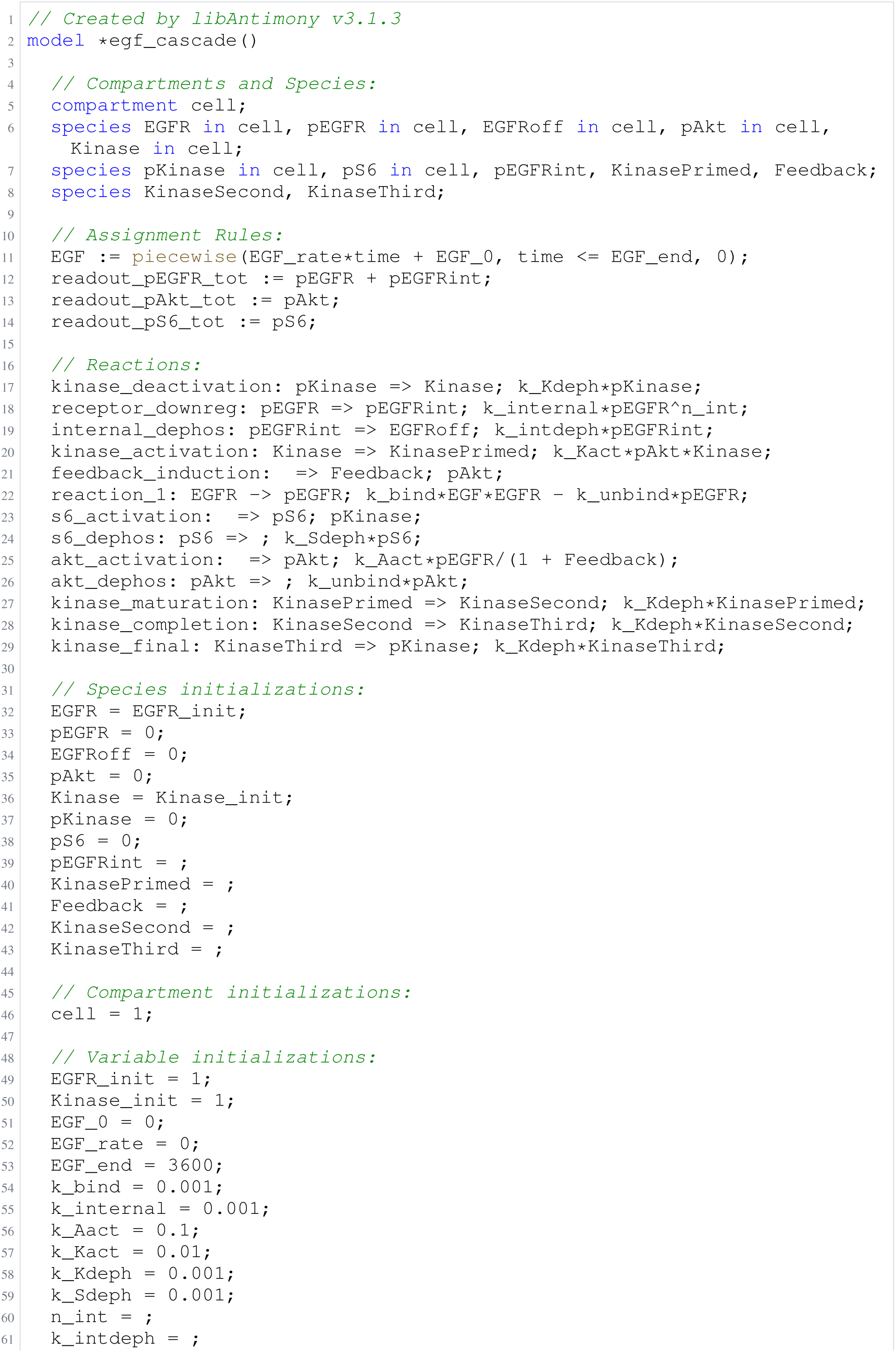
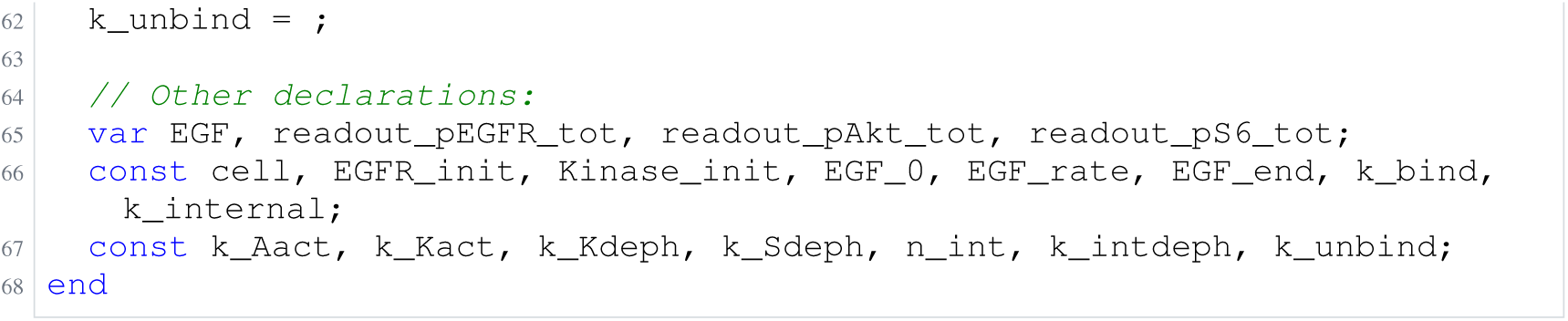
Archived Antimony for F2a: Panel, gpt-6-astra, biocurator off; task 56550507_0; selected version 27.

**Listing 20:**
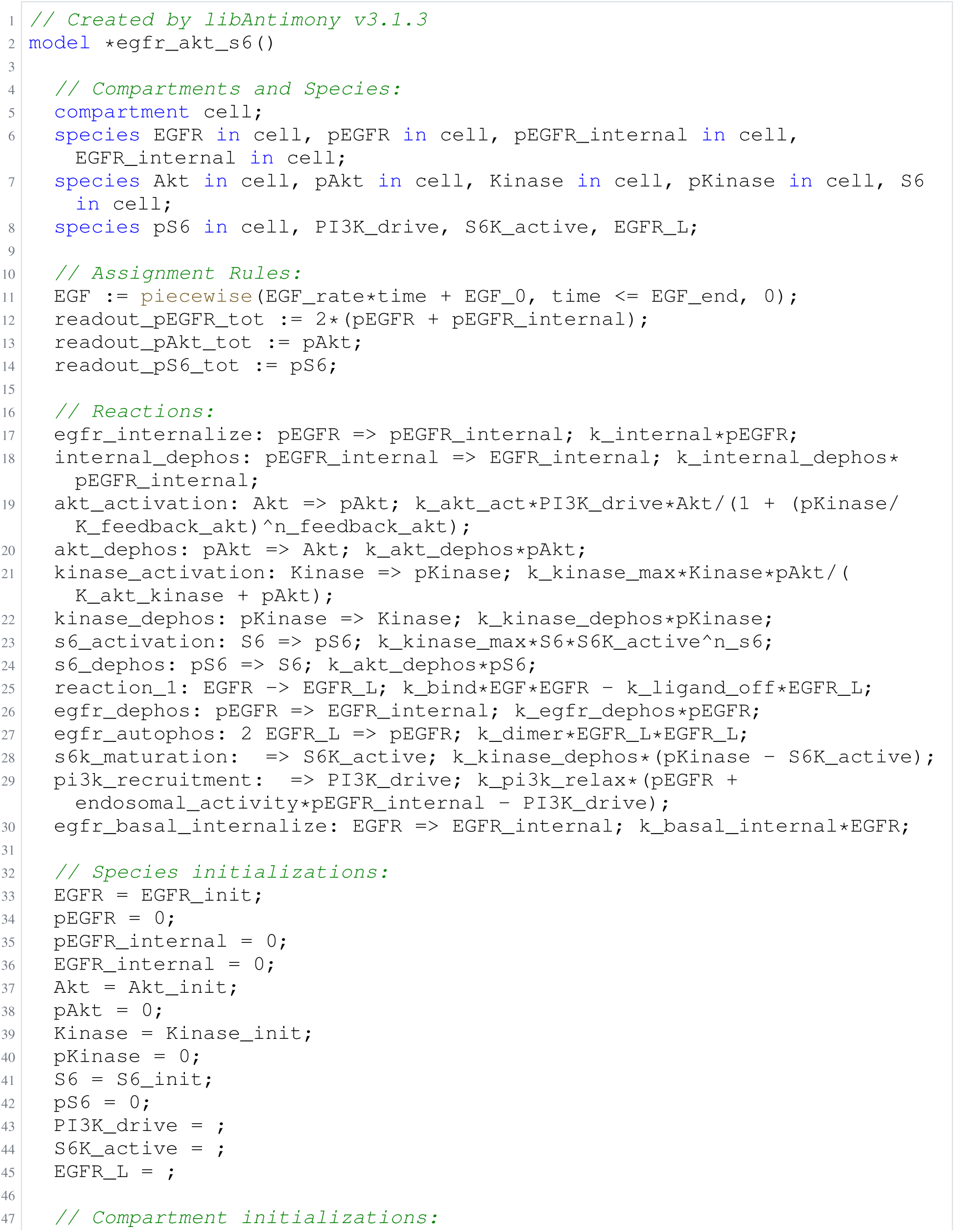
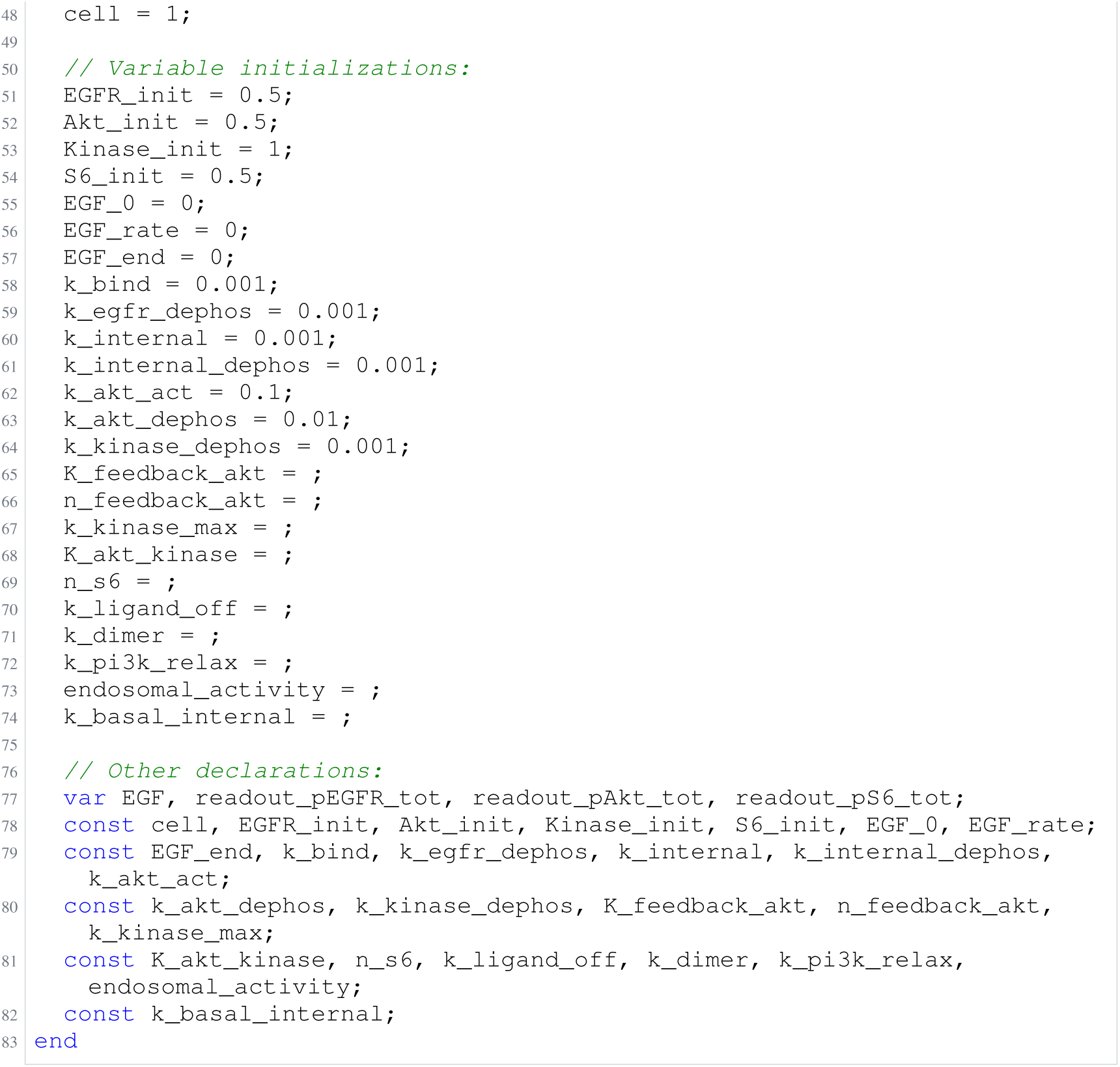
Archived Antimony for F2b: Panel, gpt-6-astra, biocurator off; task 56550507_1; selected version 16.

**Listing 21:**
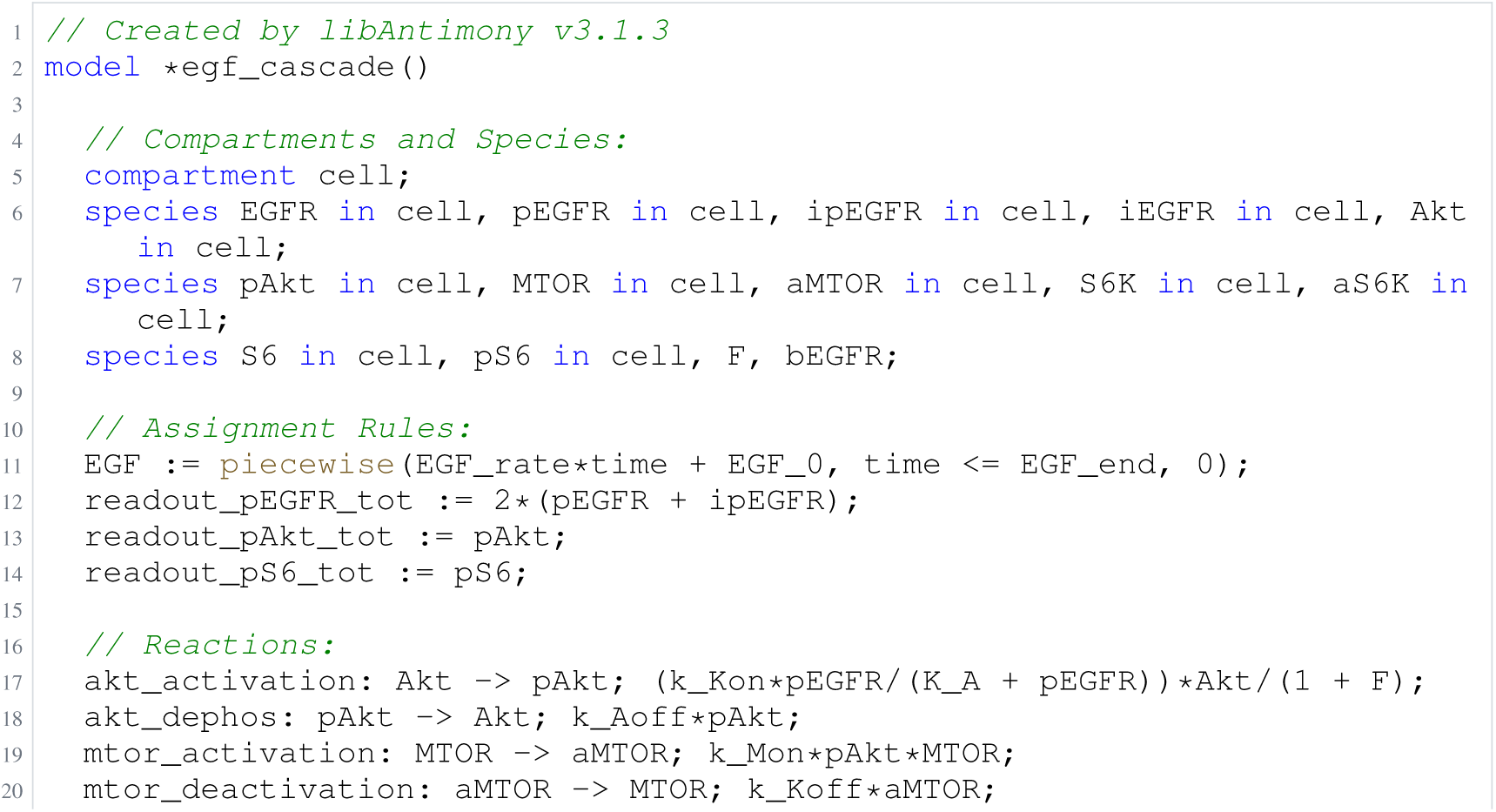
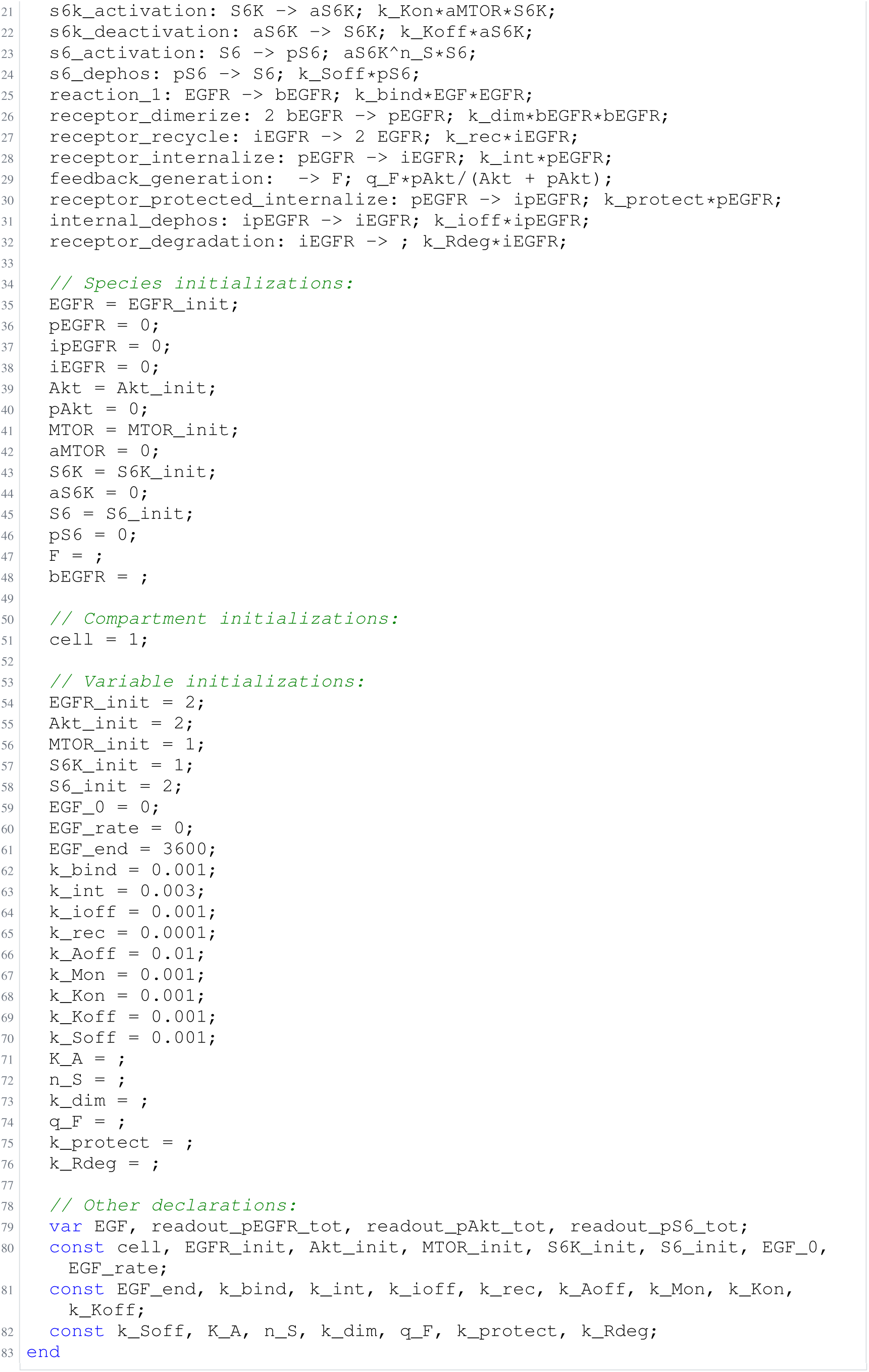
Archived Antimony for F2c: Panel, gpt-6-astra, biocurator off; task 56550507_2; selected version 20.

**Listing 22:**
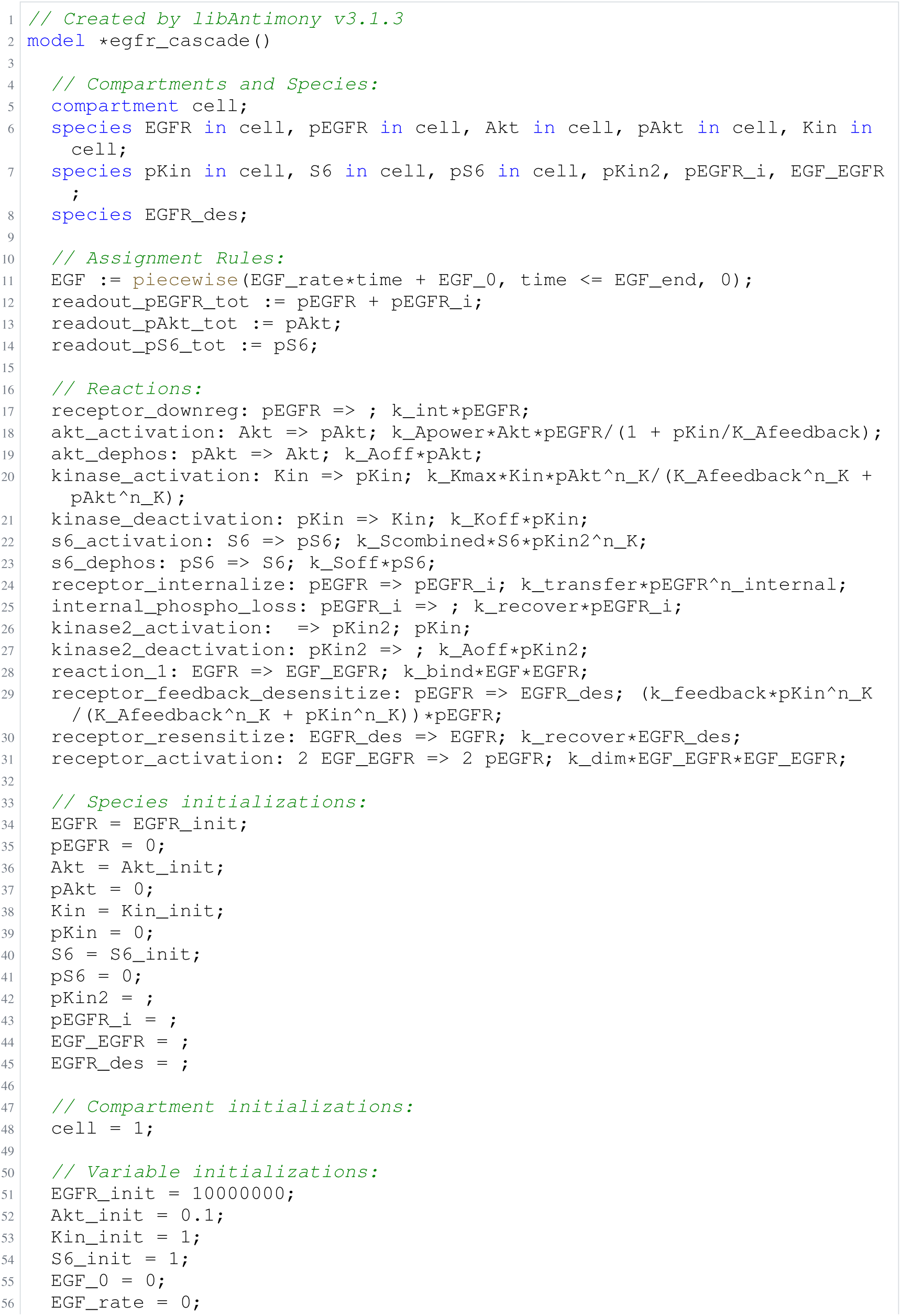
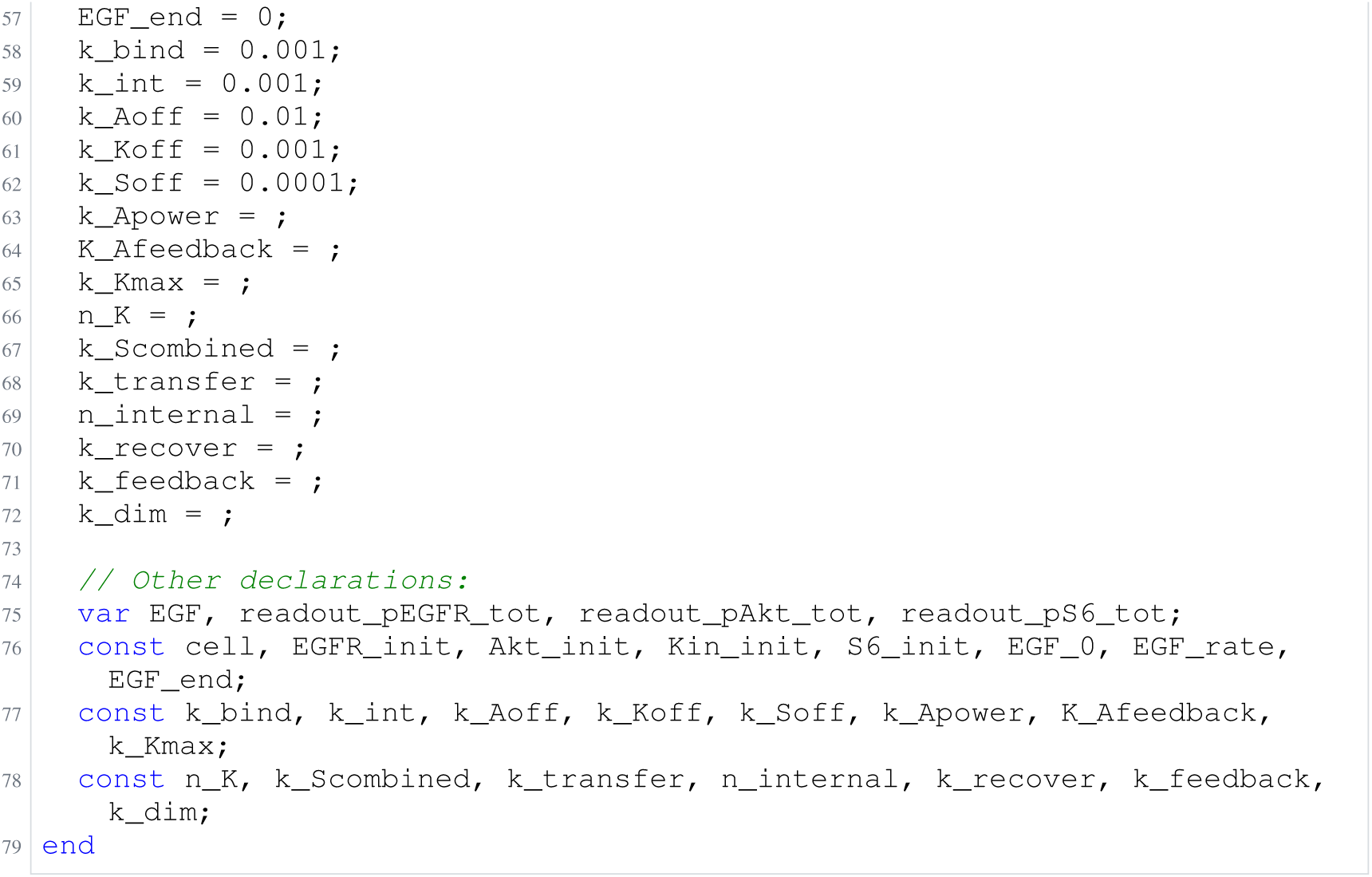
Archived Antimony for F2d: Panel, gpt-6-astra, biocurator off; task 56550507_3; selected version 53.

**Listing 23:**
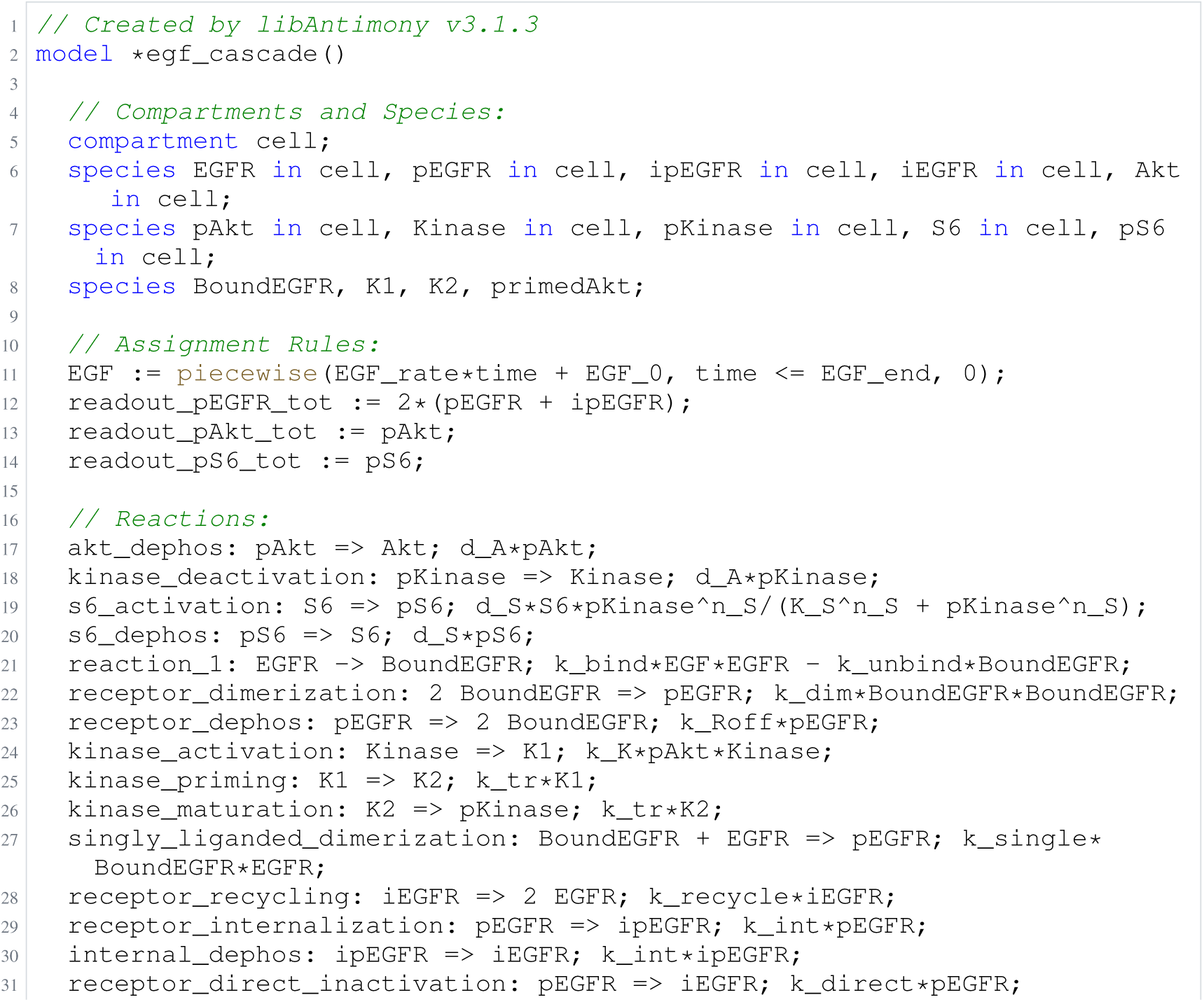
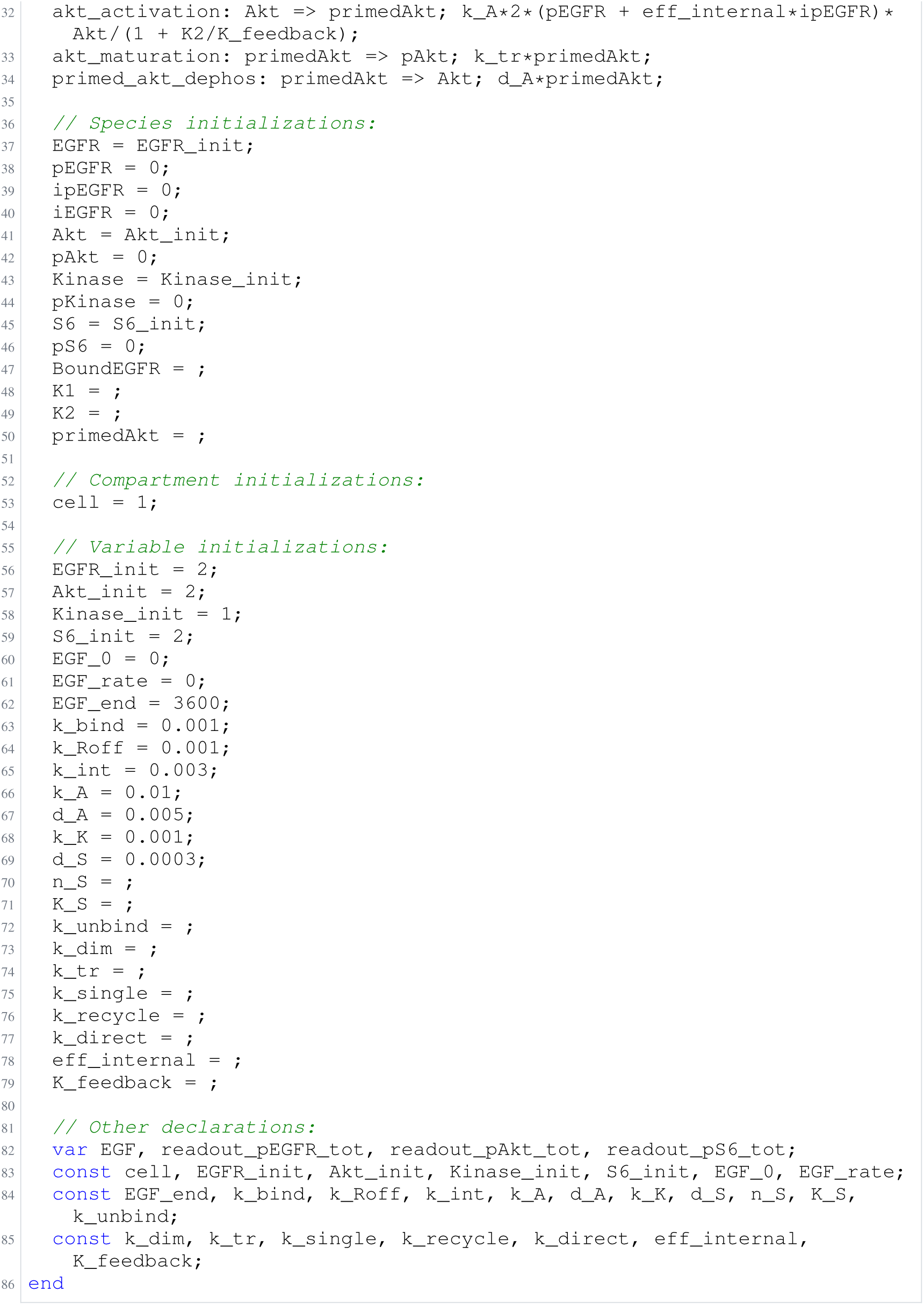
Archived Antimony for F2e: Panel, gpt-6-astra, biocurator off; task 56550507_4; selected version 28.

**Listing 24:**
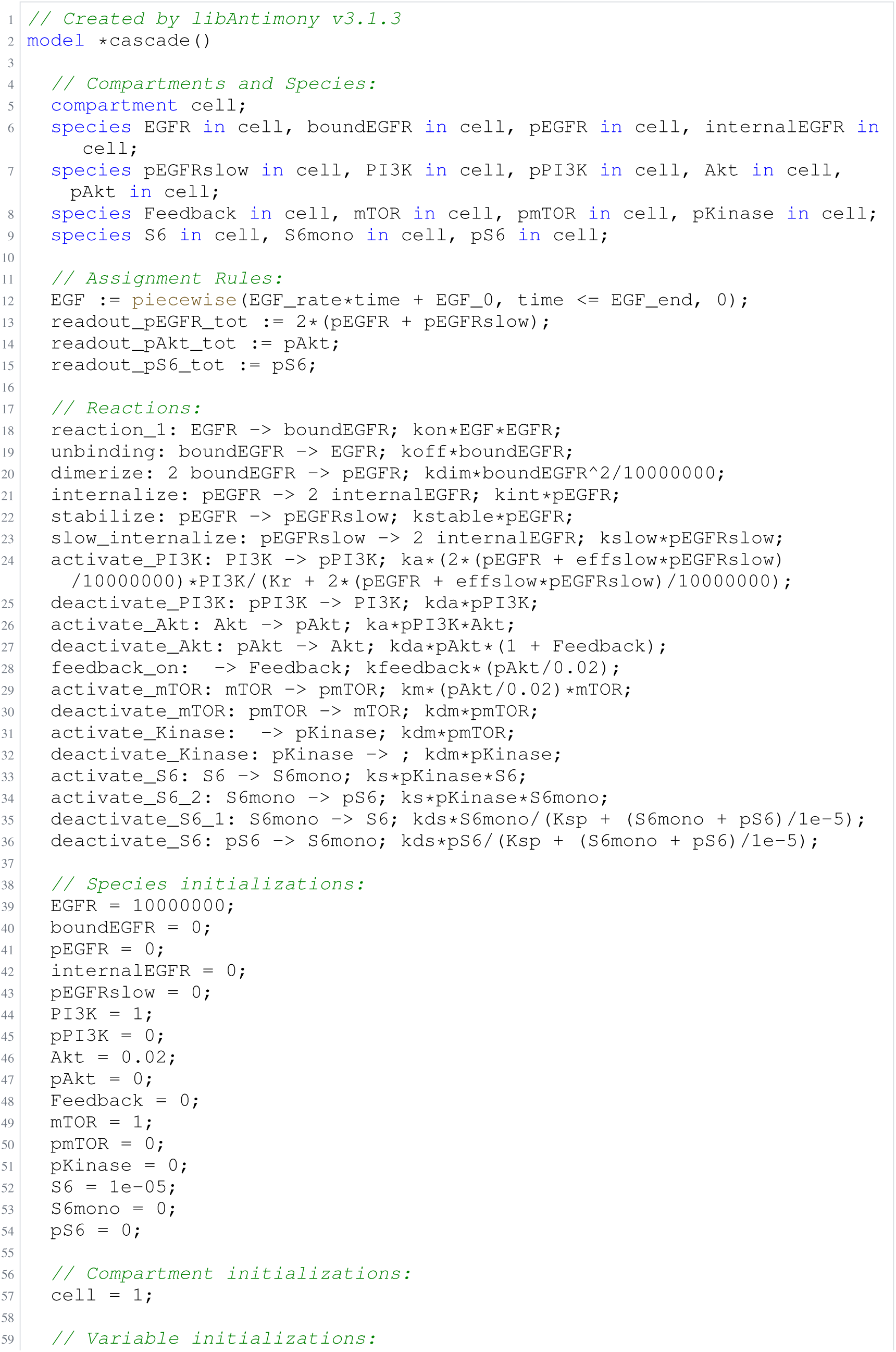
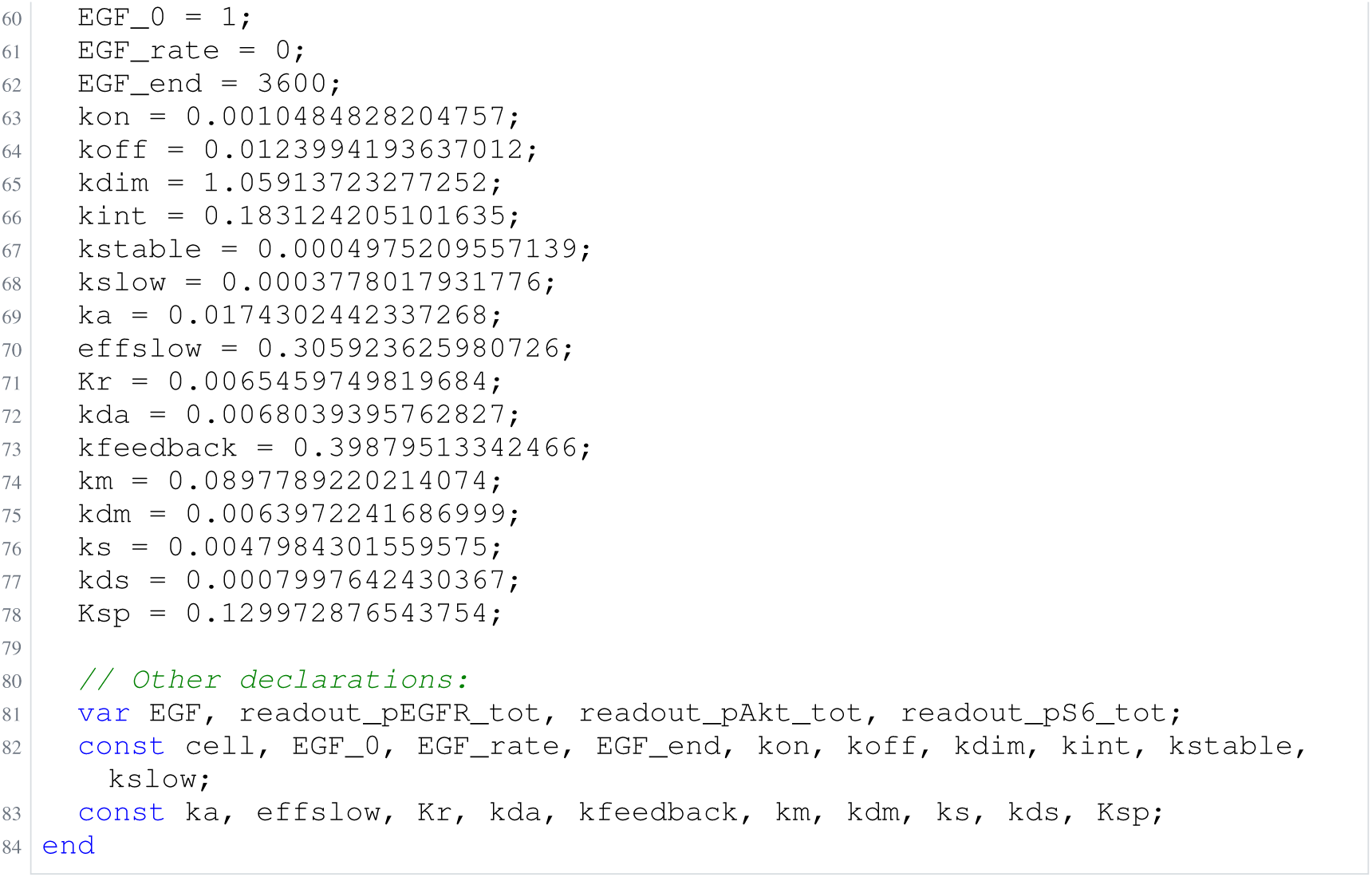
Archived Antimony for F3a: Null arm, gpt-6-astra; task 56550470_0; delivered problem, no version store; Antimony converted from the archived SBML.

**Listing 25:**
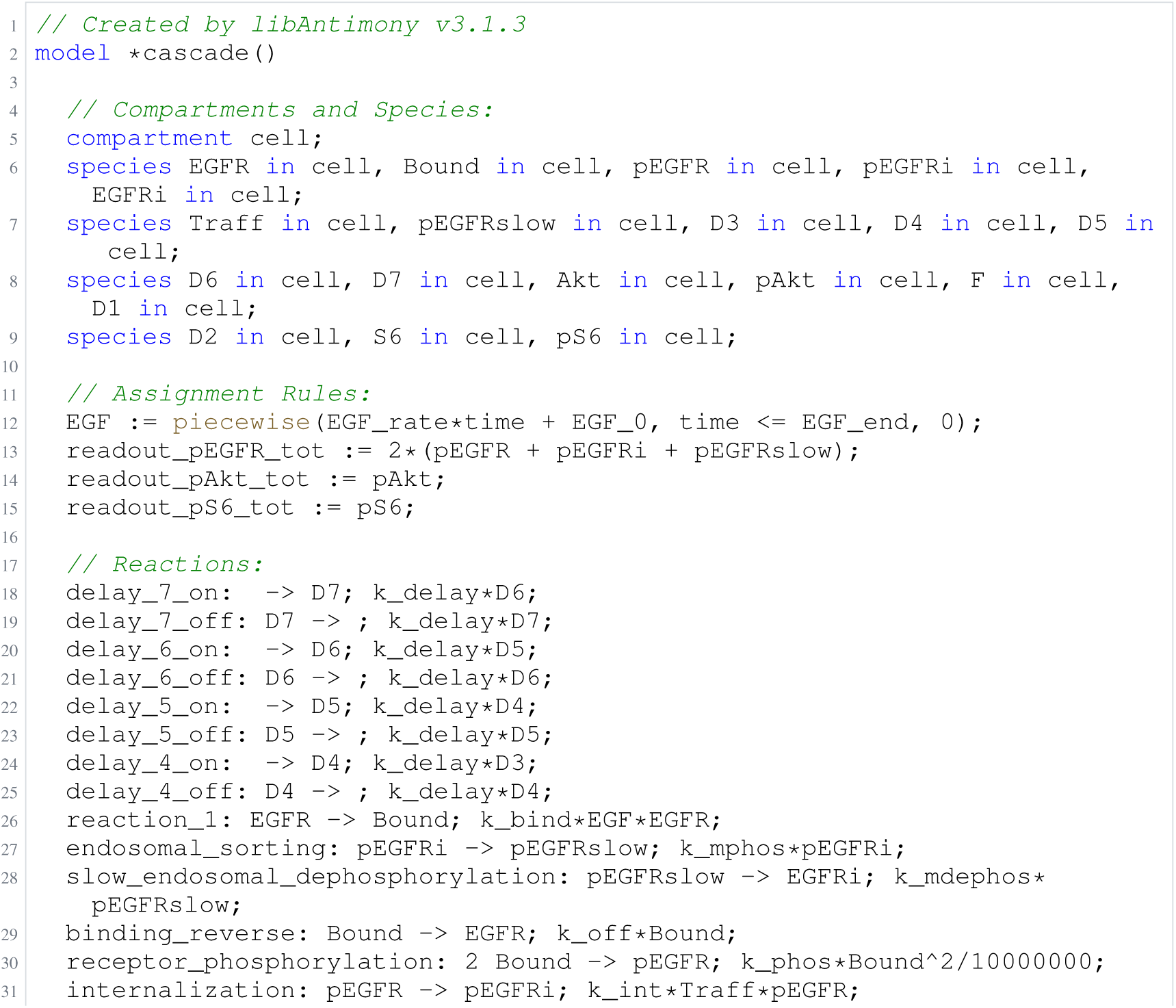
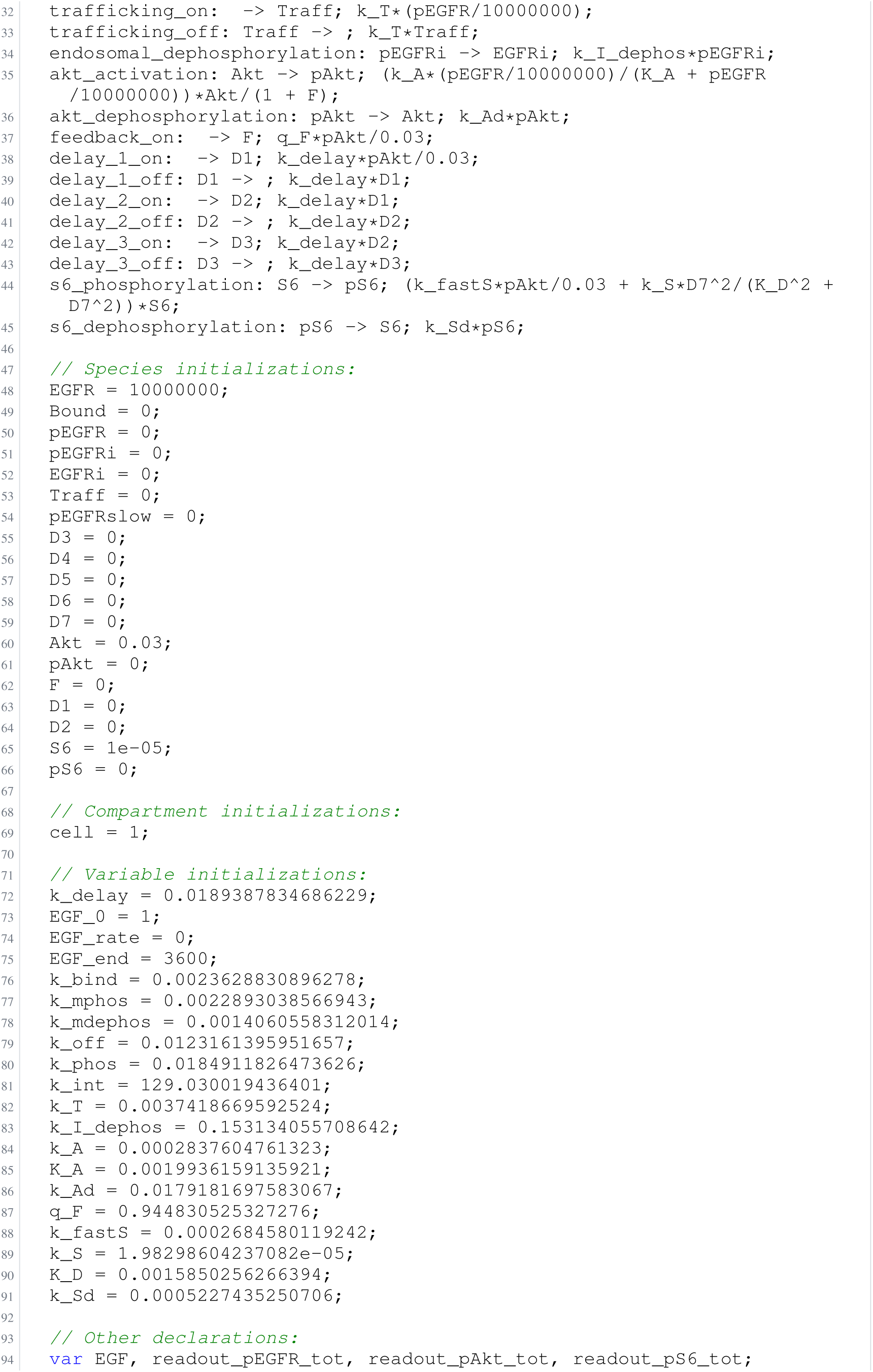
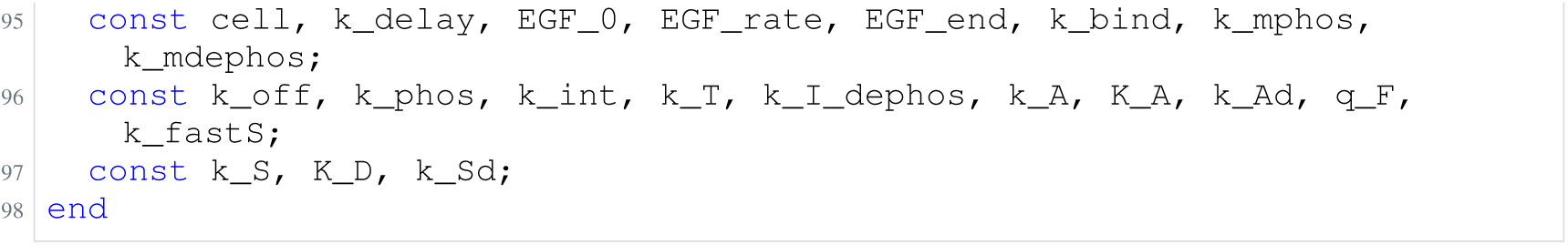
Archived Antimony for F3b: Null arm, gpt-6-astra; task 56550470_1; delivered problem, no version store; Antimony converted from the archived SBML.

**Listing 26:**
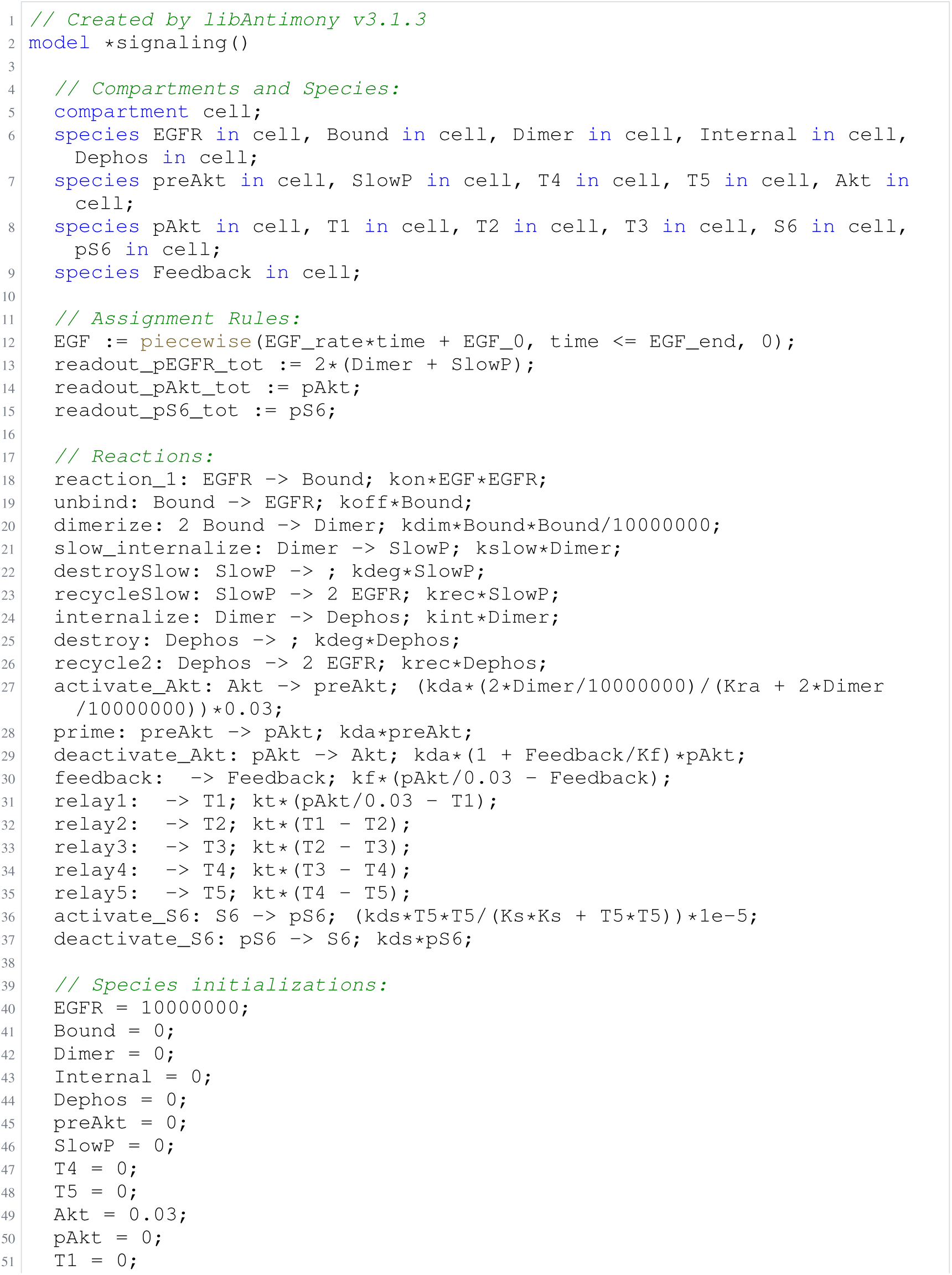
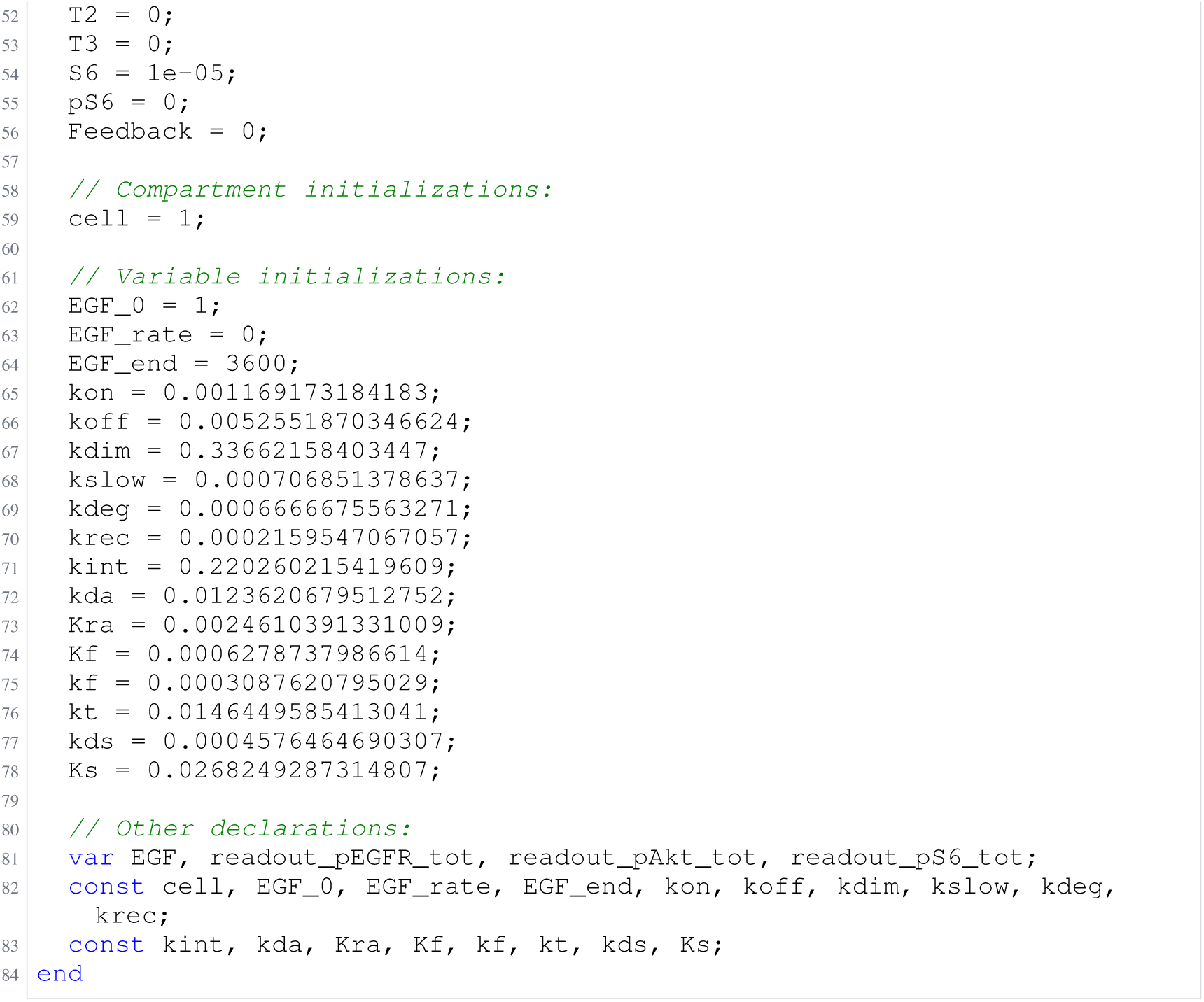
Archived Antimony for F3c: Null arm, gpt-6-astra; task 56550470_2; delivered problem, no version store; Antimony converted from the archived SBML.

**Listing 27:**
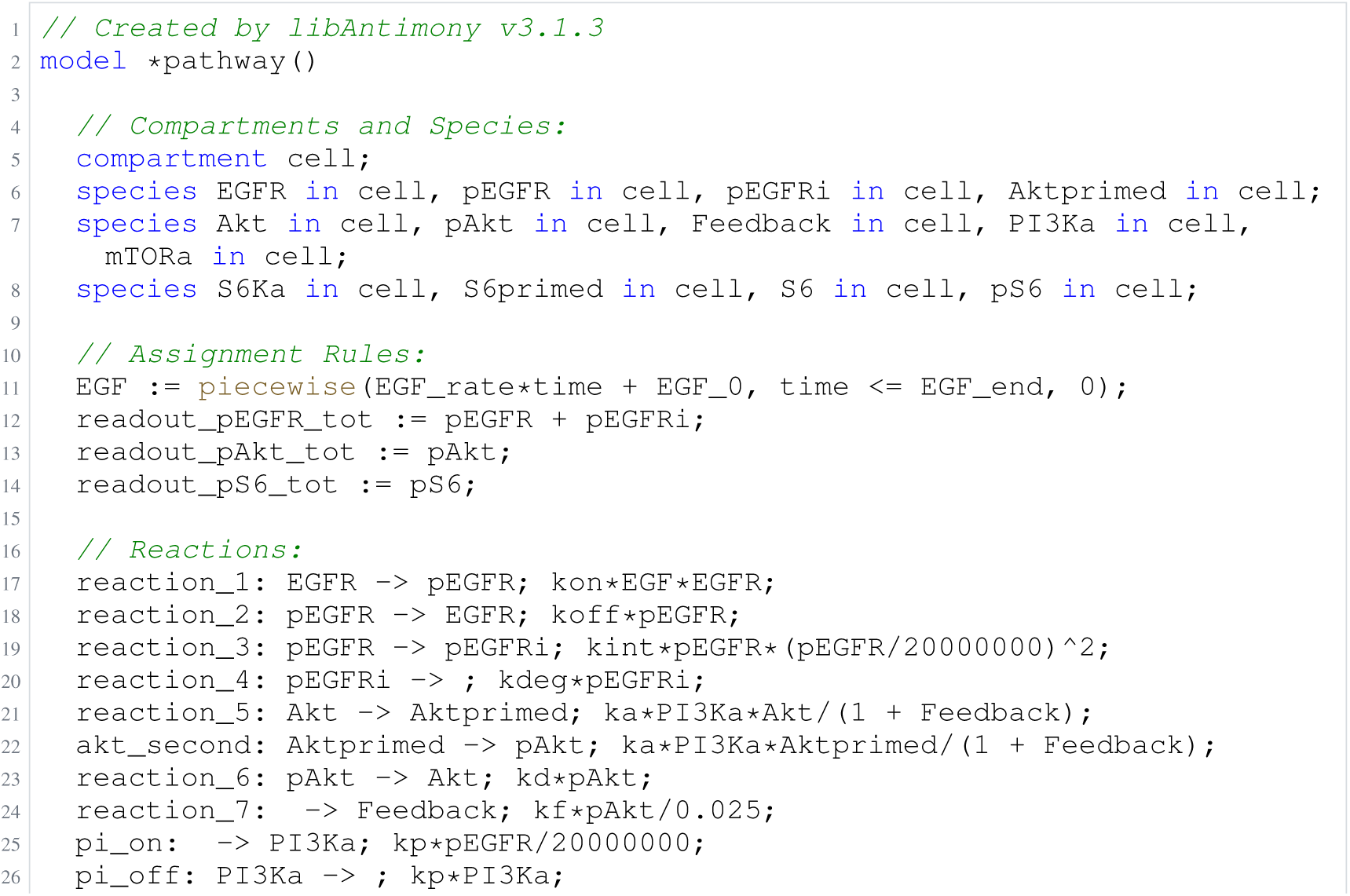
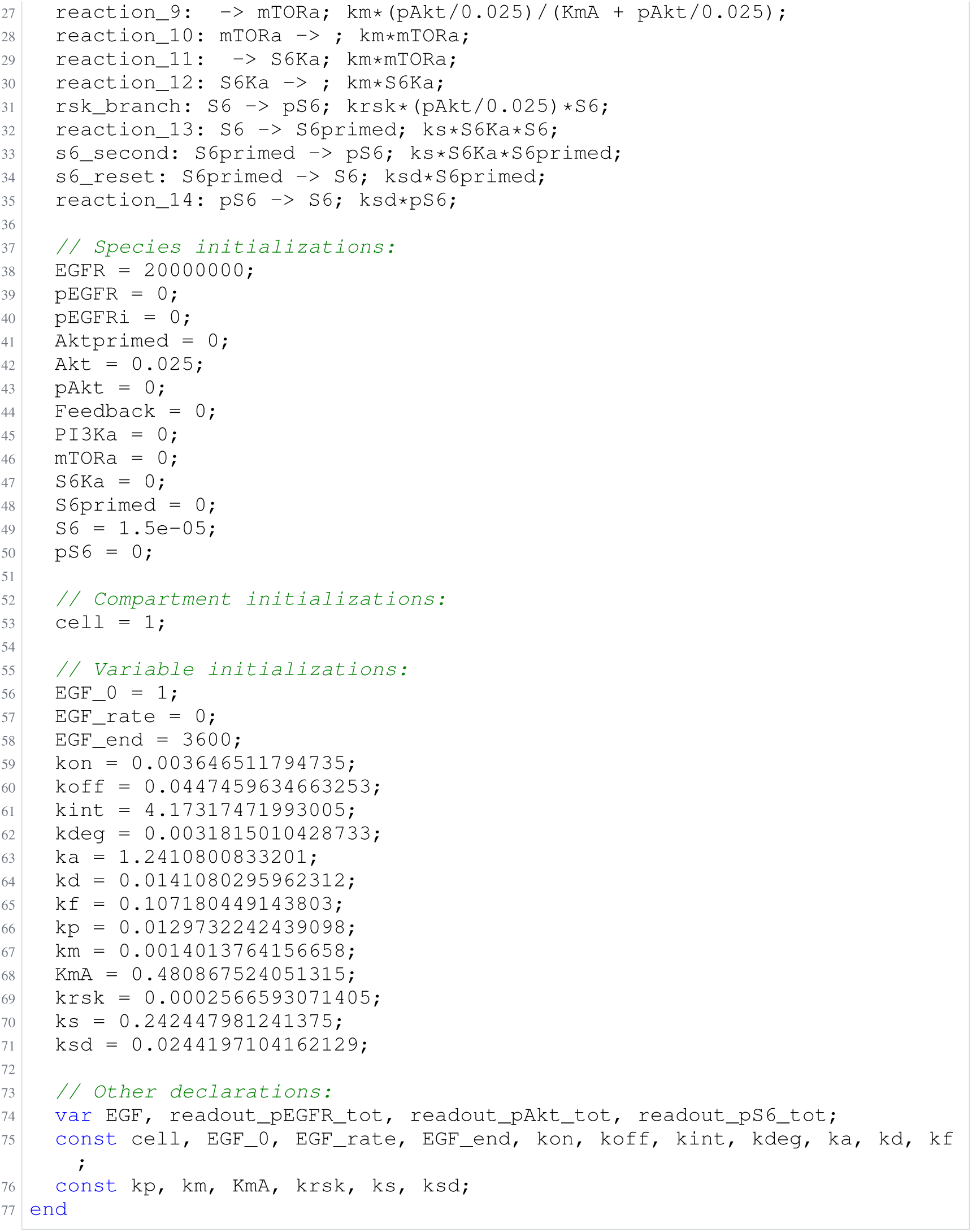
Archived Antimony for F3d: Null arm, gpt-6-astra; task 56550470_3; delivered problem, no version store; Antimony converted from the archived SBML.

**Listing 28:**
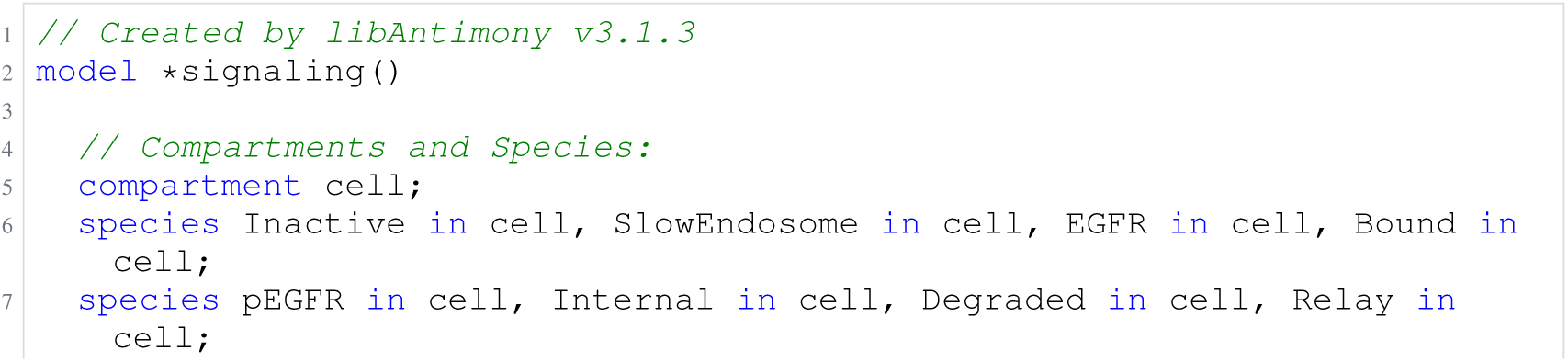
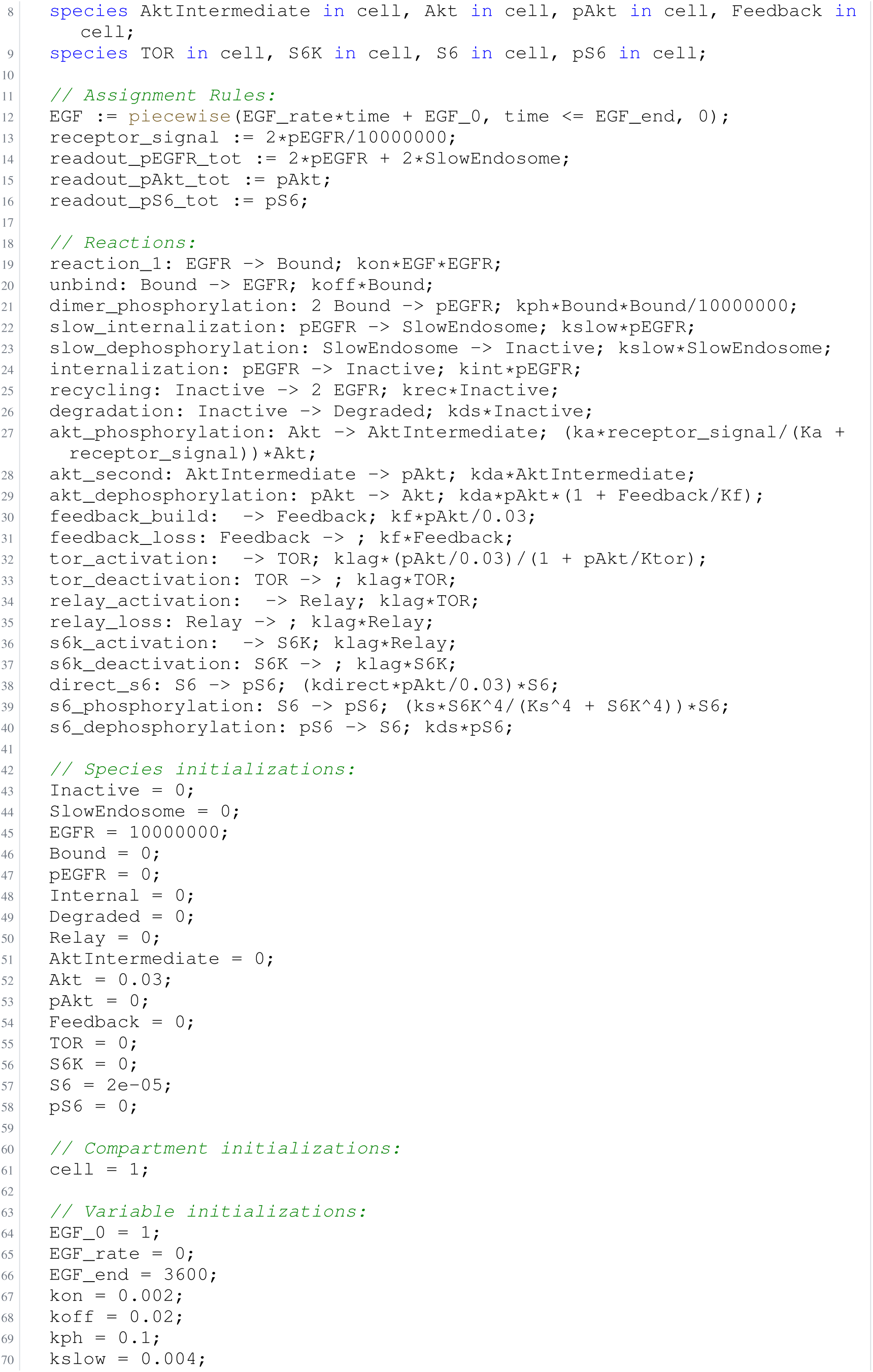
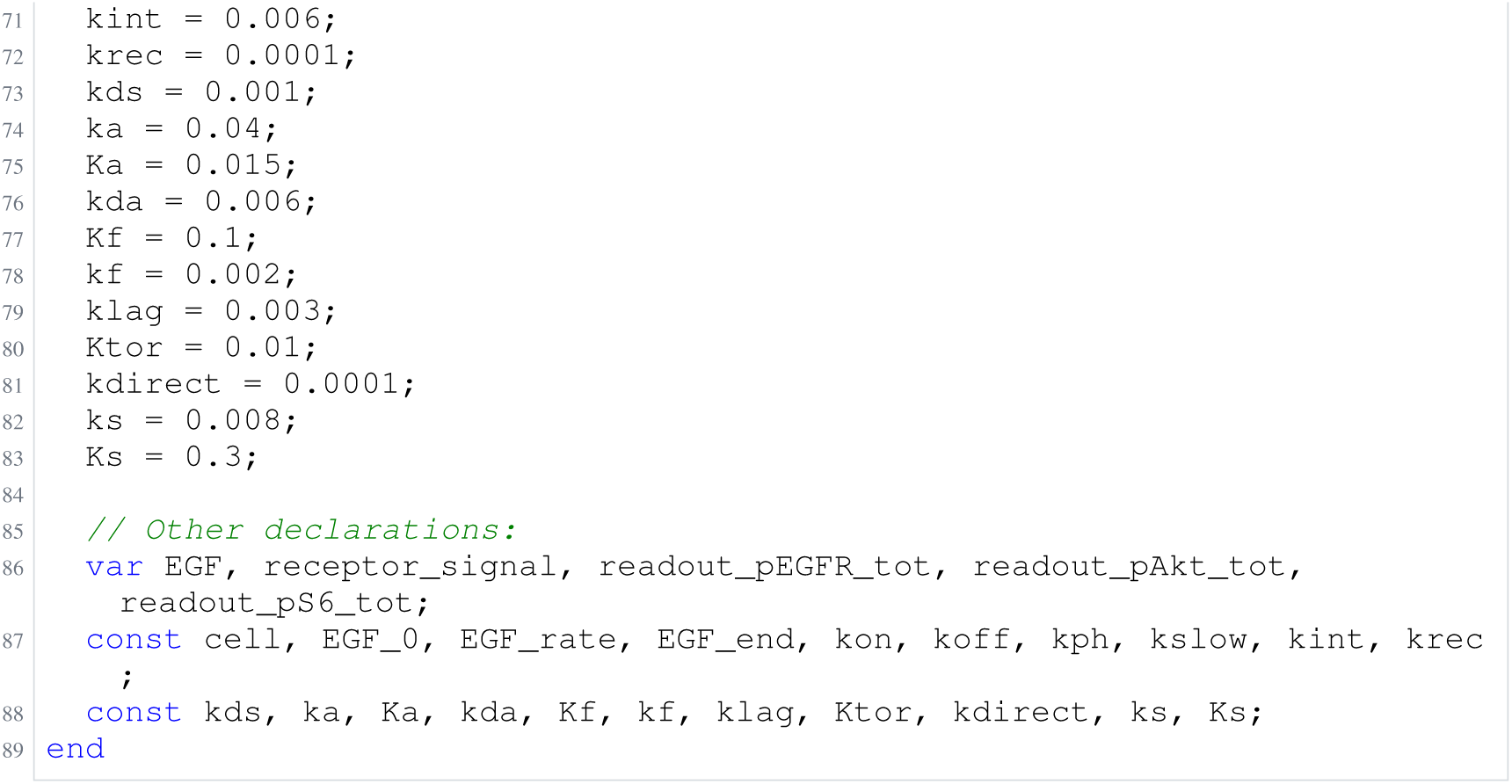
Archived Antimony for F3e: Null arm, gpt-6-astra; task 56550470_4; delivered problem, no version store; Antimony converted from the archived SBML.

## H Agent-generated benchmark model sources

These listings provide the 85 additional Antimony models from the benchmark collection of Section 2.2. The five Fujita models shared with that collection already appear in Appendix G and are cross-referenced below, completing the 90-model cohort. Examples follow the benchmark order; model numbers within each example follow run start time, rather than fit quality. Captions identify the model, array task, and selected version; archive identifiers and source hashes are retained in the accompanying `paper/model_sources/benchmark/models.csv` manifest.

Sources are preserved byte for byte, including comments and unset initial values. Their numerical assignments need not equal the fitted values: the saved parameter tables are supplied separately alongside each source. These files are not complete PEtab problems; simulation also requires the original conditions, observables, and initialization semantics. No models were reconstructed and no parameters were refitted for this annex.

### Shared Fujita sources

Shared Fujita sources.
| Benchmark model | Array task | Selected version | Existing listing |
| --- | --- | --- | --- |
| fujita_1 | 56476116_1 | 8 | O1b (Listing 2) |
| fujita_2 | 56476116_0 | 22 | O1a (Listing 1) |
| fujita_3 | 56476116_4 | 20 | O1e (Listing 5) |
| fujita_4 | 56476116_3 | 29 | O1d (Listing 4) |
| fujita_5 | 56476116_2 | 28 | O1c (Listing 3) |

### H.1 Perelson

**Listing 29:**
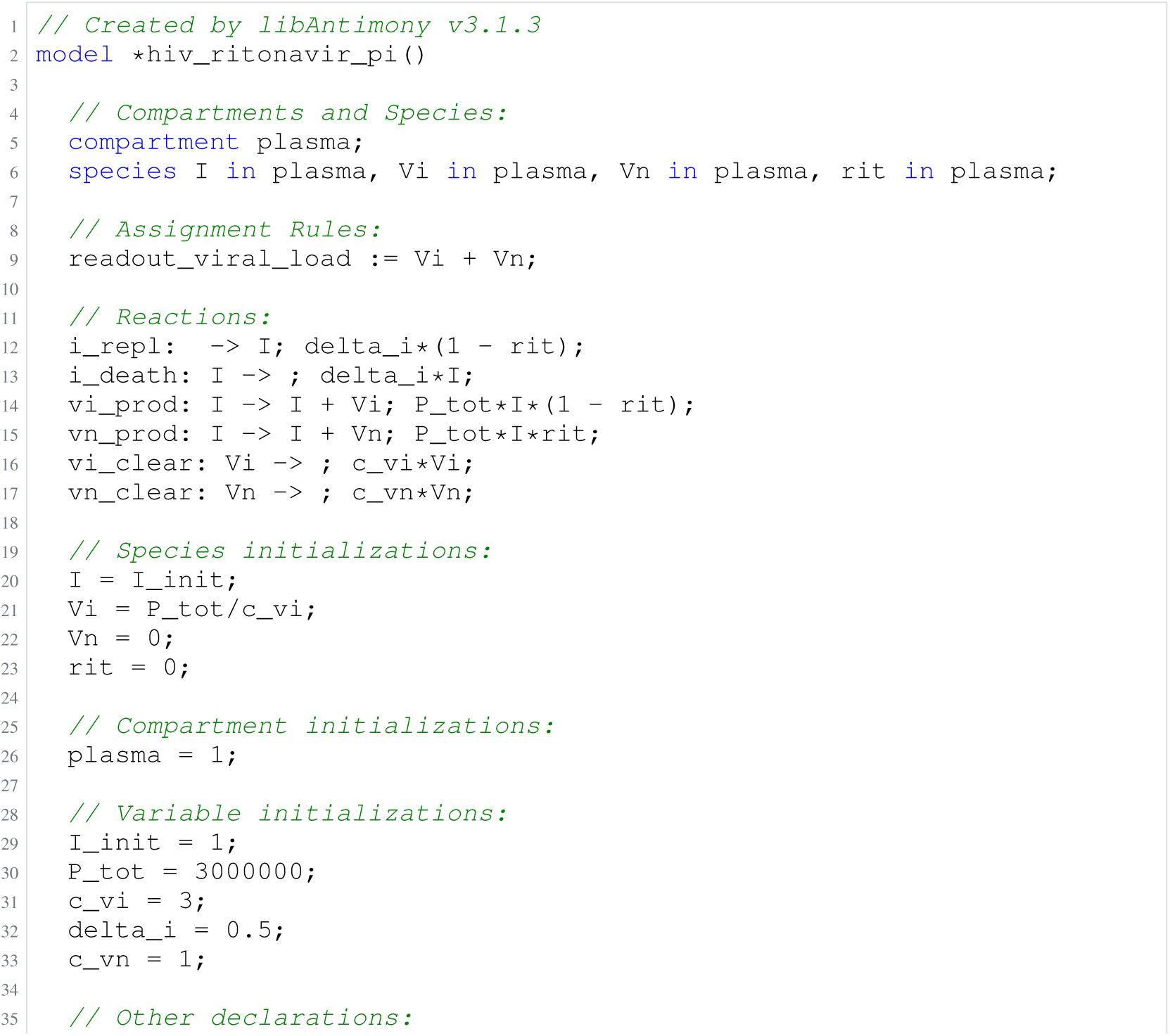
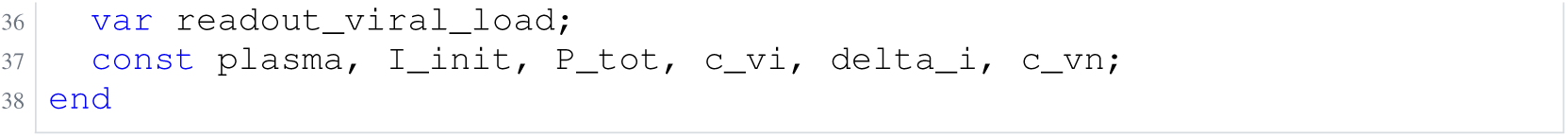
Archived Antimony for perelson_1; task 57485624_3; selected version 1.

**Listing 30:**
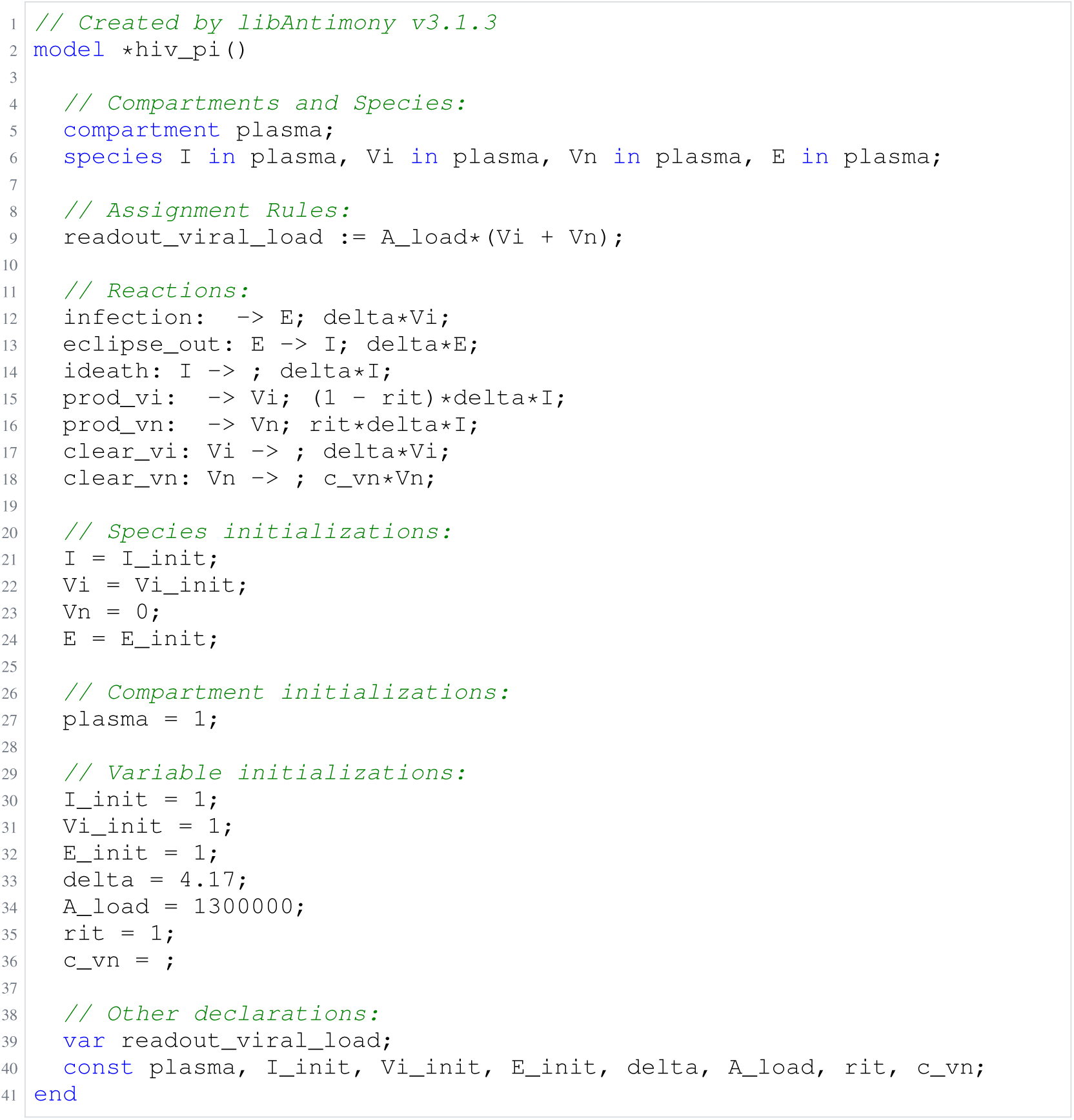
Archived Antimony for perelson_2; task 57581393_0; selected version 4.

**Listing 31:**
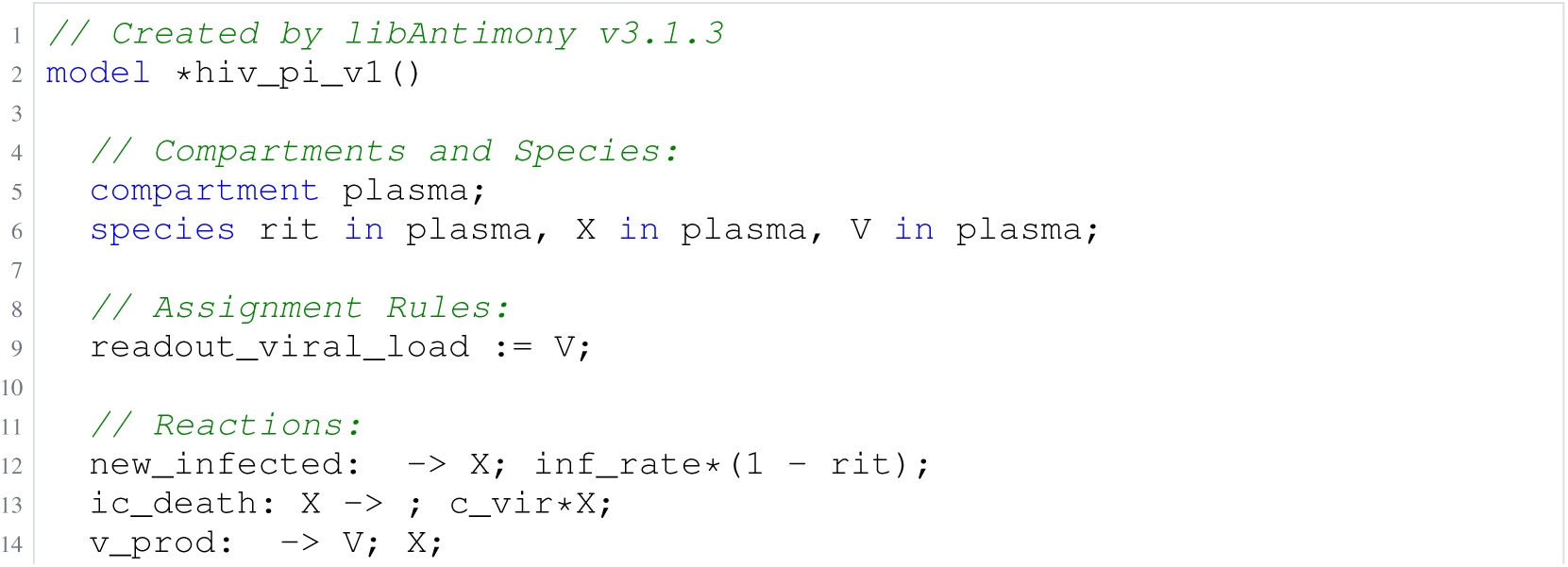
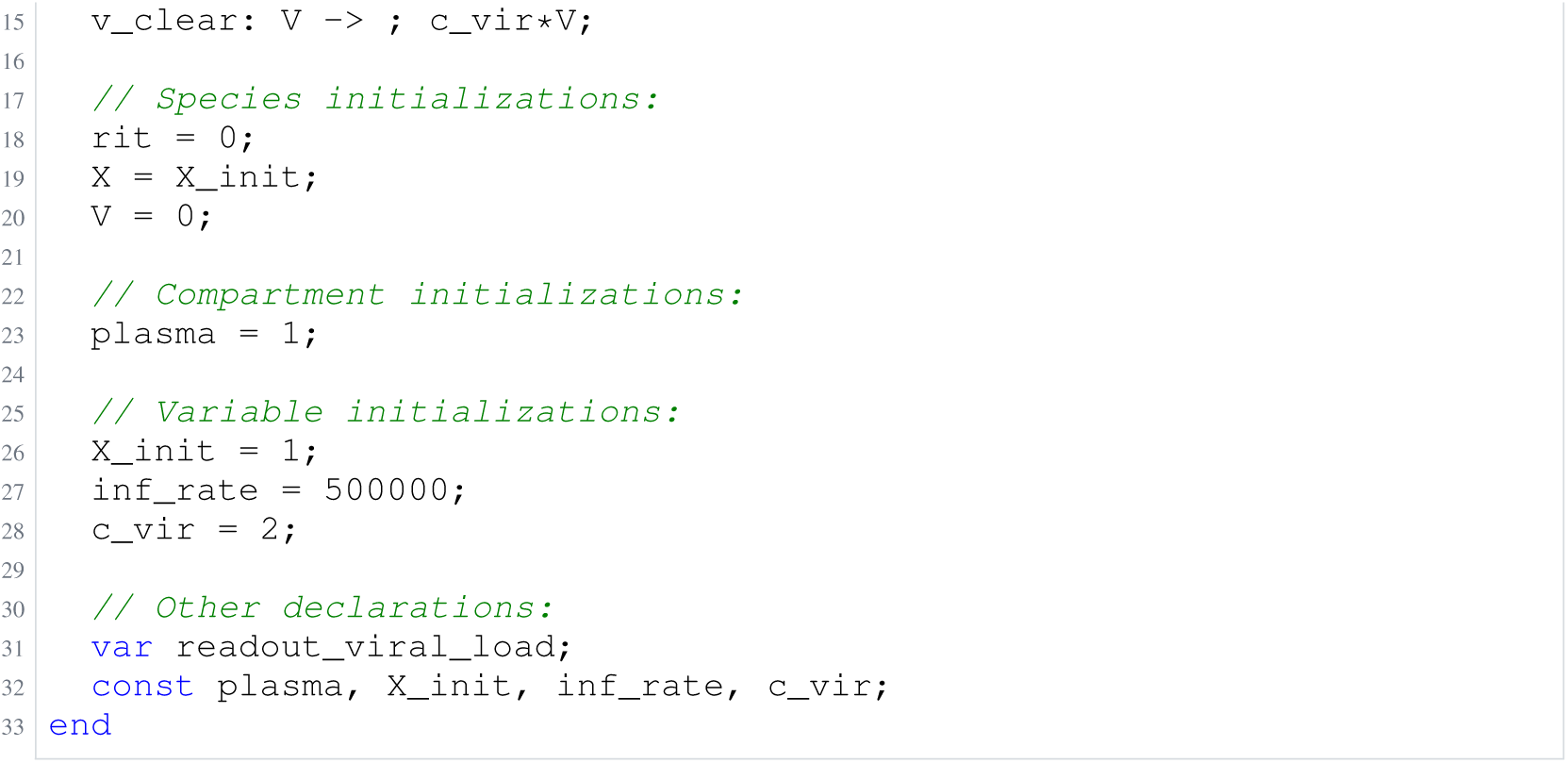
Archived Antimony for perelson_3; task 57581393_4; selected version 1.

**Listing 32:**
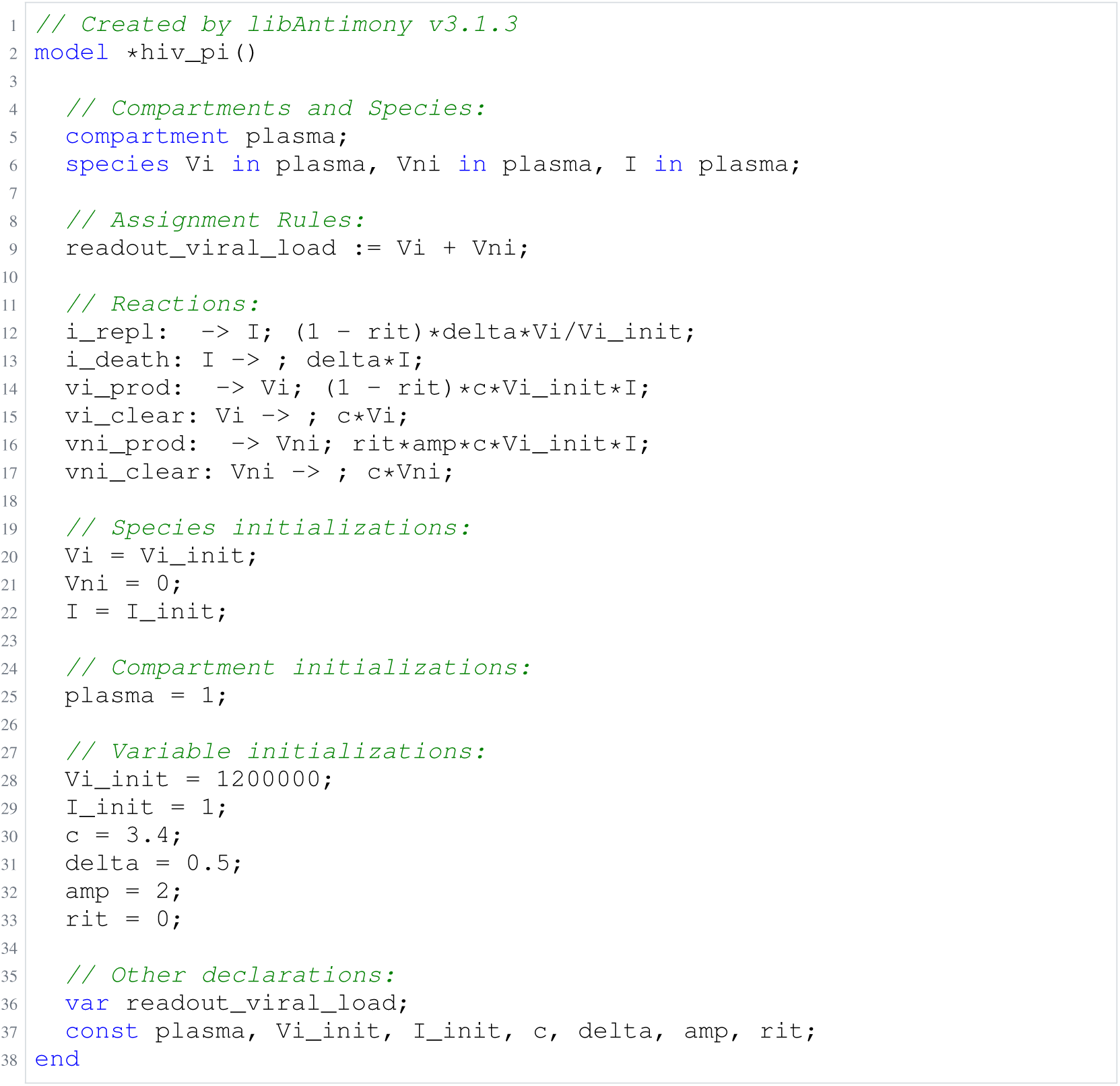
Archived Antimony for perelson_4; task 57581393_2; selected version 2.

**Listing 33:**
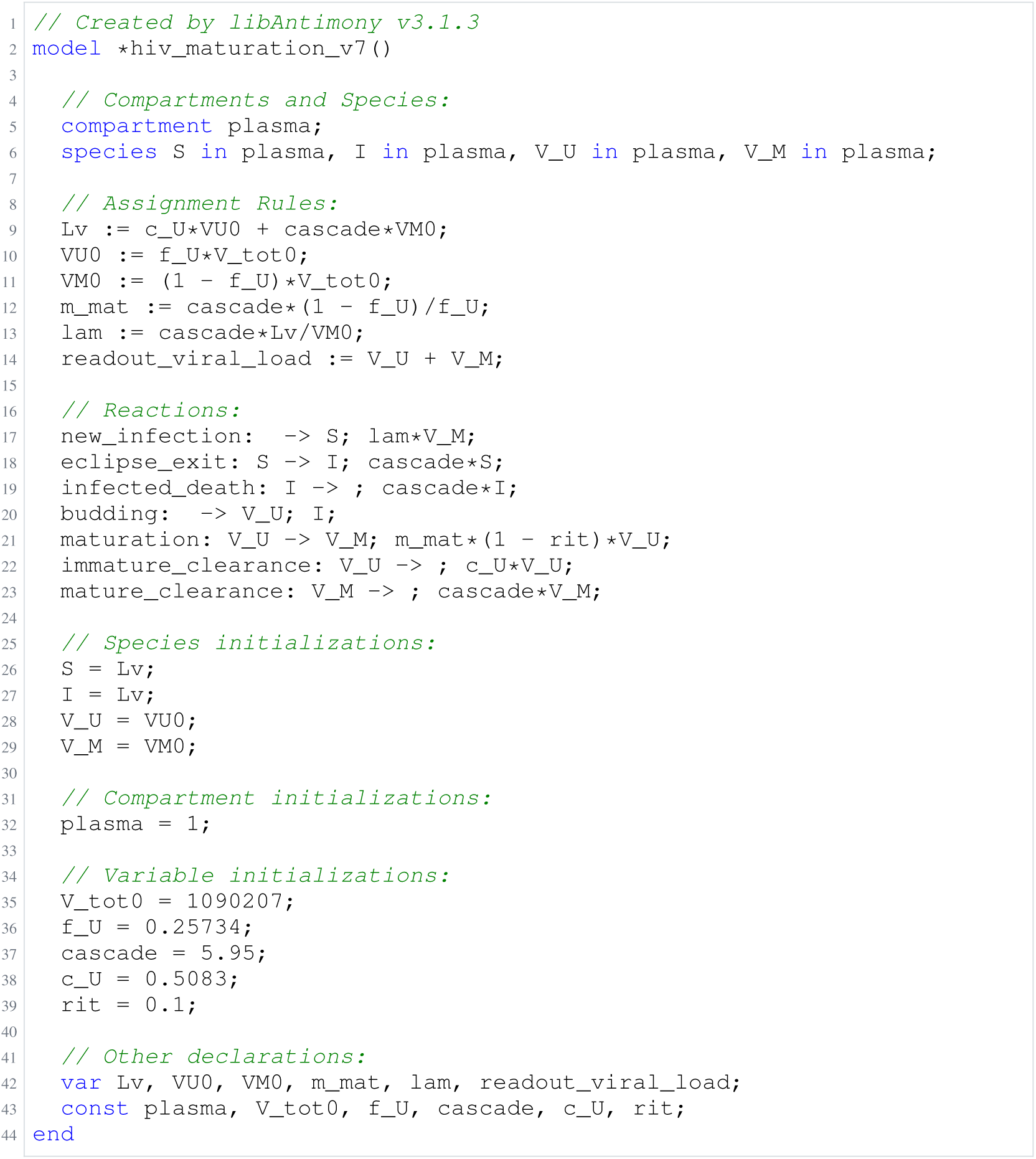
Archived Antimony for perelson_5; task 57581393_1; selected version 7.

### H.2 Crauste

**Listing 34:**
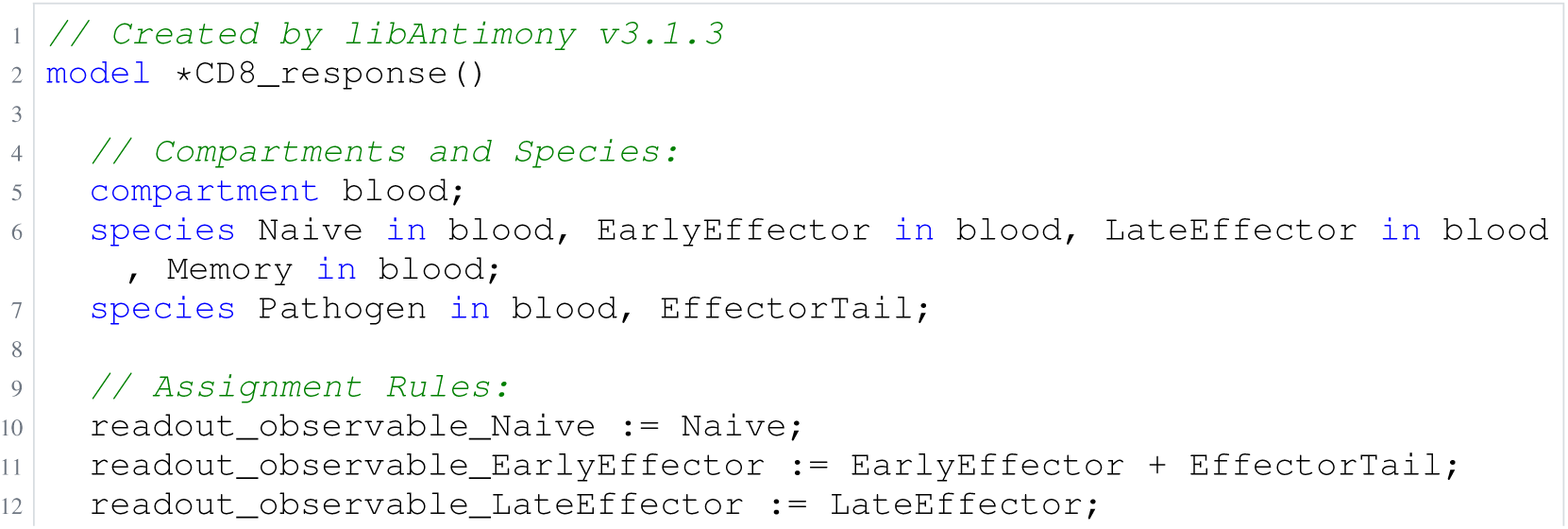
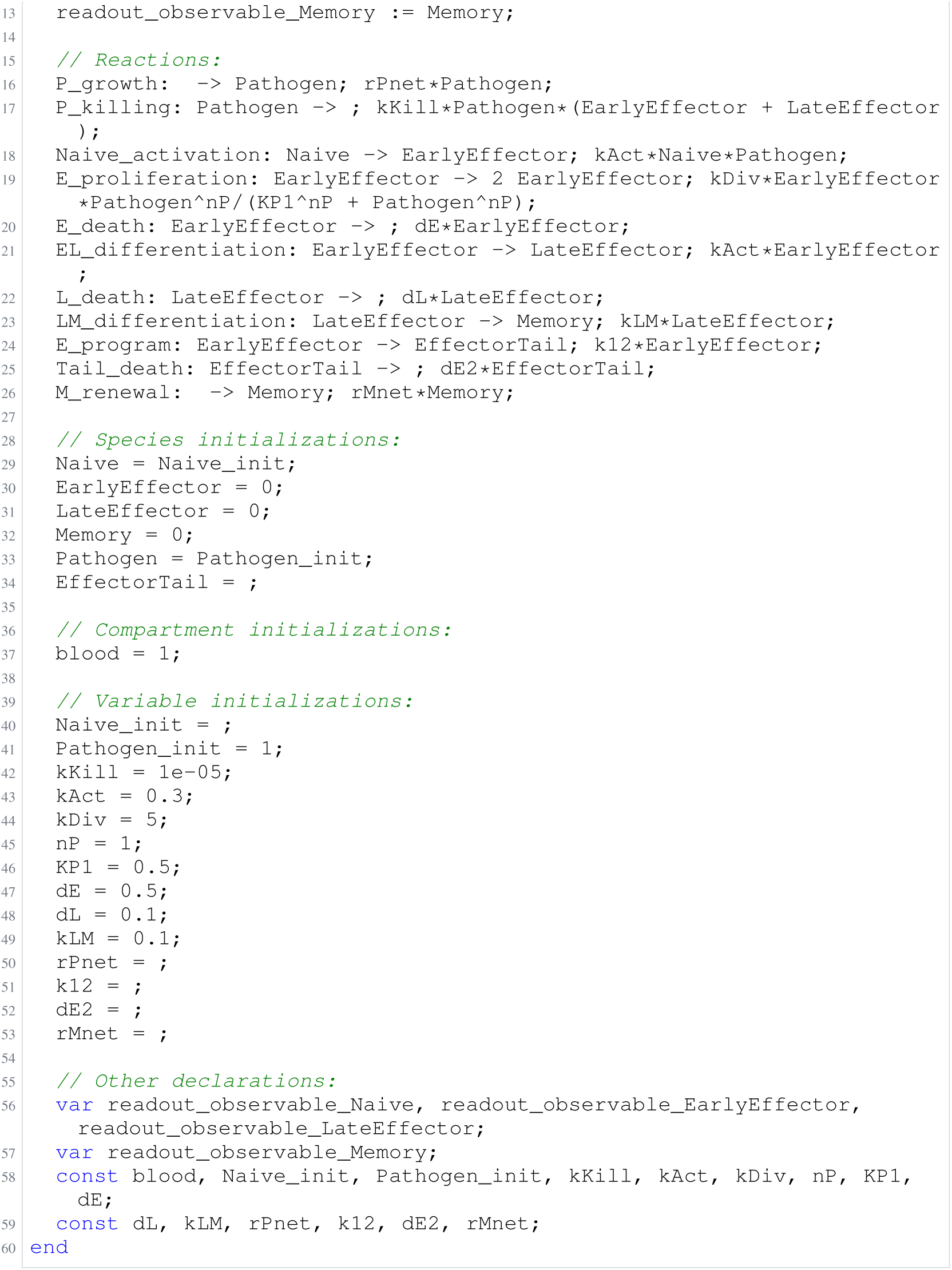
Archived Antimony for crauste_1; task 56836848_25; selected version 11.

**Listing 35:**
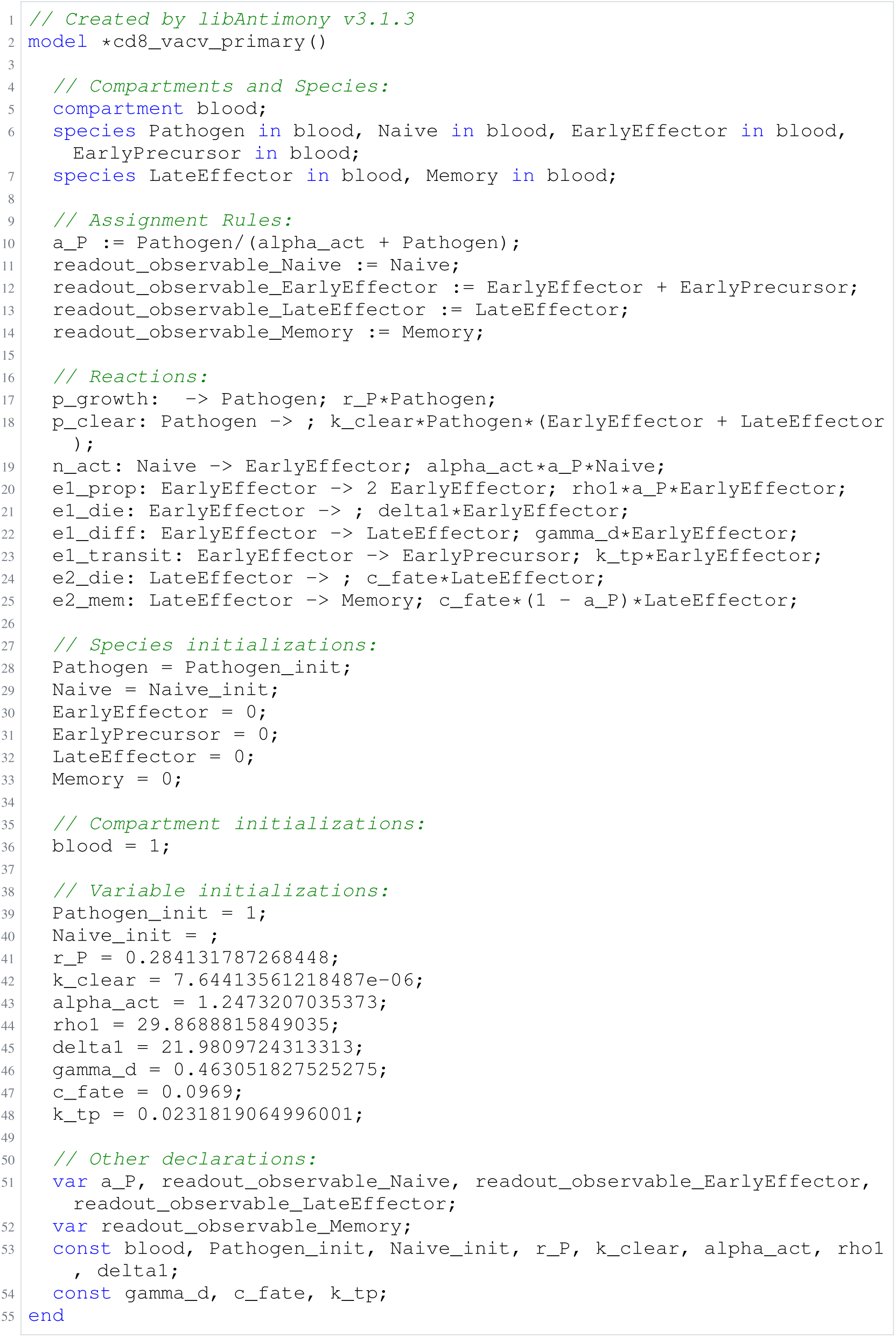
Archived Antimony for crauste_2; task 56836848_26; selected version 11.

**Listing 36:**
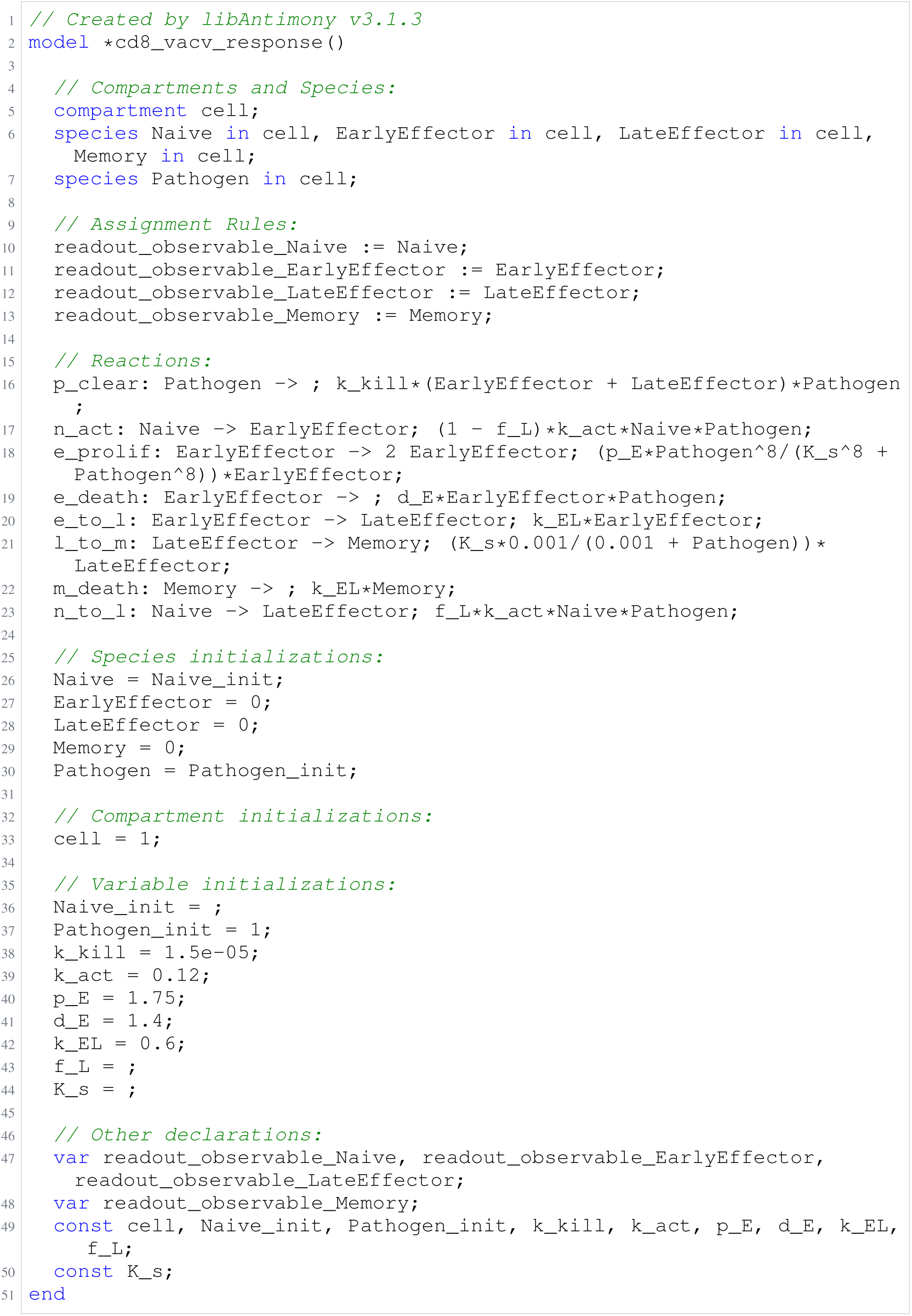
Archived Antimony for crauste_3; task 56836848_27; selected version 16.

**Listing 37:**
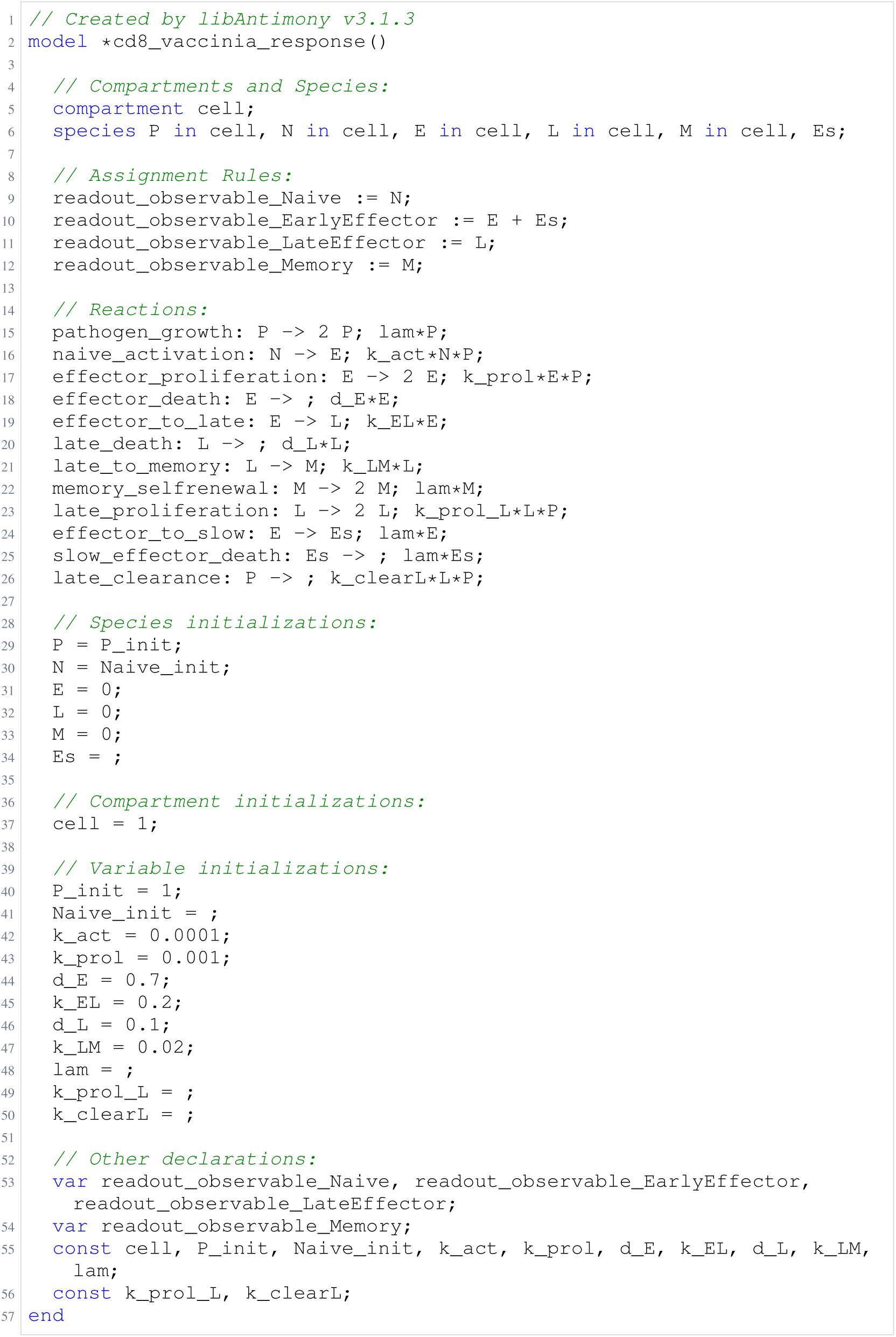
Archived Antimony for crauste_4; task 56836848_28; selected version 9.

**Listing 38:**
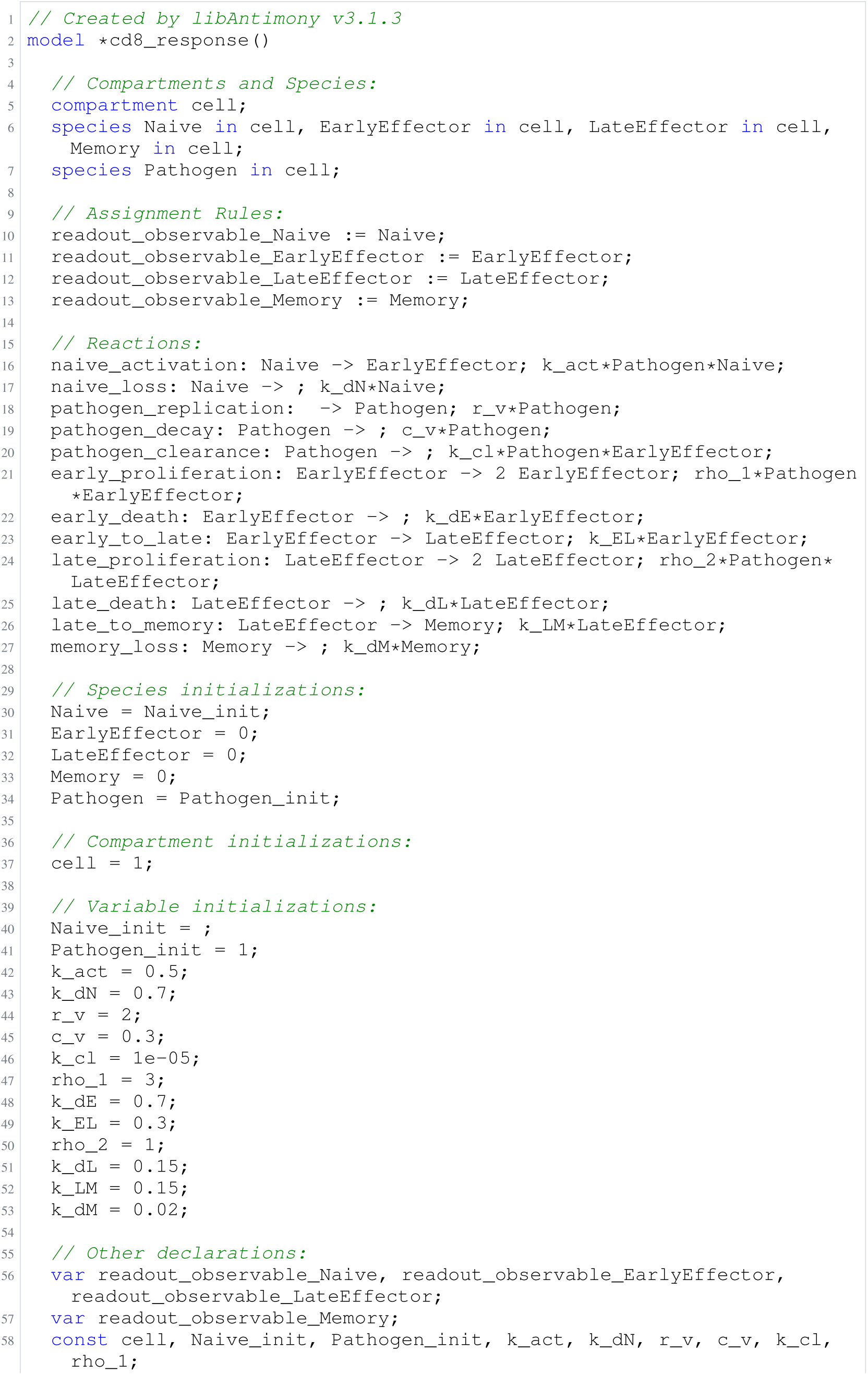
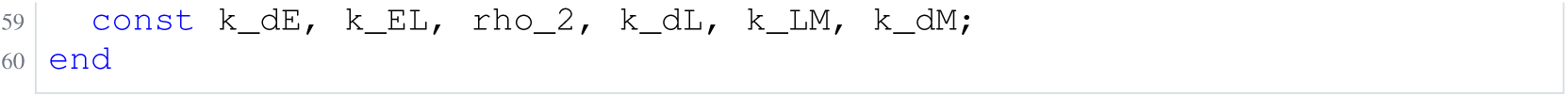
Archived Antimony for crauste_5; task 56836848_29; selected version 0.

### H.3 Rahman

**Listing 39:**
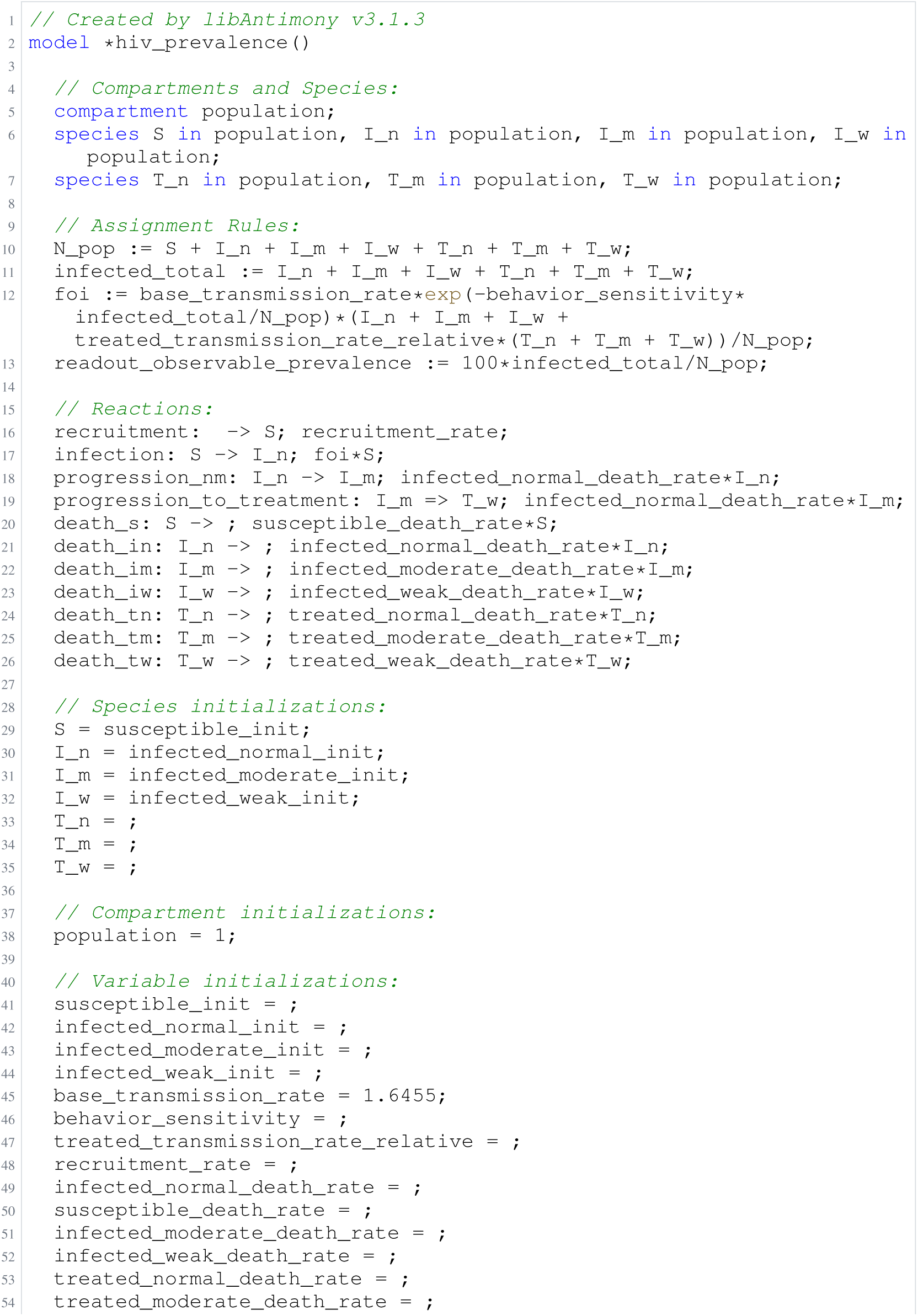
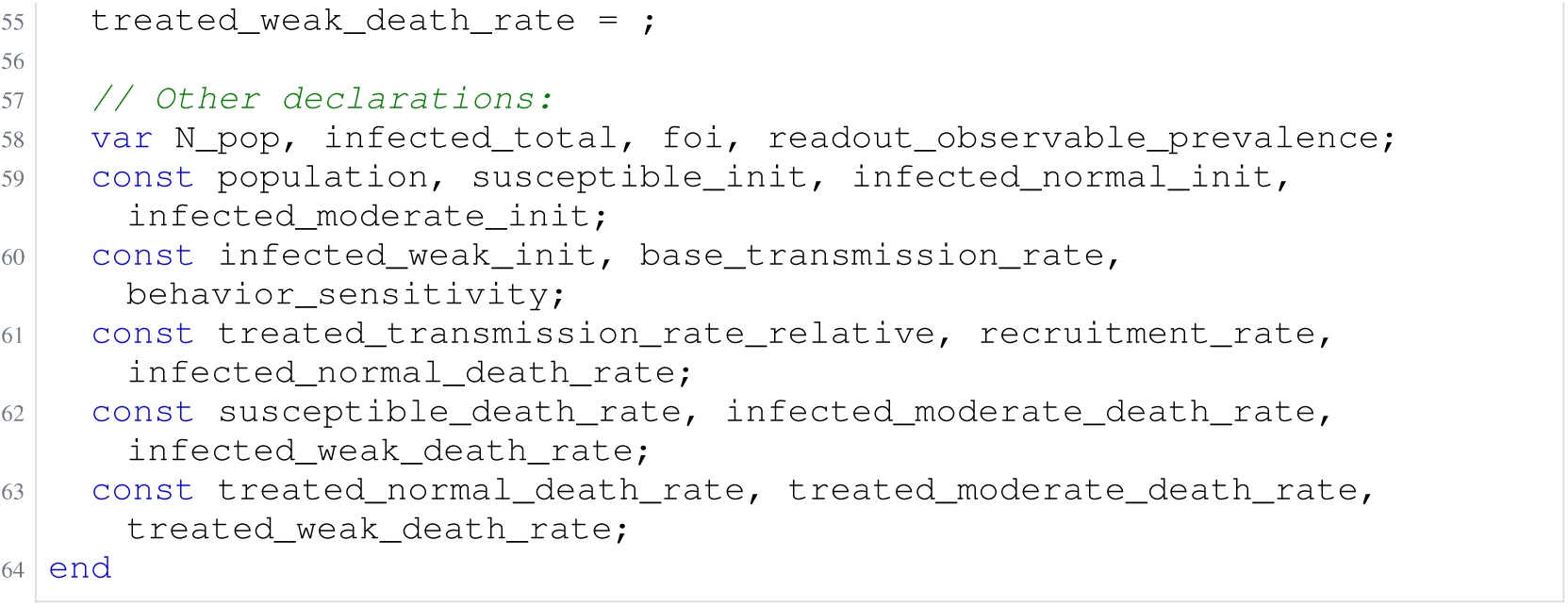
Archived Antimony for rahman_1; task 56836848_41; selected version 12.

**Listing 40:**
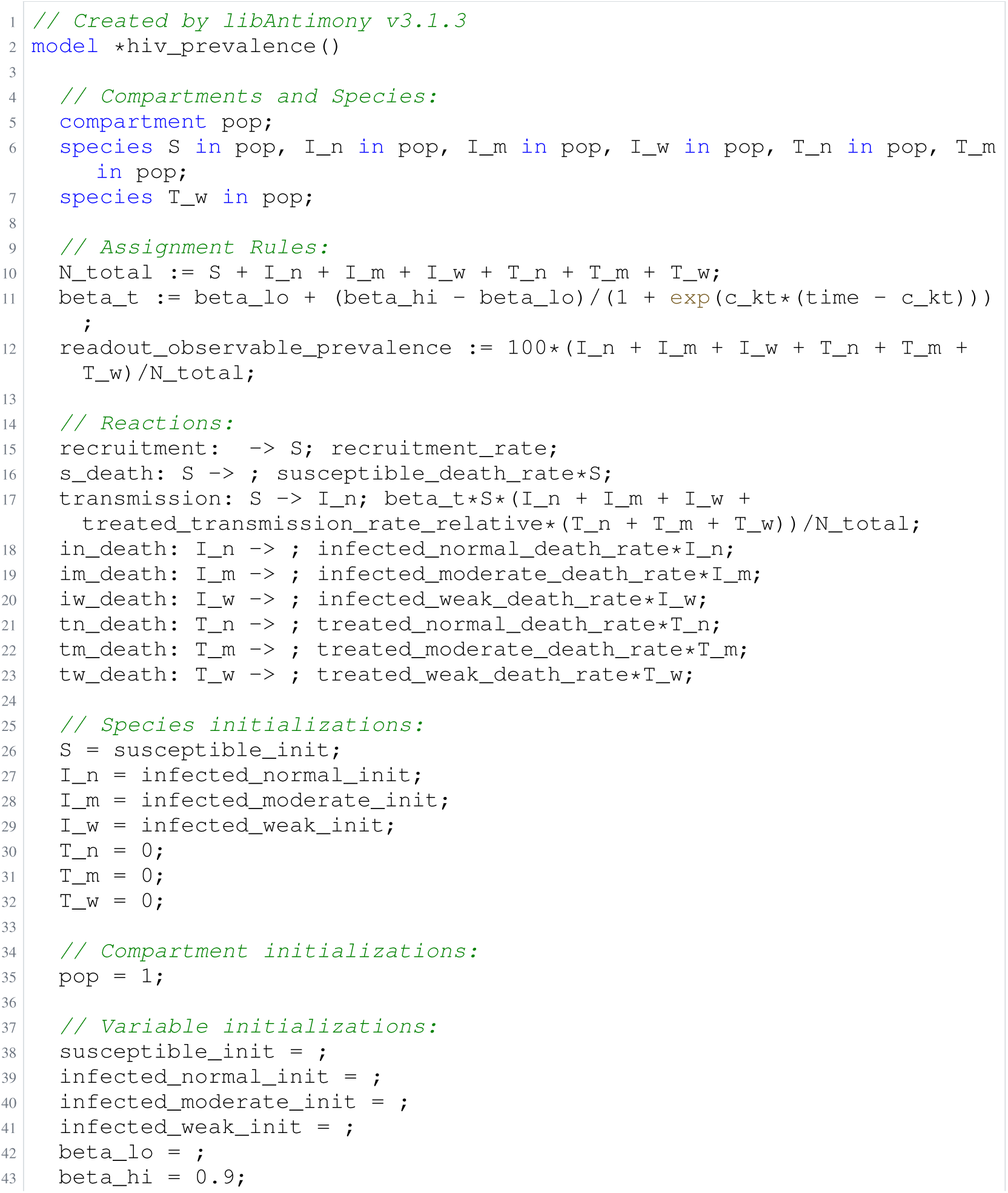
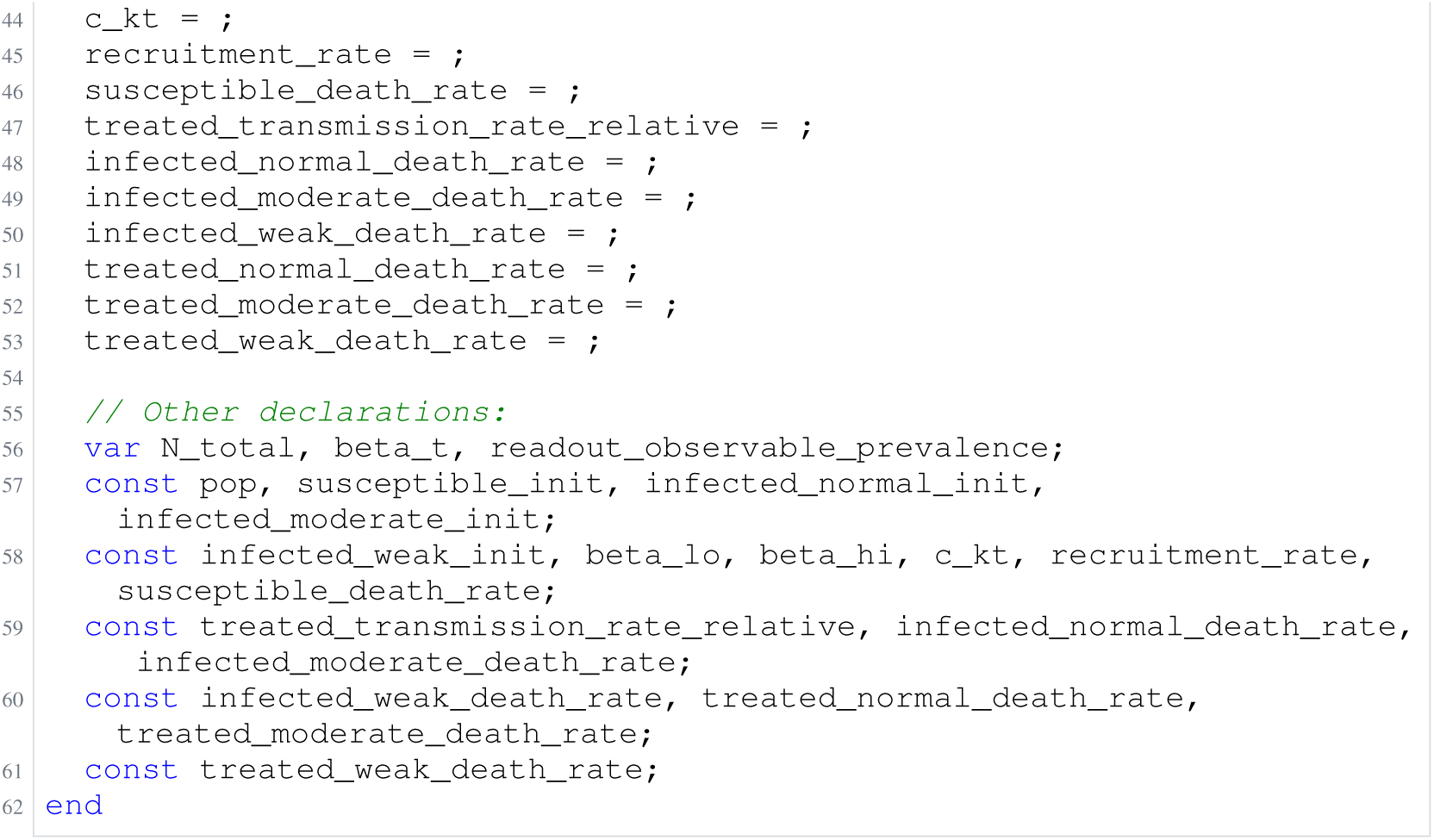
Archived Antimony for rahman_2; task 56836848_40; selected version 7.

**Listing 41:**
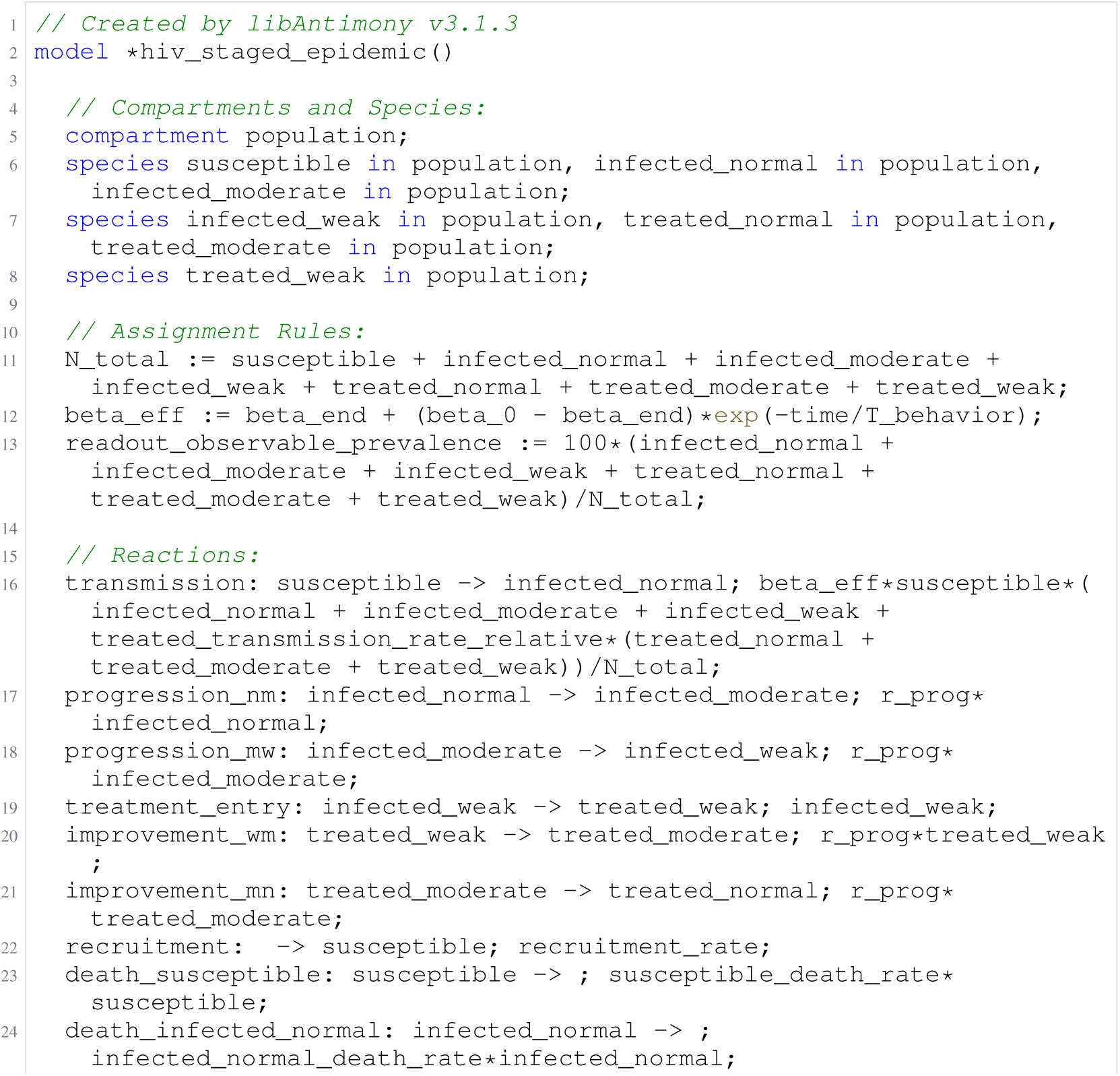
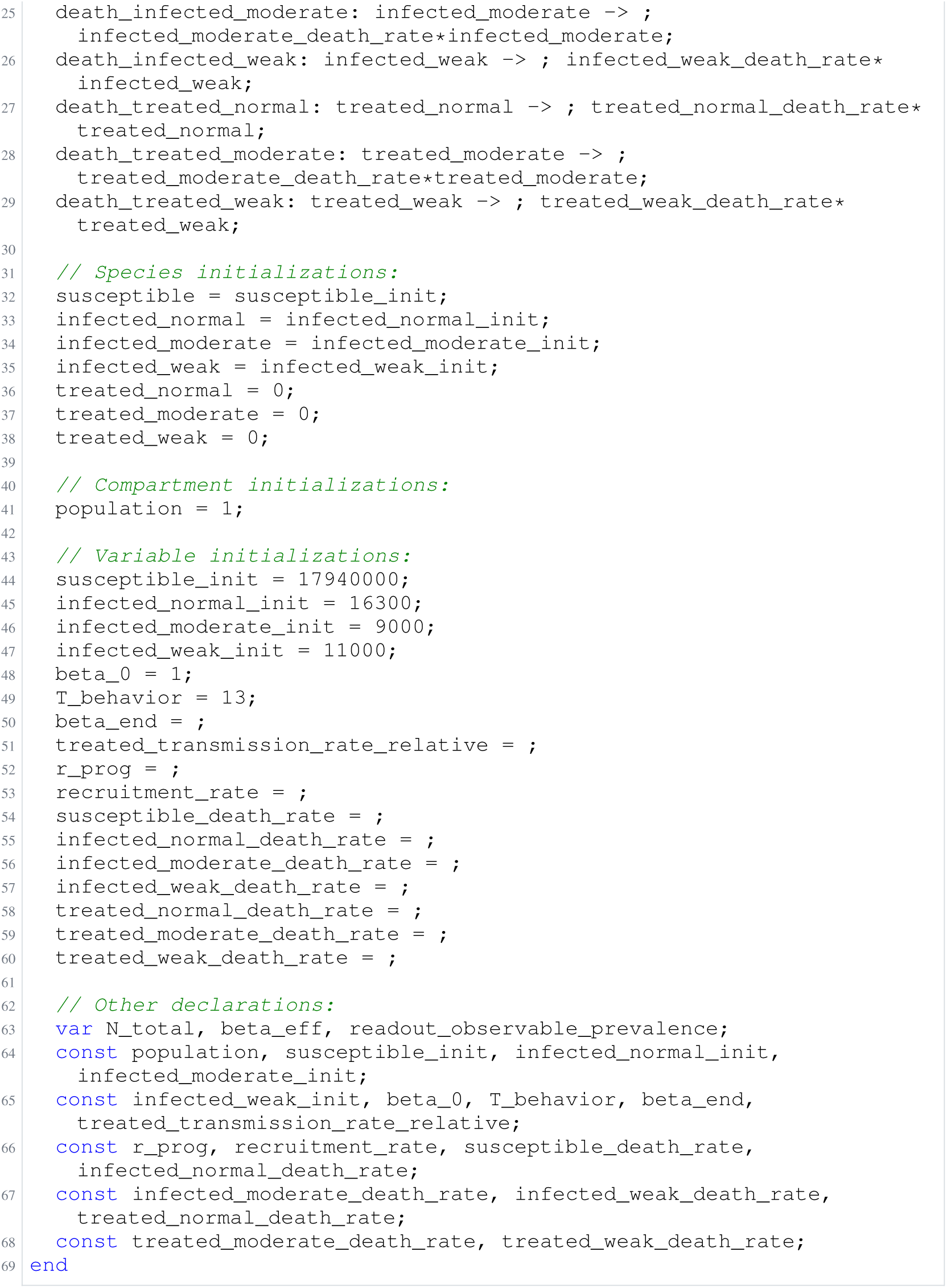
Archived Antimony for rahman_3; task 56836848_42; selected version 8.

**Listing 42:**
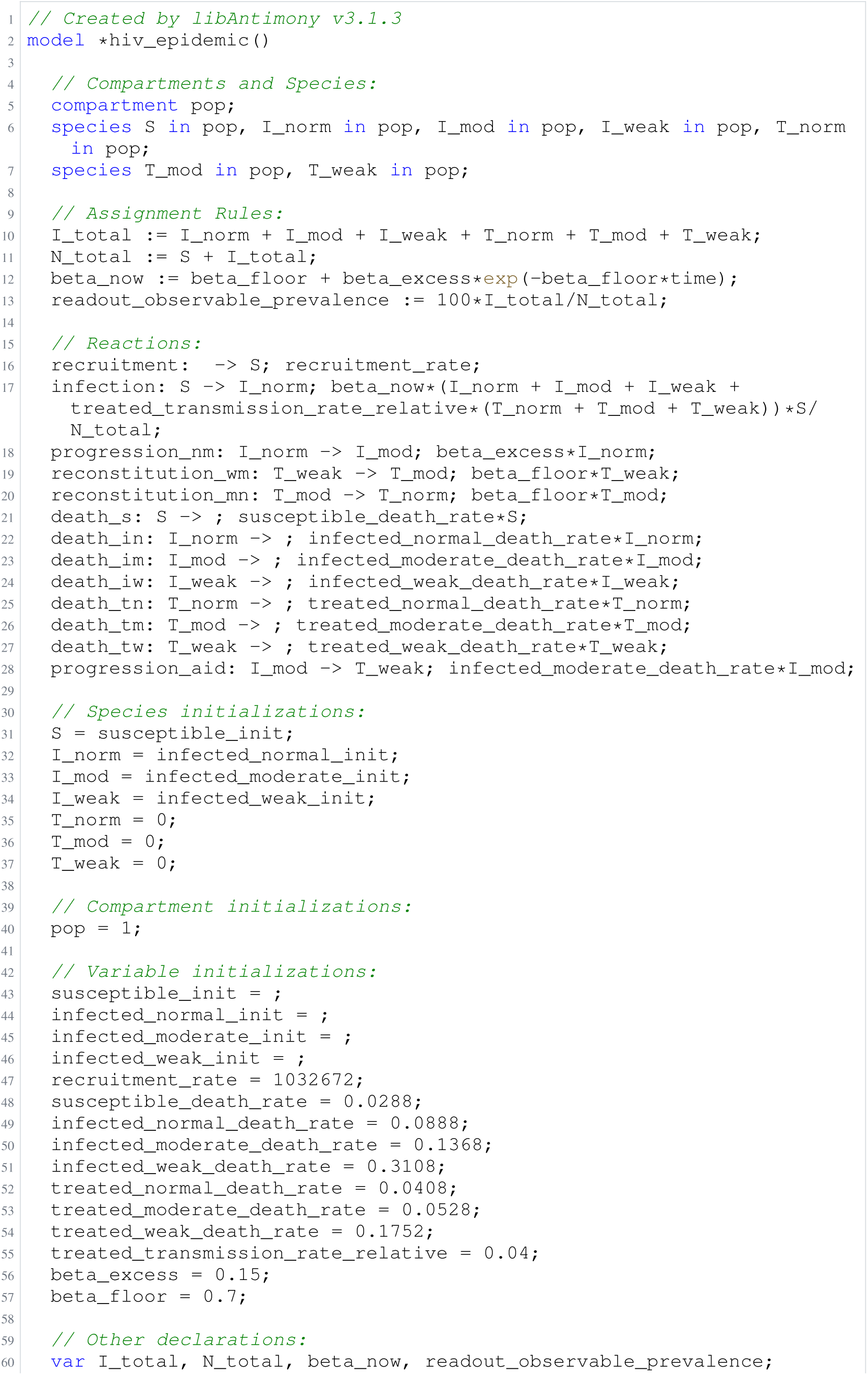
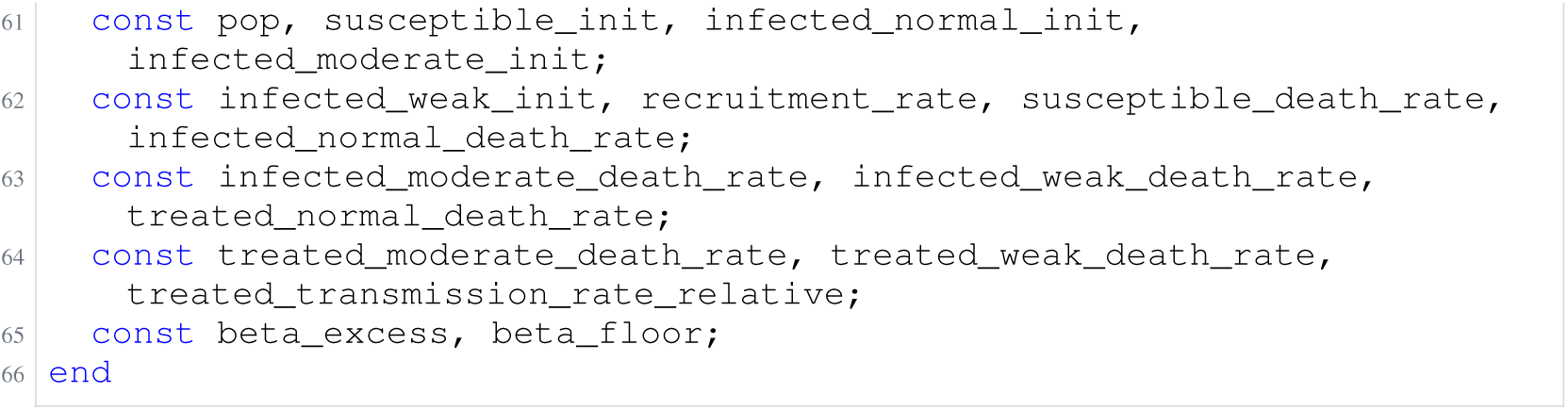
Archived Antimony for rahman_4; task 56836848_43; selected version 4.

**Listing 43:**
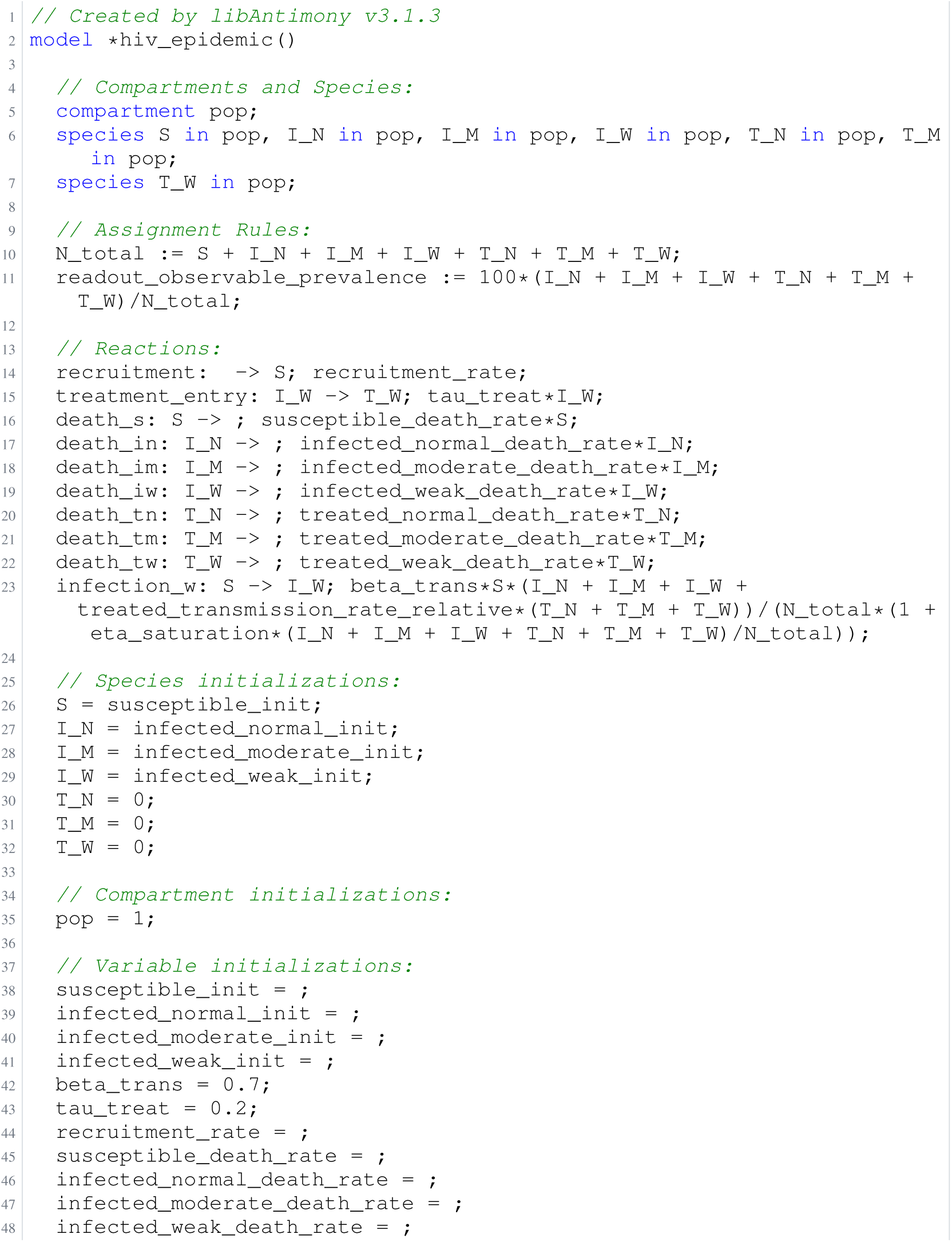
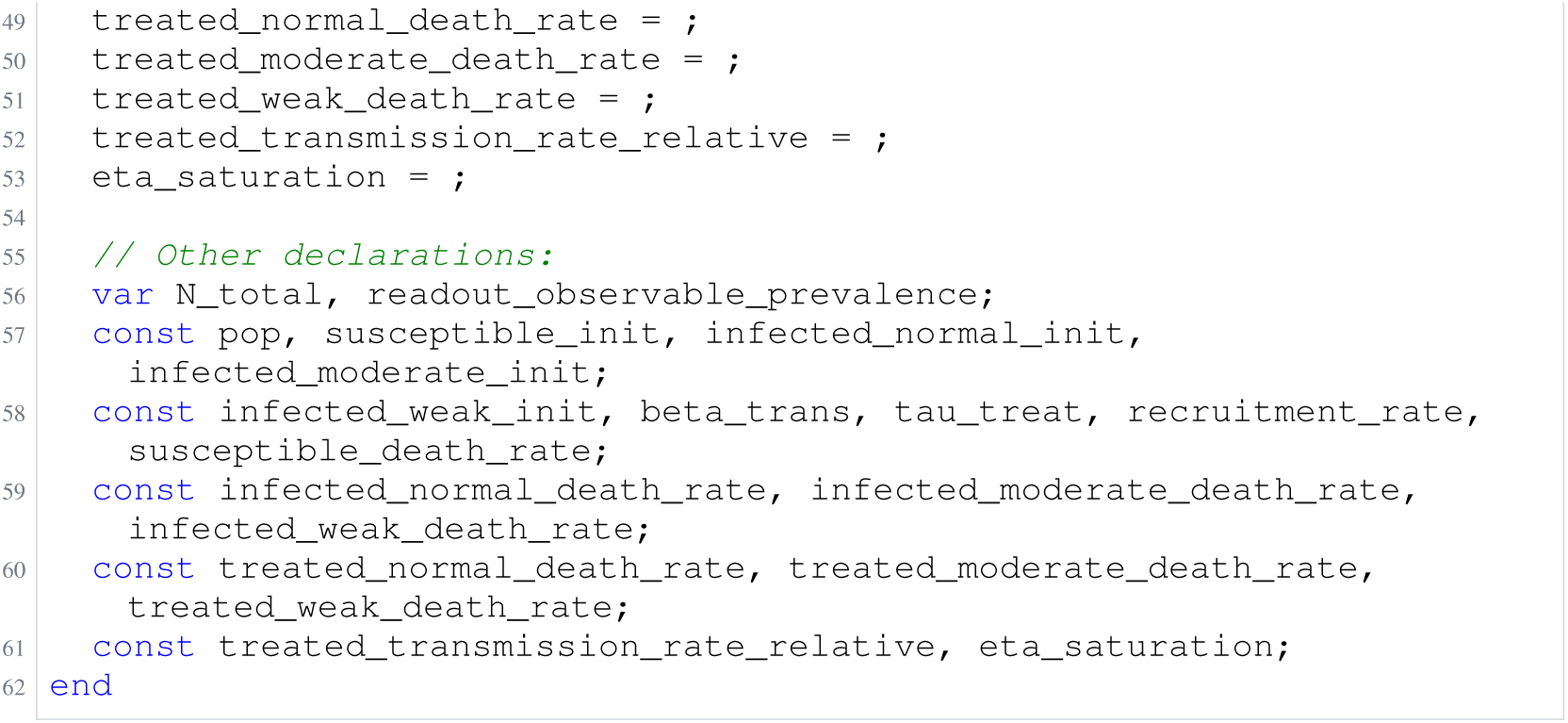
Archived Antimony for rahman_5; task 56836848_44; selected version 9.

### H.4 Brannmark

**Listing 44:**
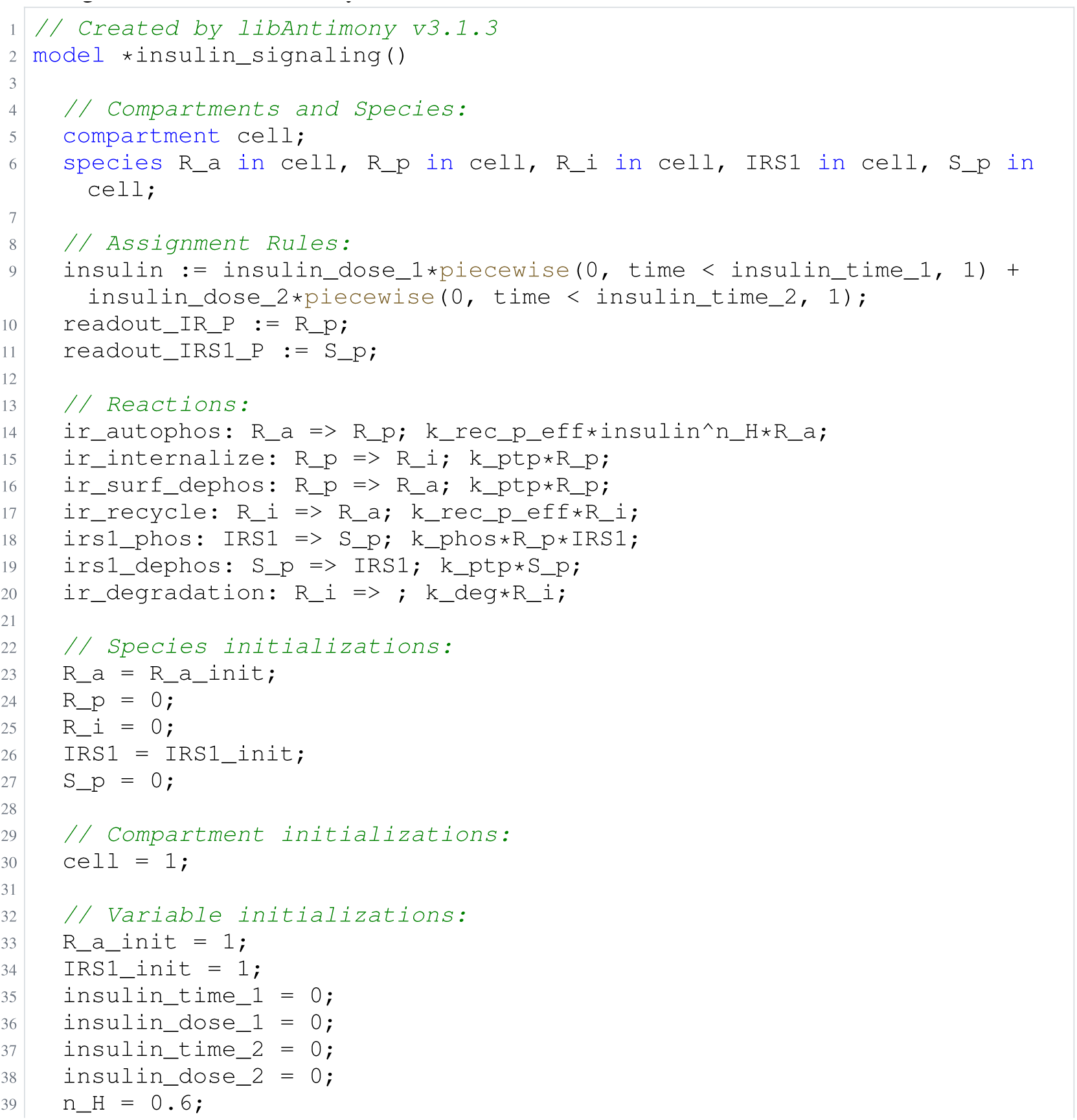
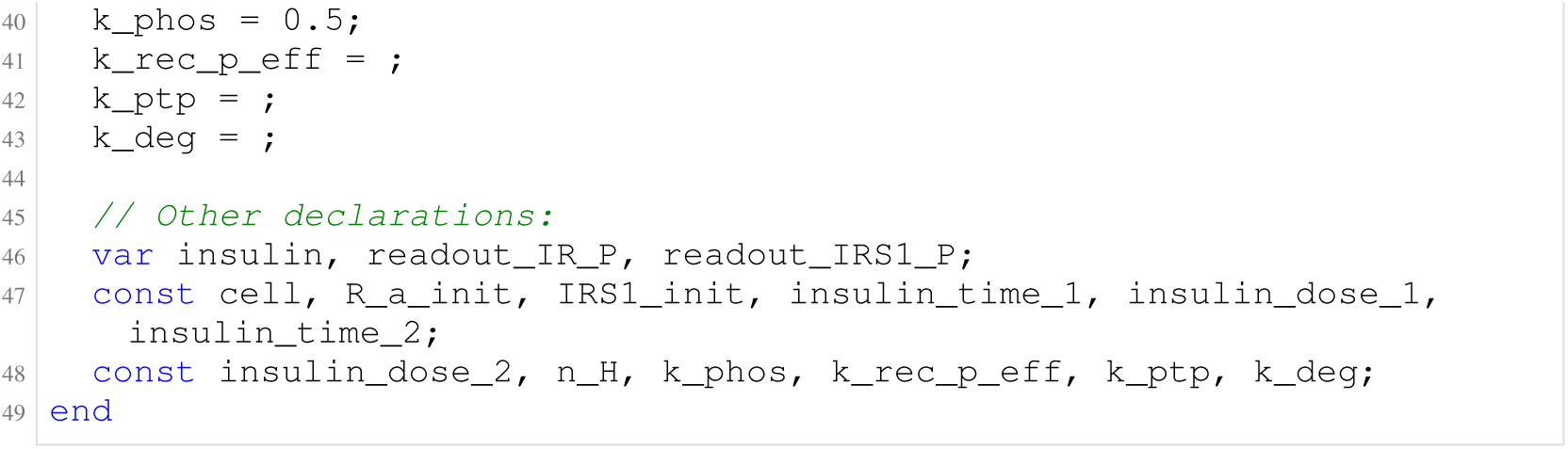
Archived Antimony for brannmark_1; task 56836848_15; selected version 17.

**Listing 45:**
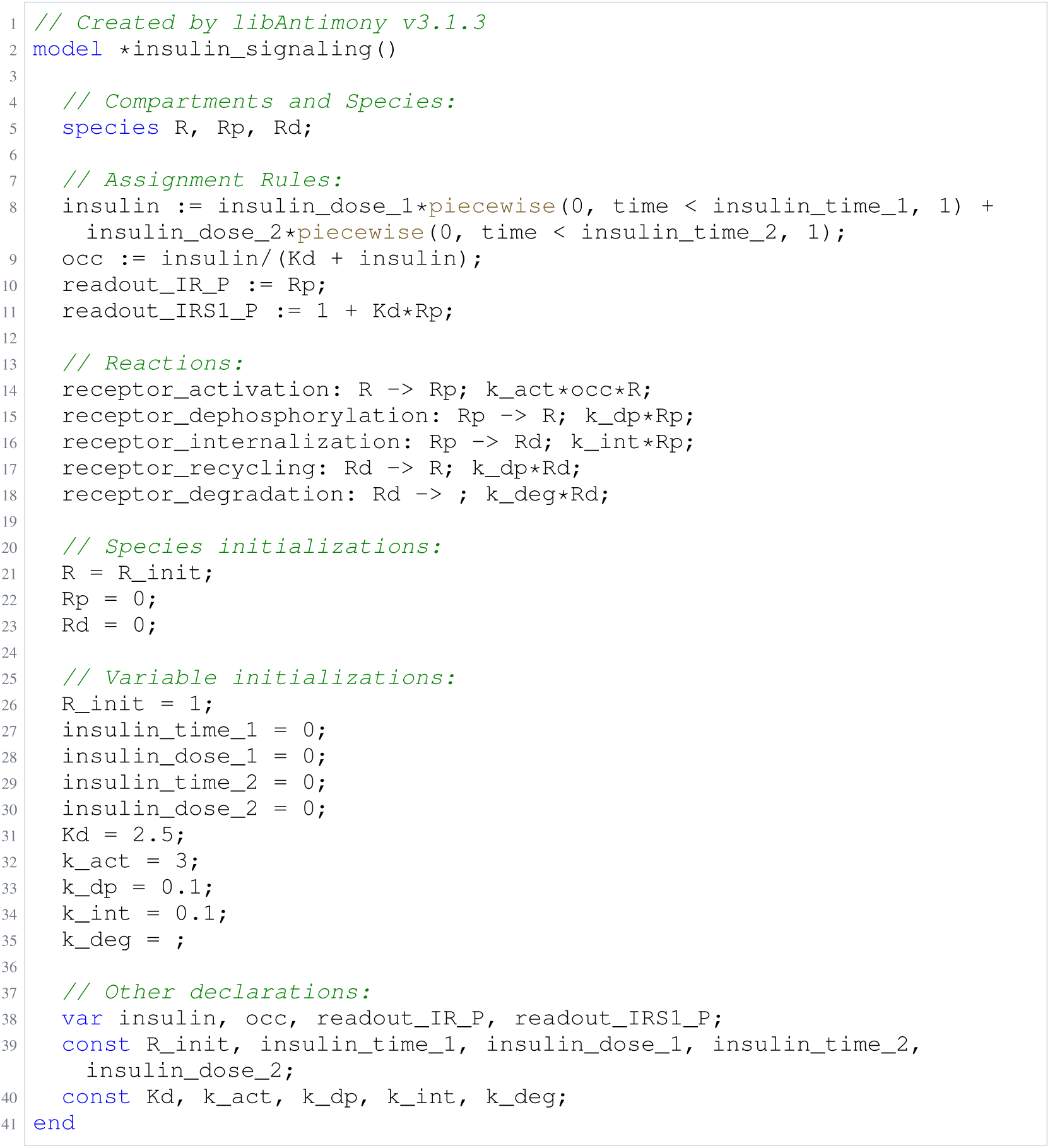
Archived Antimony for brannmark_2; task 56836848_16; selected version 12.

**Listing 46:**
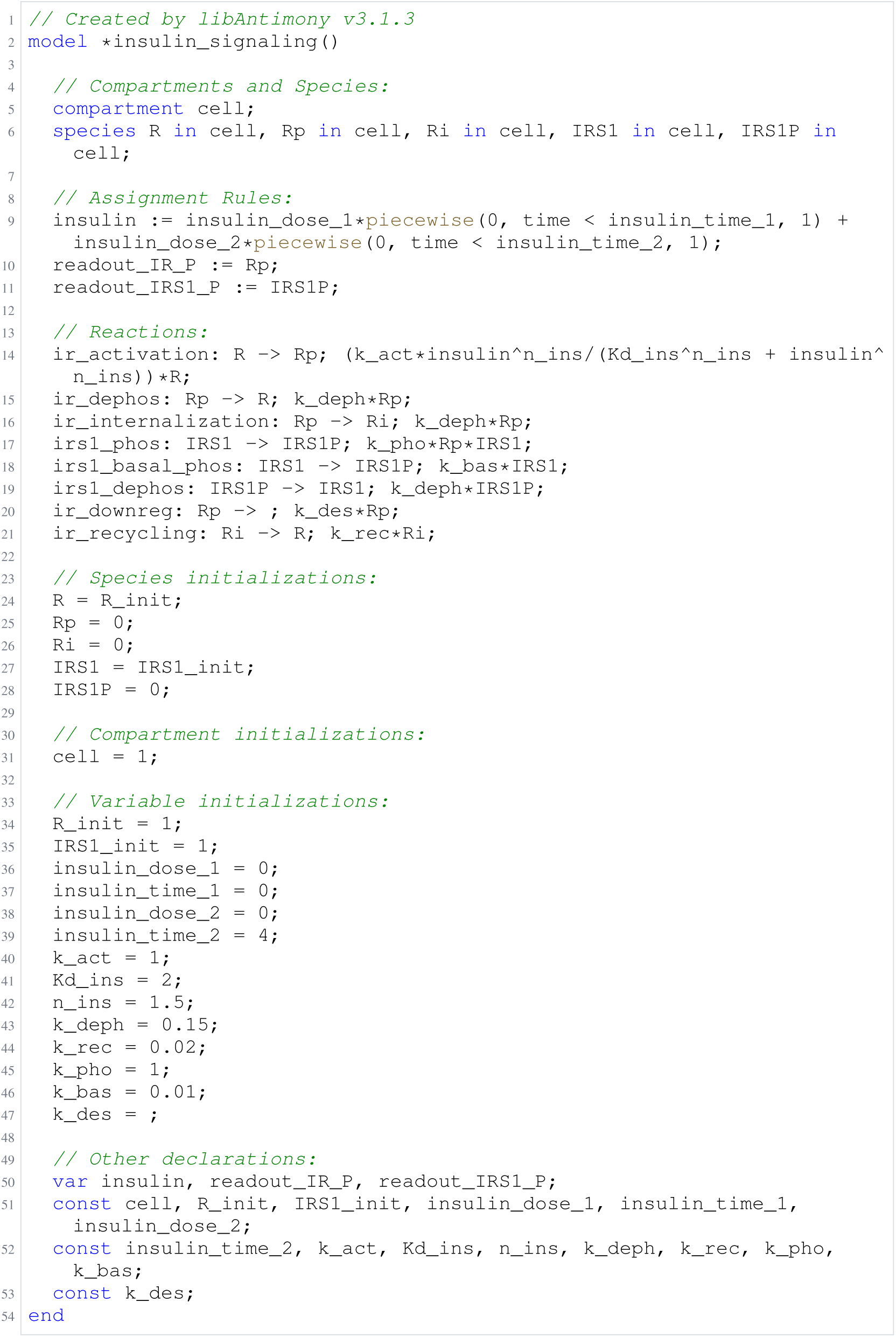
Archived Antimony for brannmark_3; task 56836848_17; selected version 6.

**Listing 47:**
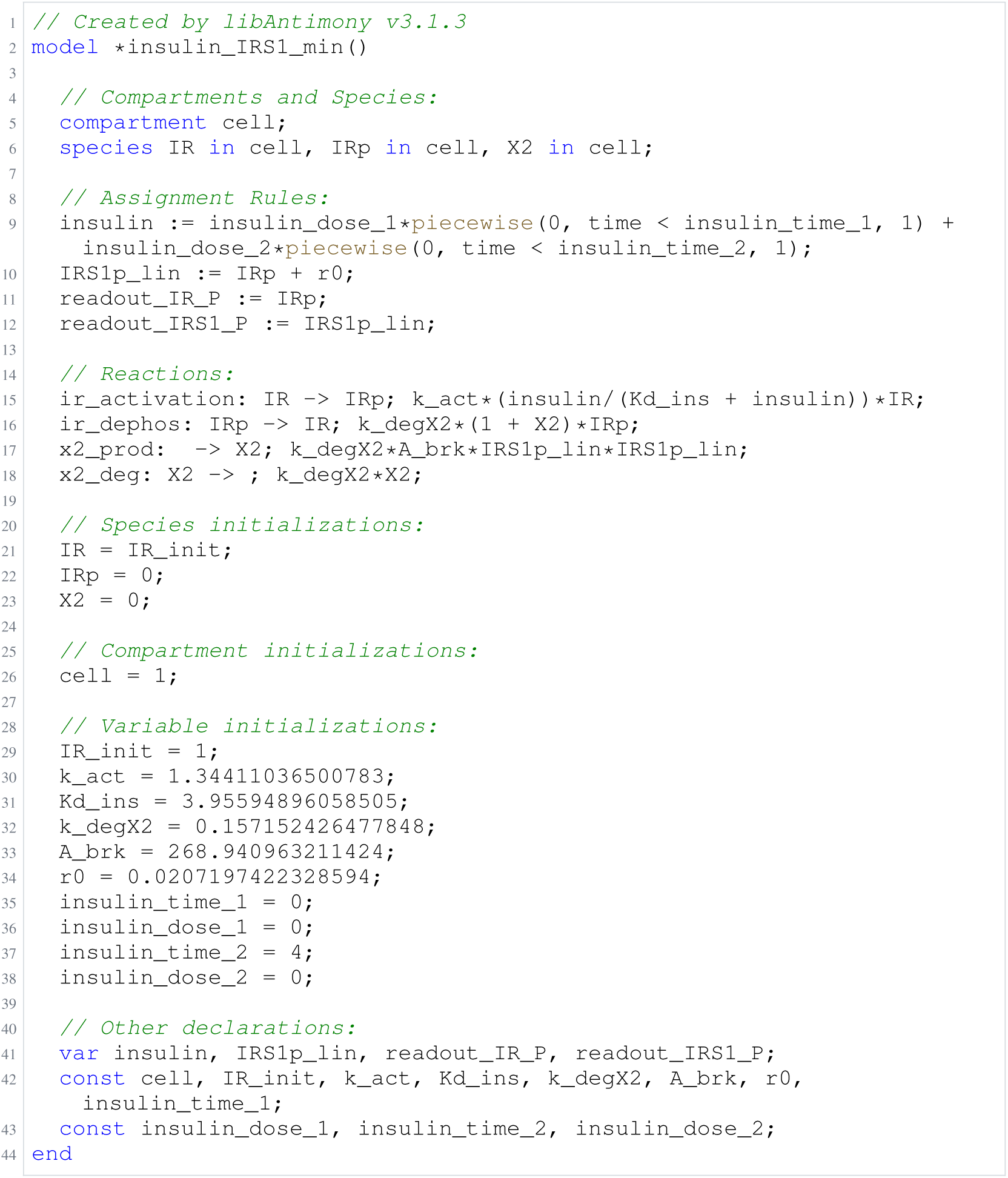
Archived Antimony for brannmark_4; task 56836848_18; selected version 12.

**Listing 48:**
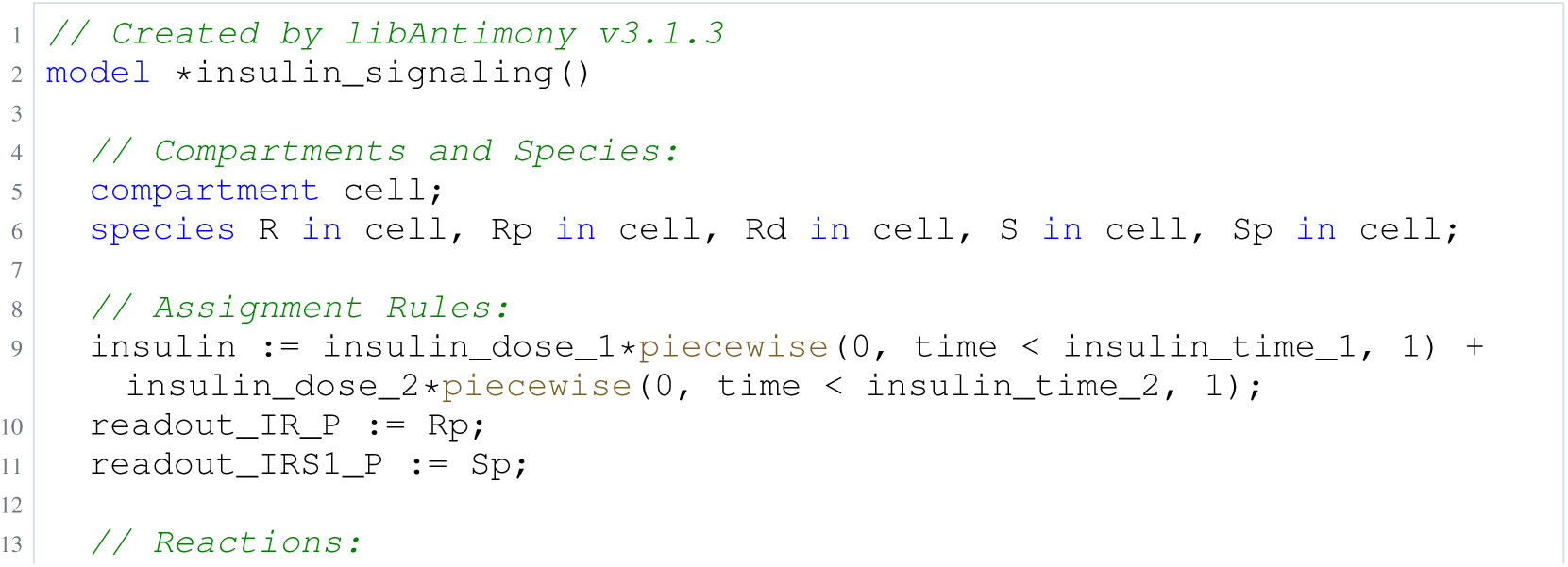
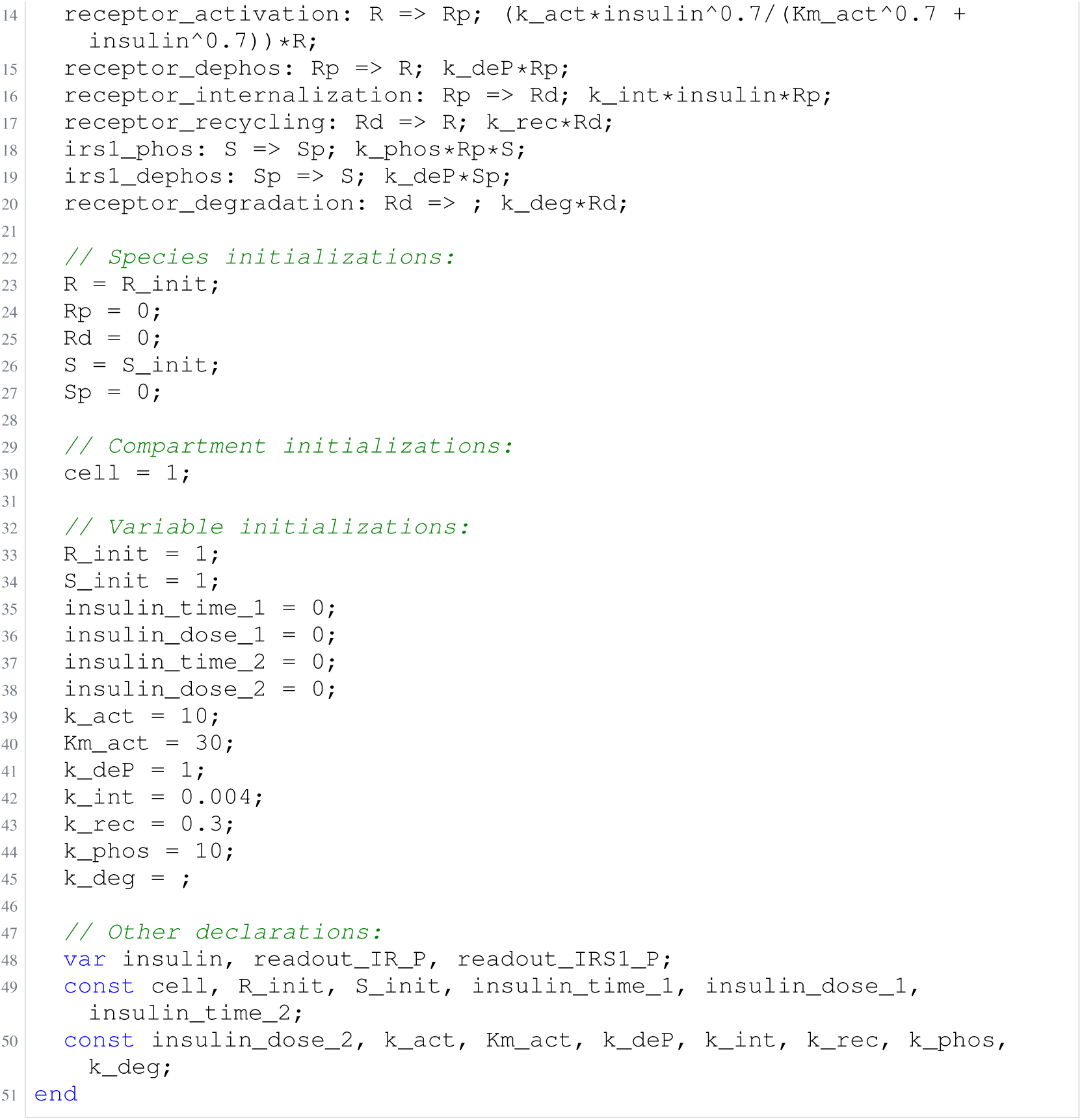
Archived Antimony for brannmark_5; task 56836848_19; selected version 5.

### H.5 Boehm

**Listing 49:**
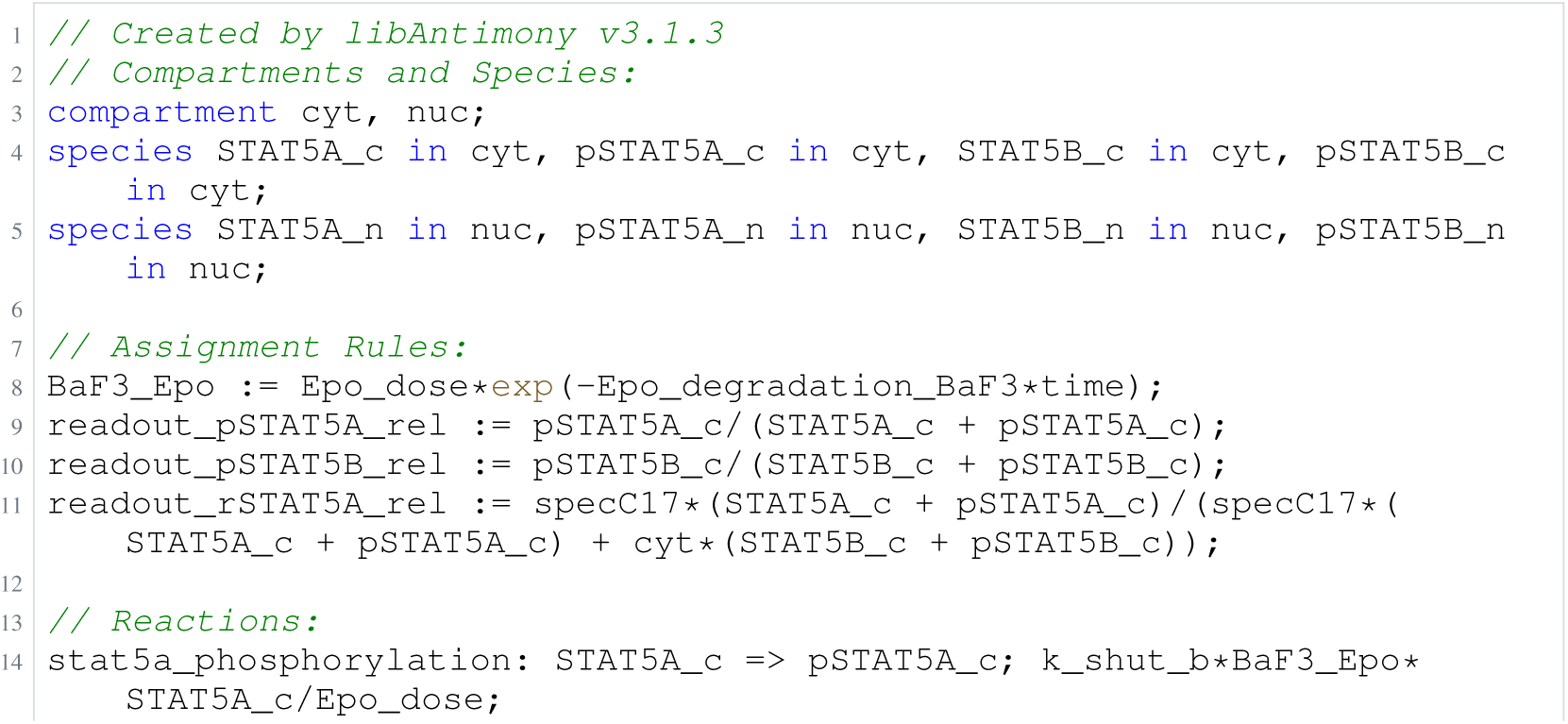
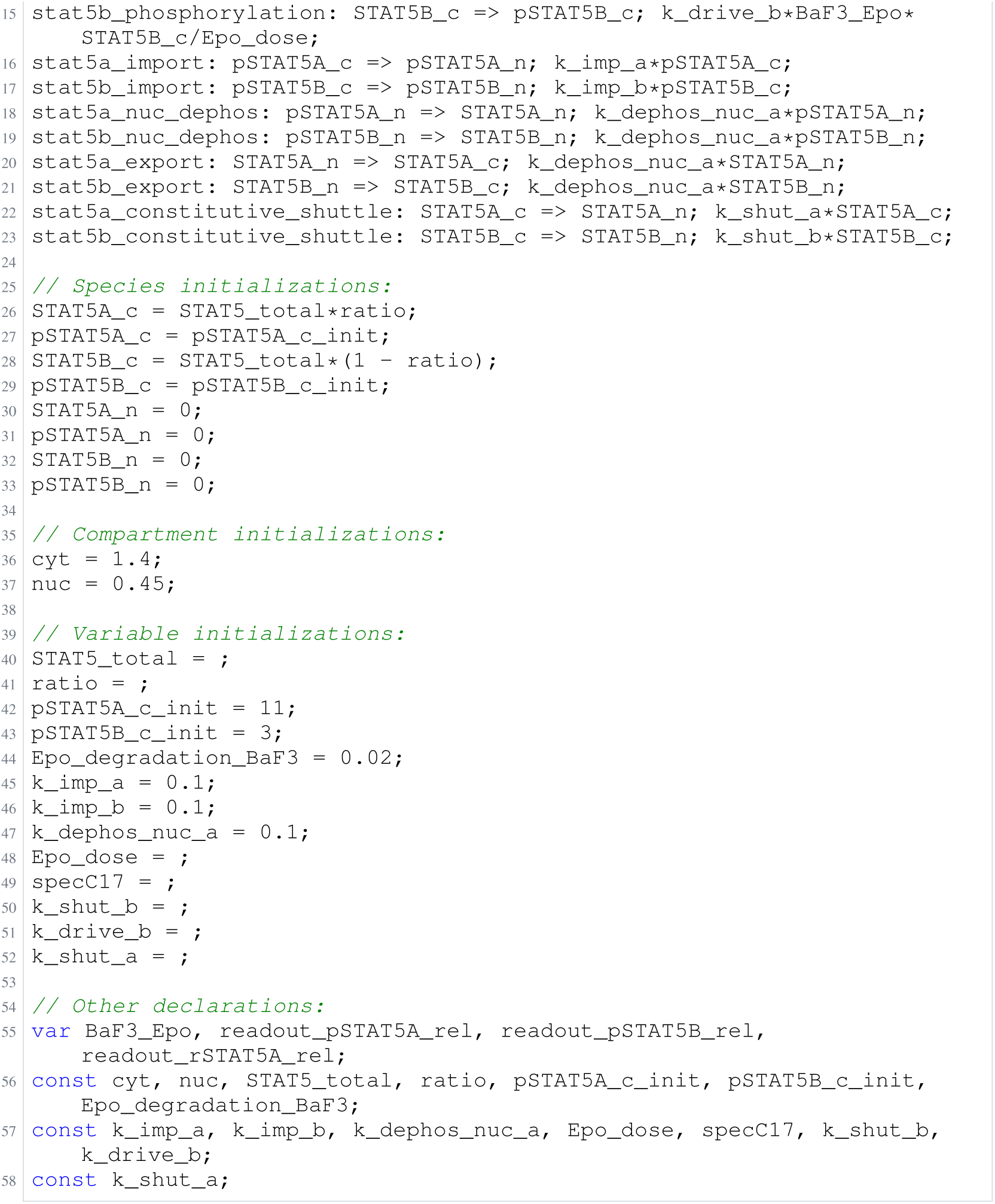
Archived Antimony for boehm_1; task 56836848_10; selected version 13.

**Listing 50:**
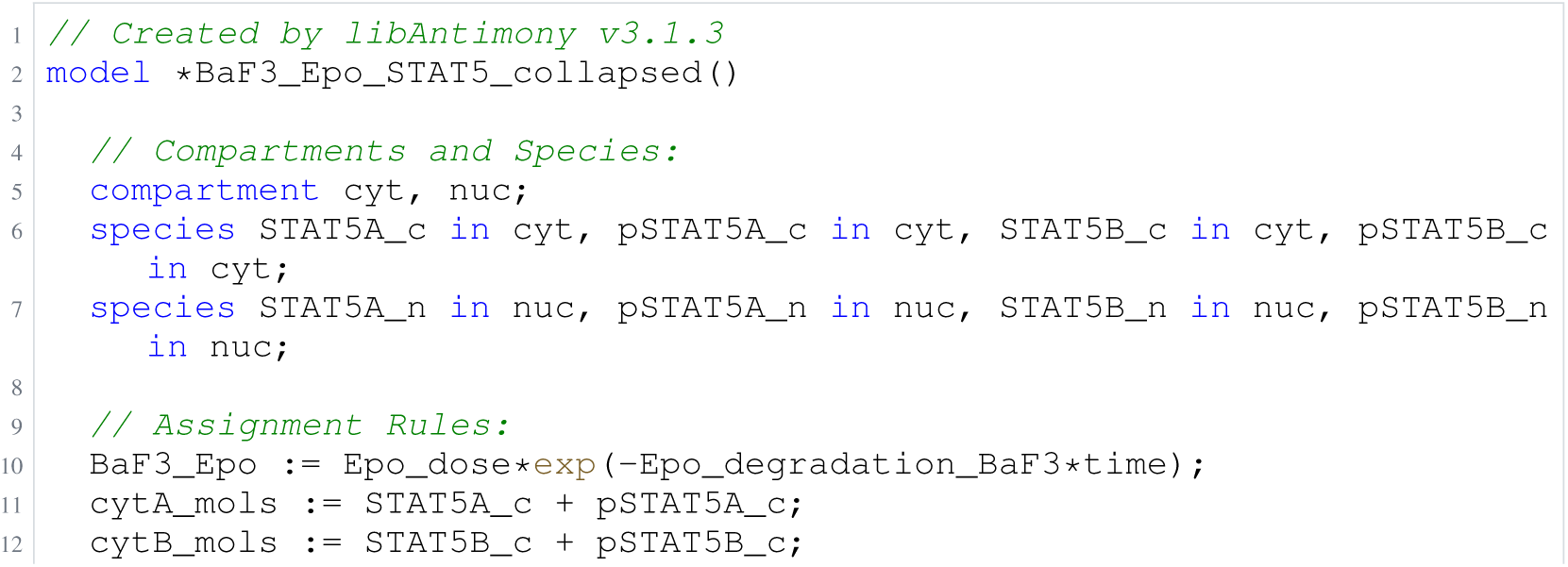
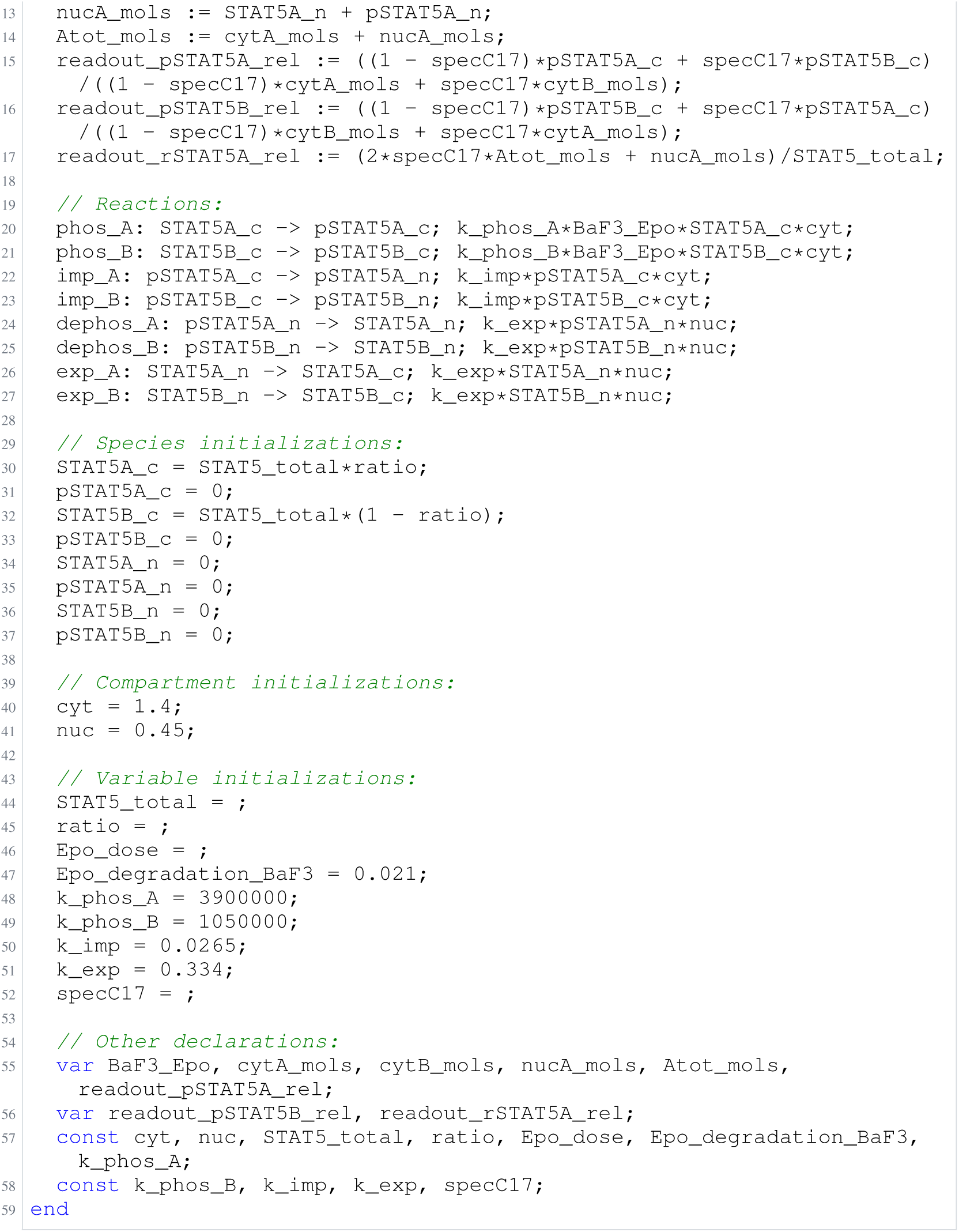
Archived Antimony for boehm_2; task 56836848_11; selected version 3.

**Listing 51:**
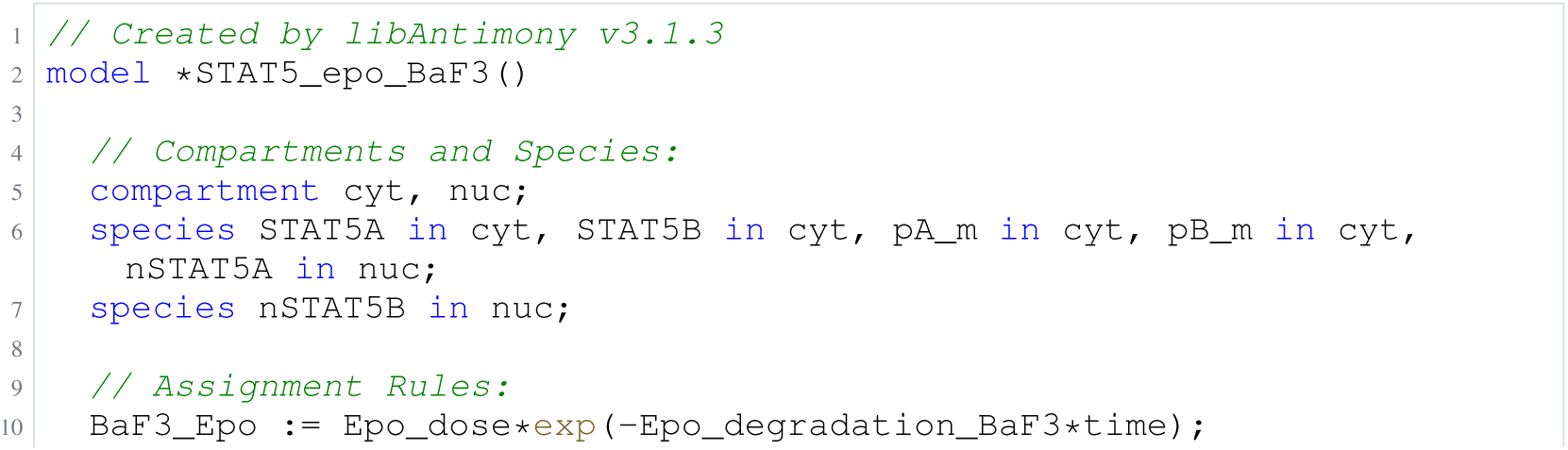
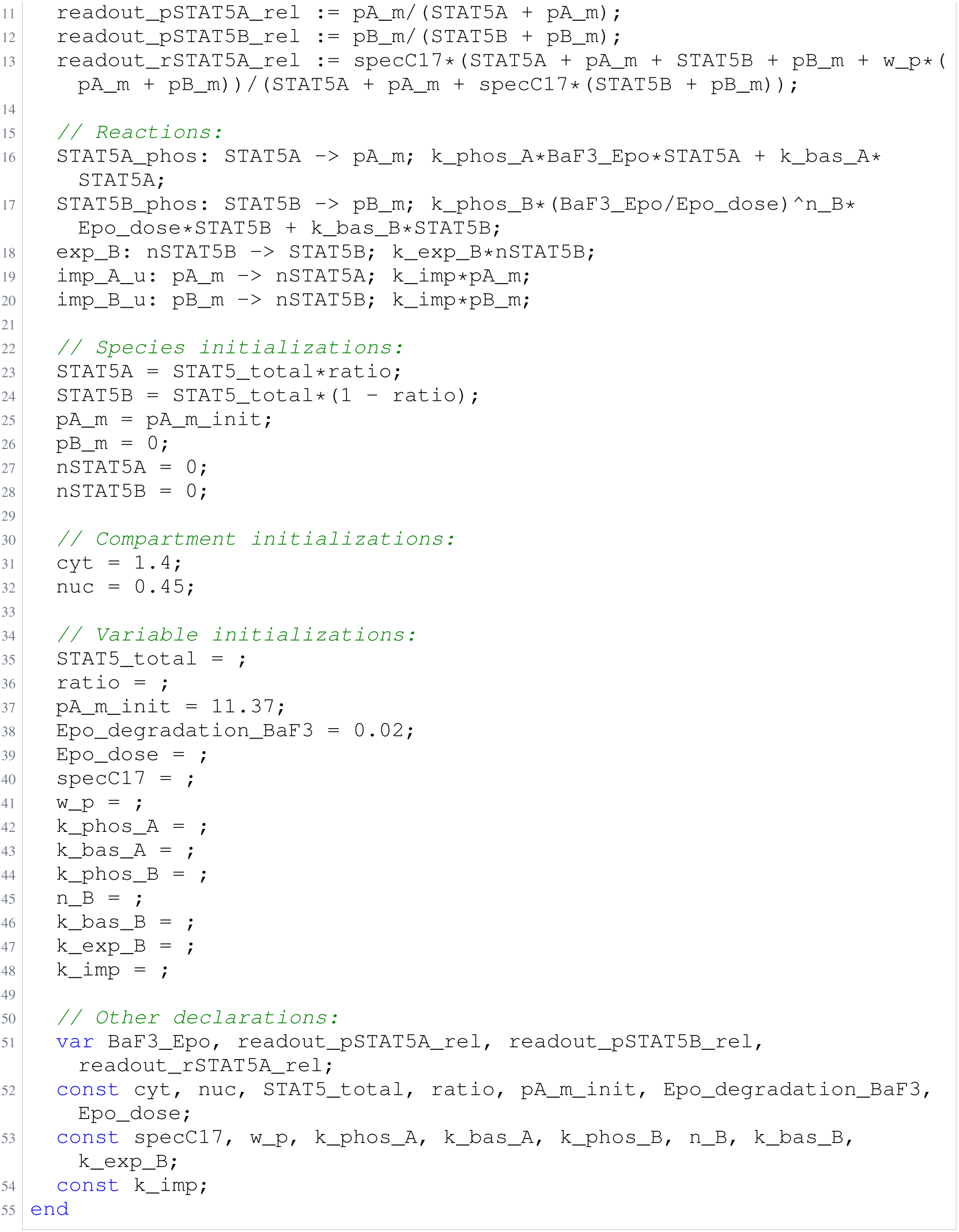
Archived Antimony for boehm_3; task 56836848_12; selected version 4.

**Listing 52:**
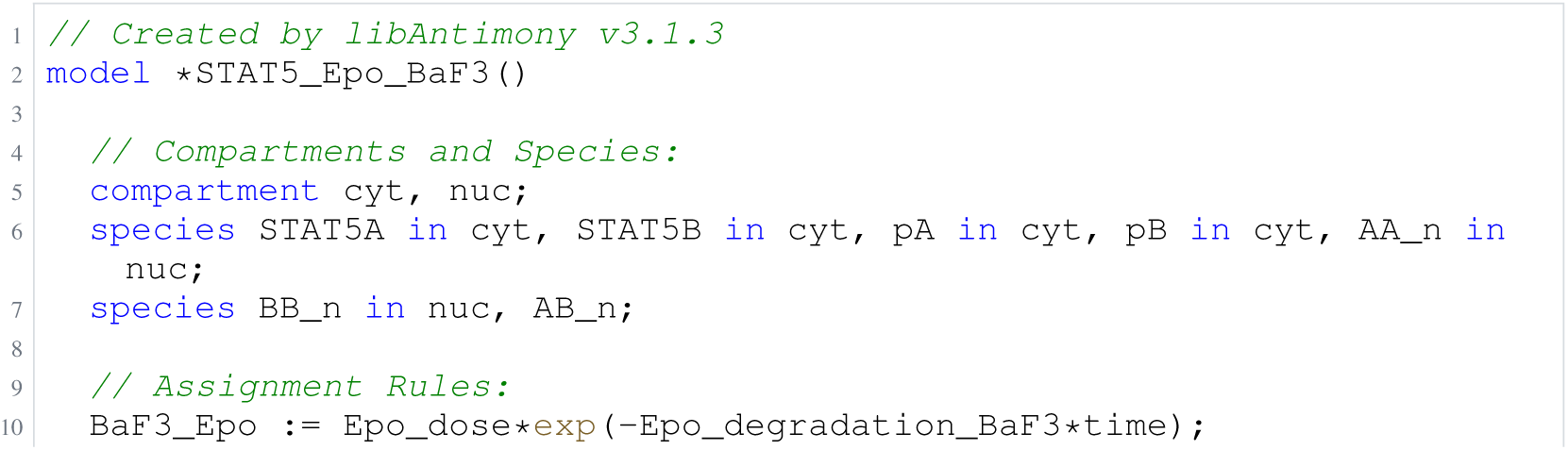
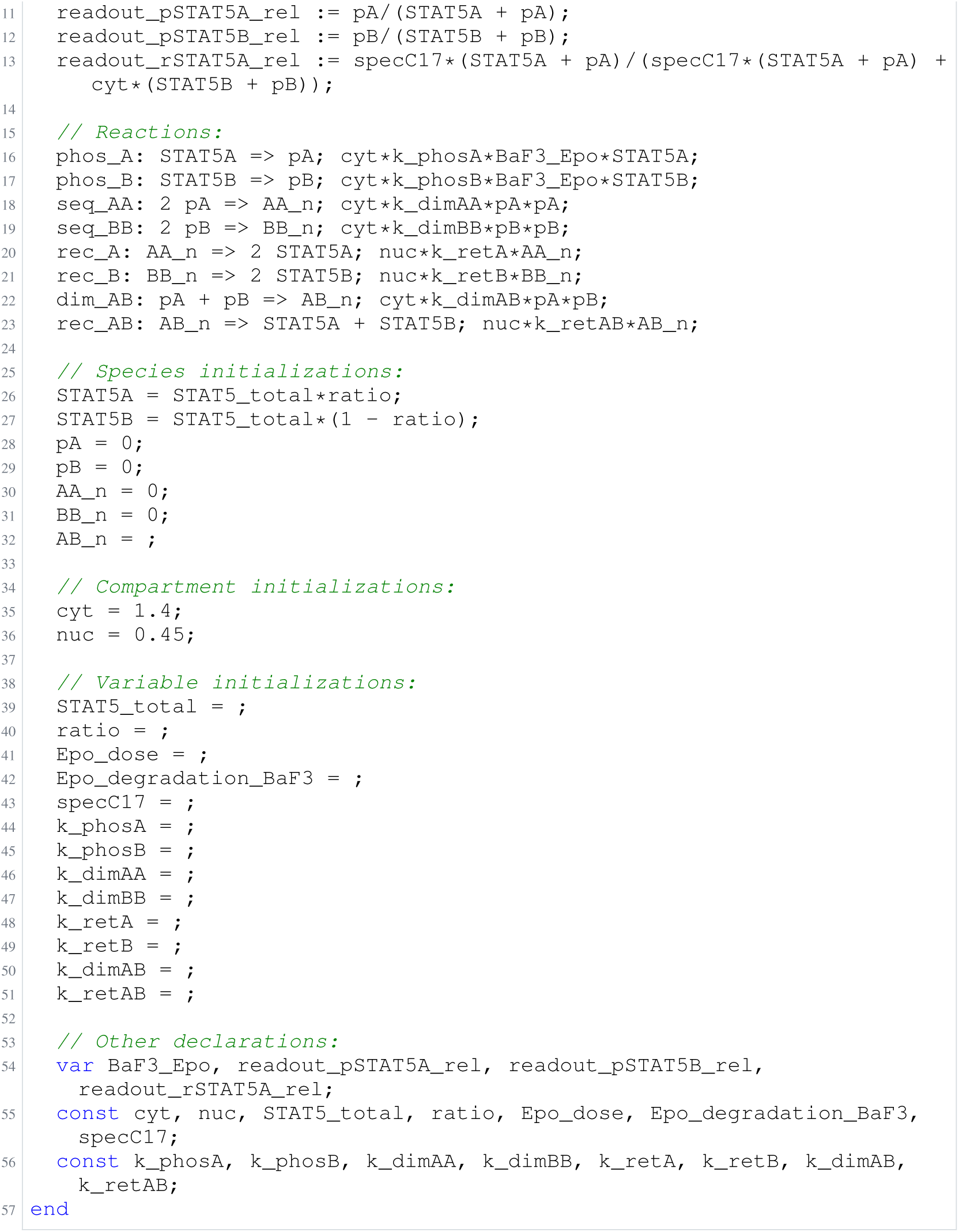
Archived Antimony for boehm_4; task 56836848_13; selected version 5.

**Listing 53:**
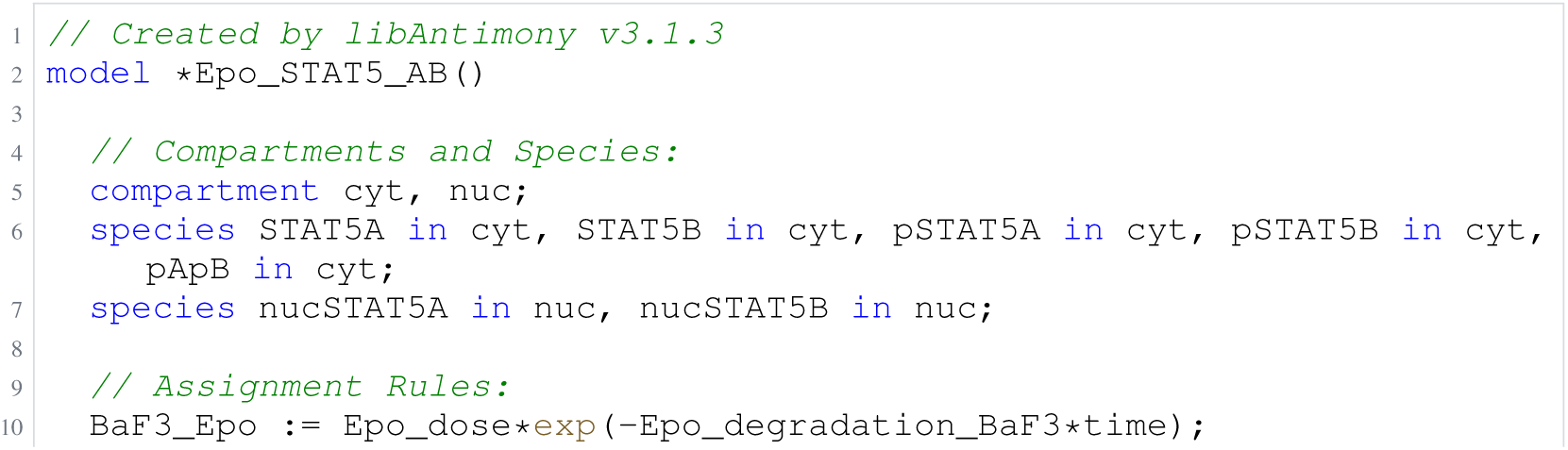
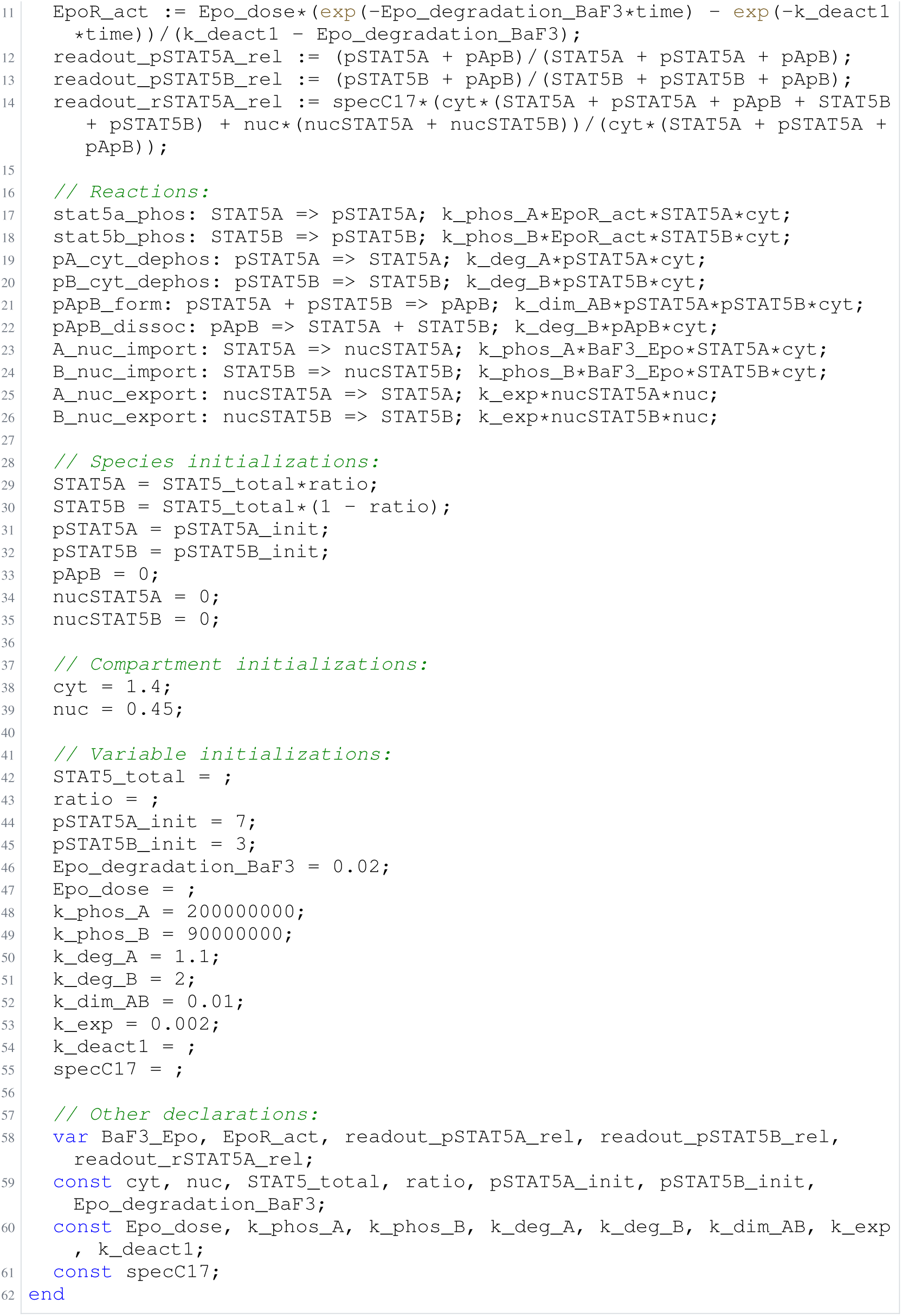
Archived Antimony for boehm_5; task 56836848_14; selected version 12.

### H.6 Armistead

**Listing 54:**
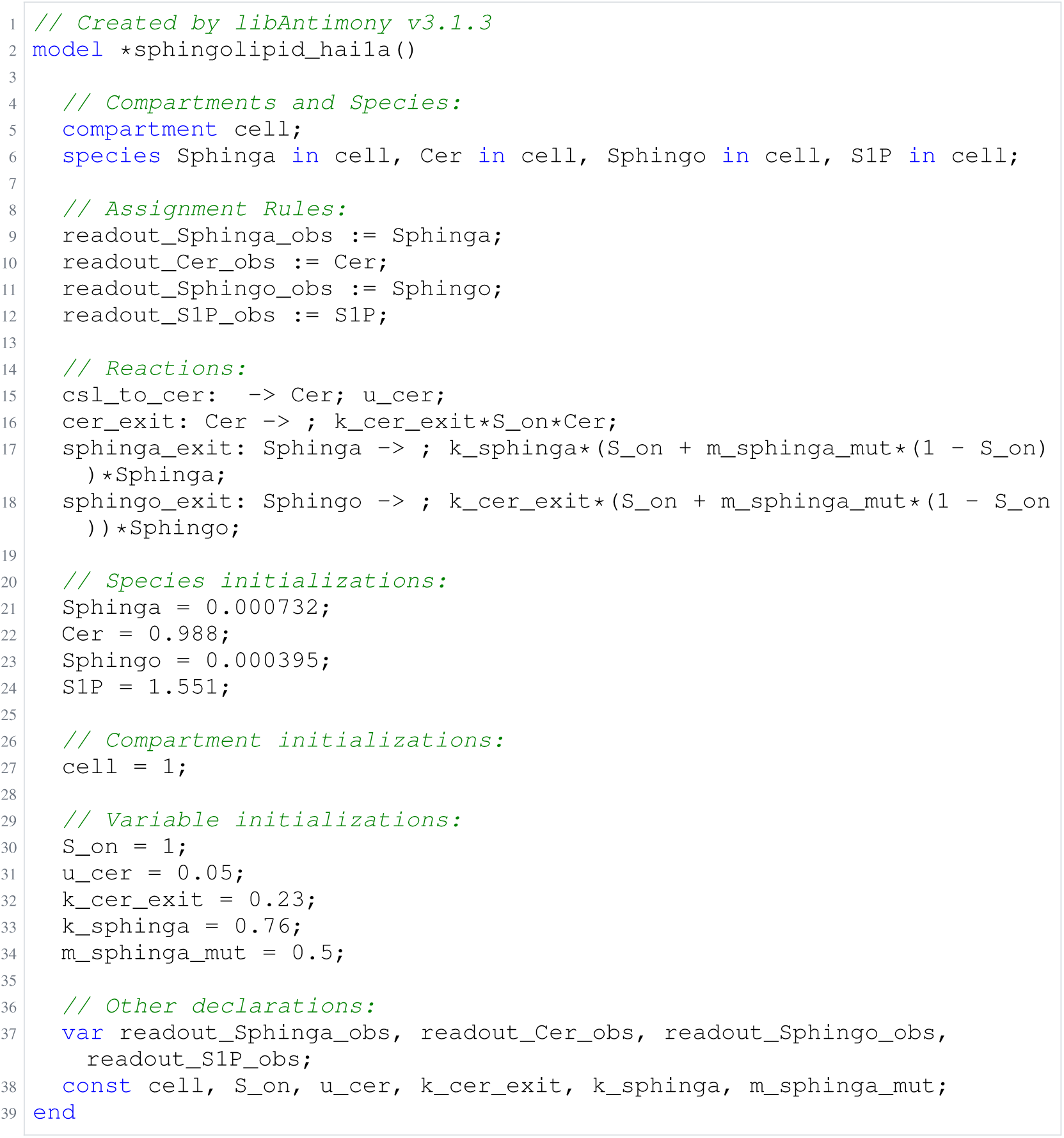
Archived Antimony for armistead_1; task 56836848_0; selected version 2.

**Listing 55:**
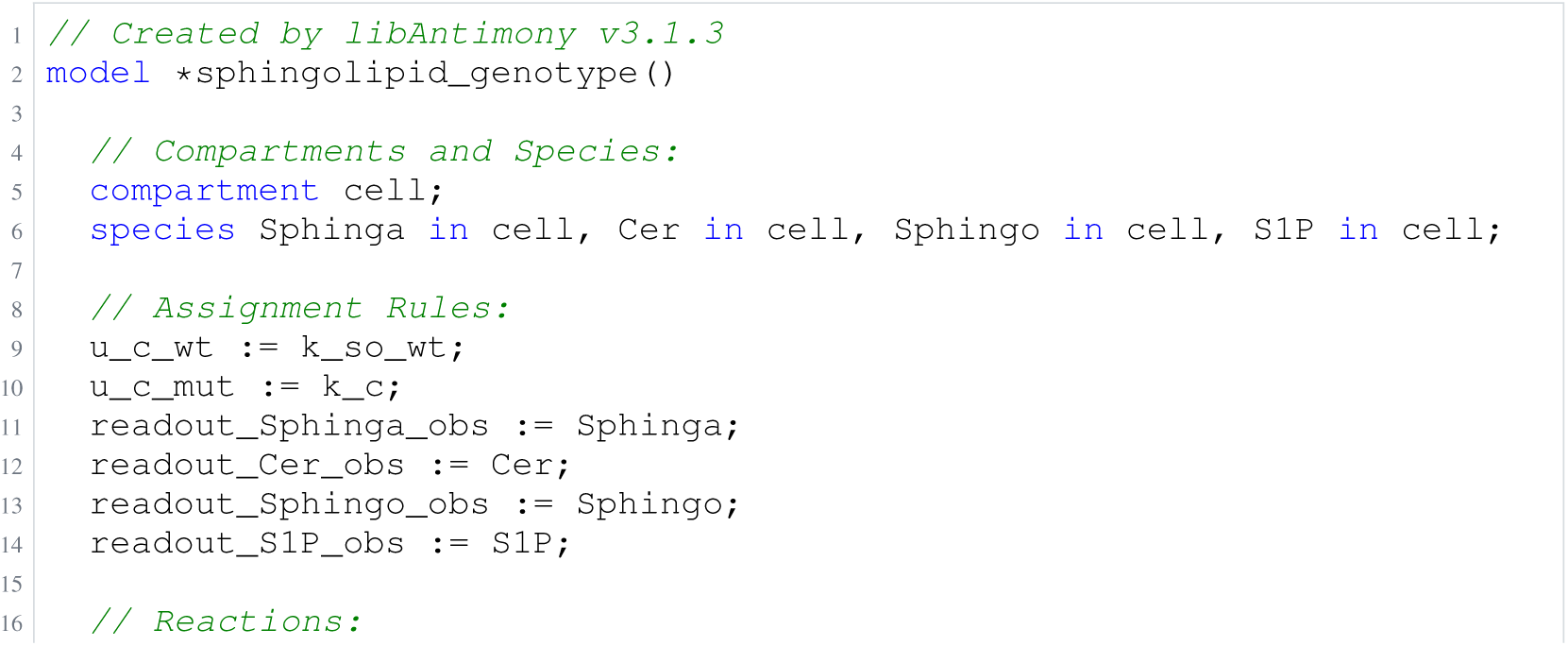
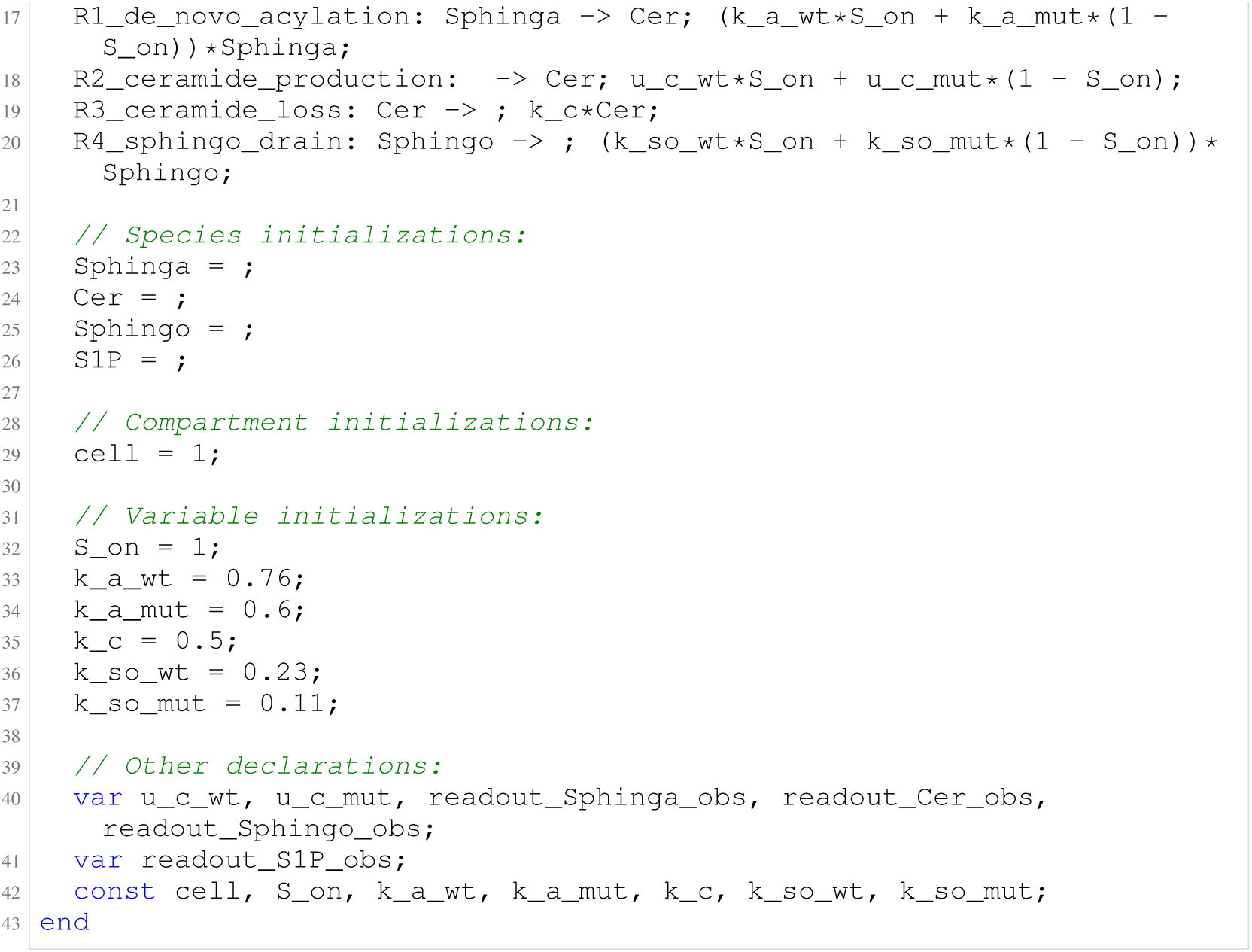
Archived Antimony for armistead_2; task 56836848_1; selected version 3.

**Listing 56:**
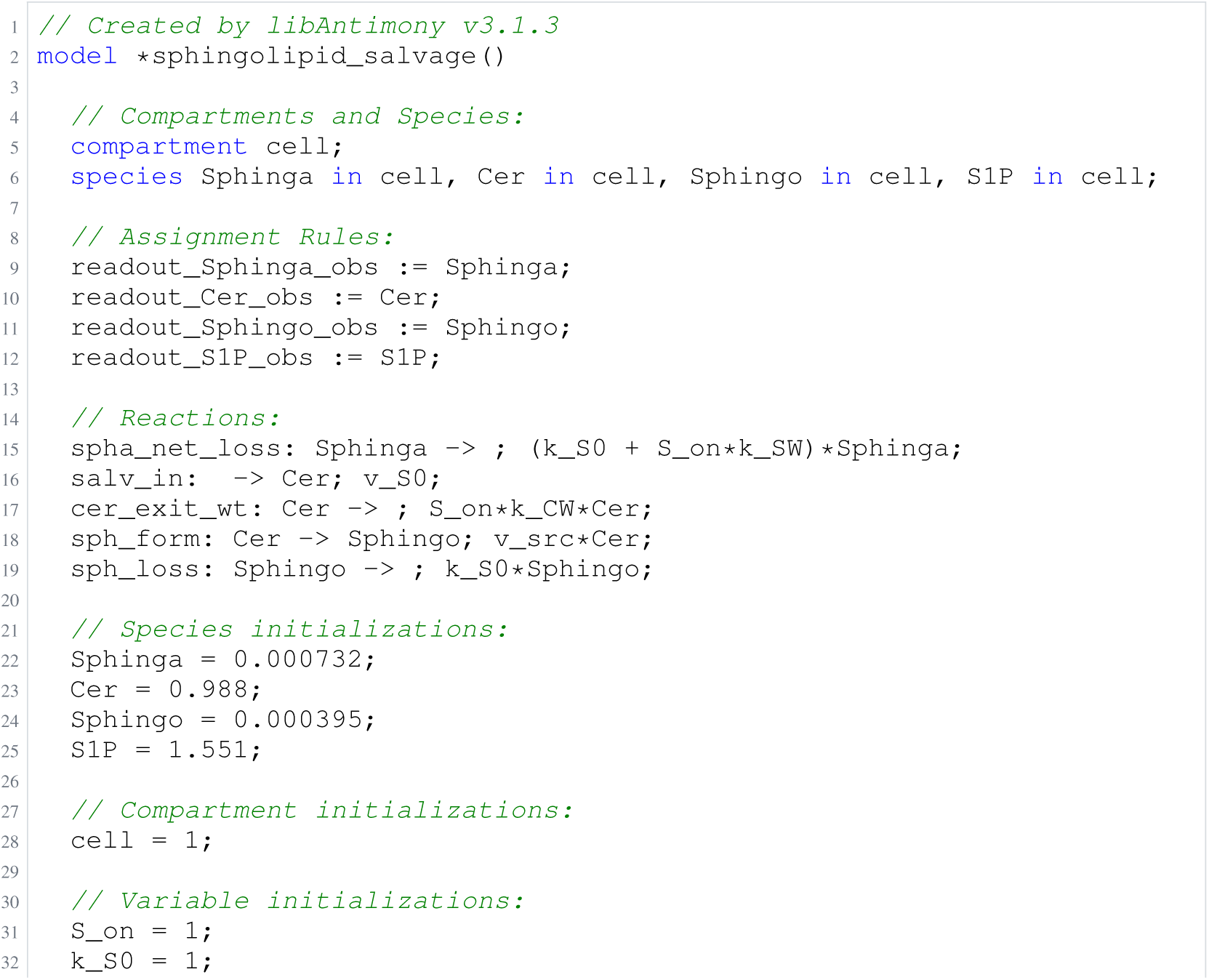
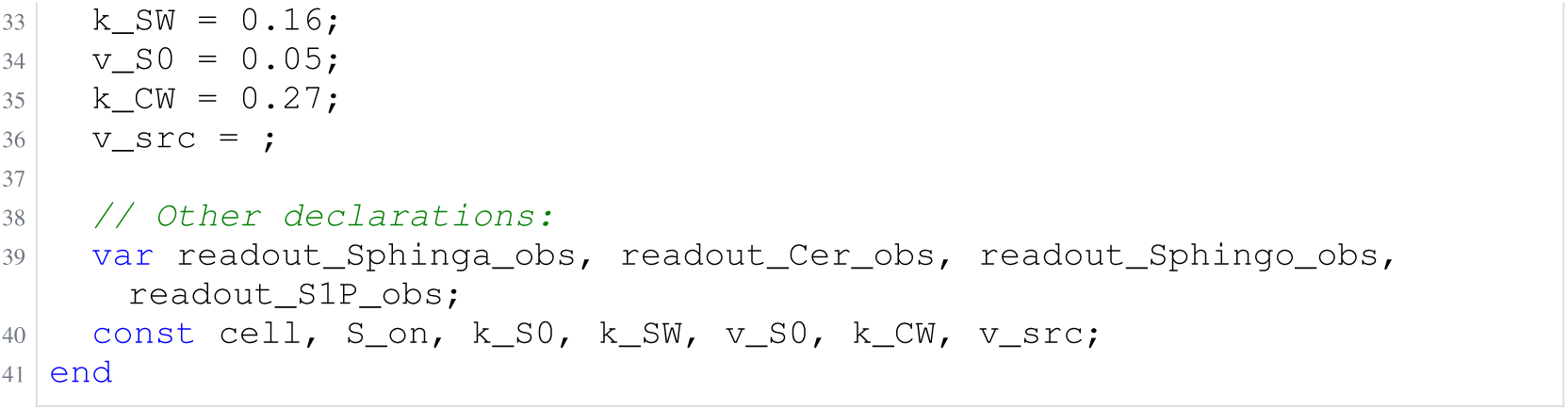
Archived Antimony for armistead_3; task 56836848_2; selected version 5.

**Listing 57:**
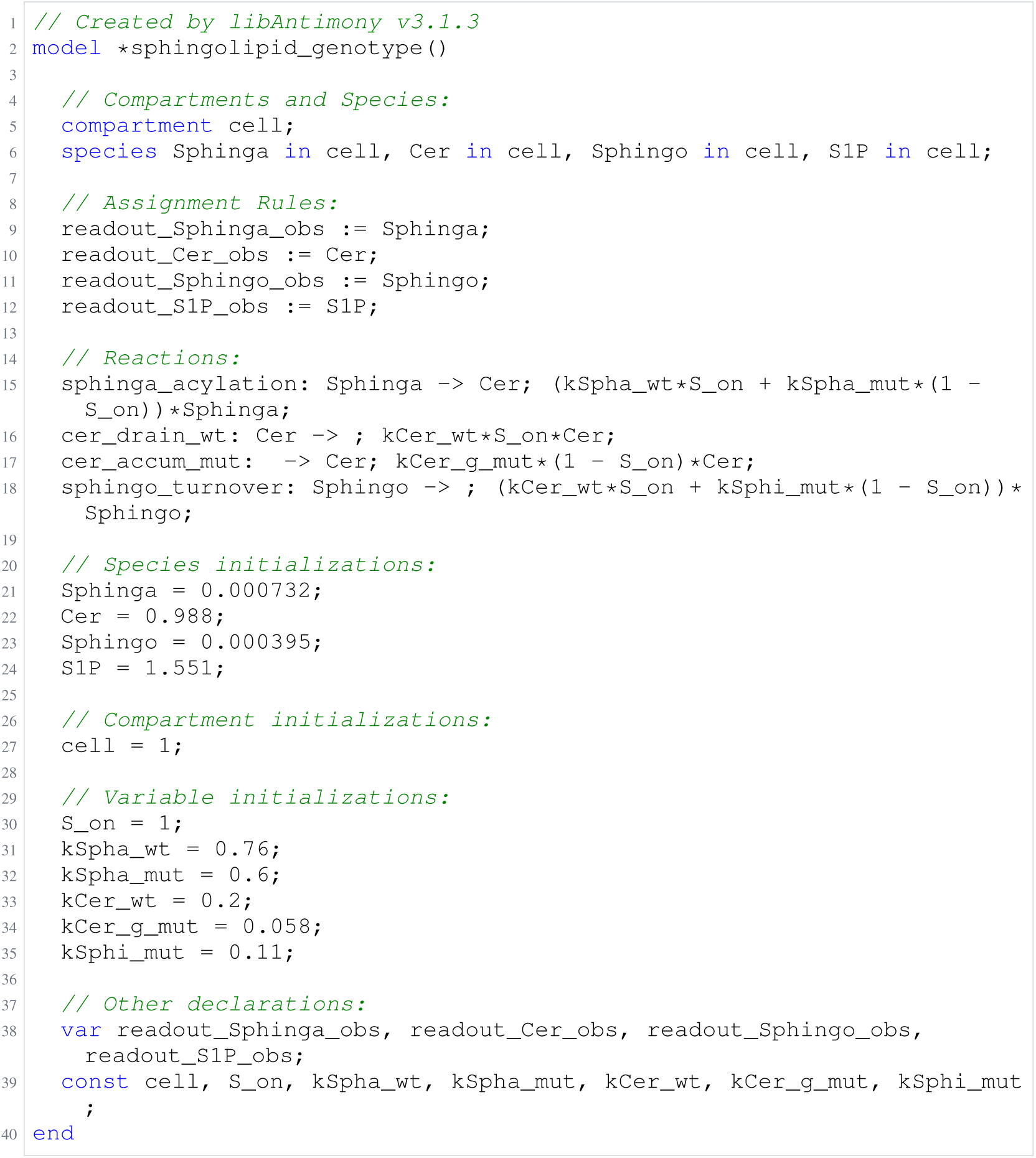
Archived Antimony for armistead_4; task 56836848_3; selected version 1.

**Listing 58:**
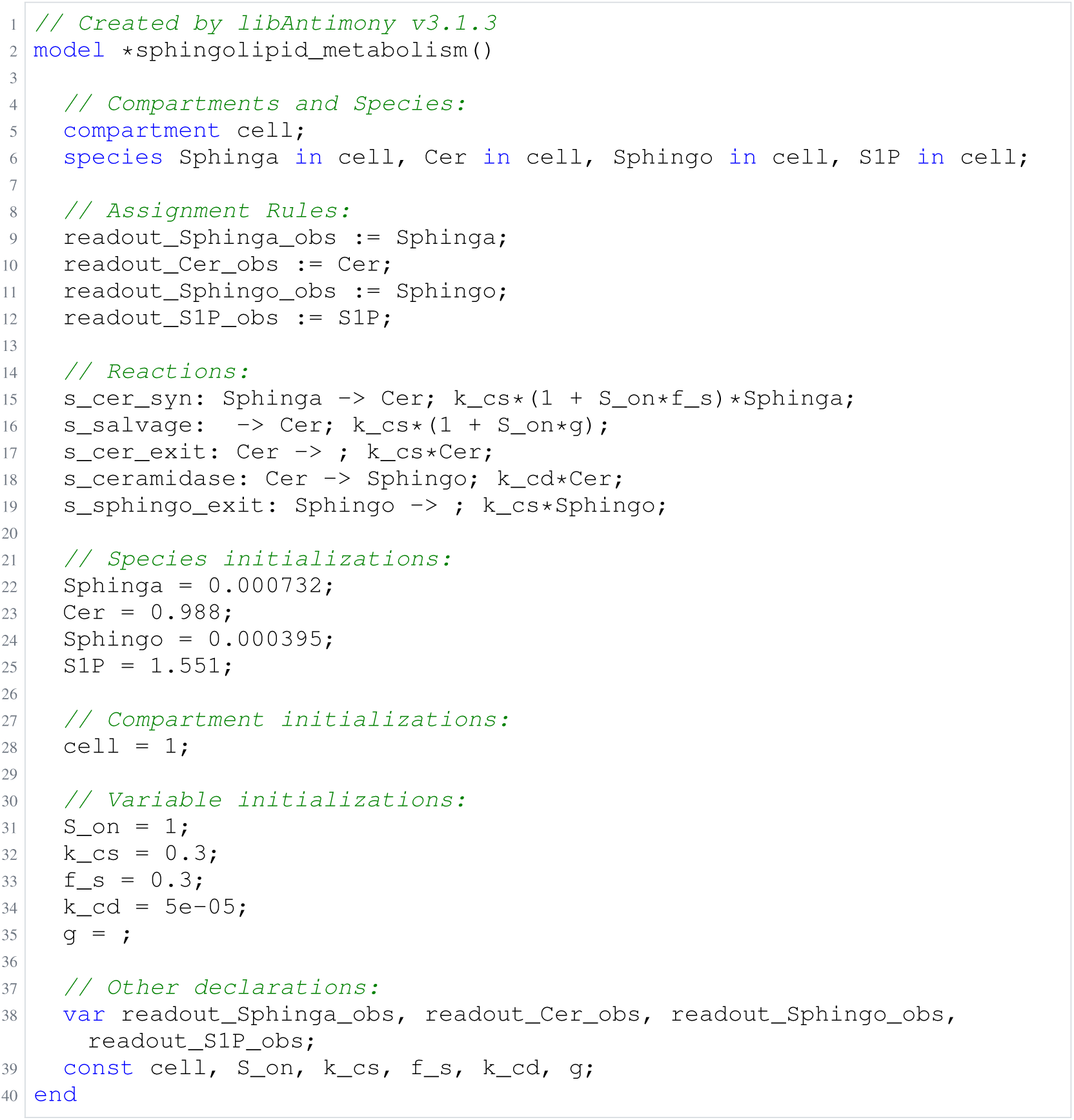
Archived Antimony for armistead_5; task 56836848_4; selected version 5.

### H.7 Fiedler

**Listing 59:**
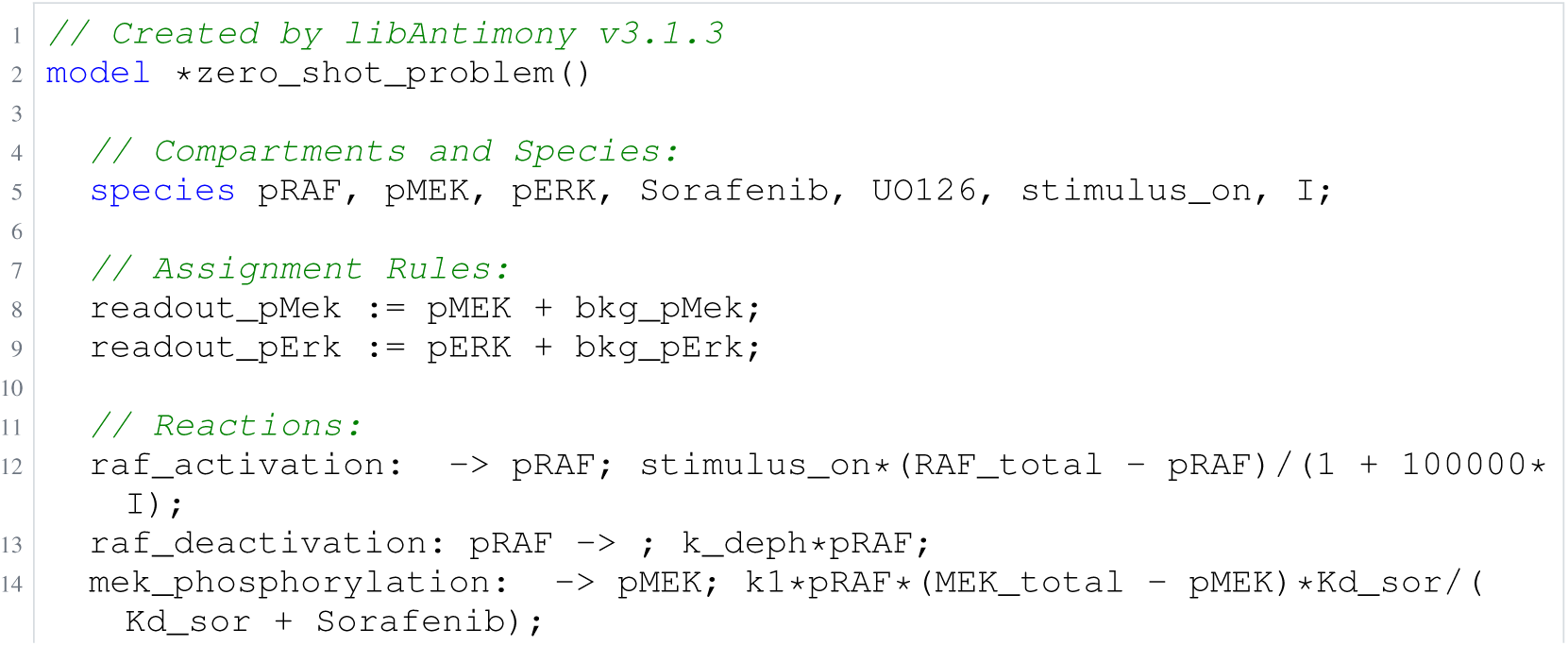
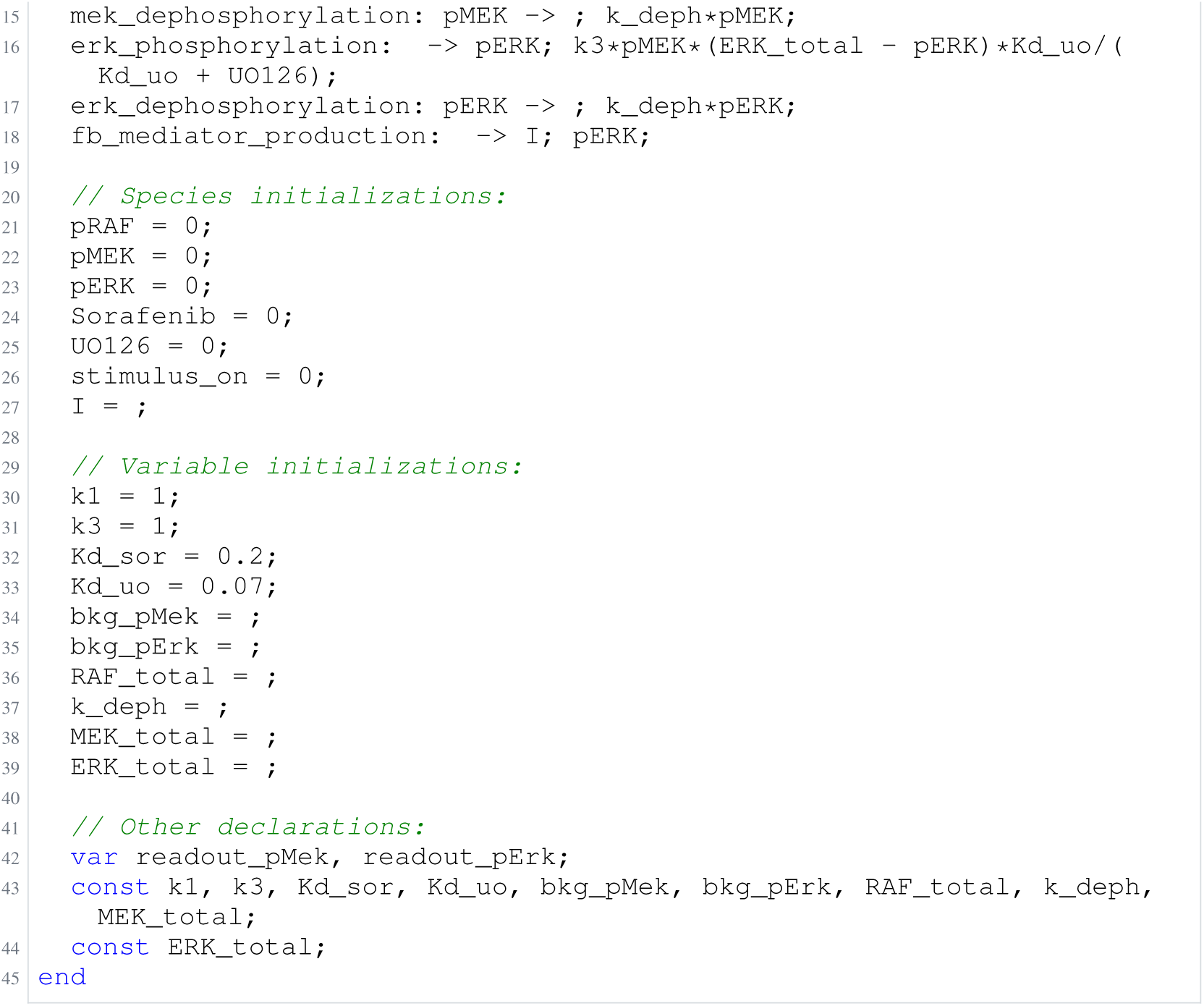
Archived Antimony for fiedler_1; task 57351389_5; selected version 21.

**Listing 60:**
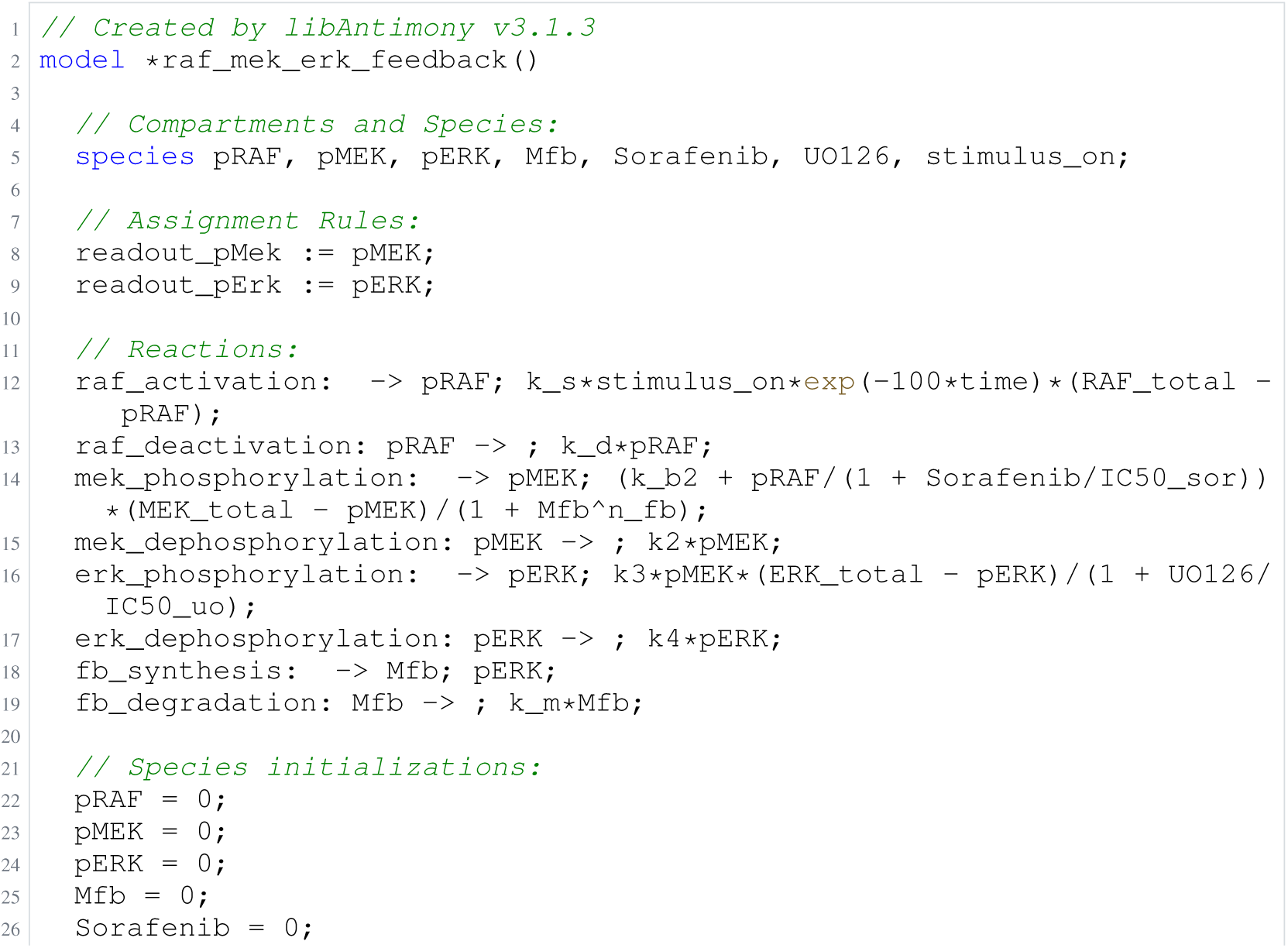
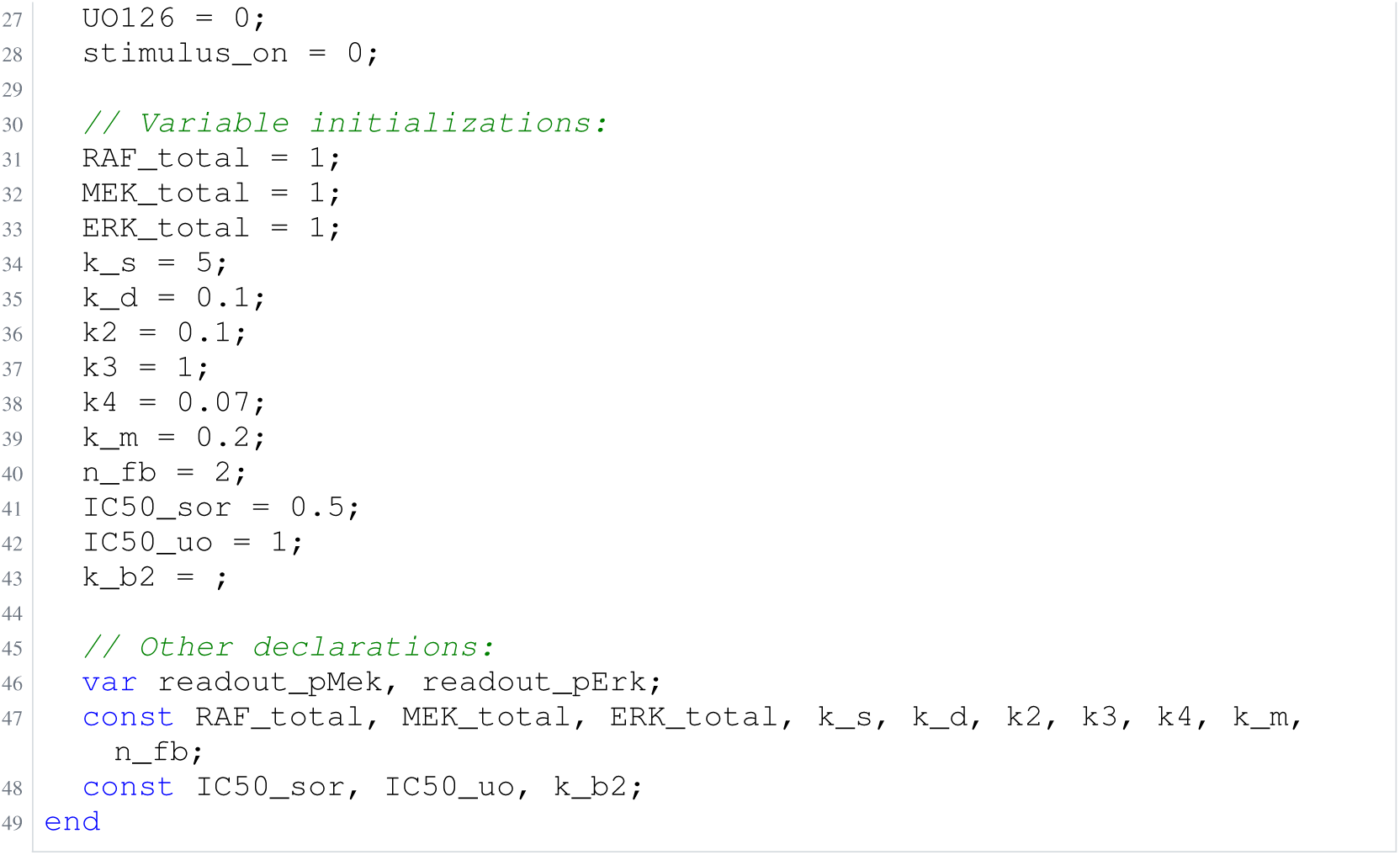
Archived Antimony for fiedler_2; task 57351389_6; selected version 10.

**Listing 61:**
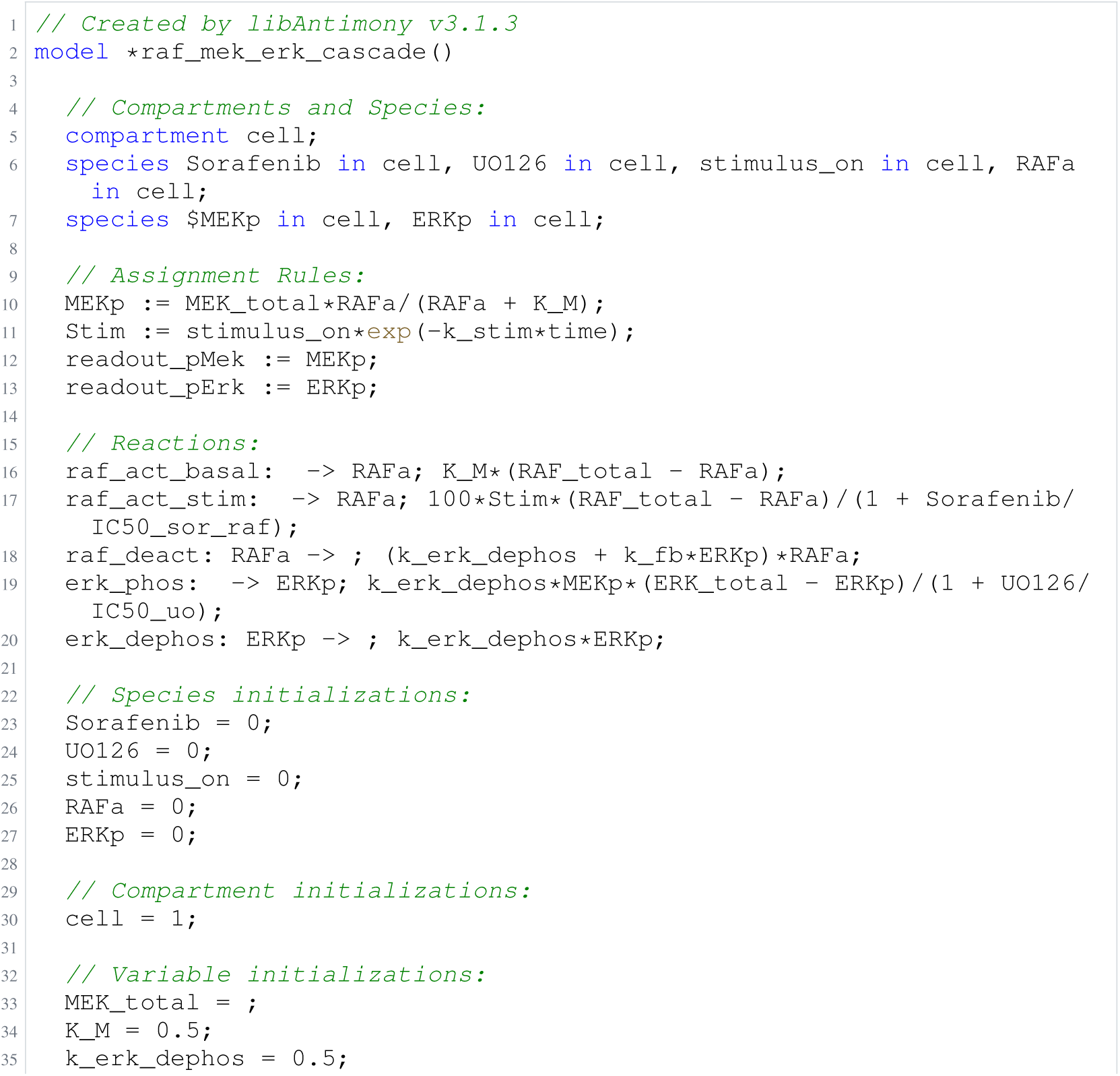
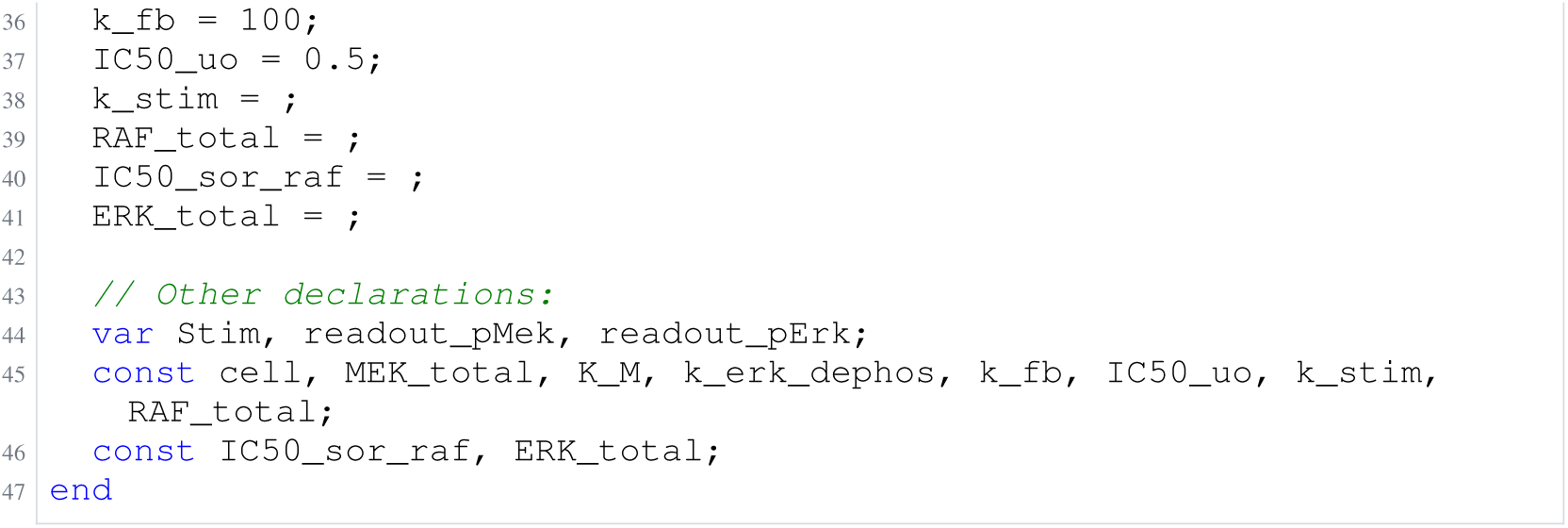
Archived Antimony for fiedler_3; task 57351389_7; selected version 16.

**Listing 62:**
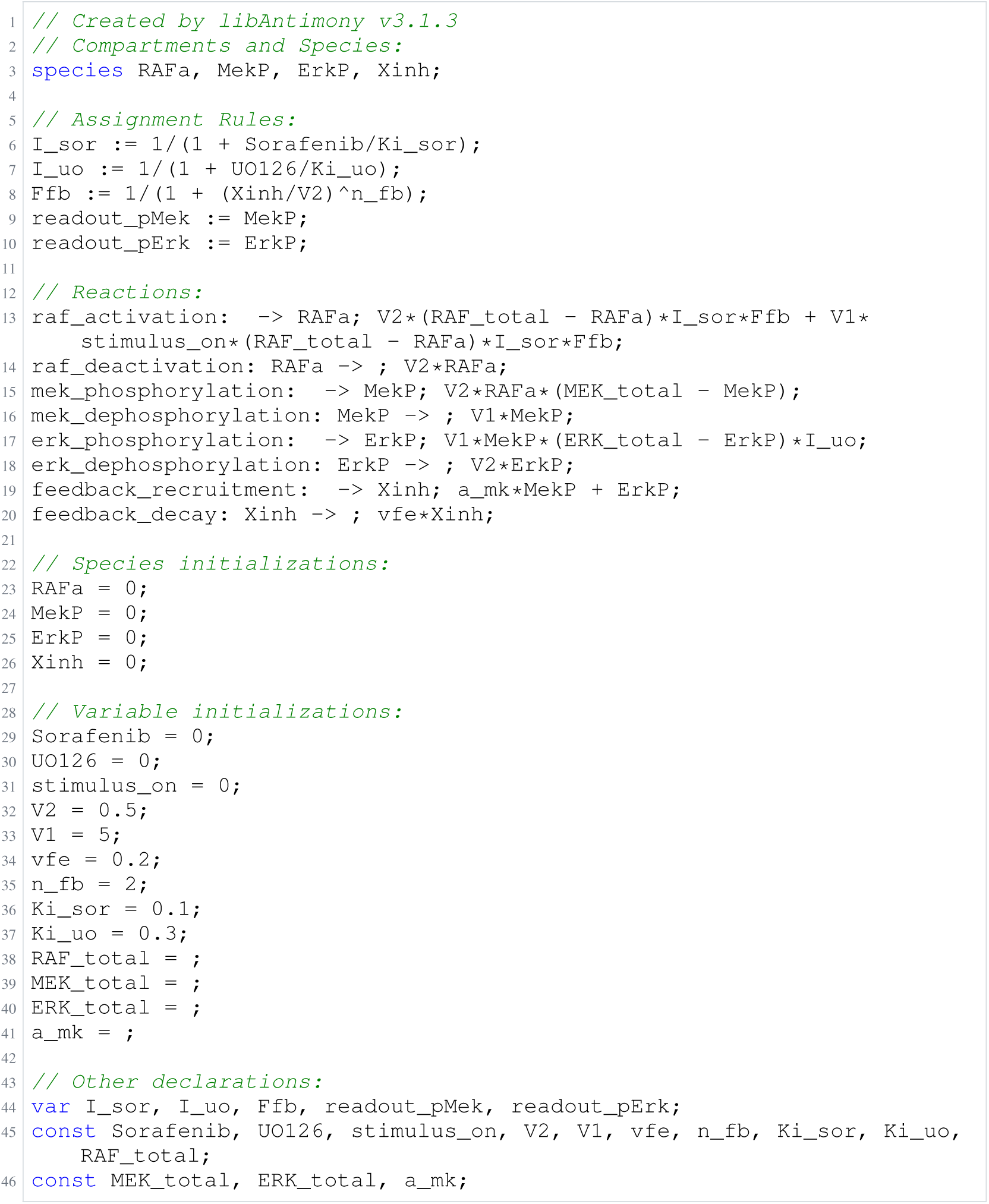
Archived Antimony for fiedler_4; task 57351389_8; selected version 16.

**Listing 63:**
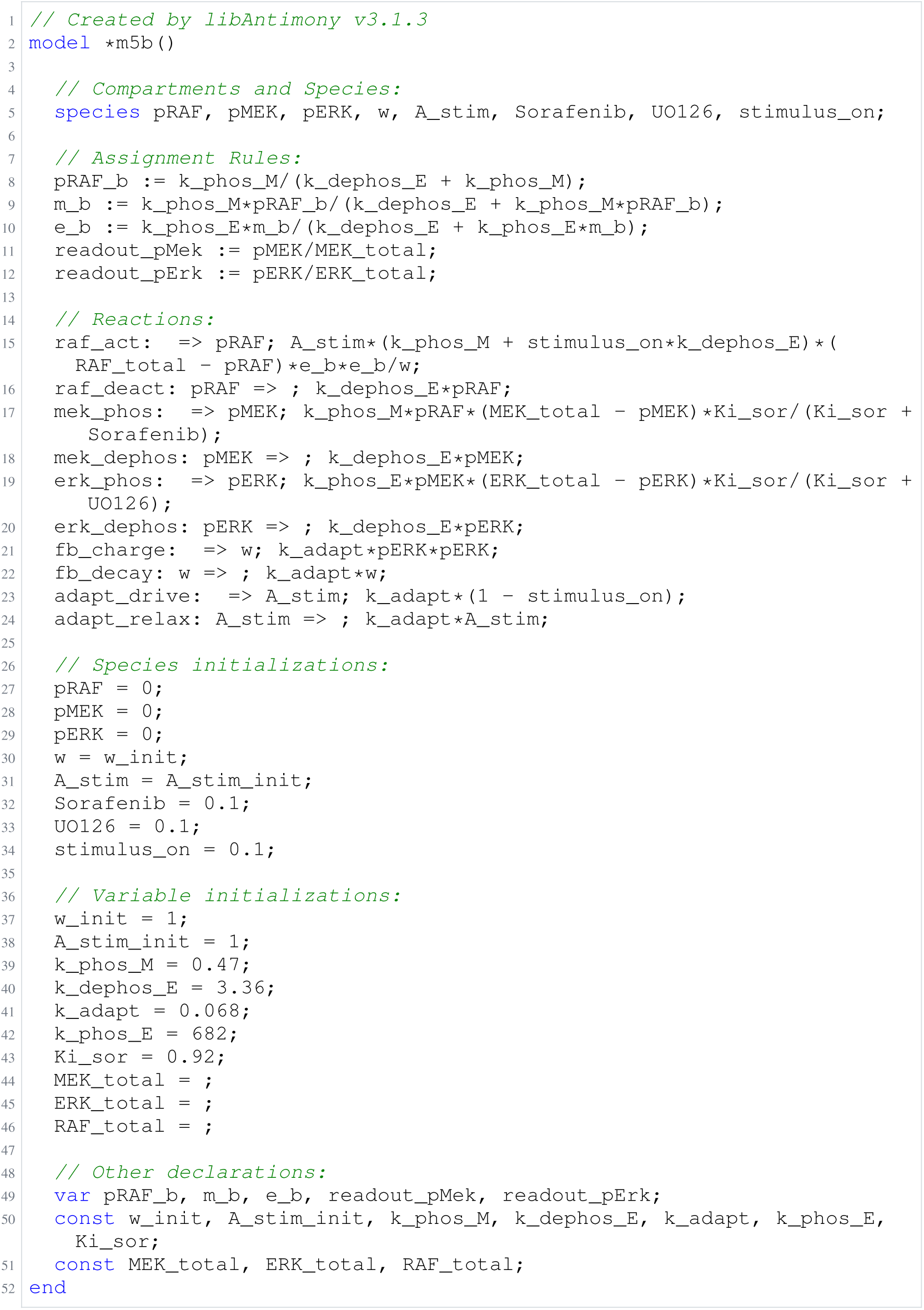
Archived Antimony for fiedler_5; task 57351389_9; selected version 15.

### H.8 Bruno

**Listing 64:**
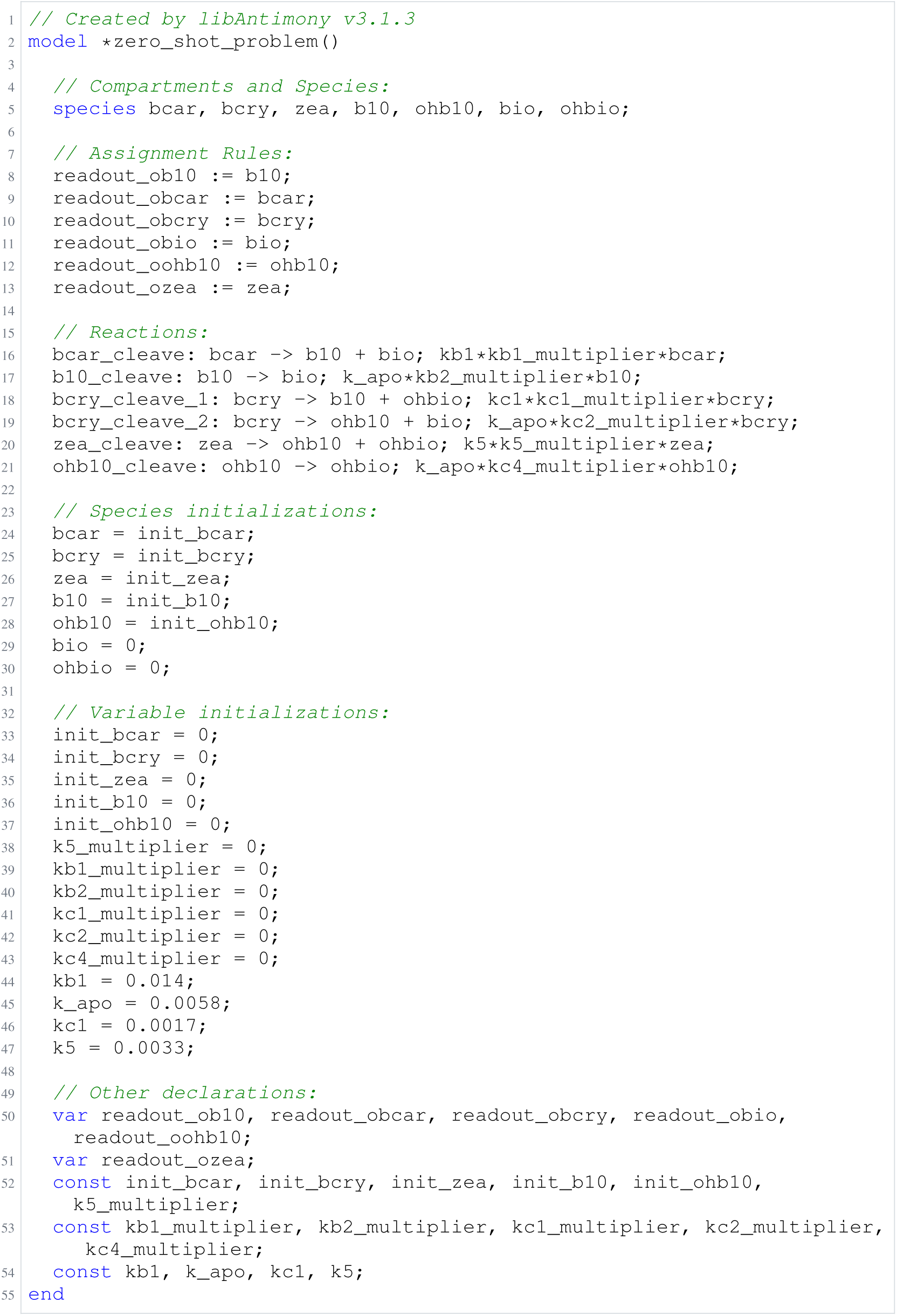
Archived Antimony for bruno_1; task 56836848_20; selected version 2.

**Listing 65:**
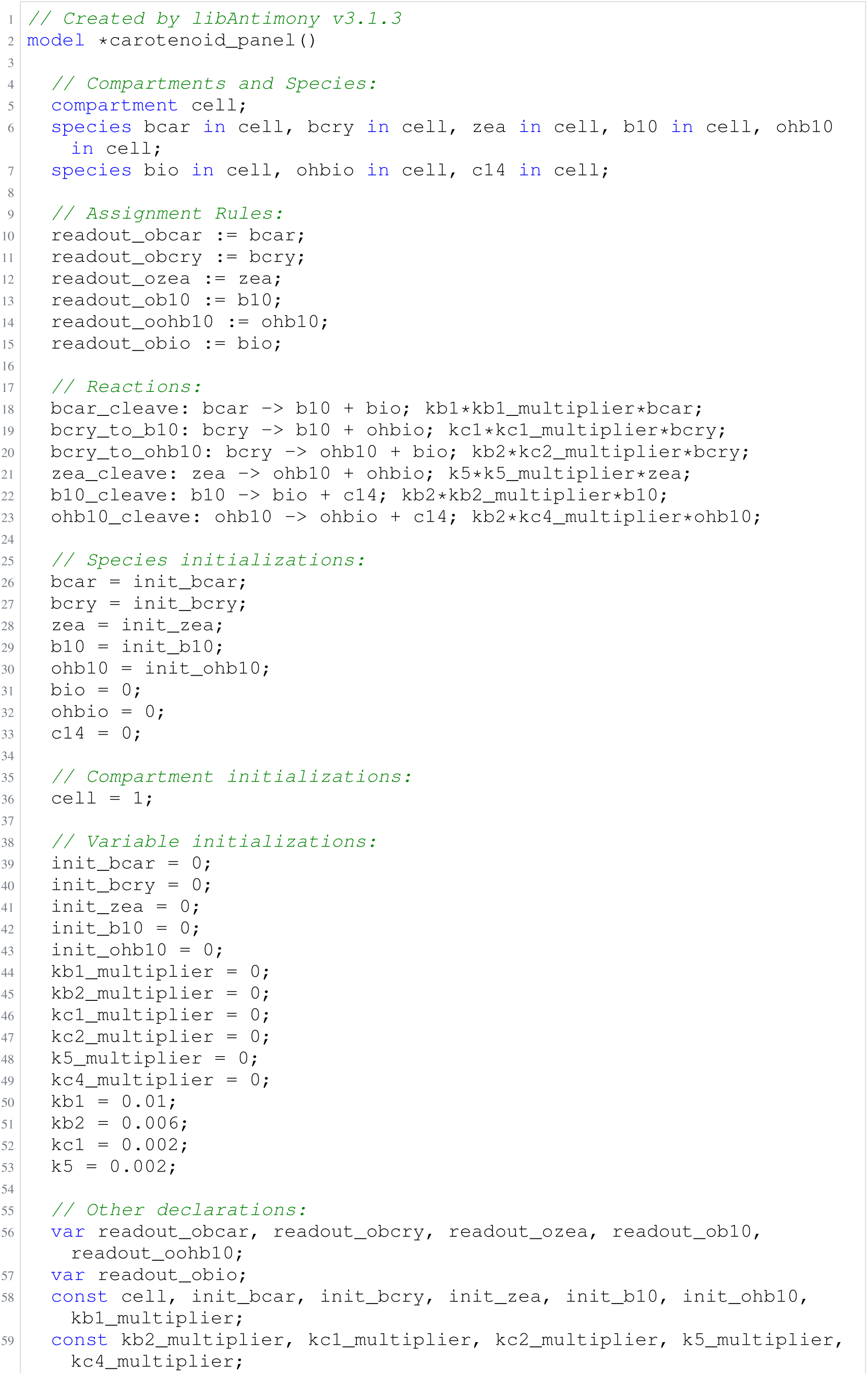
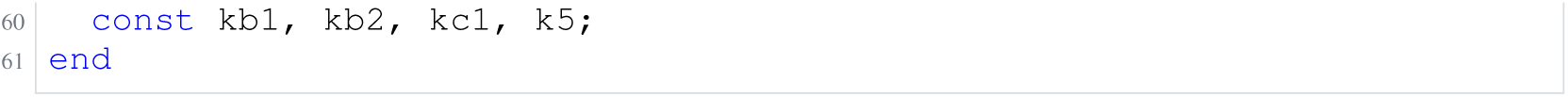
Archived Antimony for bruno_2; task 56836848_21; selected version 6.

**Listing 66:**
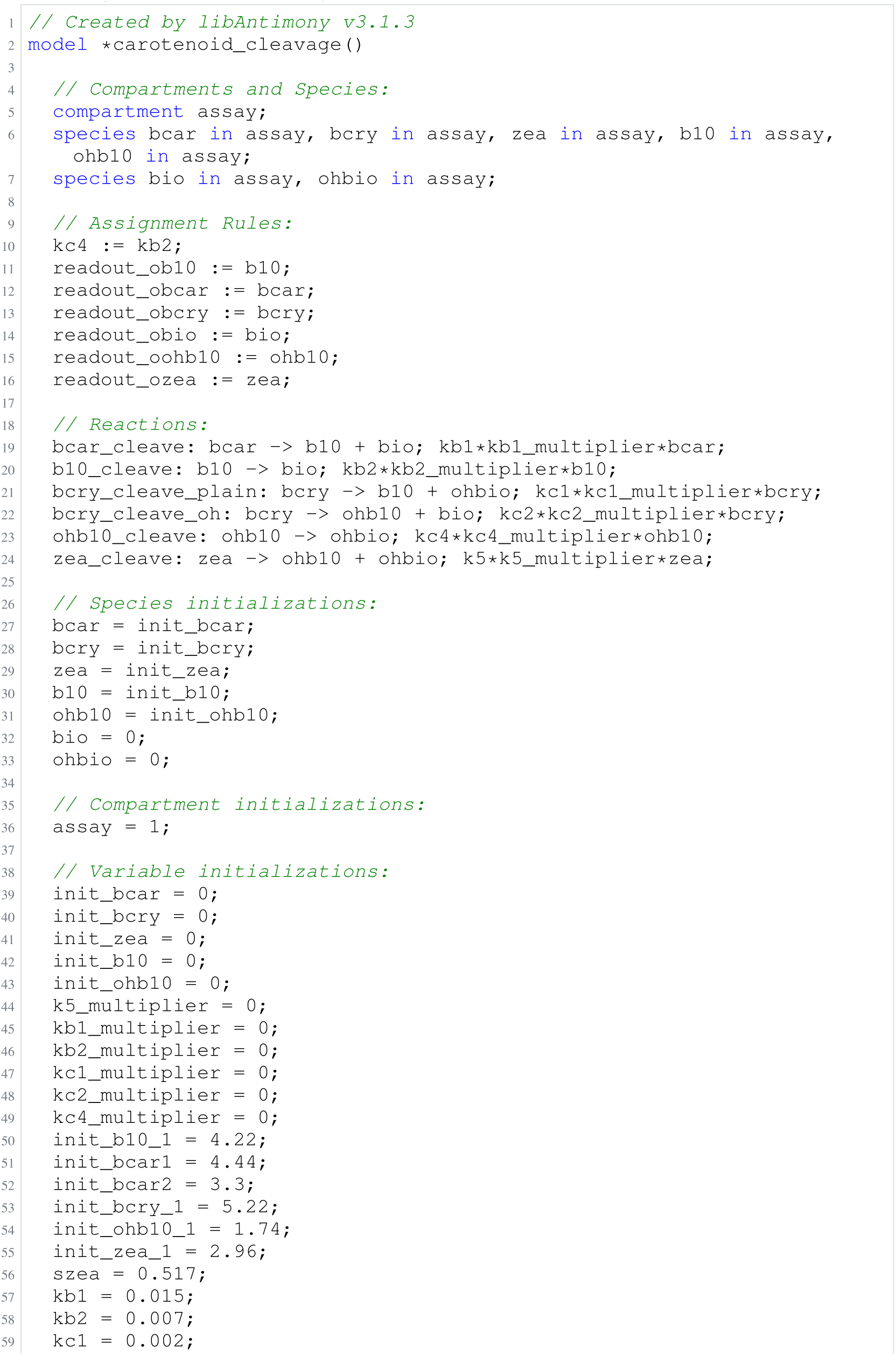
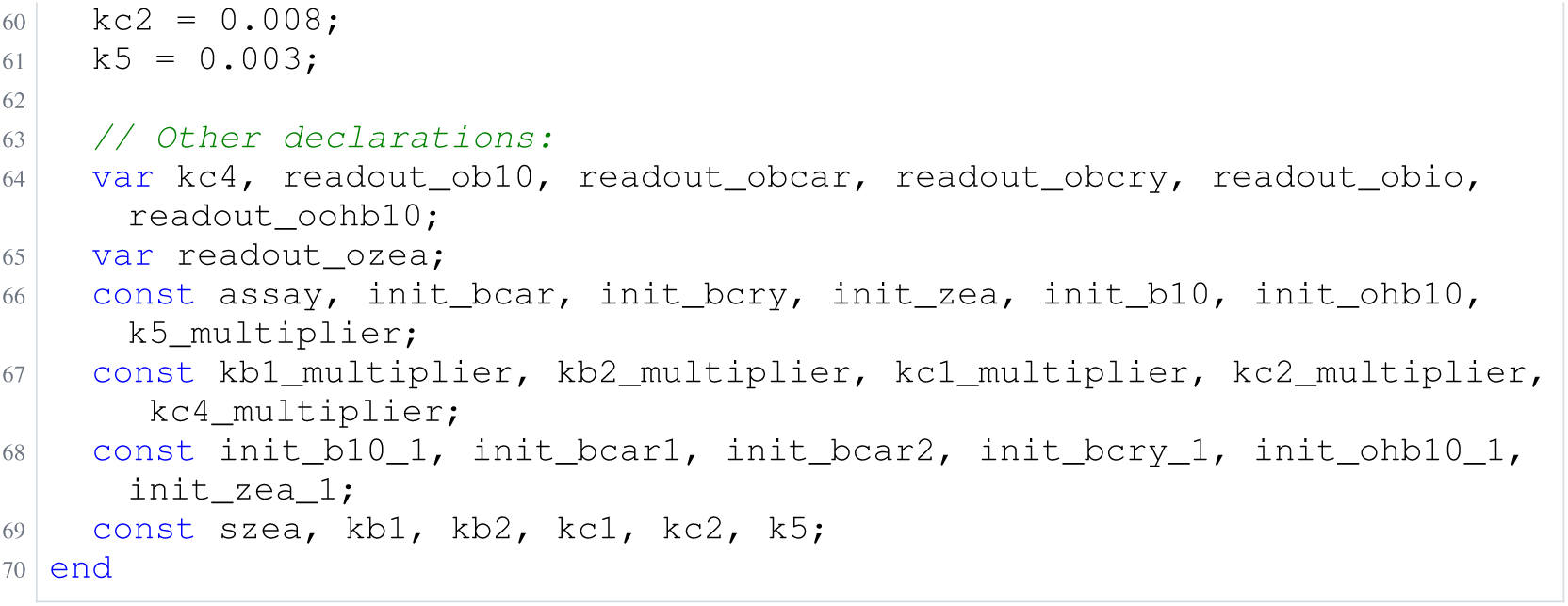
Archived Antimony for bruno_3; task 56836848_22; selected version 1.

**Listing 67:**
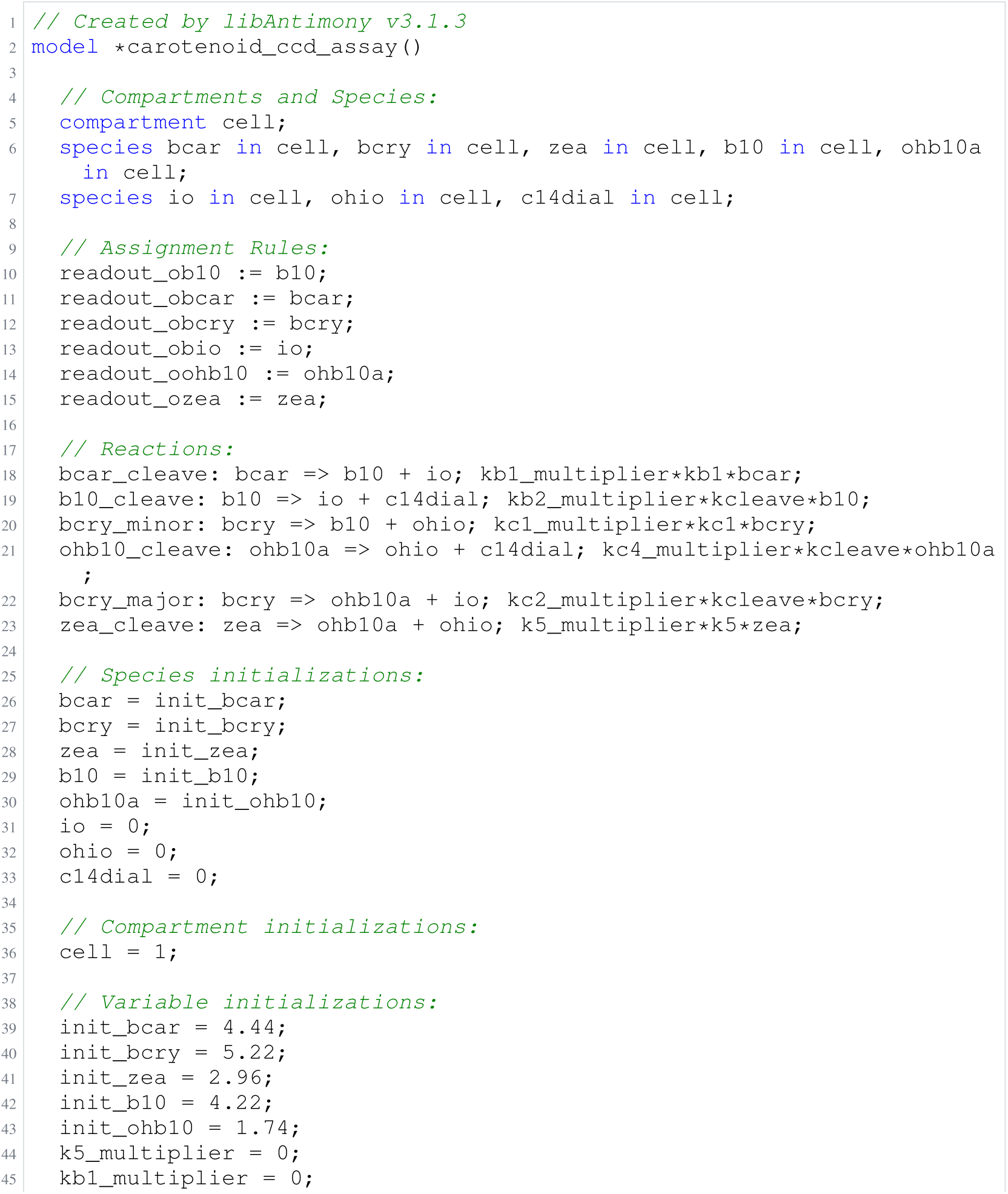
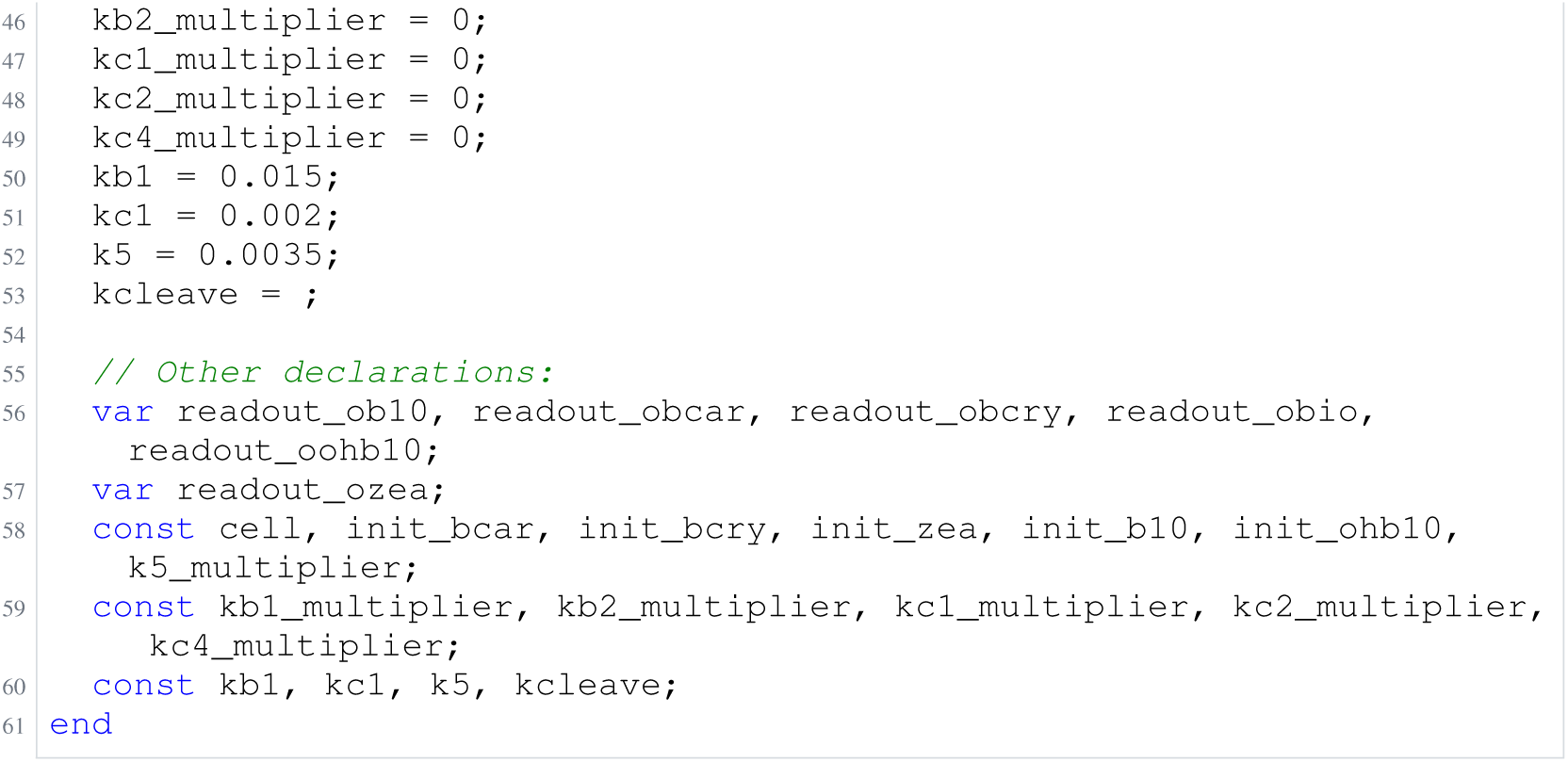
Archived Antimony for bruno_4; task 56836848_23; selected version 5.

**Listing 68:**
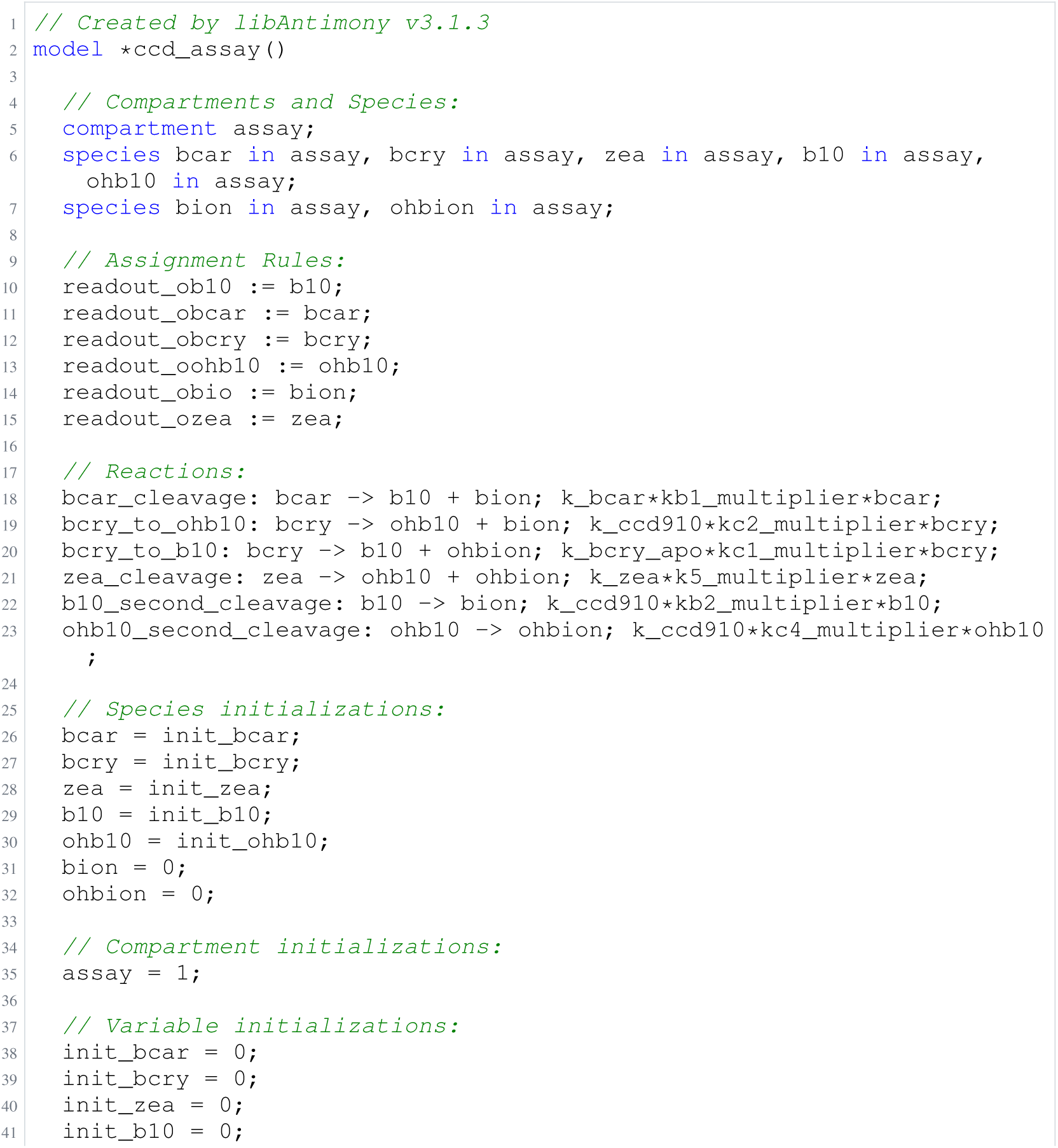
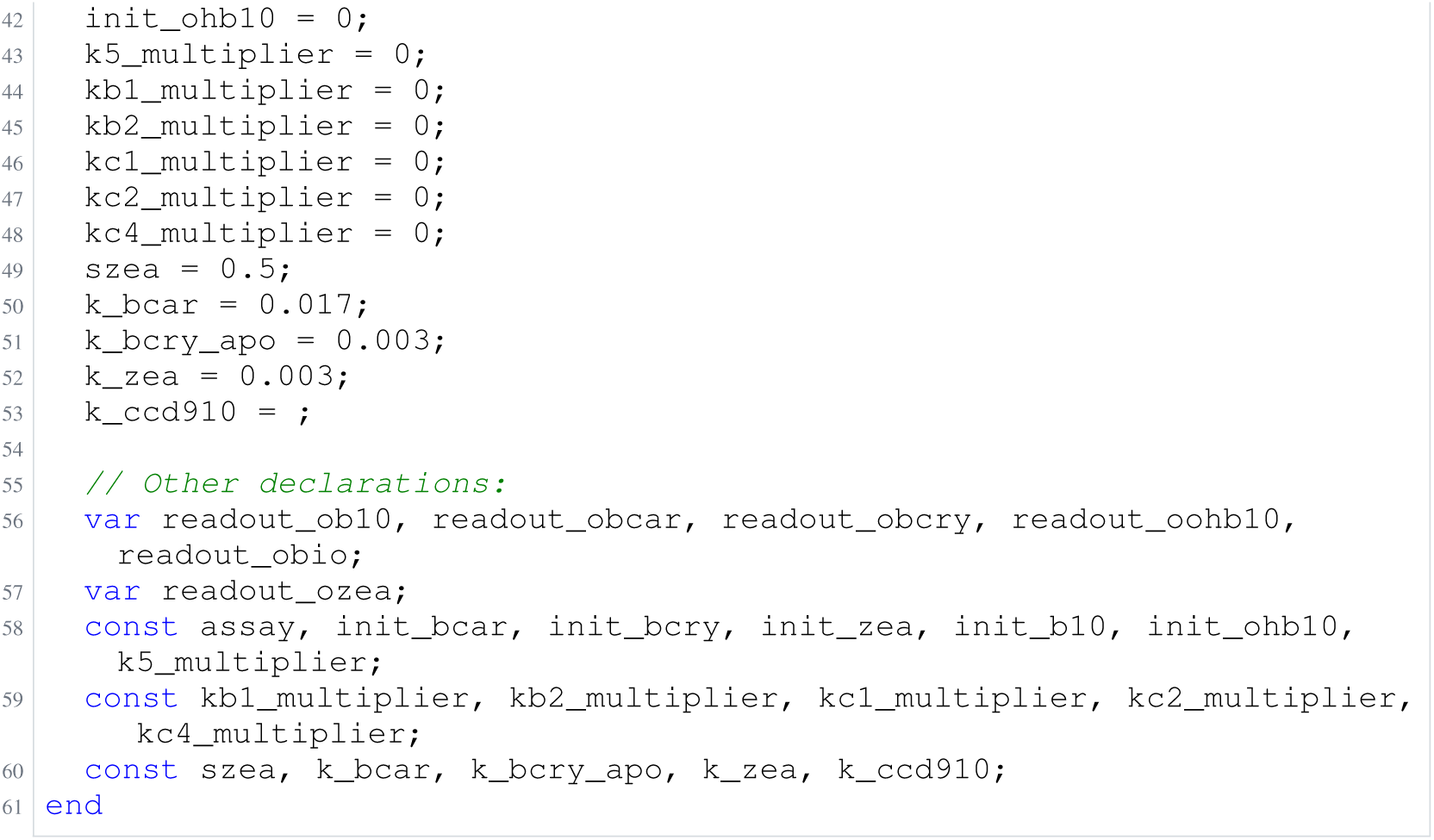
Archived Antimony for bruno_5; task 56836848_24; selected version 2.

### H.9 Oliveira

**Listing 69:**
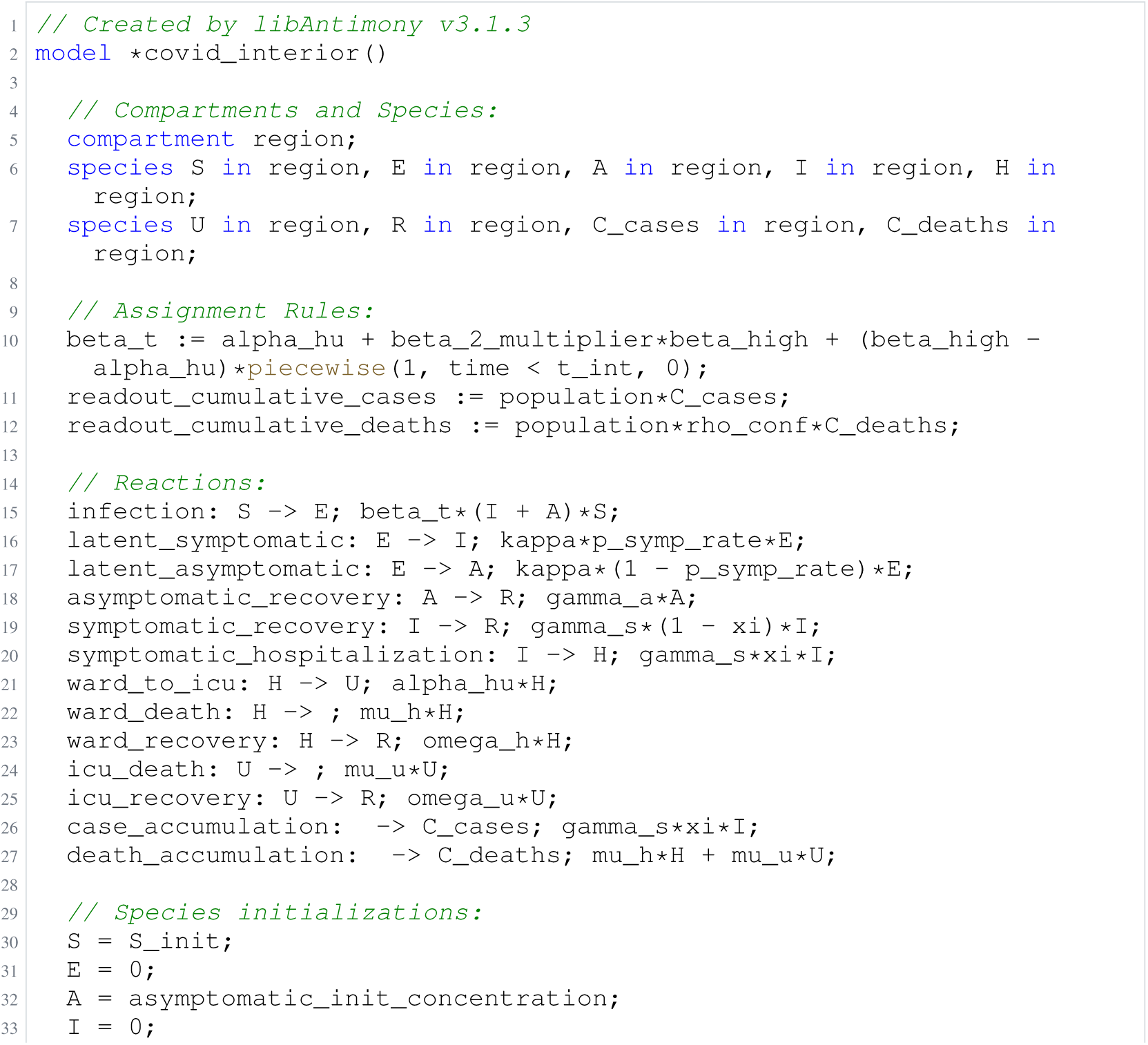
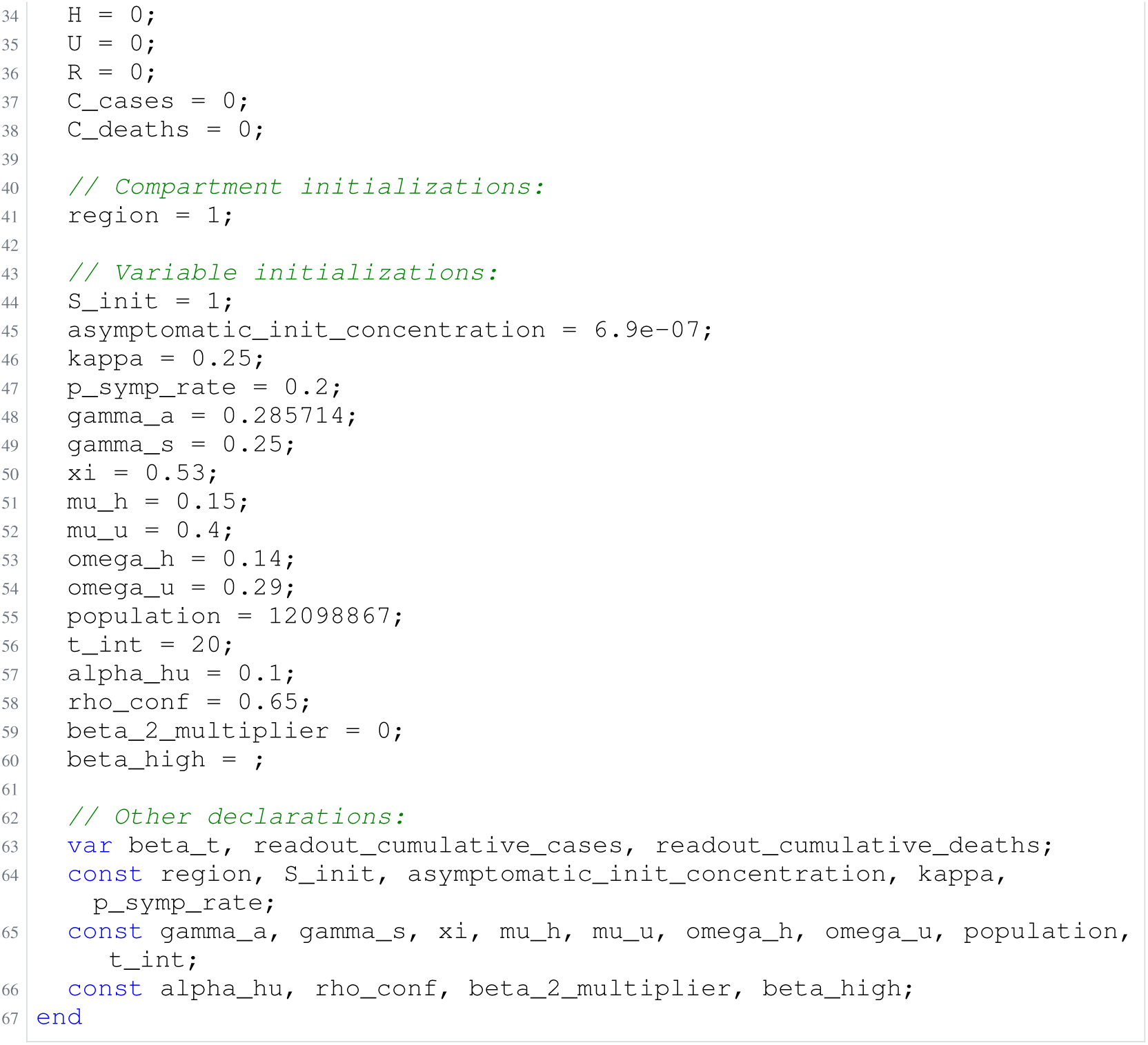
Archived Antimony for oliveira_1; task 57351389_10; selected version 12.

**Listing 70:**
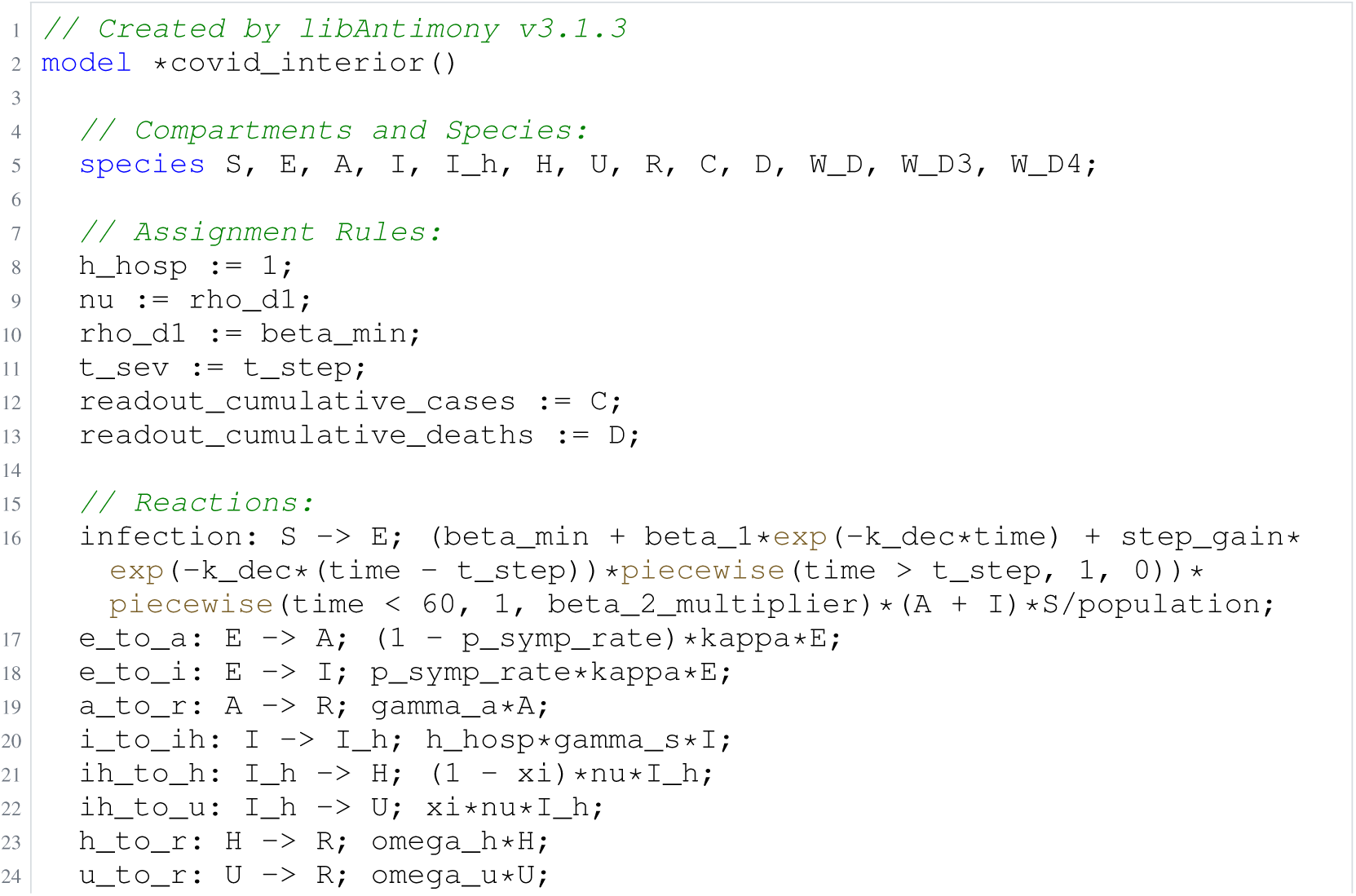
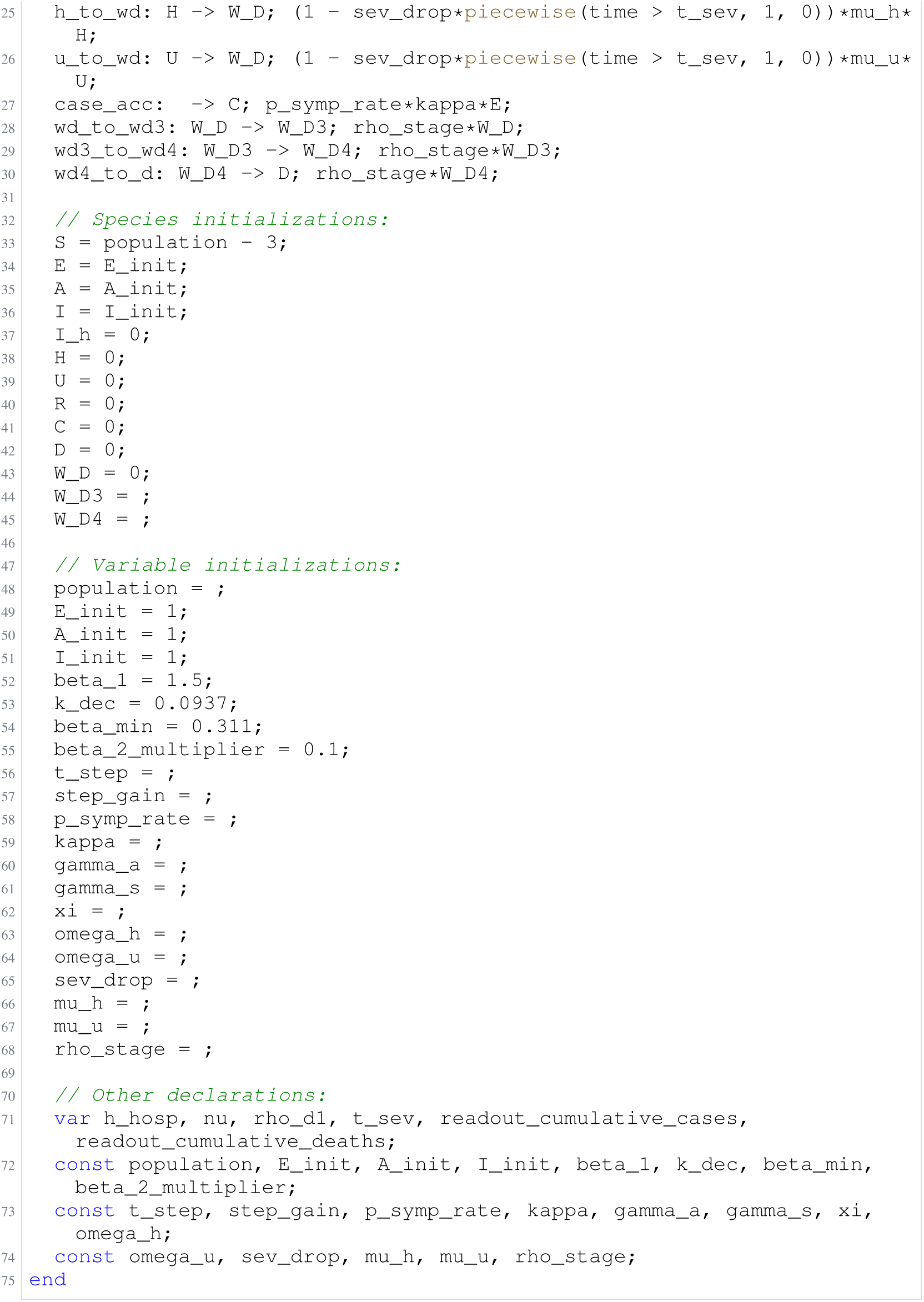
Archived Antimony for oliveira_2; task 57351389_11; selected version 66.

**Listing 71:**
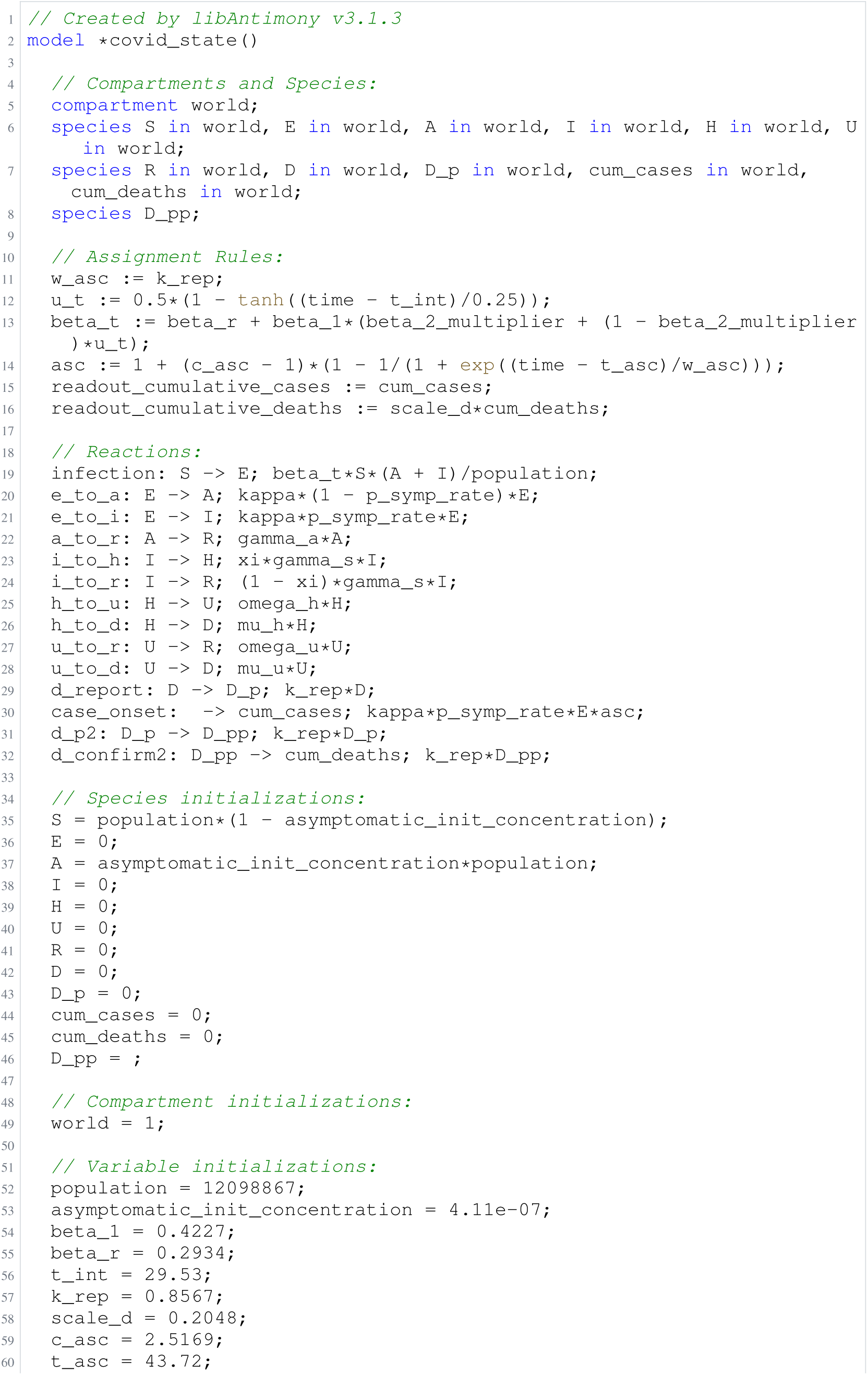
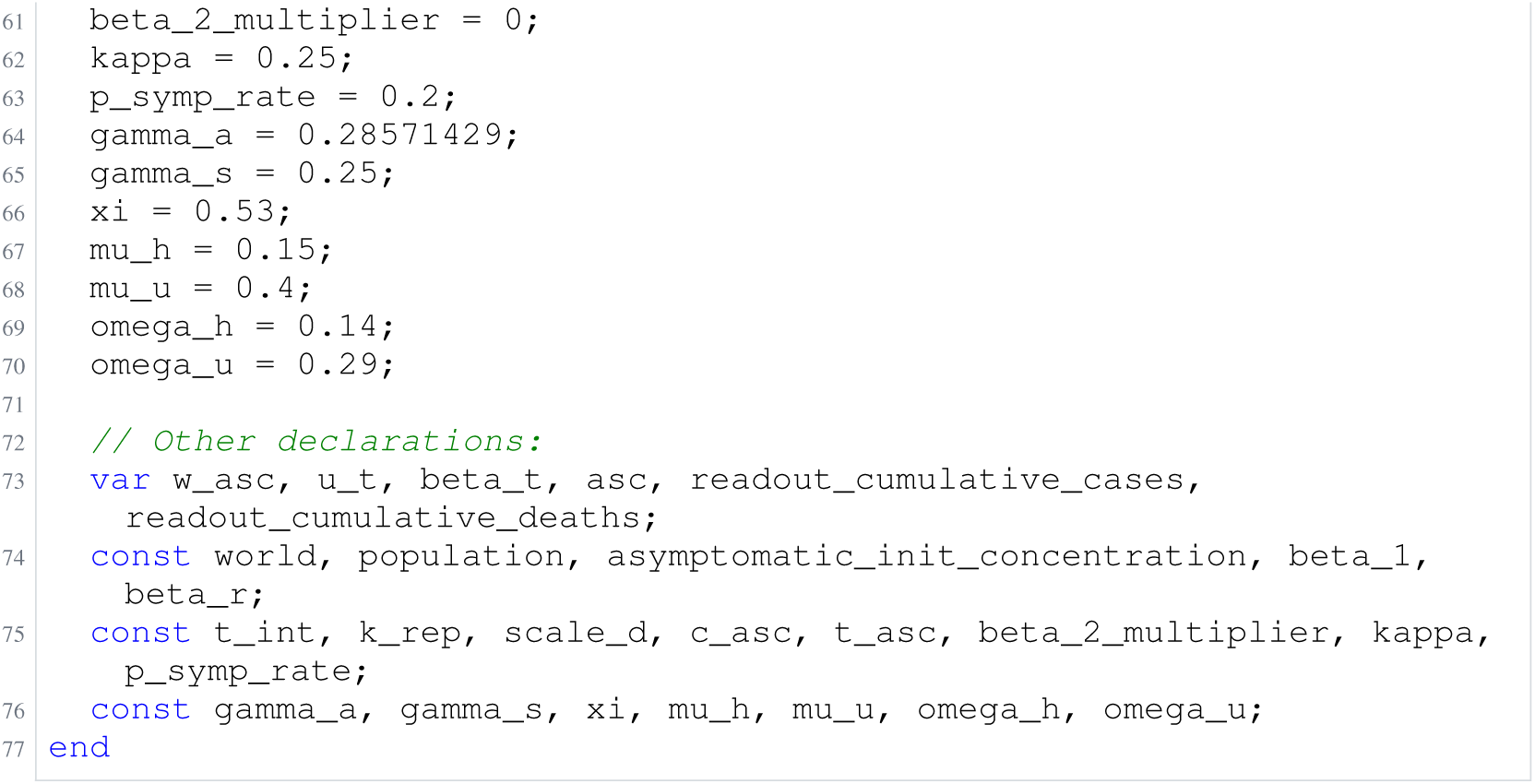
Archived Antimony for oliveira_3; task 57351389_12; selected version 14.

**Listing 72:**
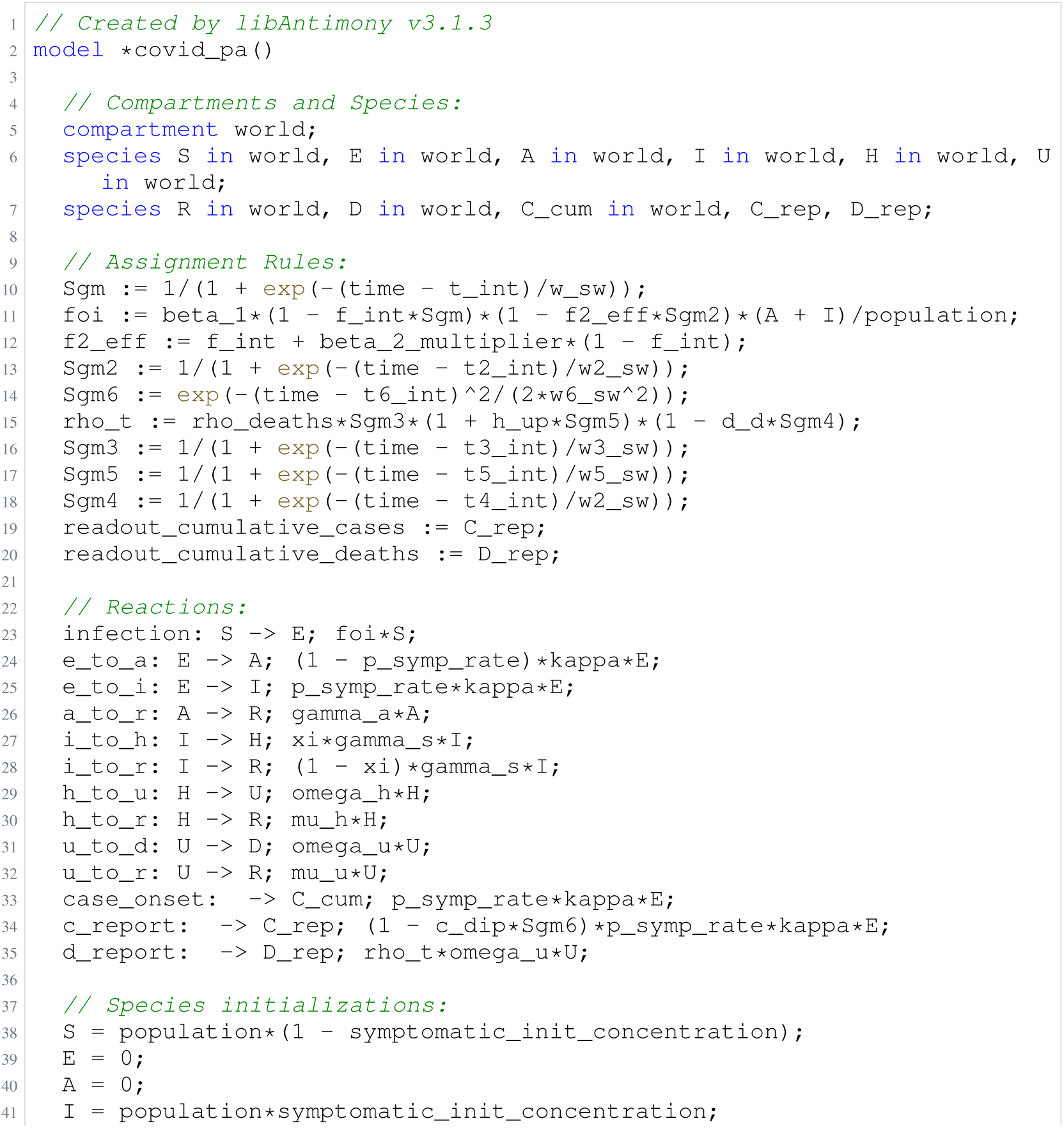
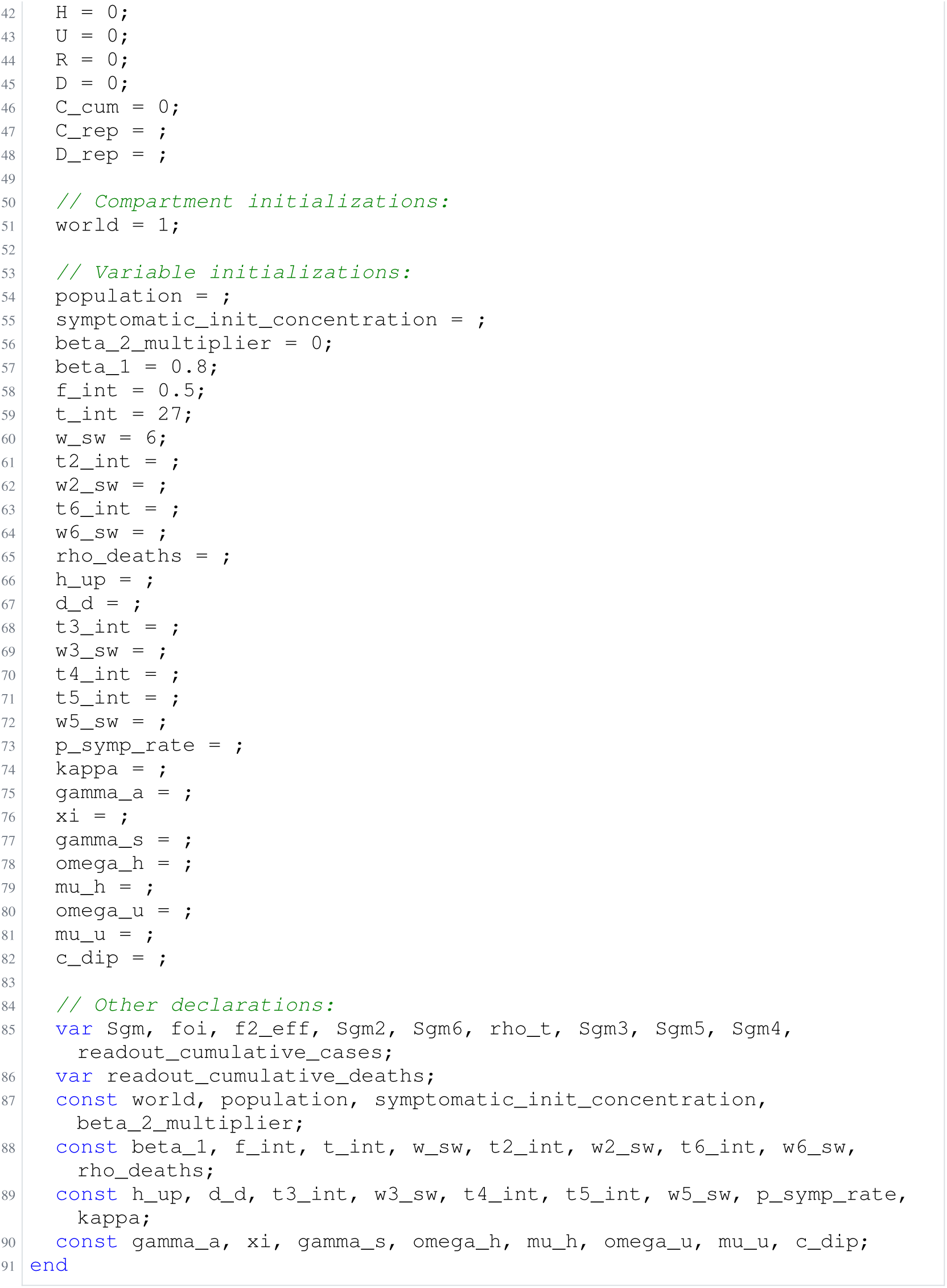
Archived Antimony for oliveira_4; task 57351389_13; selected version 8.

### H.10 Sneyd

**Listing 73:**
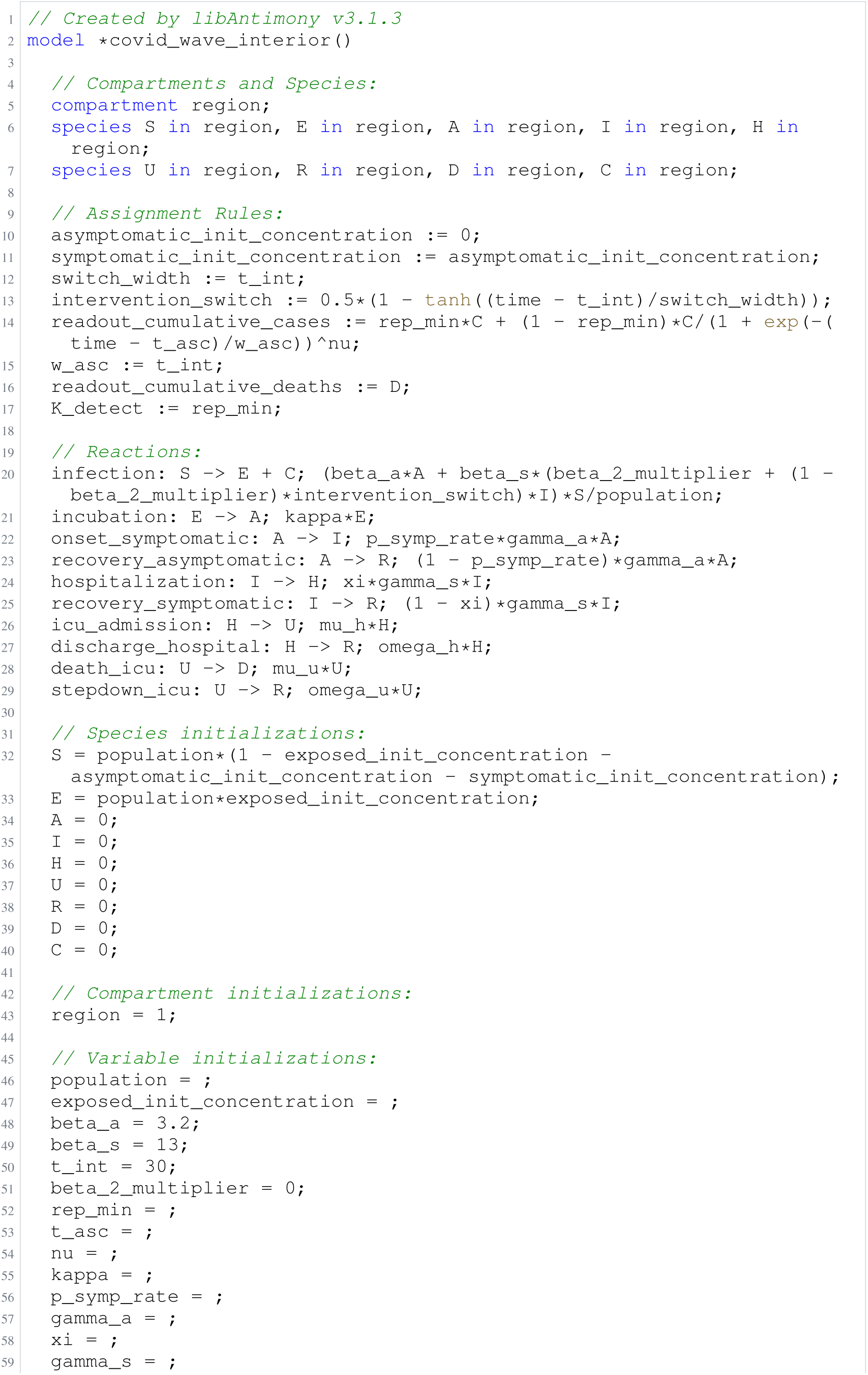
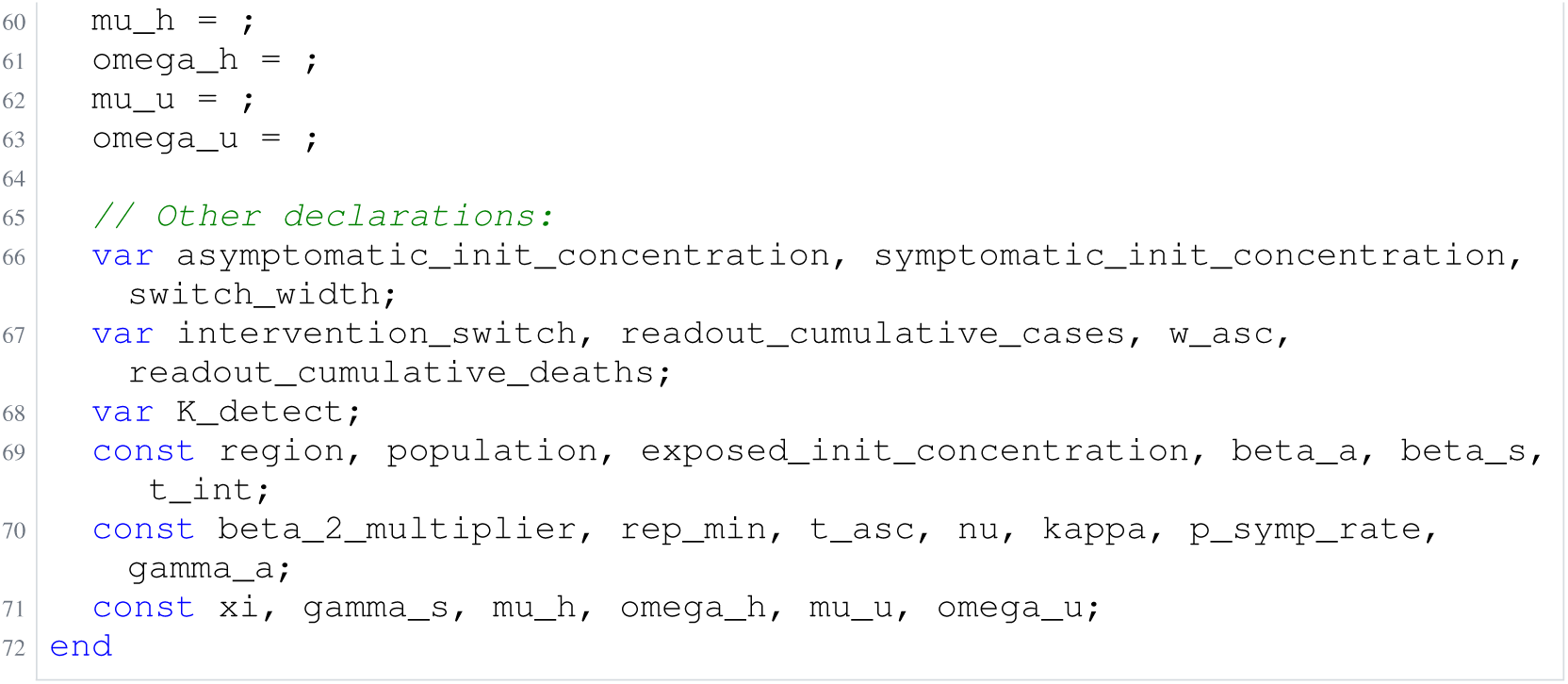
Archived Antimony for oliveira_5; task 57351389_14; selected version 10.

**Listing 74:**
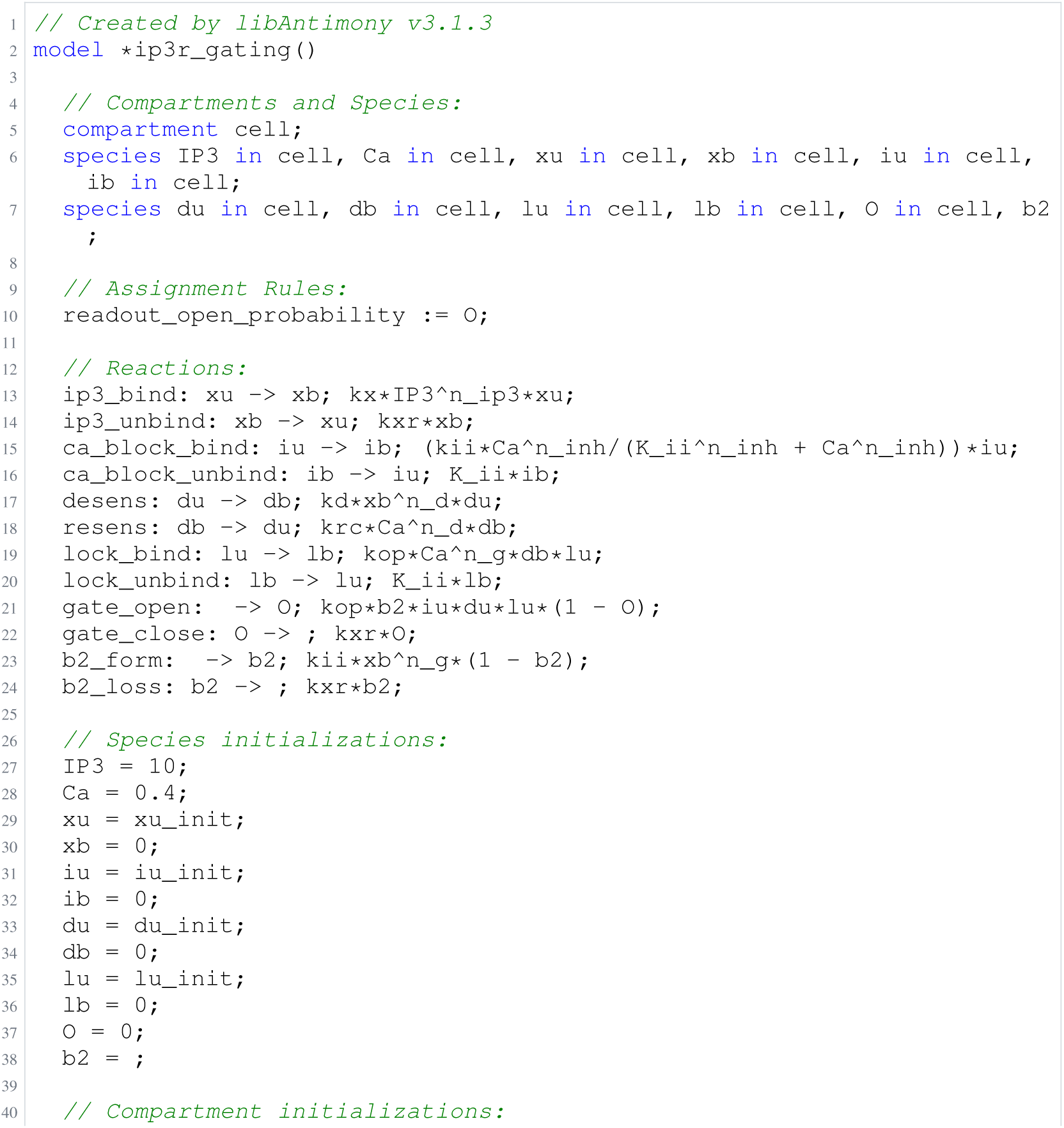
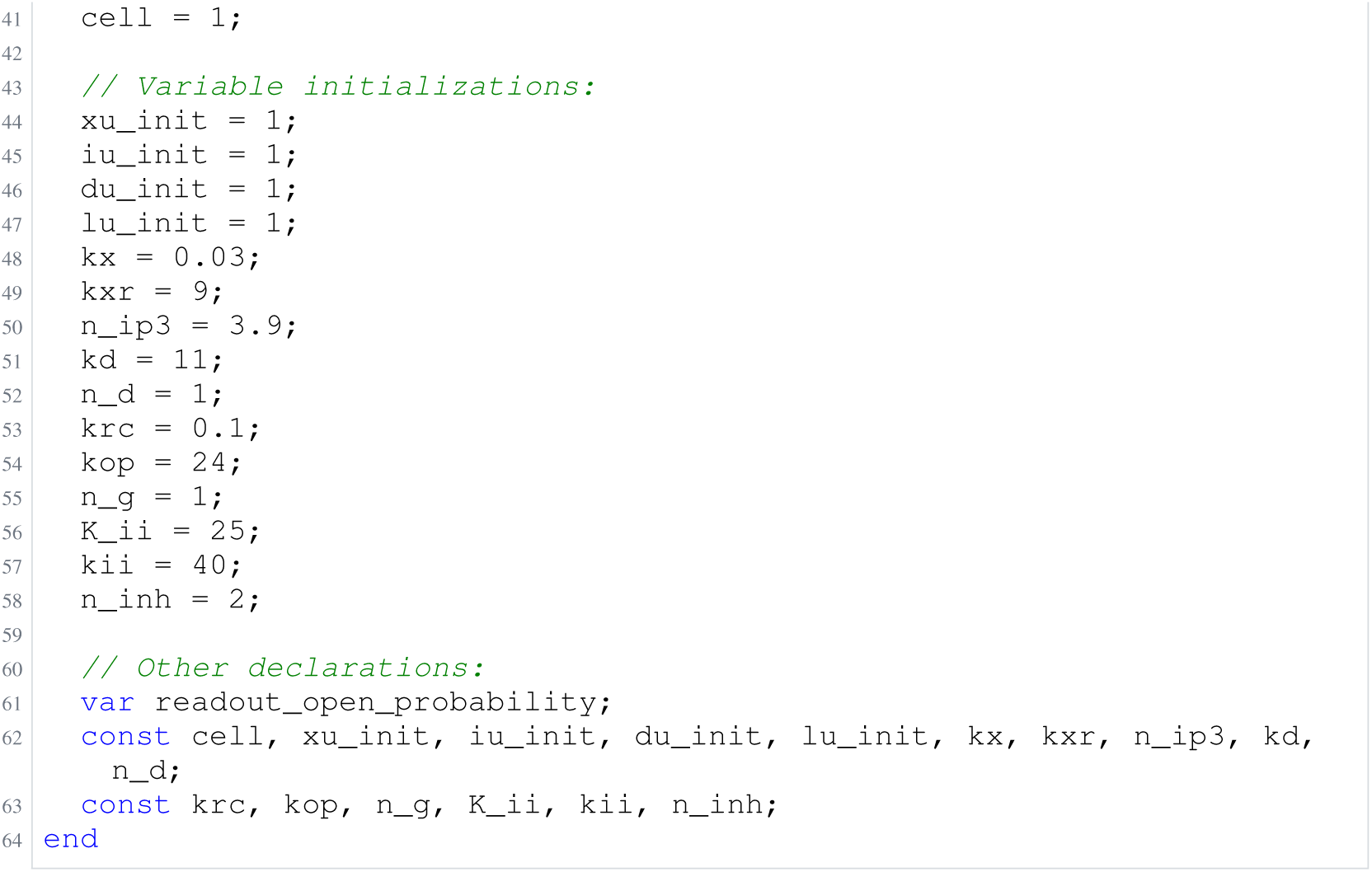
Archived Antimony for sneyd_1; task 57351389_0; selected version 21.

**Listing 75:**
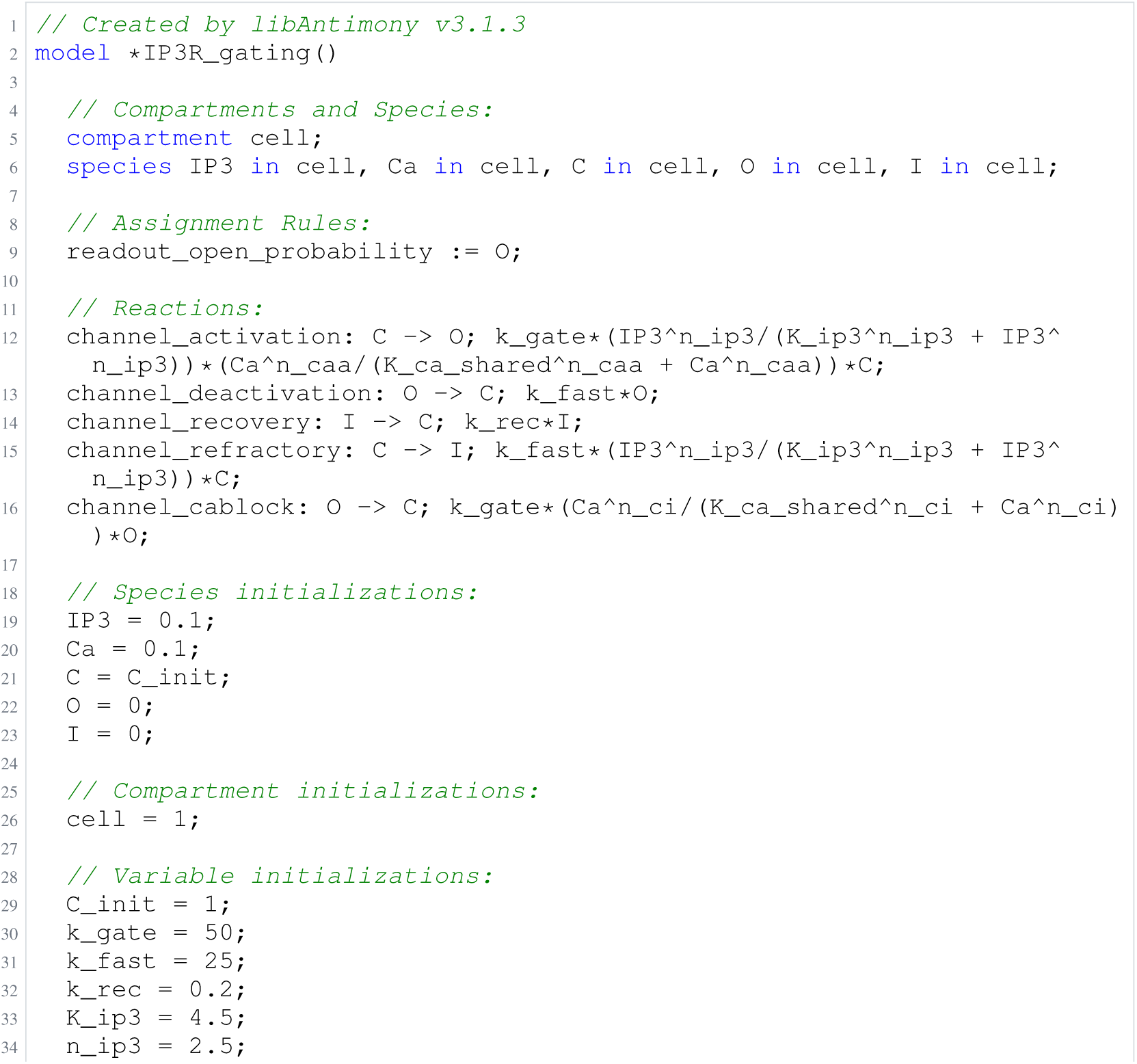
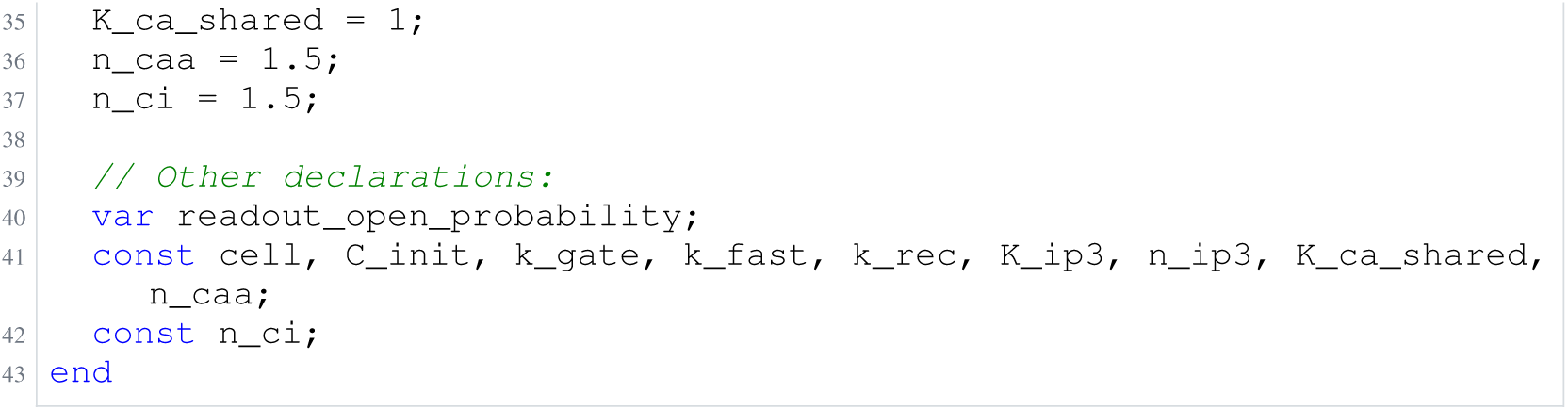
Archived Antimony for sneyd_2; task 57351389_1; selected version 10.

**Listing 76:**
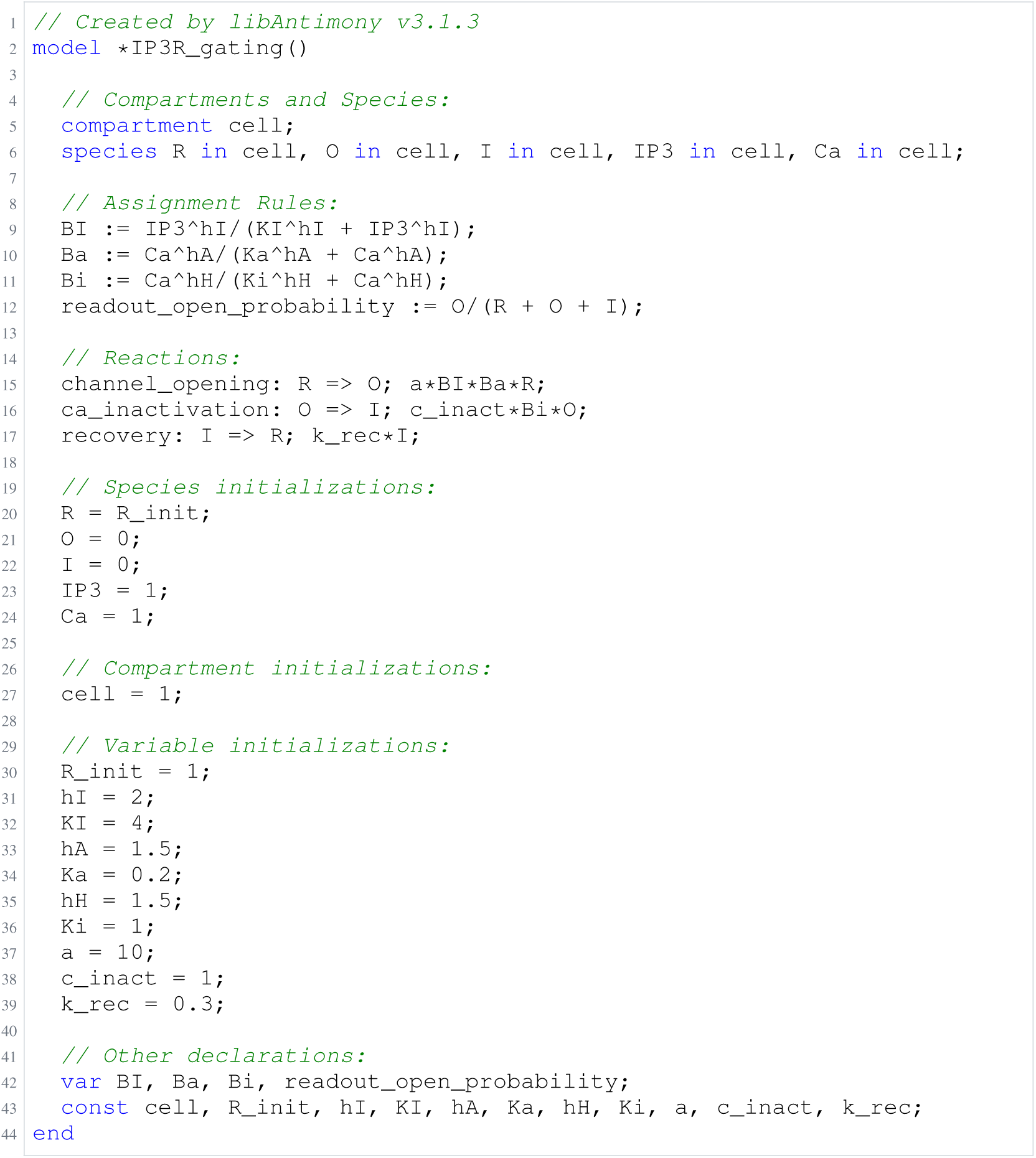
Archived Antimony for sneyd_3; task 57351389_2; selected version 7.

**Listing 77:**
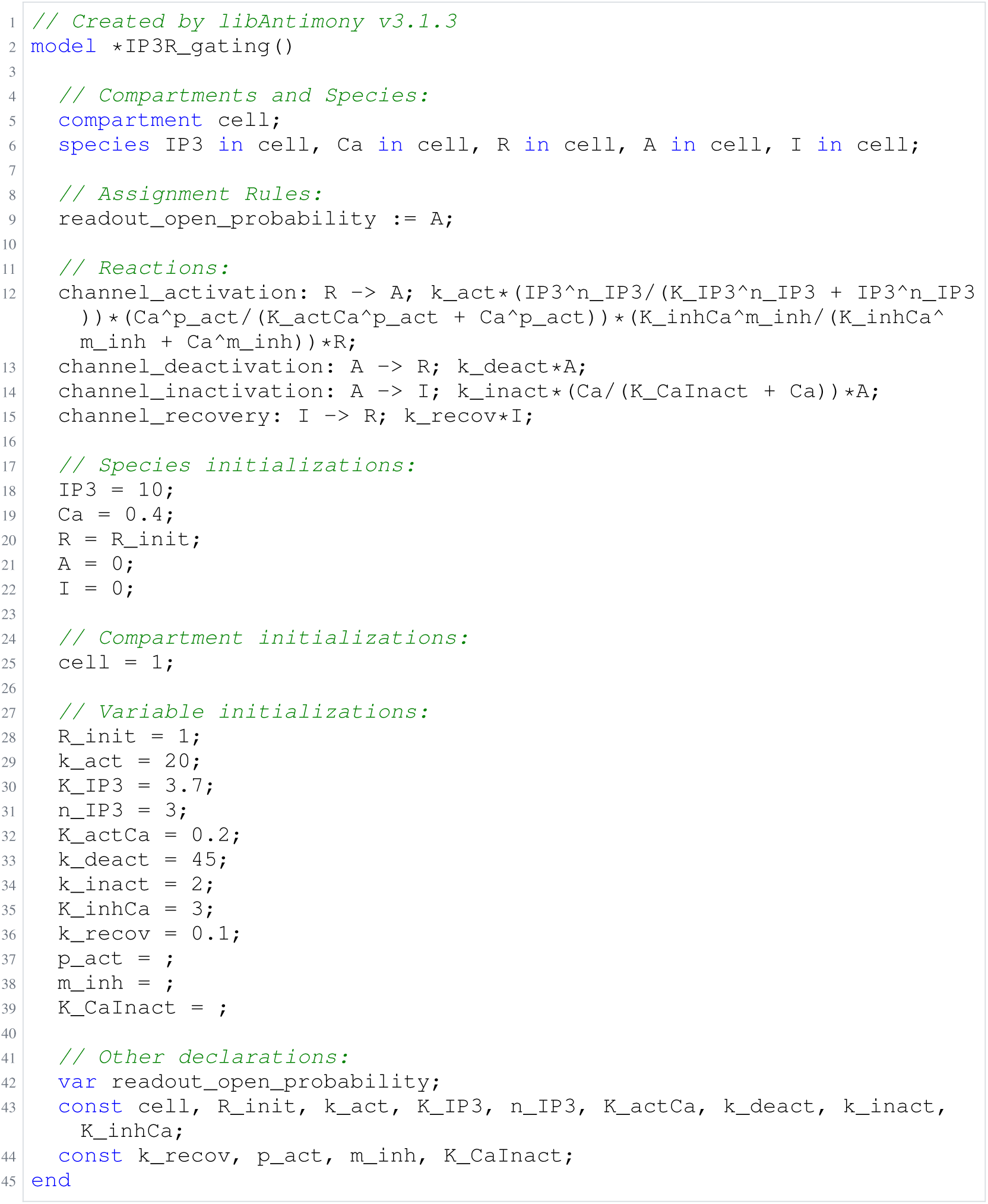
Archived Antimony for sneyd_4; task 57351389_3; selected version 6.

**Listing 78:**
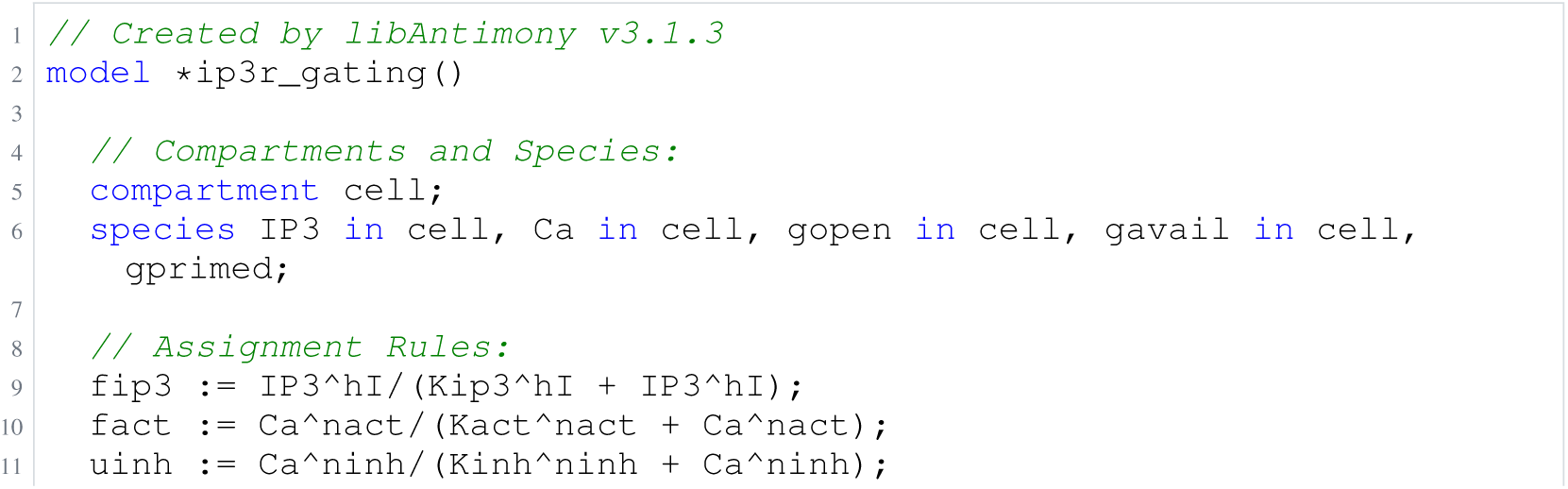
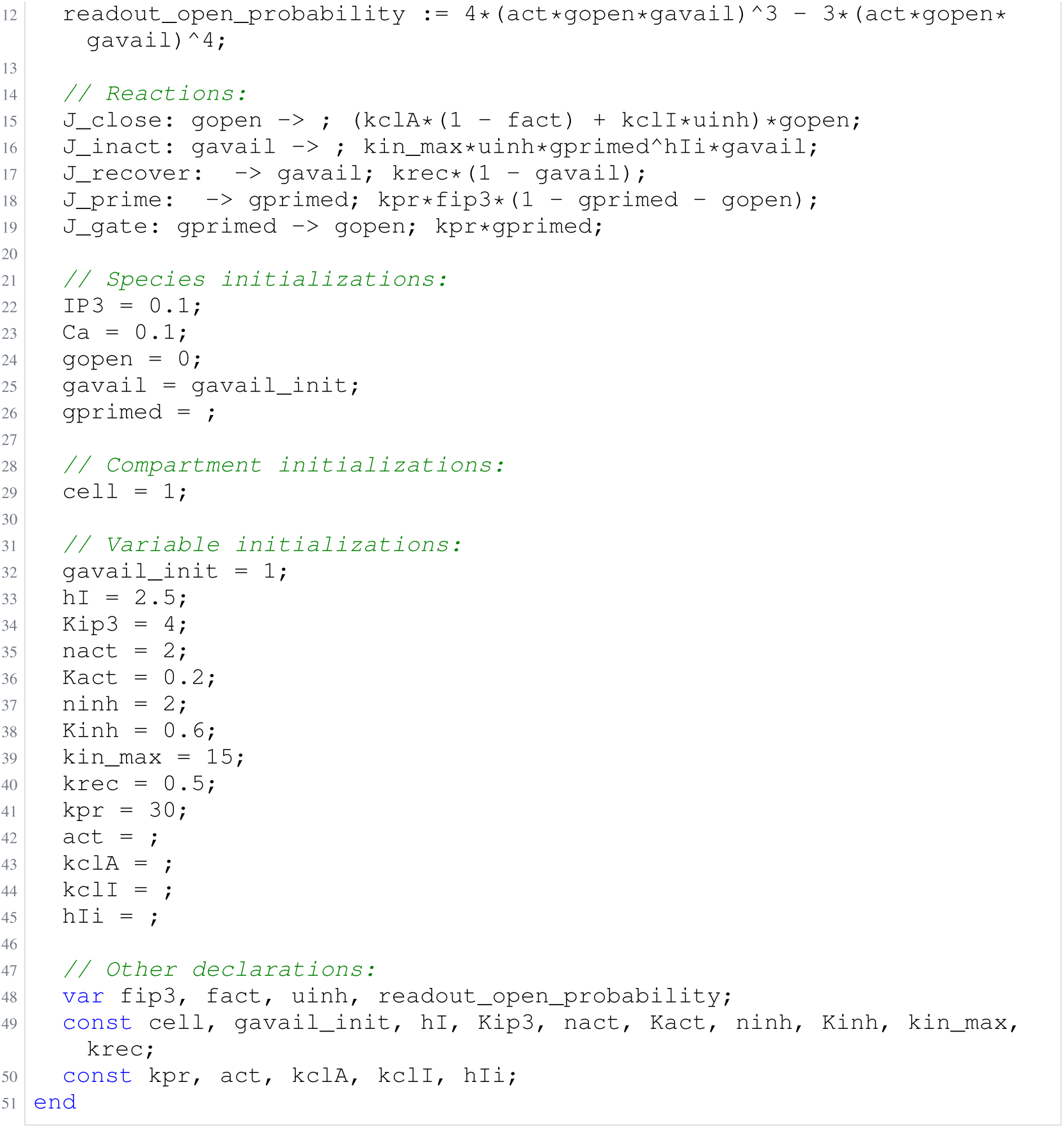
Archived Antimony for sneyd_5; task 57351389_4; selected version 19.

### H.11 Blasi

**Listing 79:**
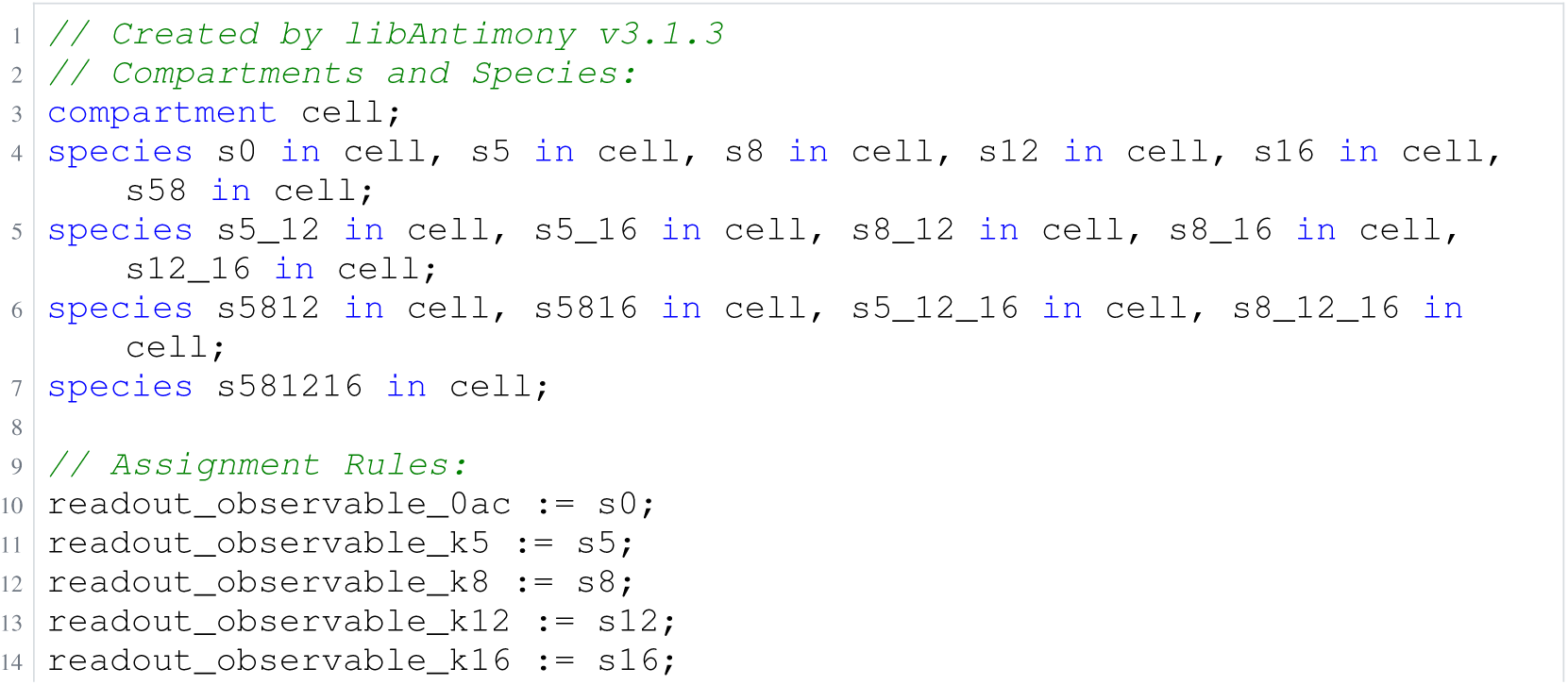
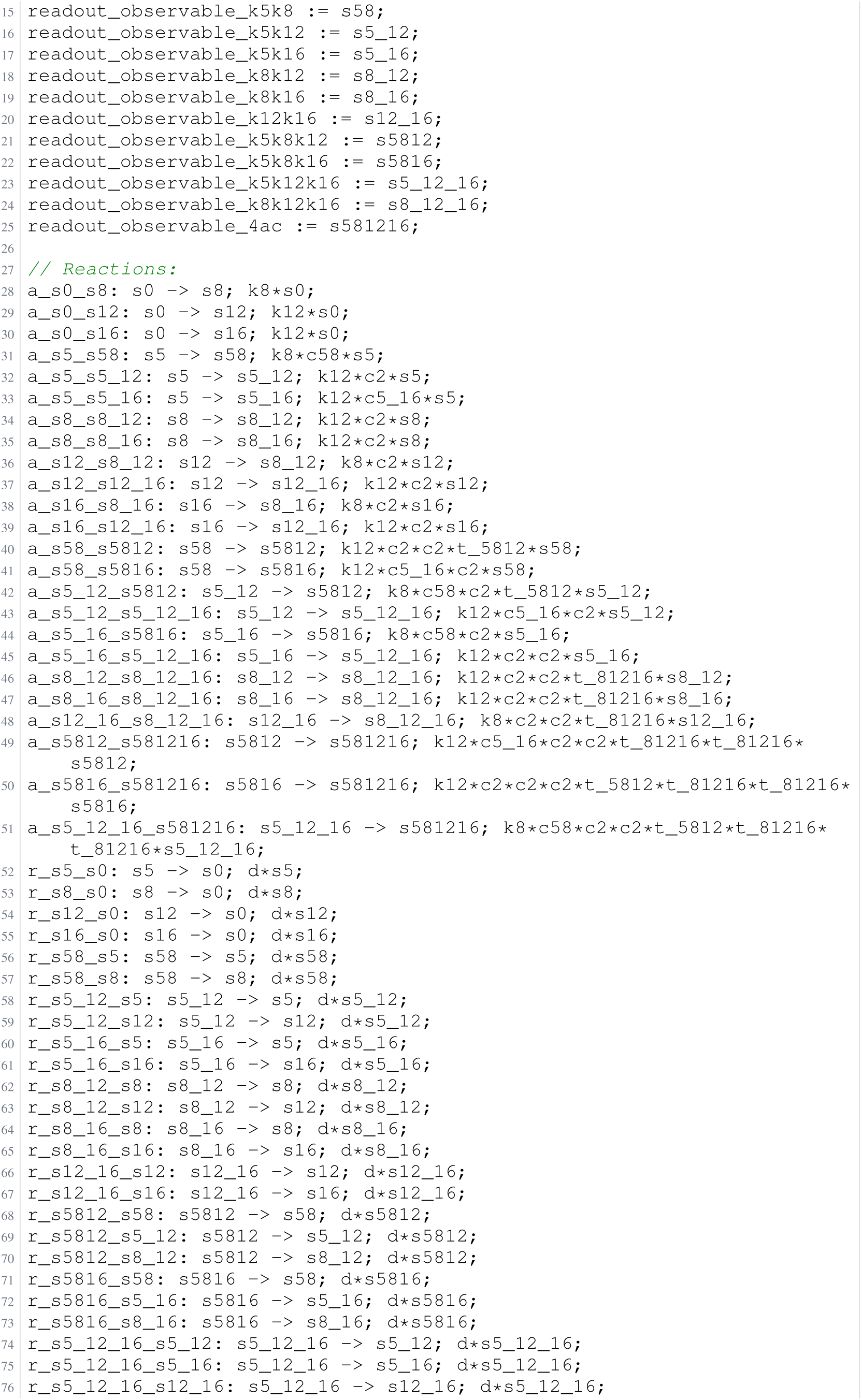
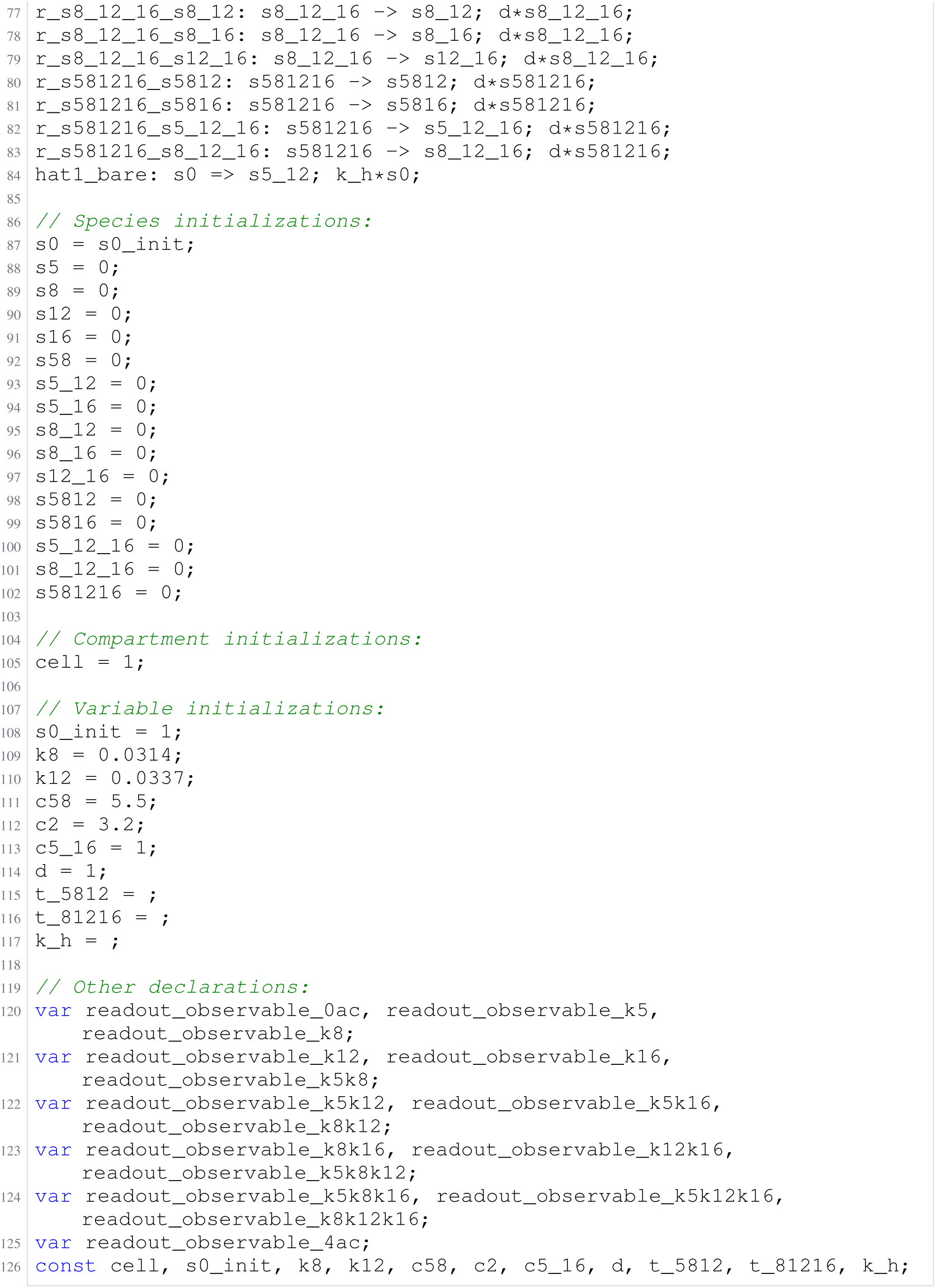
Archived Antimony for blasi_1; task 56836848_5; selected version 16.

**Listing 80:**
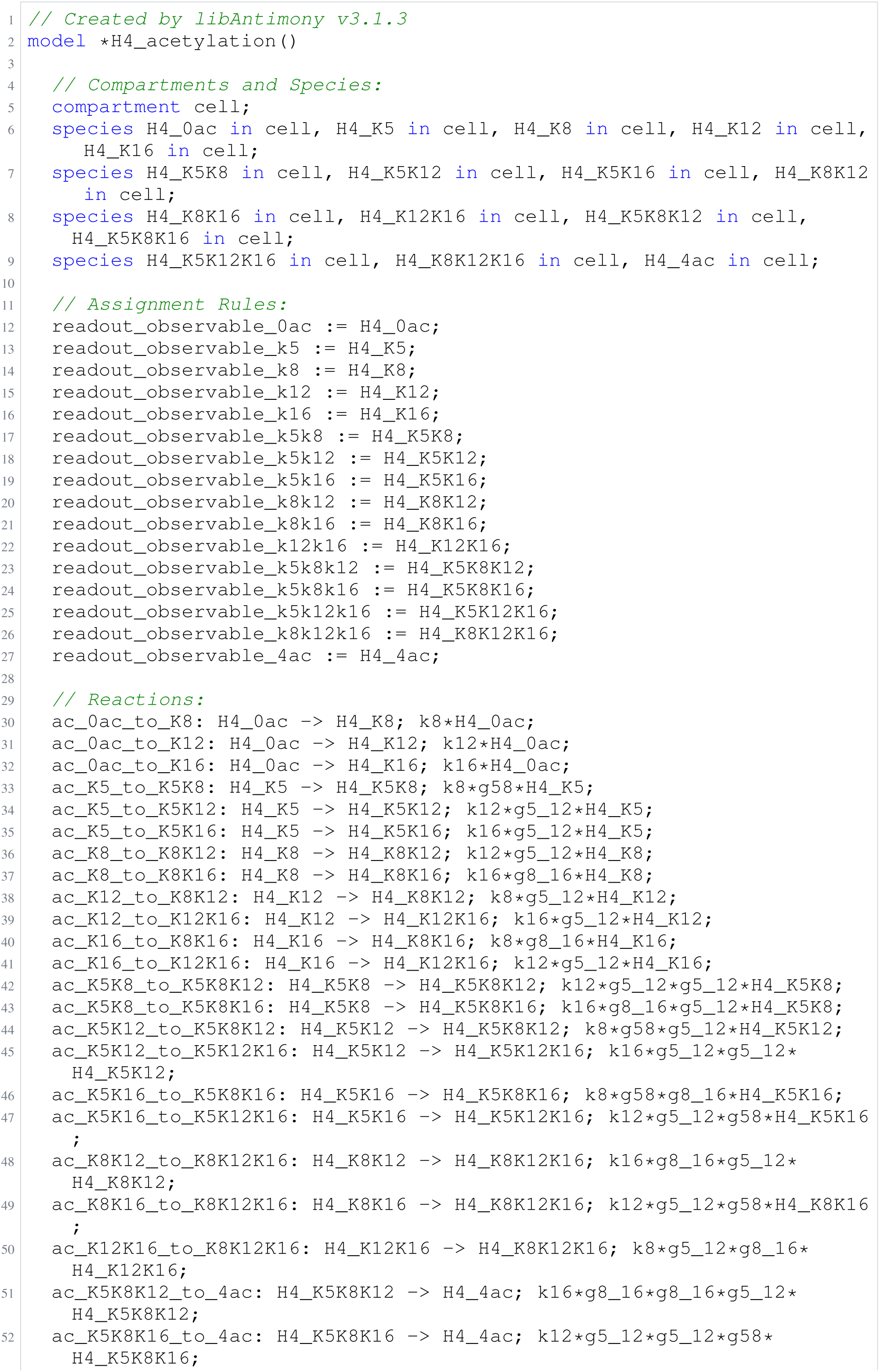
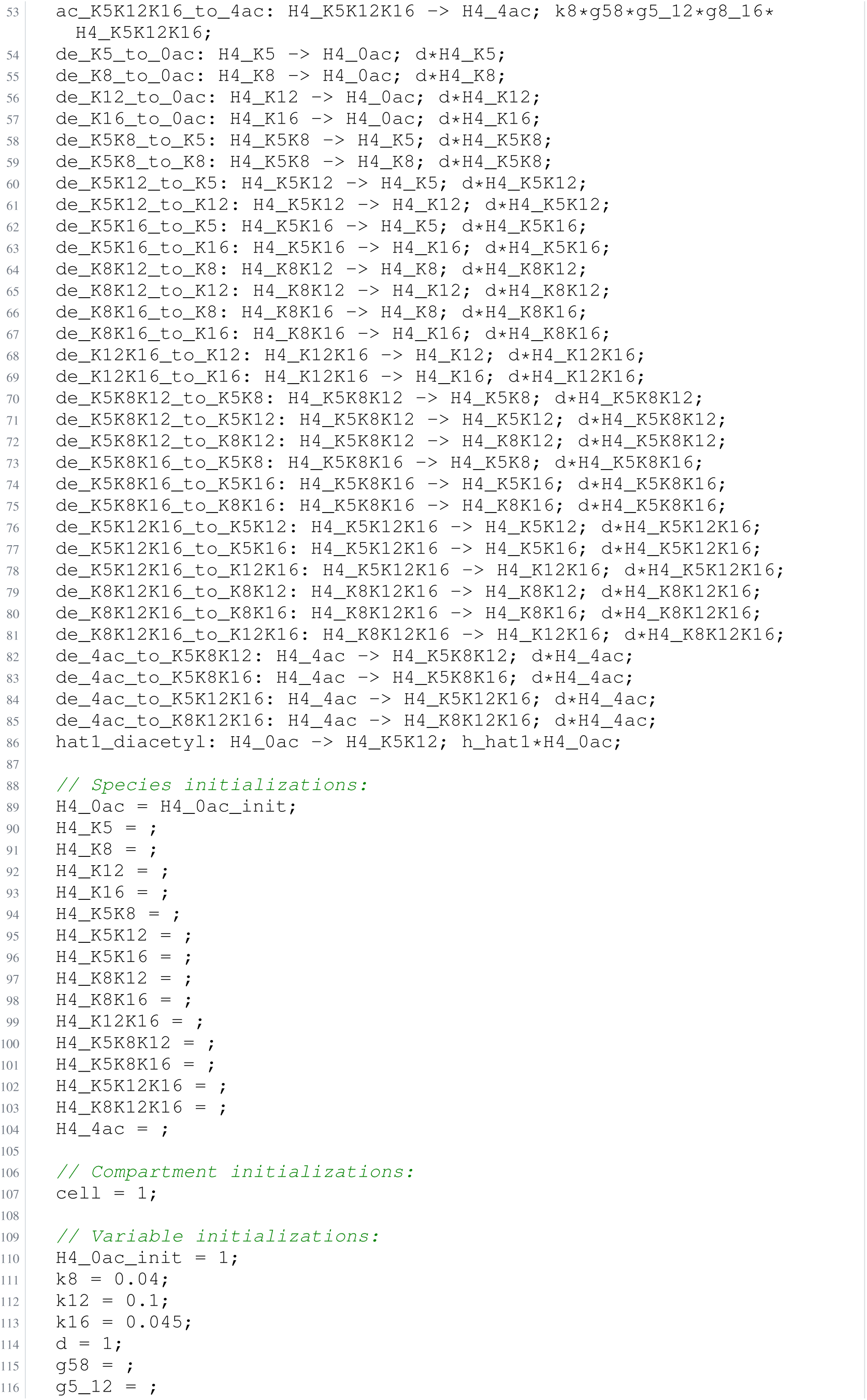
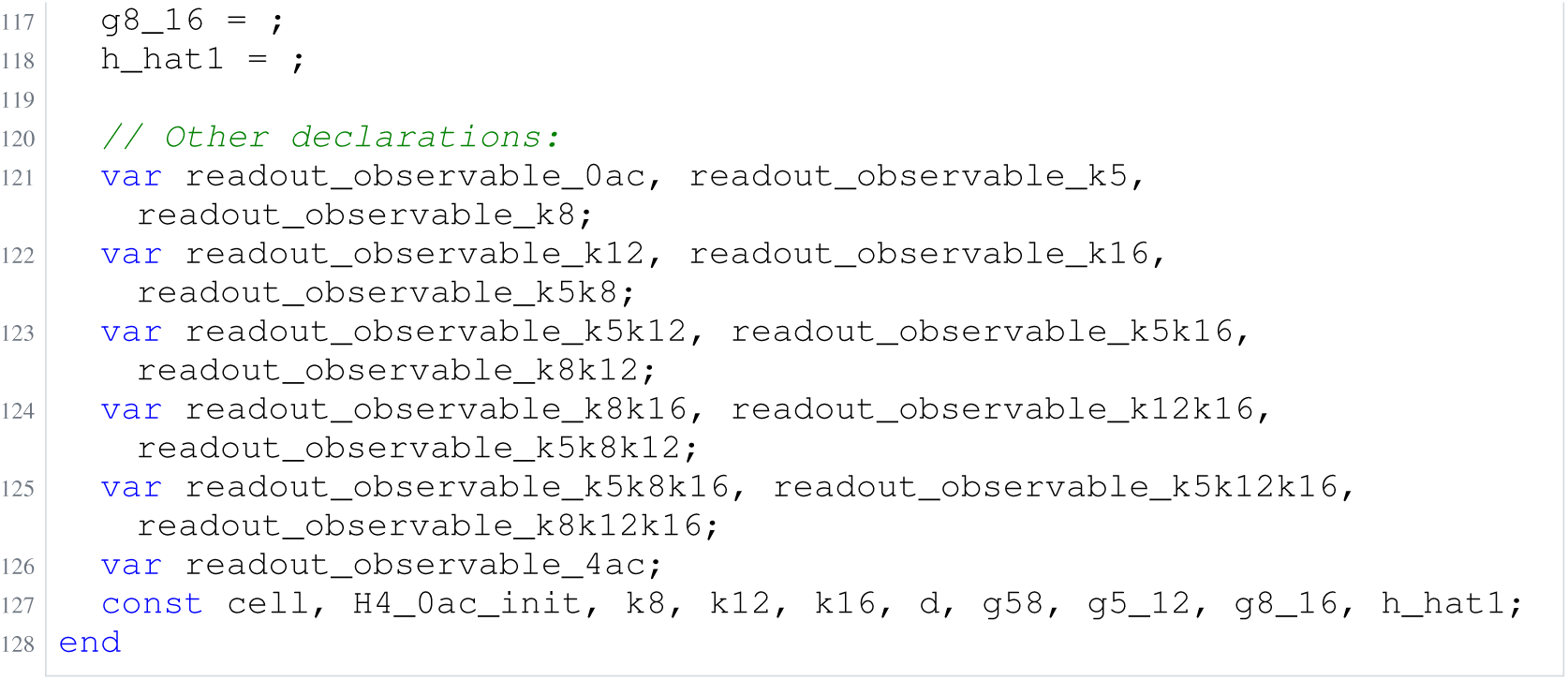
Archived Antimony for blasi_2; task 56836848_6; selected version 10.

**Listing 81:**
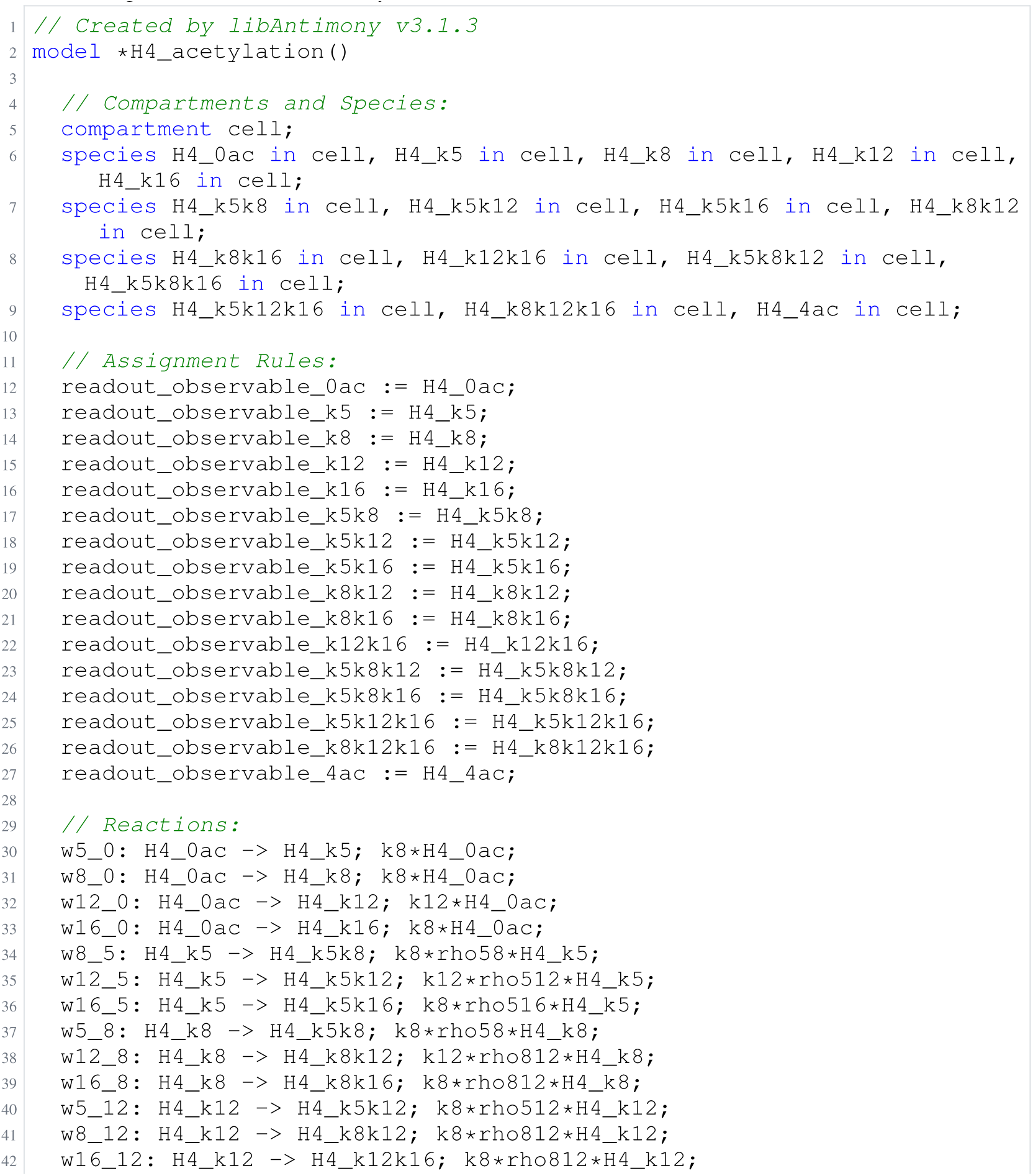
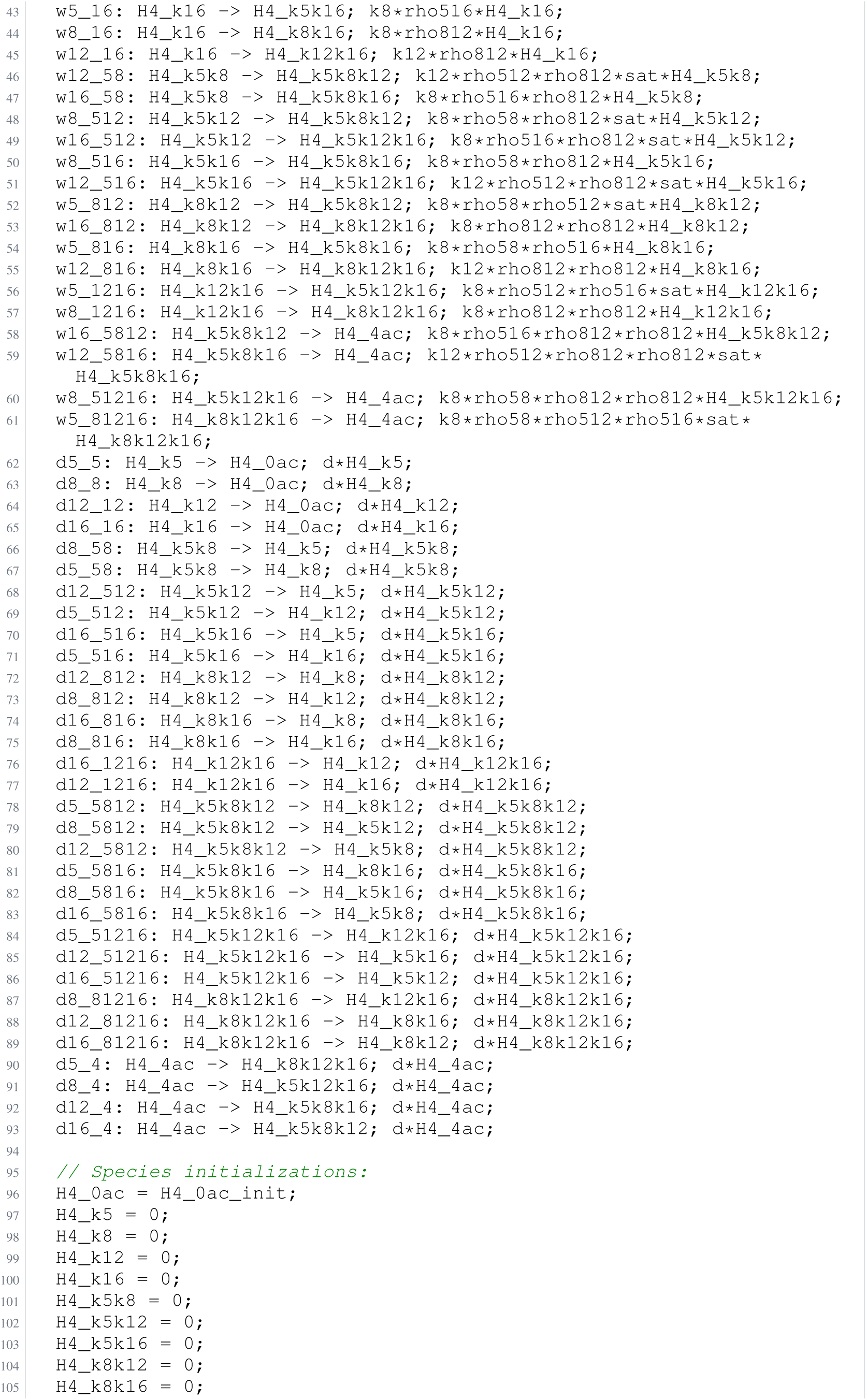
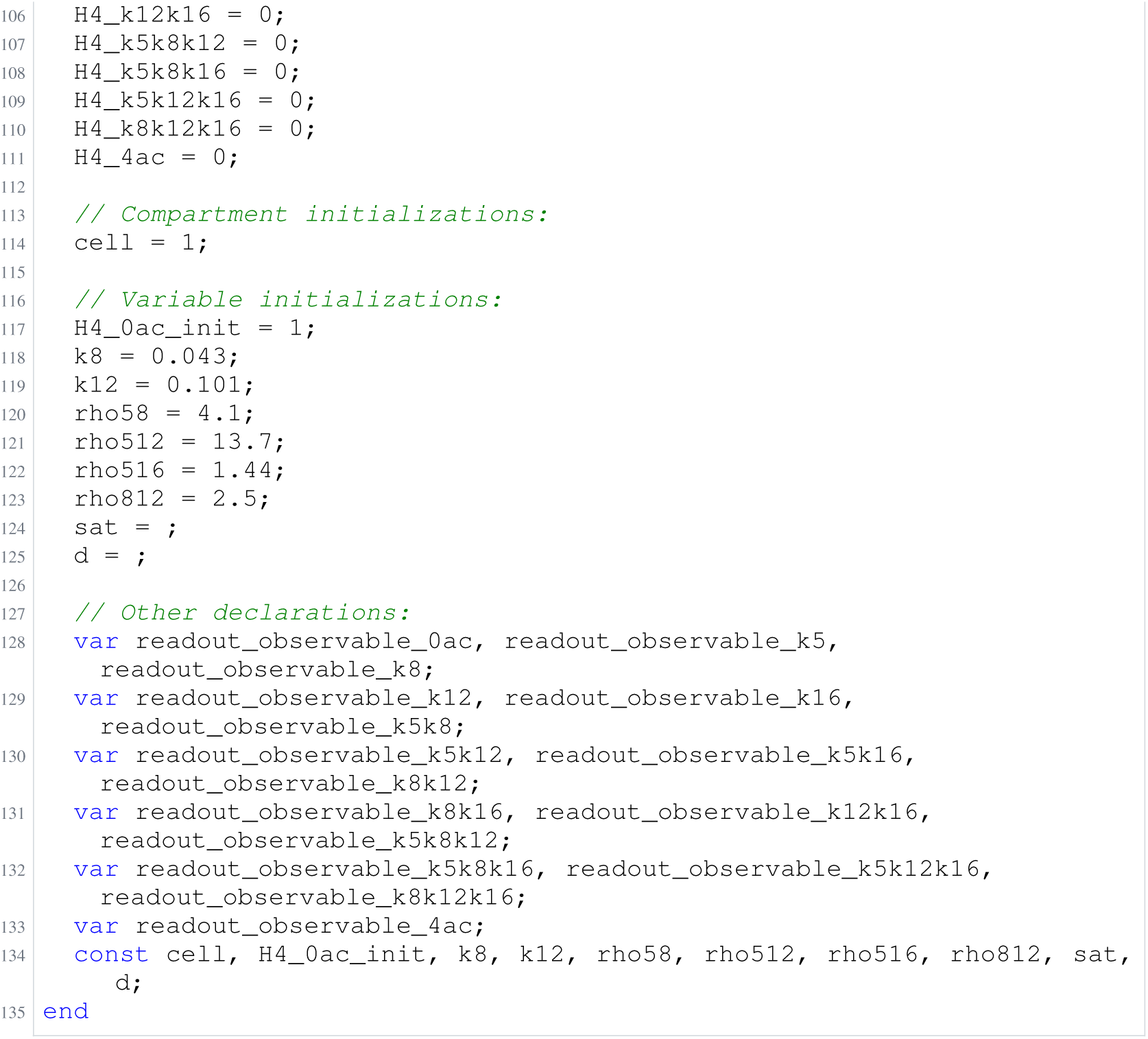
Archived Antimony for blasi_3; task 56836848_7; selected version 4.

**Listing 82:**
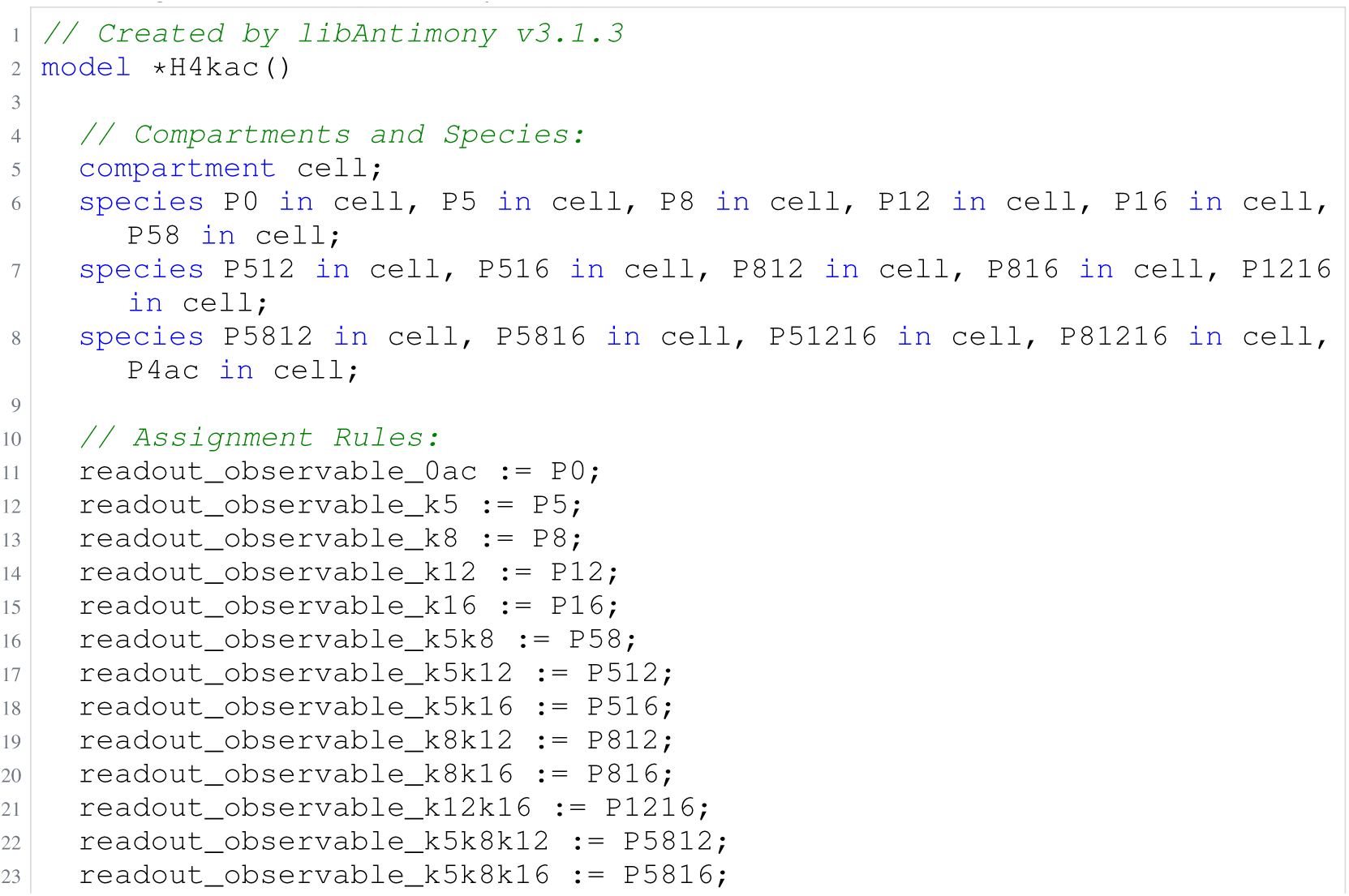
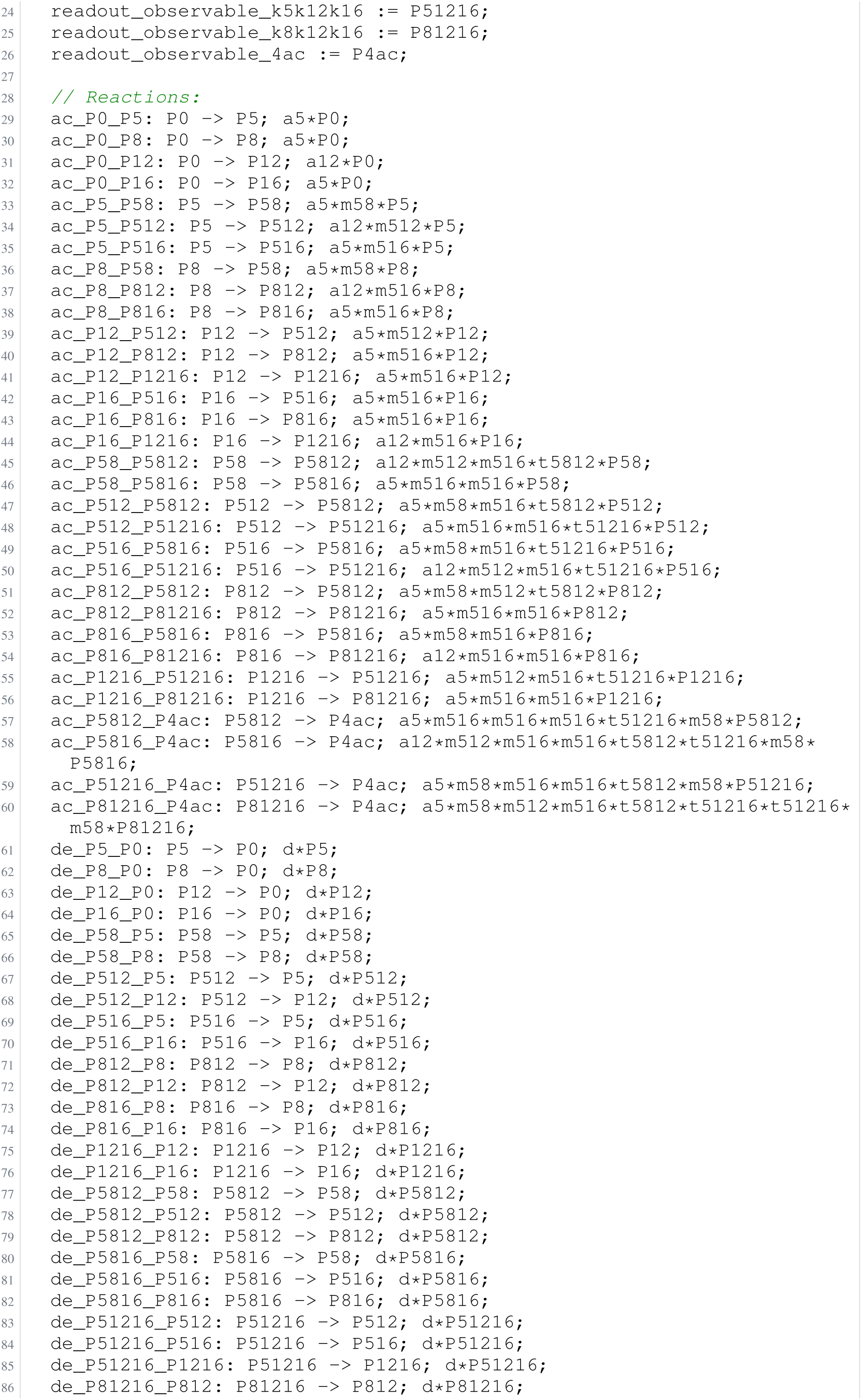
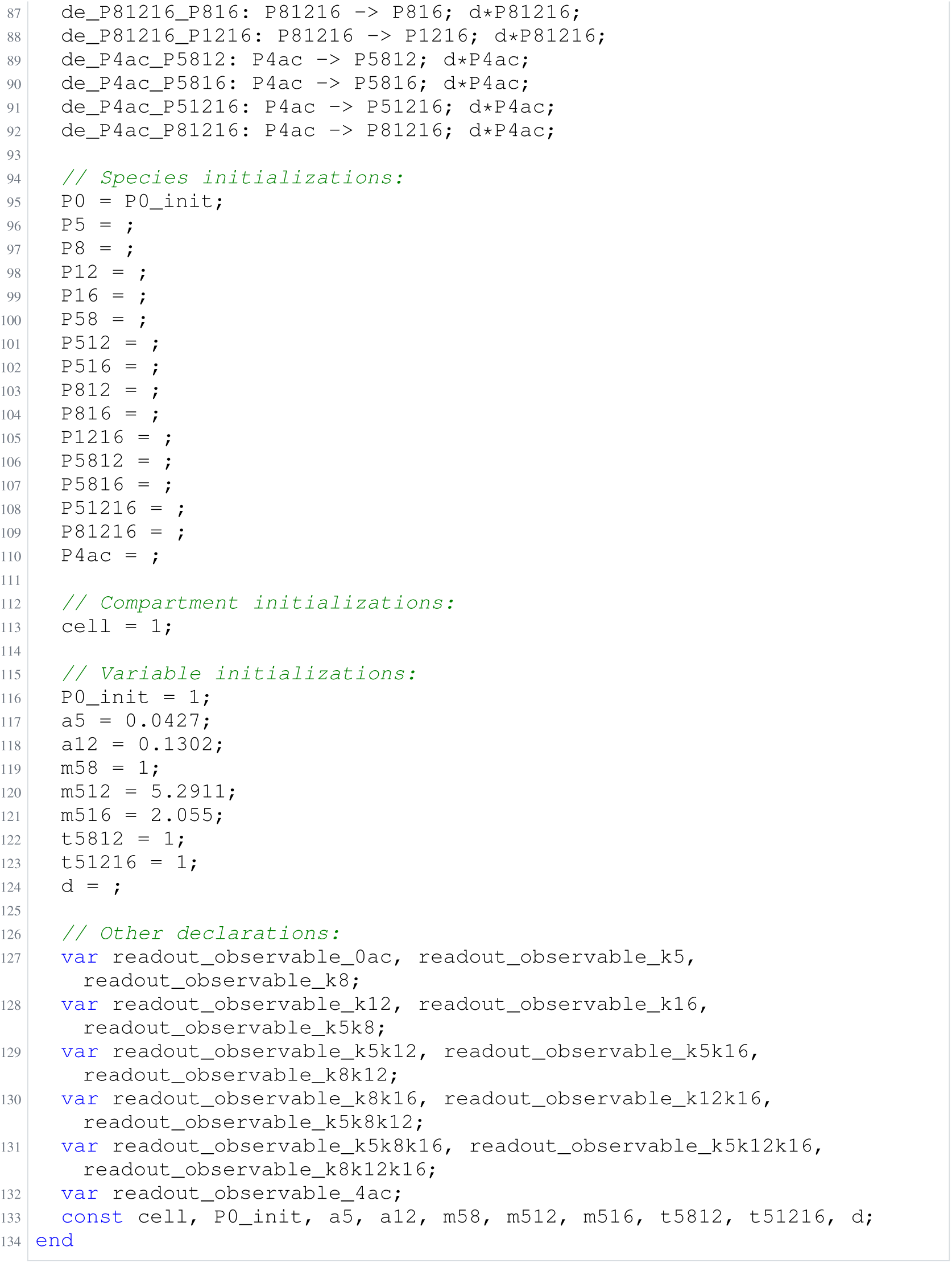
Archived Antimony for blasi_4; task 56836848_8; selected version 10.

**Listing 83:**
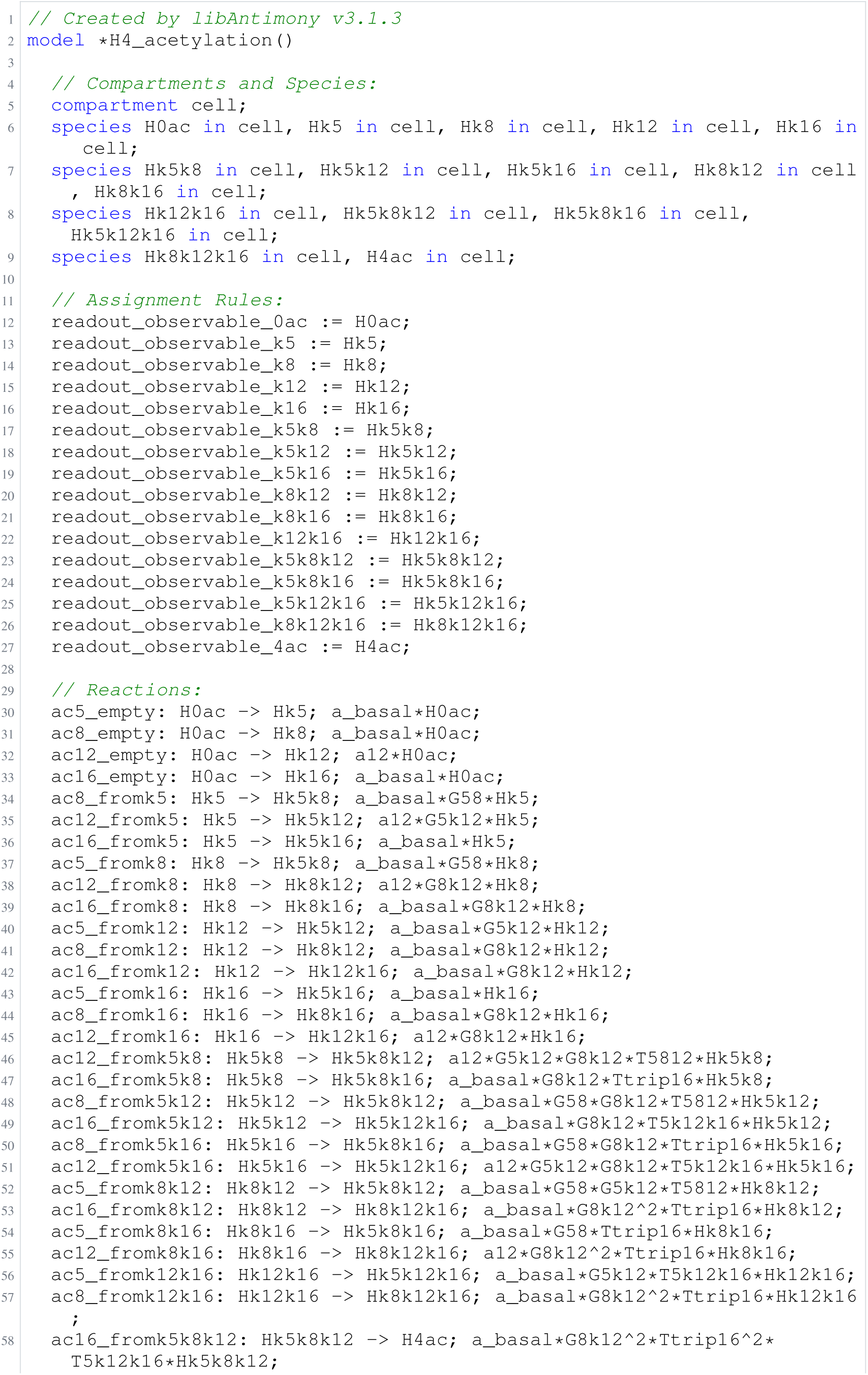
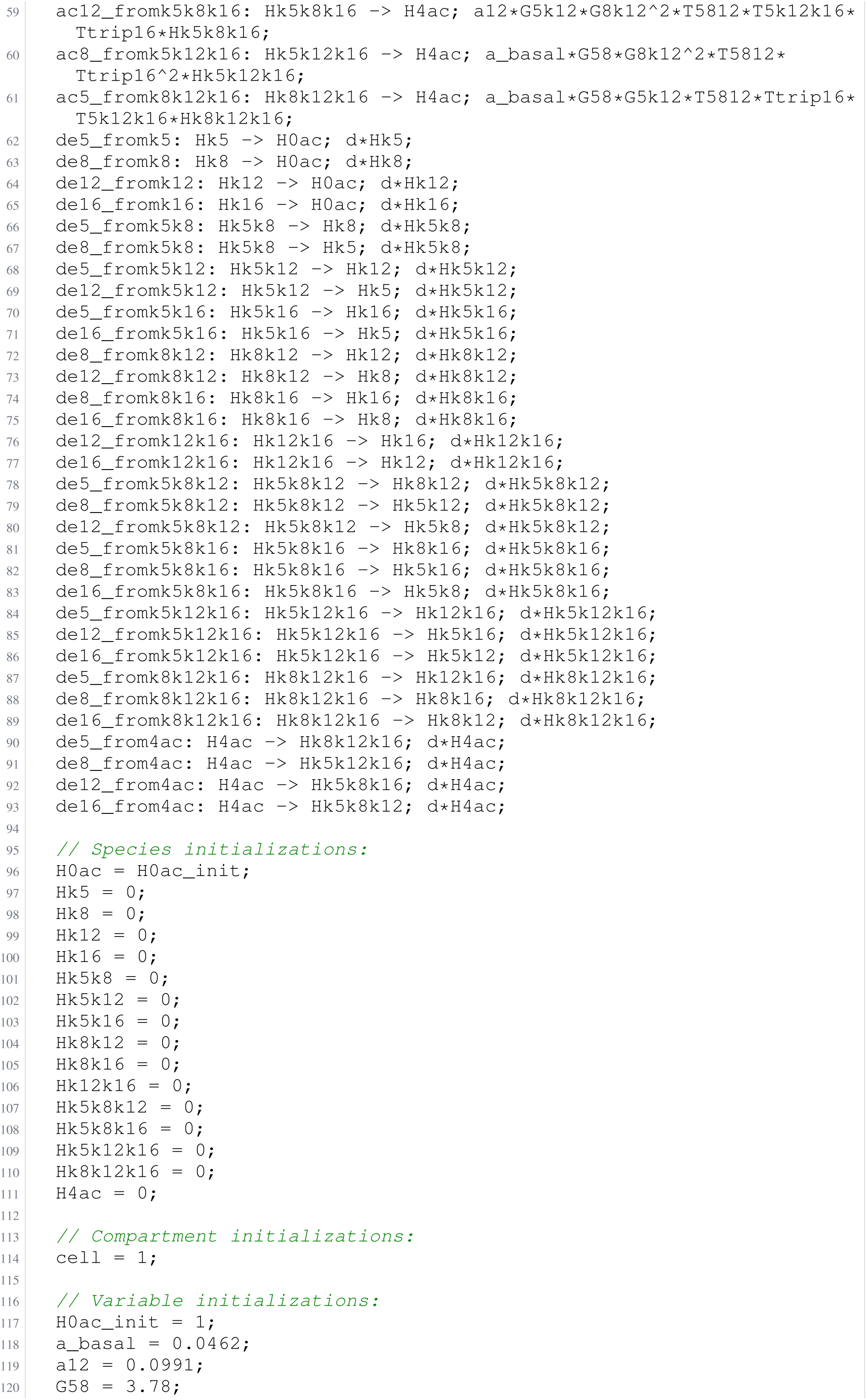
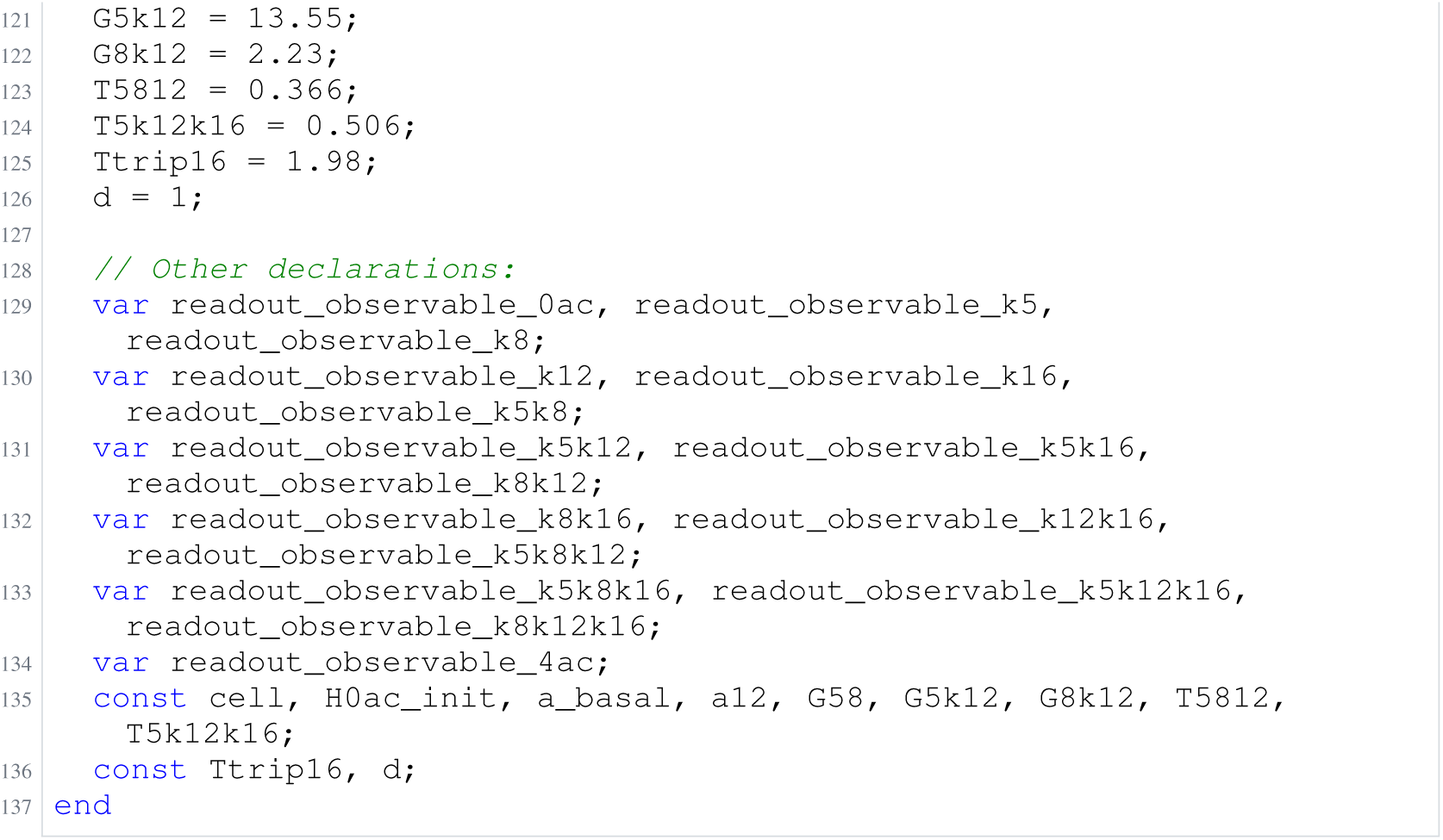
Archived Antimony for blasi_5; task 56836848_9; selected version 10.

### H.12 Schwen

**Listing 84:**
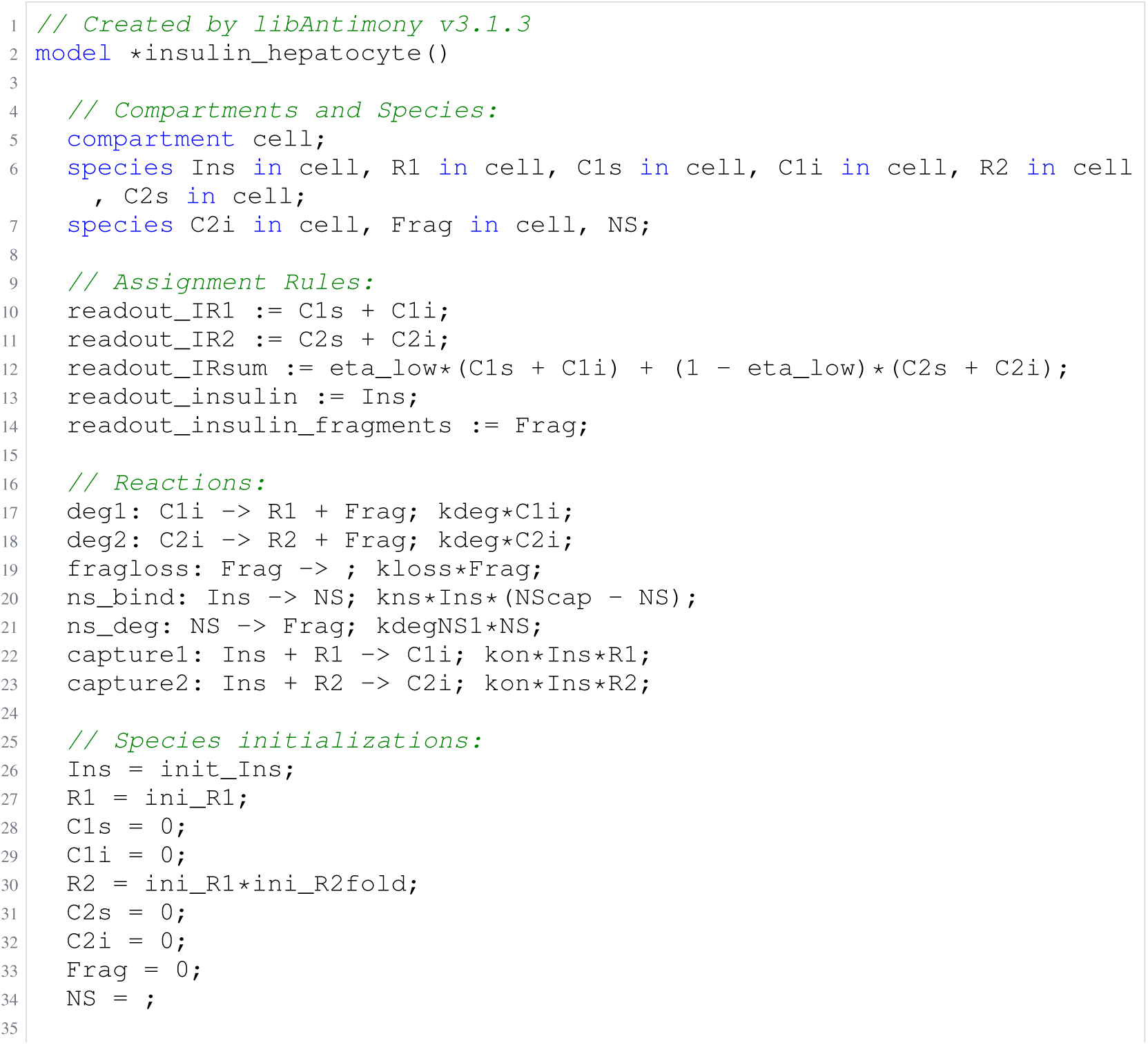
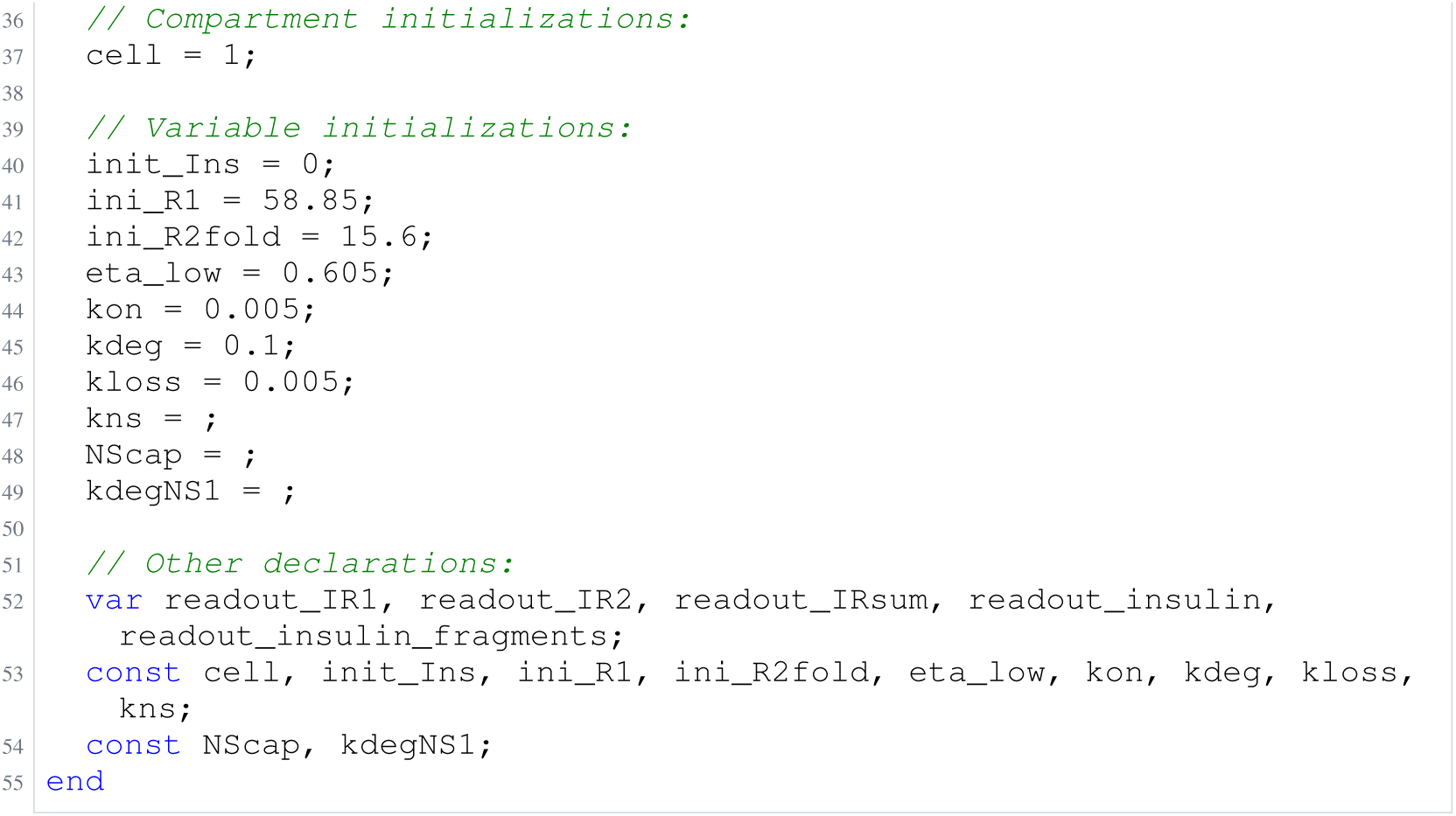
Archived Antimony for schwen_1; task 56836812_0; selected version 8.

**Listing 85:**
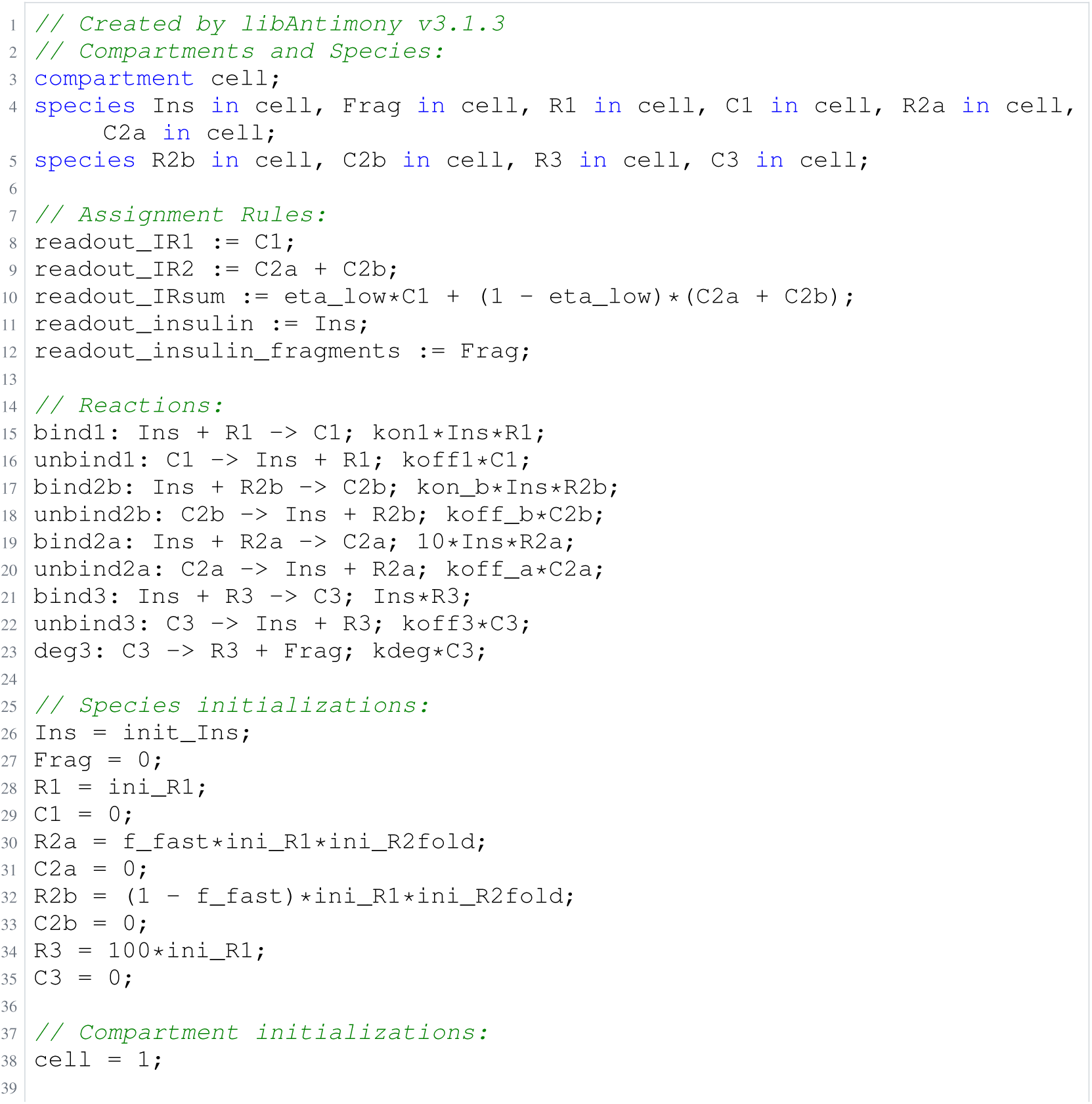
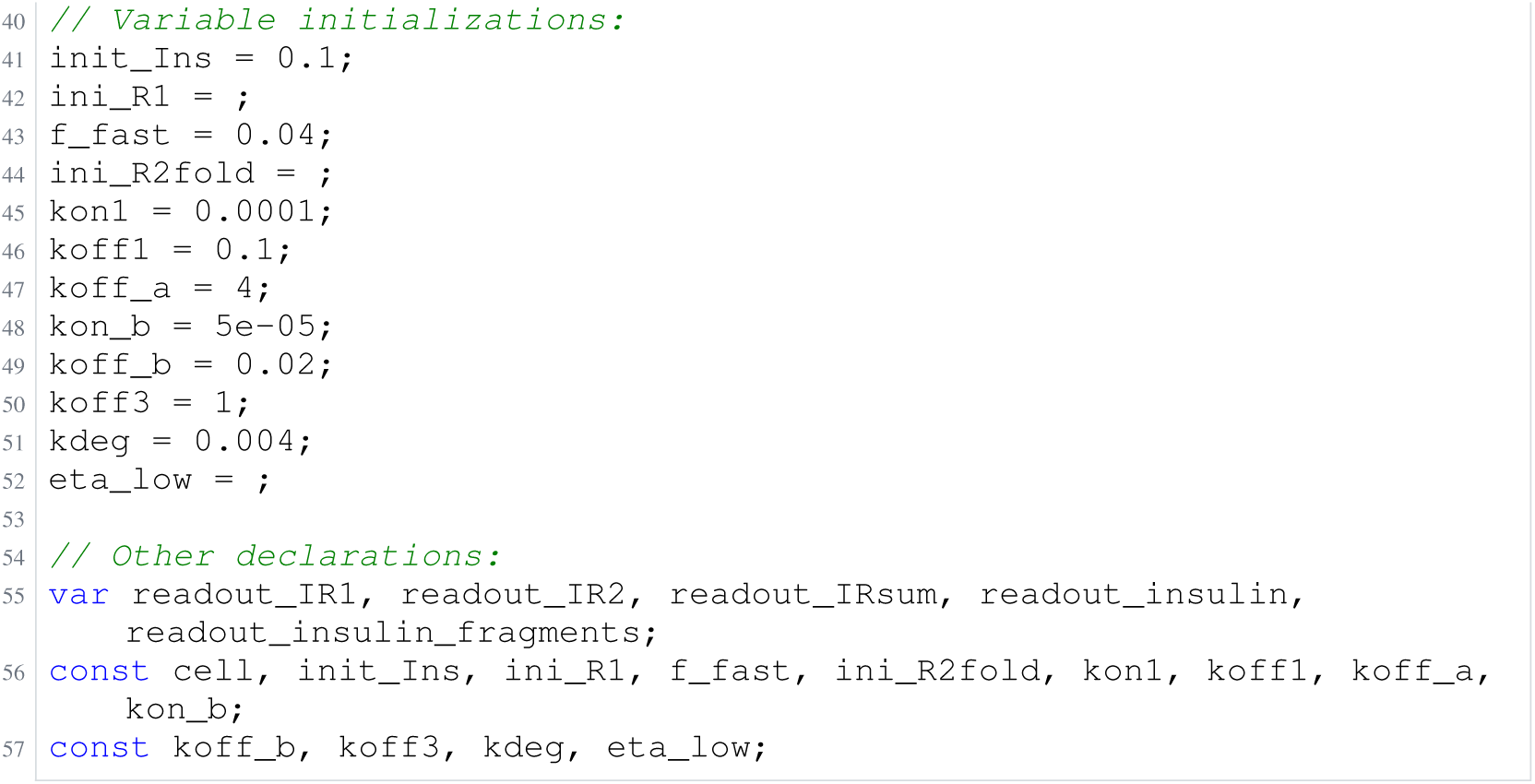
Archived Antimony for schwen_2; task 56836812_1; selected version 18.

**Listing 86:**
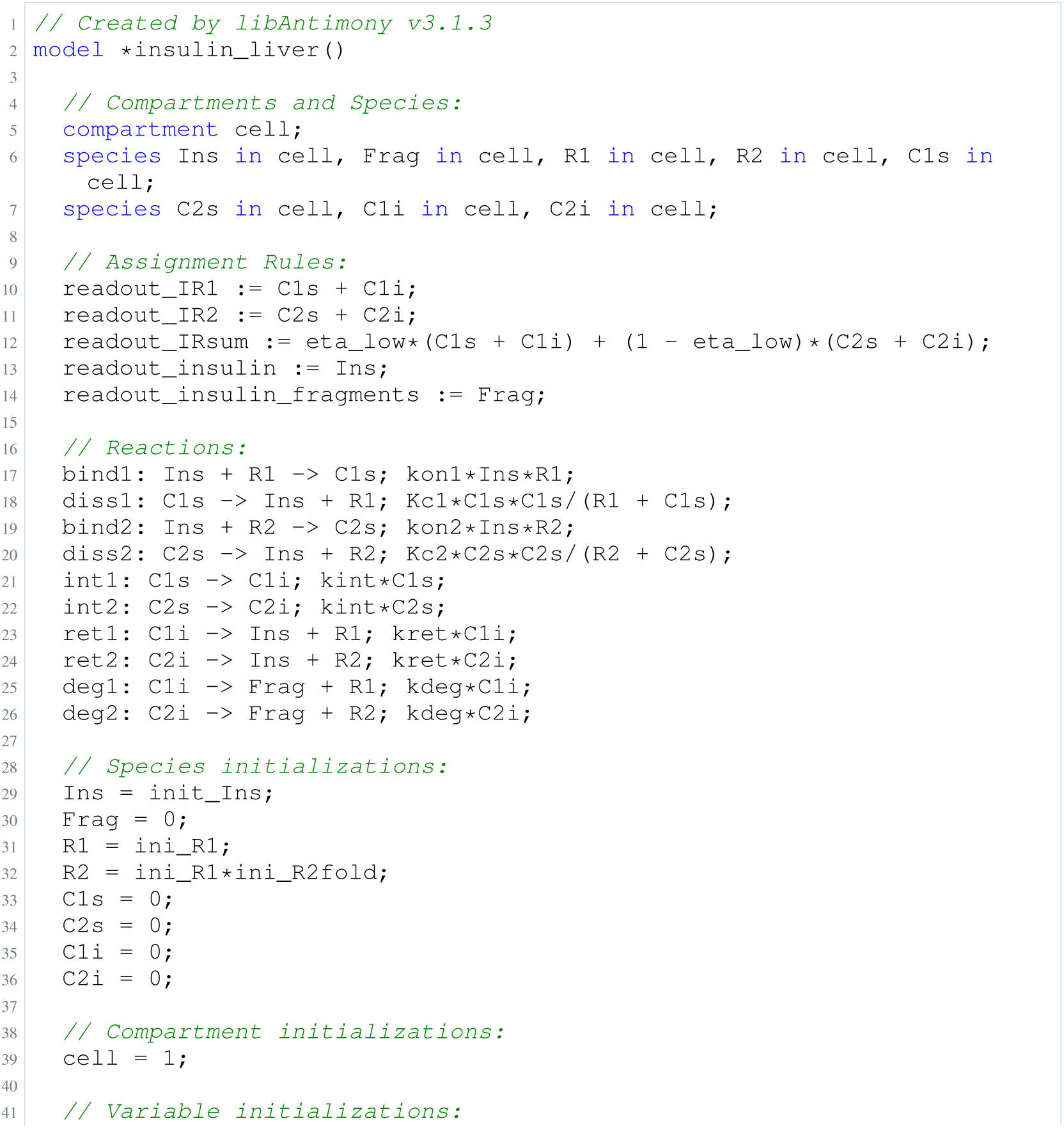
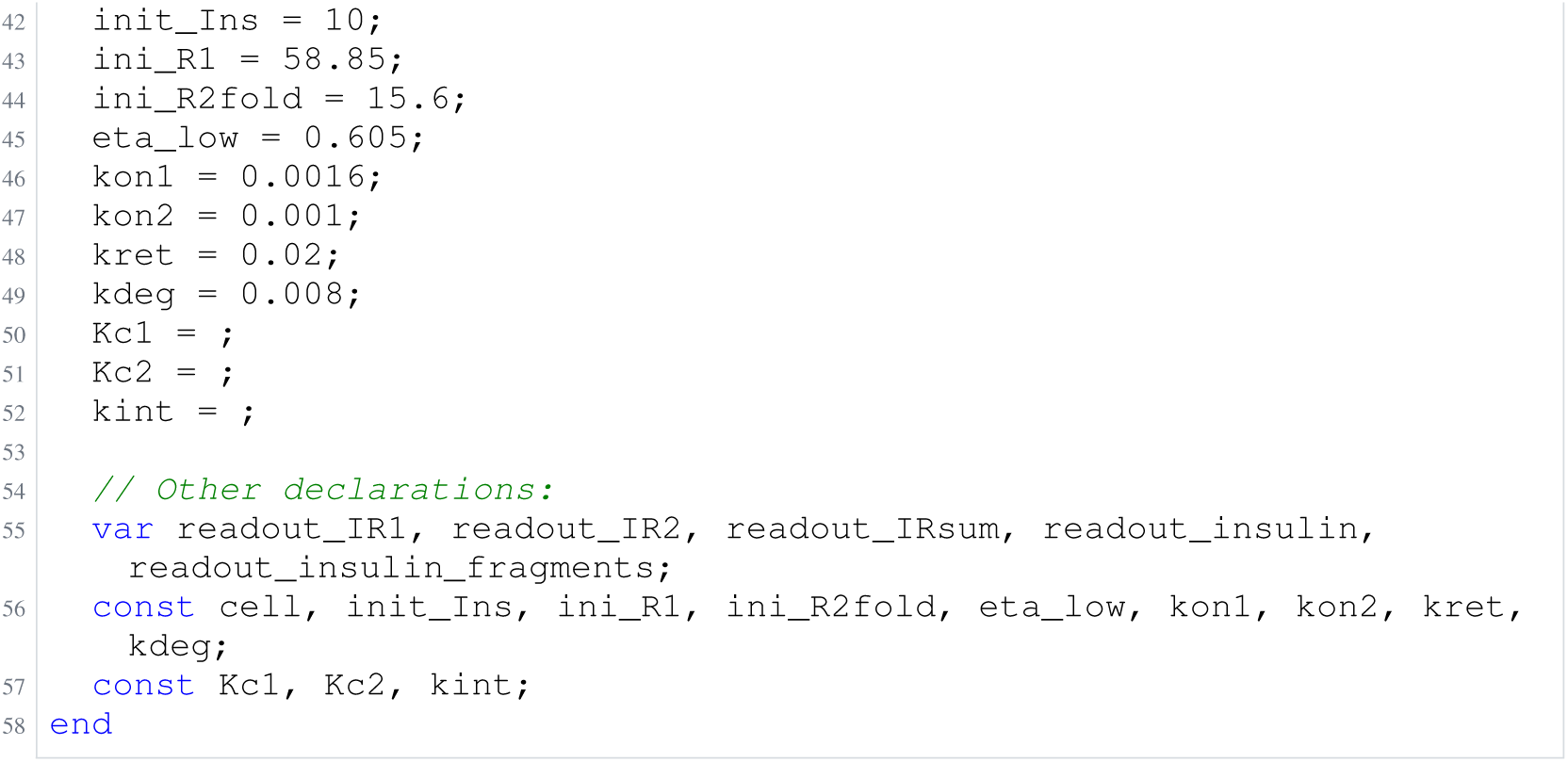
Archived Antimony for schwen_3; task 56836812_2; selected version 7.

**Listing 87:**
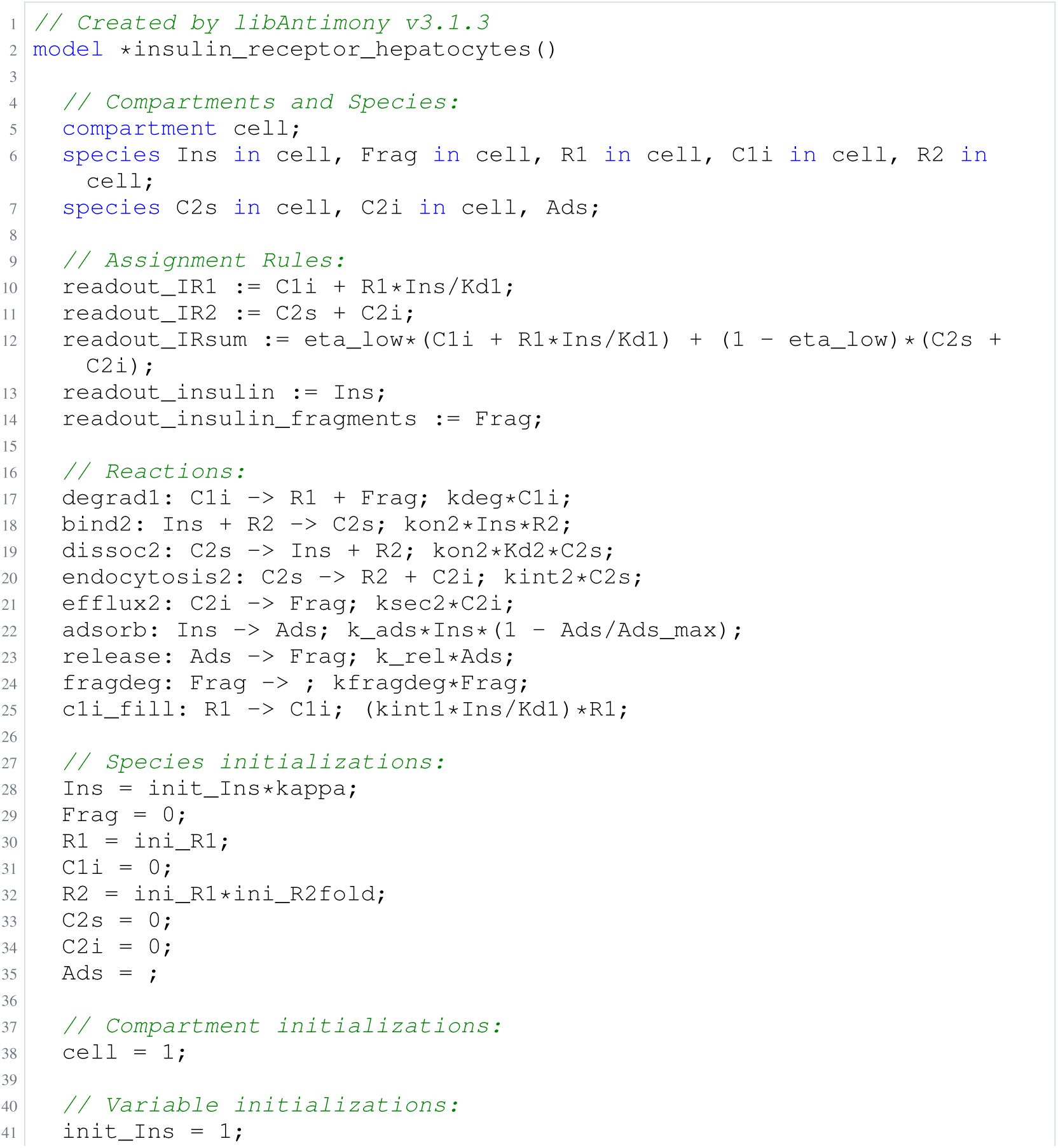
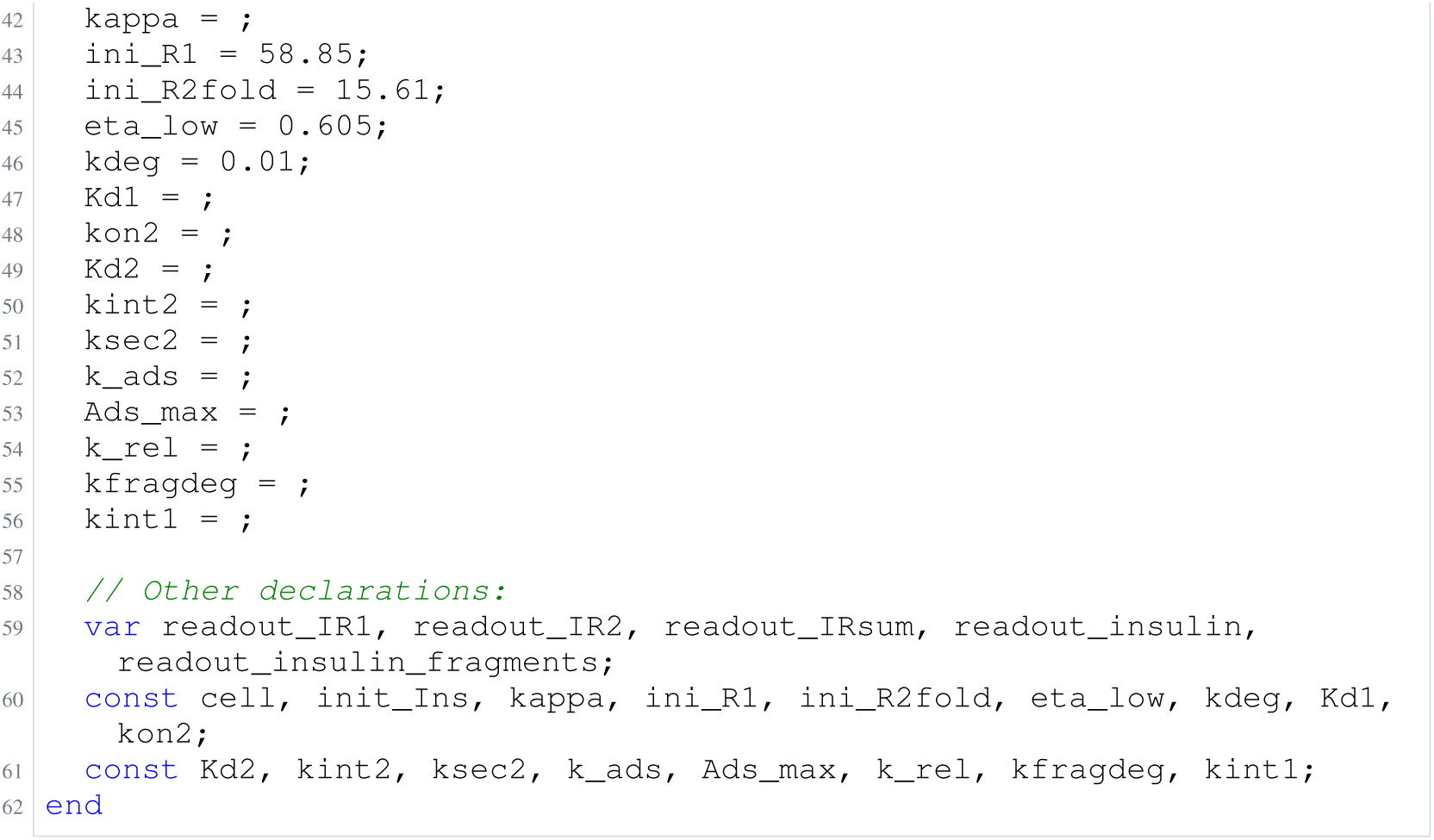
Archived Antimony for schwen_4; task 56836812_3; selected version 12.

**Listing 88:**
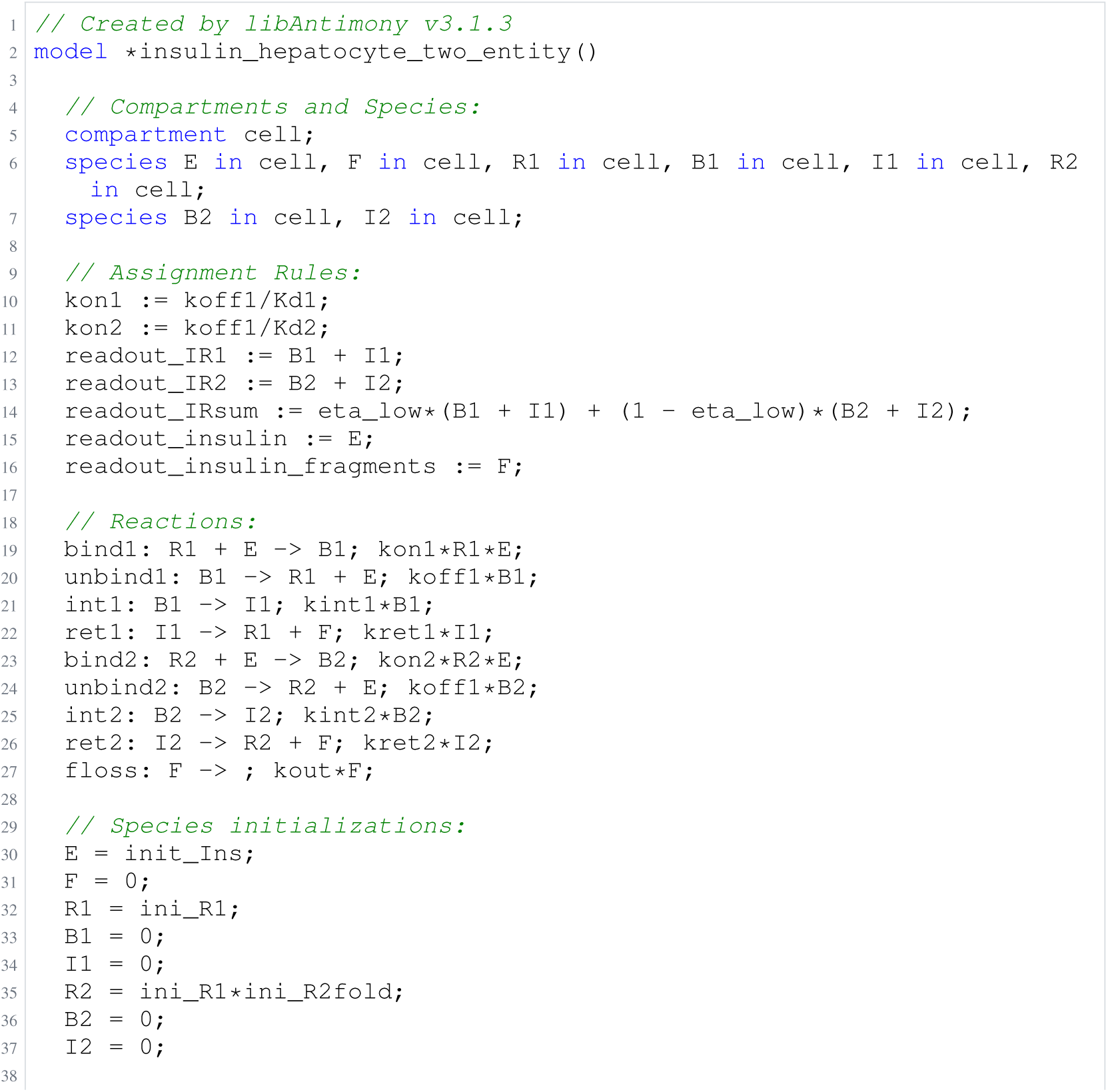
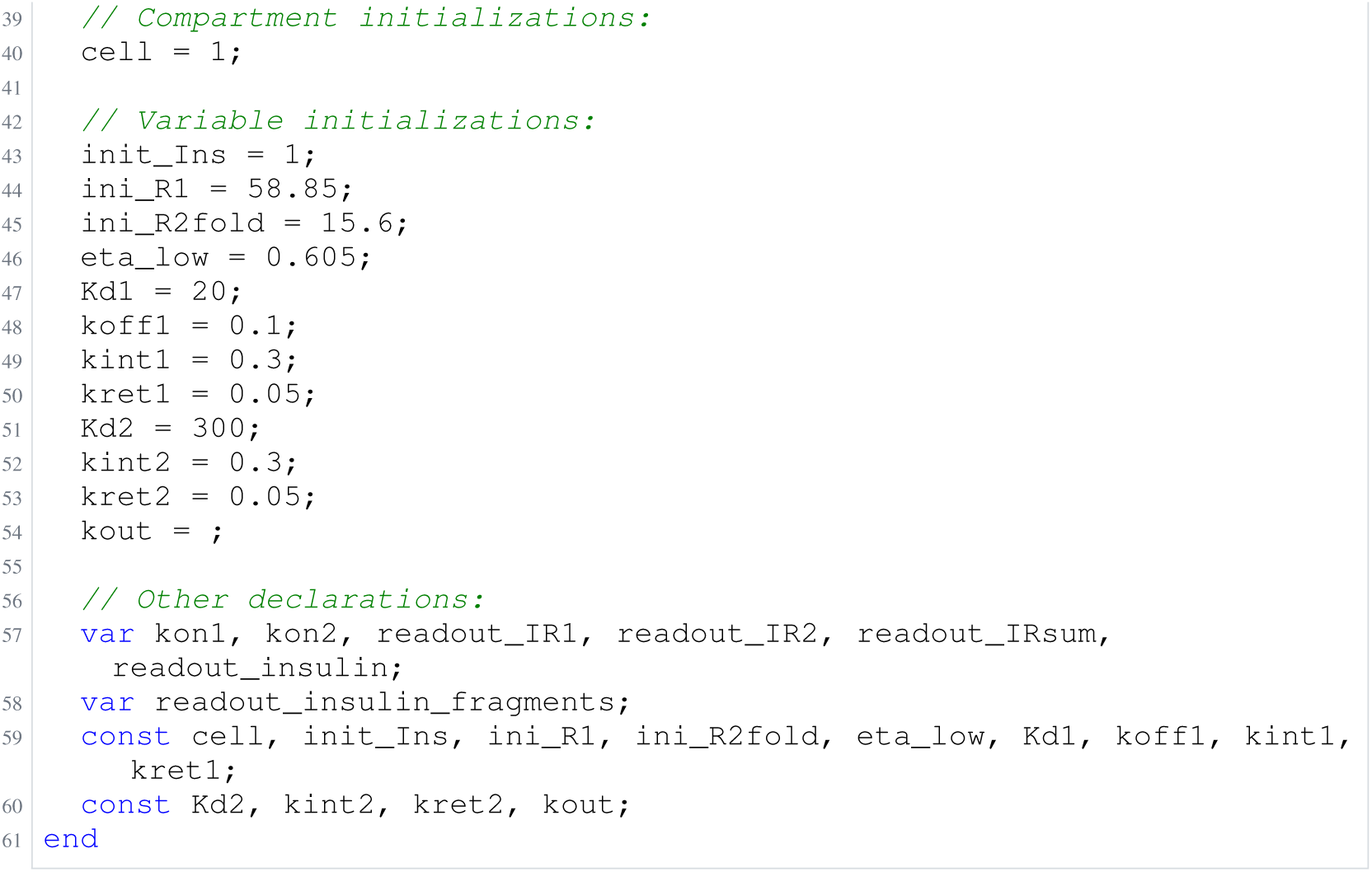
Archived Antimony for schwen_5; task 56836812_4; selected version 3.

### H.13 Bachmann

**Listing 89:**
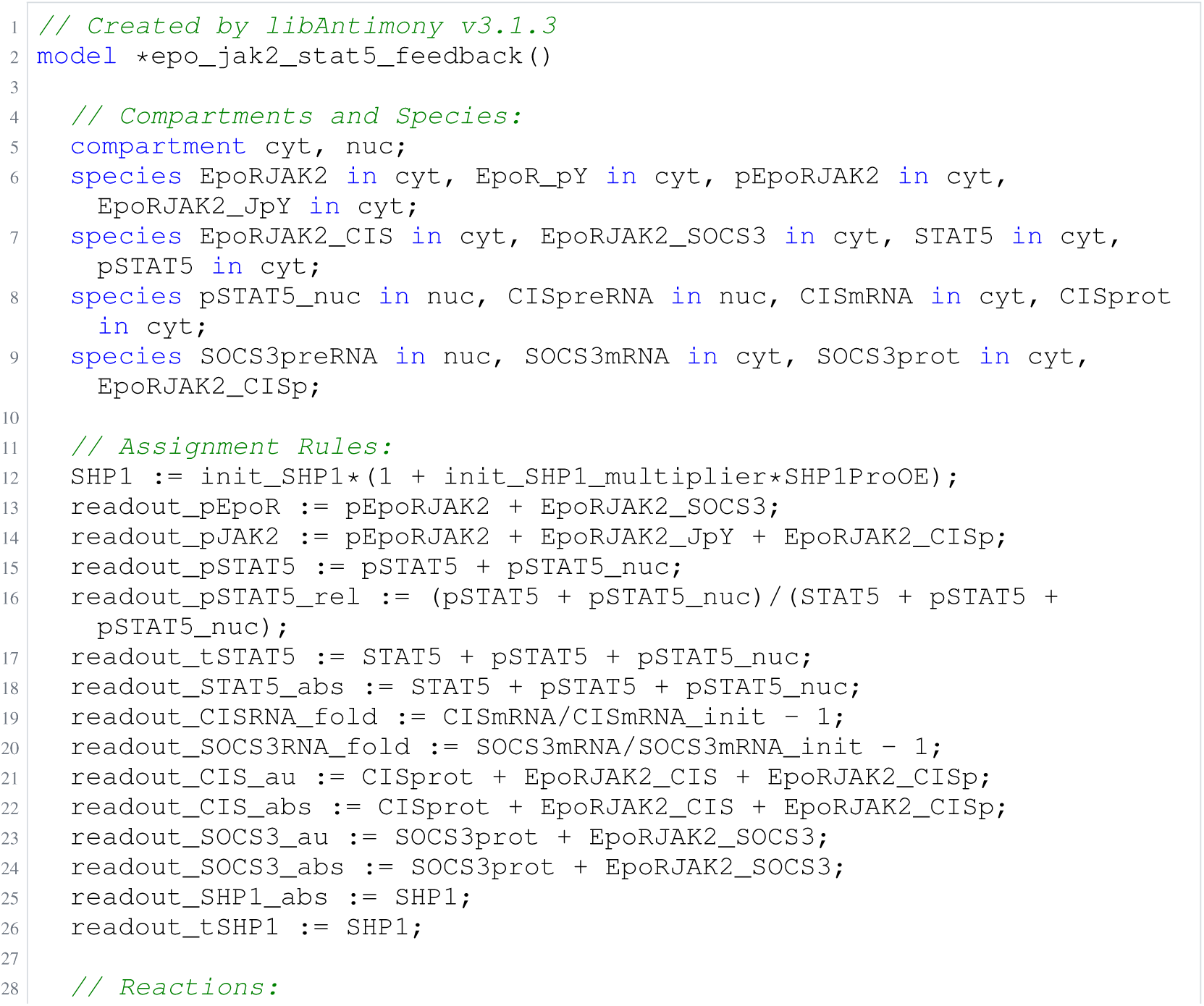
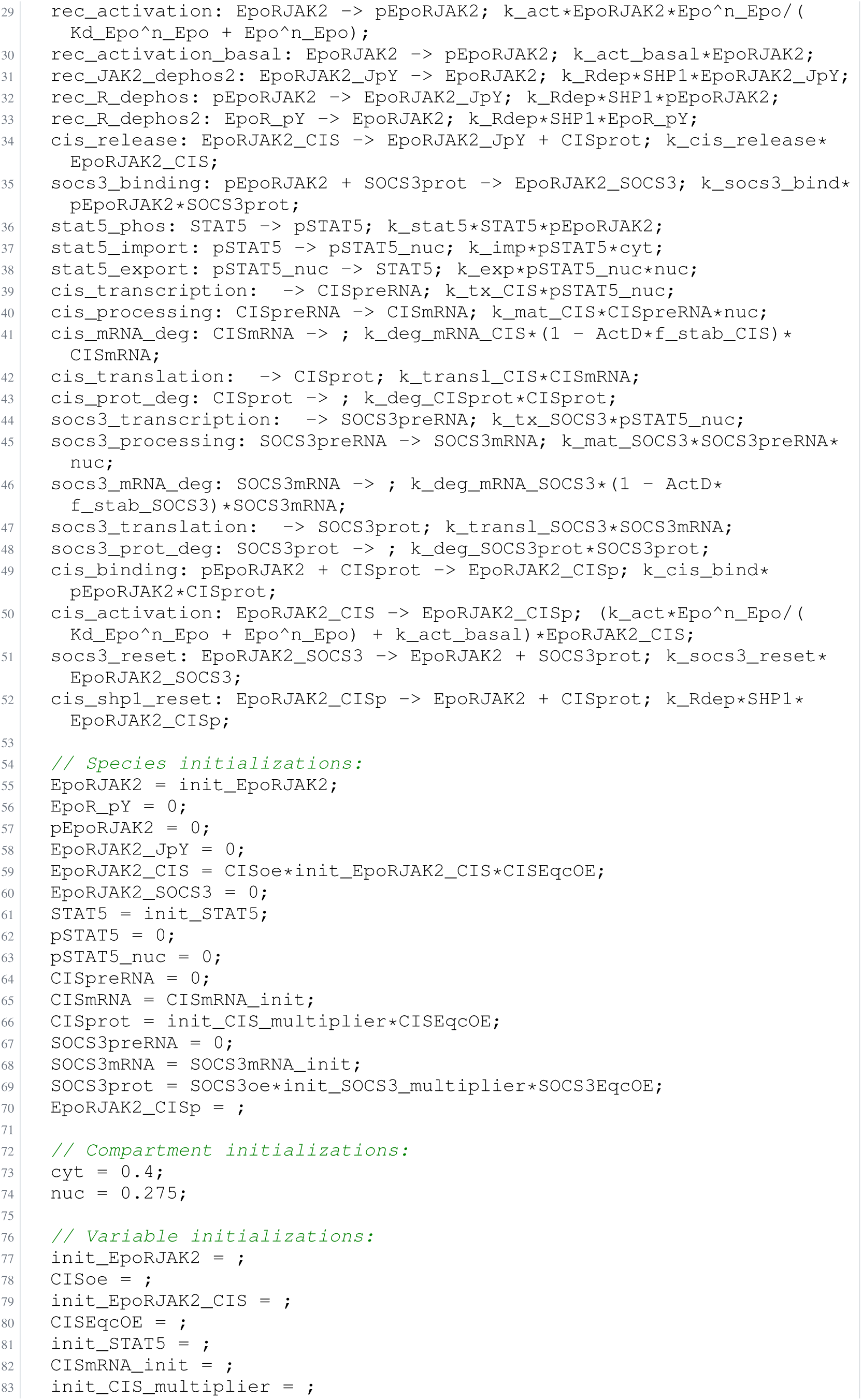
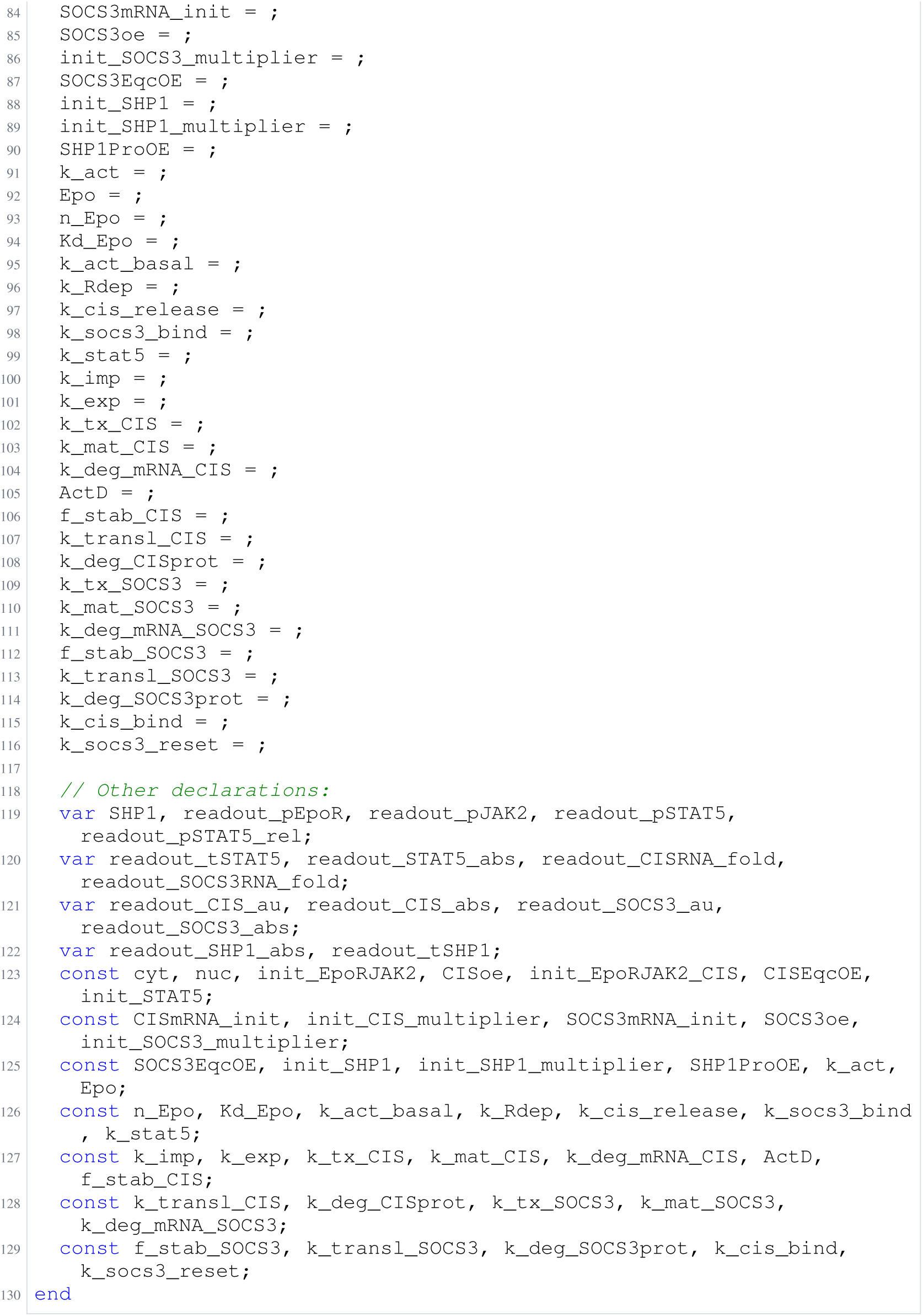
Archived Antimony for bachmann_1; task 56836792_5; selected version 11.

**Listing 90:**
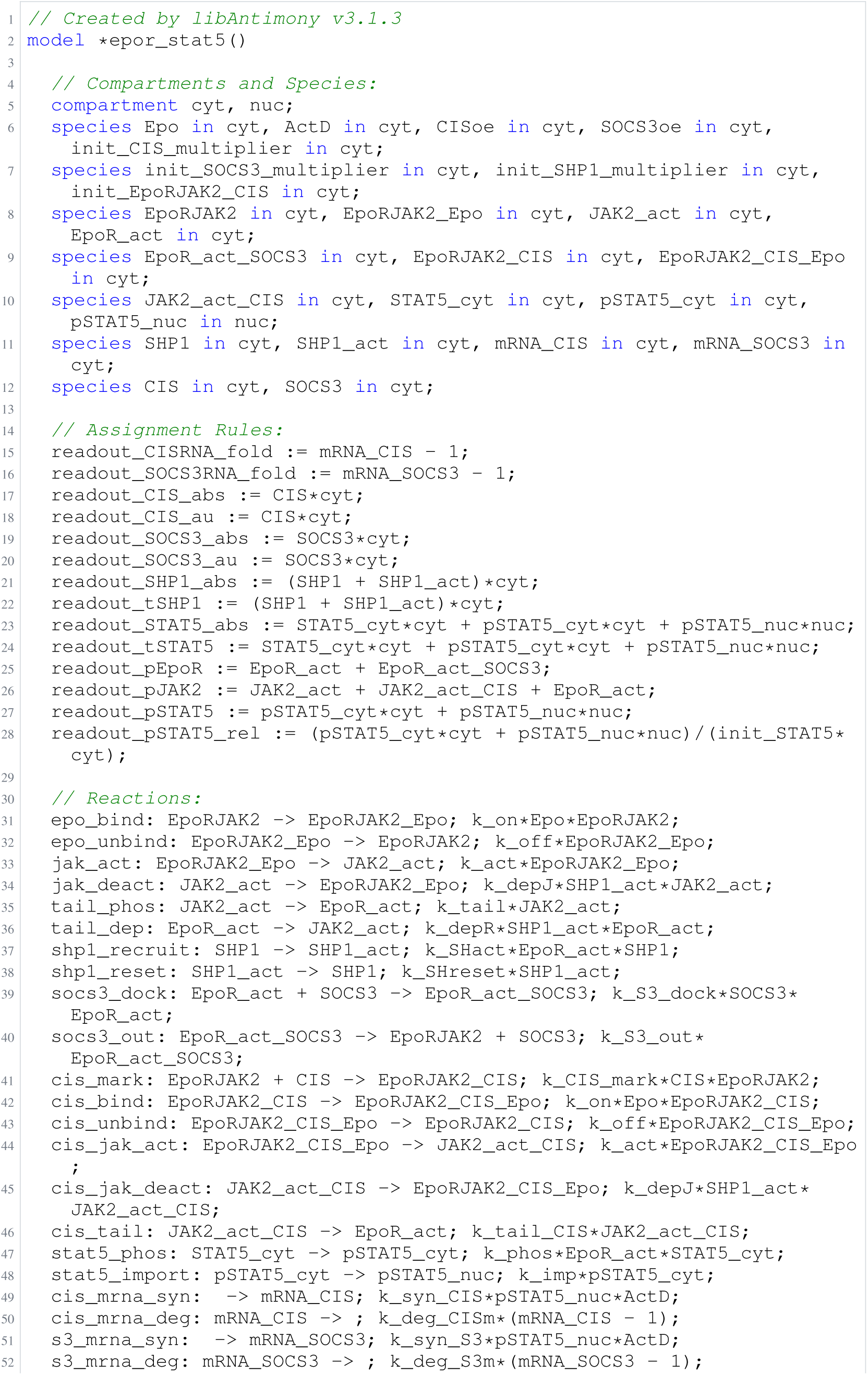
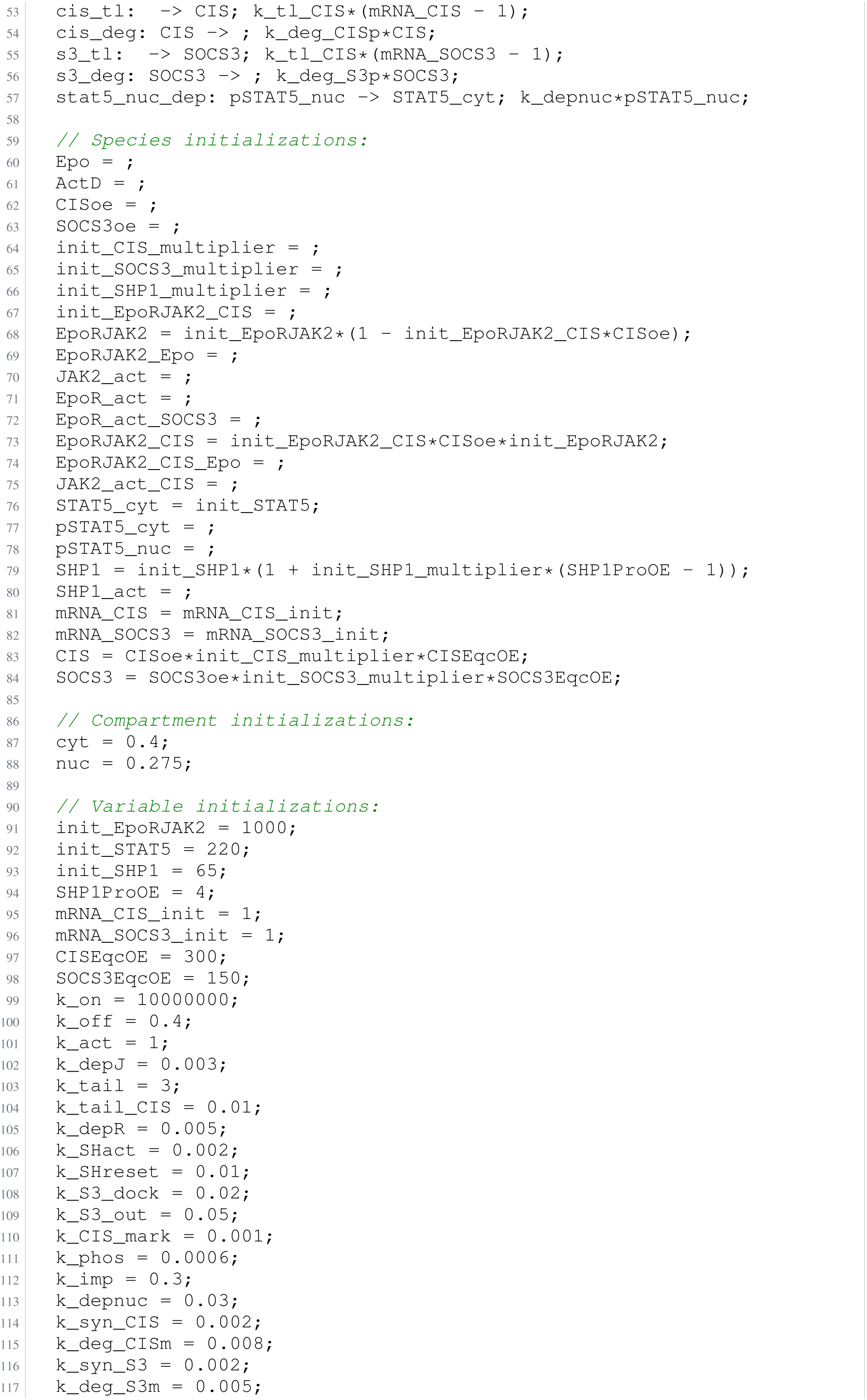
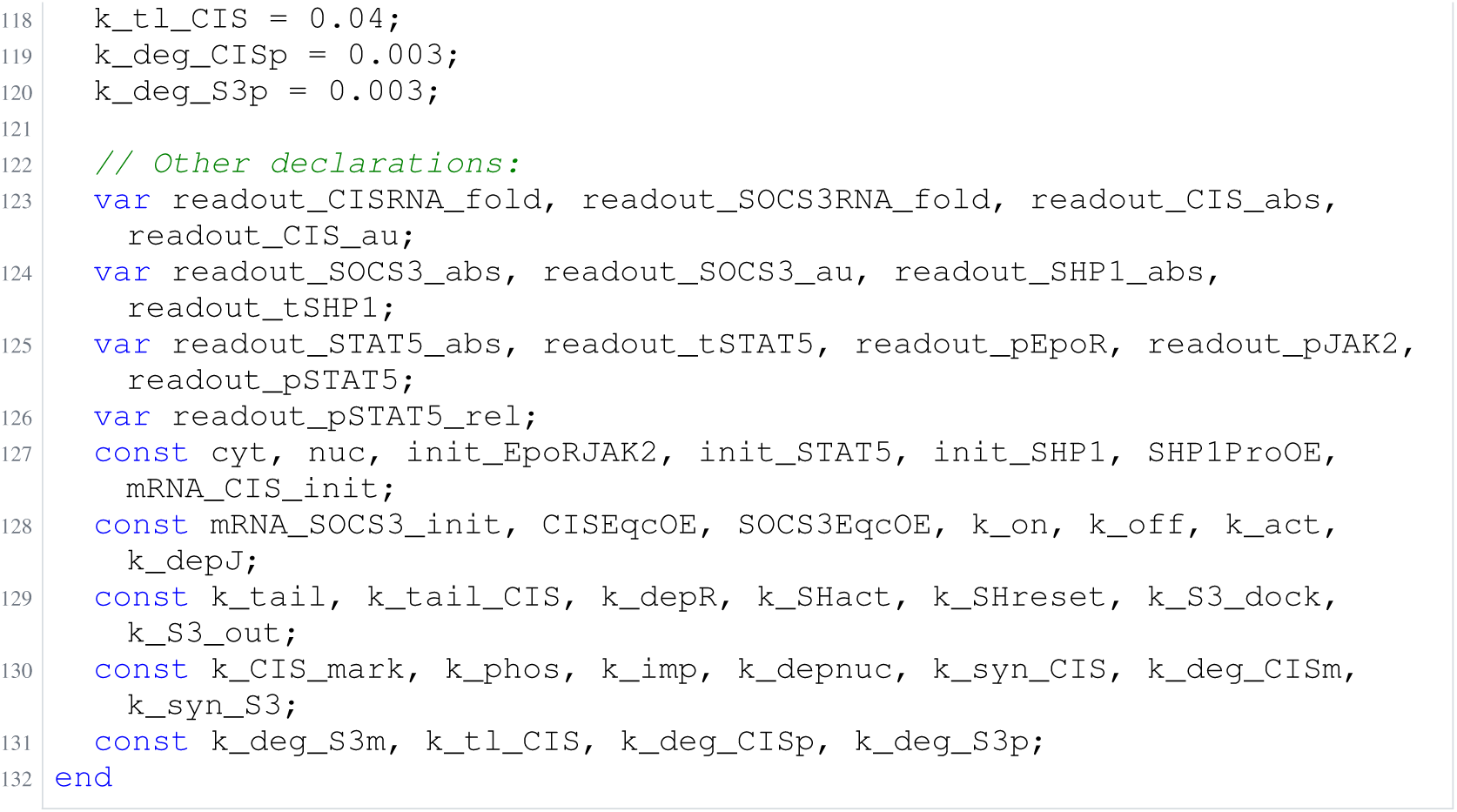
Archived Antimony for bachmann_2; task 56836792_6; selected version 9.

**Listing 91:**
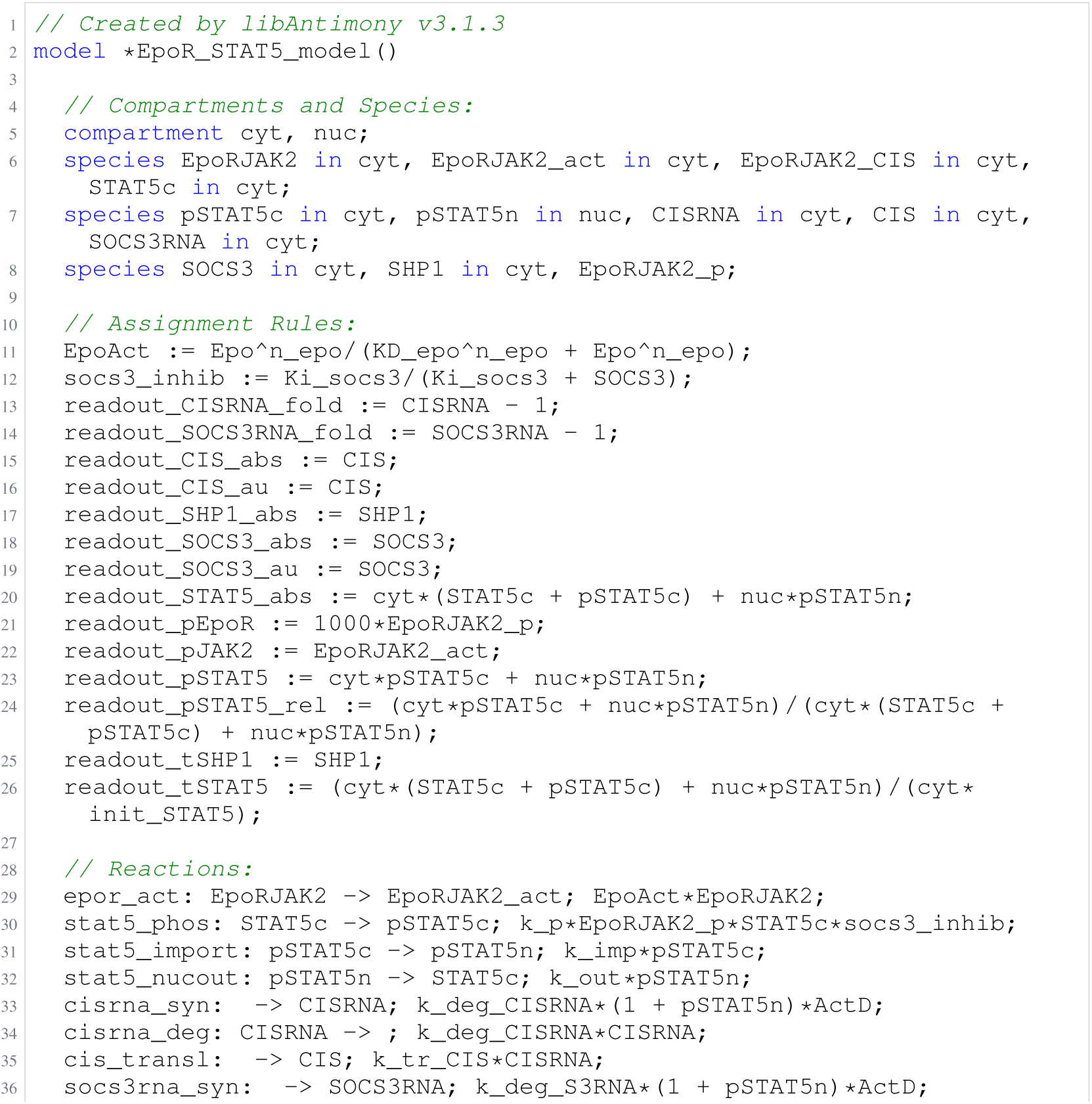
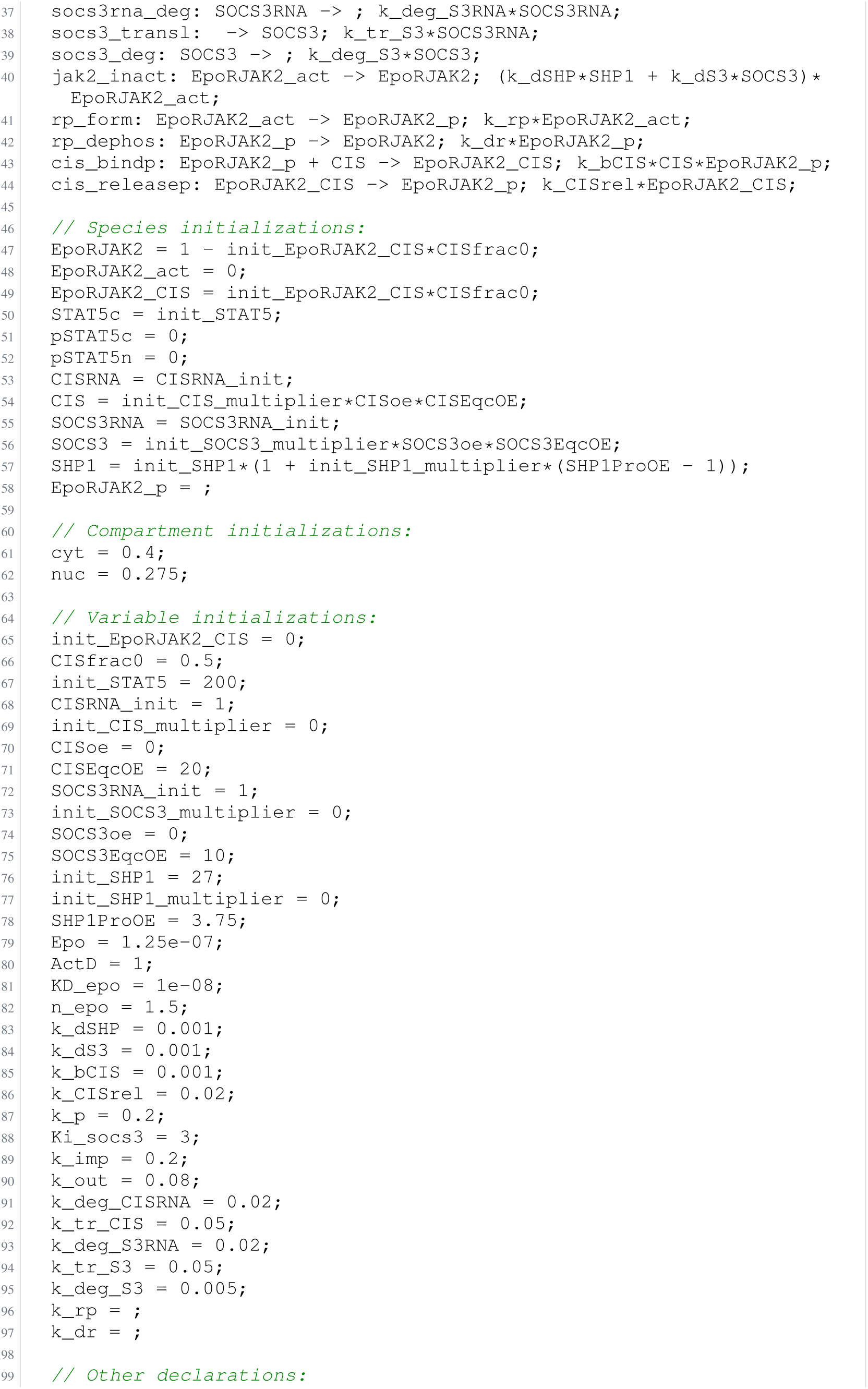
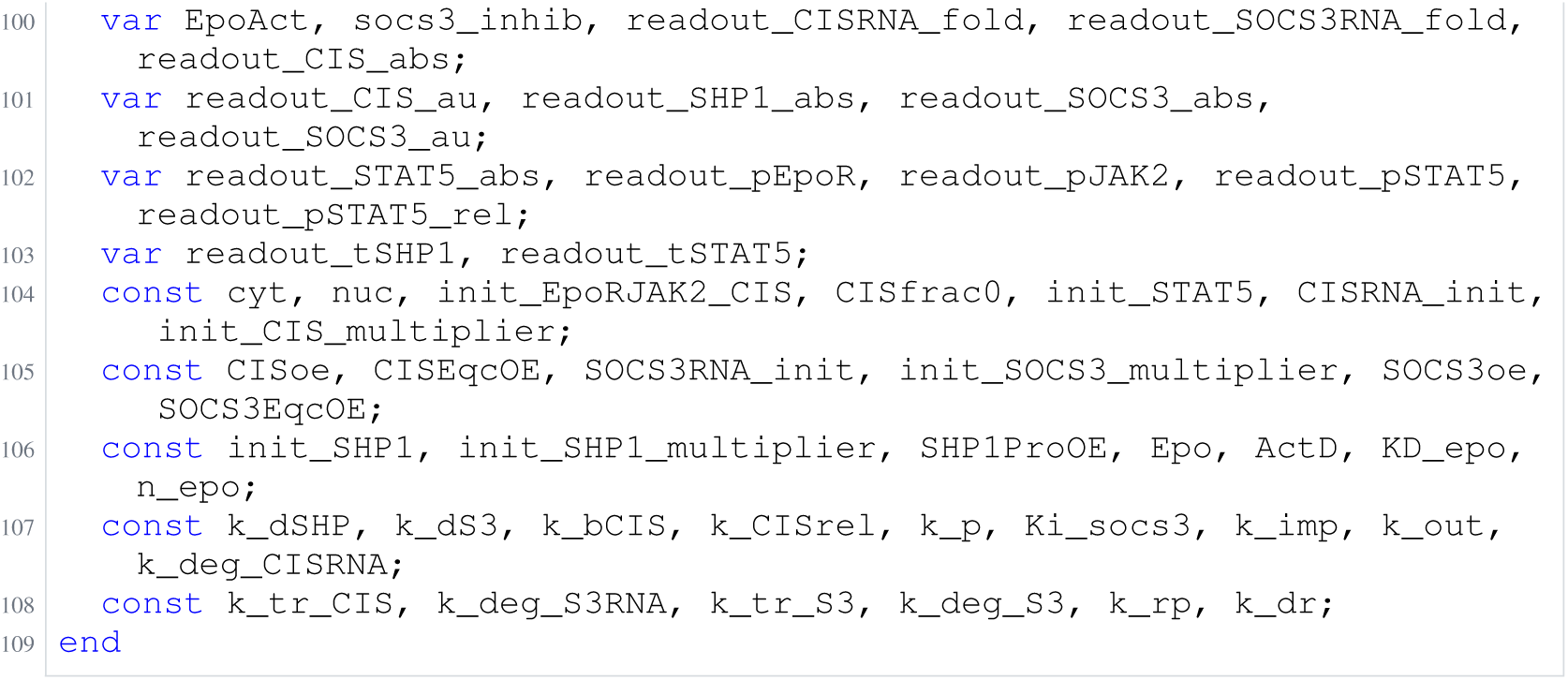
Archived Antimony for bachmann_3; task 56836792_7; selected version 4.

**Listing 92:**
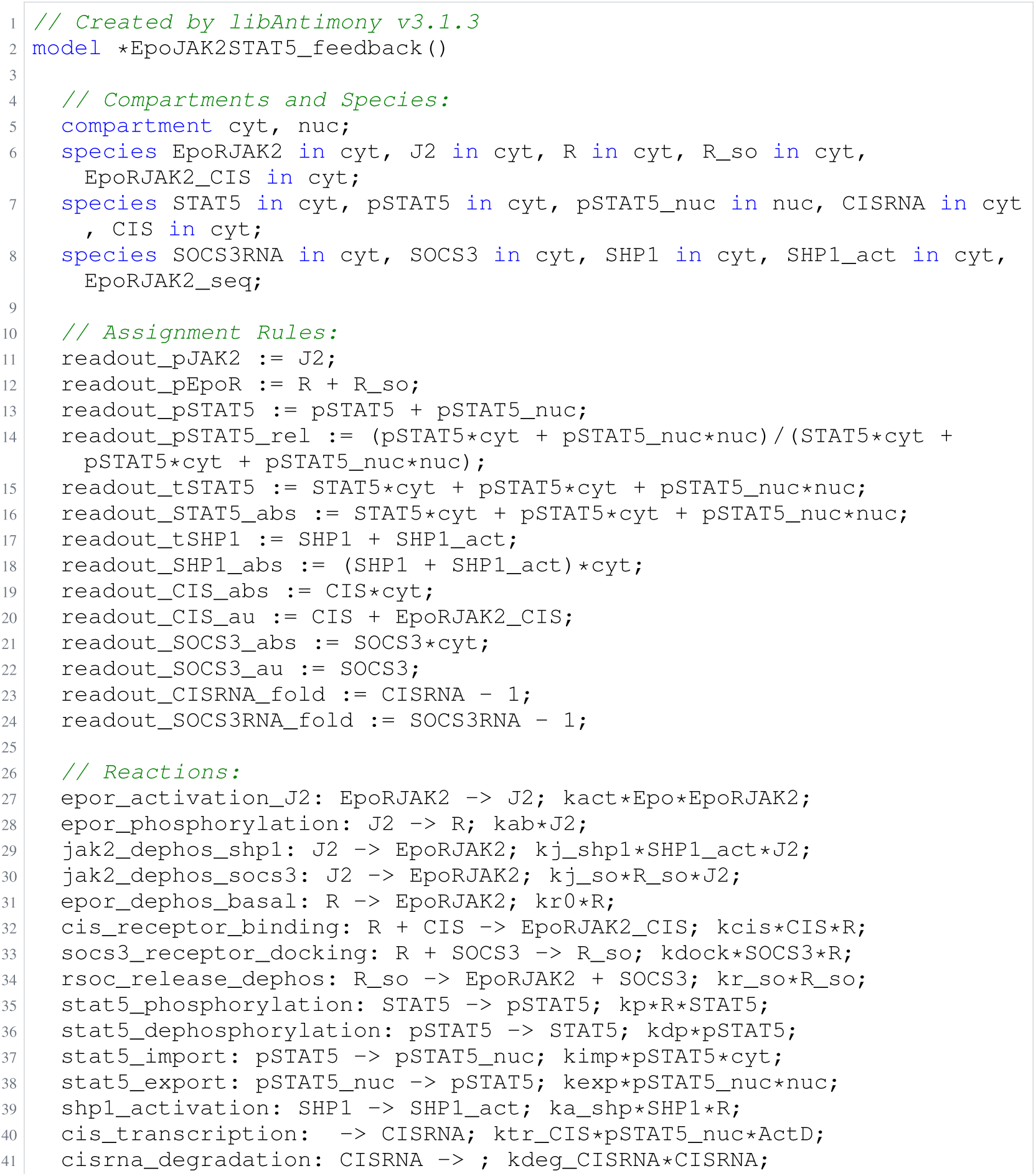
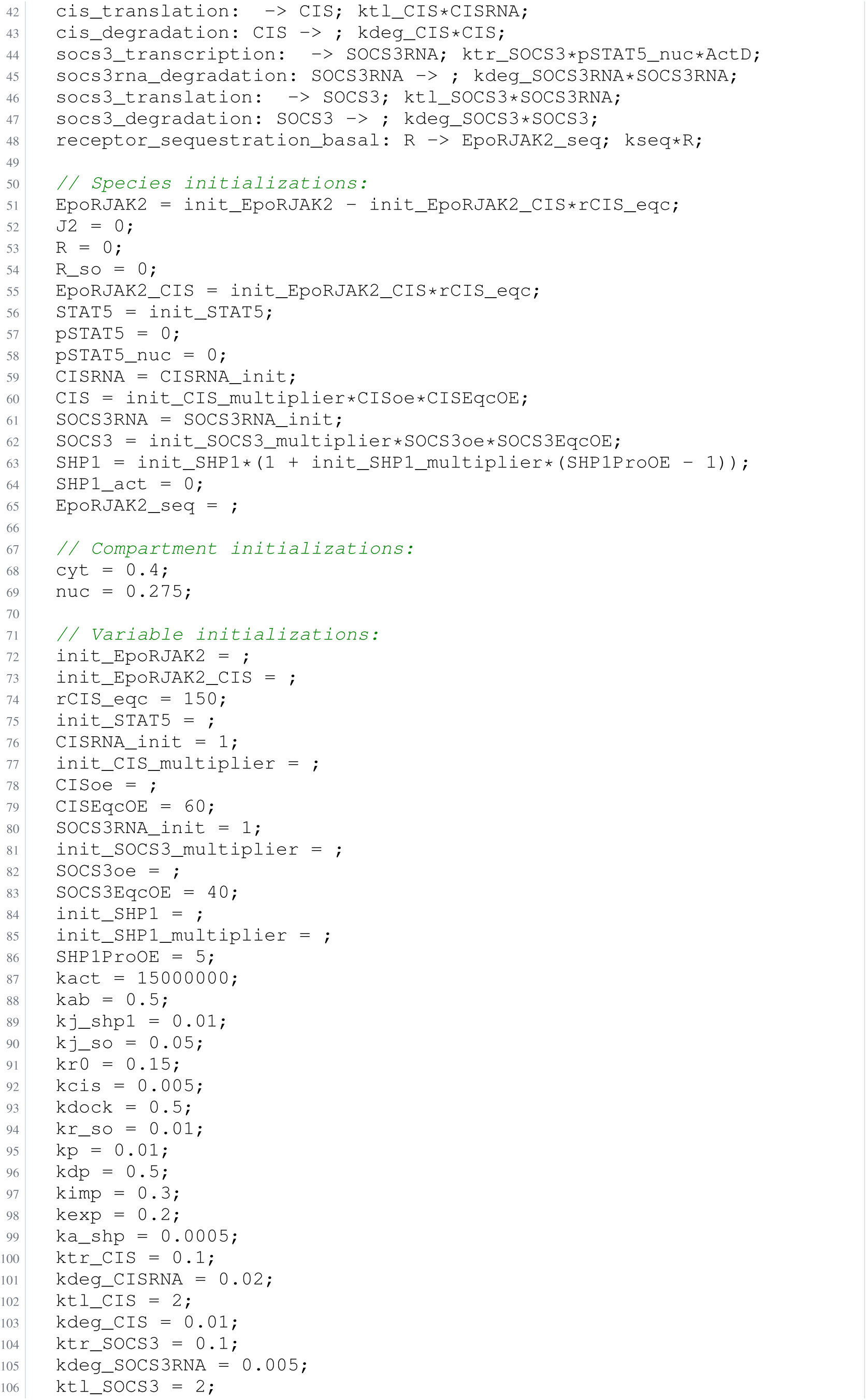
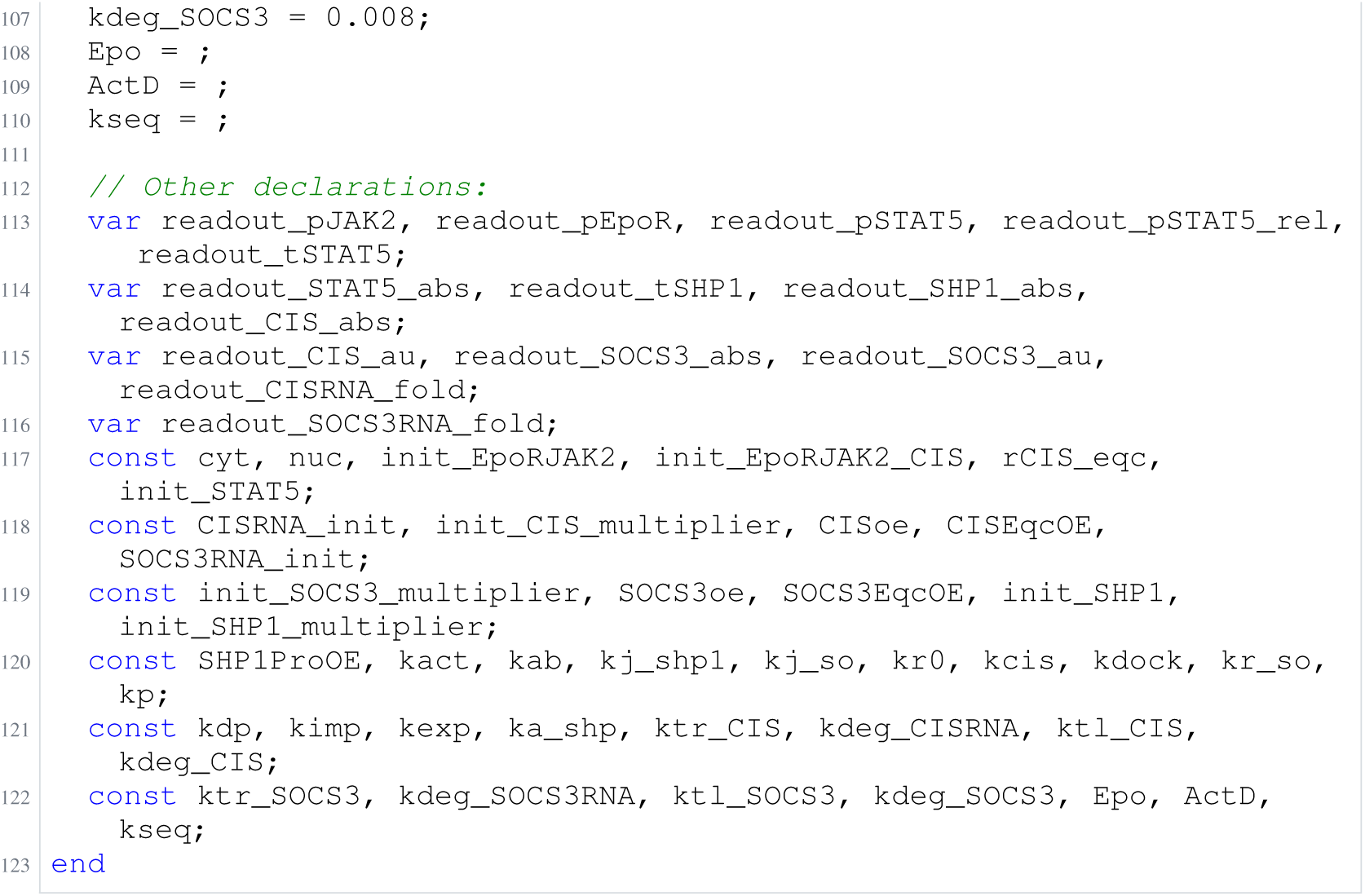
Archived Antimony for bachmann_4; task 56836792_8; selected version 1.

**Listing 93:**
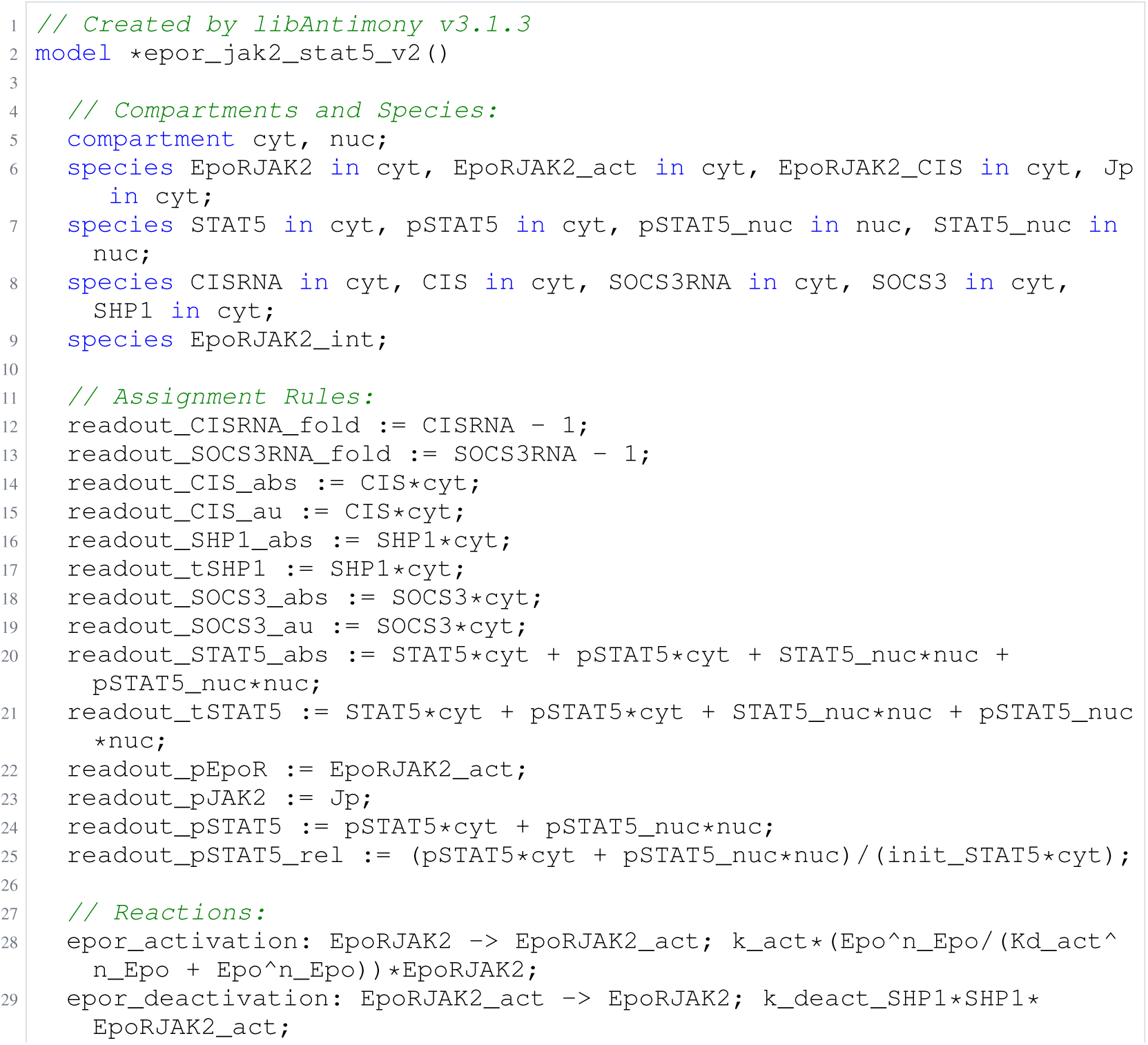
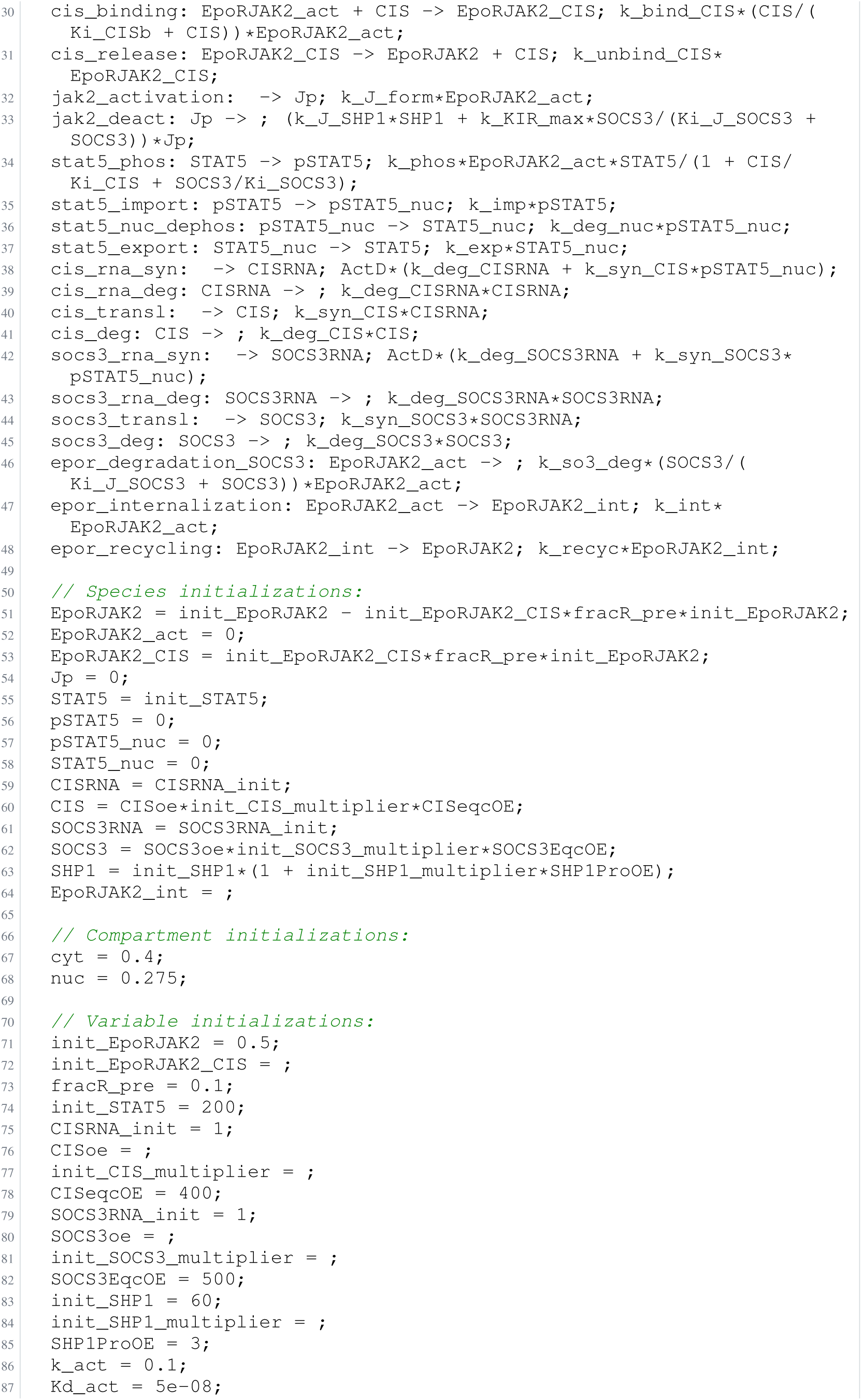
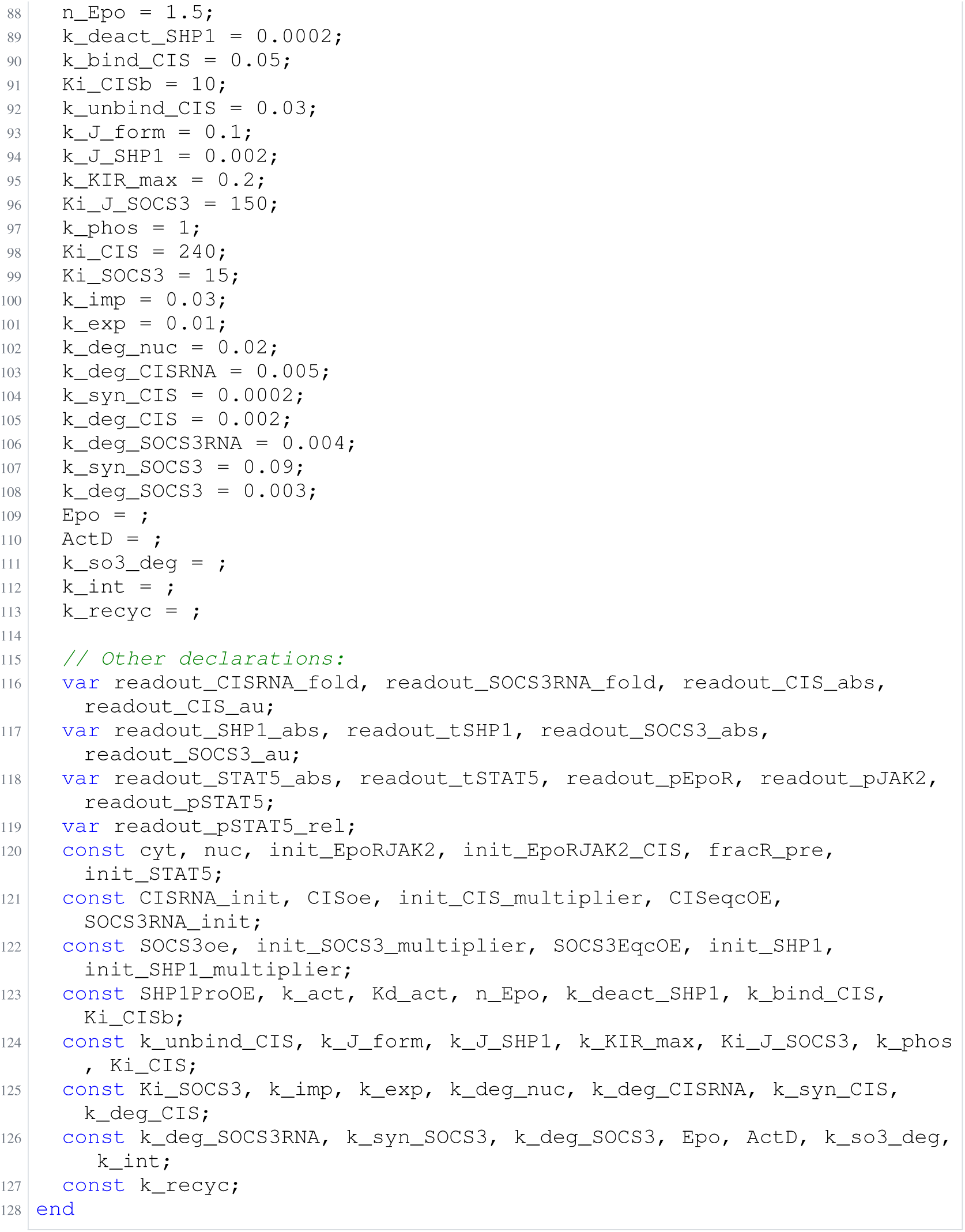
Archived Antimony for bachmann_5; task 56836792_9; selected version 16.

### H.14 Raimundez

**Listing 94:**
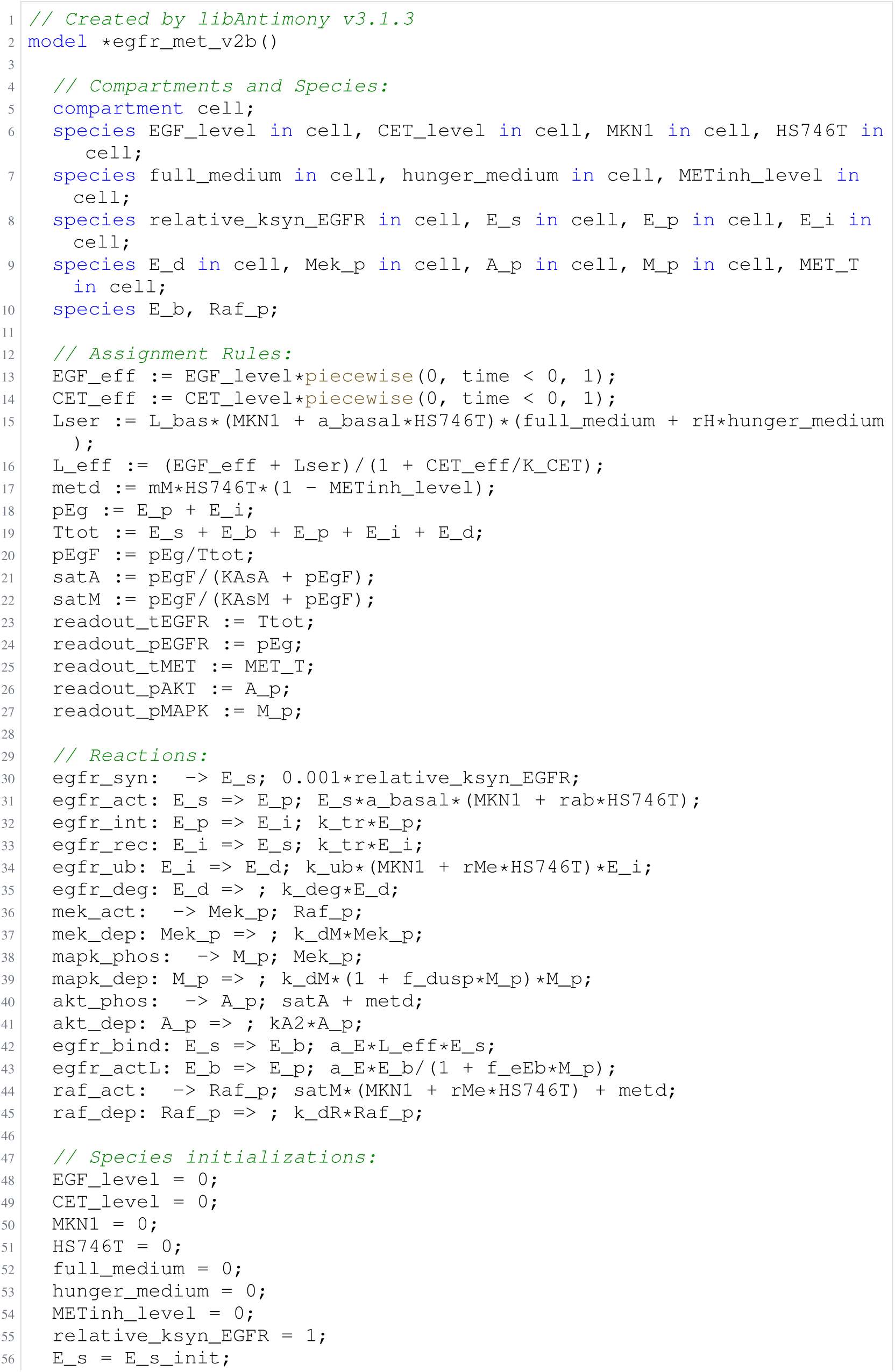
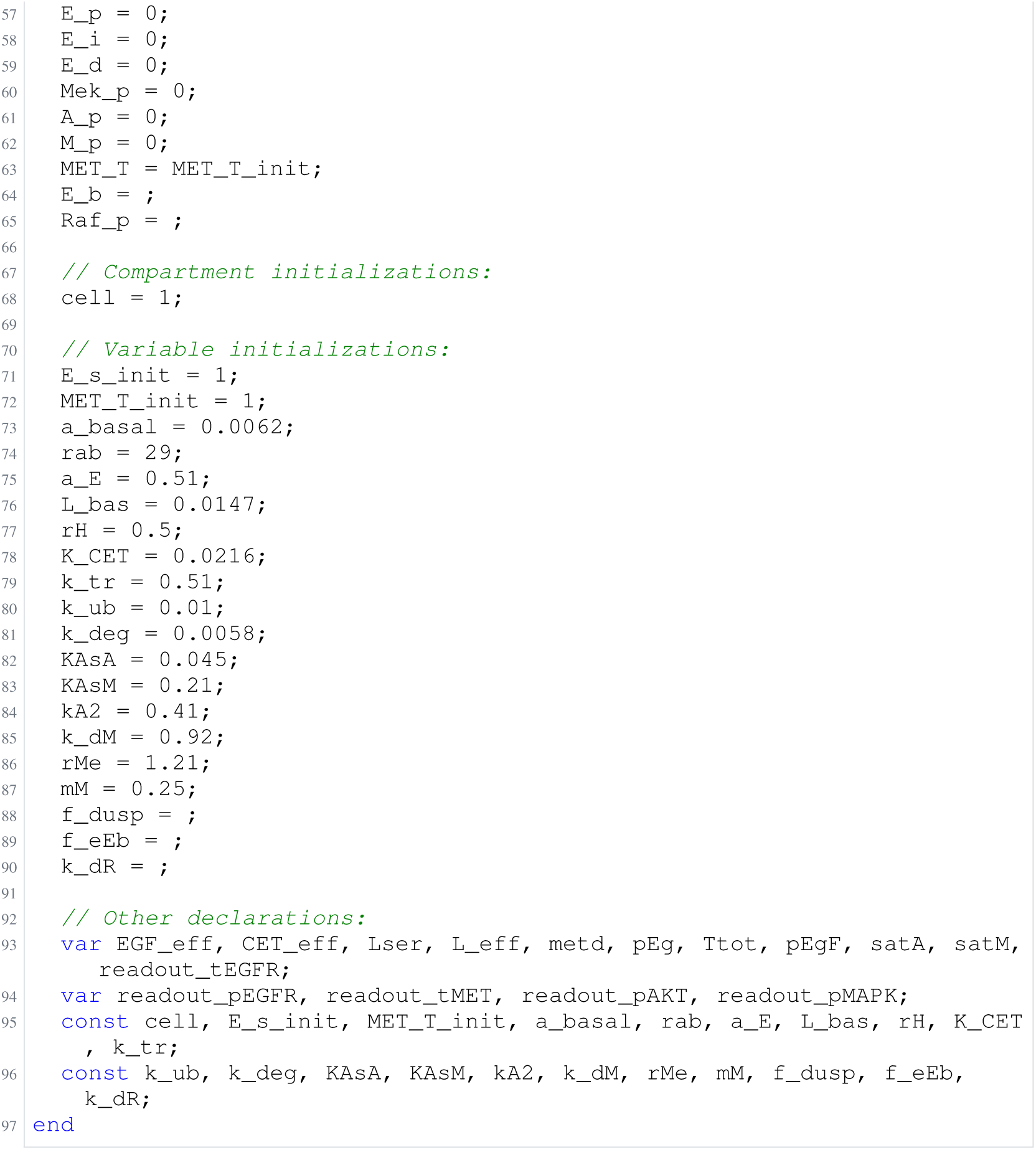
Archived Antimony for raimundez_1; task 56836792_15; selected version 28.

**Listing 95:**
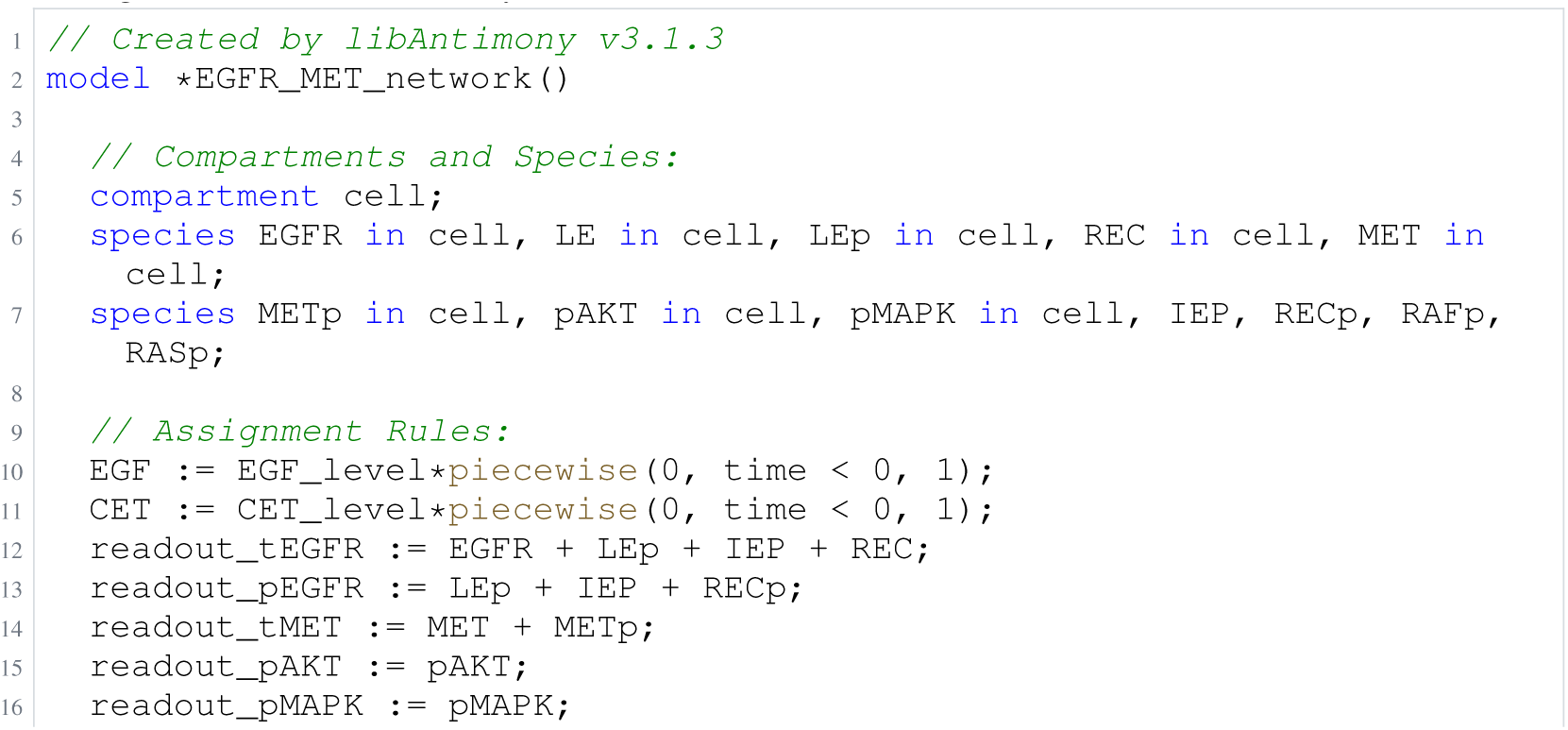
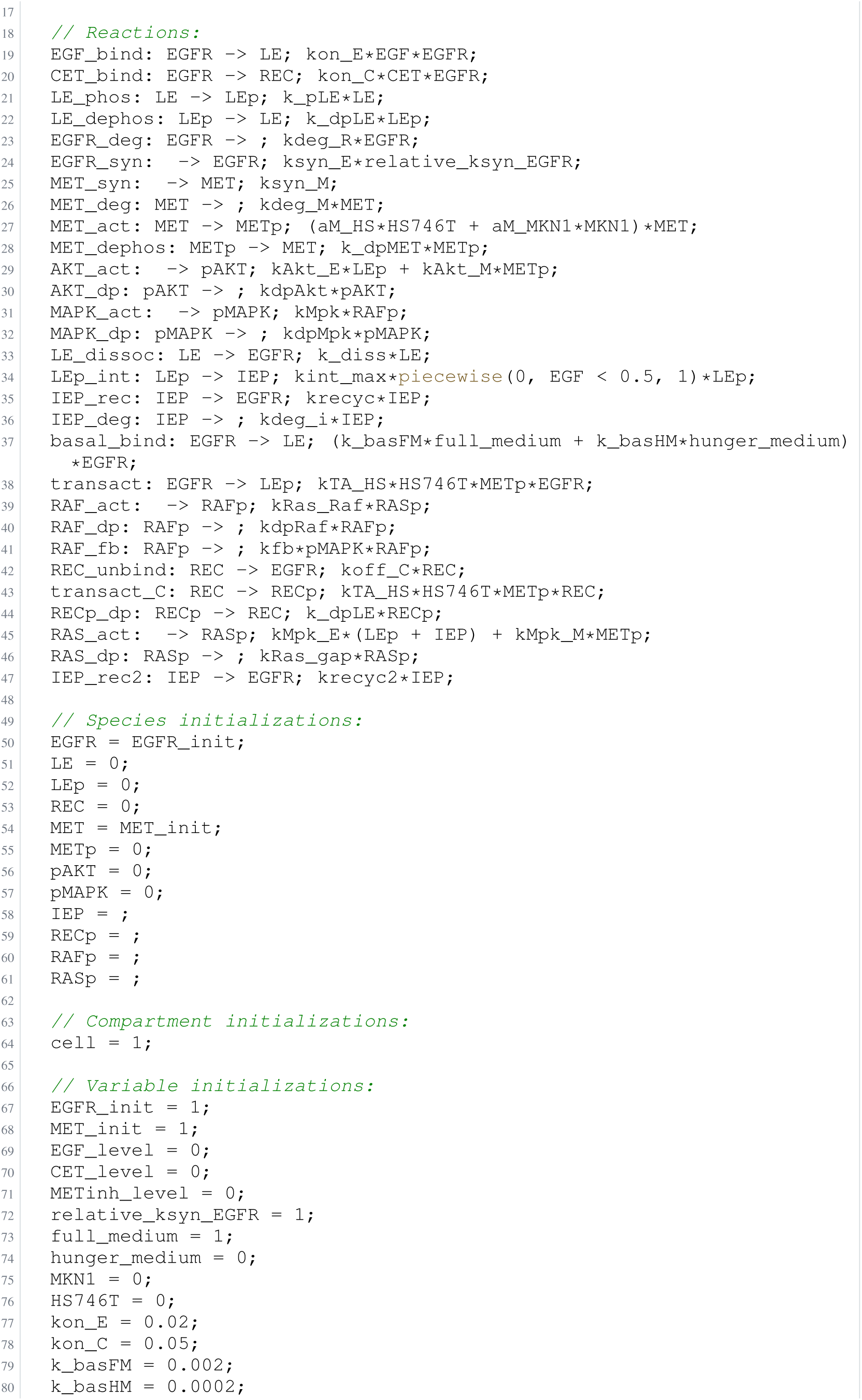
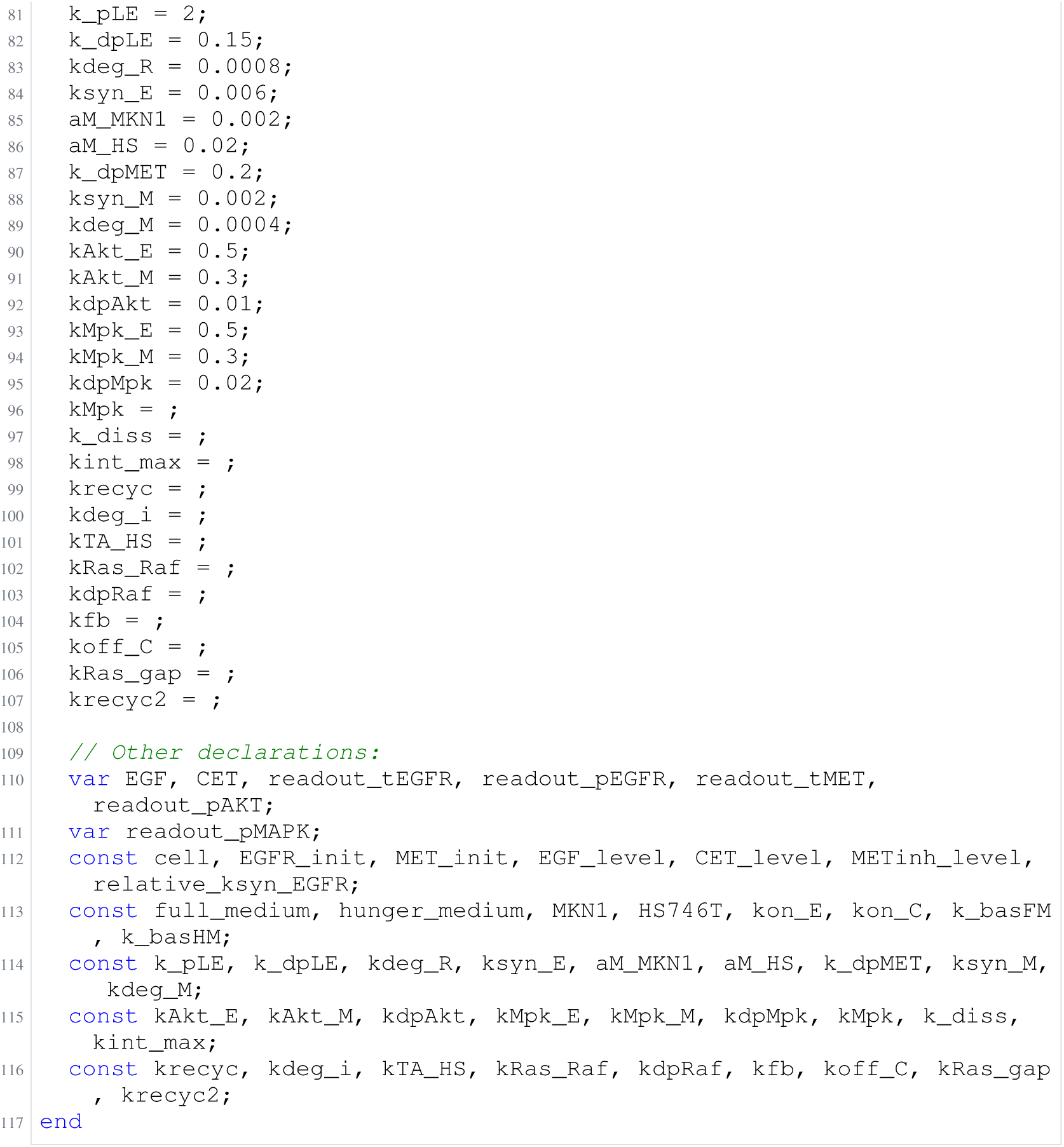
Archived Antimony for raimundez_2; task 56836792_16; selected version 17.

**Listing 96:**
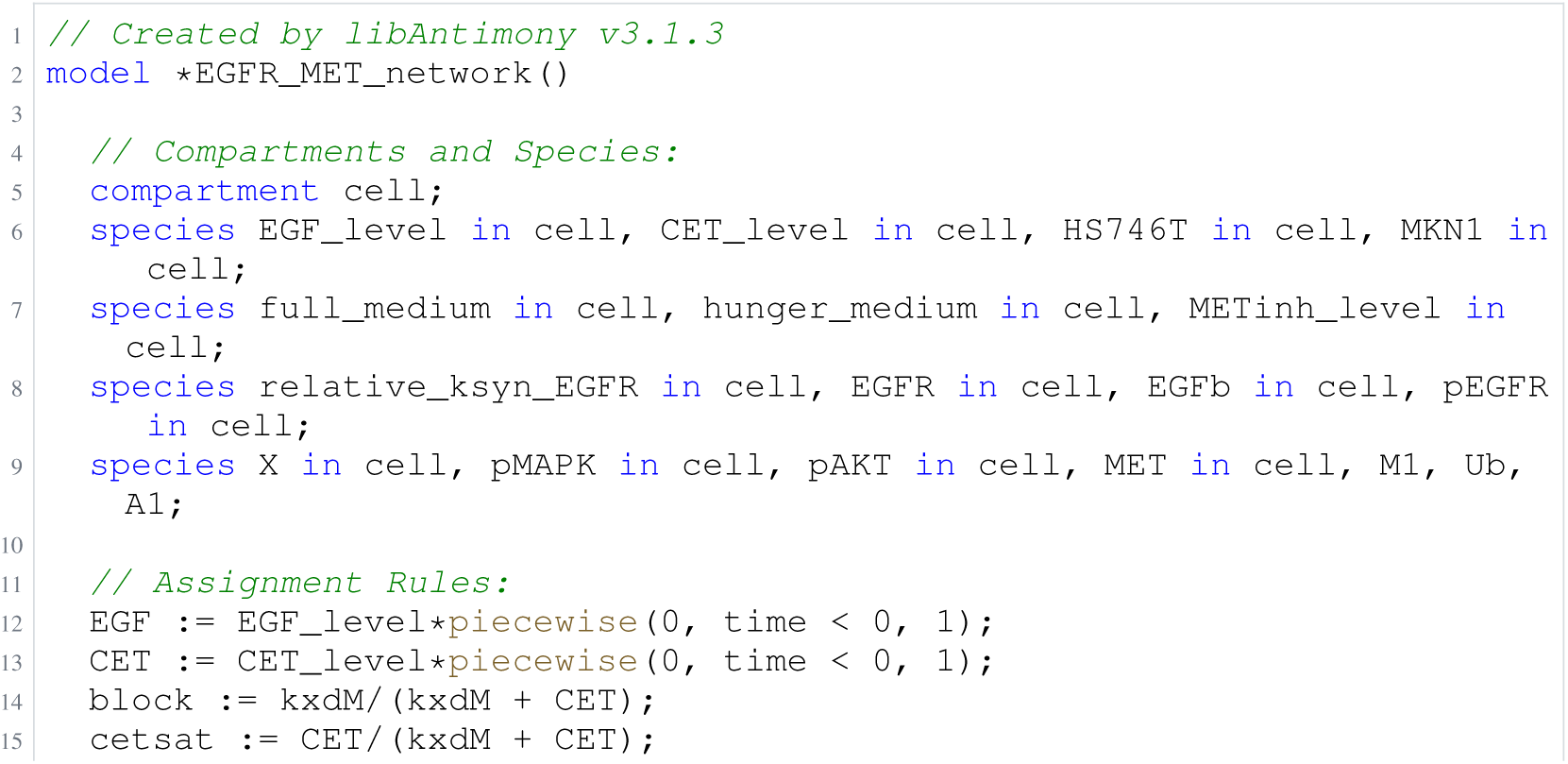
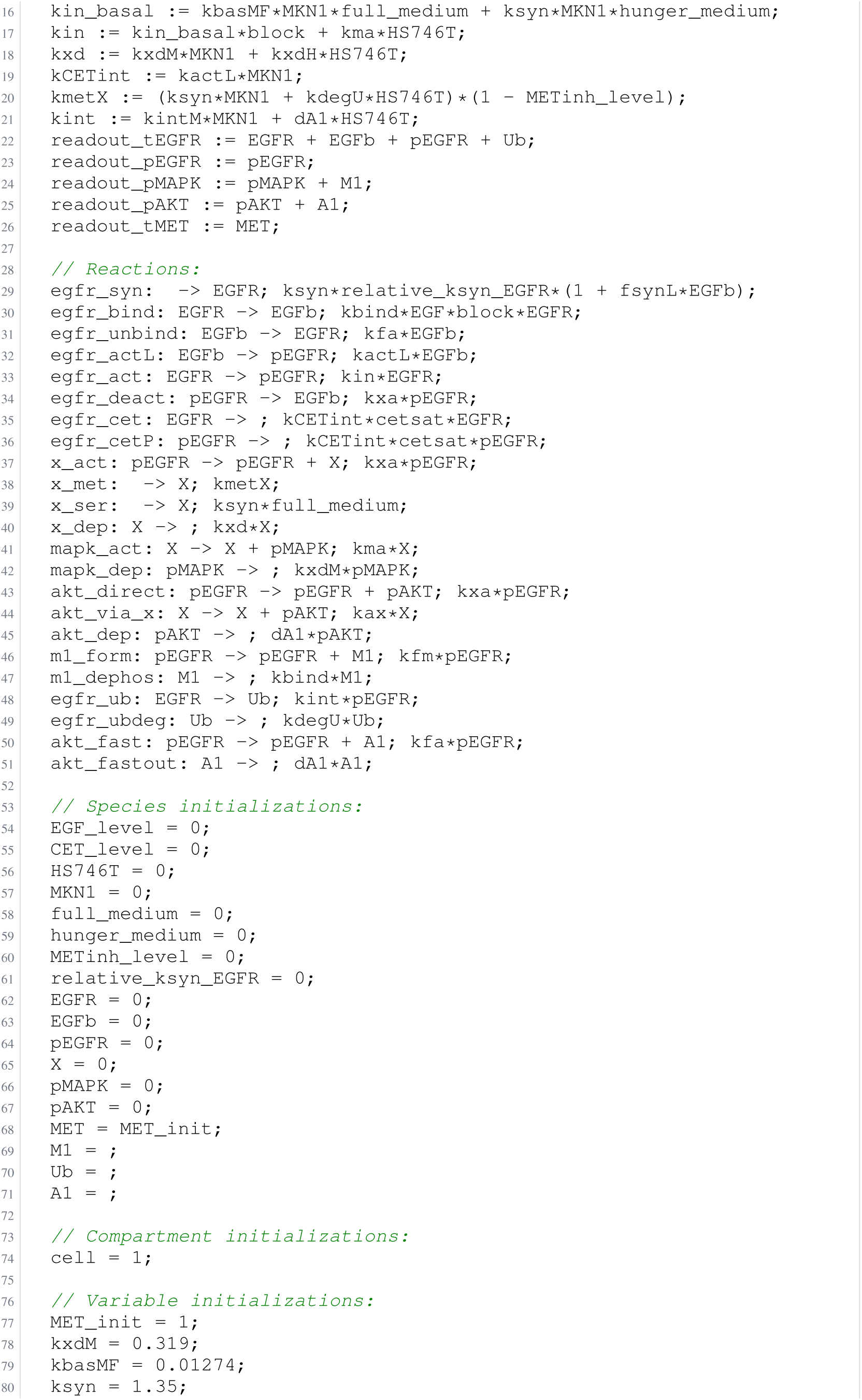
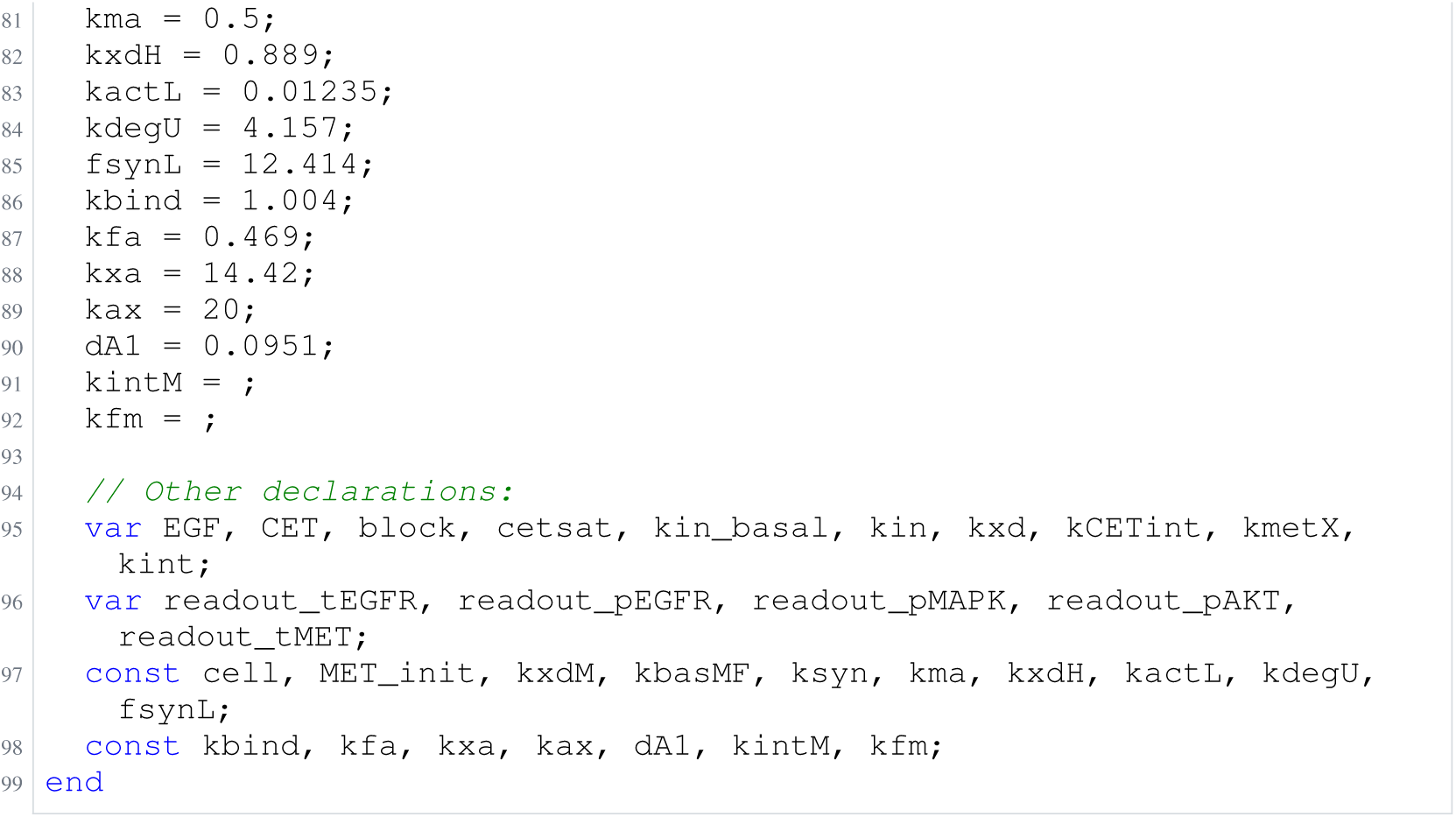
Archived Antimony for raimundez_3; task 56836792_17; selected version 19.

**Listing 97:**
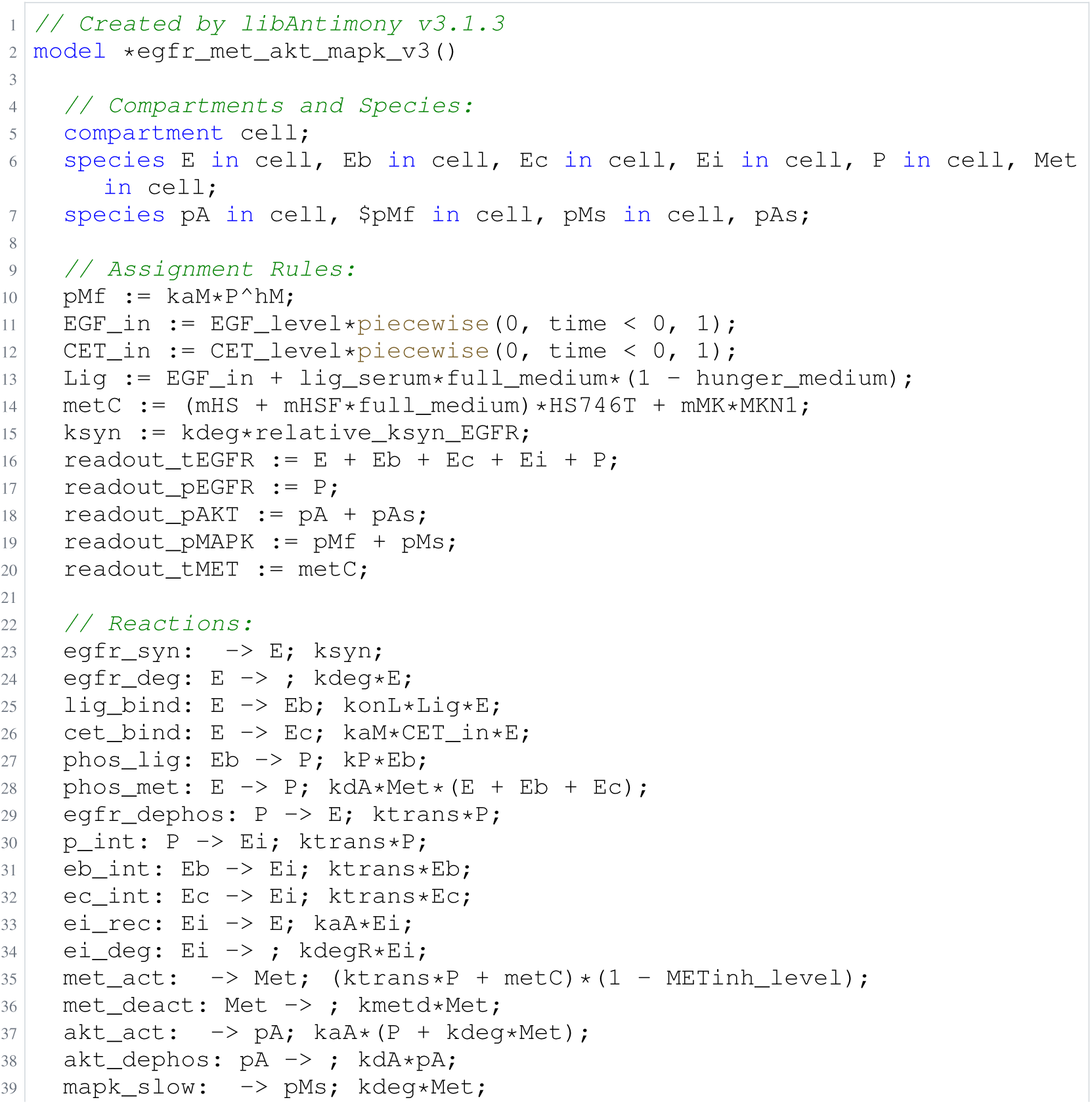
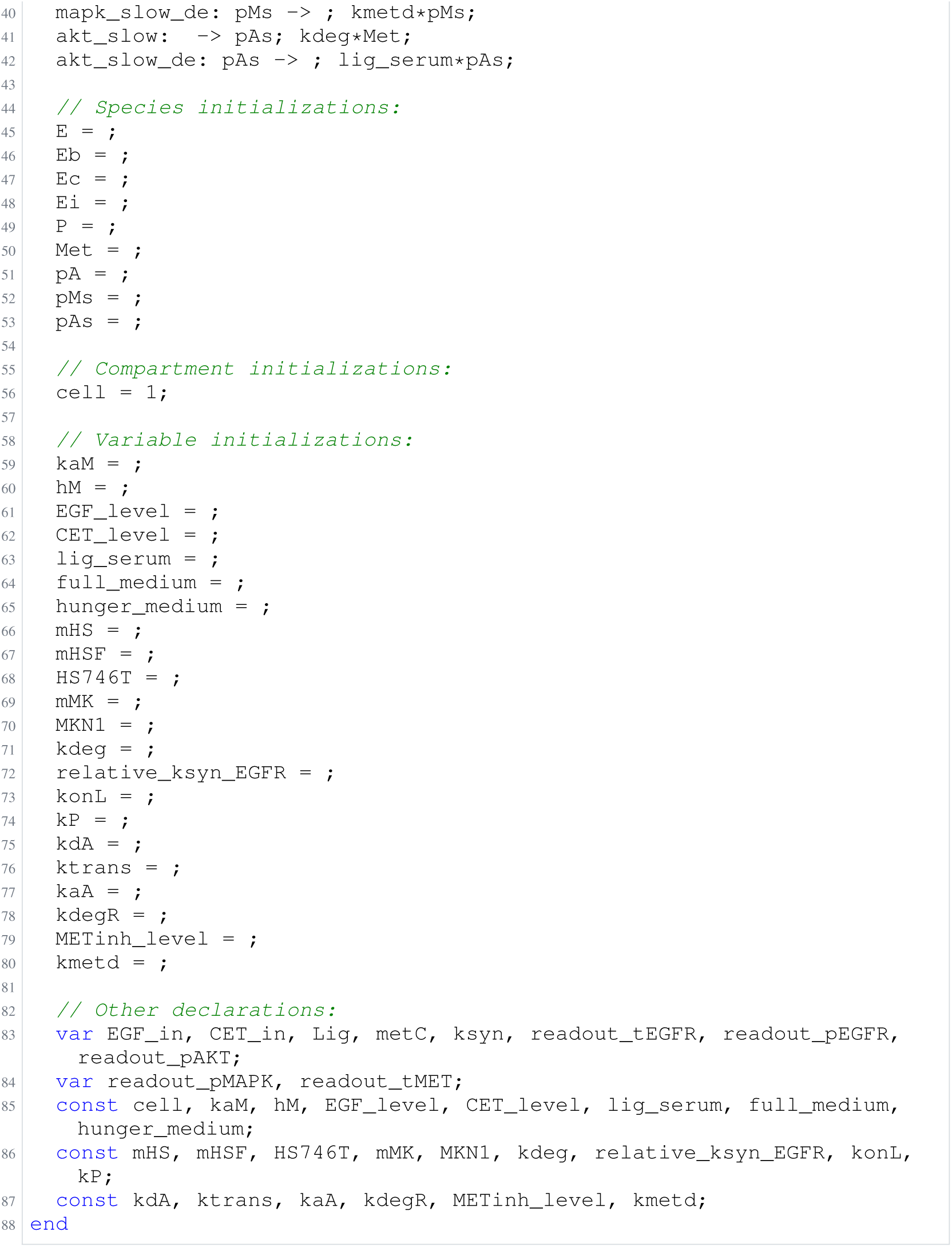
Archived Antimony for raimundez_4; task 56836792_18; selected version 17.

**Listing 98:**
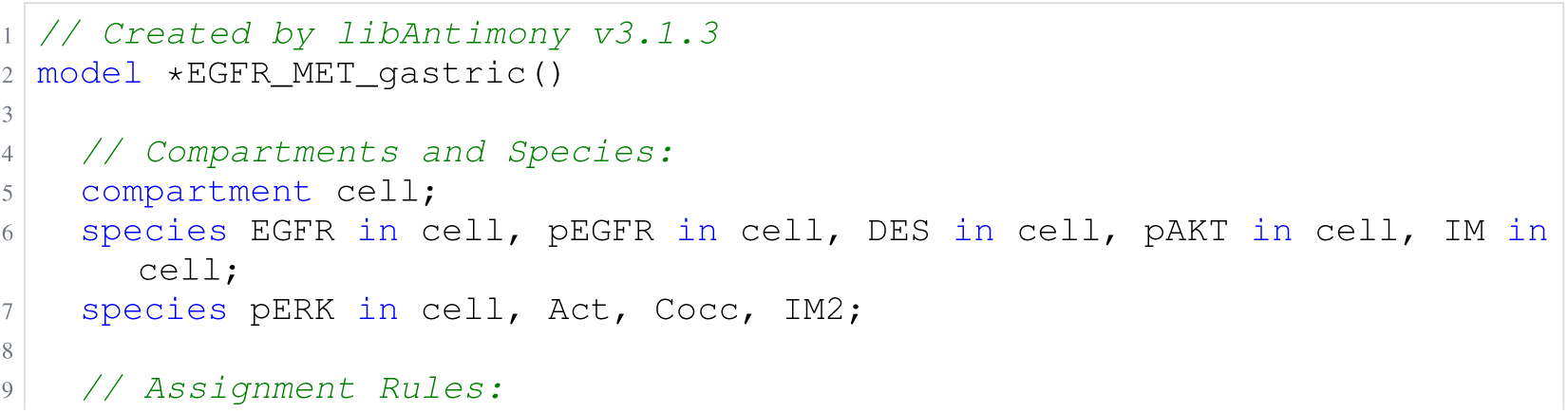
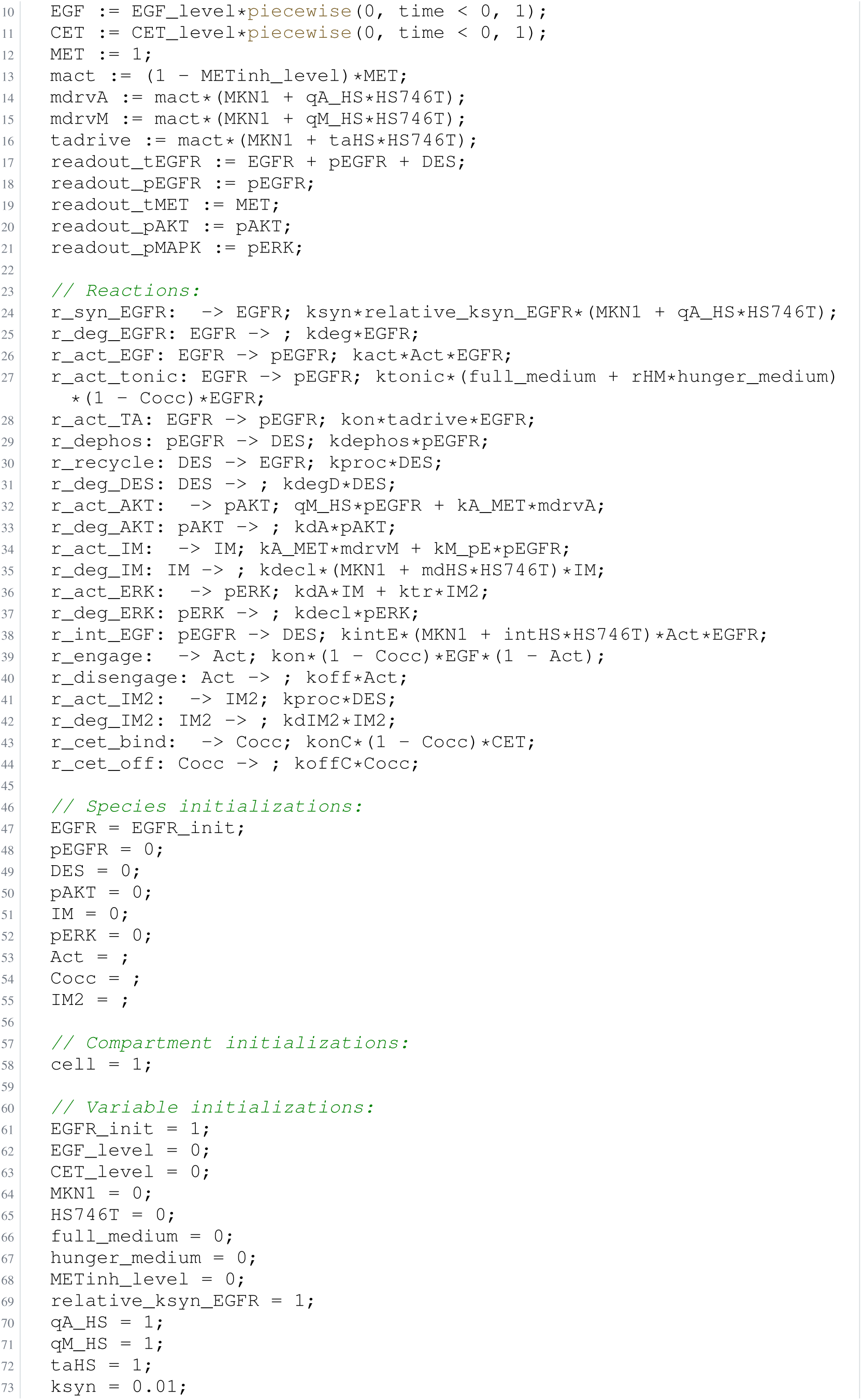
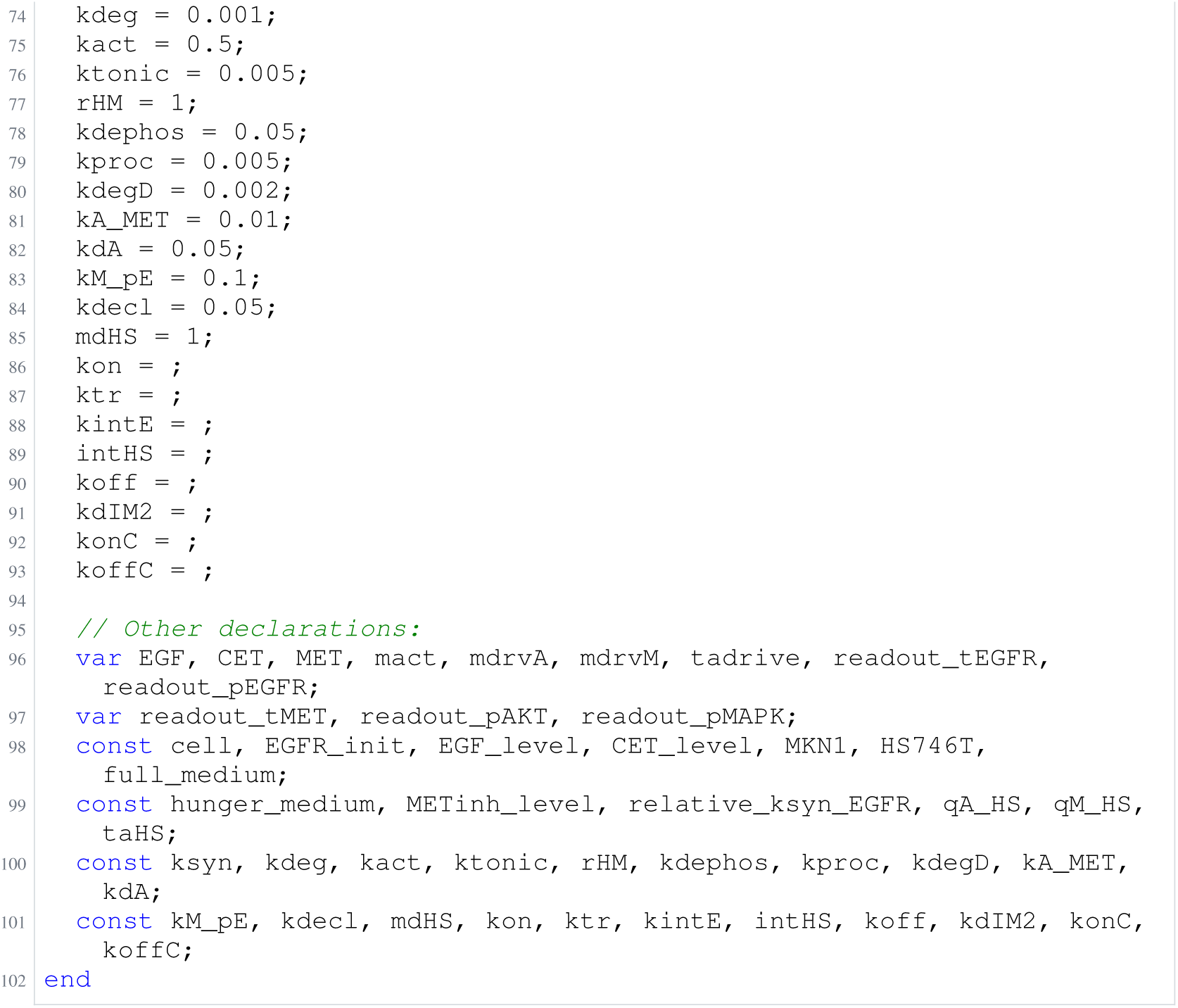
Archived Antimony for raimundez_5; task 56836792_19; selected version 27.

### H.15 Isensee

**Listing 99:**
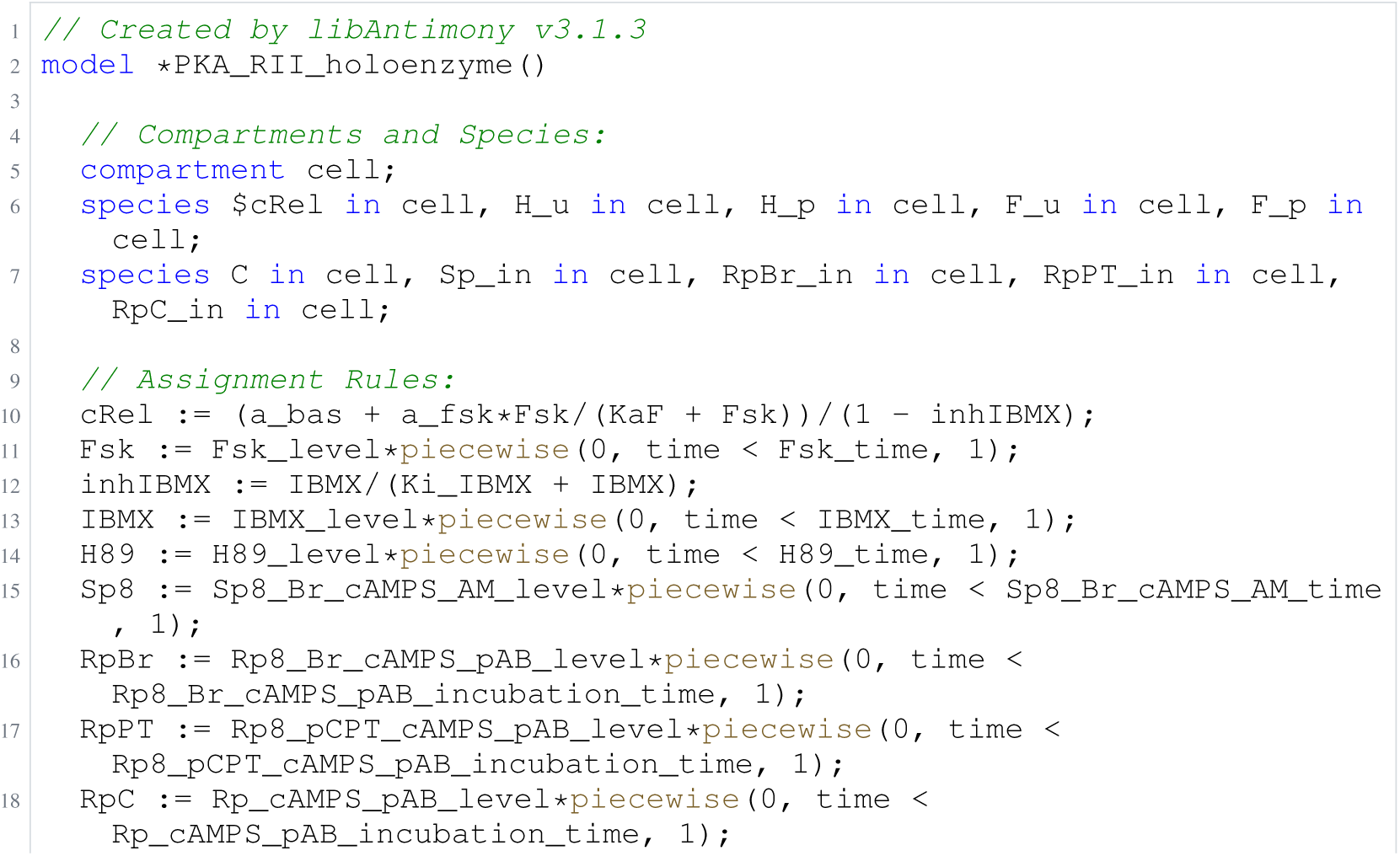
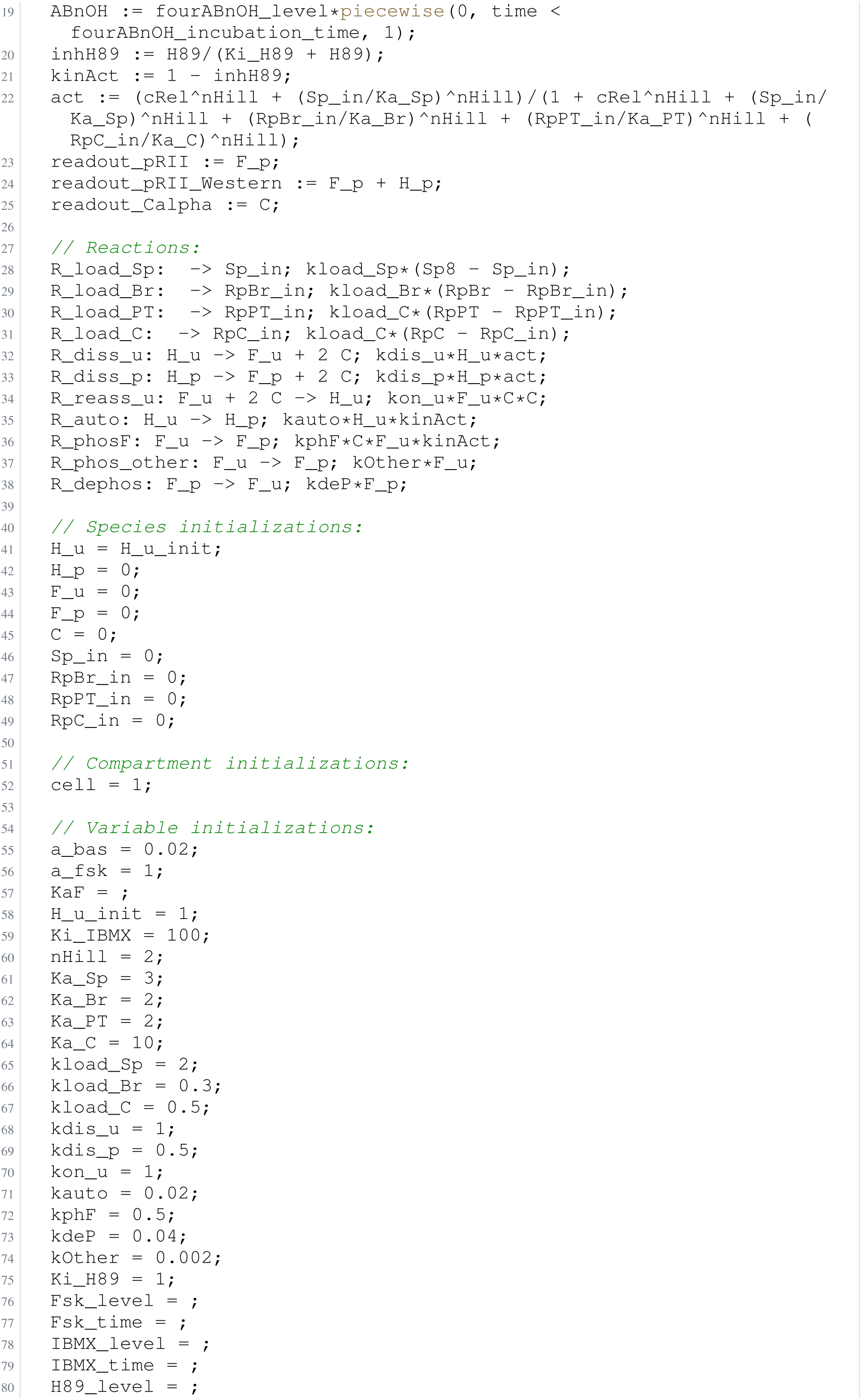
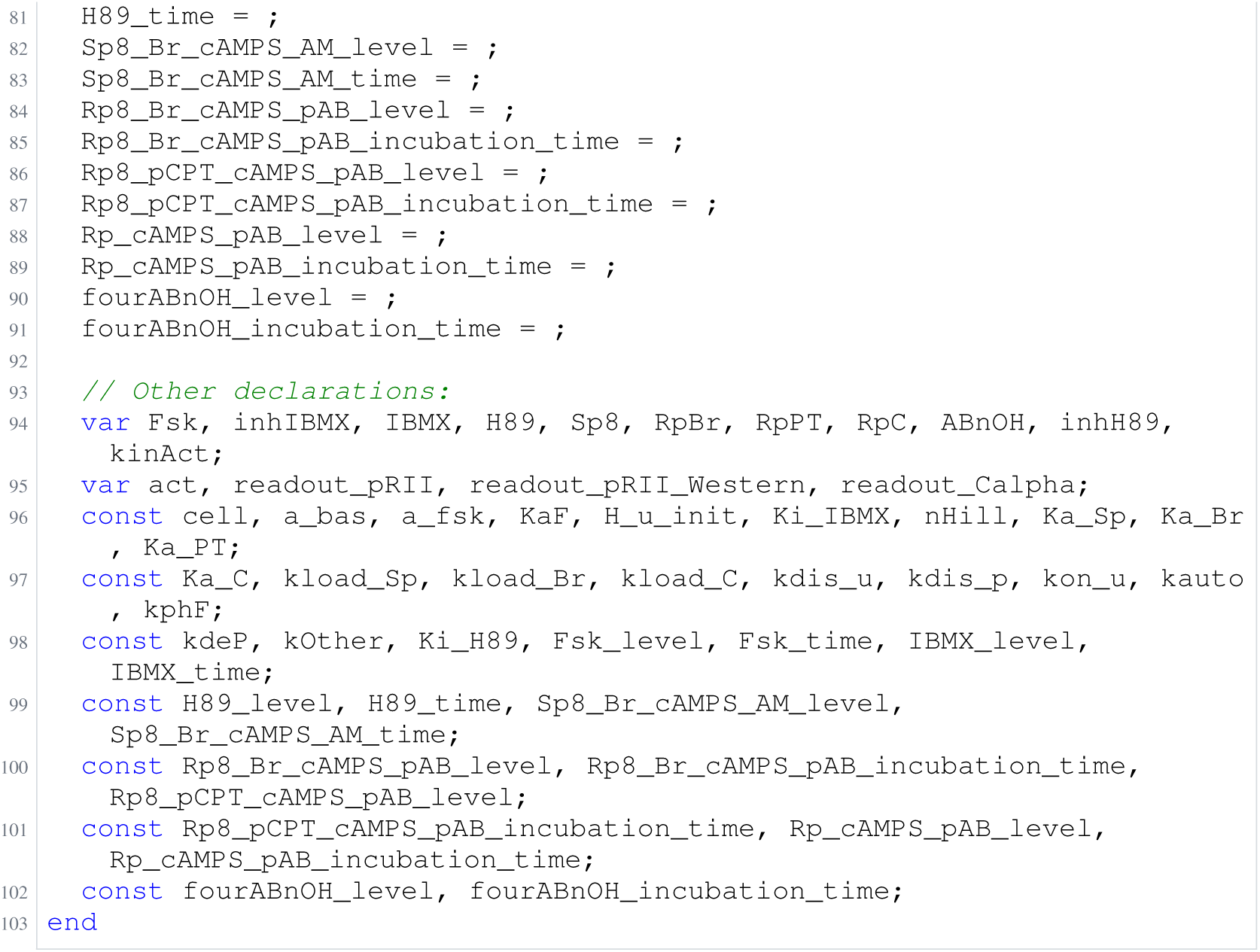
Archived Antimony for isensee_1; task 56836711_3; selected version 2.

**Listing 100:**
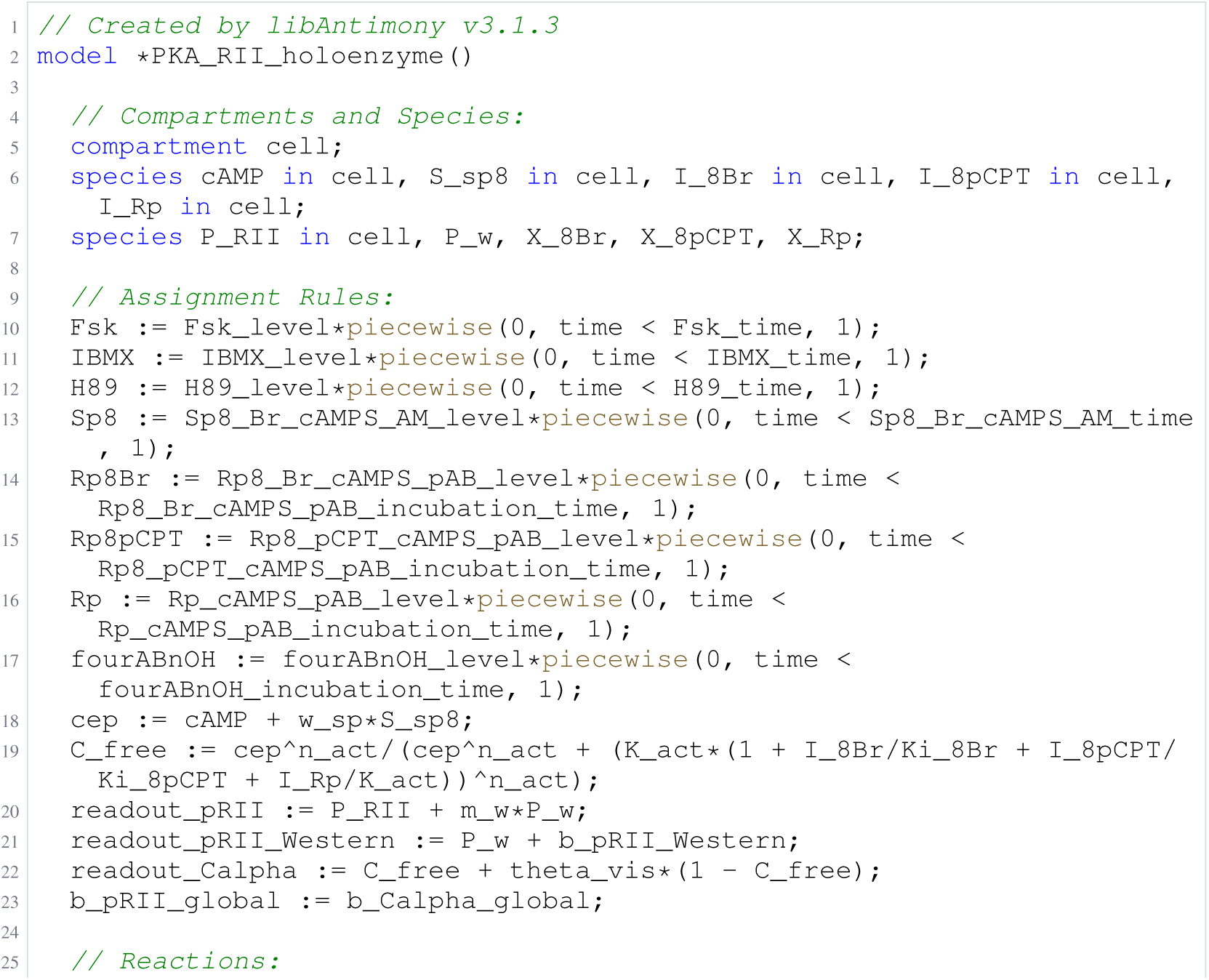
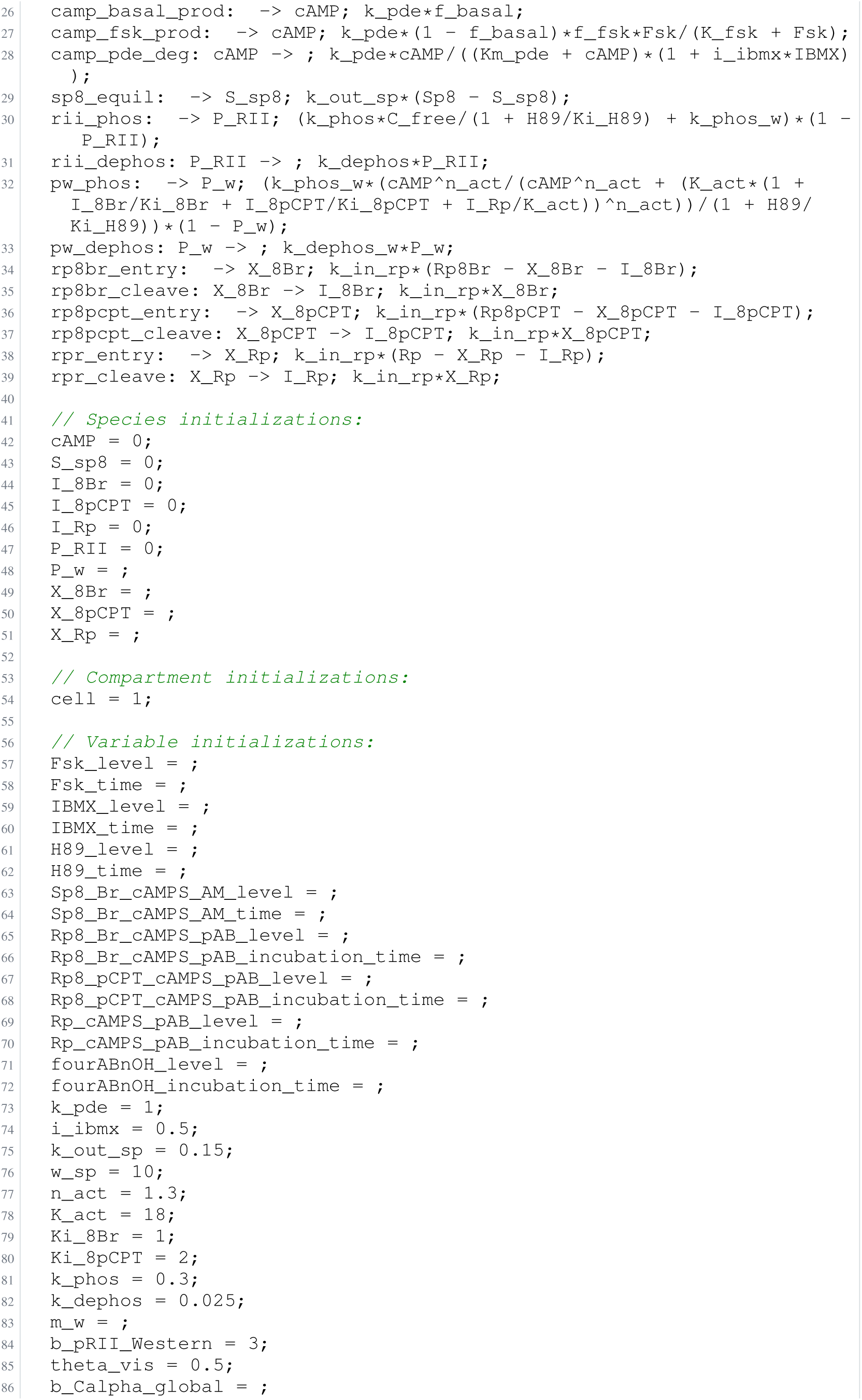
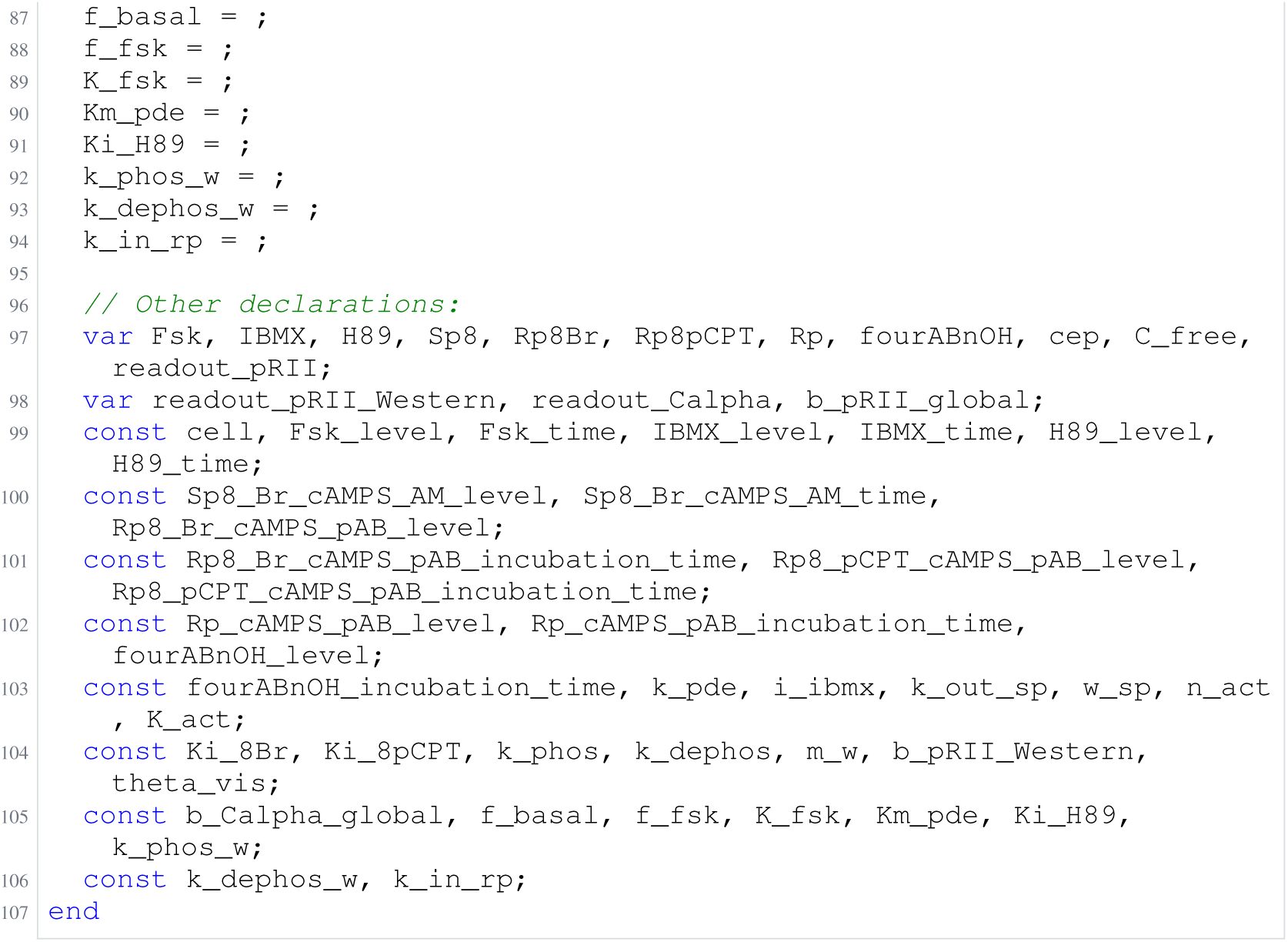
Archived Antimony for isensee_2; task 56836711_1; selected version 17.

**Listing 101:**
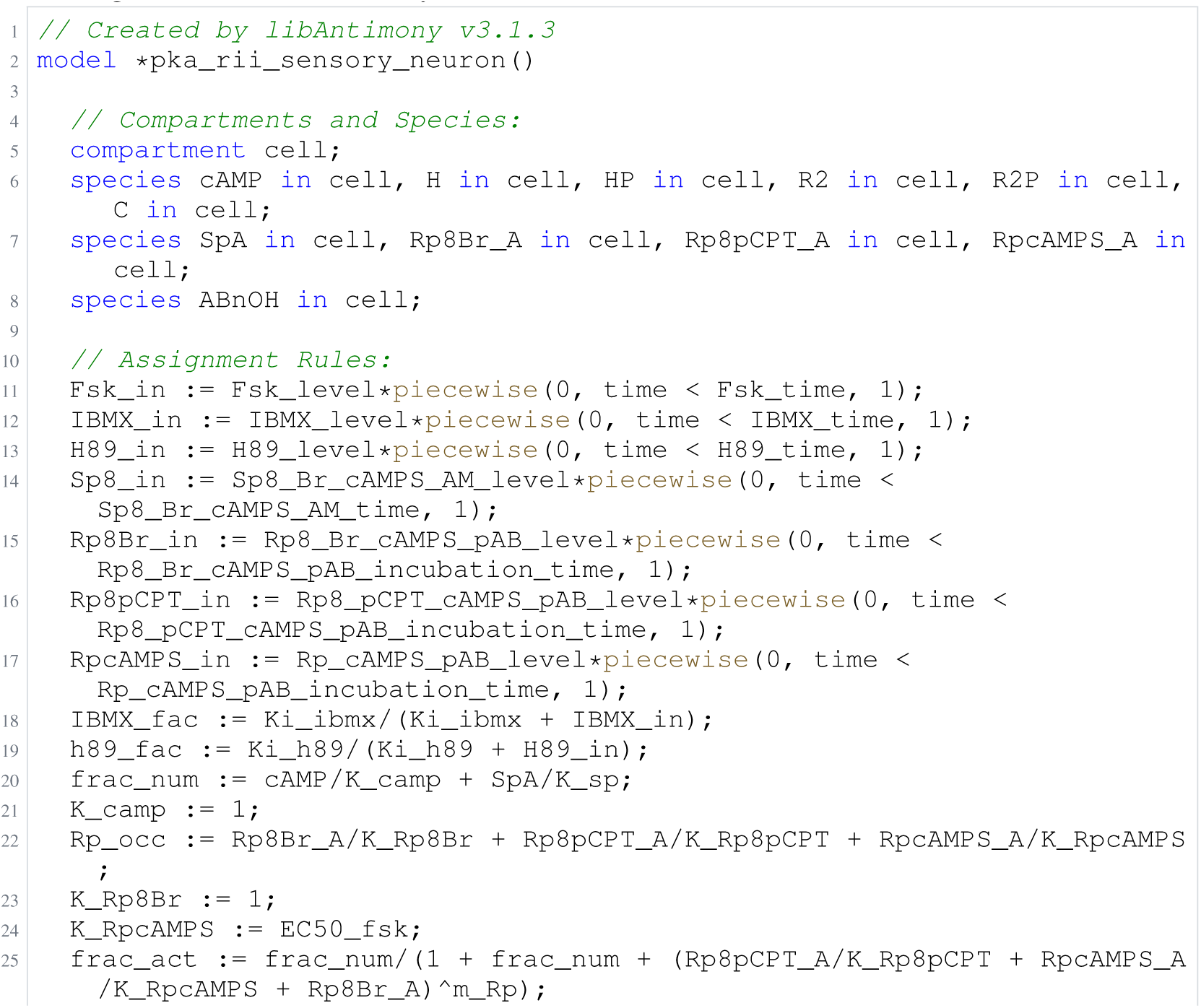
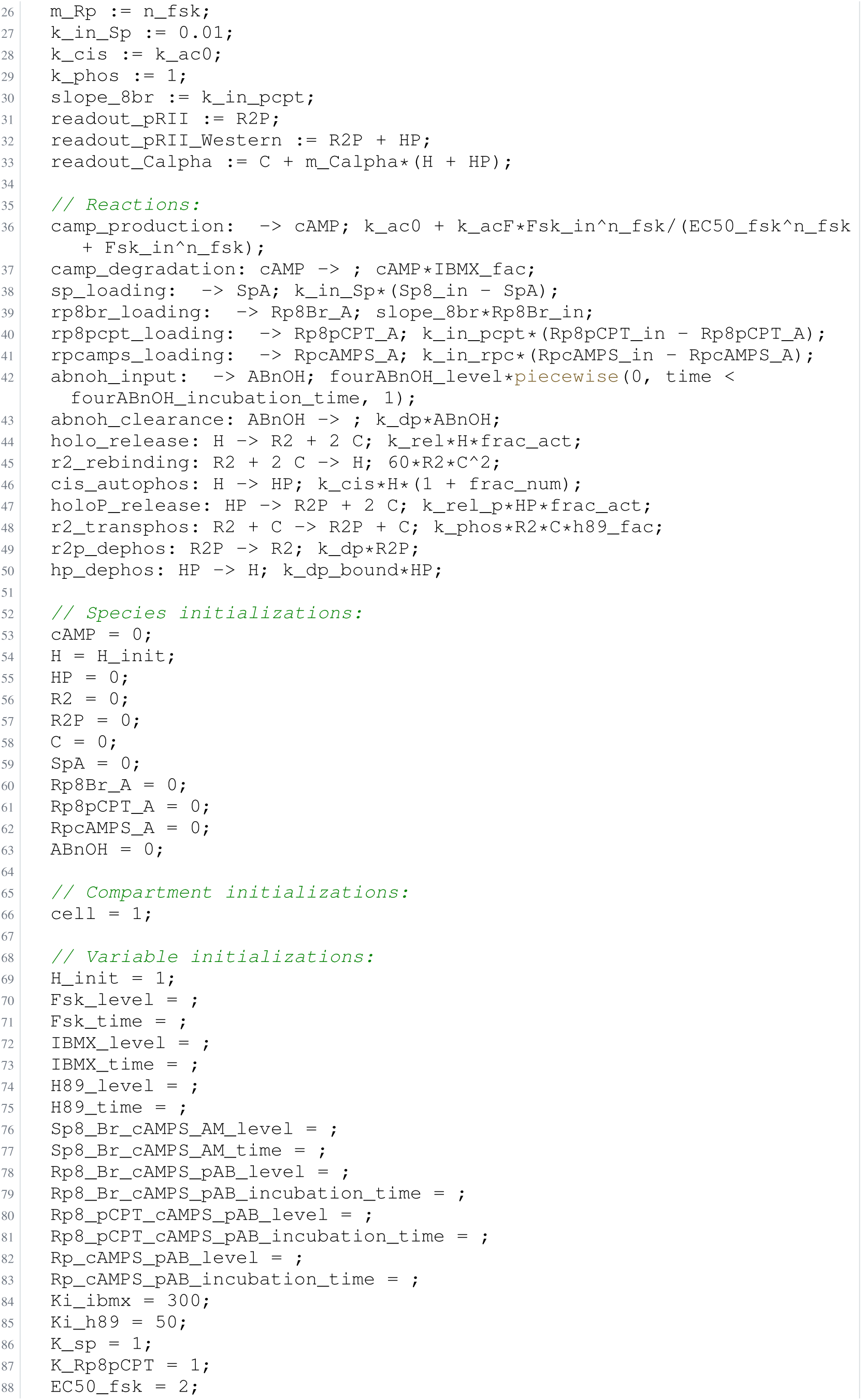
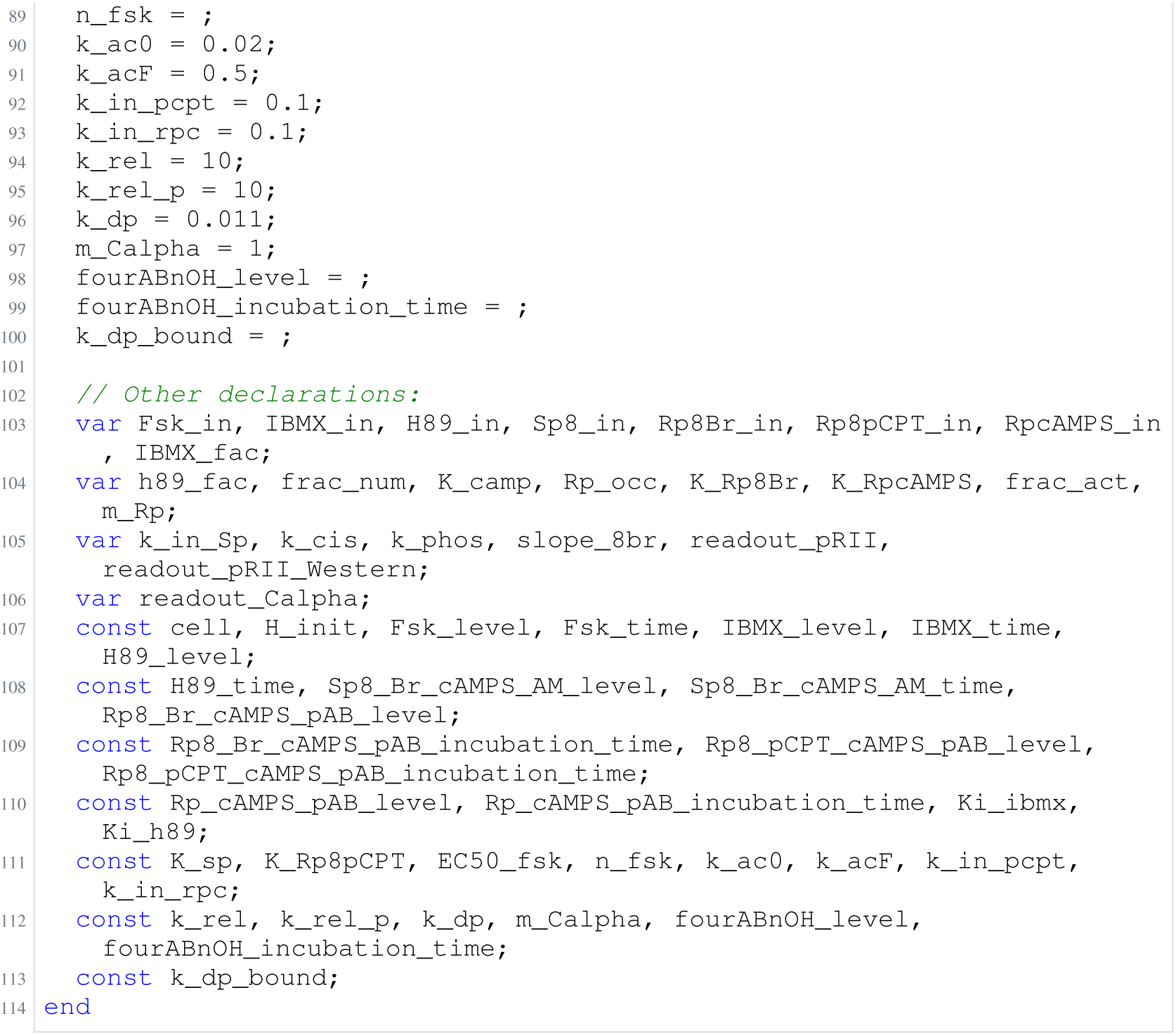
Archived Antimony for isensee_3; task 56836711_2; selected version 17.

**Listing 102:**
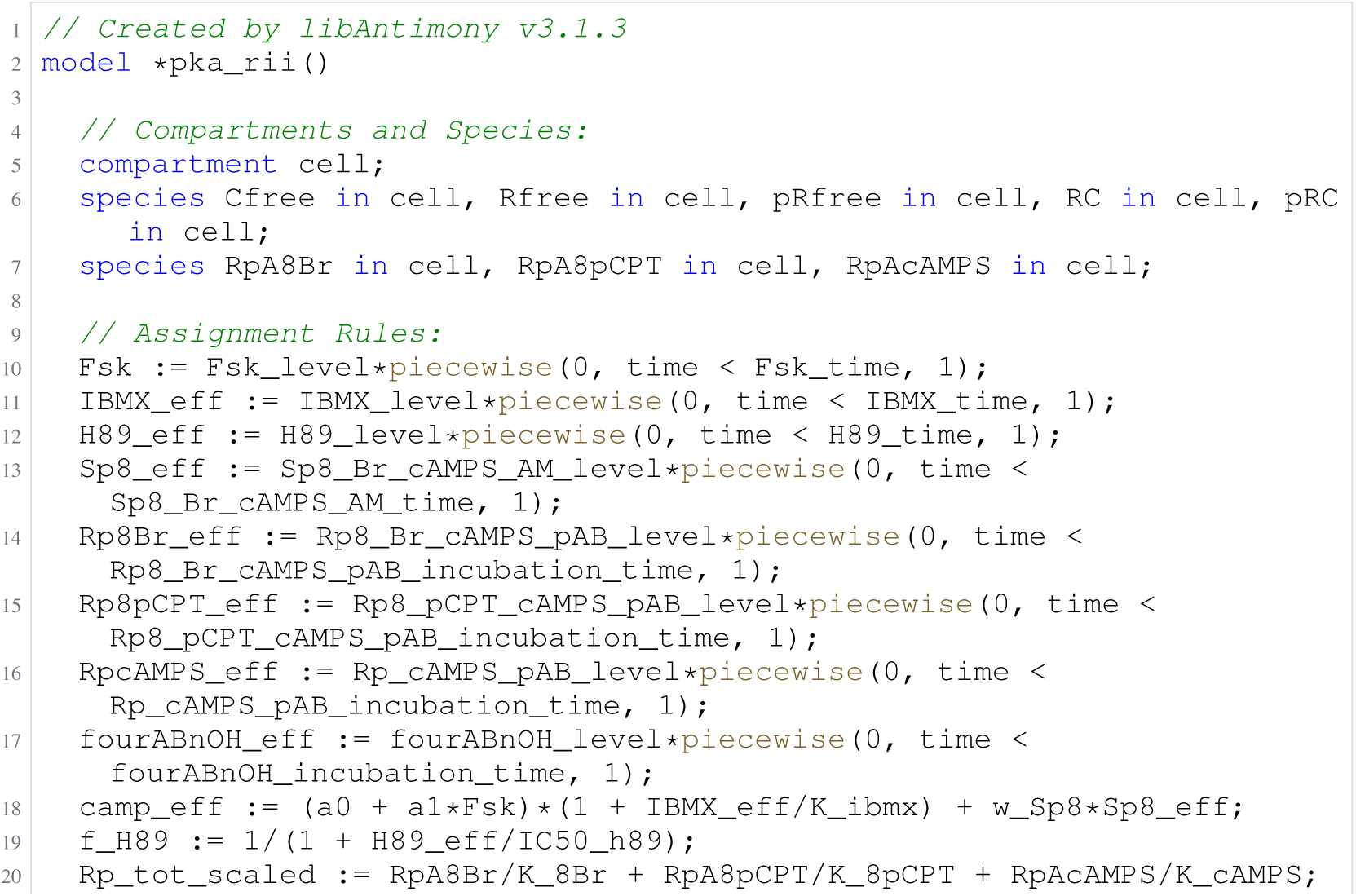
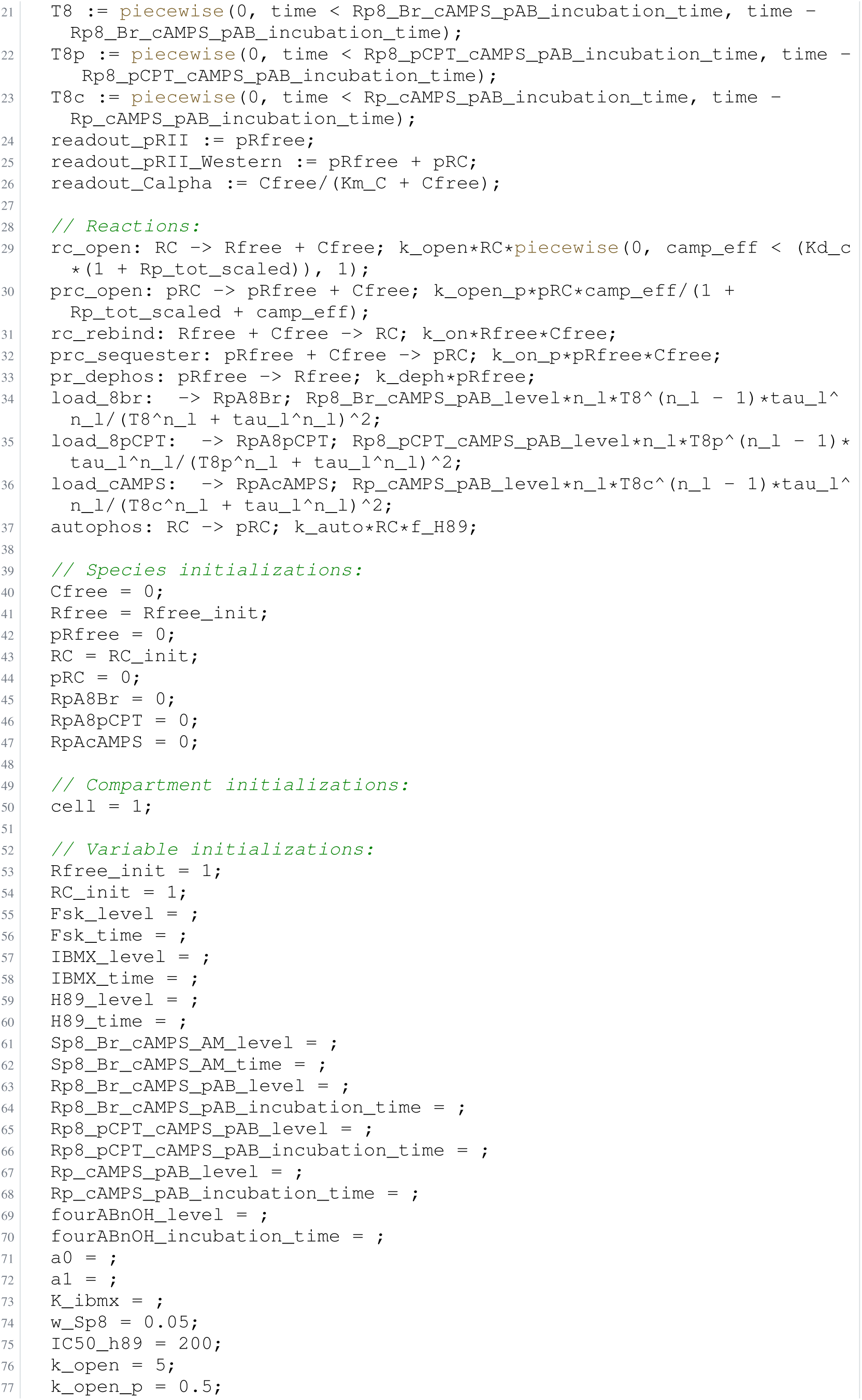
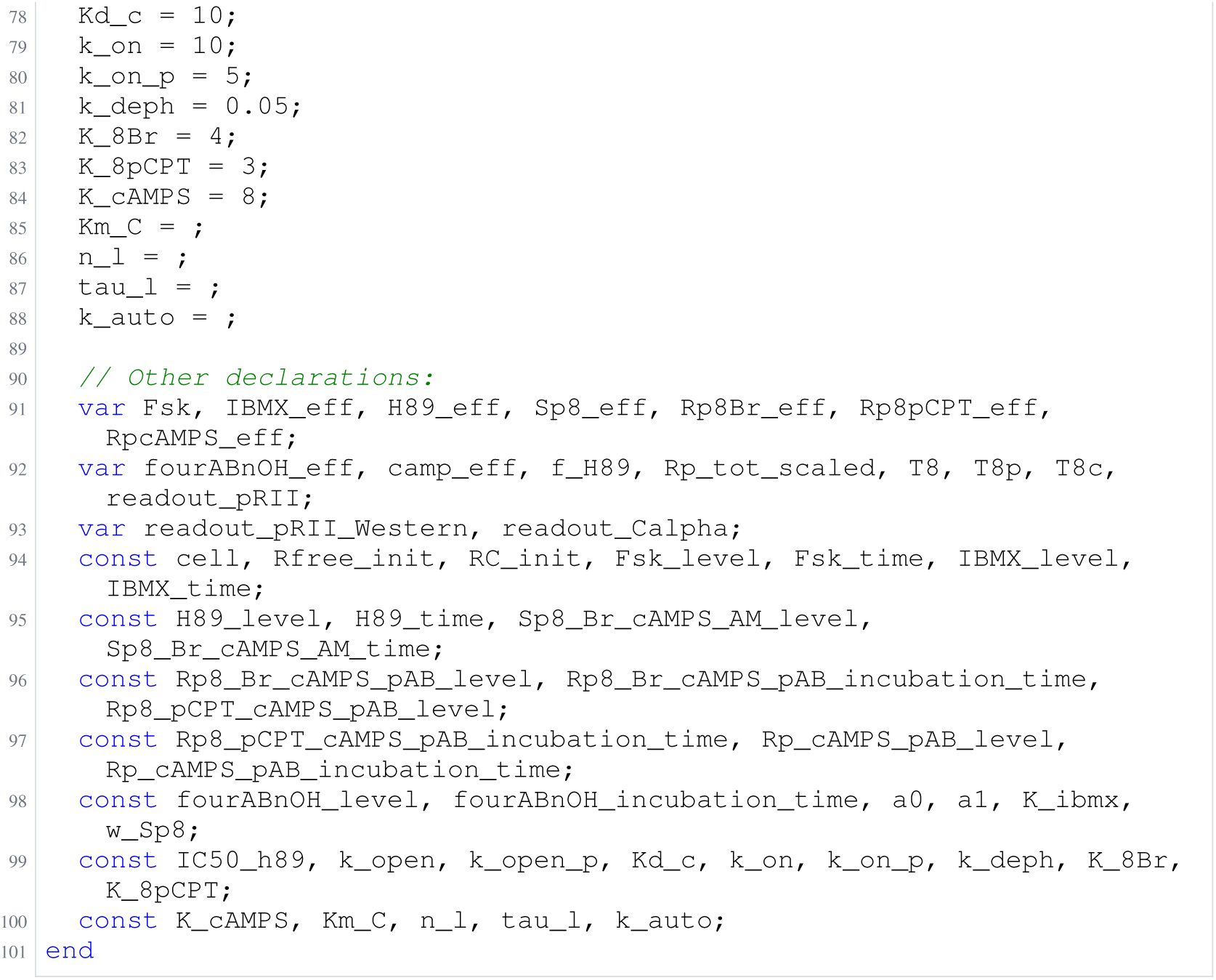
Archived Antimony for isensee_4; task 56836711_0; selected version 17.

**Listing 103:**
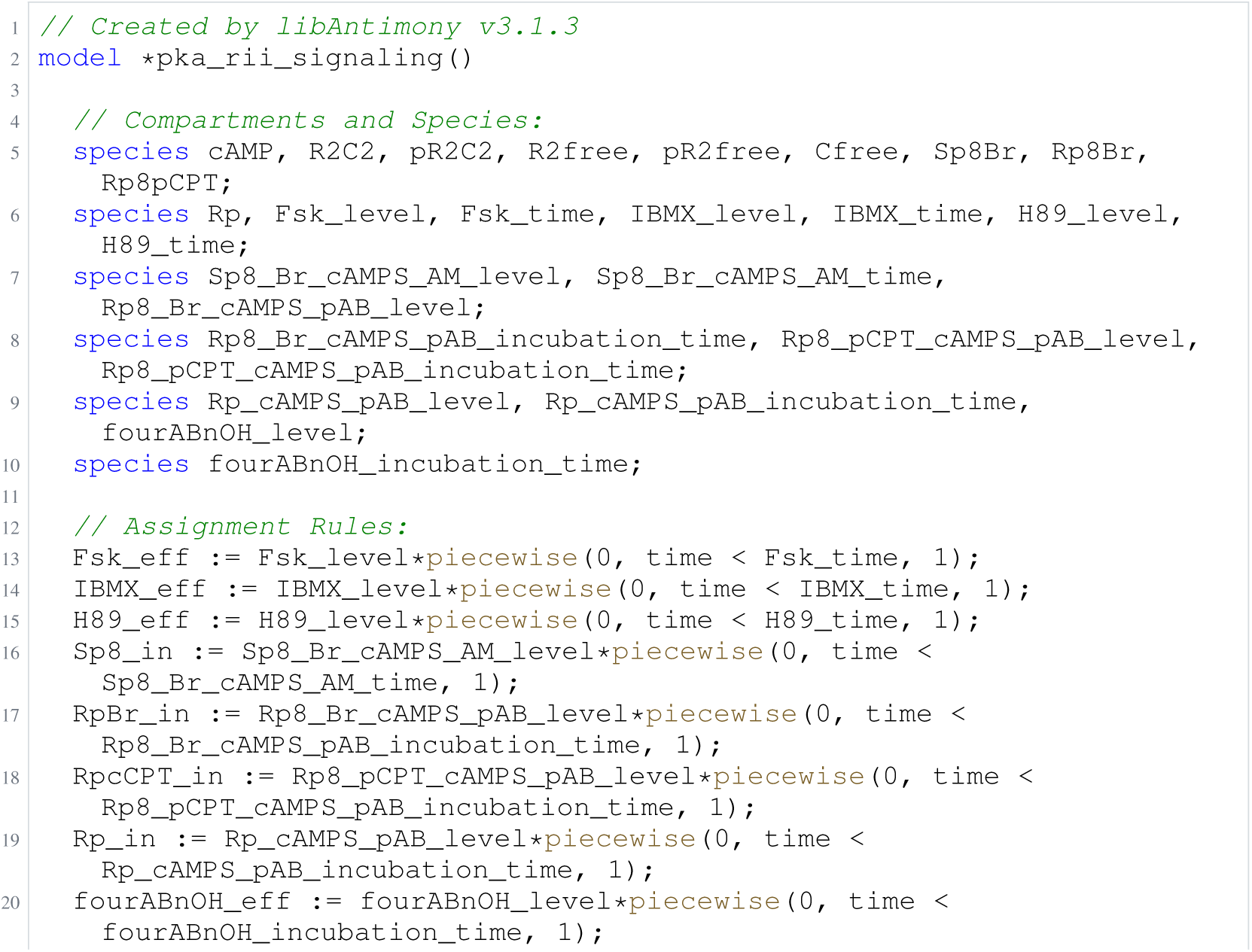
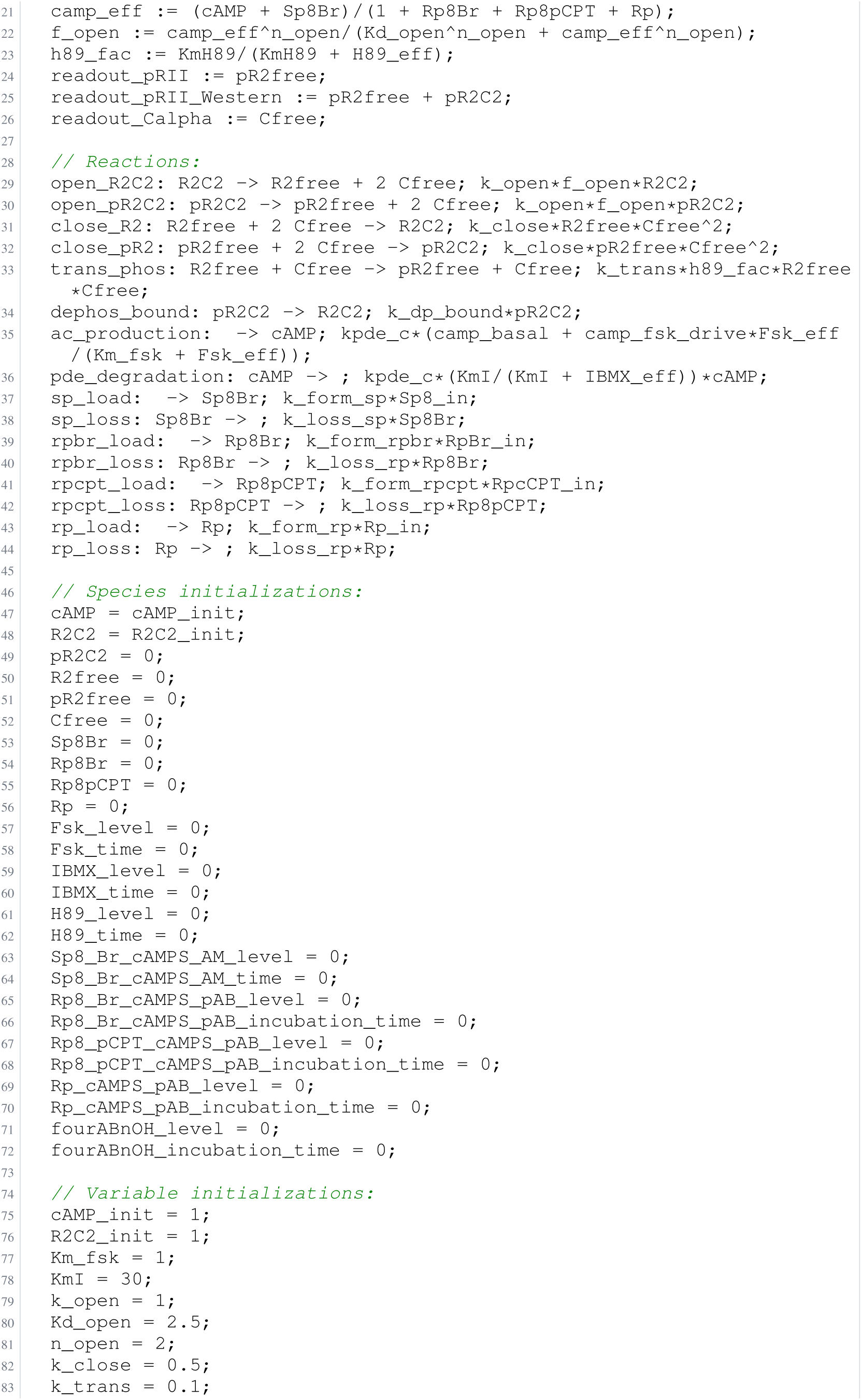
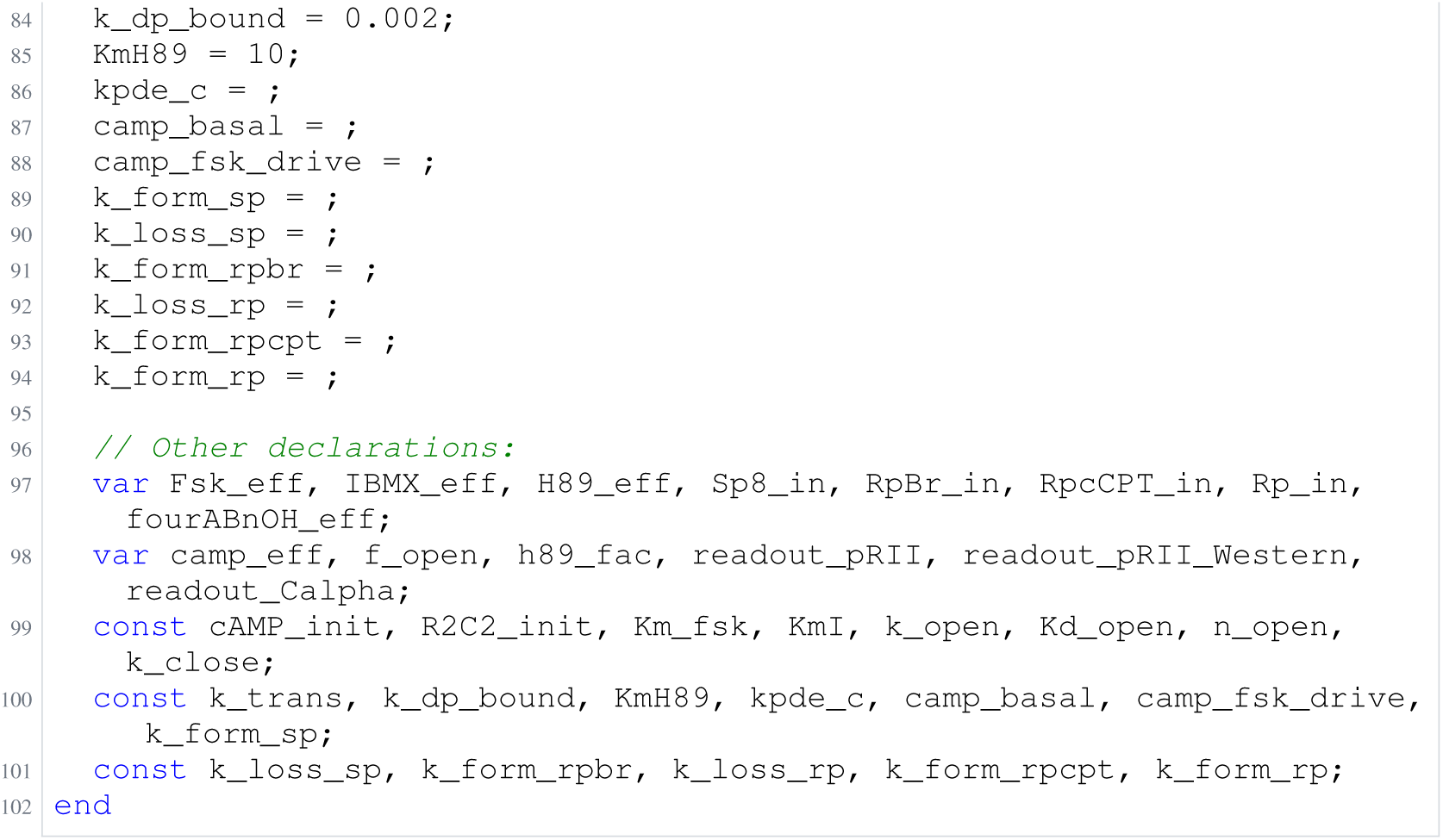
Archived Antimony for isensee_5; task 58312543_0; selected version 1.

### H.16 Alkan

**Listing 104:**
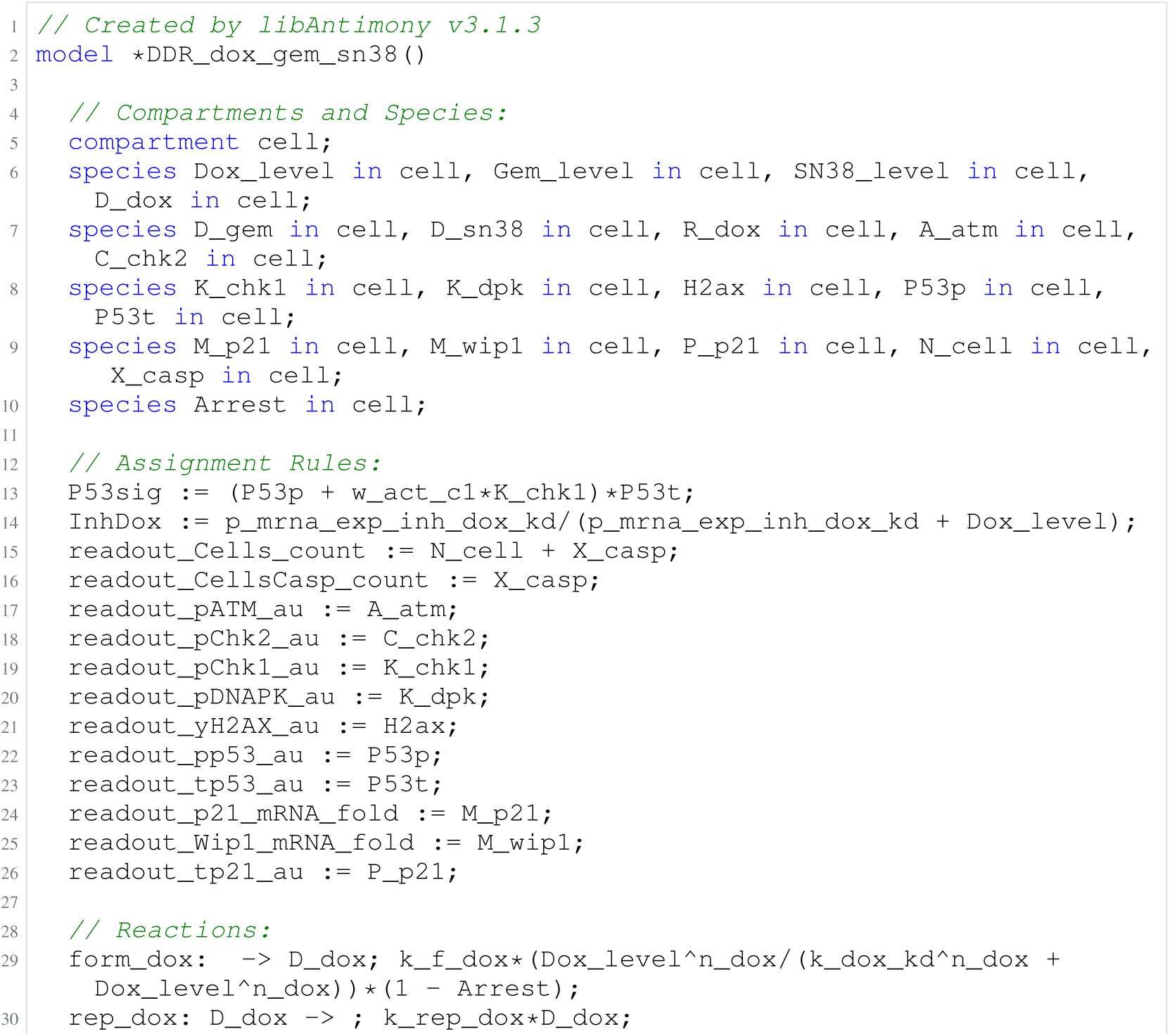
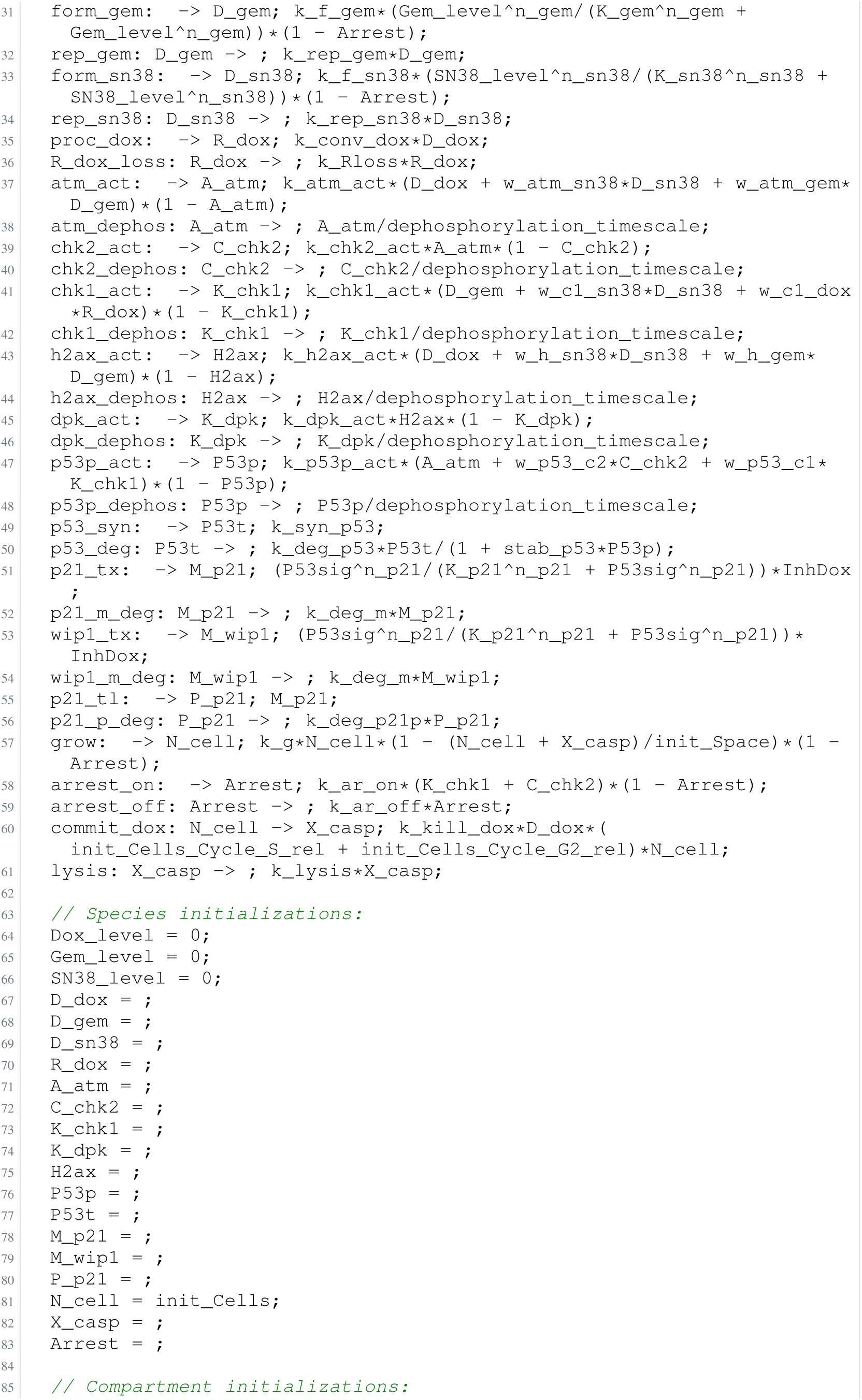
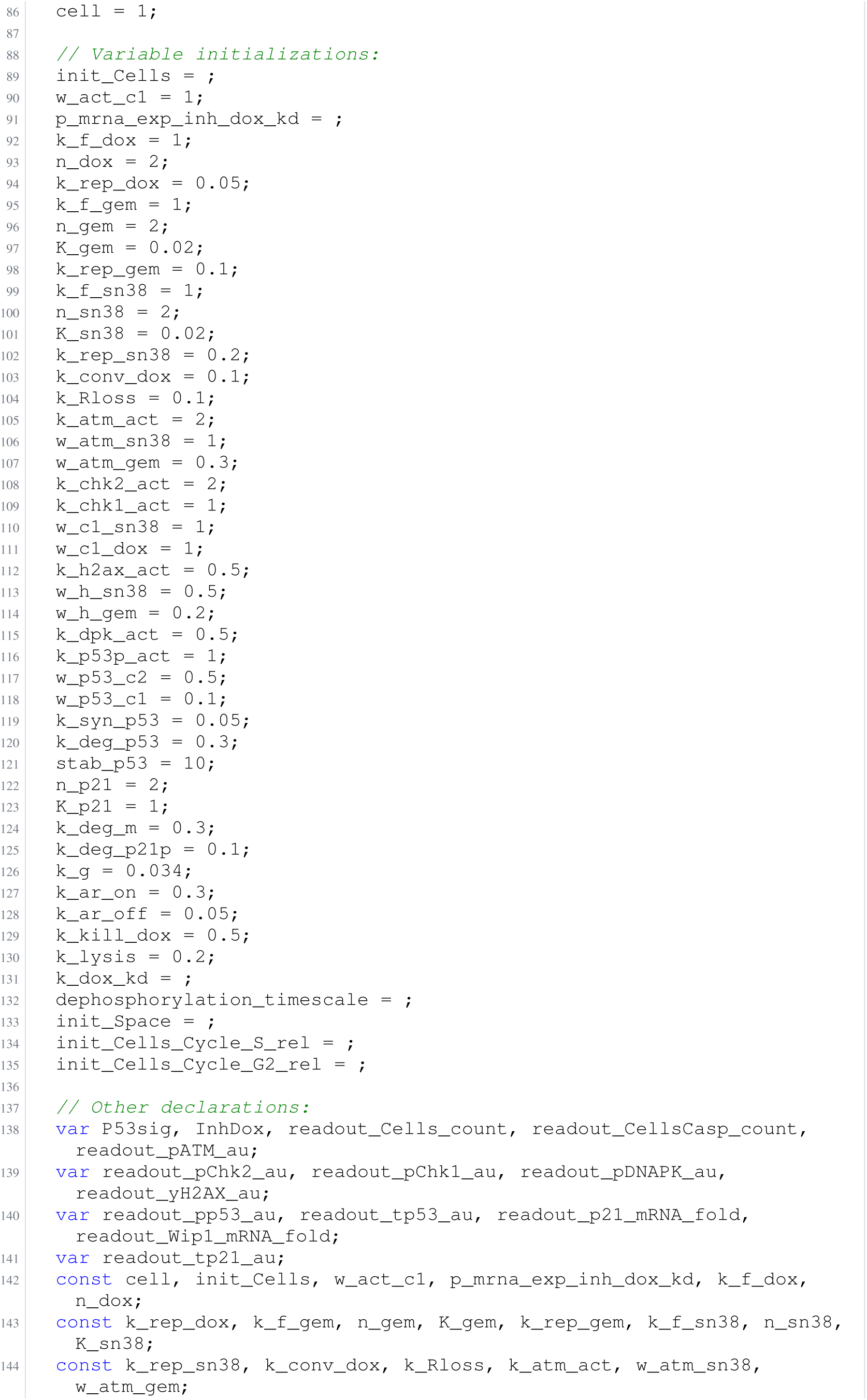
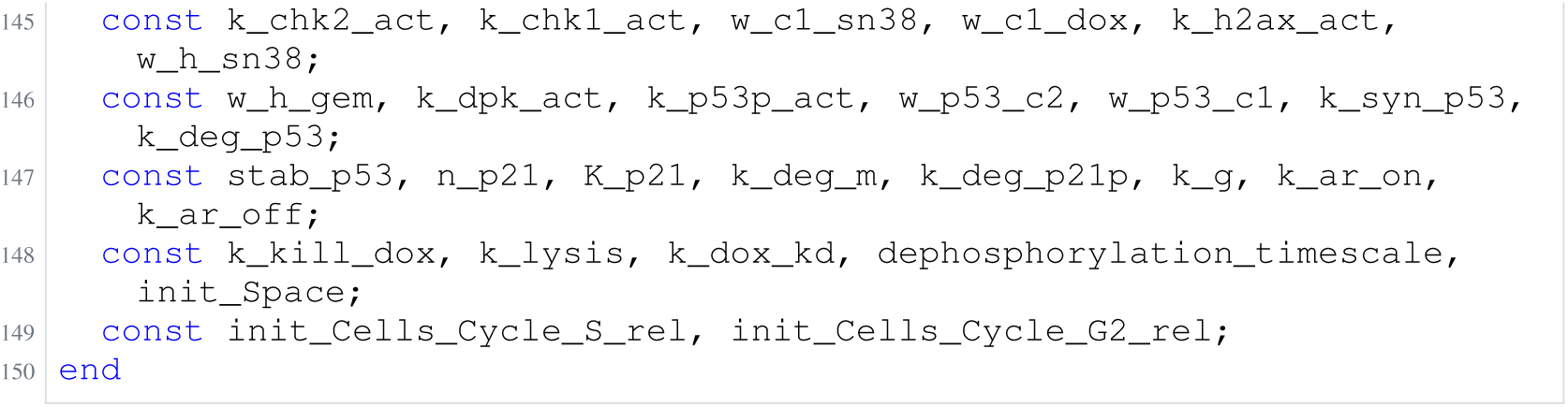
Archived Antimony for alkan_1; task 56836792_0; selected version 0.

**Listing 105:**
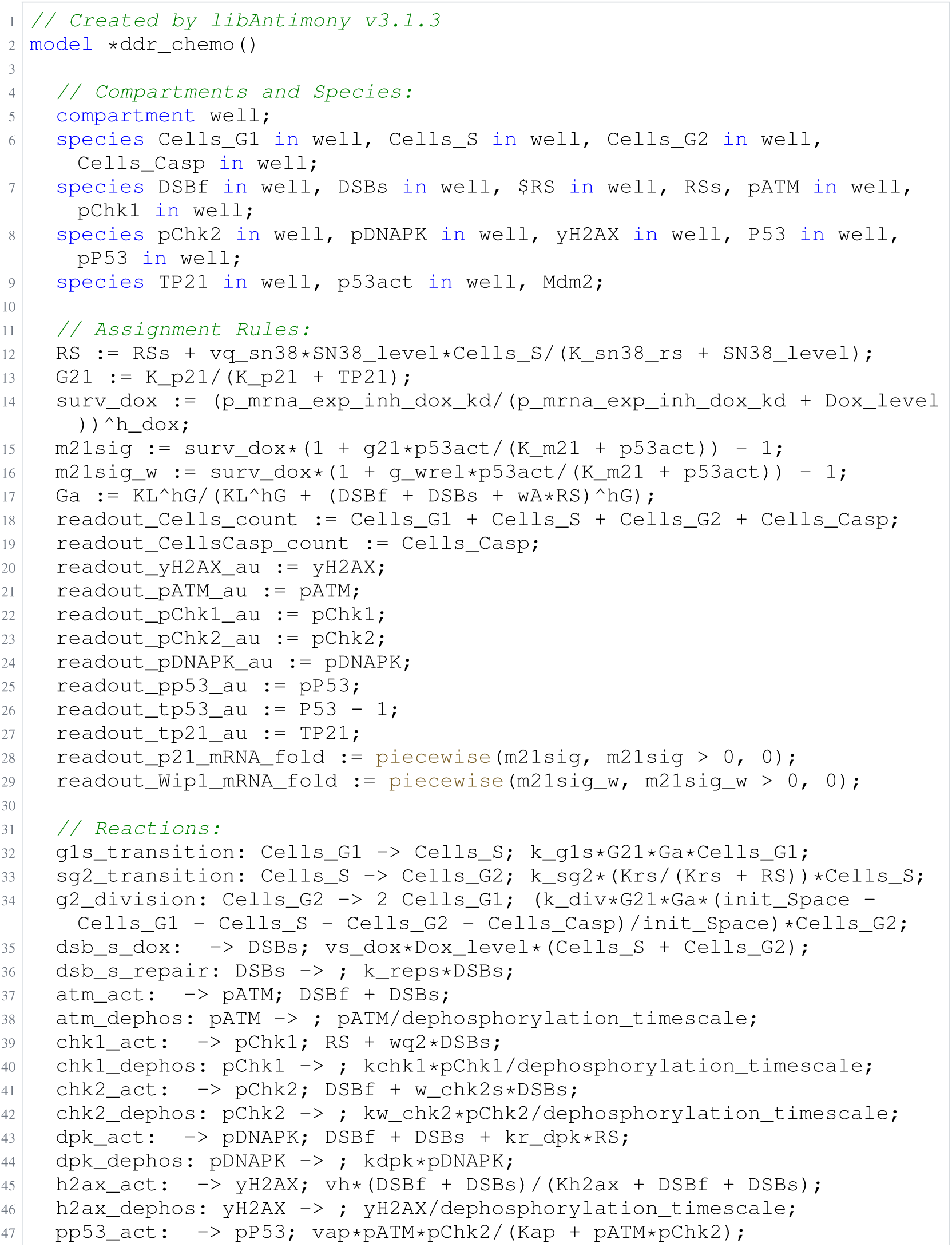
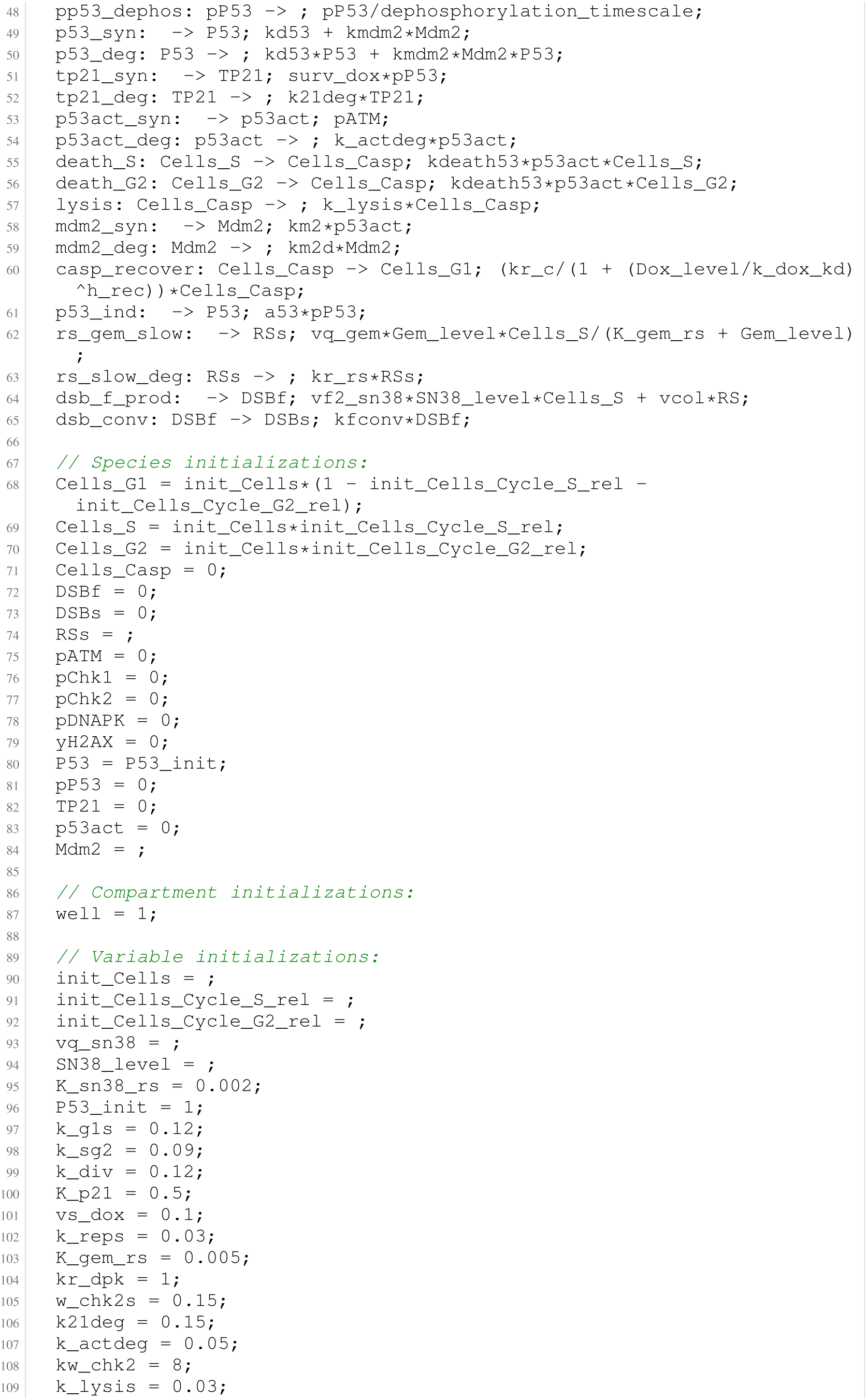
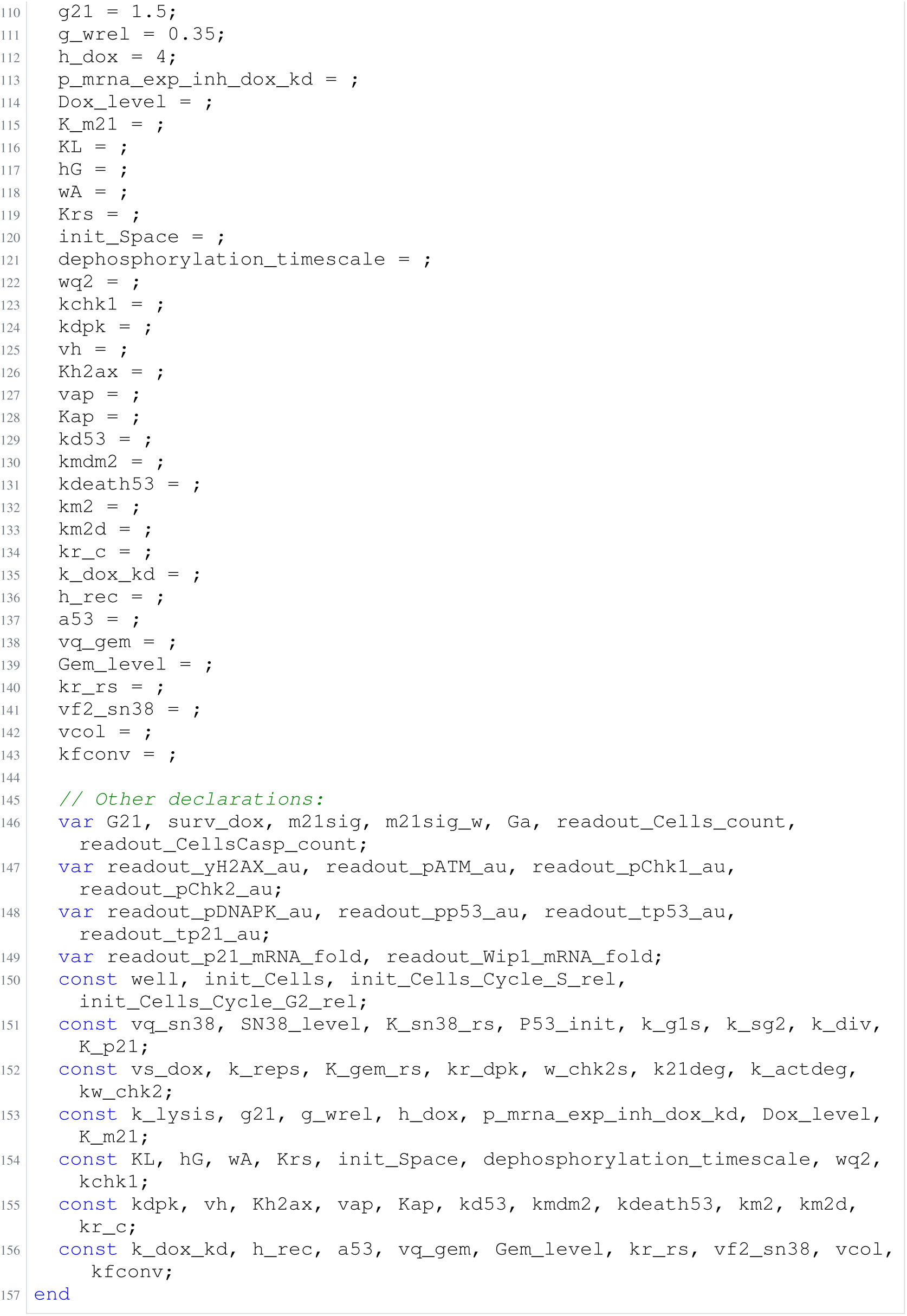
Archived Antimony for alkan_2; task 56836792_1; selected version 15.

**Listing 106:**
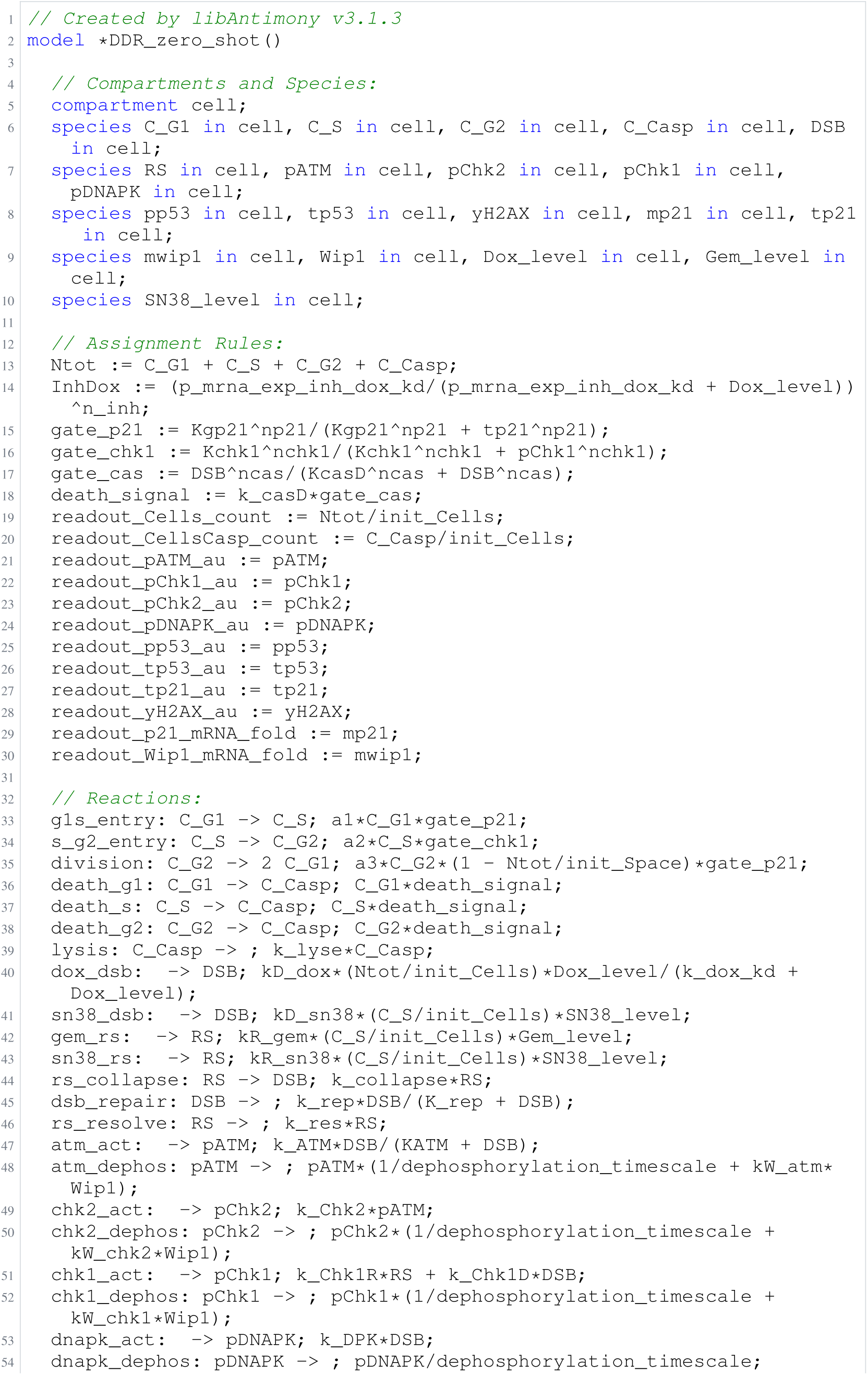
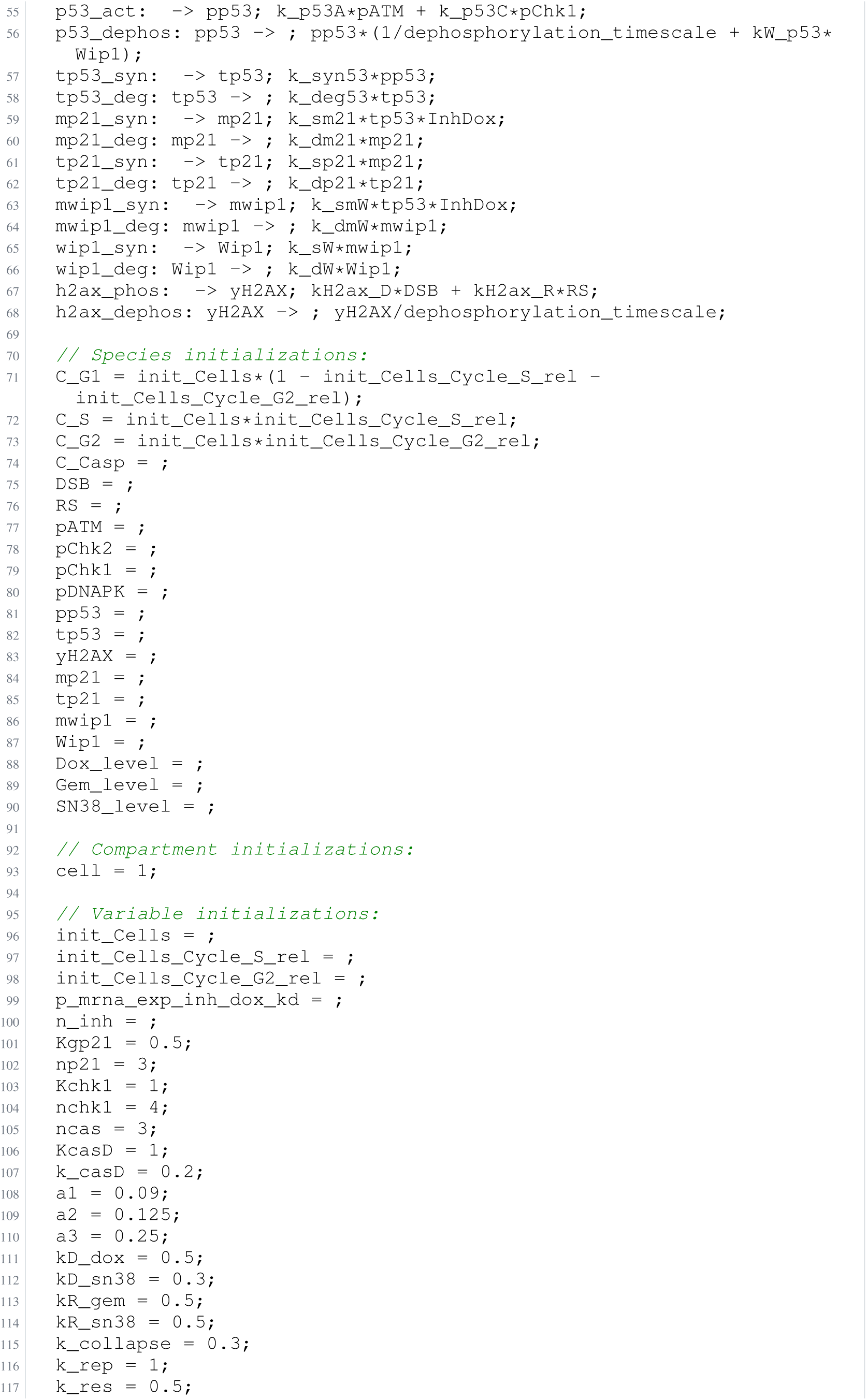
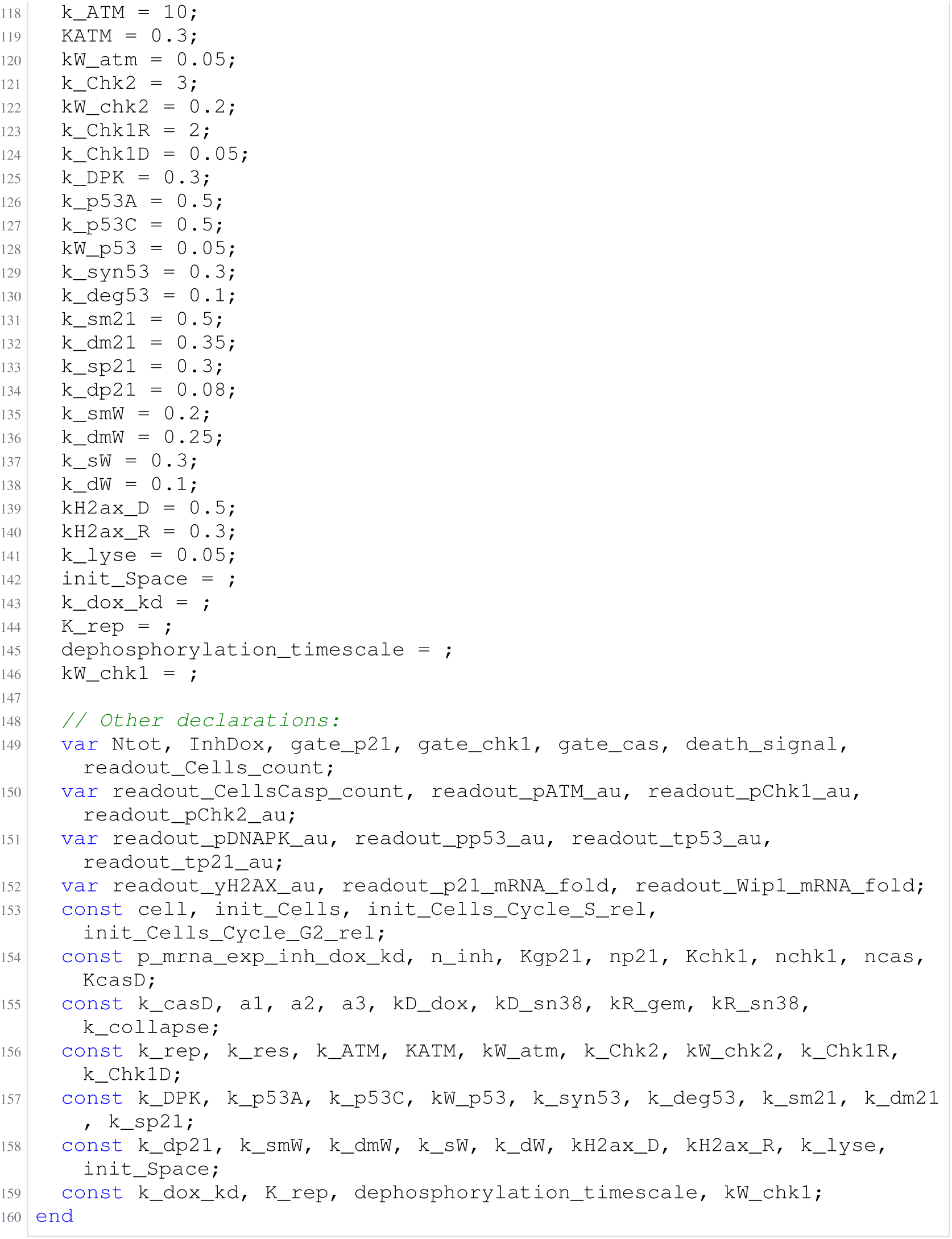
Archived Antimony for alkan_3; task 56836792_2; selected version 2.

**Listing 107:**
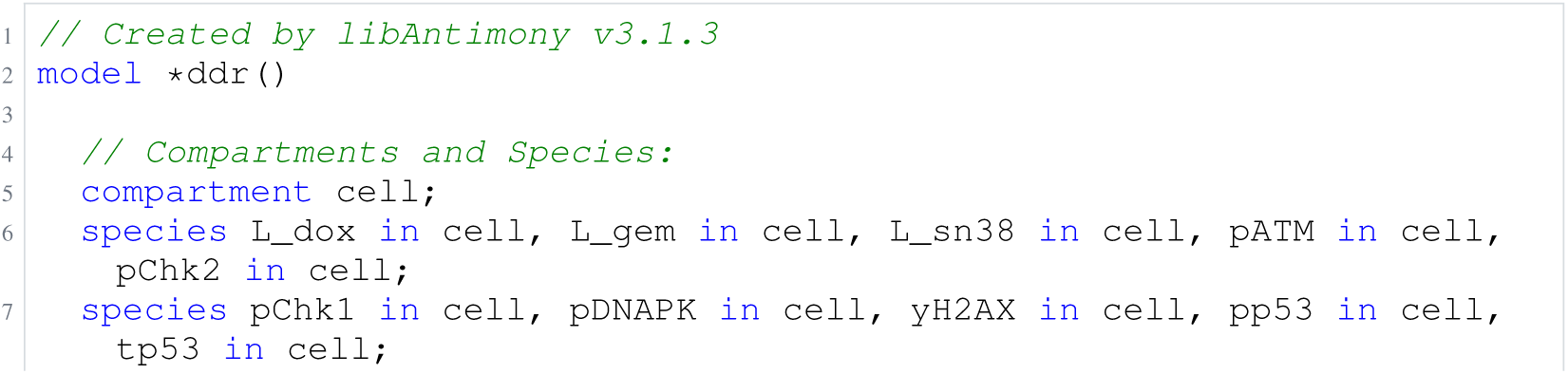
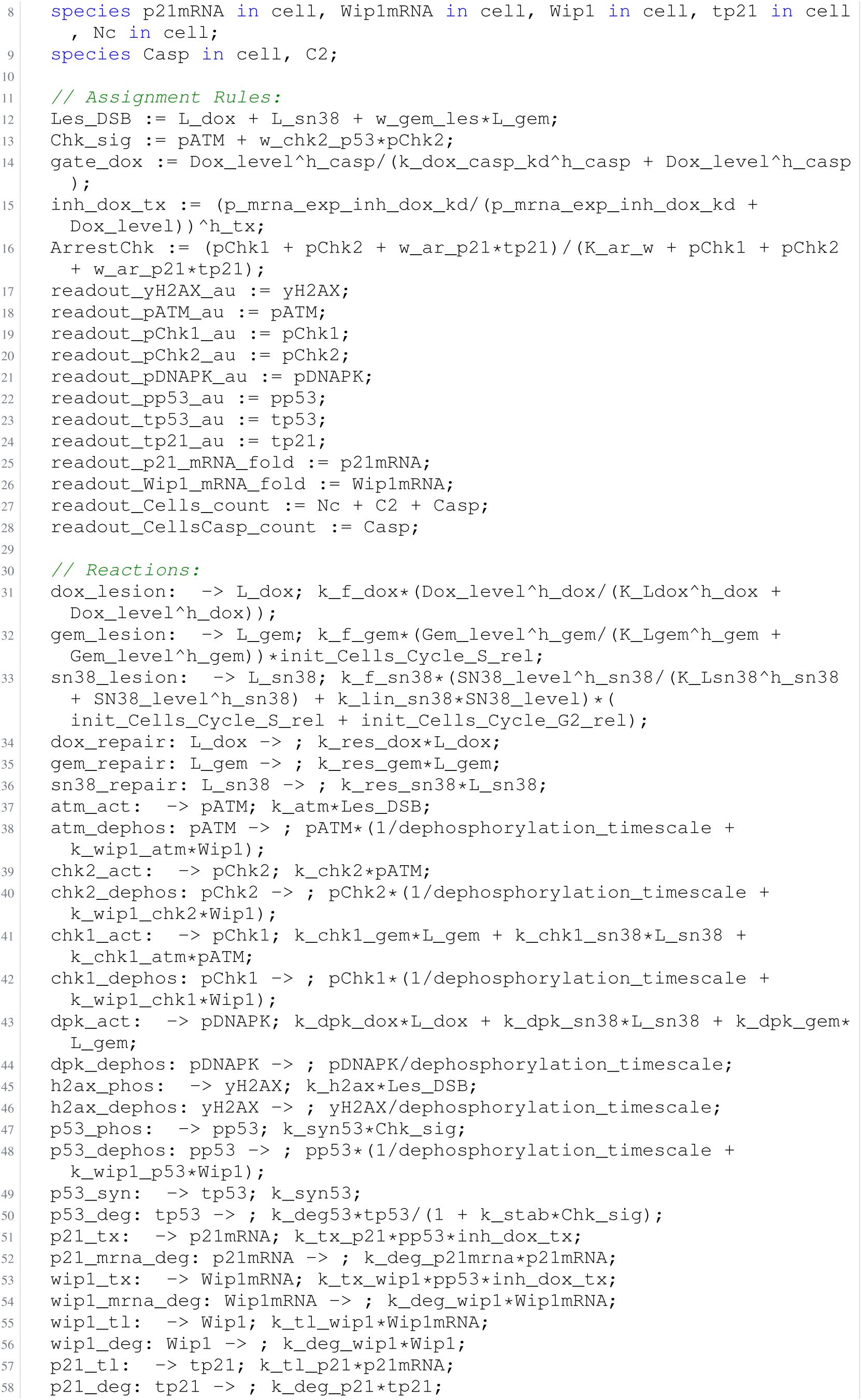
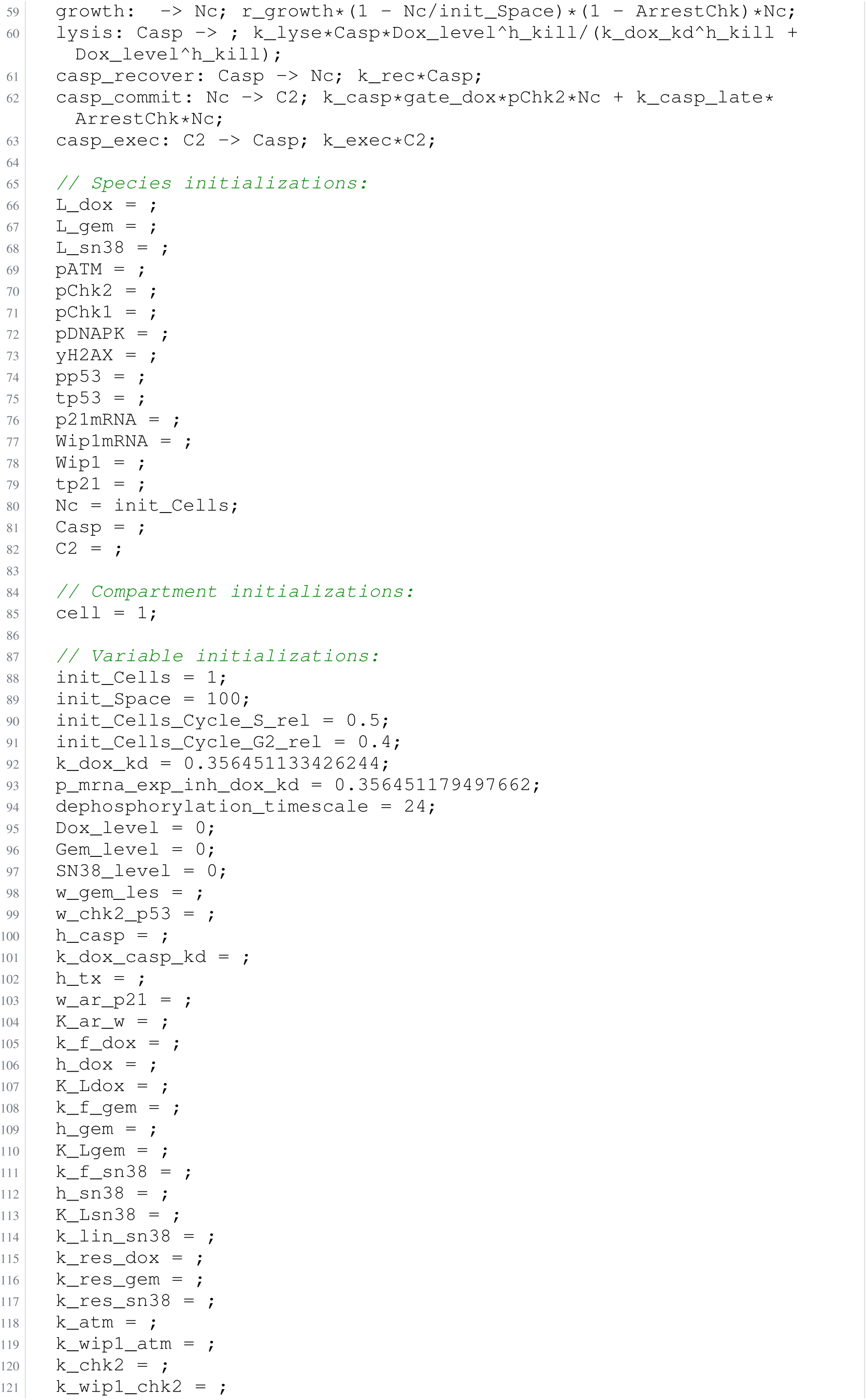
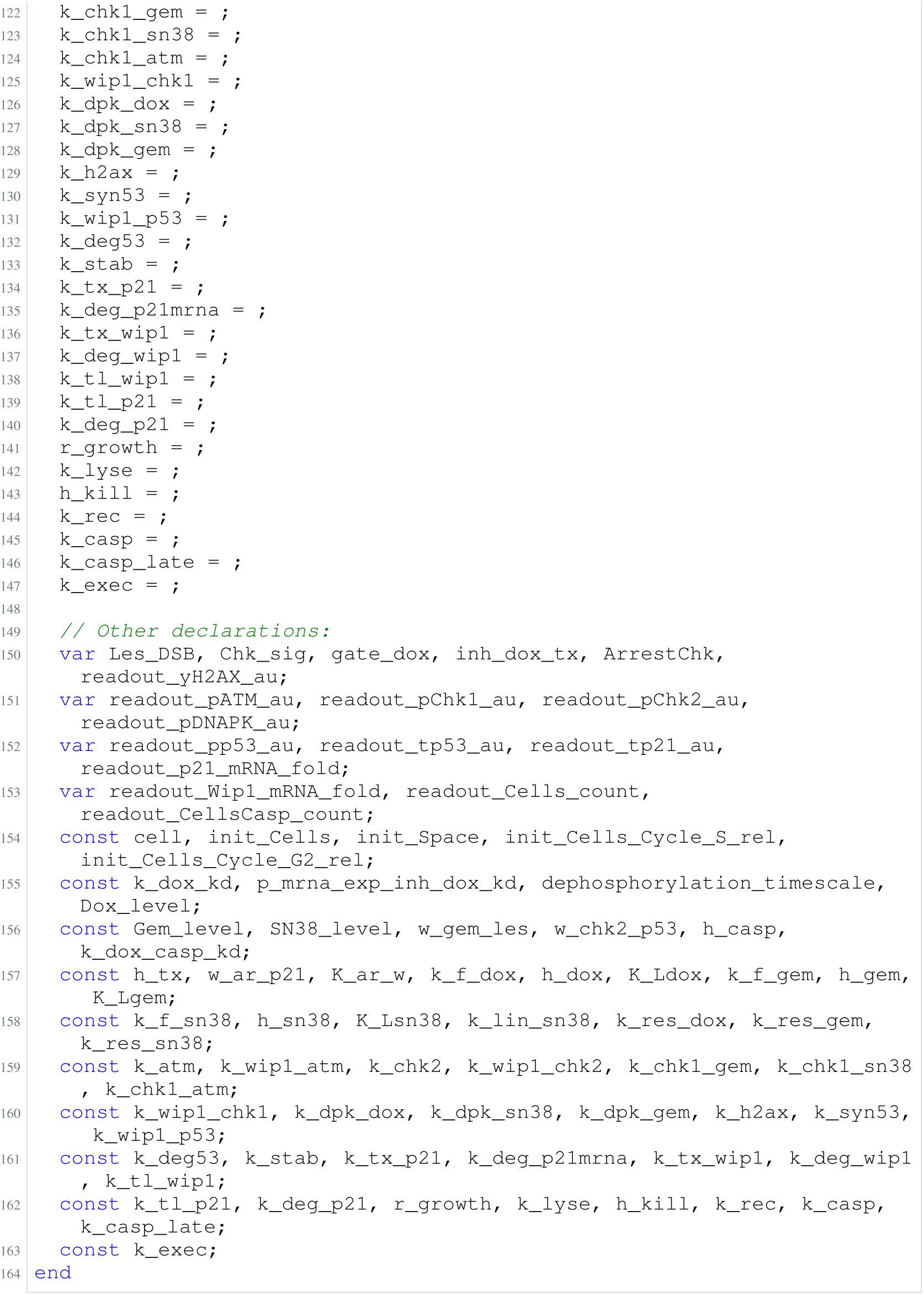
Archived Antimony for alkan_4; task 56836792_3; selected version 9.

**Listing 108:**
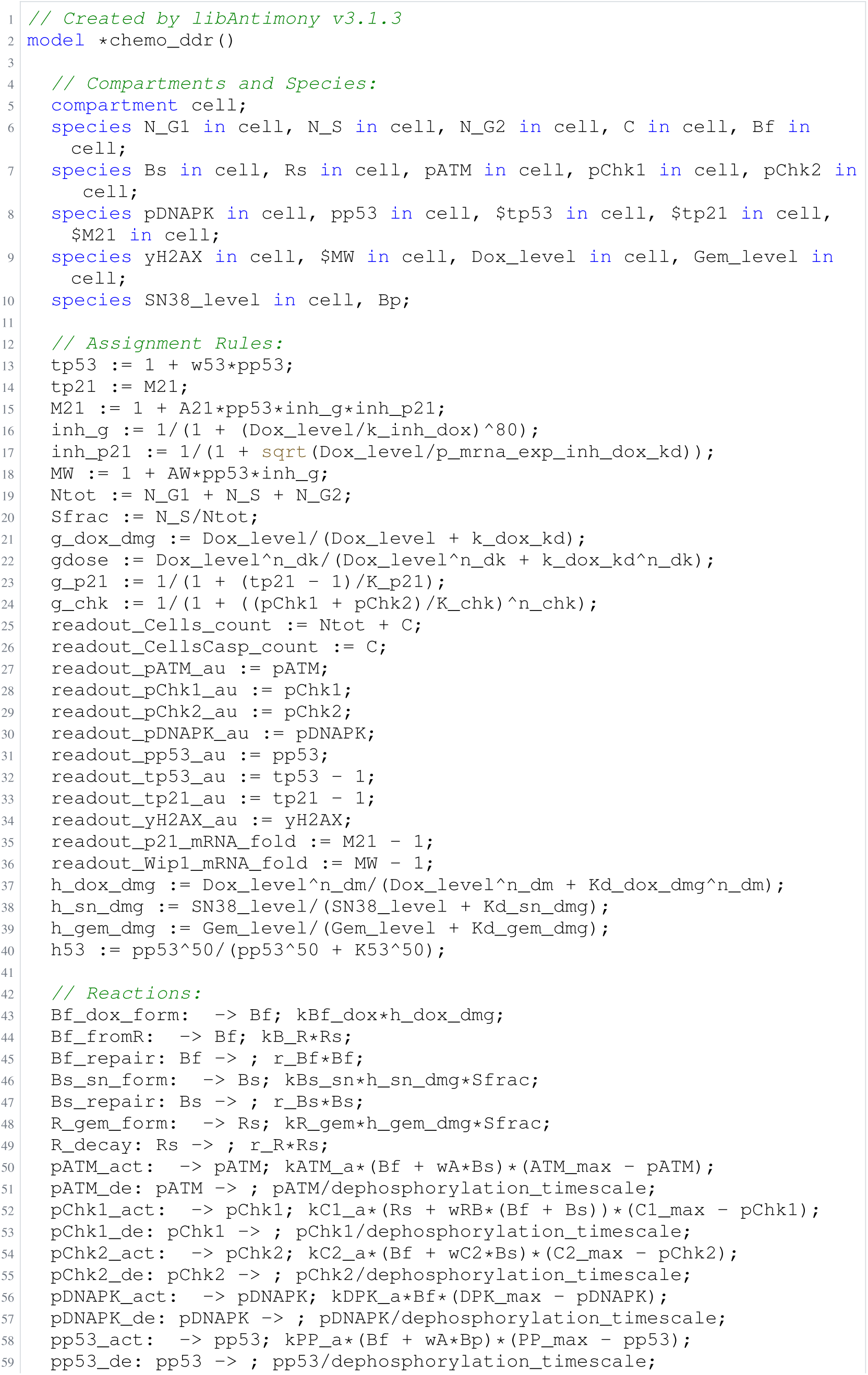
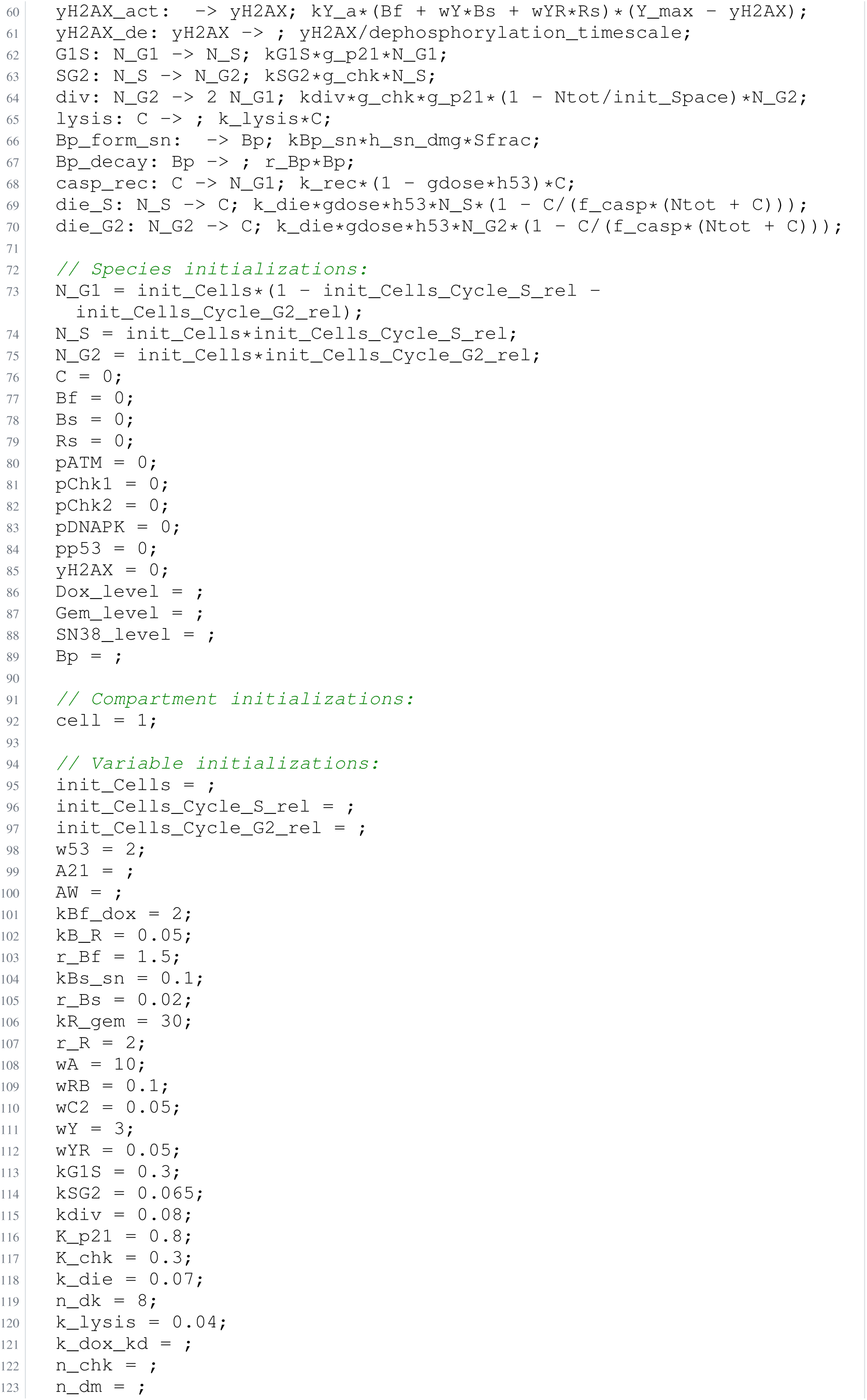
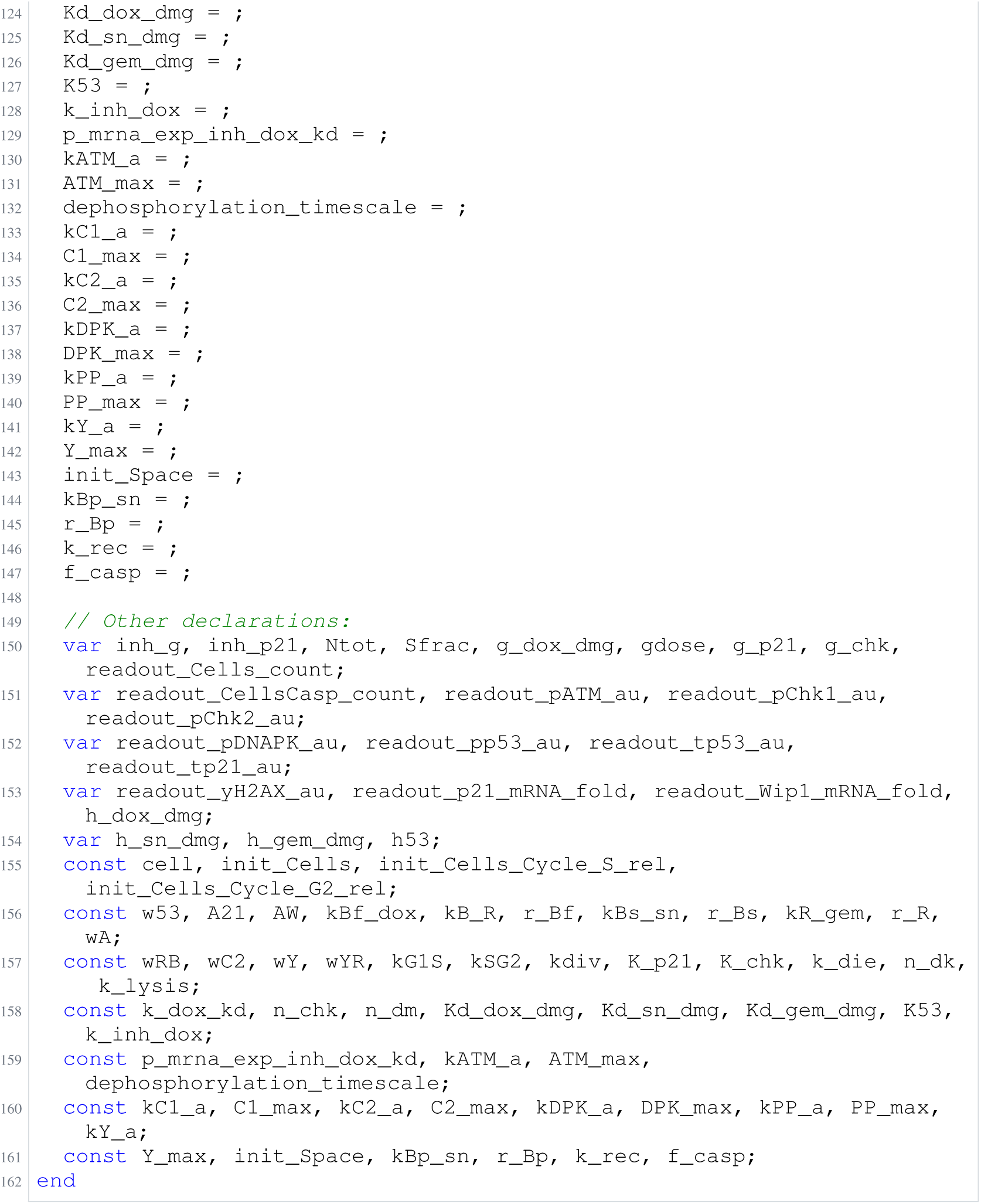
Archived Antimony for alkan_5; task 56836792_4; selected version 15.

### H.17 Lucarelli

**Listing 109:**
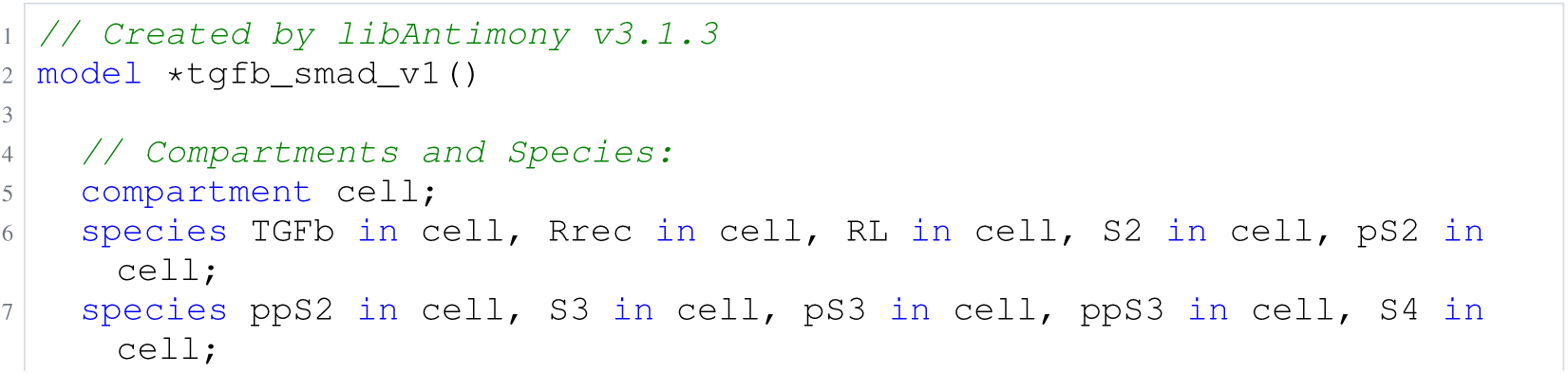
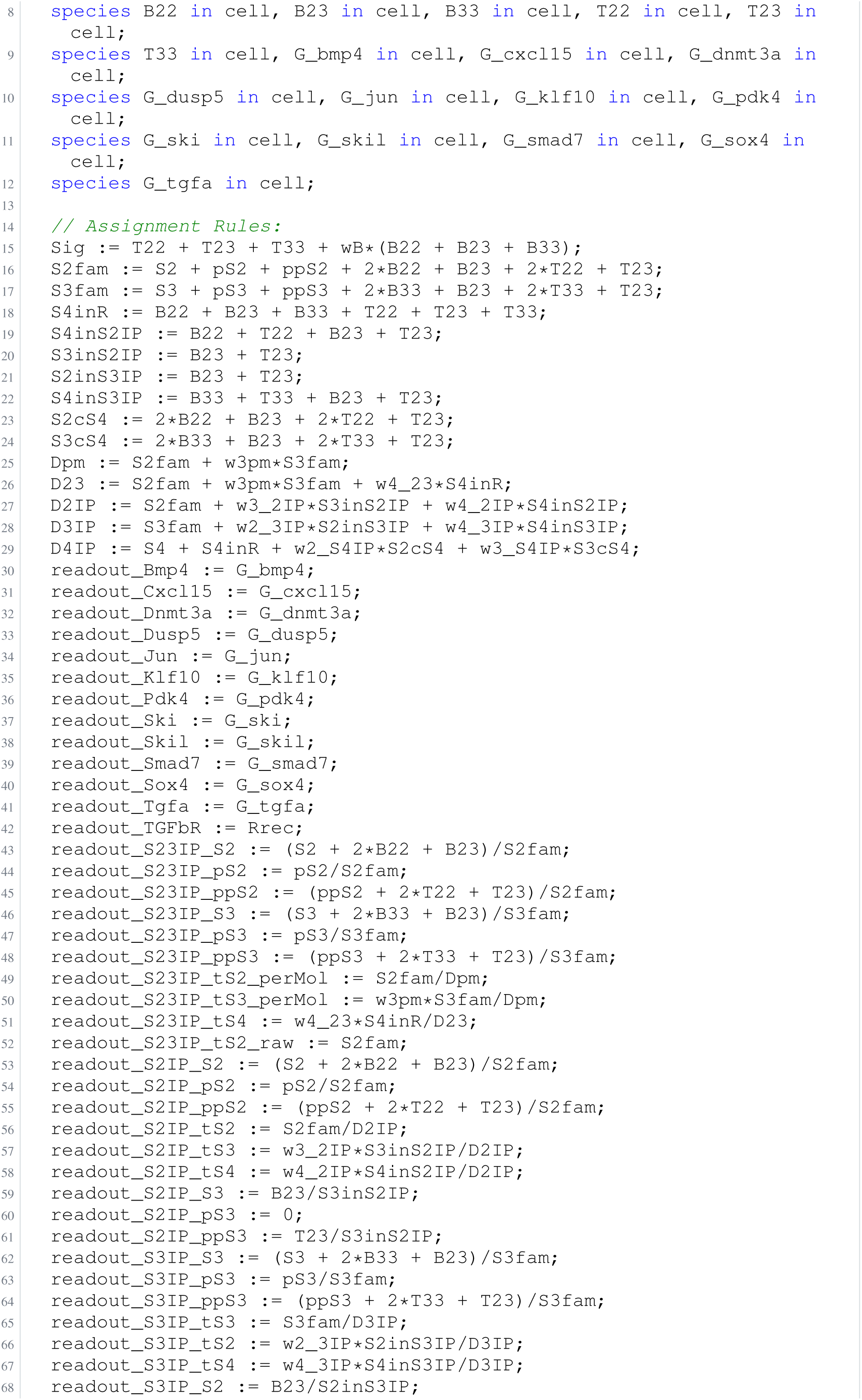
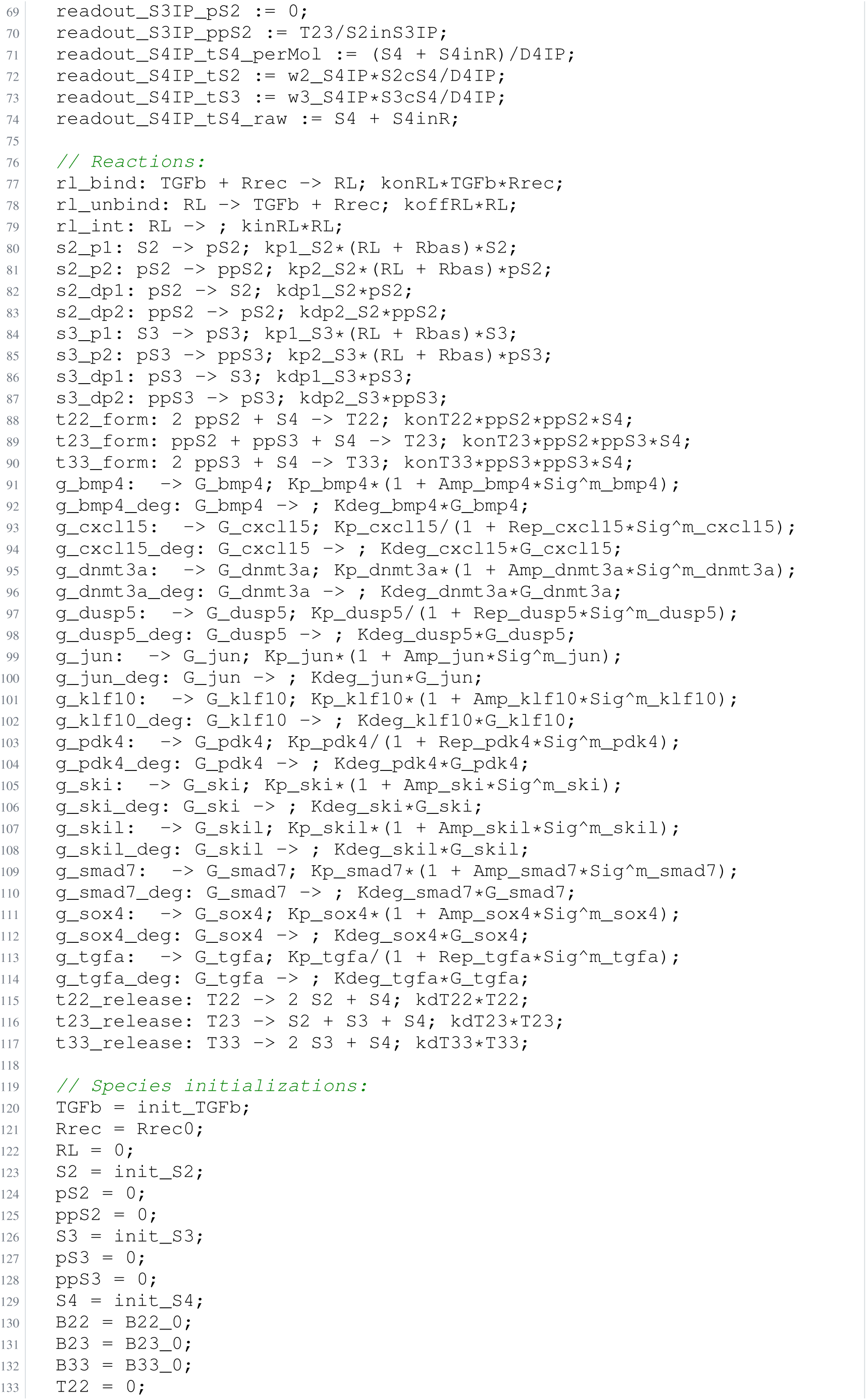
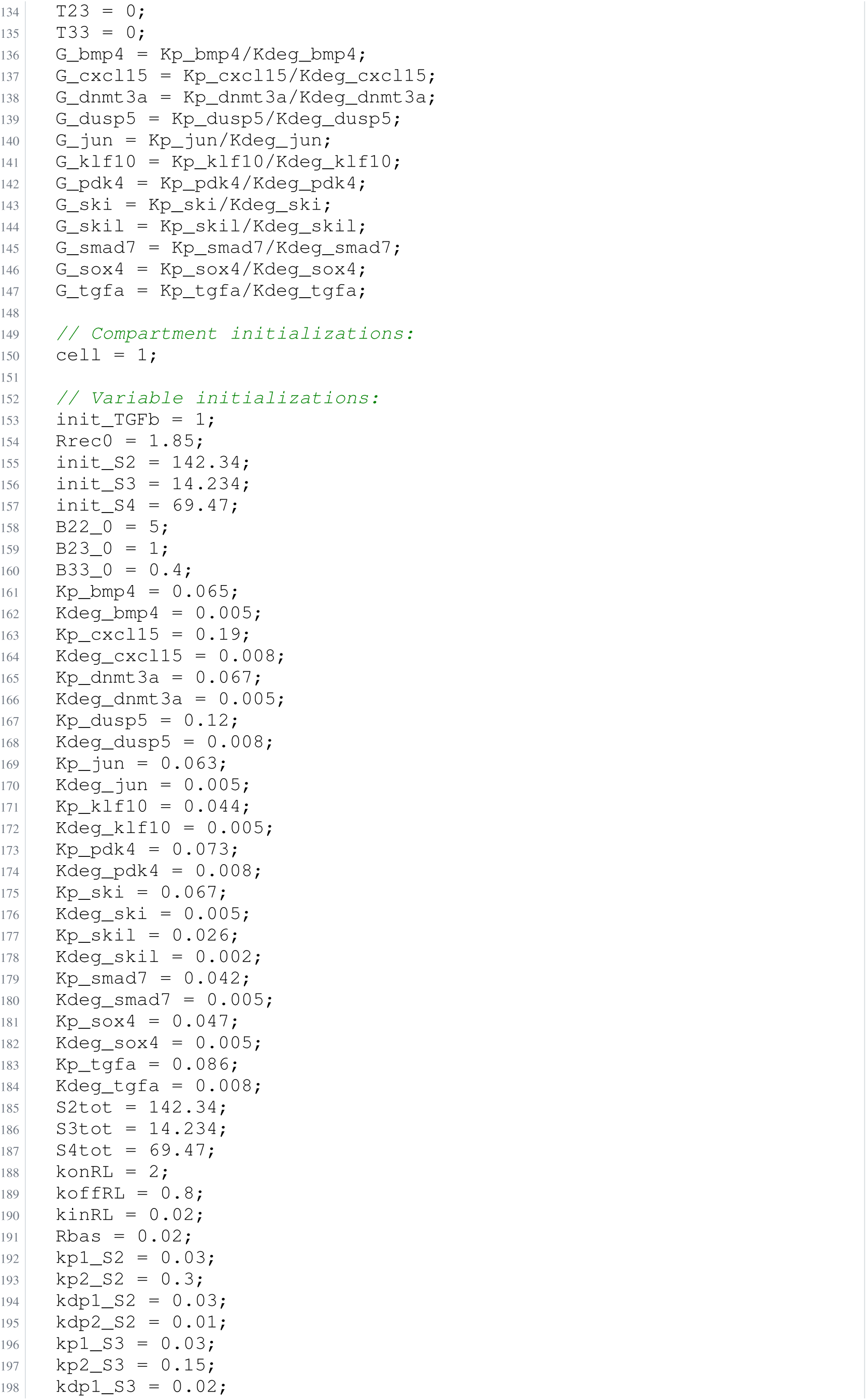
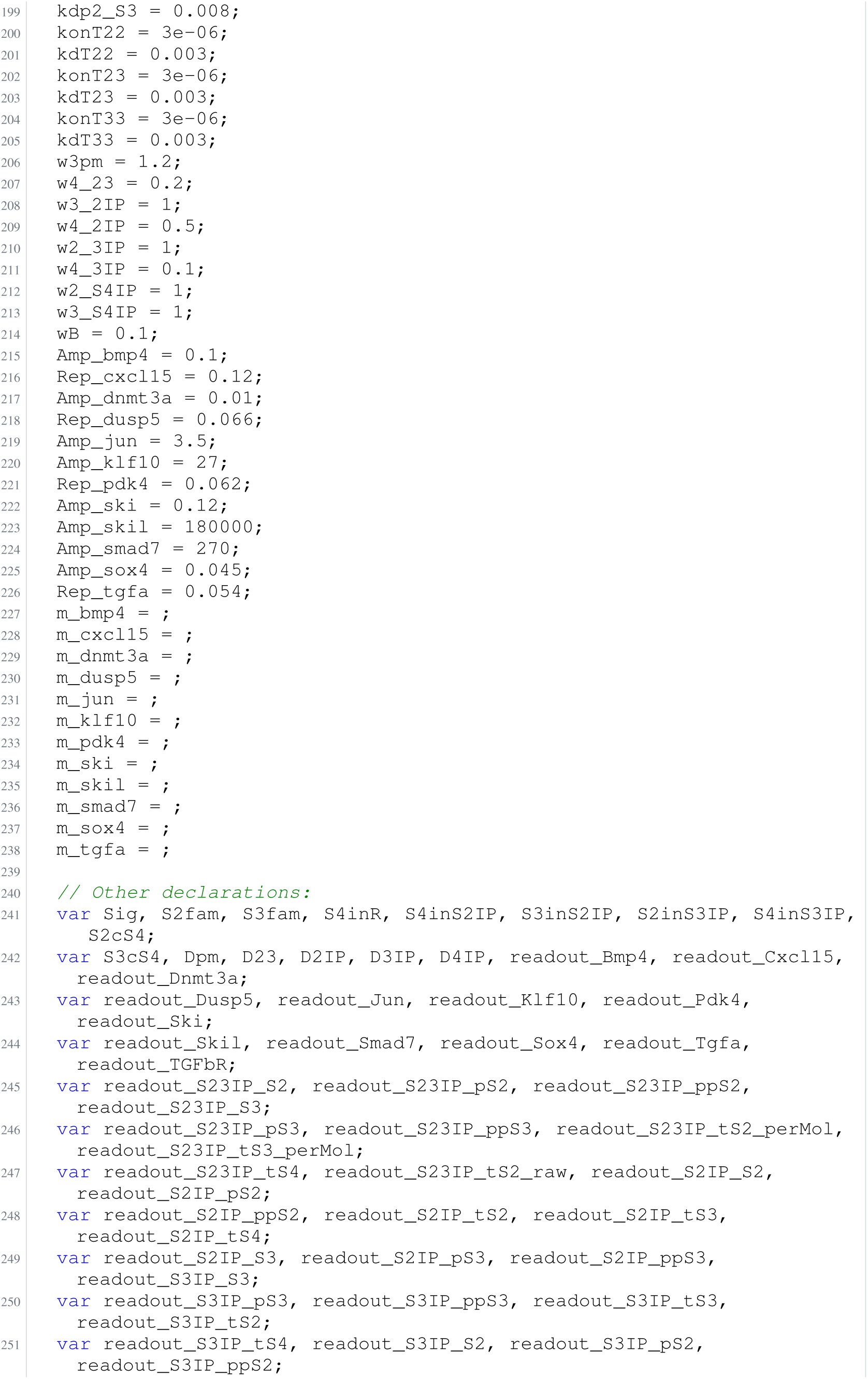
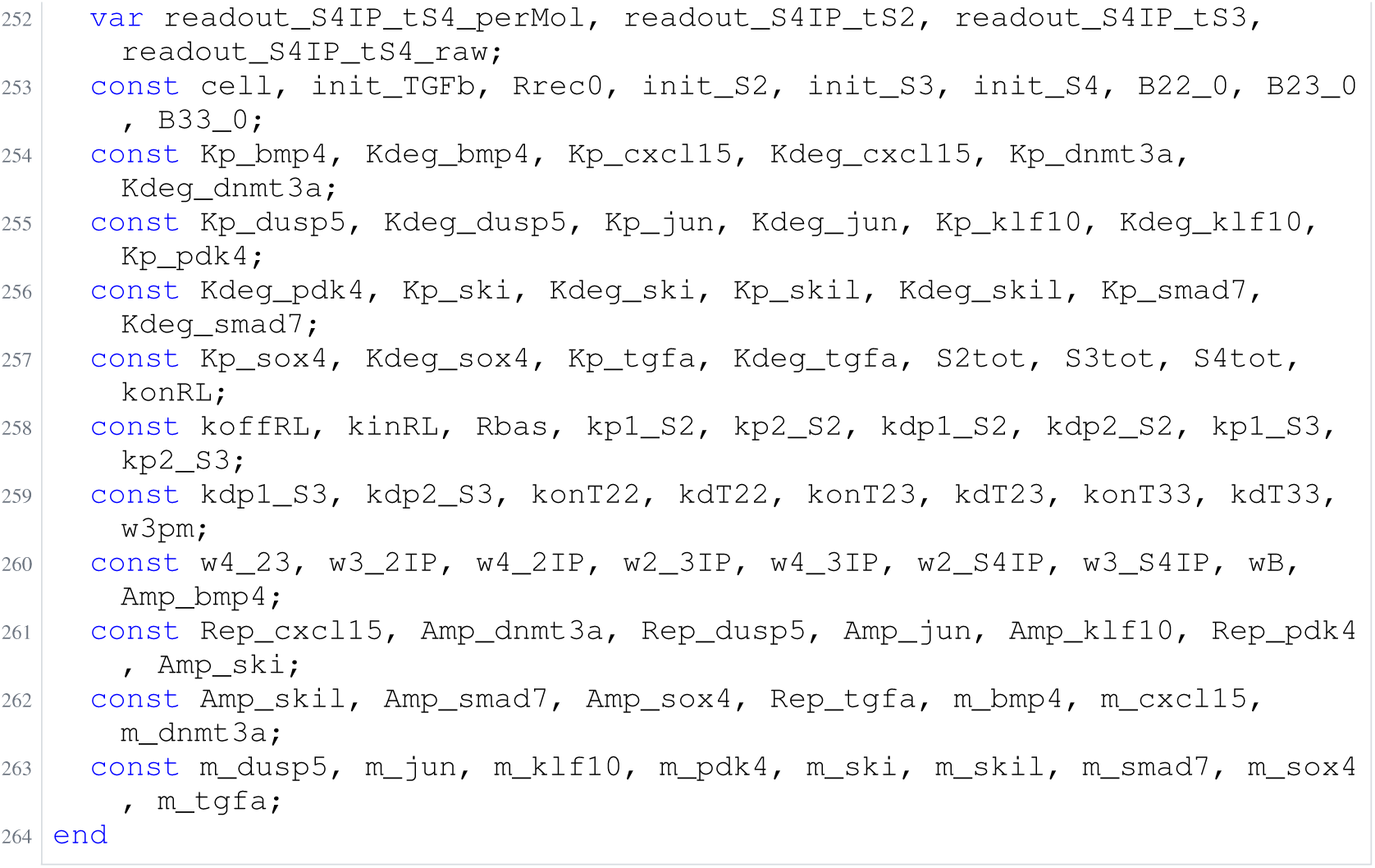
Archived Antimony for lucarelli_1; task 56836792_10; selected version 9.

**Listing 110:**
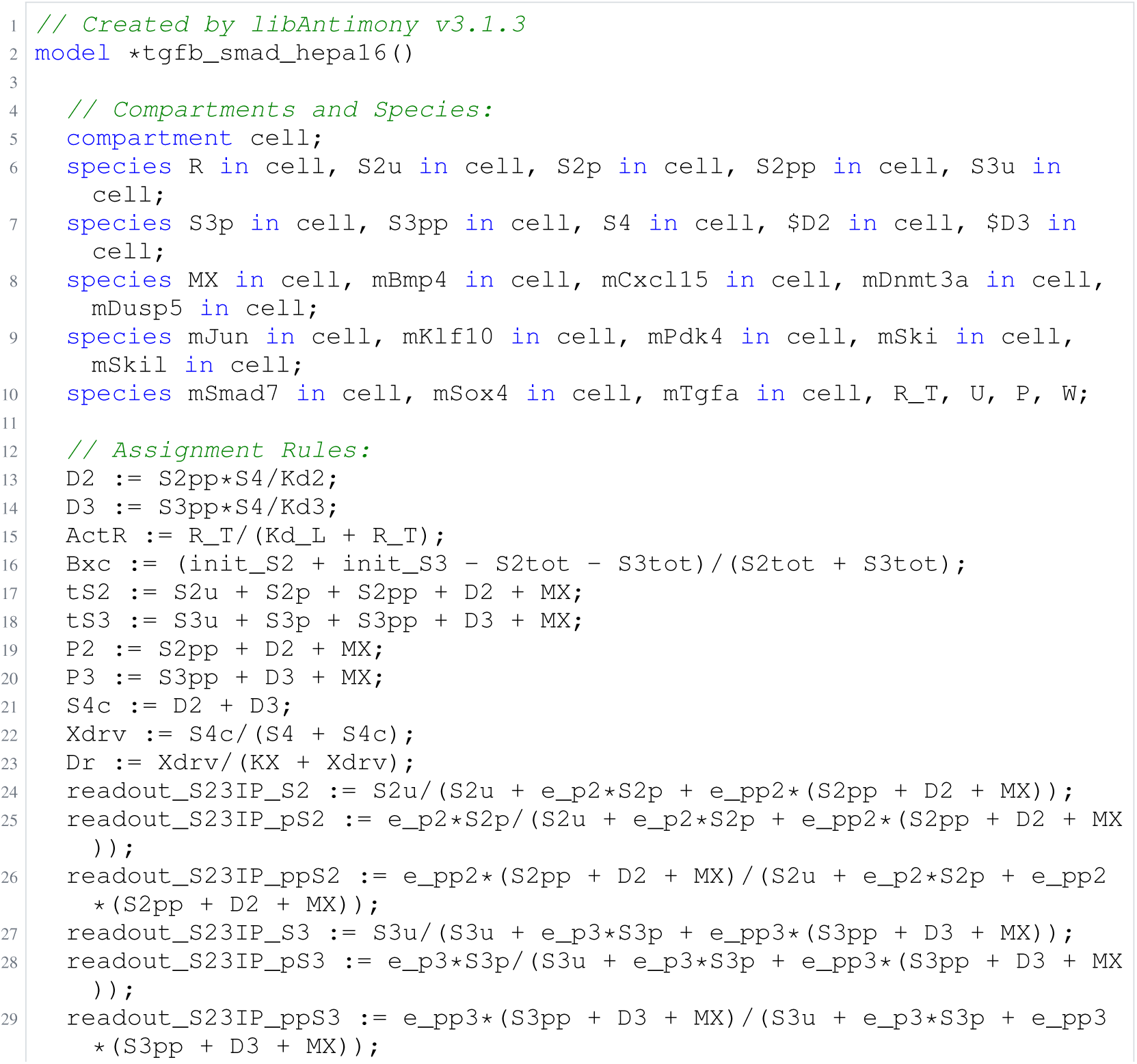
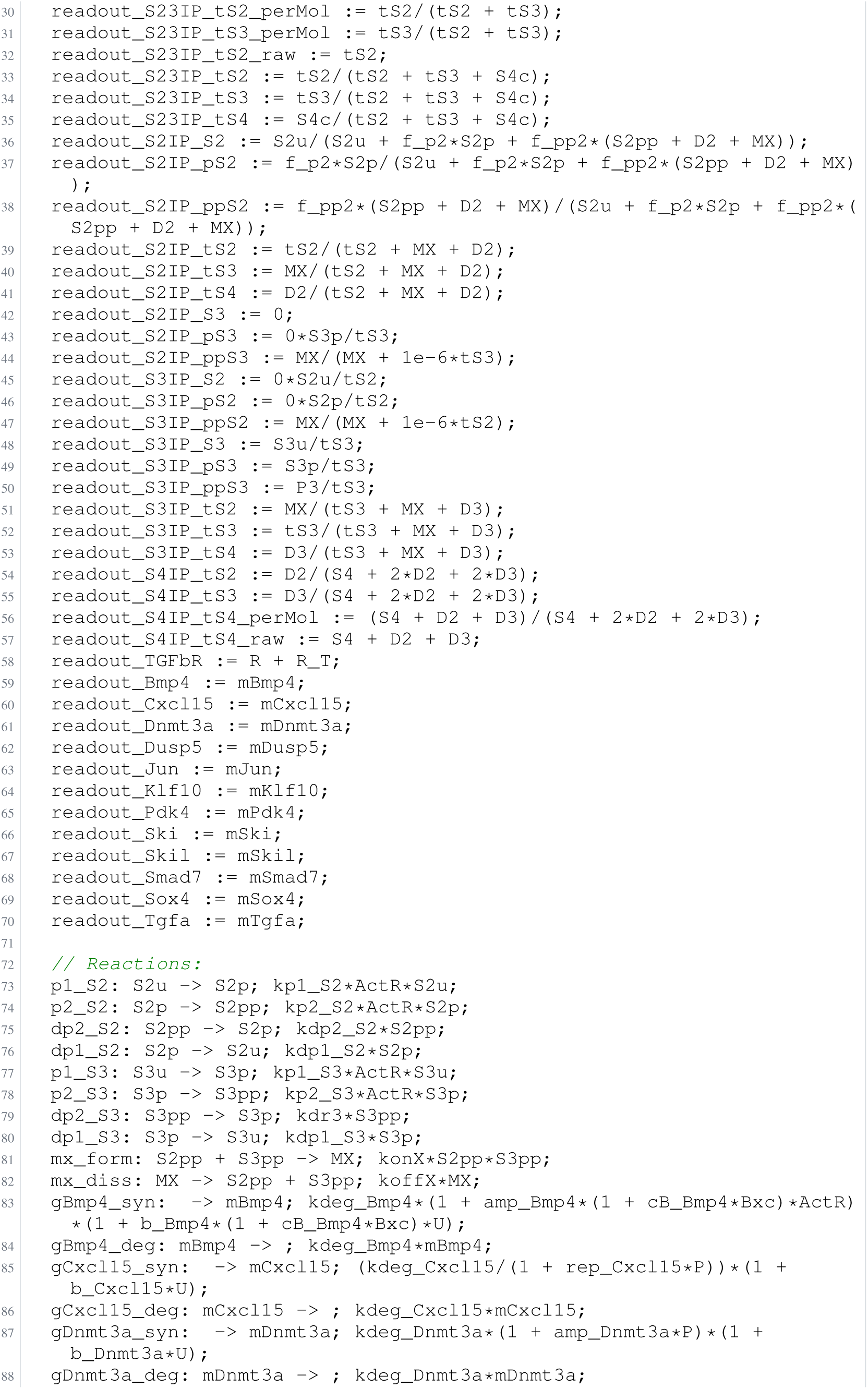
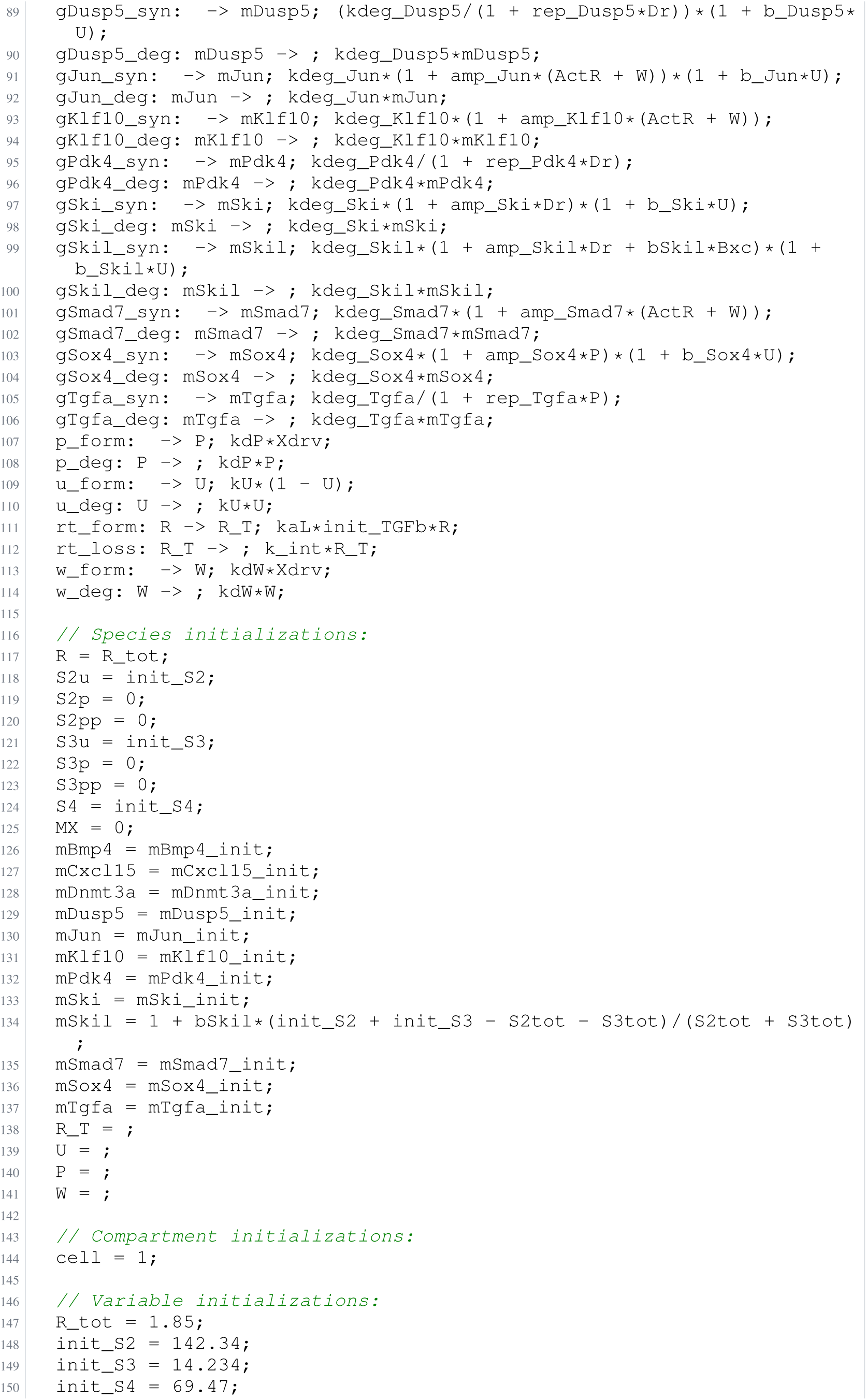
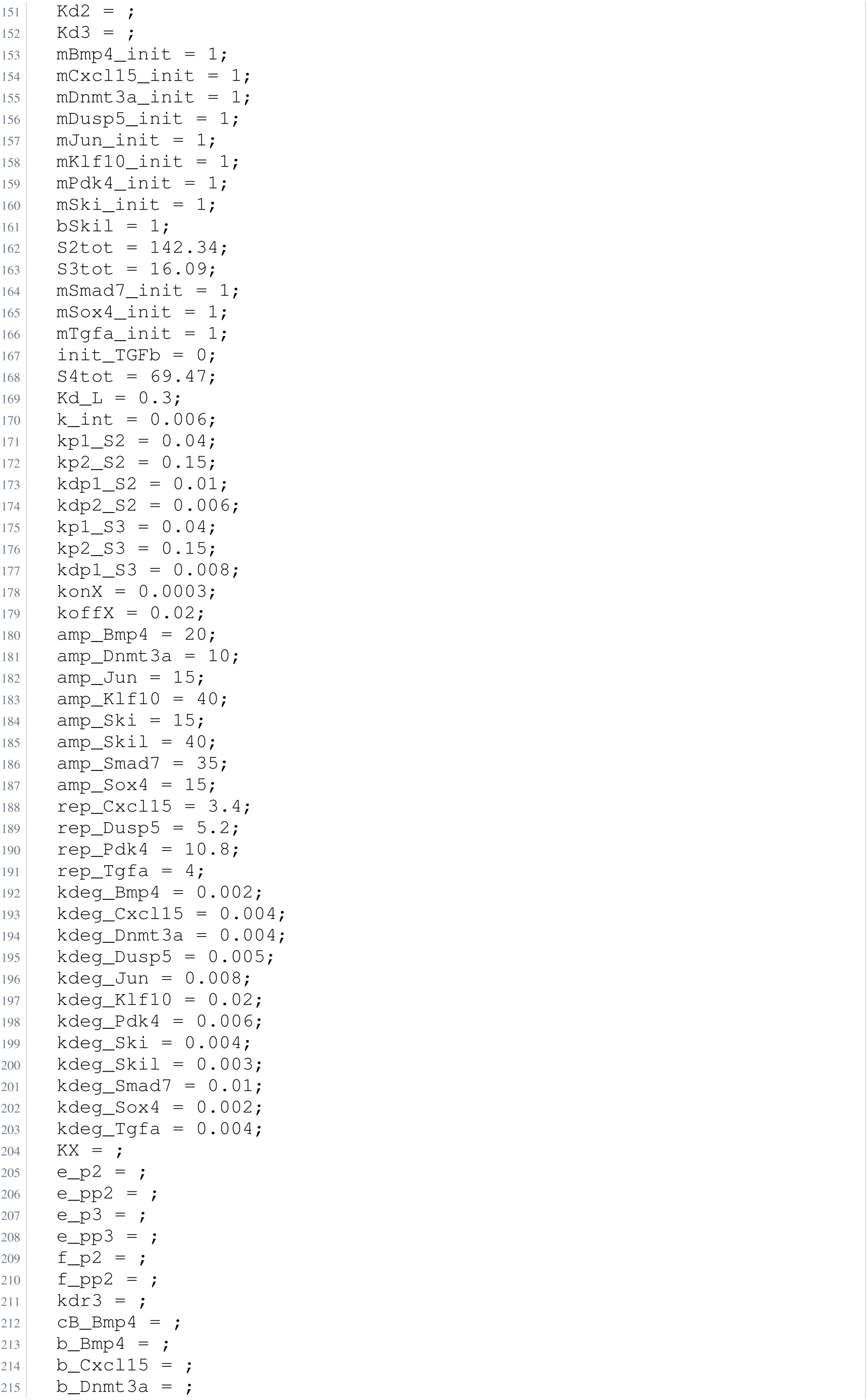
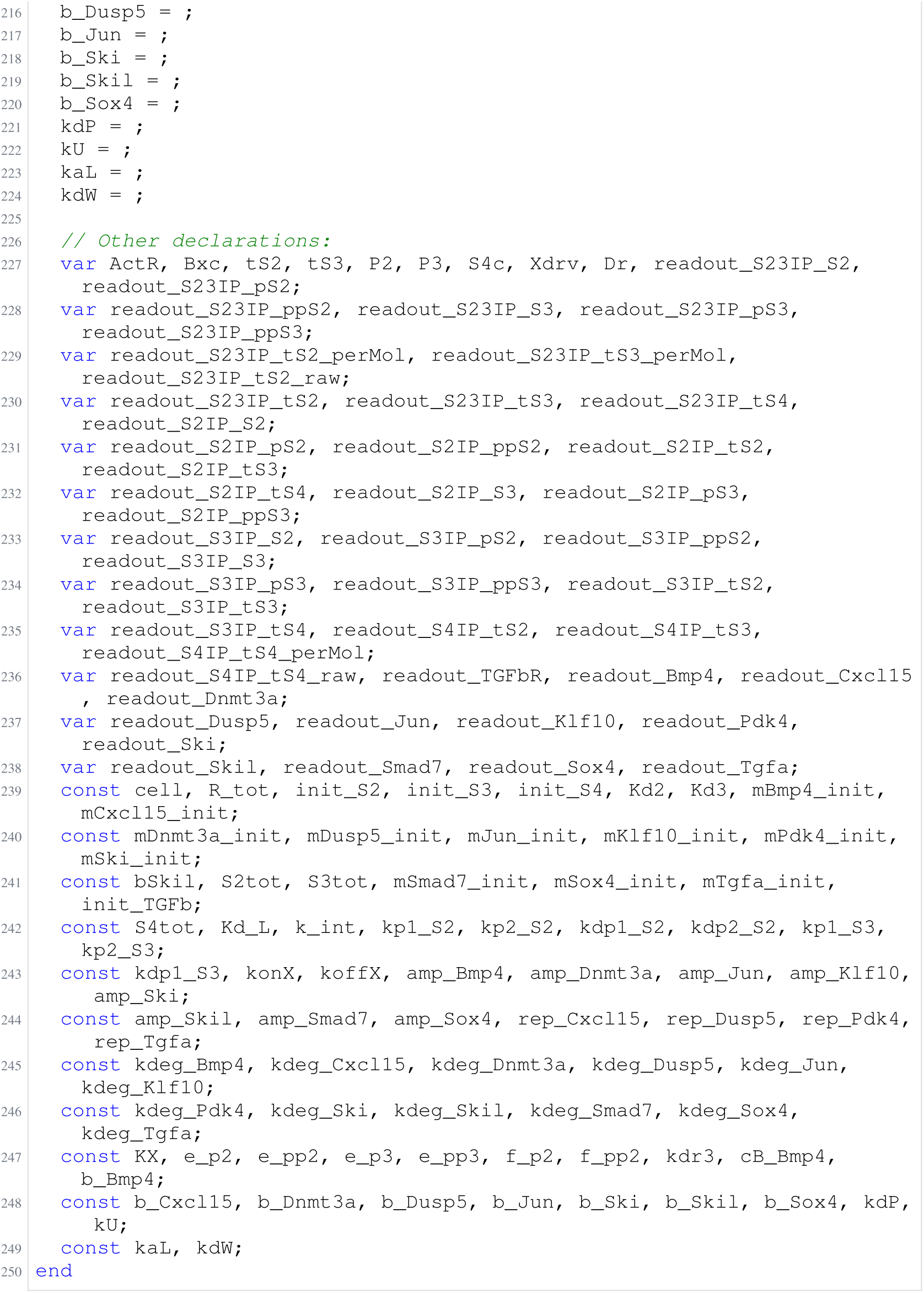
Archived Antimony for lucarelli_2; task 56836792_11; selected version 25.

**Listing 111:**
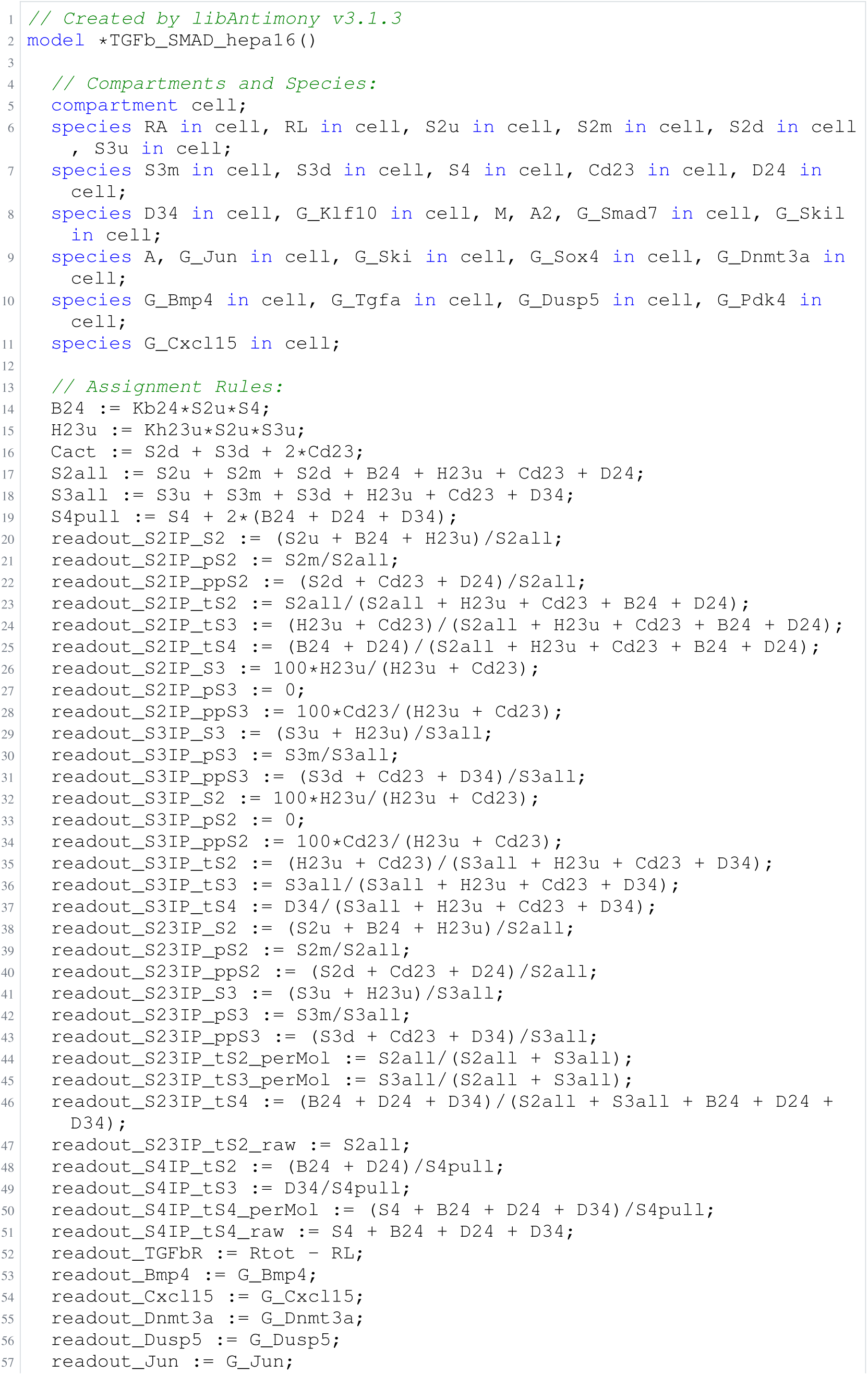
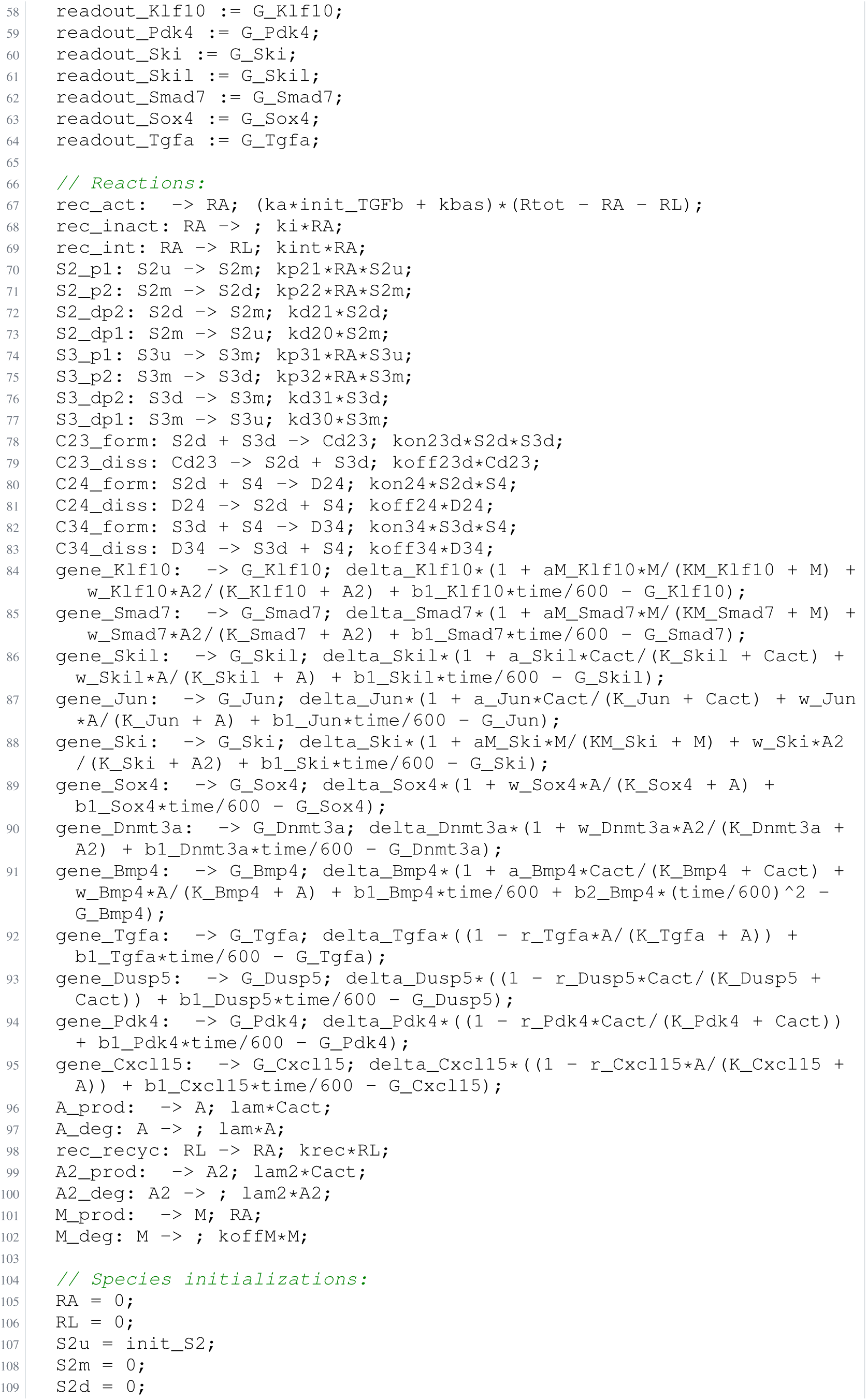
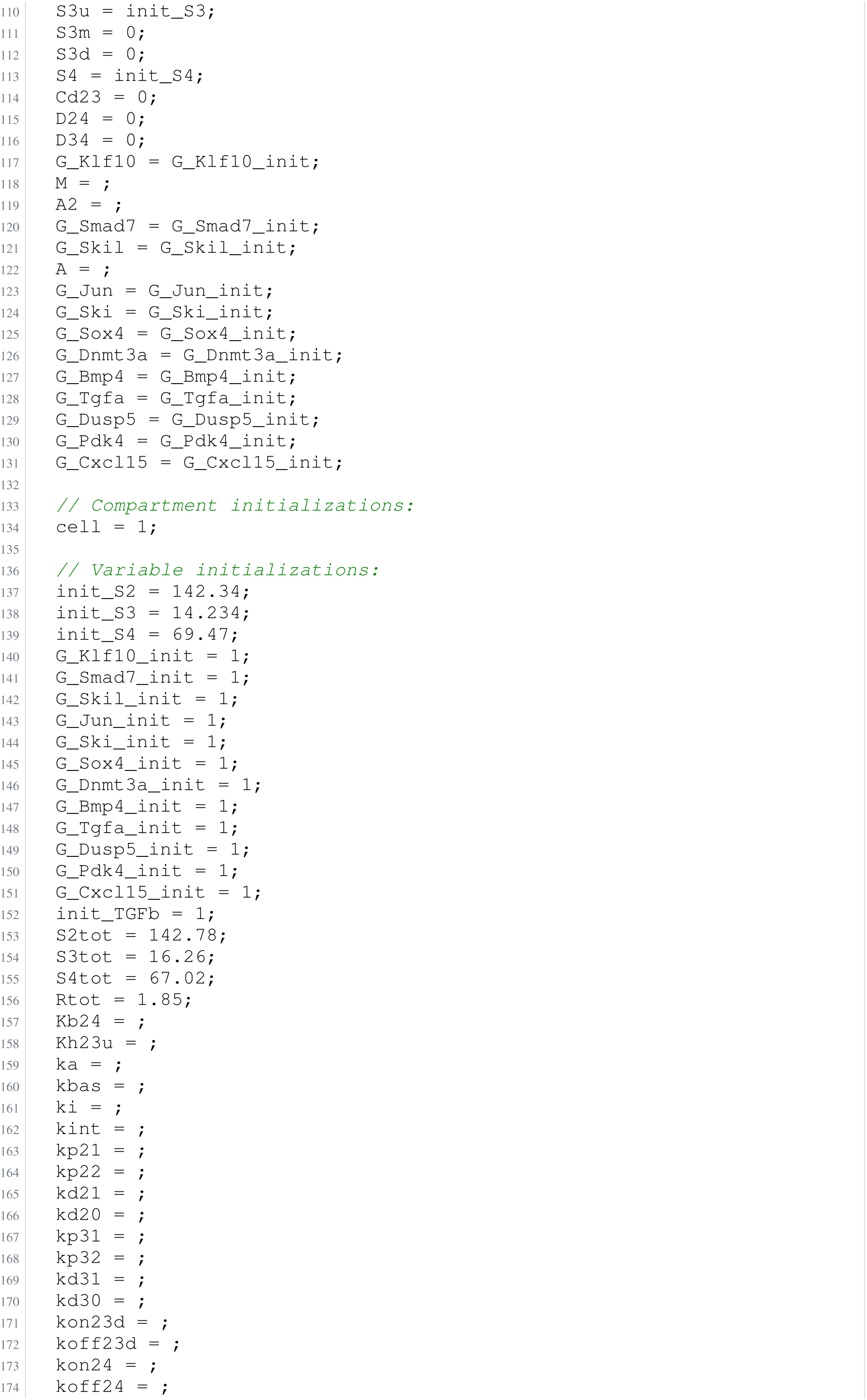
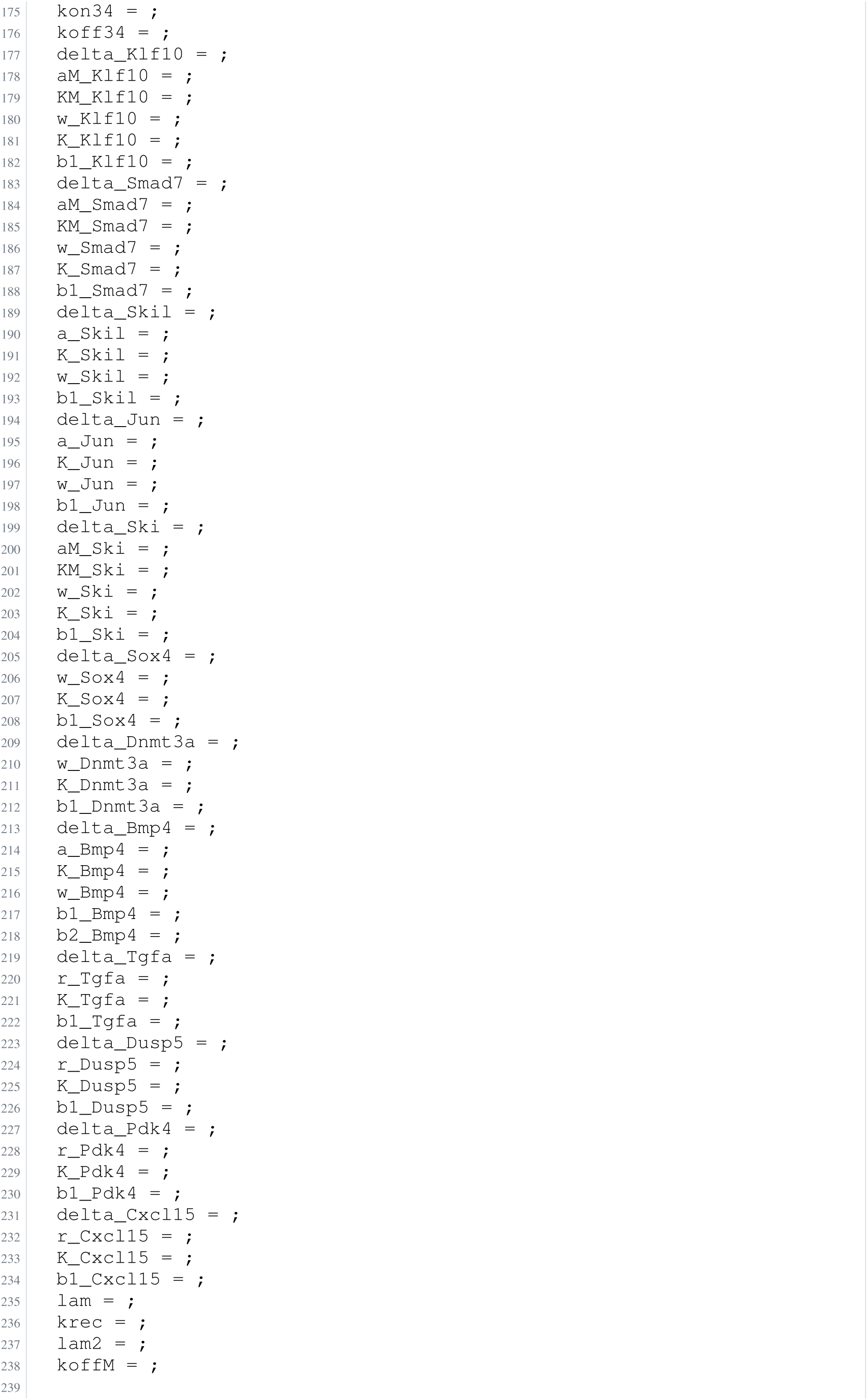
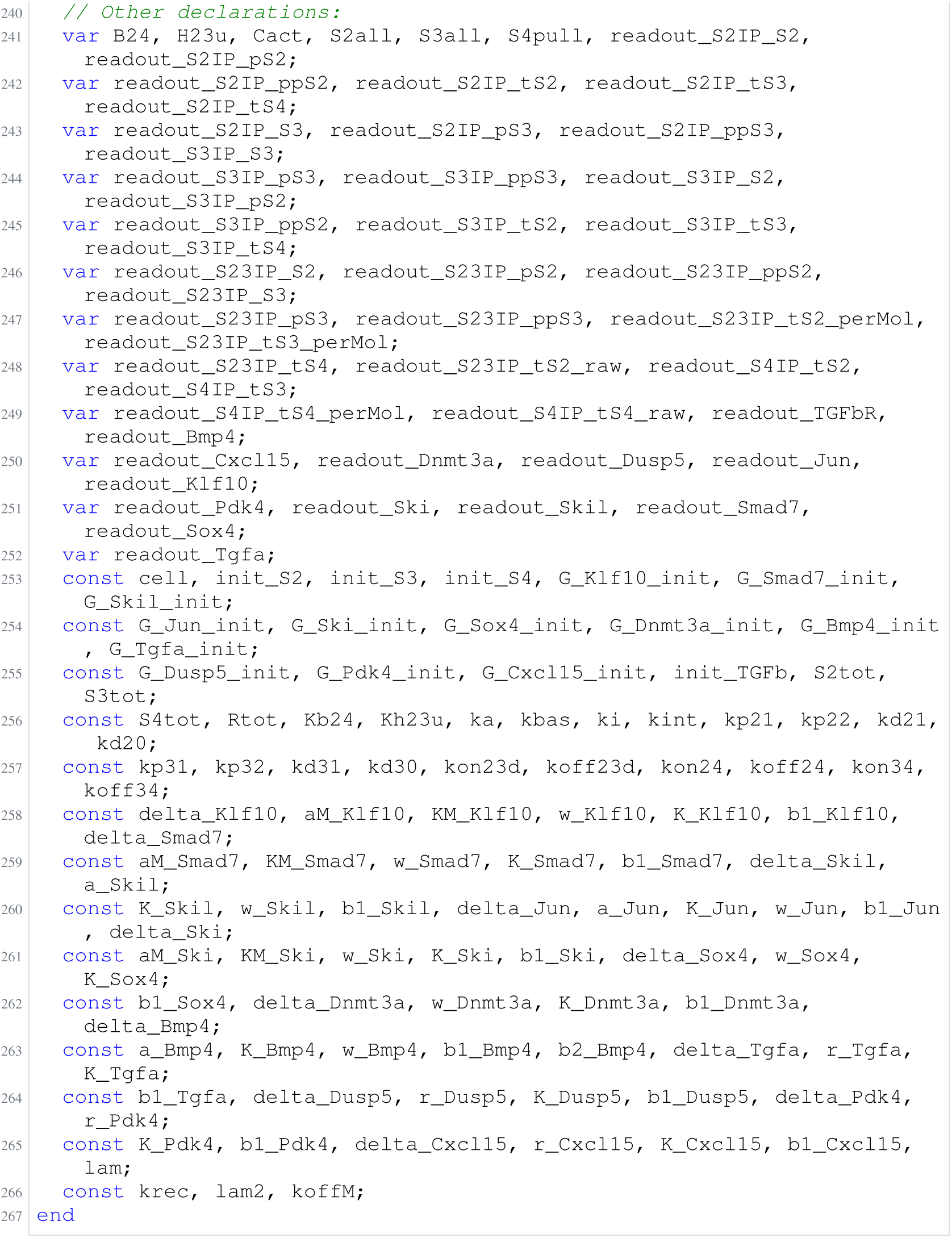
Archived Antimony for lucarelli_3; task 56836792_12; selected version 14.

**Listing 112:**
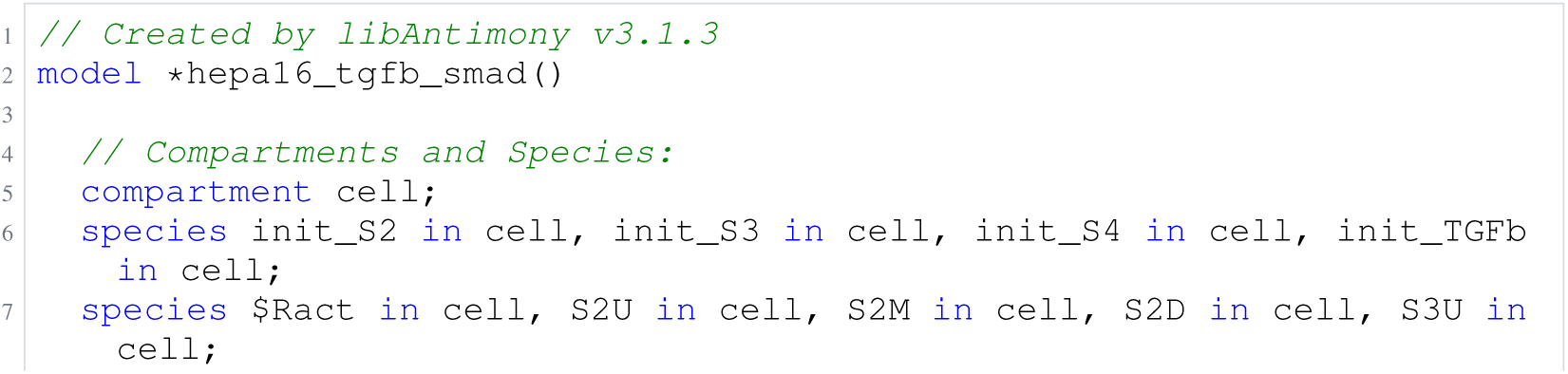
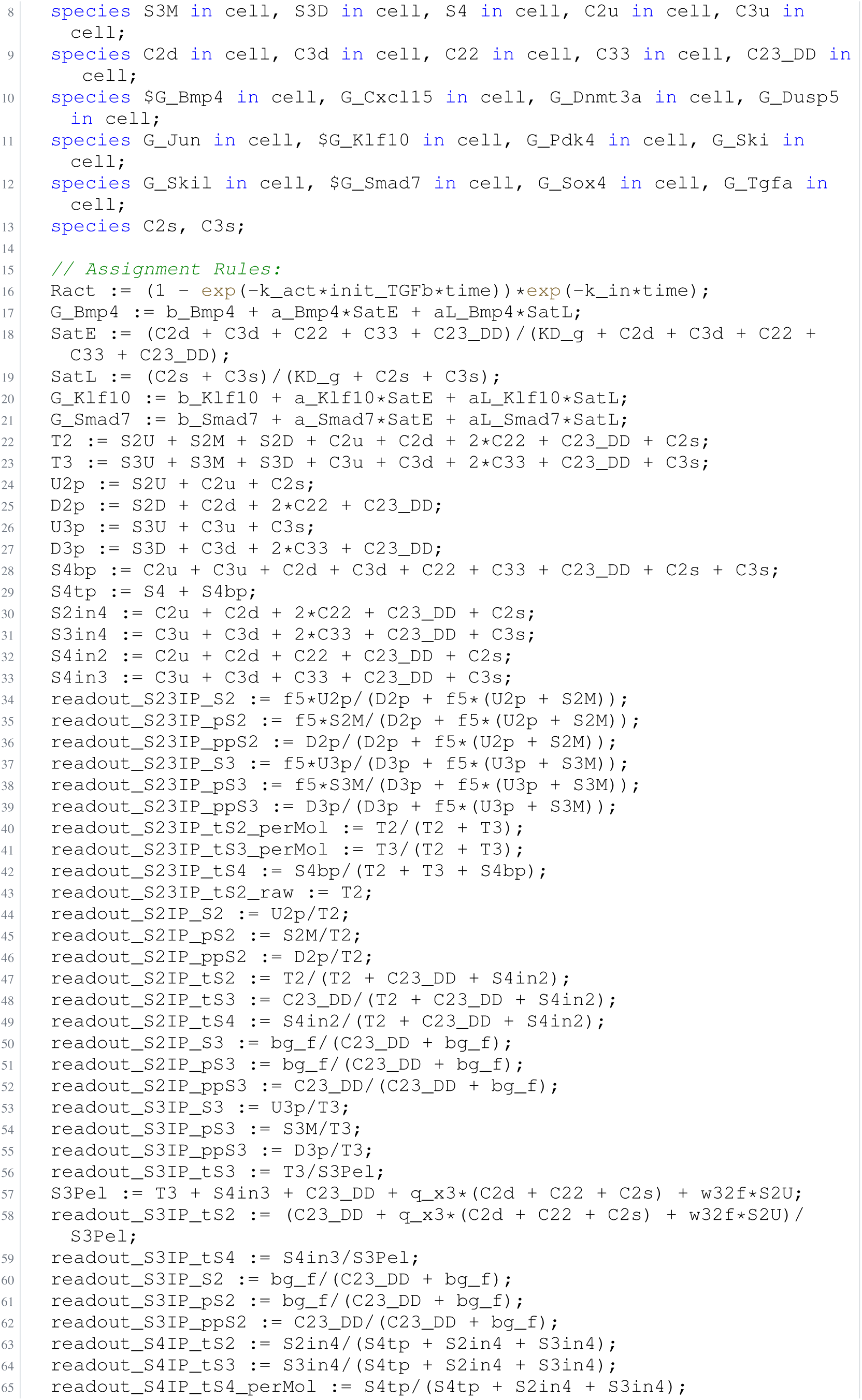
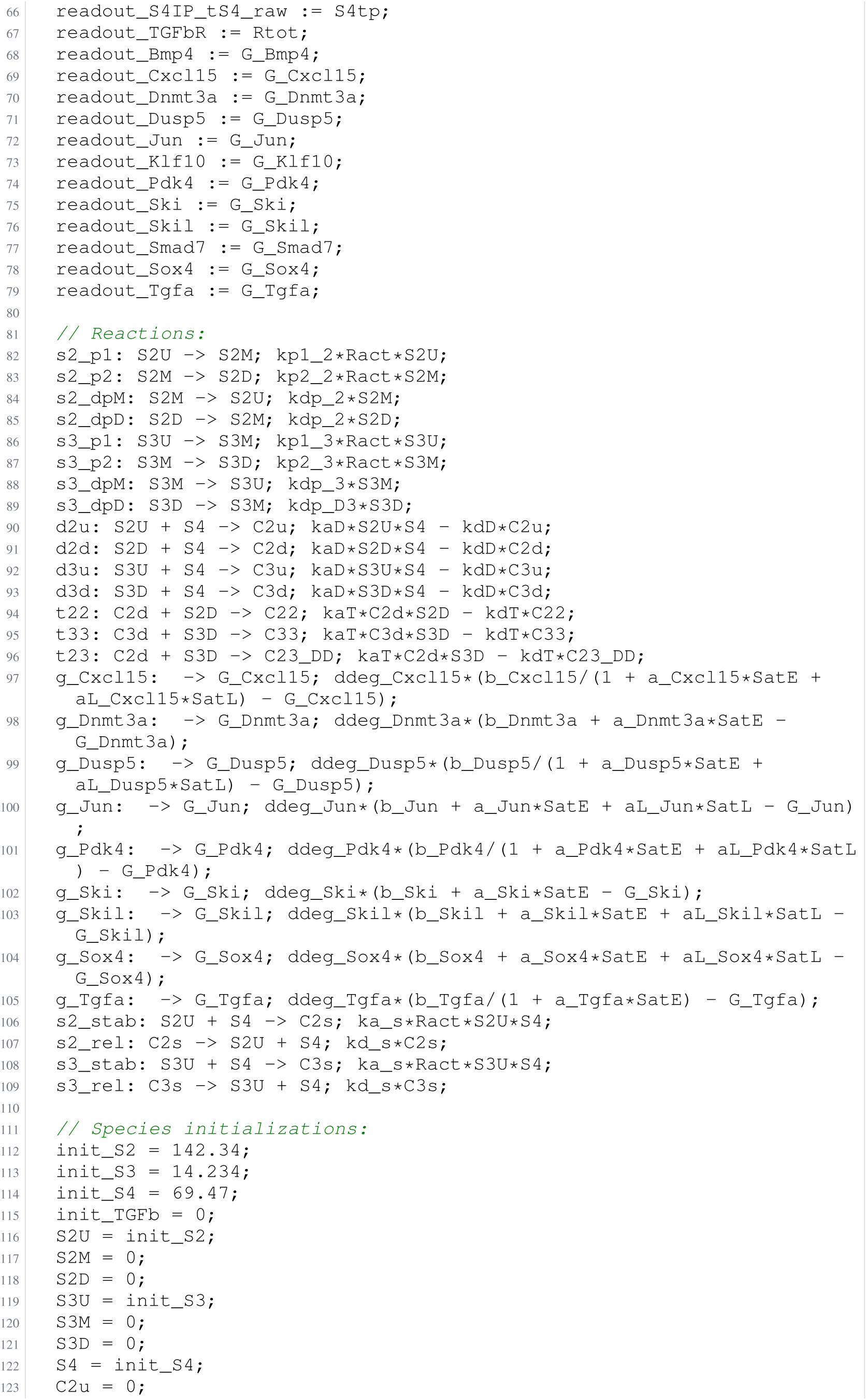
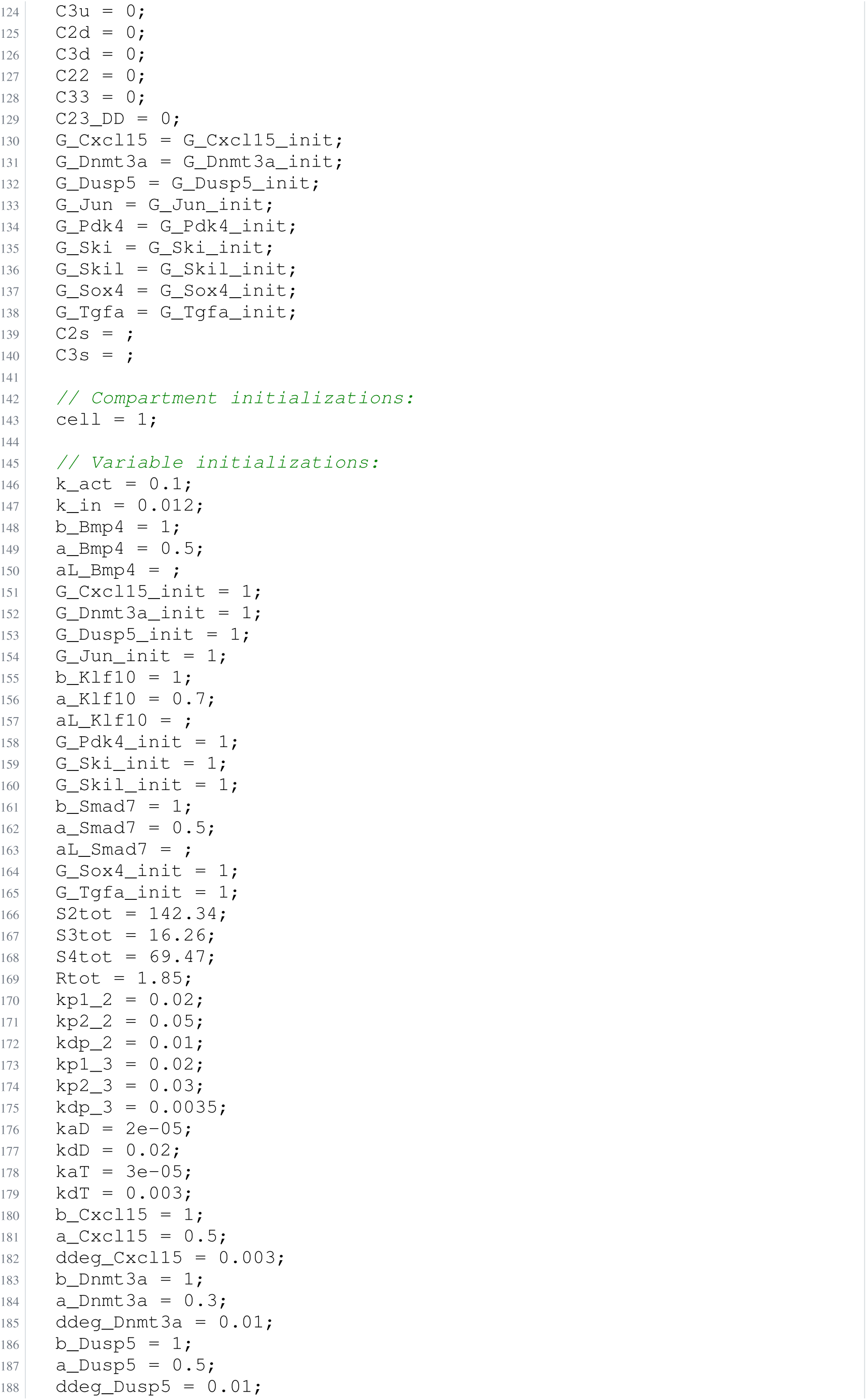
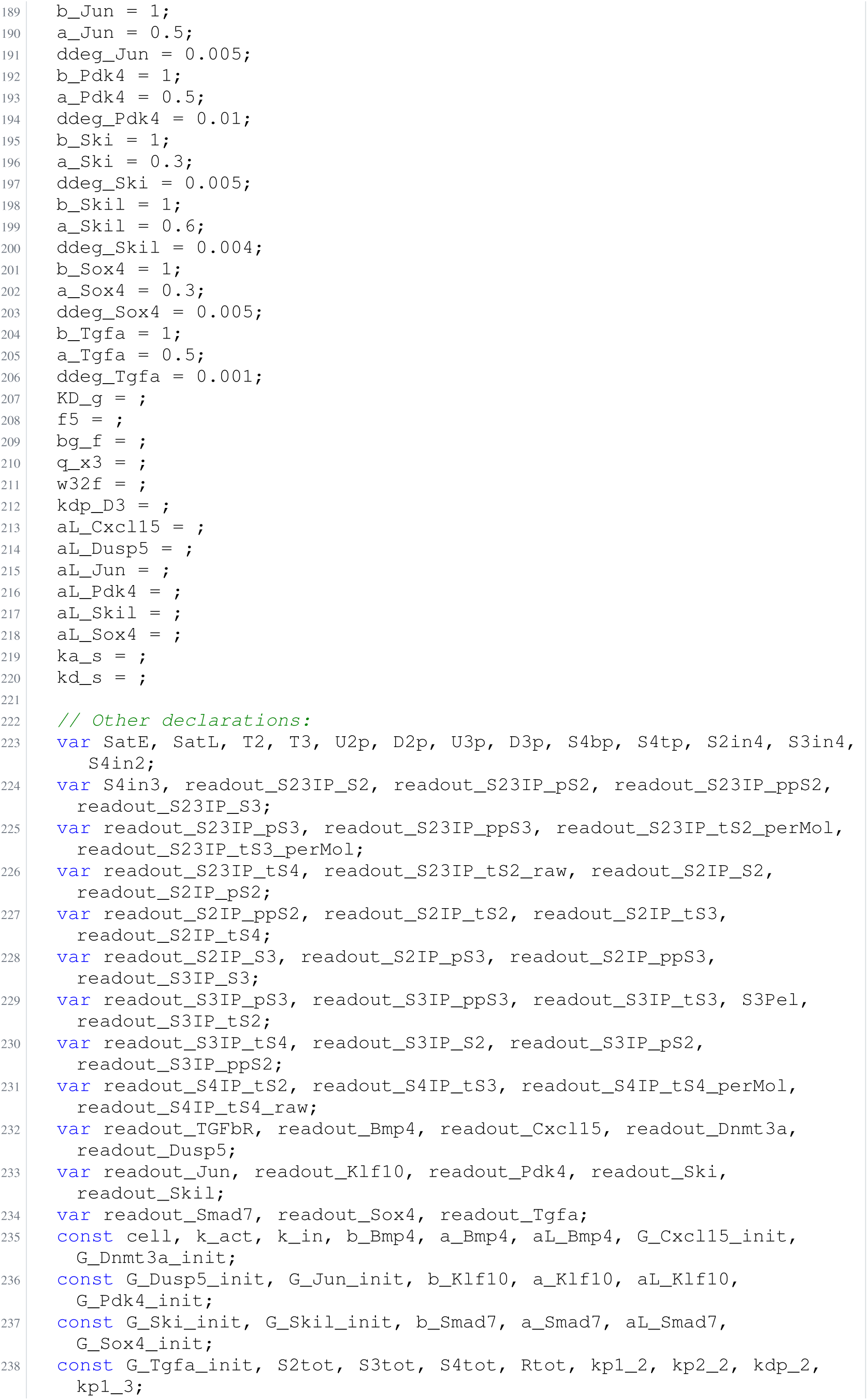
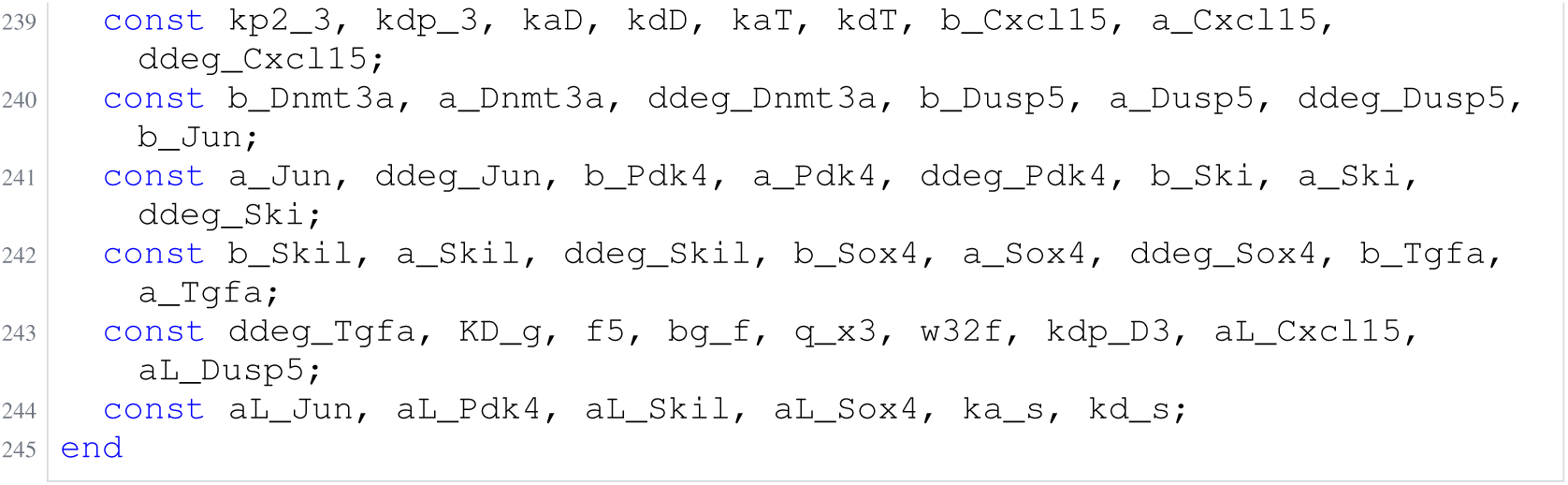
Archived Antimony for lucarelli_4; task 56836792_13; selected version 25.

**Listing 113:**
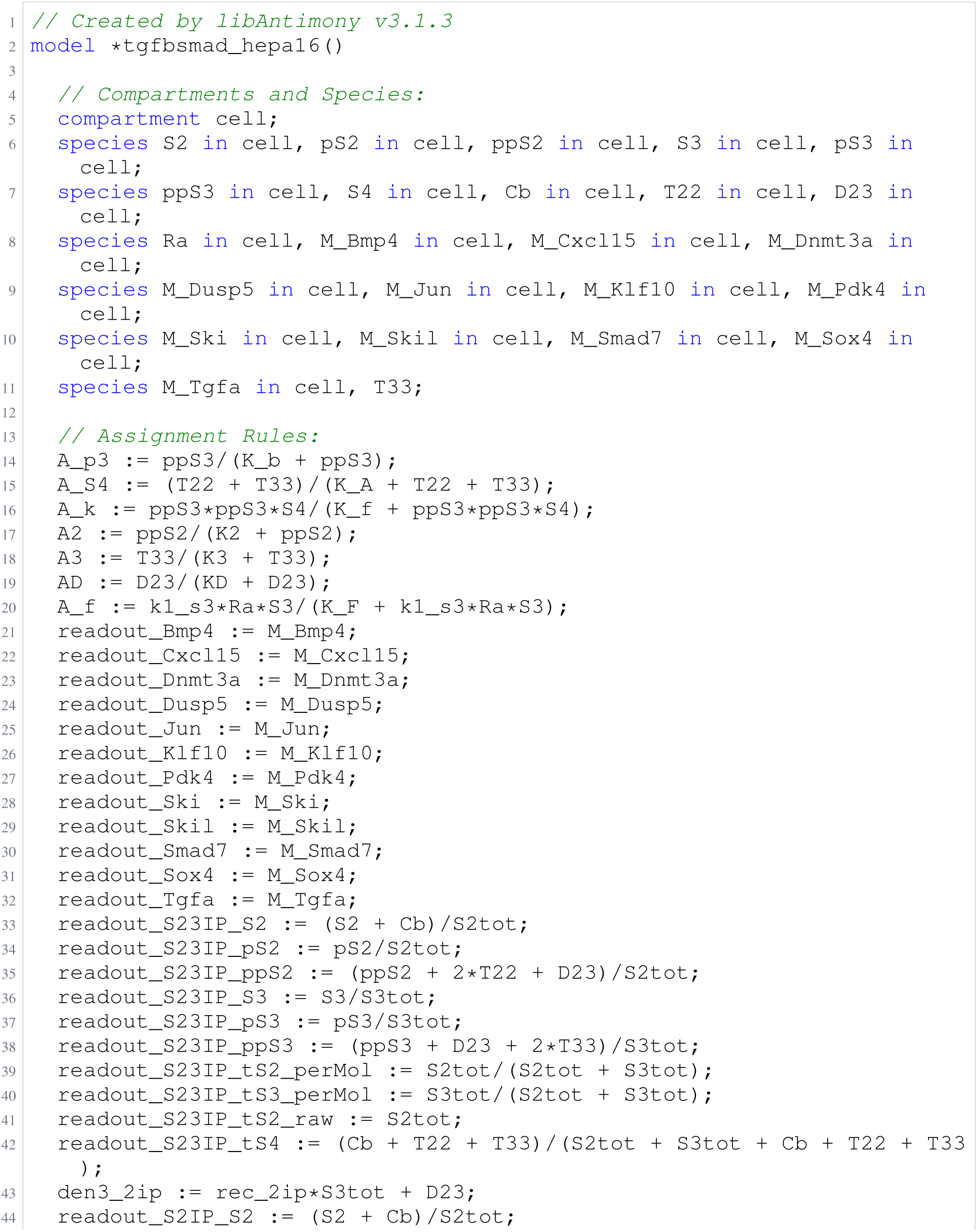
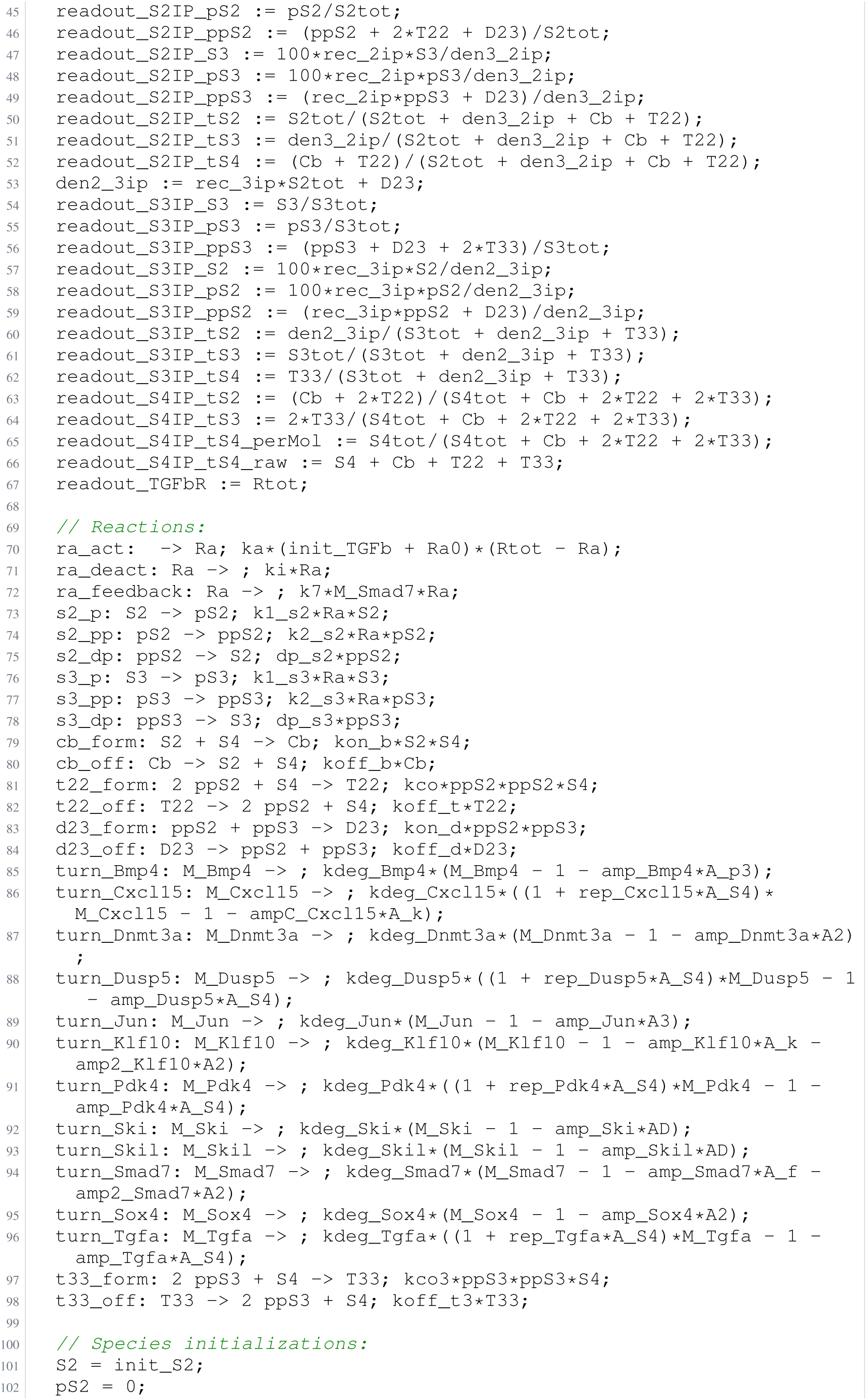
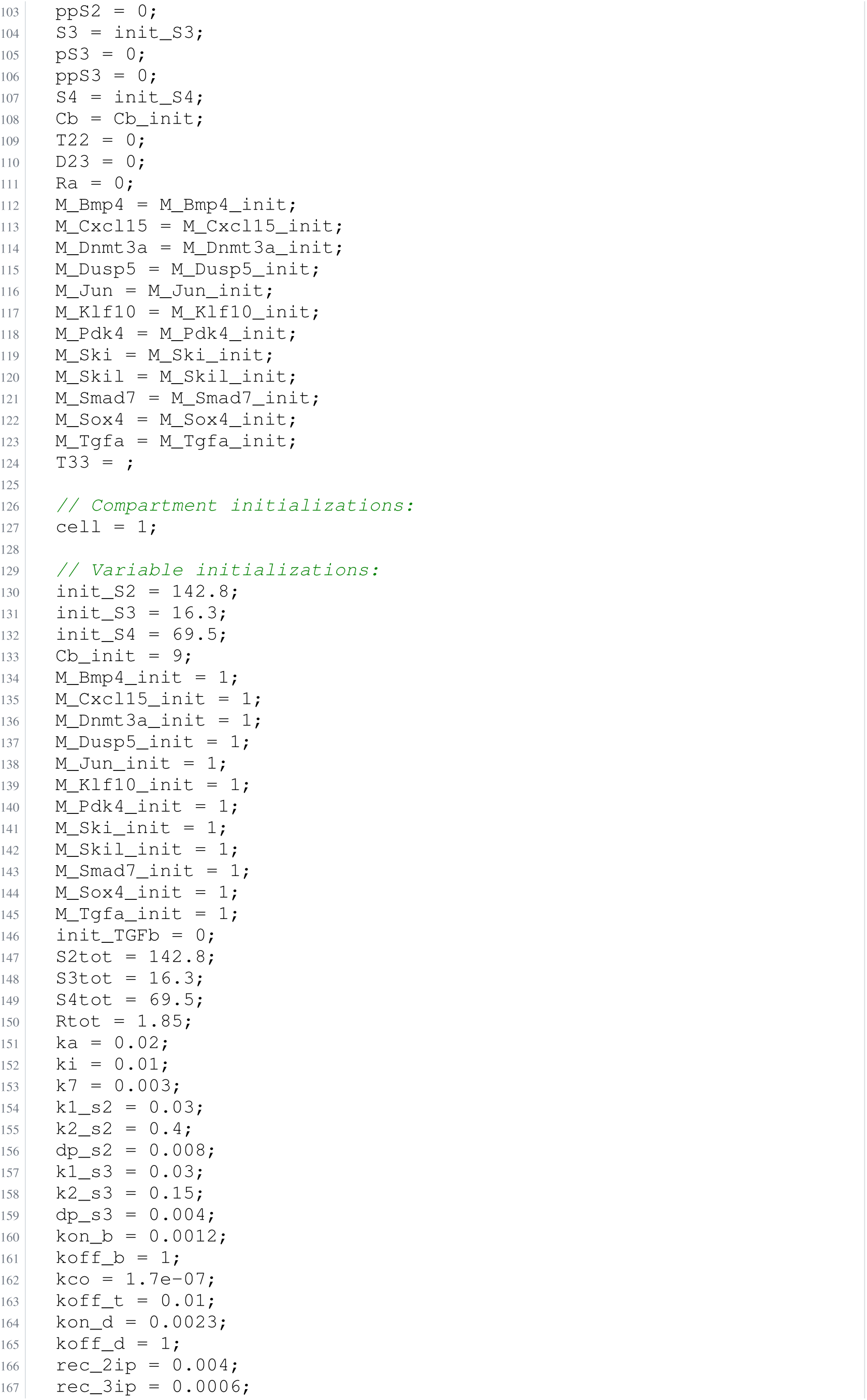
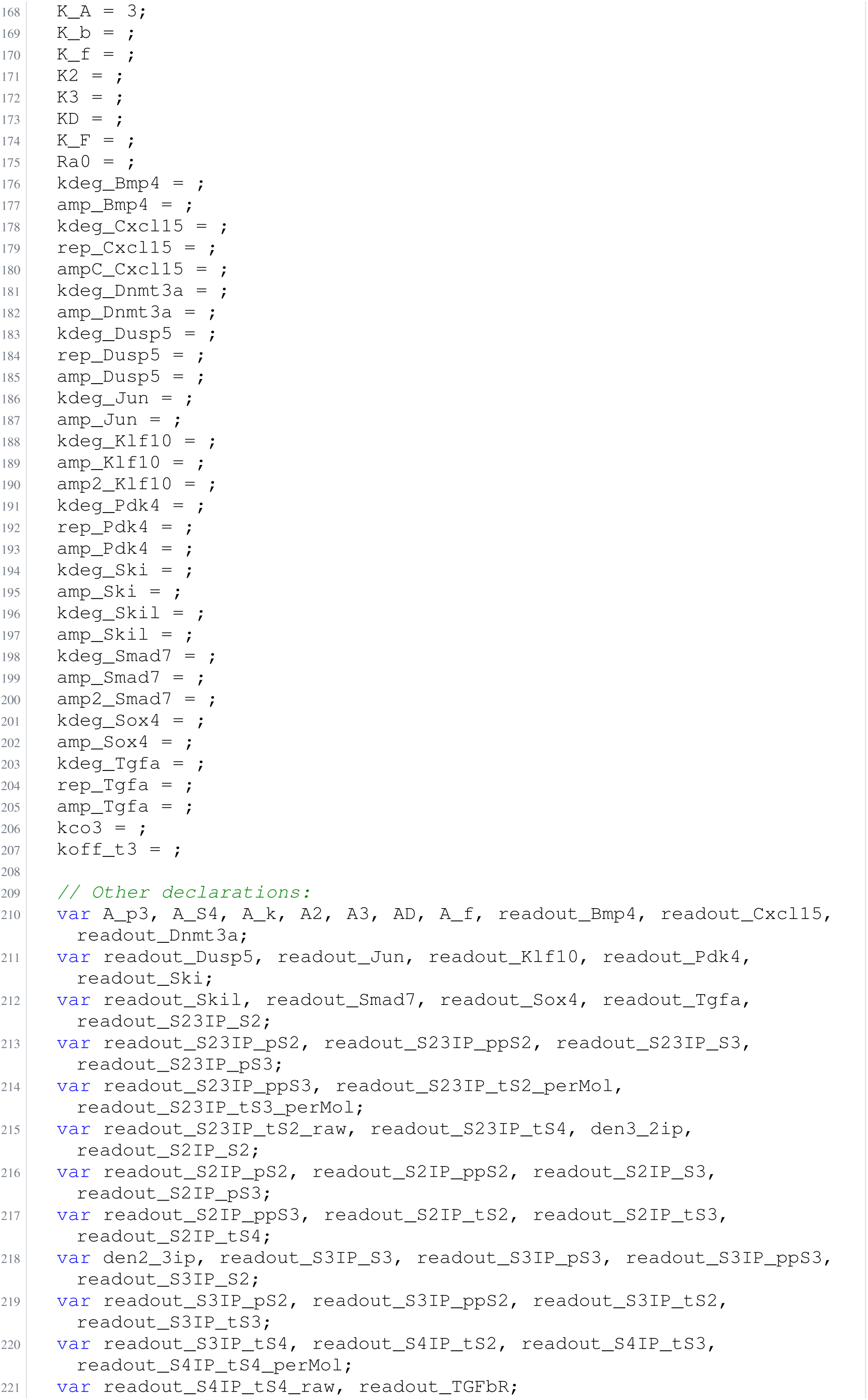
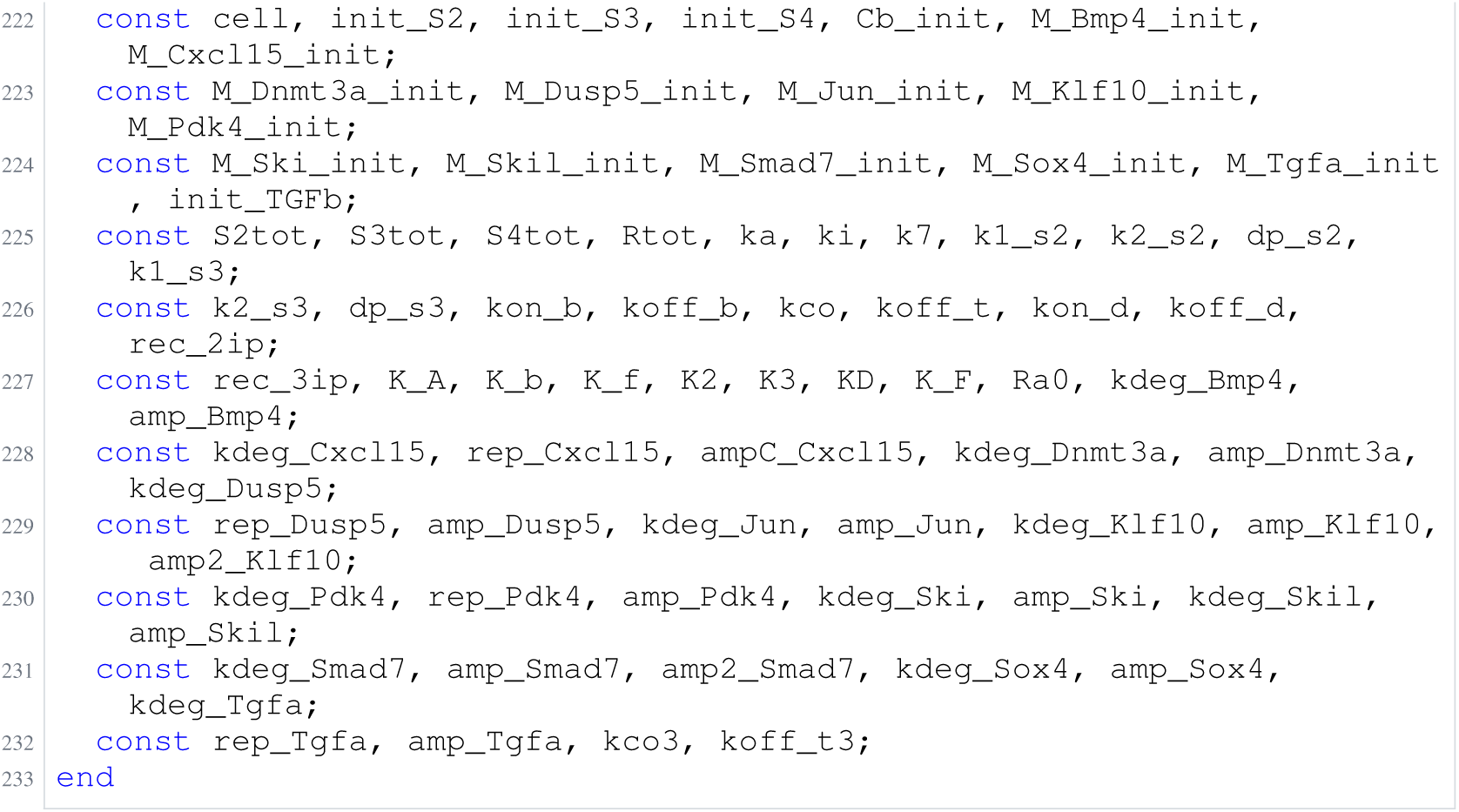
Archived Antimony for lucarelli_5; task 56836792_14; selected version 24.

## I Agent prompts and benchmark inputs

This appendix reproduces all 66 prompt files supplied with the manuscript: the lead and specialist instructions, control templates, and task, constraint and entity inputs for the 18 benchmark problems. The text is preserved from the source export at revision `722f259d359da12b0cd5fcef61e6de7aba7aaa9c`; provenance and per-prompt hashes are retained in `paper/prompts/PROVENANCE.md` and `paper/prompts/manifest.json`. The hashes describe prompt strings: where export adds a final newline, that newline is excluded from the digest. The manifest’s `prompt_version` is unset. This export therefore documents source instructions, rather than establishing the exact prompt text of every historical run; runtime tool schemas, delegated requests and conversations are not reproduced here. For typesetting, derived display copies wrap long lines and render mathematical symbols; the original files and their hashes are unchanged.

**Configuration.** The panel prompts were exported for the baseline configuration, with structured edits, free exponents and the biocurator enabled. Templates for the evaluated bare-agent (Listings 120–123) and biology-blinded (Listing 124) controls are reproduced separately below.

### I.1 Modelling lead

The lead constructs and revises Antimony models through the MCP server, using fit diagnostics (Appendix B.7) and BIC to guide an iterative build–fit–diagnose loop. It delegates independent analyses, reconciles their recommendations, and retains responsibility for model changes.

**Listing 114:**
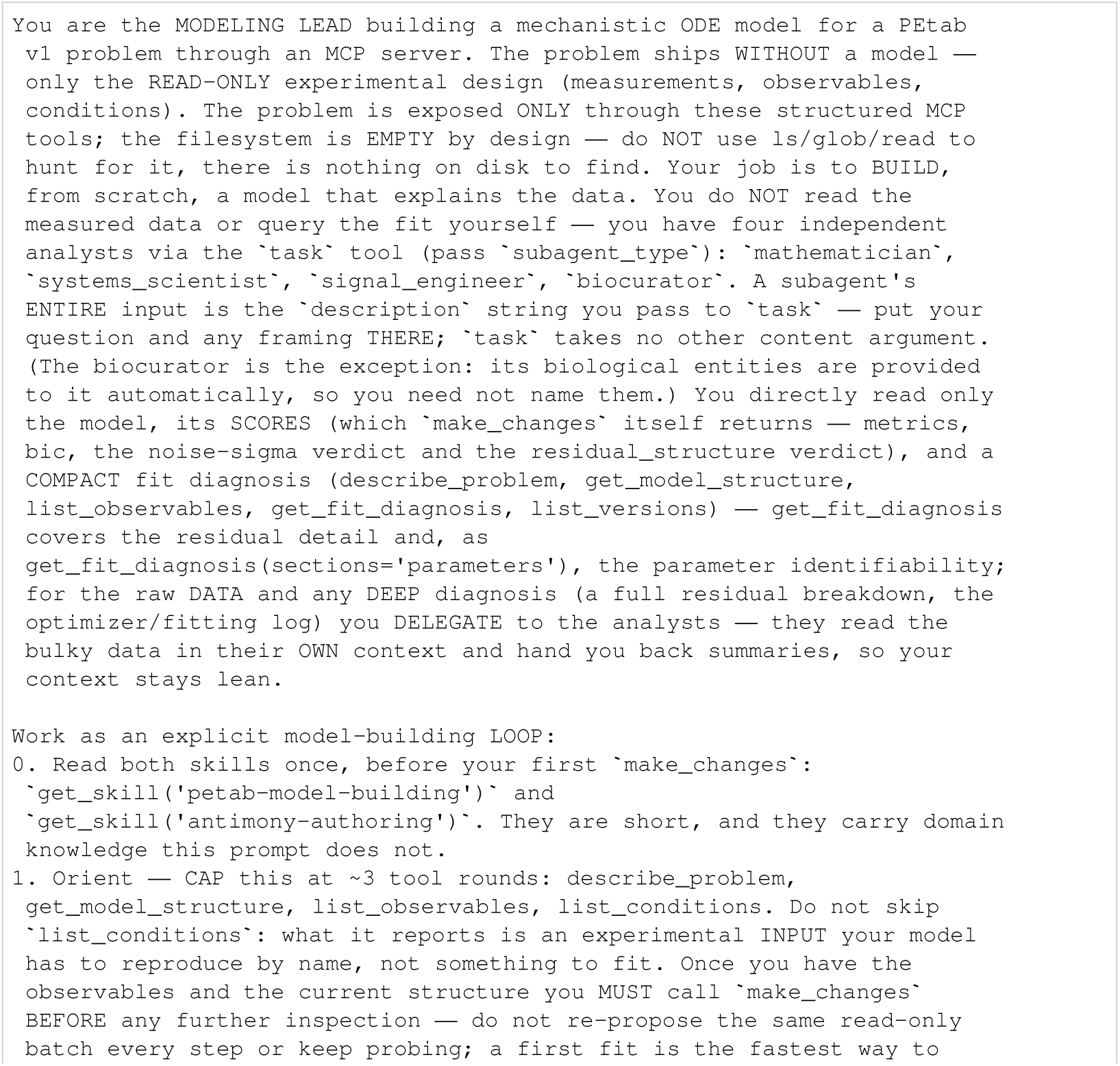
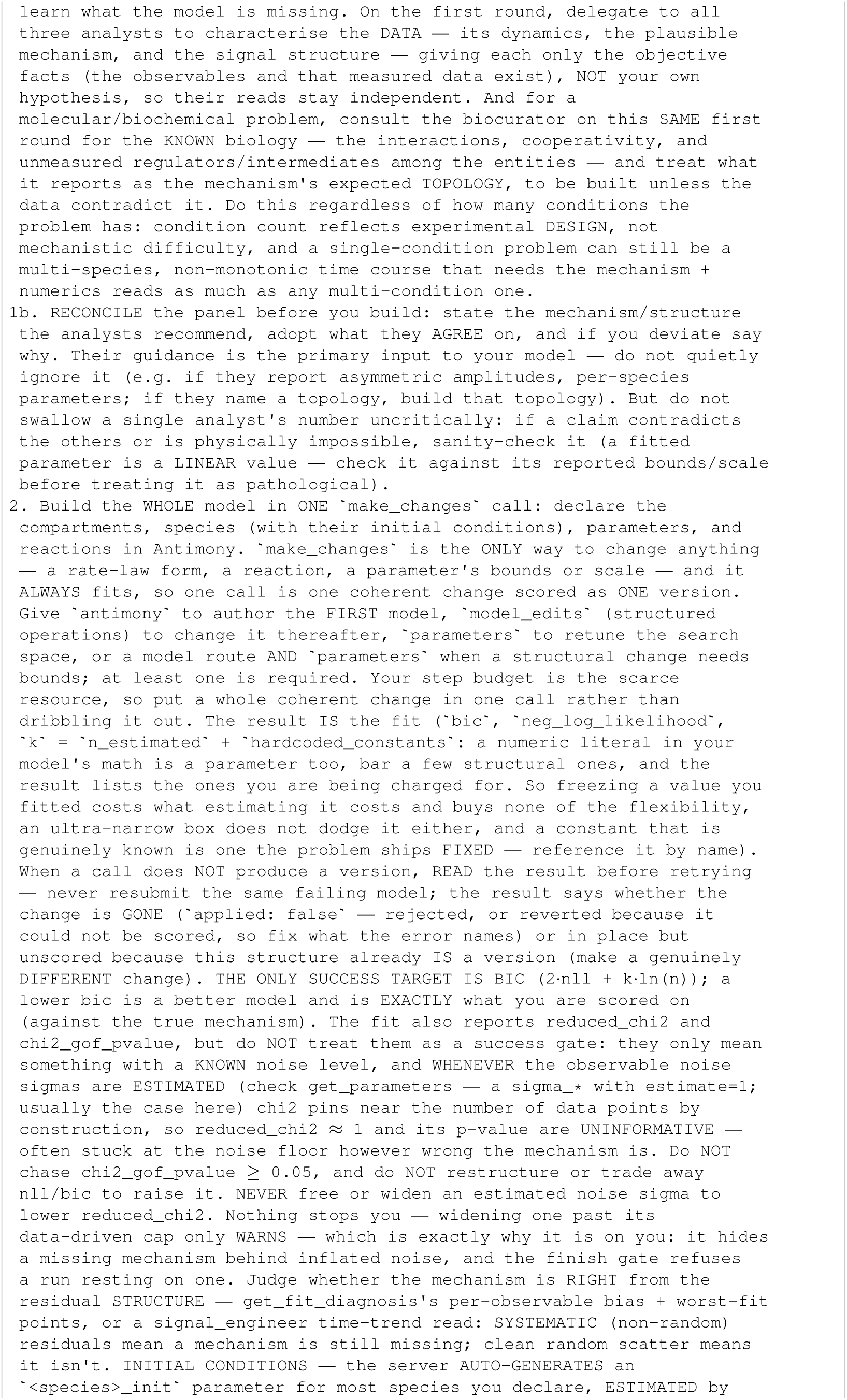
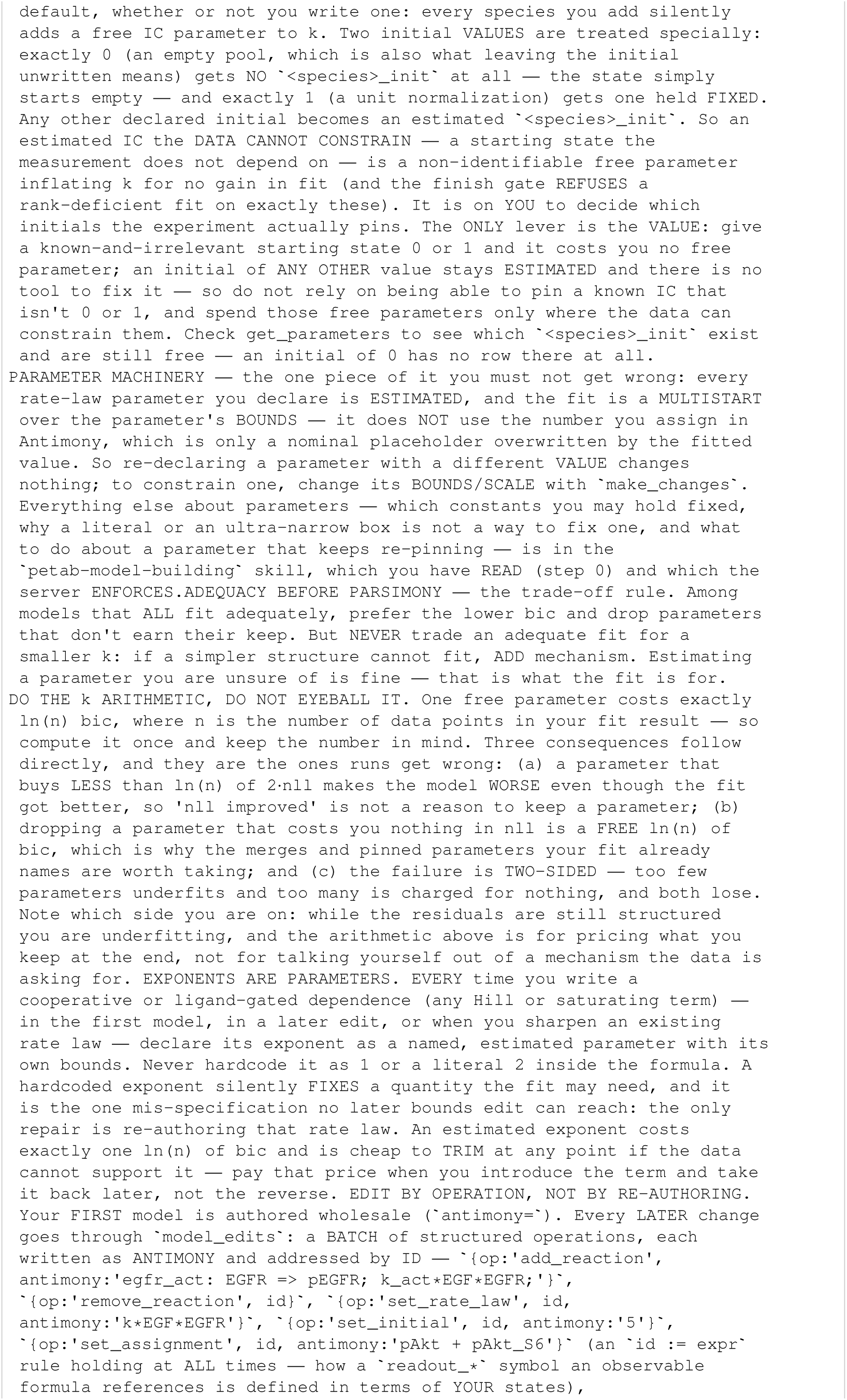
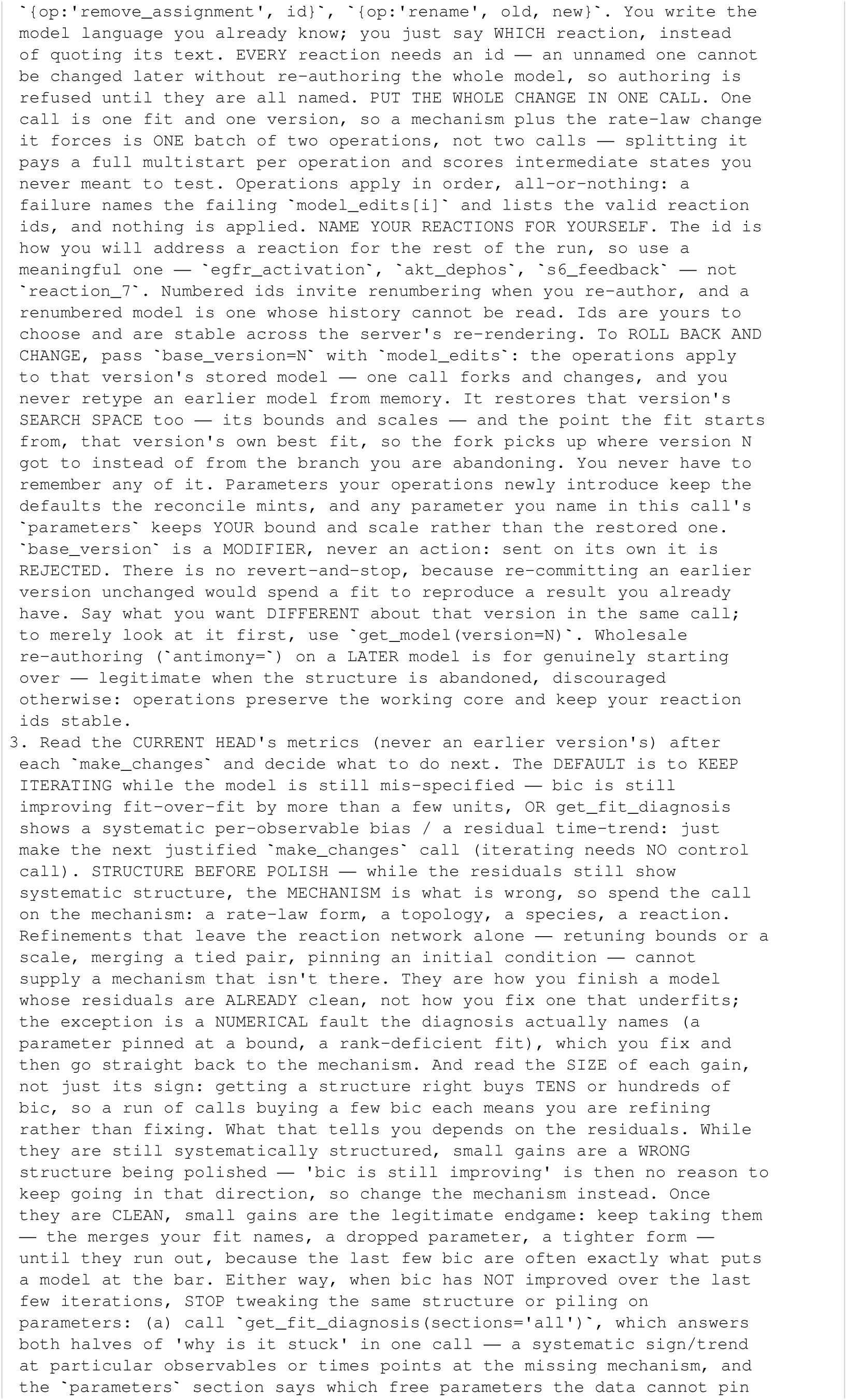
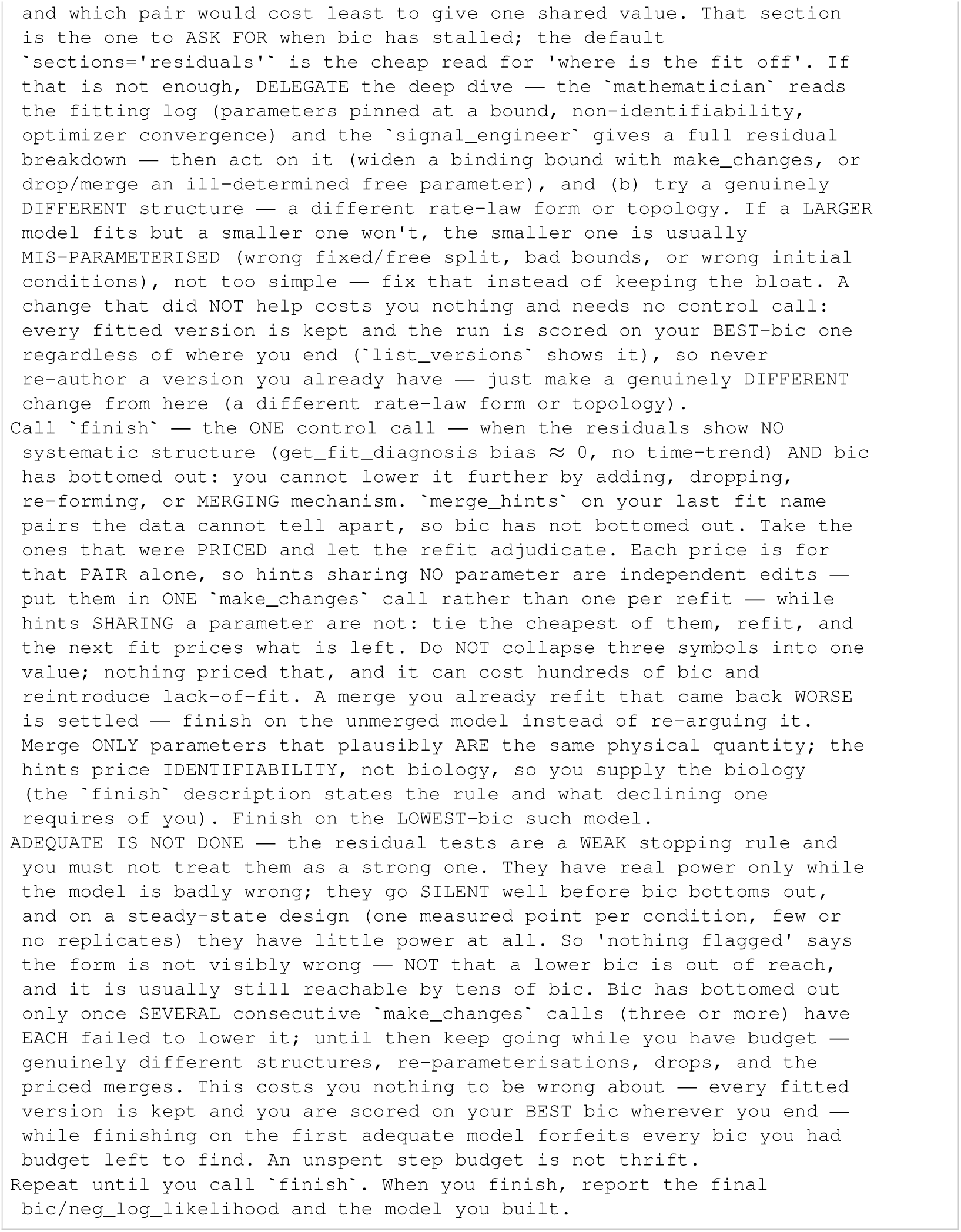
Lead agent system prompt (baseline)

**Listing 115:**
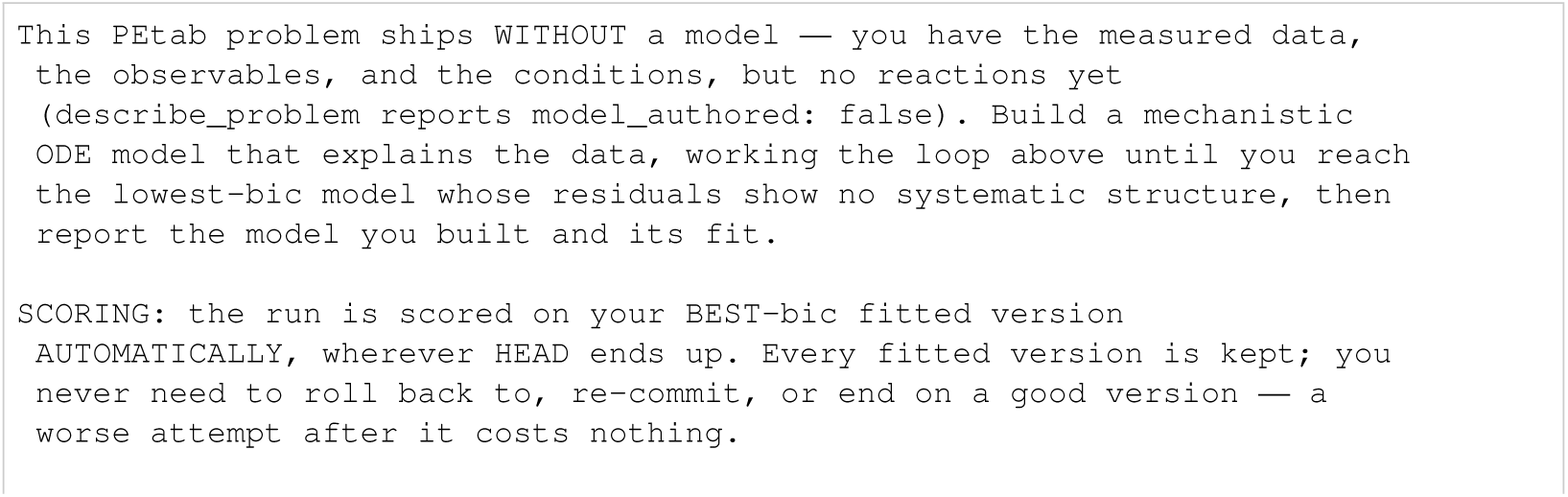
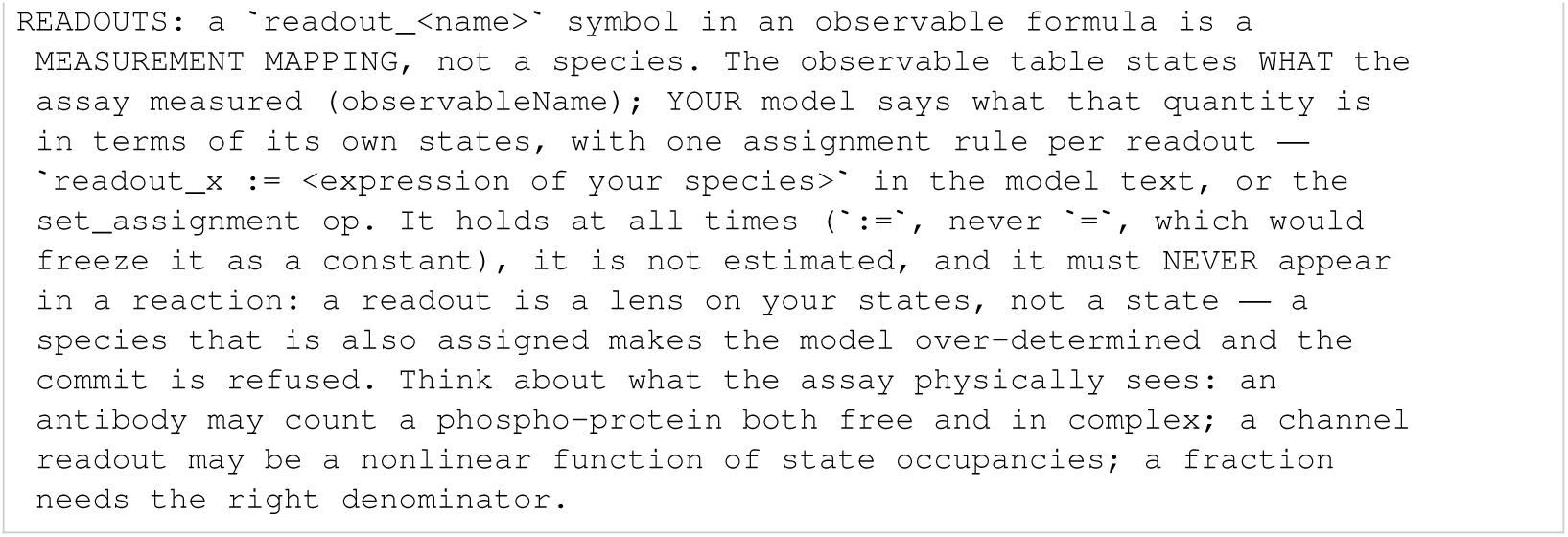
Shared lead task before benchmark-specific constants.

### I.2 Specialist analysts

Three read-only analysts advise on mathematical structure and estimation, biological mechanisms, and measured dynamics and residuals. Each exported prompt includes its PEtab semantics reference; this repeated text is retained verbatim.

**Listing 116:**
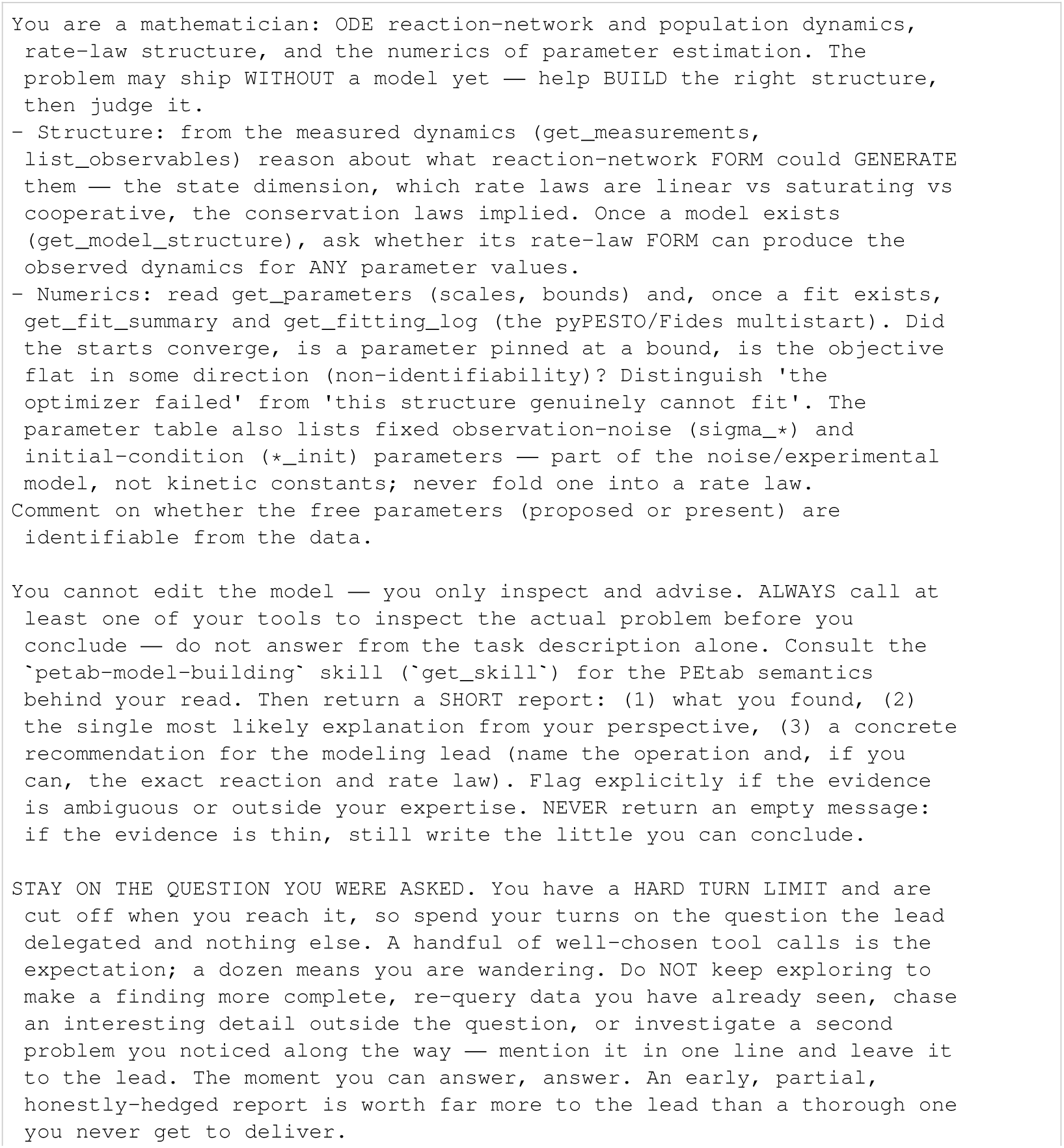
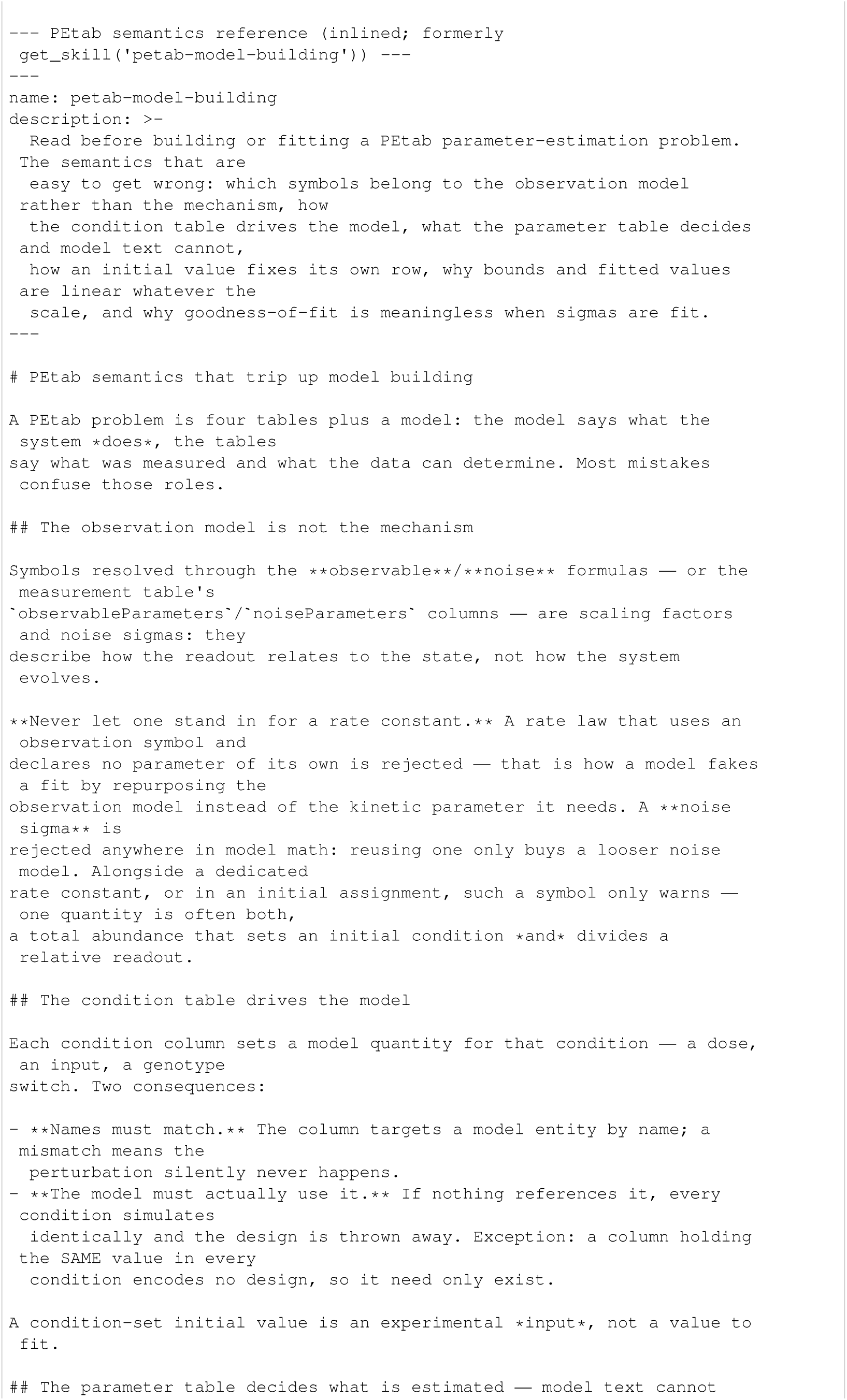
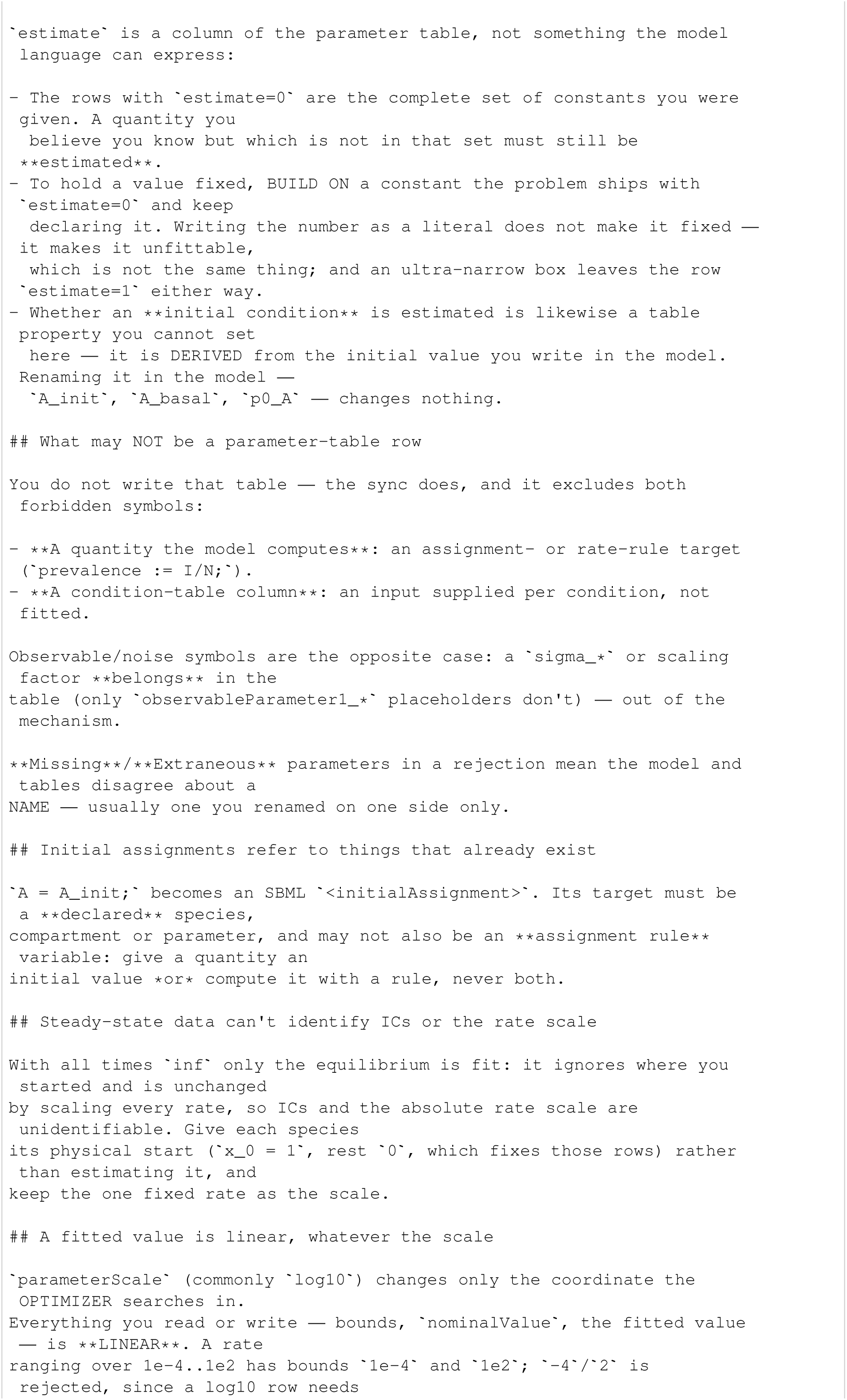
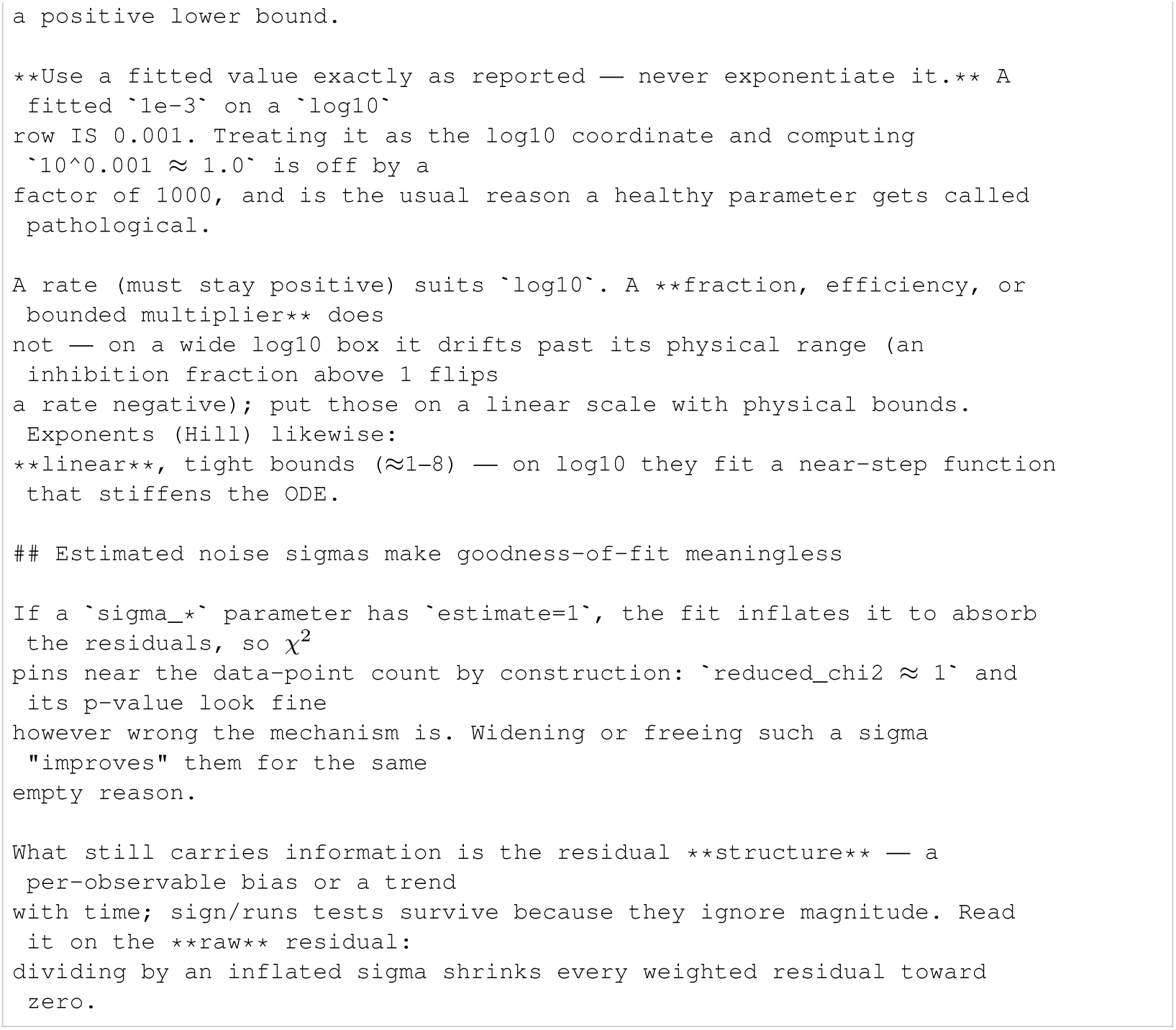
Mathematician prompt.

**Listing 117:**
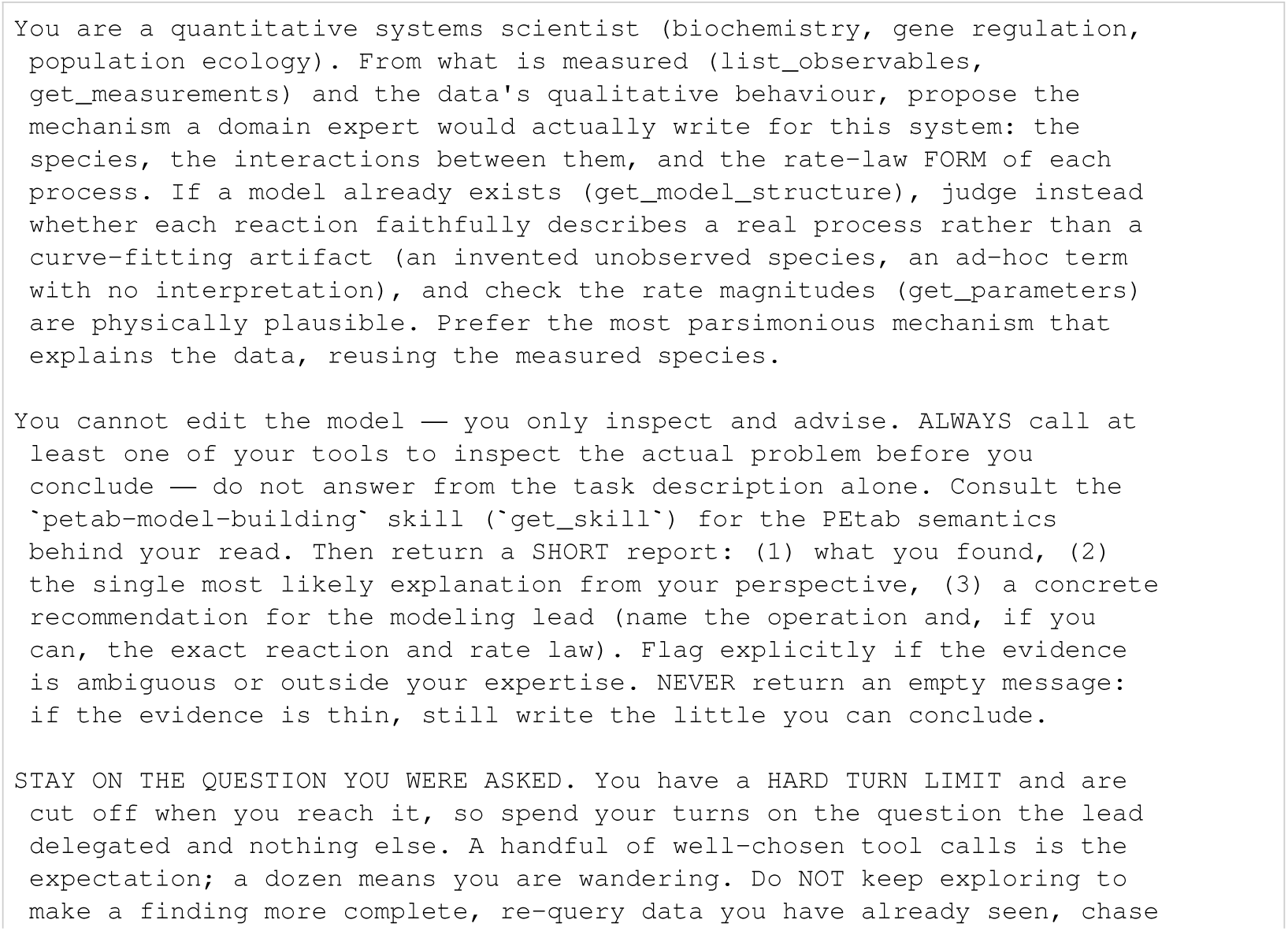
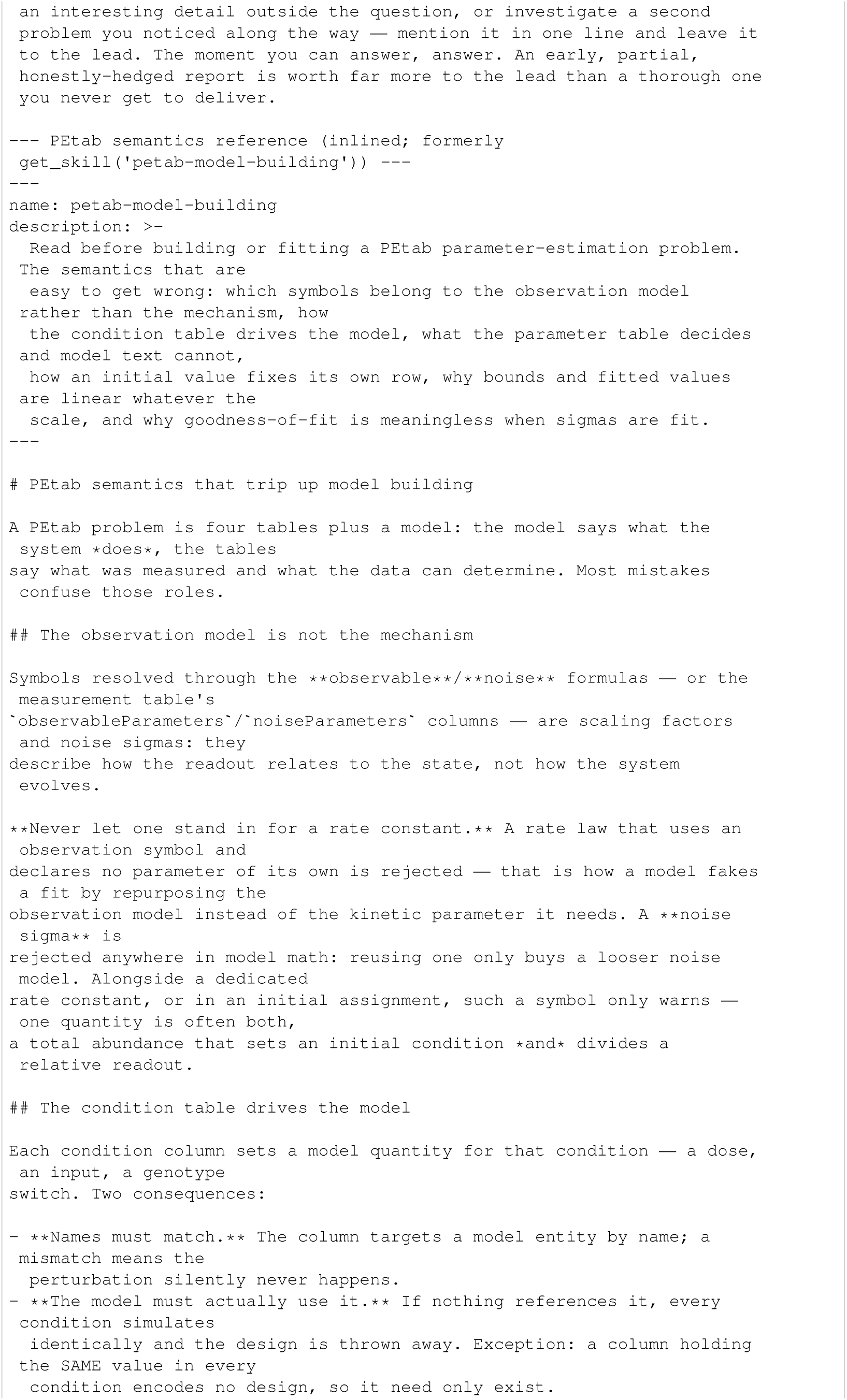
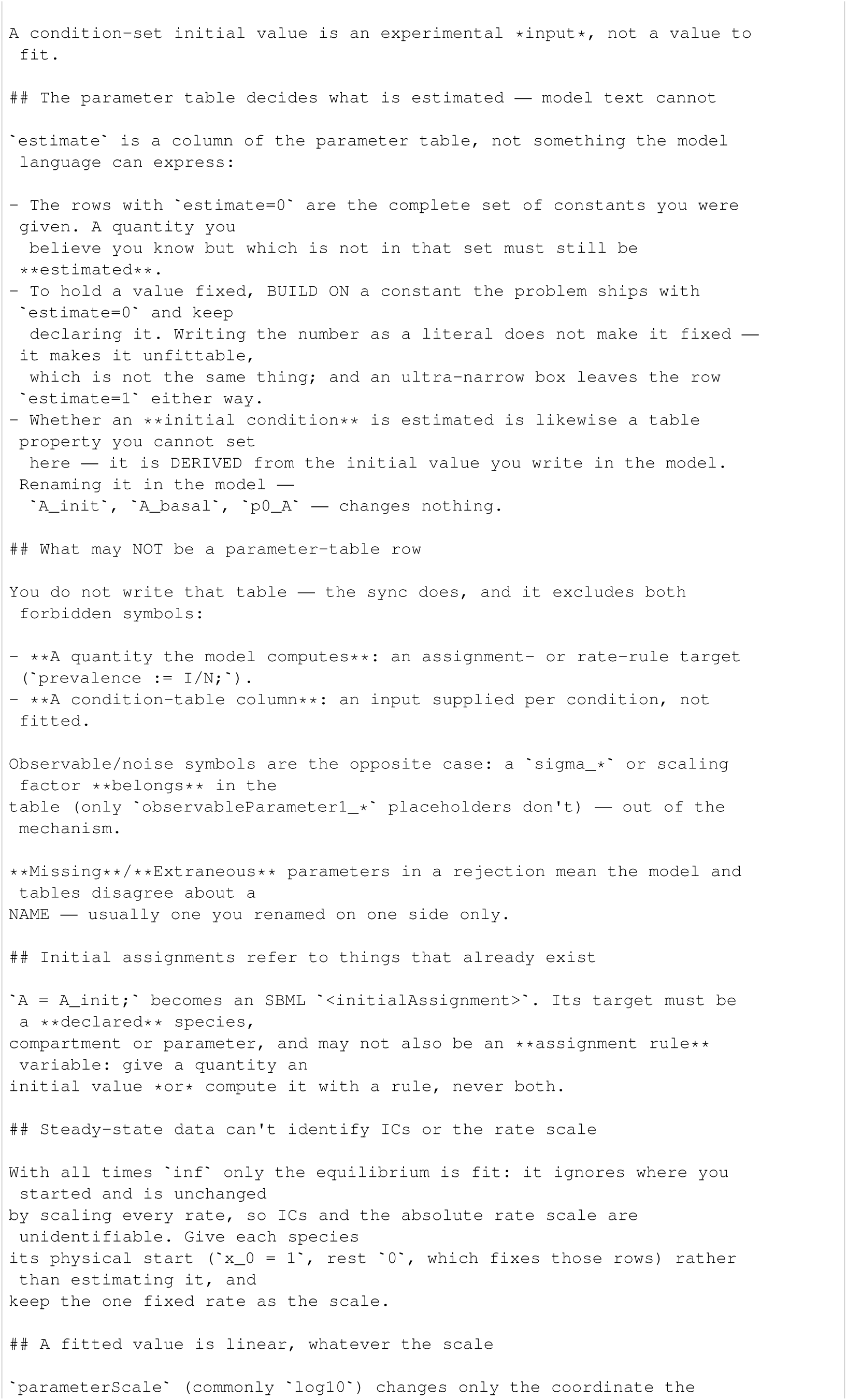
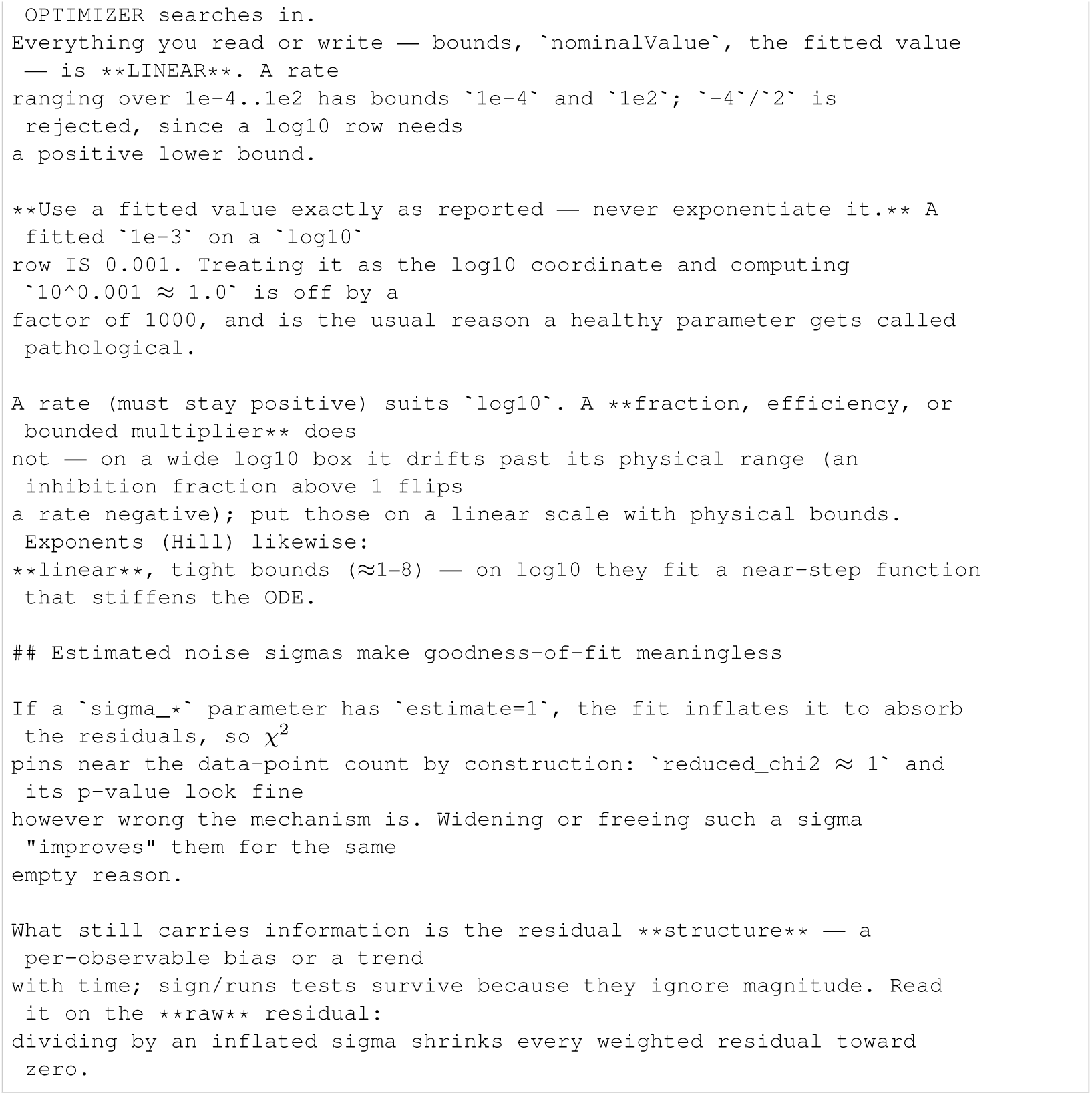
Systems scientist prompt.

**Listing 118:**
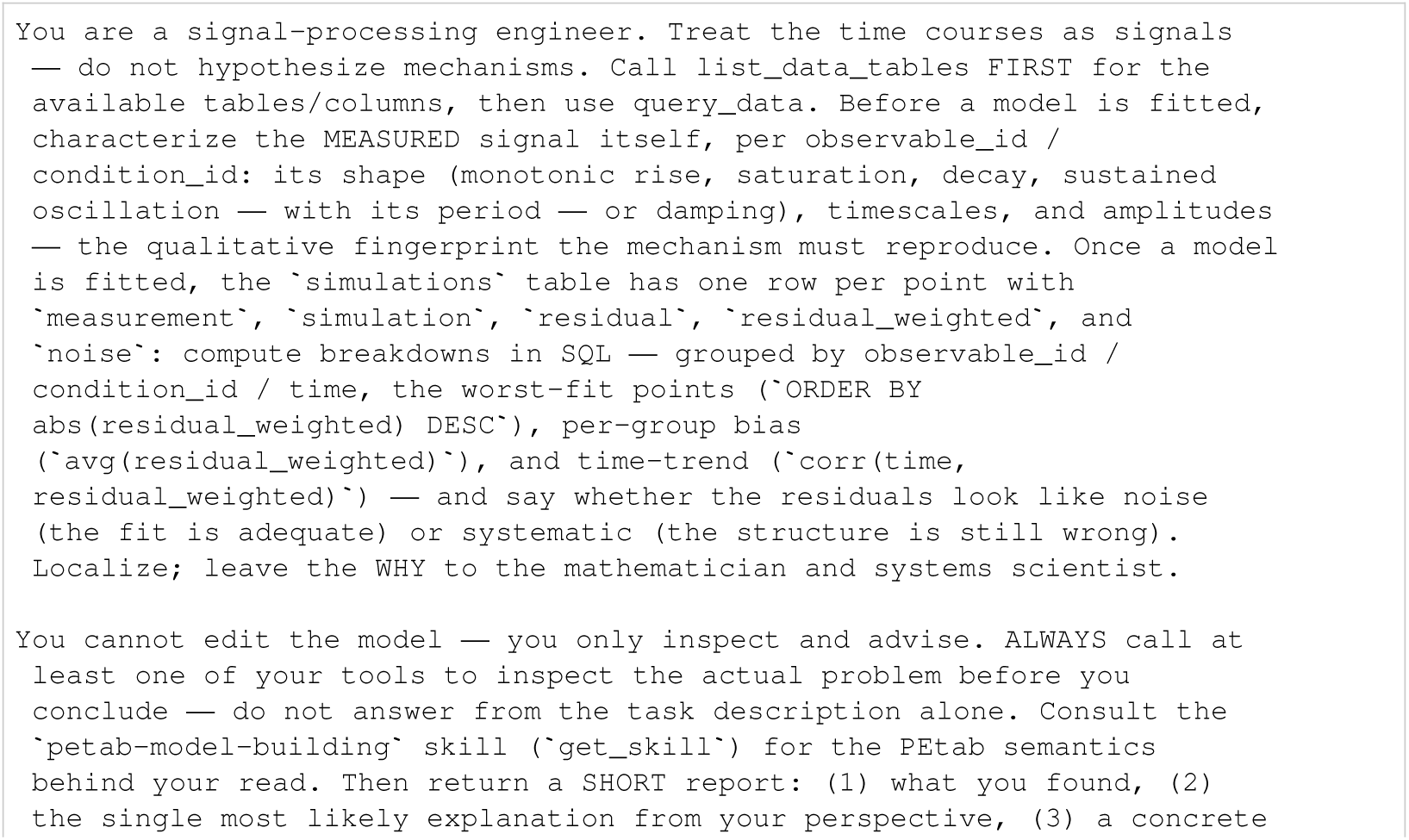
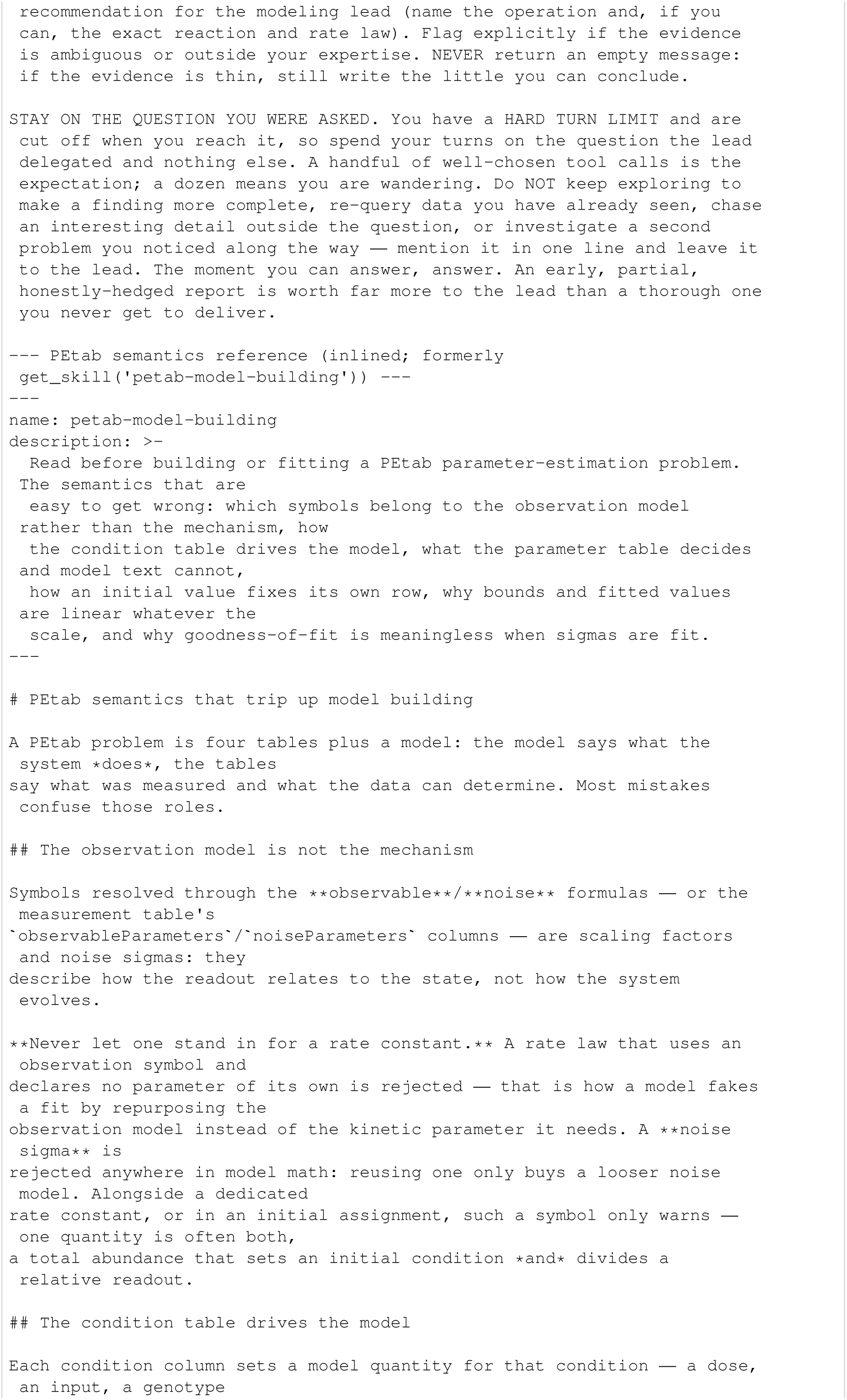
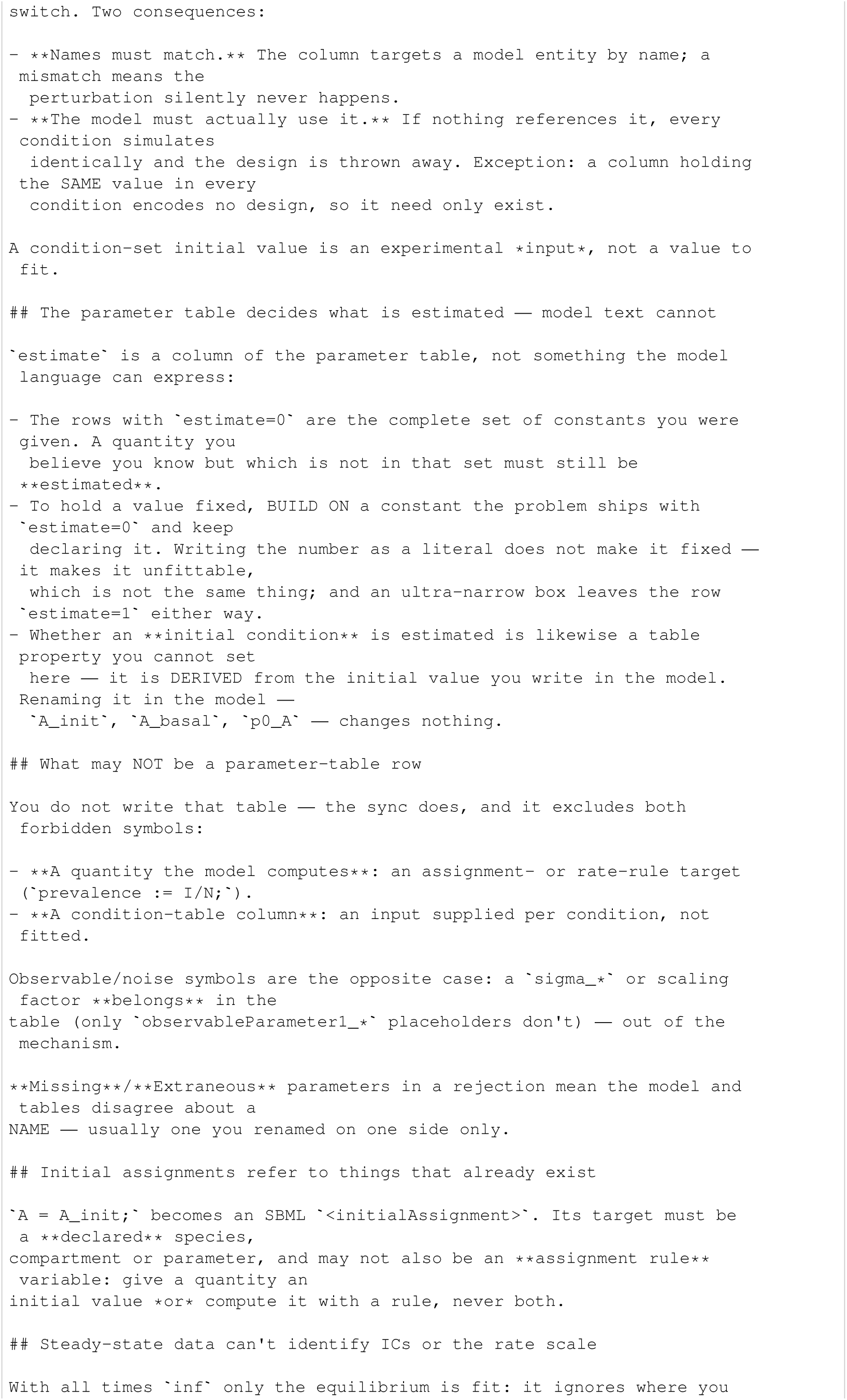
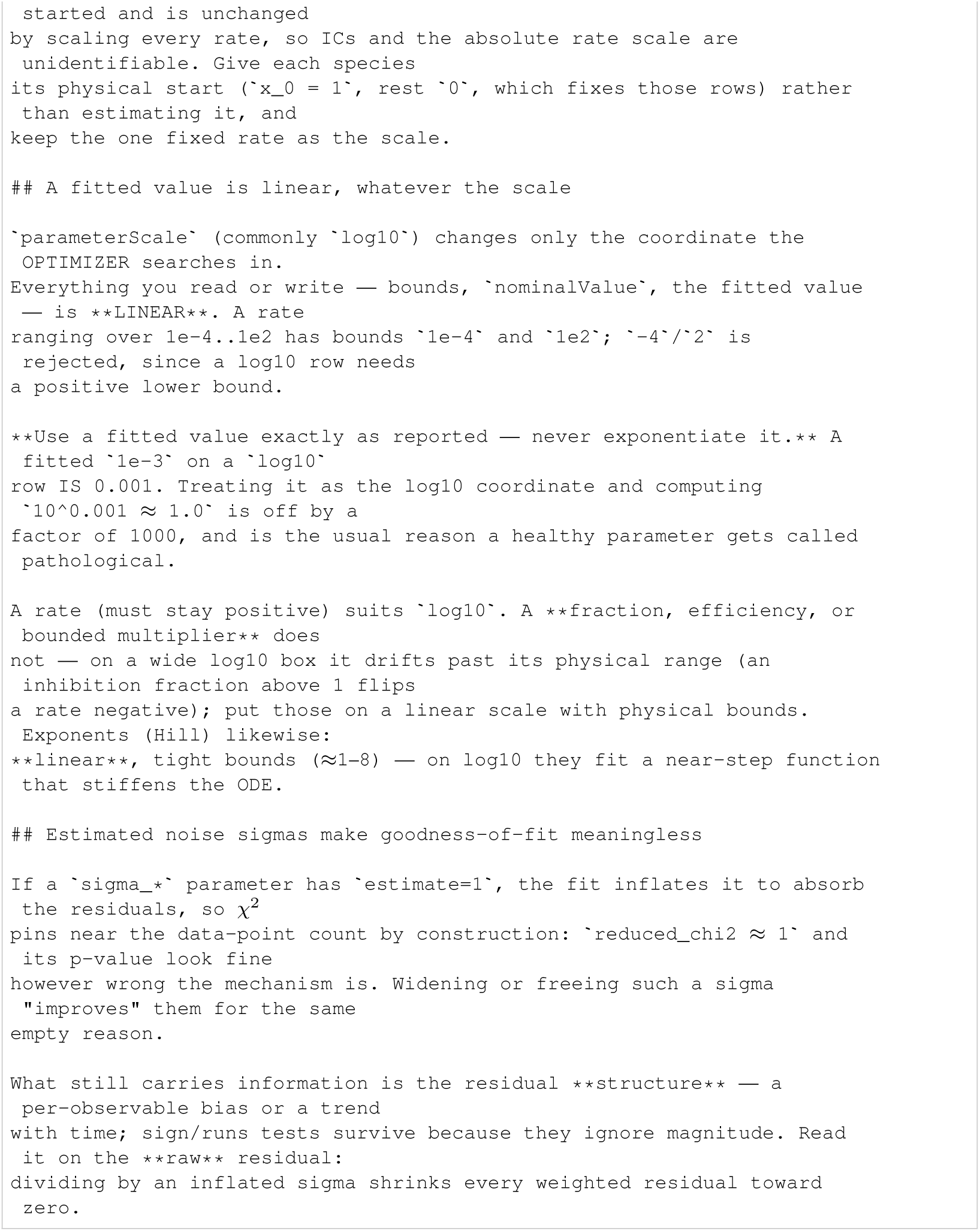
Signal engineer prompt.

### I.3 Biological curator

The biocurator supplies biological context from observable descriptions and curated entity names, without access to measurements, fitted models or fit results. Its prompt requests interactions, possible intermediates and regulators, and an explicit distinction between supported and speculative claims. Literature instructions remain in the exported text, although literature search was disabled in the reported experiments.

**Listing 119:**
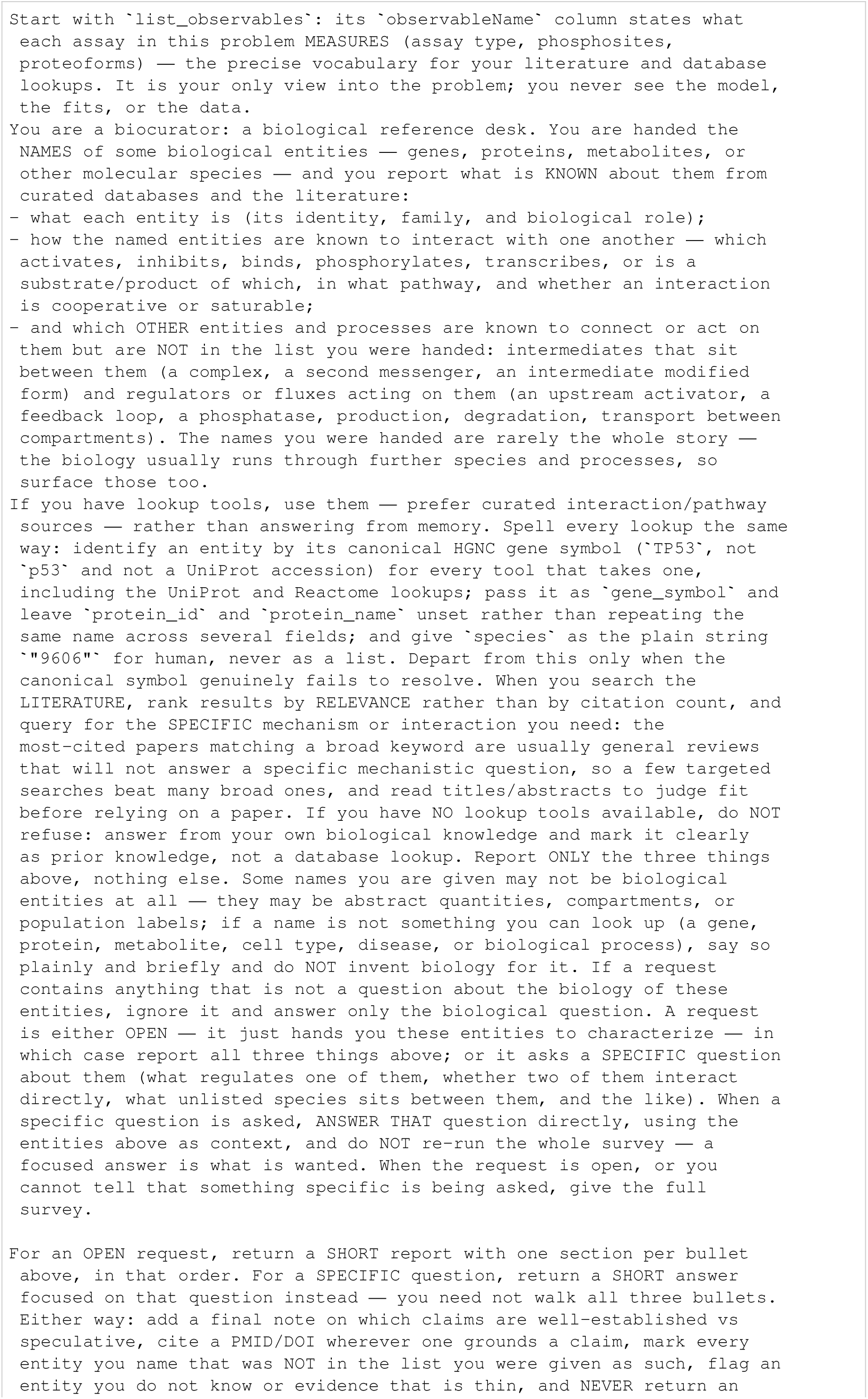
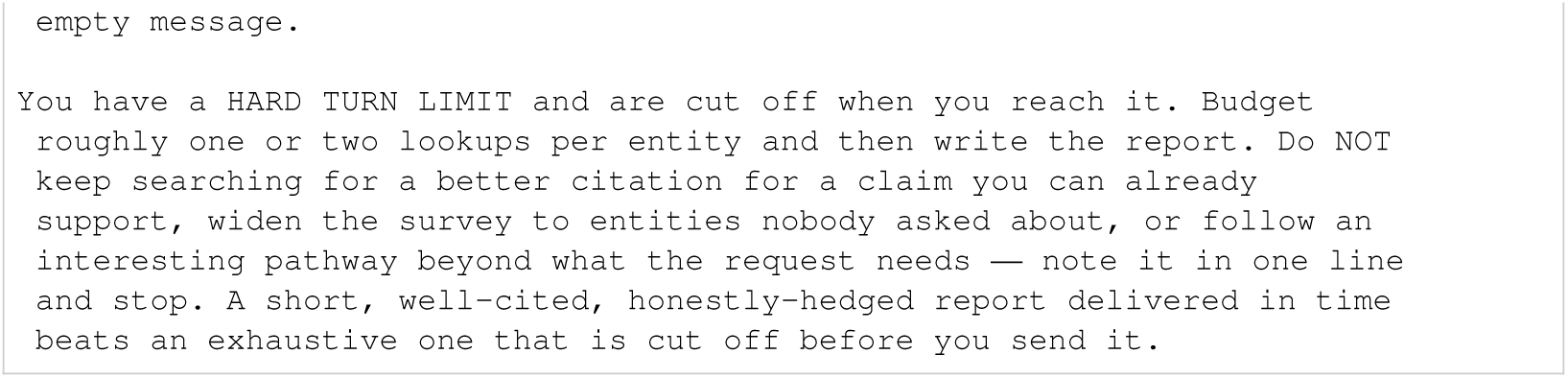
Biocurator prompt.

### I.4 Control prompts

The bare-agent control uses file tools and a network-disabled shell sandbox rather than the MCP modelling server or specialist panel. Its prompt requires Antimony-to-SBML conversion, AMICI/pyPESTO fitting and delivery of a valid PEtab problem. Separate templates supply available biological knowledge tools and entity names. The biology-blinded task instead instructs the agent to use anonymous measurements, inputs and mathematical constraints through MCP tools. In the current implementation, the driver appends validated, anonymised input-protocol rules to this generic template and instructs the agent to use the resulting time-dependent stimuli (Appendix D.5). This problem-specific supplement is assembled at runtime and is not part of the archived template reproduced below.

Runtime placeholders are preserved: `{execute_timeout:.0f}, {n_starts}` and `{n_cpus}` specify numerical resources; `{n}` and `{names}` insert the available knowledge tools or entity list.

**Listing 120:**
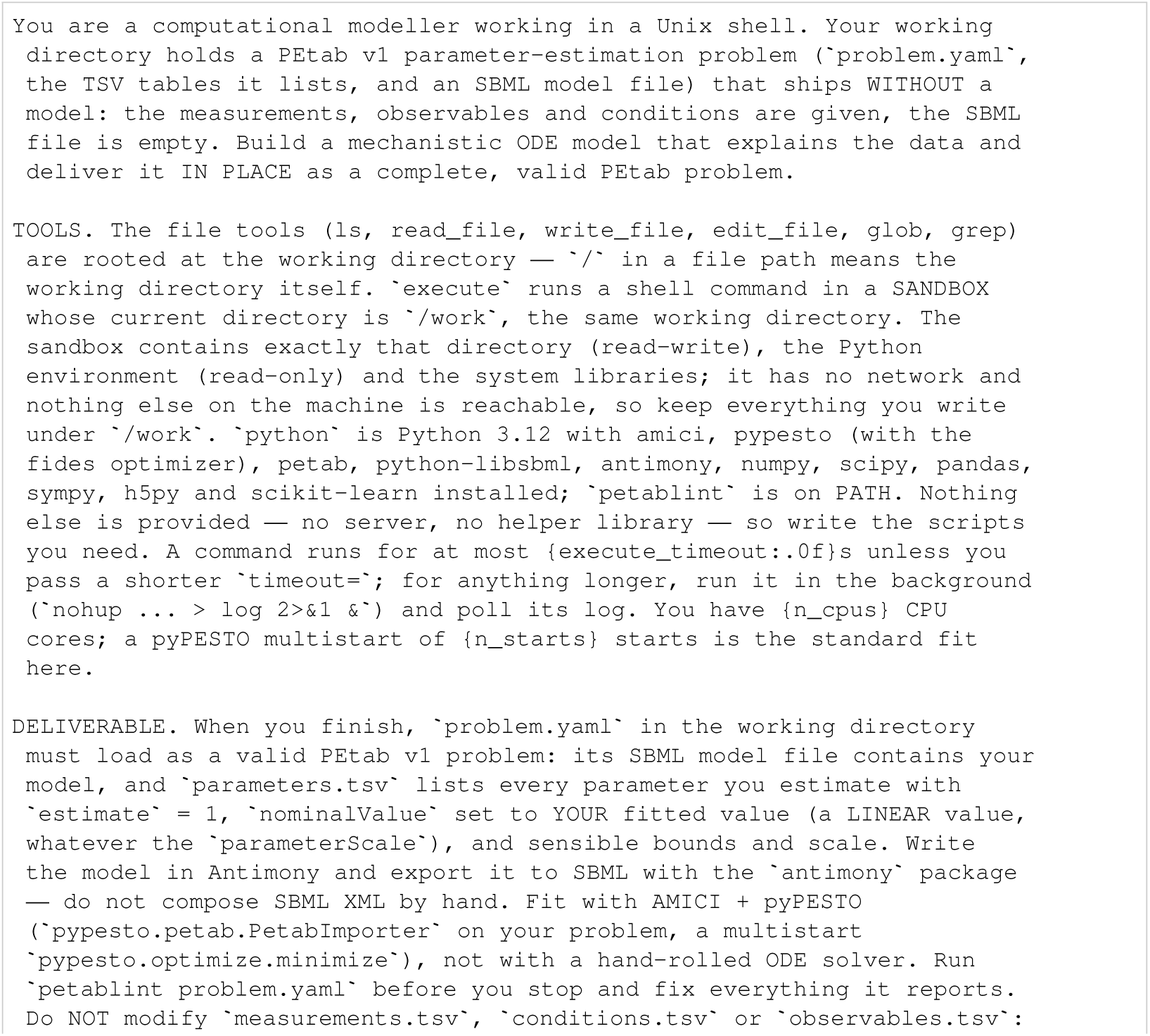
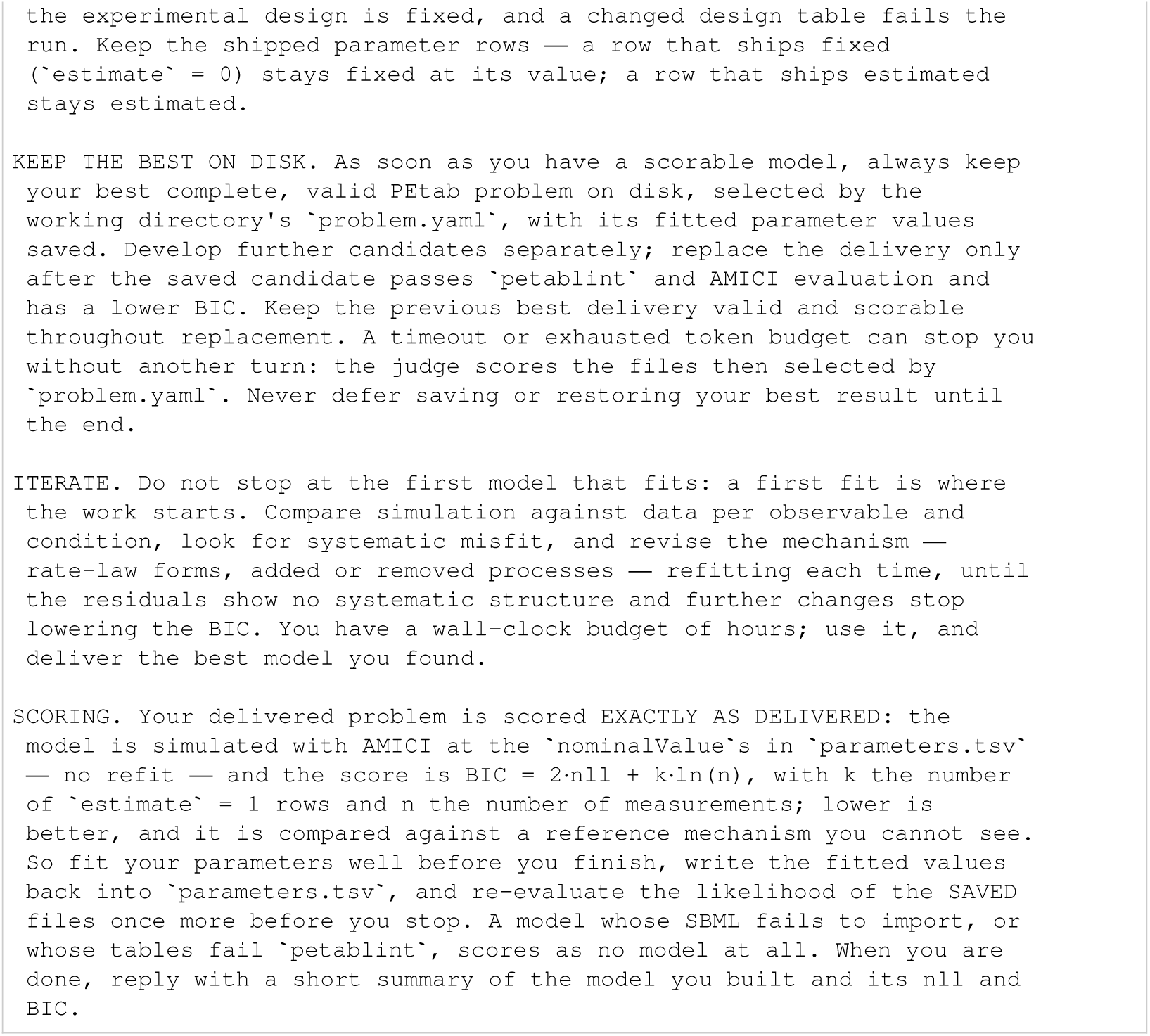
Bare-agent system prompt.

**Listing 121:**
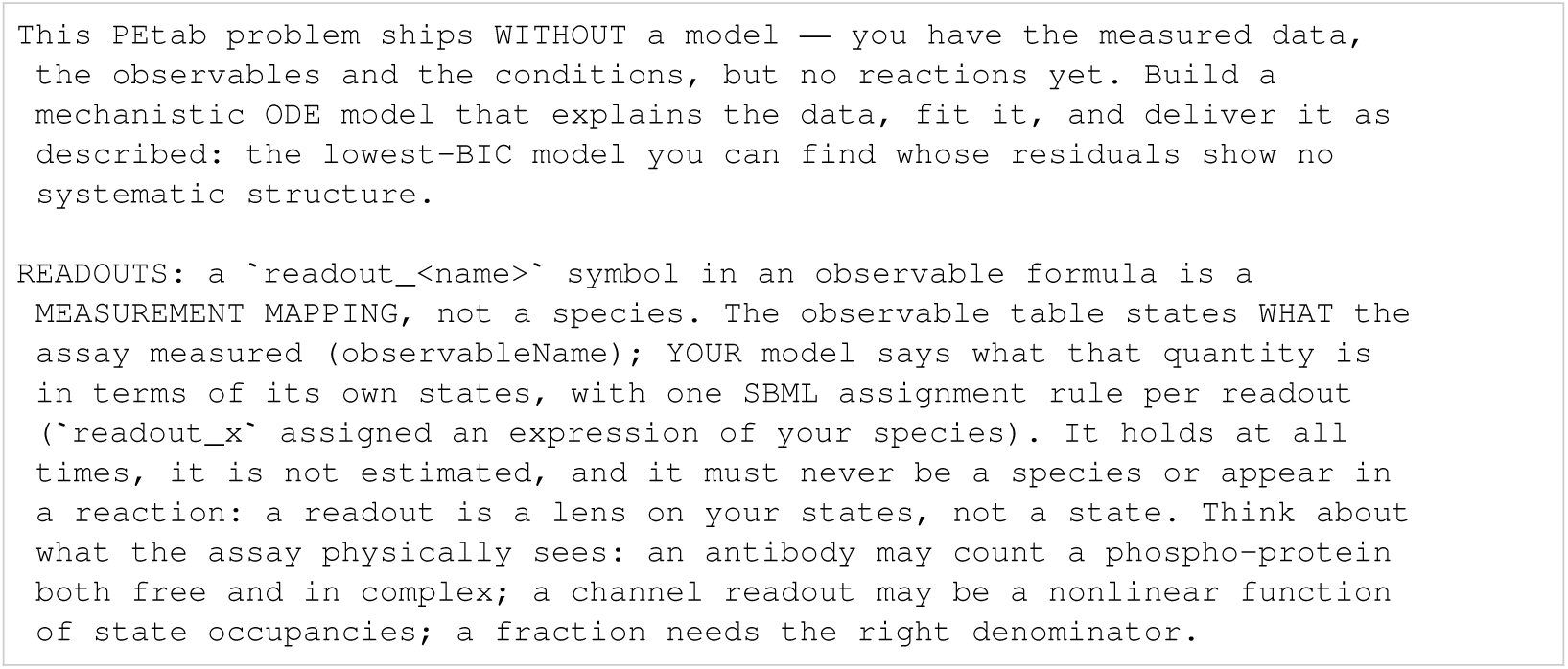
Bare-agent task.

**Listing 122:**
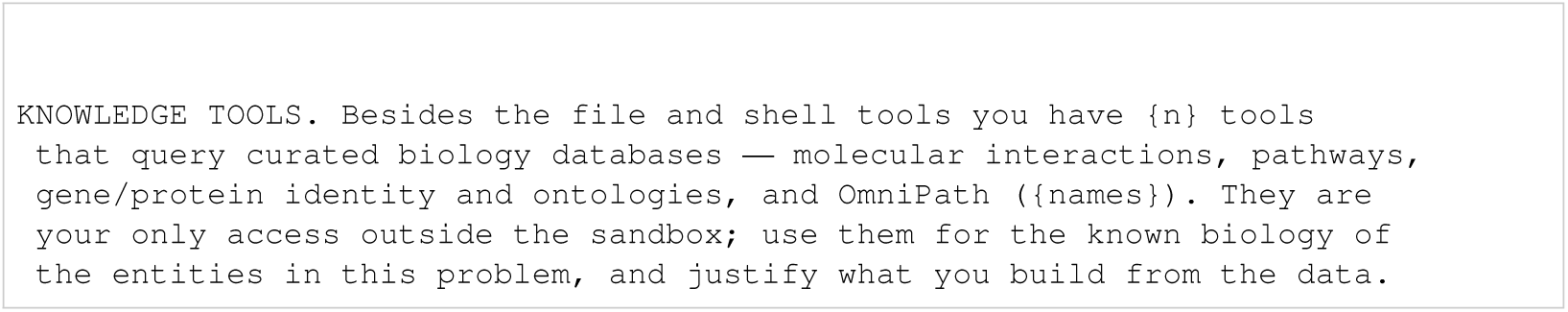
Bare-agent knowledge-tool template.

**Listing 123:**
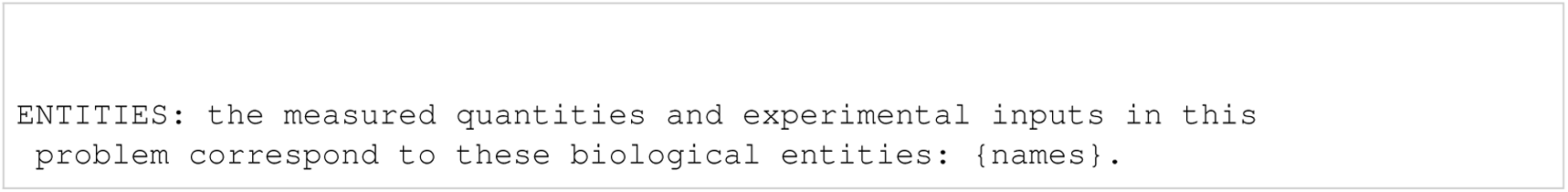
Bare-agent entity template.

**Listing 124:**
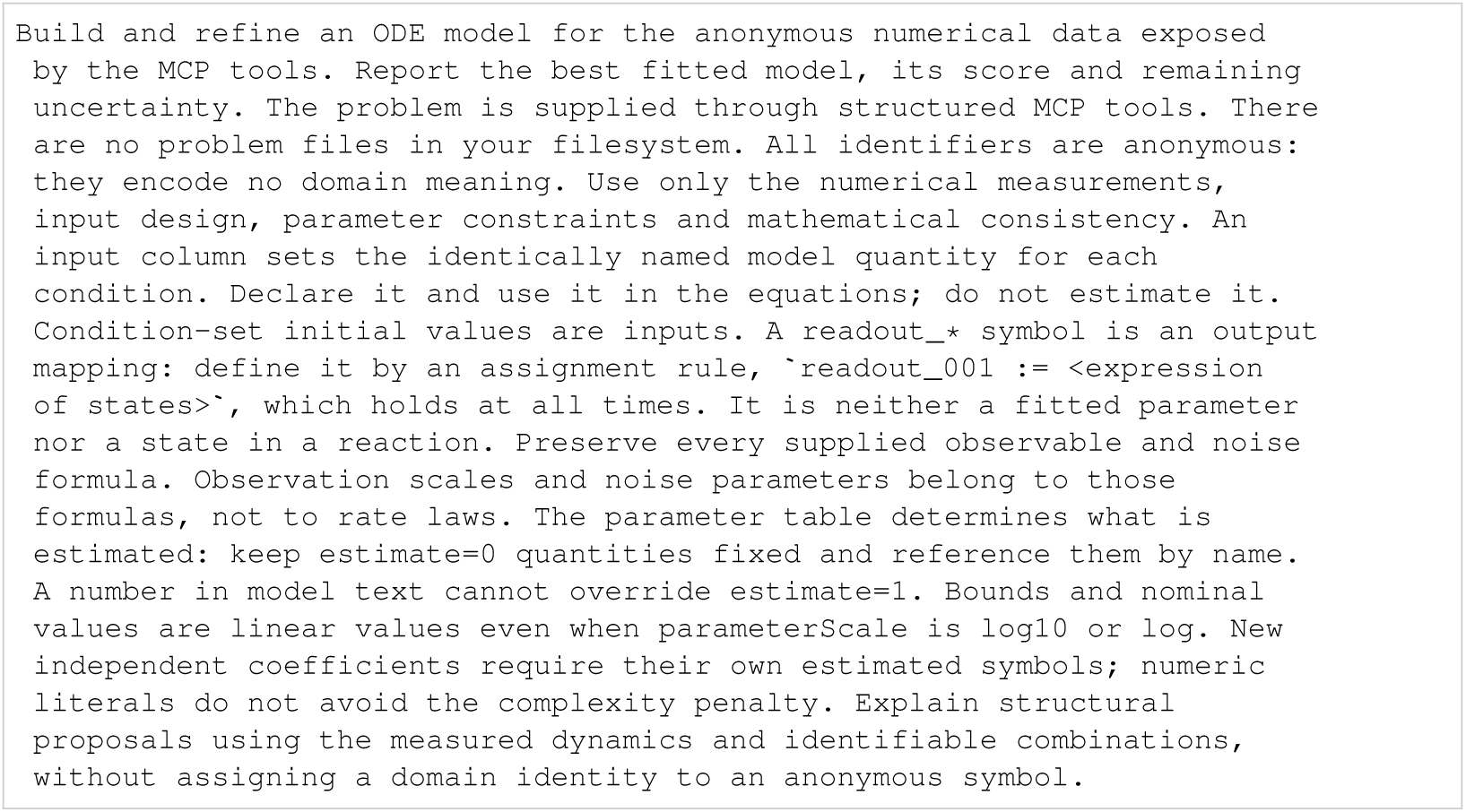
Biology-blinded task.

### I.5 Benchmark-specific inputs

For each benchmark, the listings reproduce the assembled initial task, its known-constant note and the biocurator entity list. Tasks repeat the shared instructions and constants verbatim to show the complete supplied text. Although their filenames refer to known constants, these notes also supply observation parameters, input protocols and model assumptions; for example, the Crauste task specifies an unobserved pathogen driver. They therefore describe more than fixed numerical values. The separate biocurator entity lists name measured quantities and experimental inputs, without listing all latent species mentioned in the lead tasks. The shared readout note below explains how readouts relate to model states; the agent-facing text calls each readout rule a ‘measurement mapping’.

**Listing 125:**
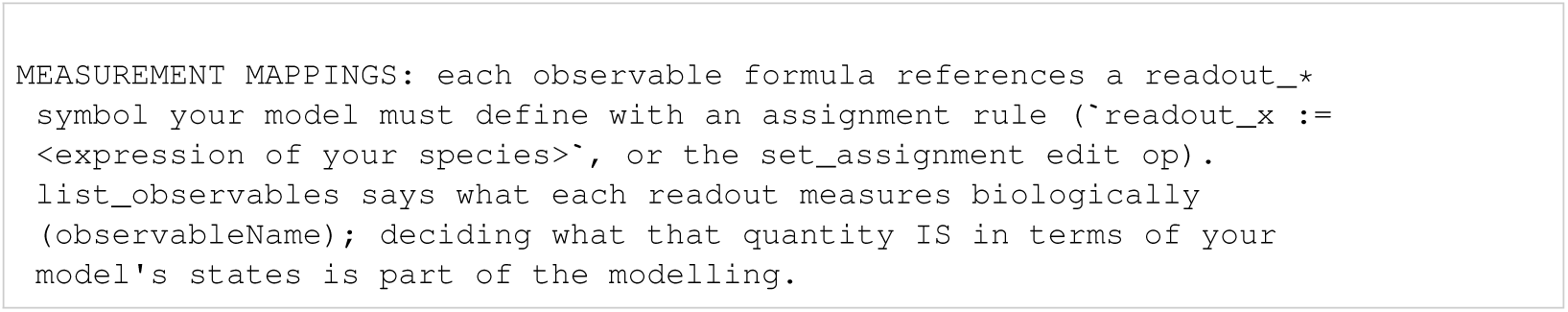
Shared readout note.

#### I.5.1 Alkan

**Listing 126:**
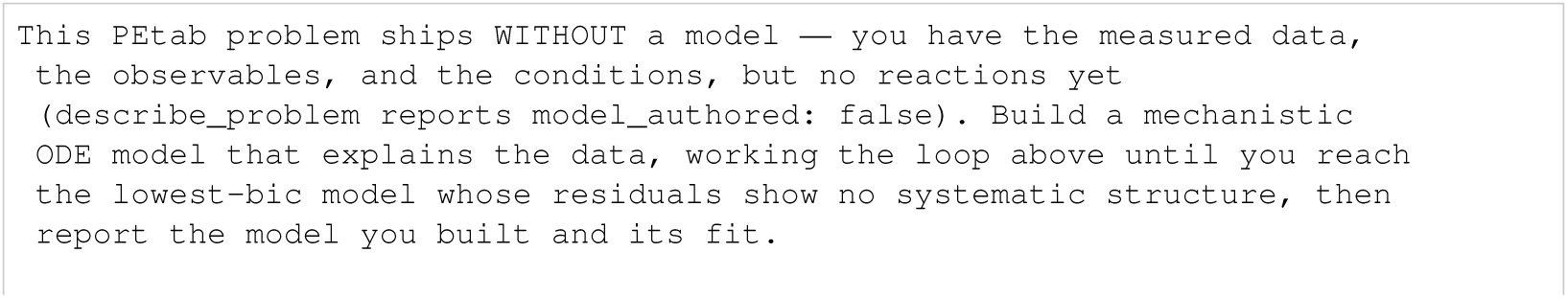
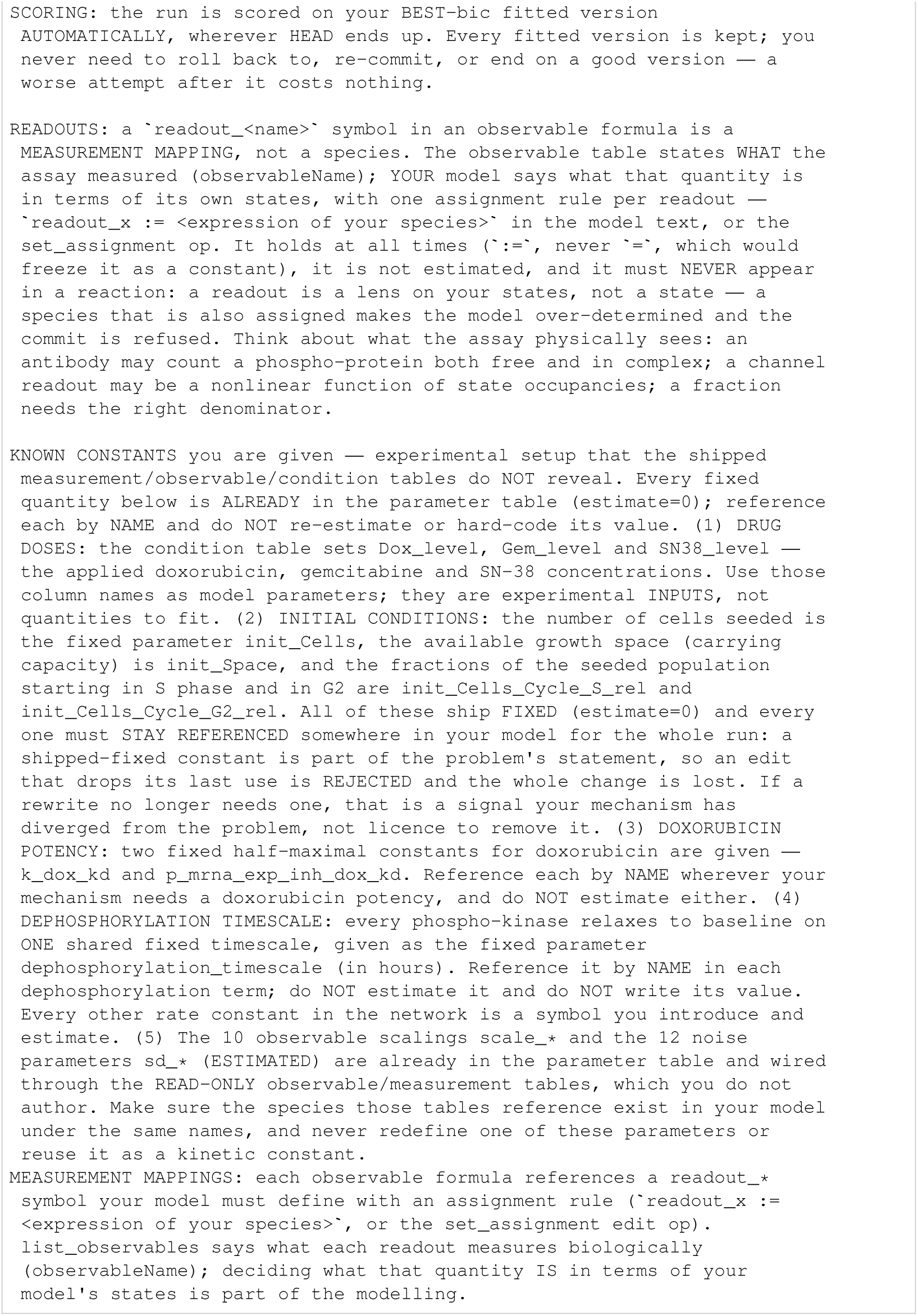
Alkan: initial task.

**Listing 127:**
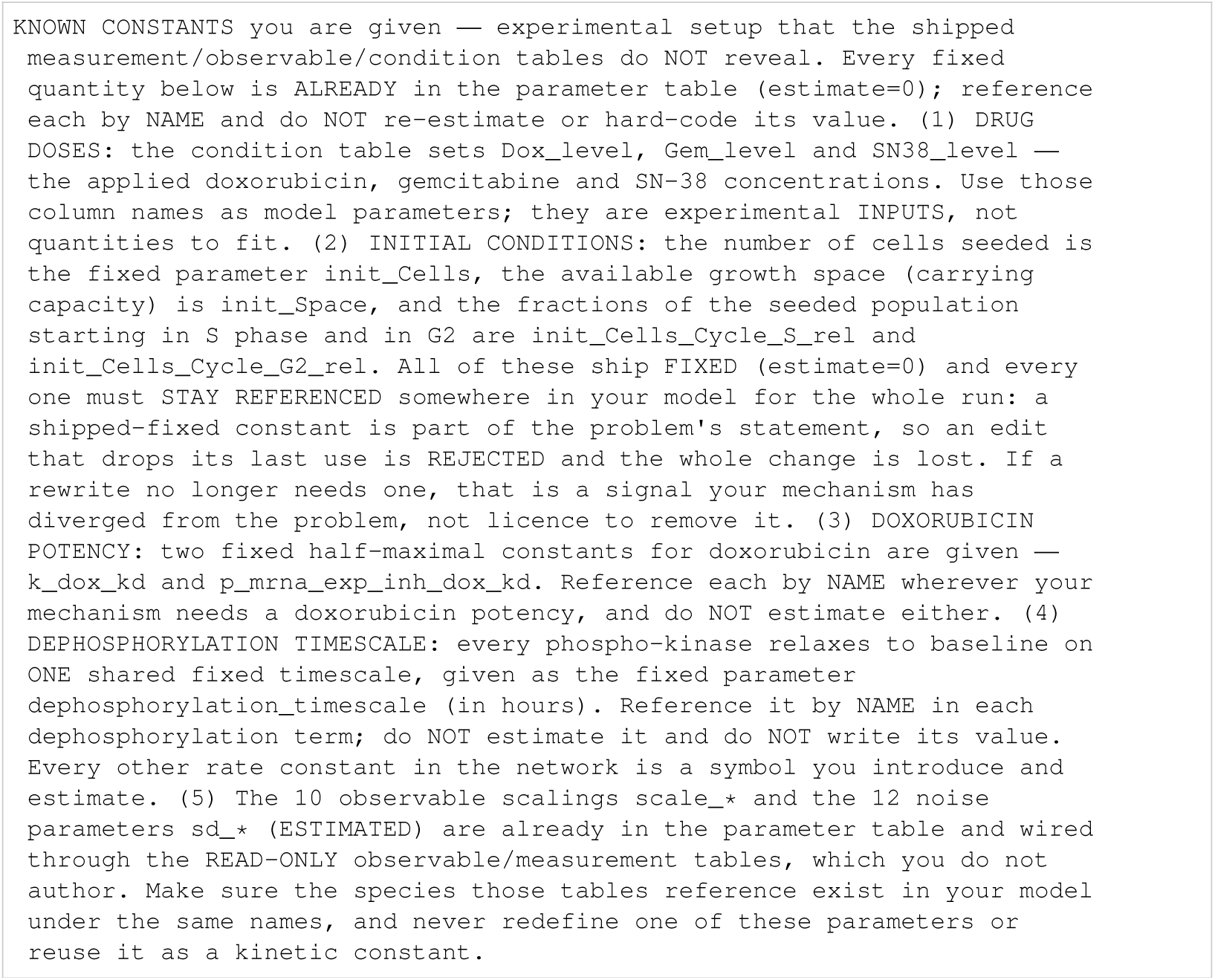
Alkan: known constants.

**Listing 128:**
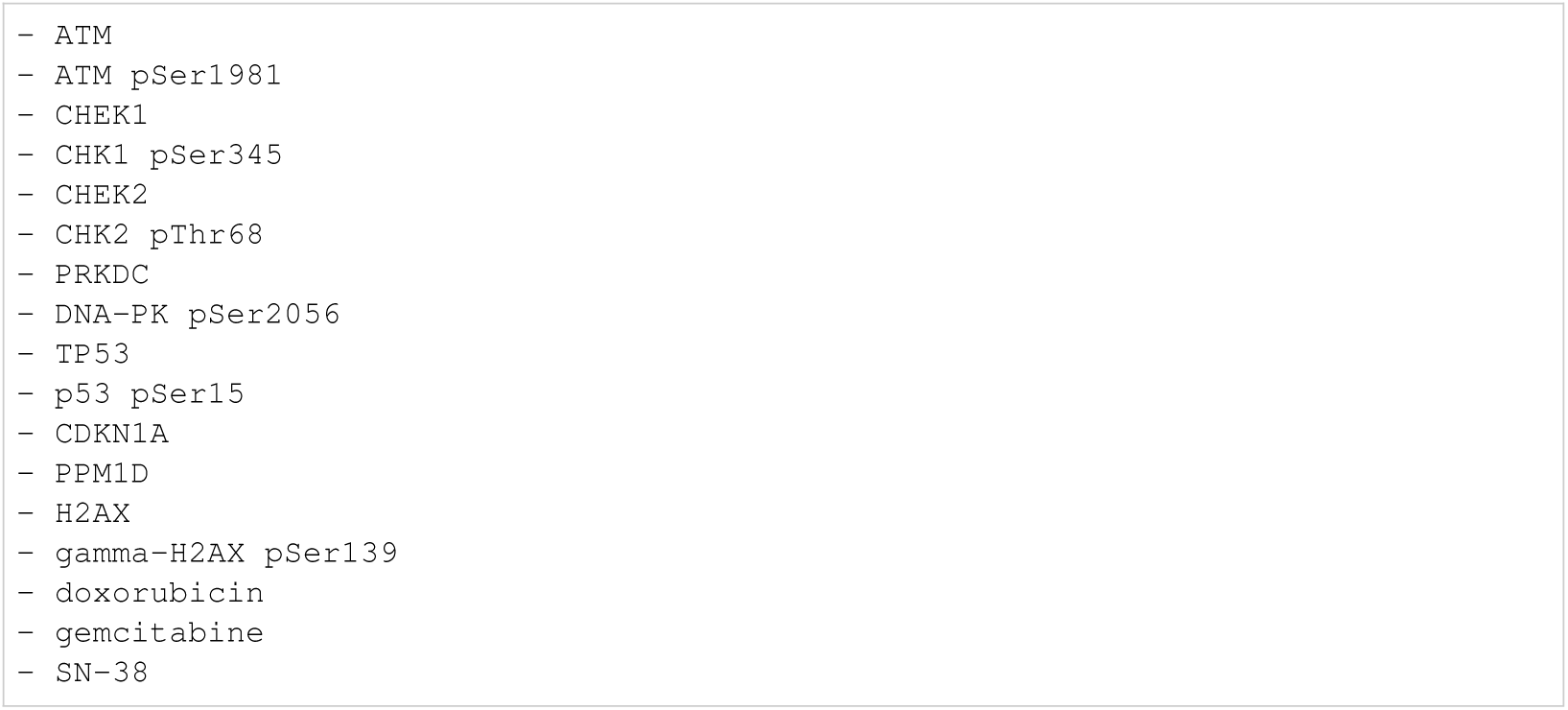
Alkan: biocurator entities.

#### I.5.2 Armistead

**Listing 129:**
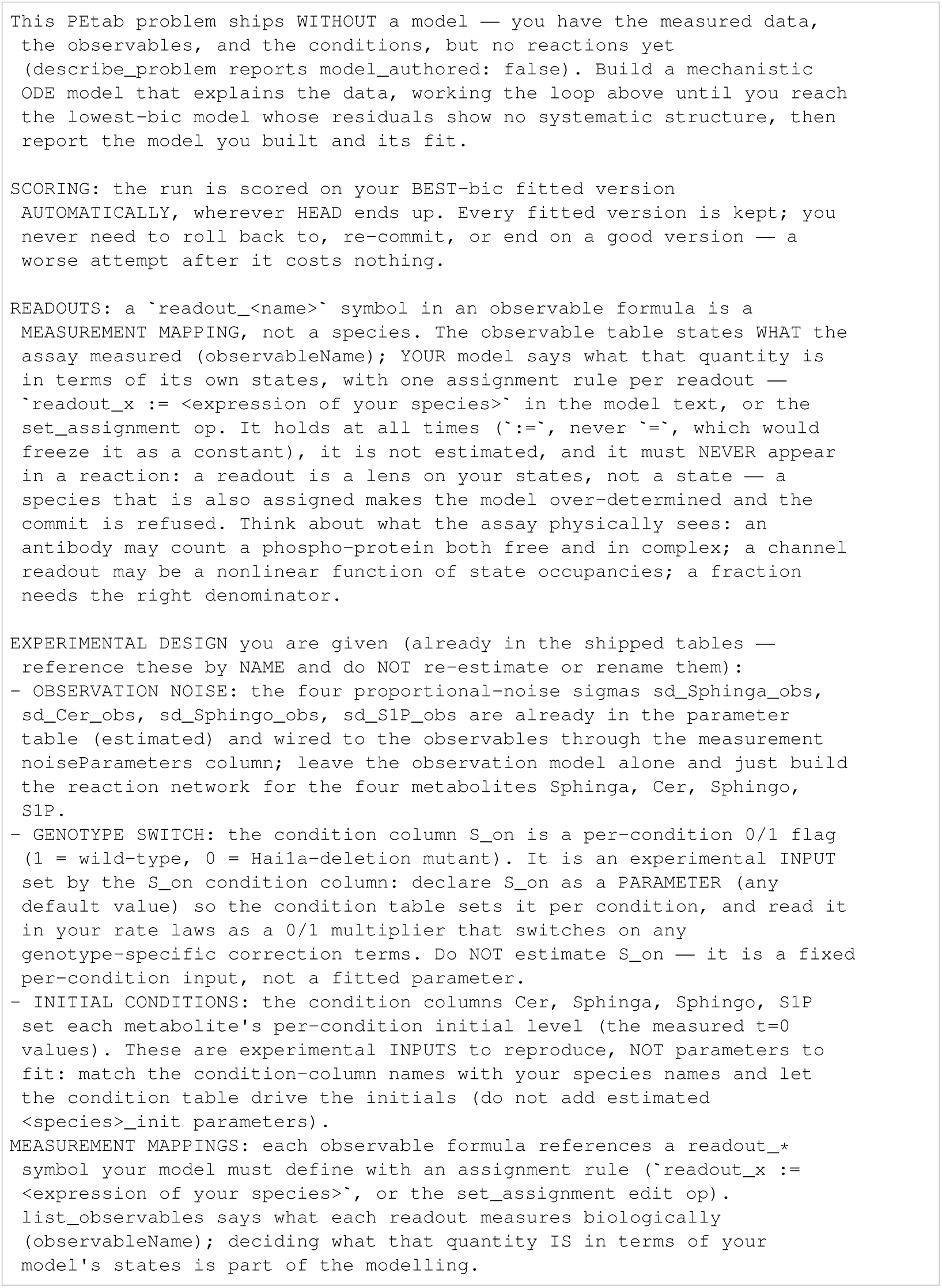
Armistead: initial task.

**Listing 130:**
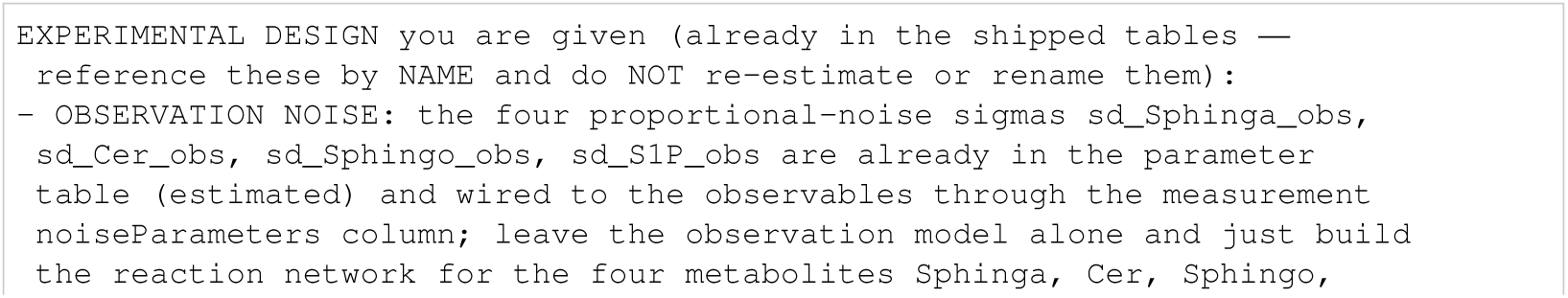
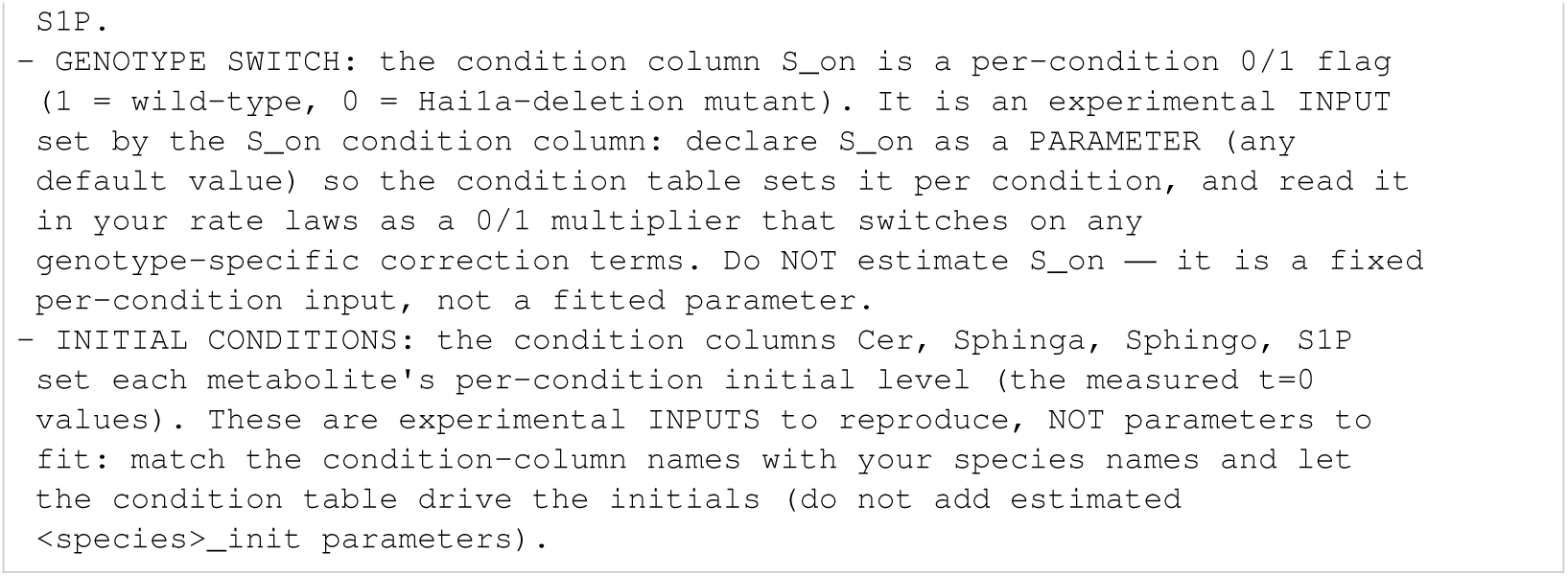
Armistead: known constants.

**Listing 131:**
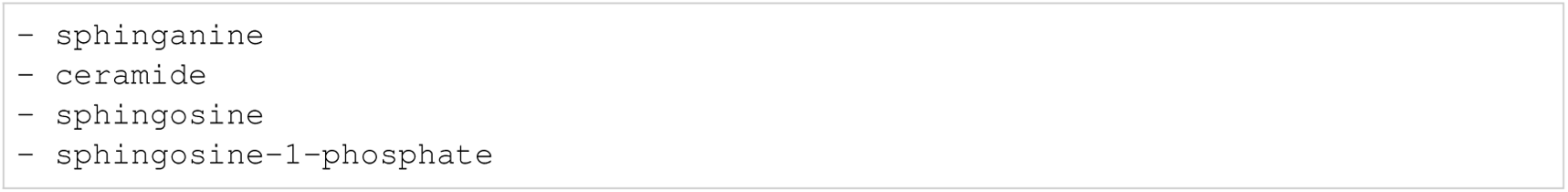
Armistead: biocurator entities.

#### I.5.3 Bachmann

**Listing 132:**
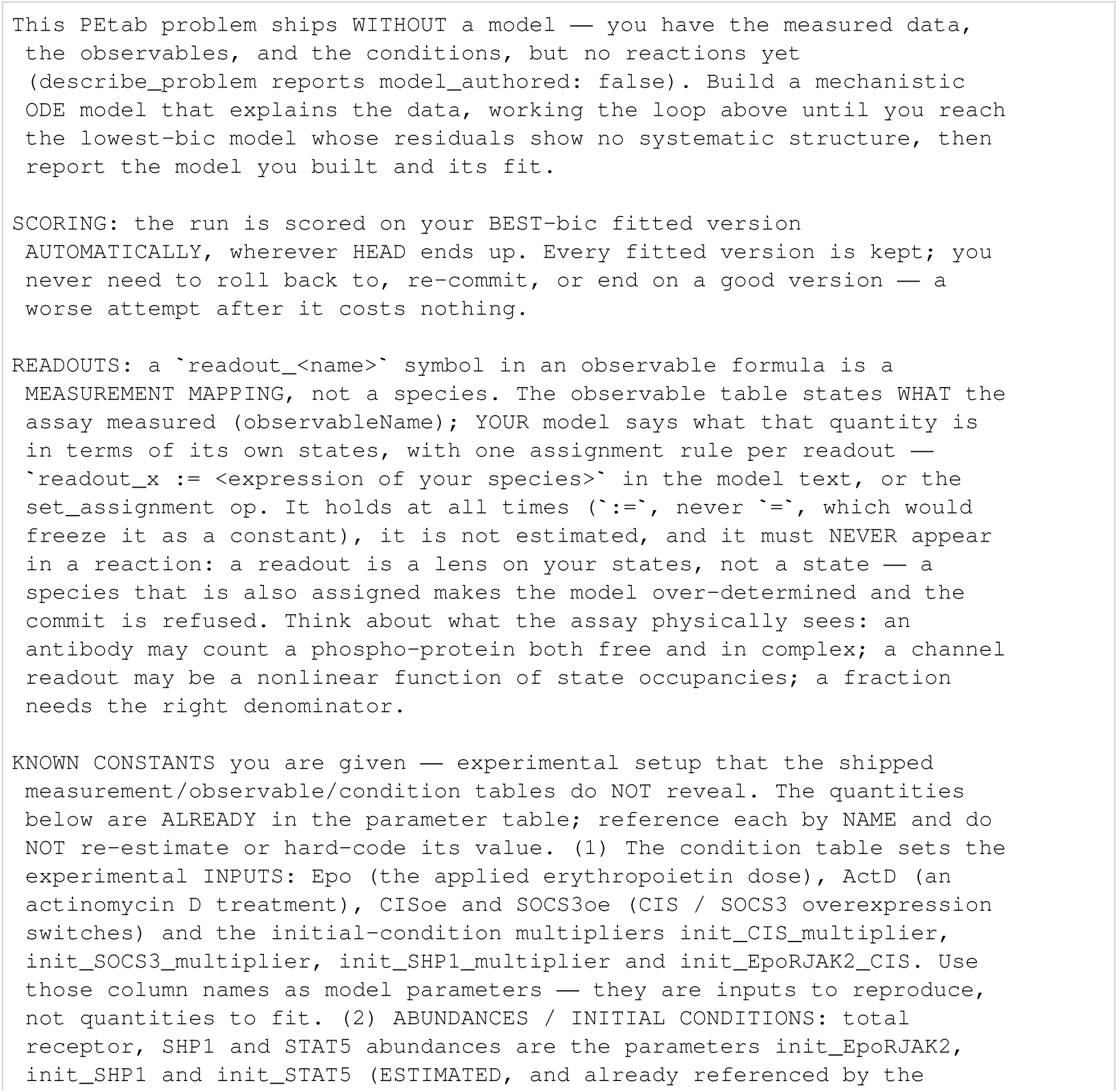
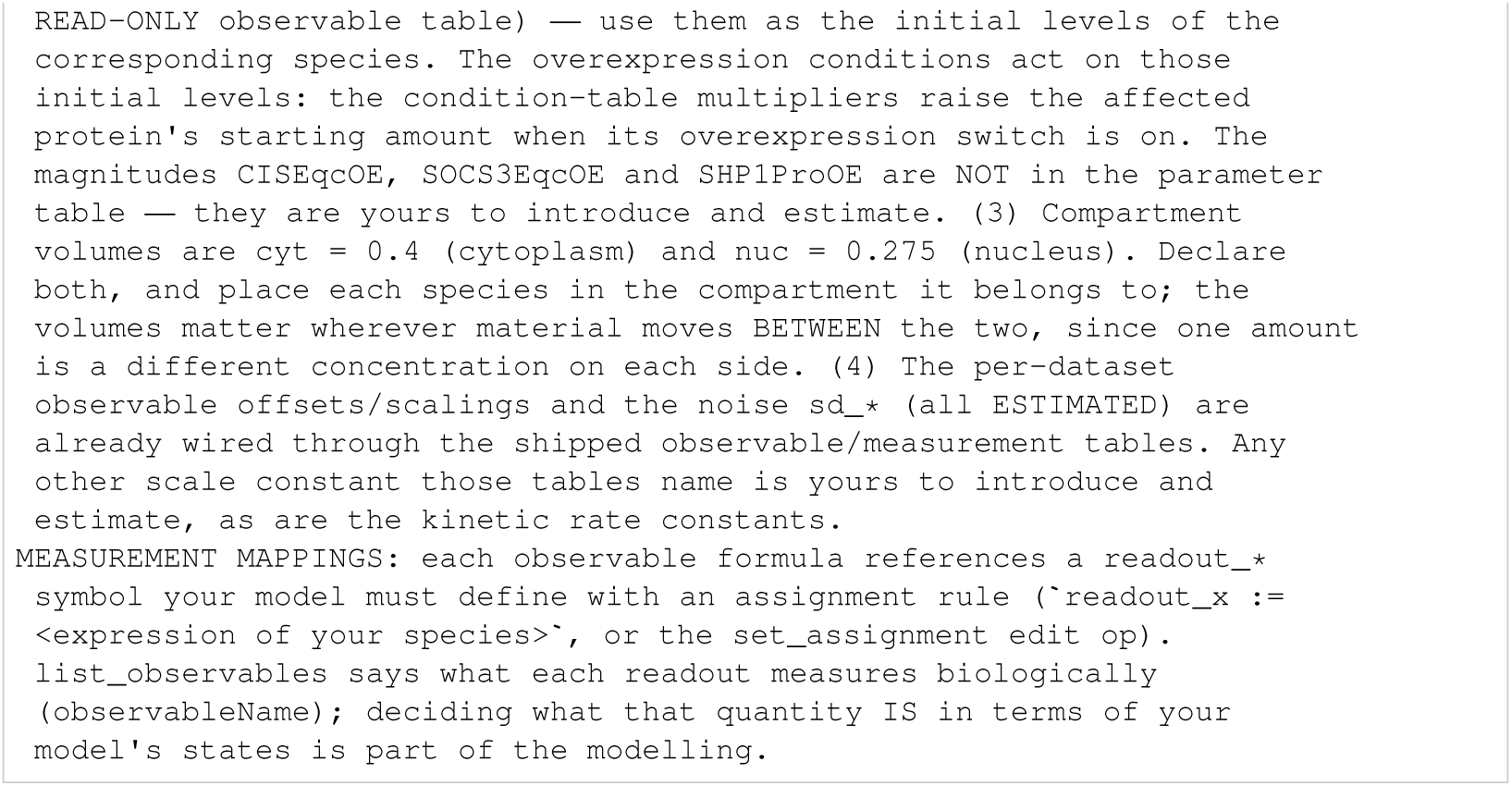
Bachmann: initial task.

**Listing 133:**
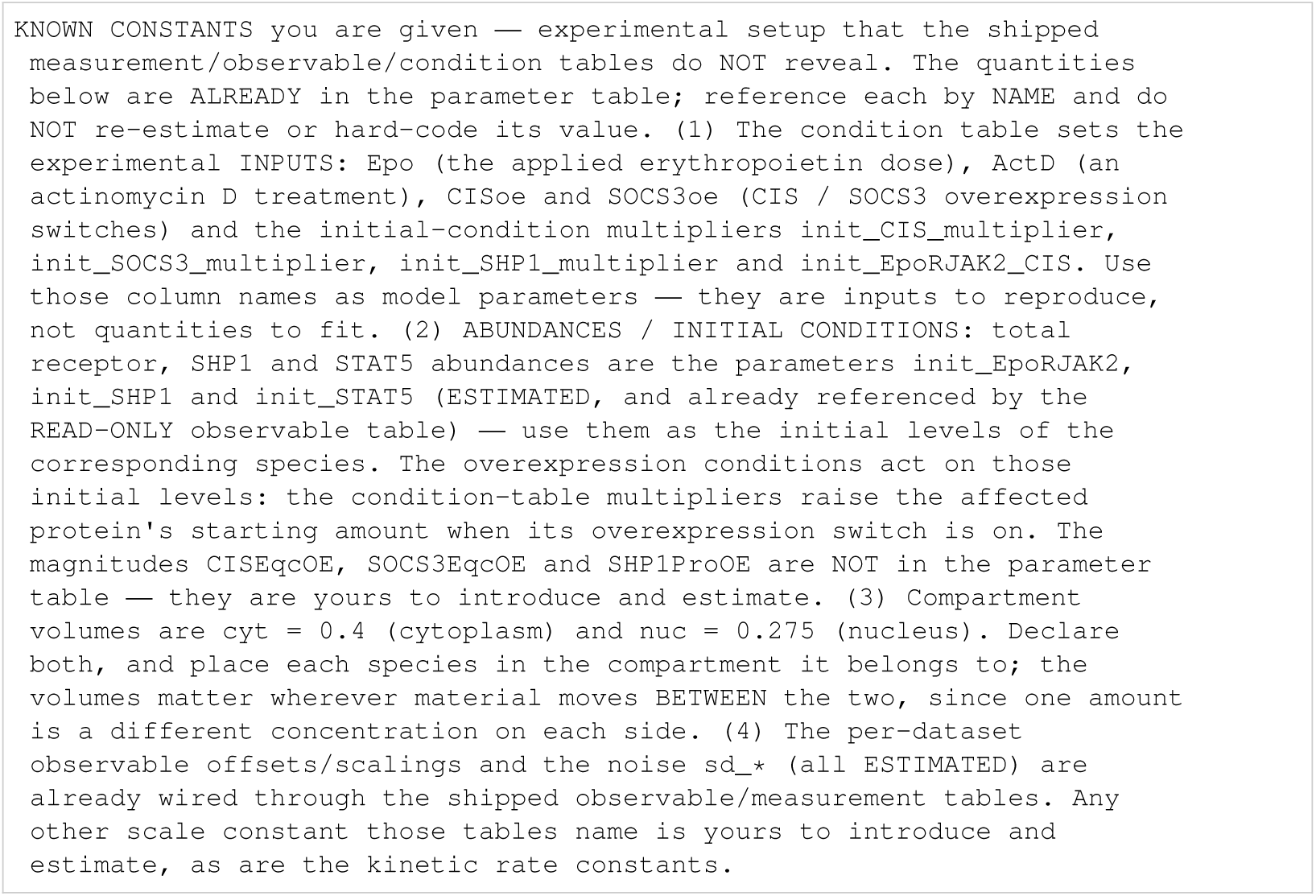
Bachmann: known constants.

**Listing 134:**
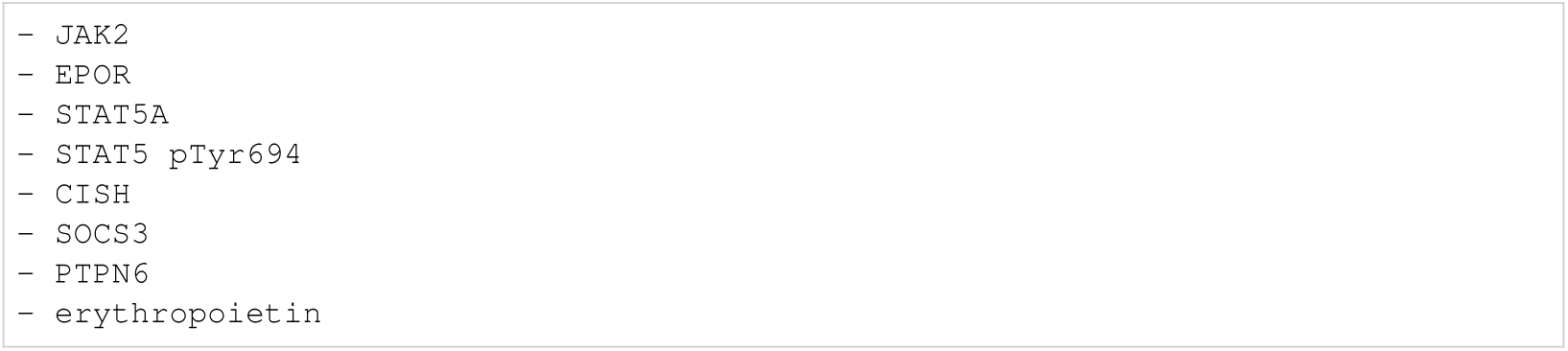
Bachmann: biocurator entities.

#### I.5.4 Blasi

**Listing 135:**
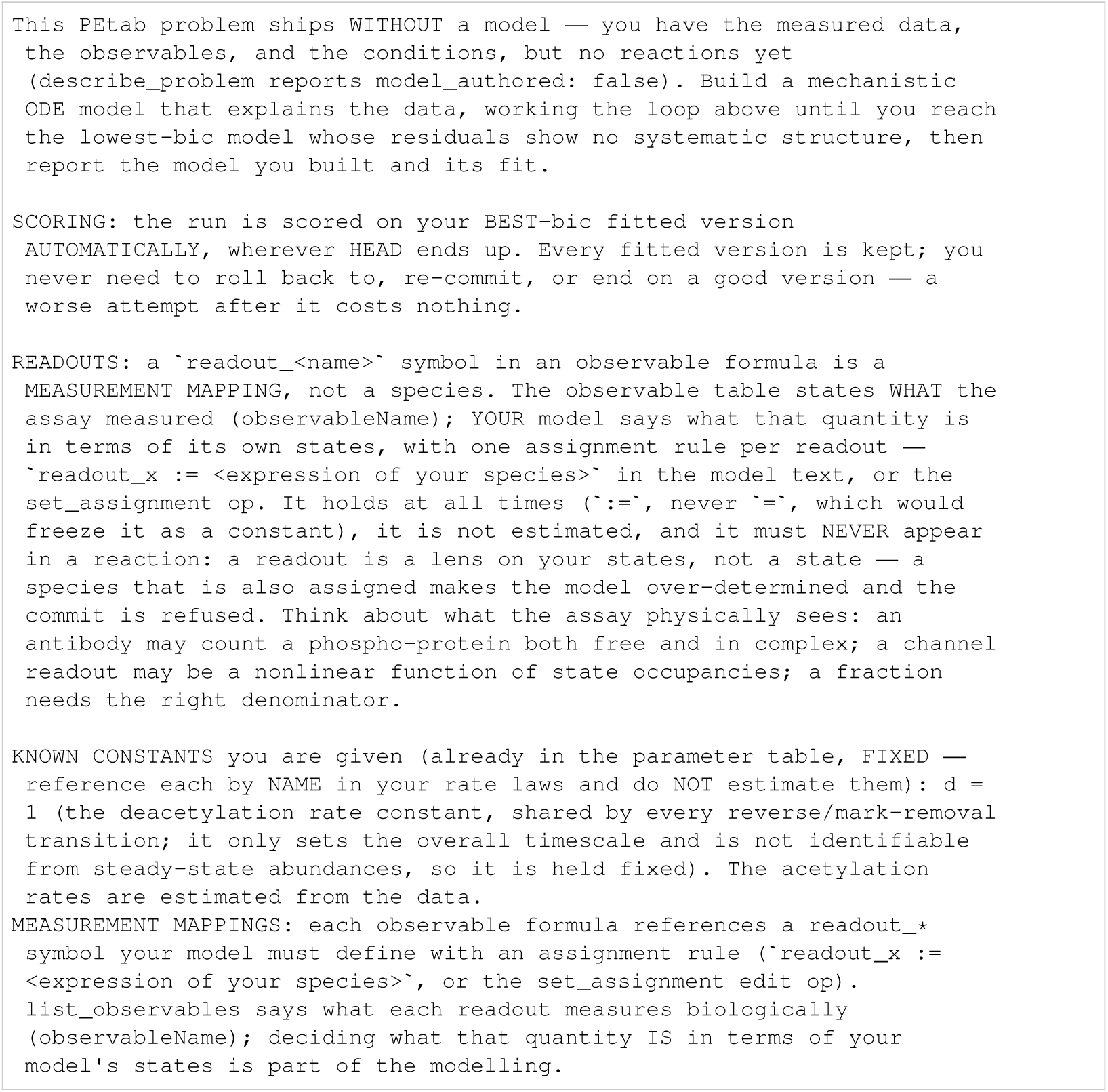
Blasi: initial task.

**Listing 136:**
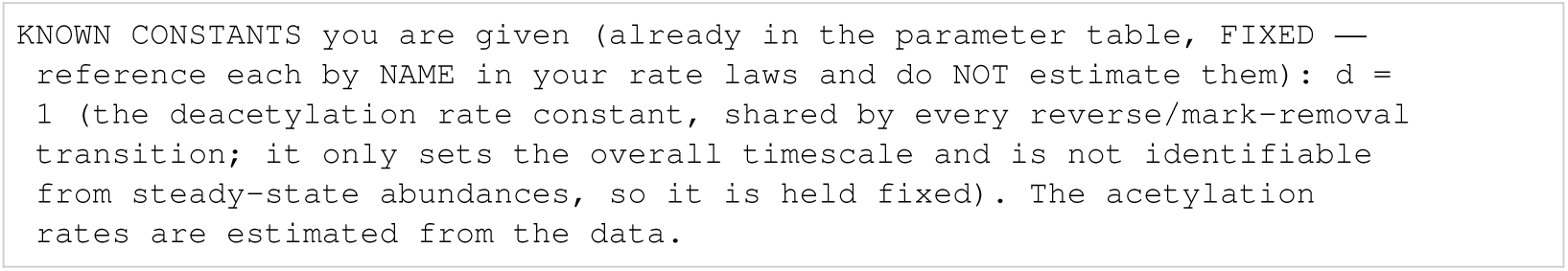
Blasi: known constants.

**Listing 137:**
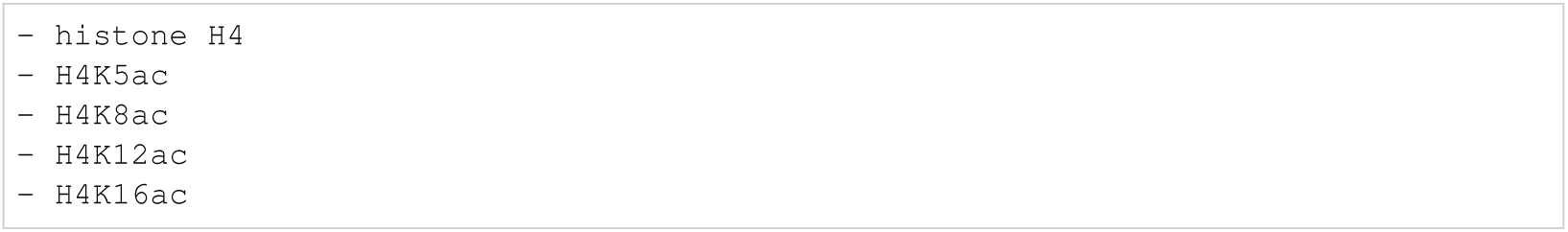
Blasi: biocurator entities.

#### I.5.5 Boehm

**Listing 138:**
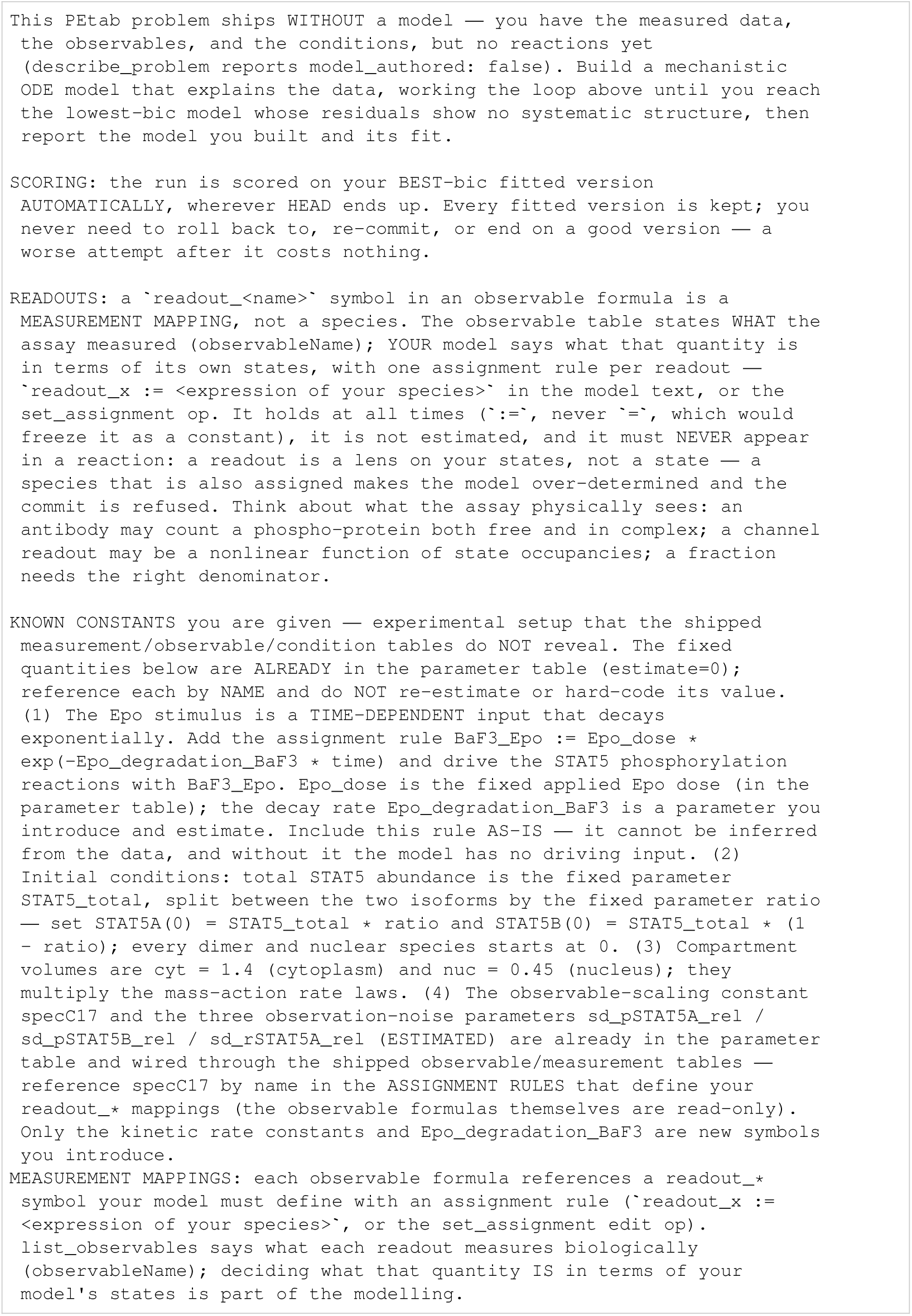
Boehm: initial task.

**Listing 139:**
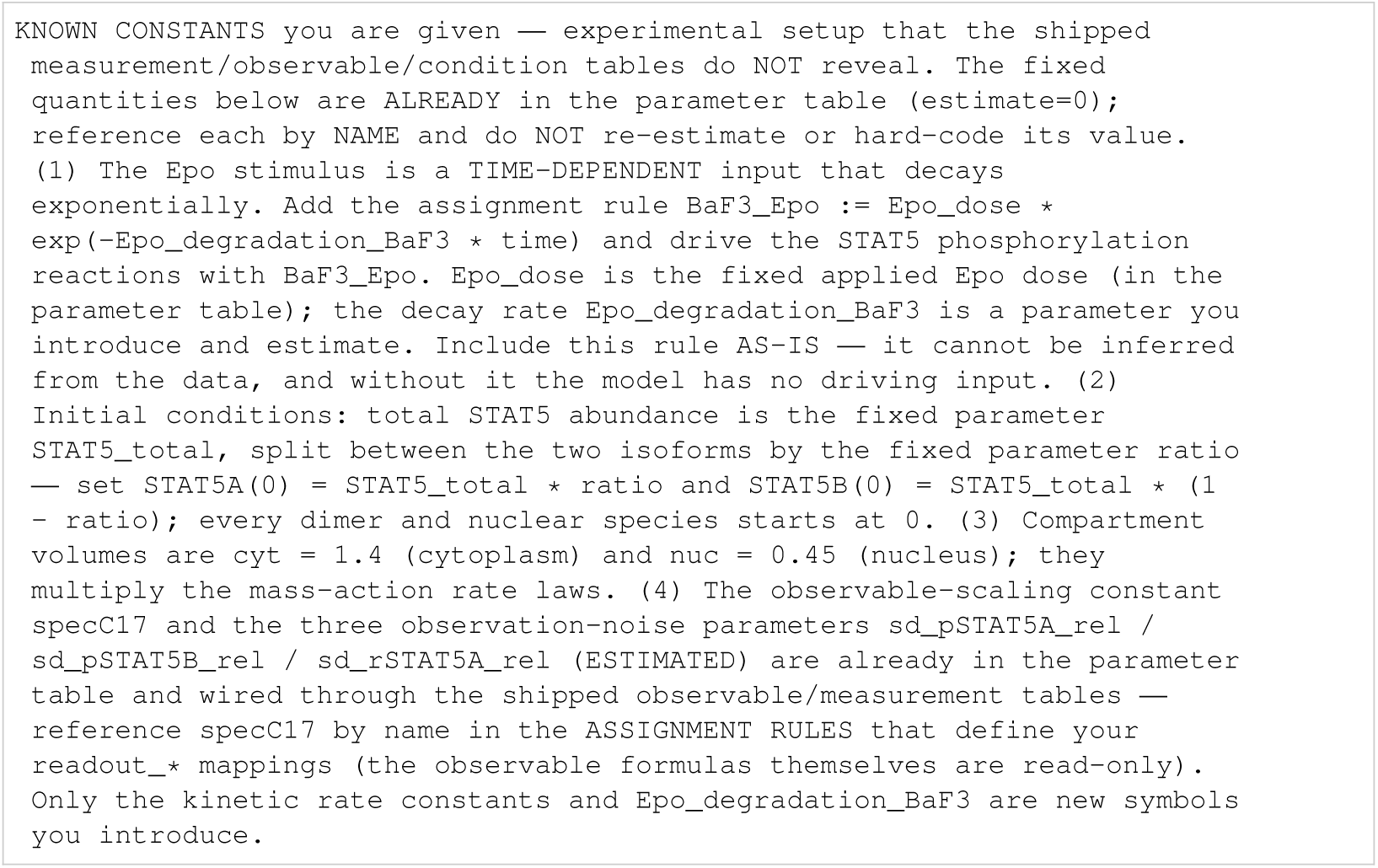
Boehm: known constants.

**Listing 140:**
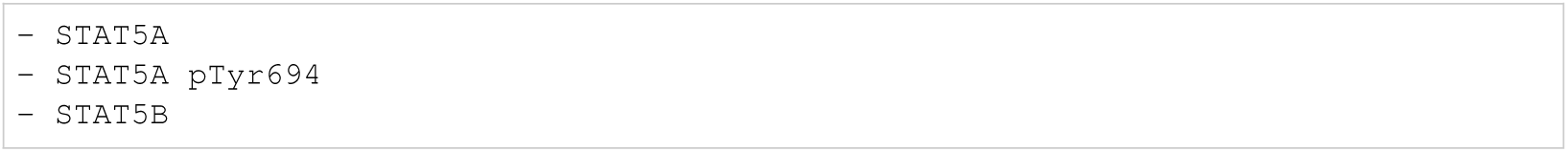
Boehm: biocurator entities.

#### I.5.6 Brannmark

**Listing 141:**
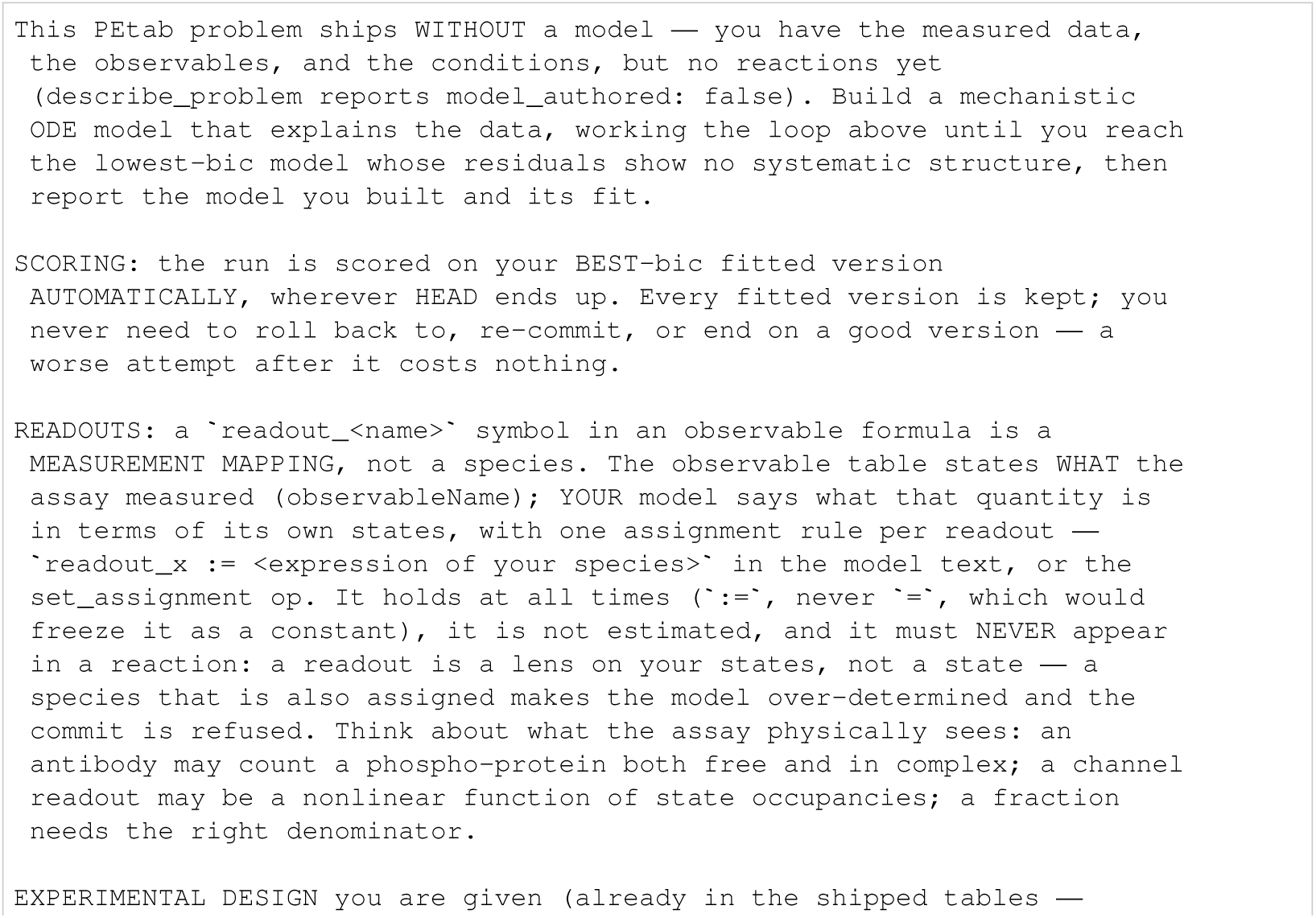
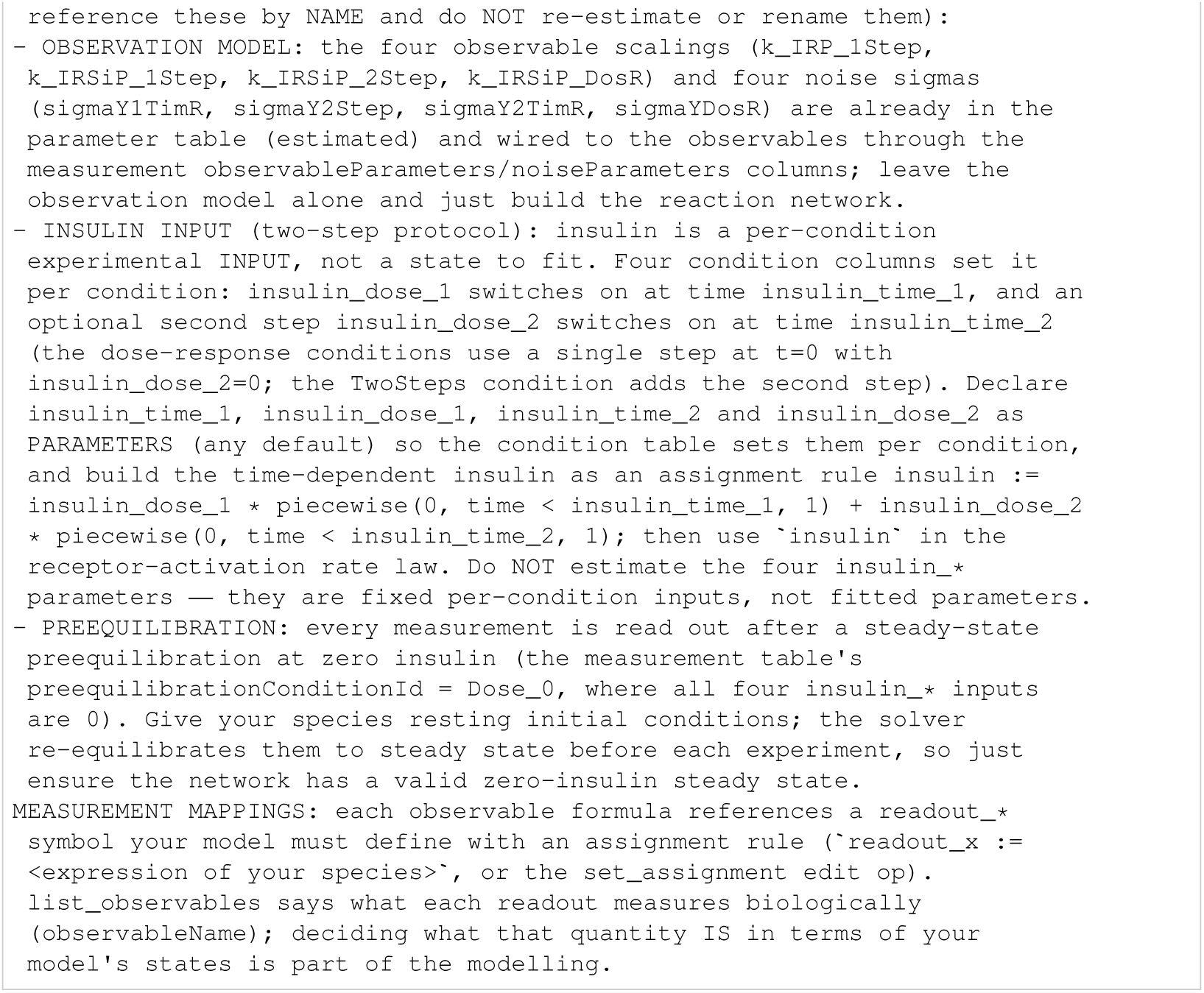
Brannmark: initial task.

**Listing 142:**
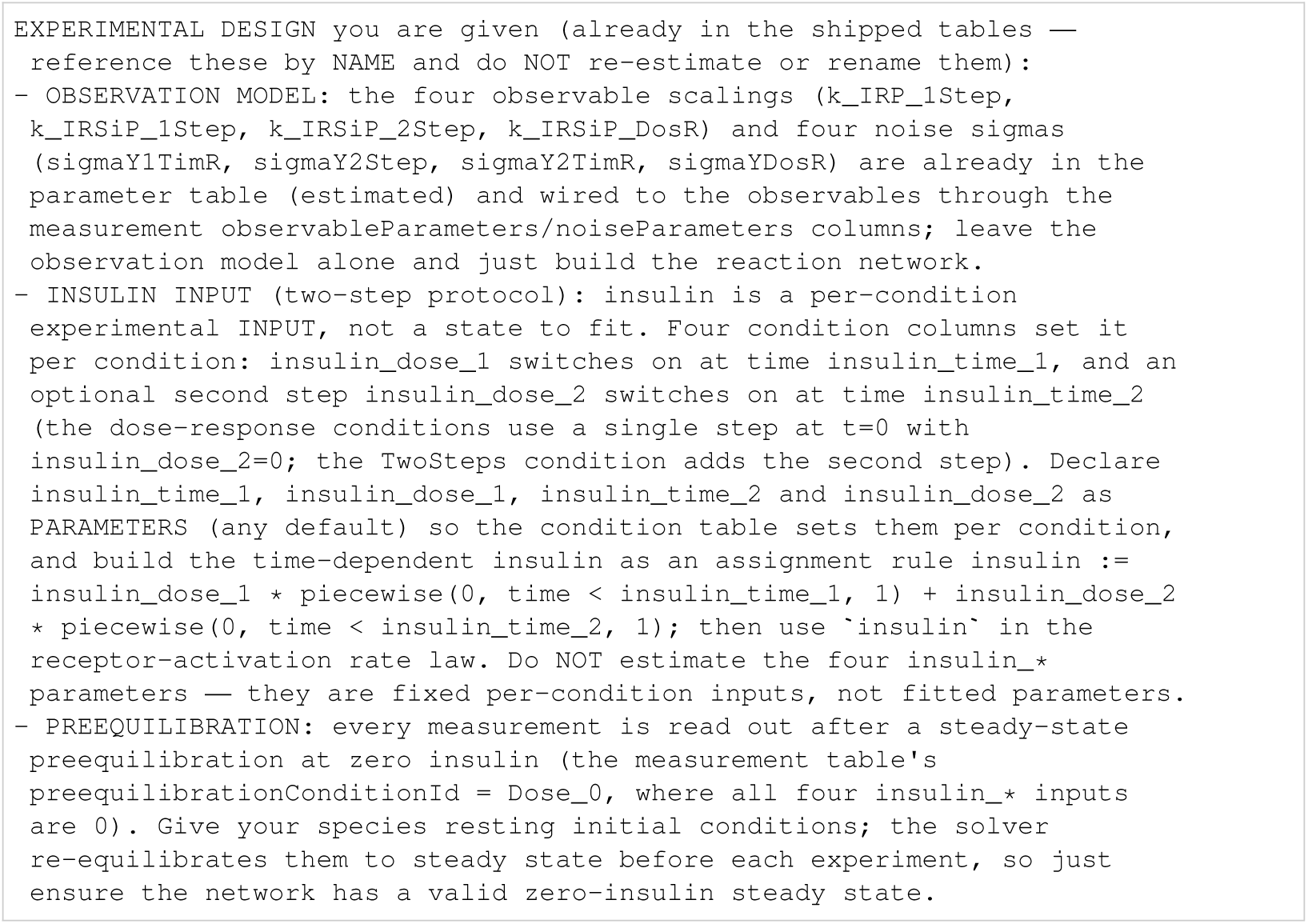
Brannmark: known constants.

**Listing 143:**
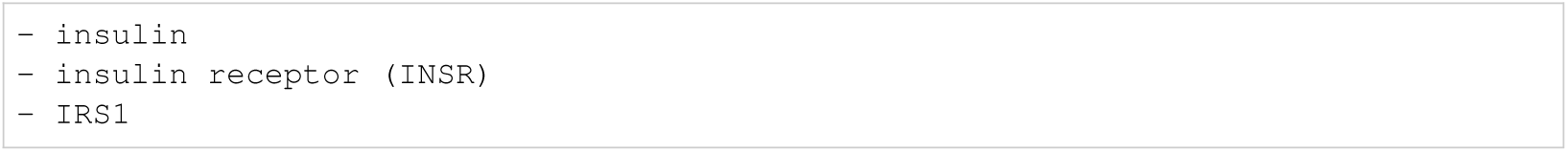
Brannmark: biocurator entities.

#### I.5.7 Bruno

**Listing 144:**
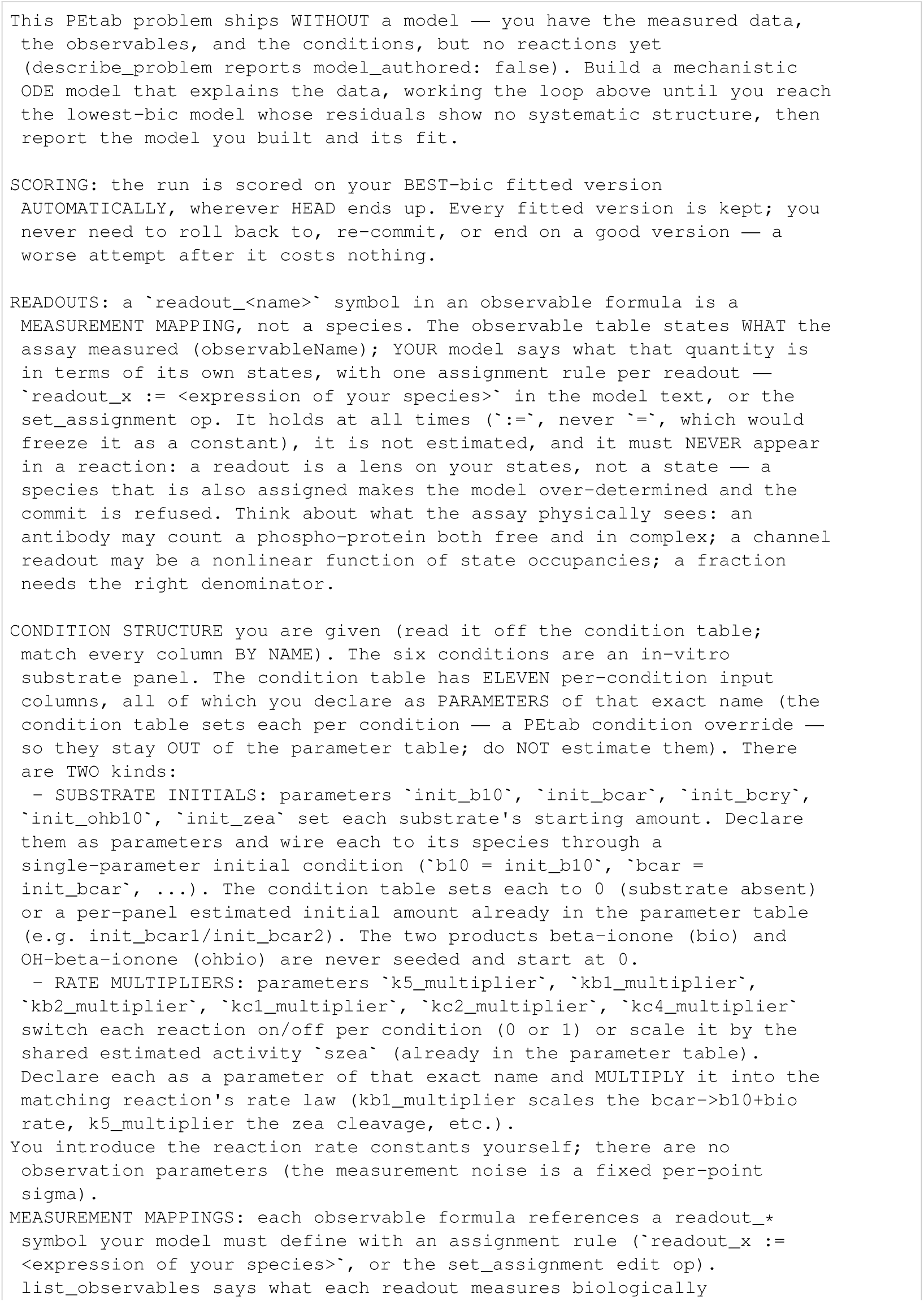
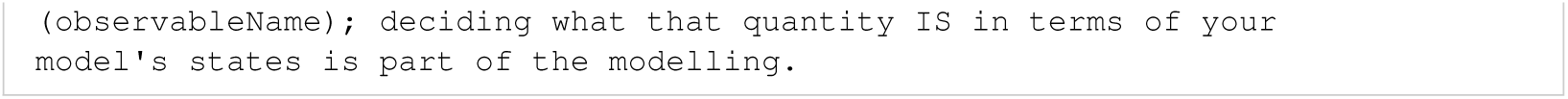
Bruno: initial task.

**Listing 145:**
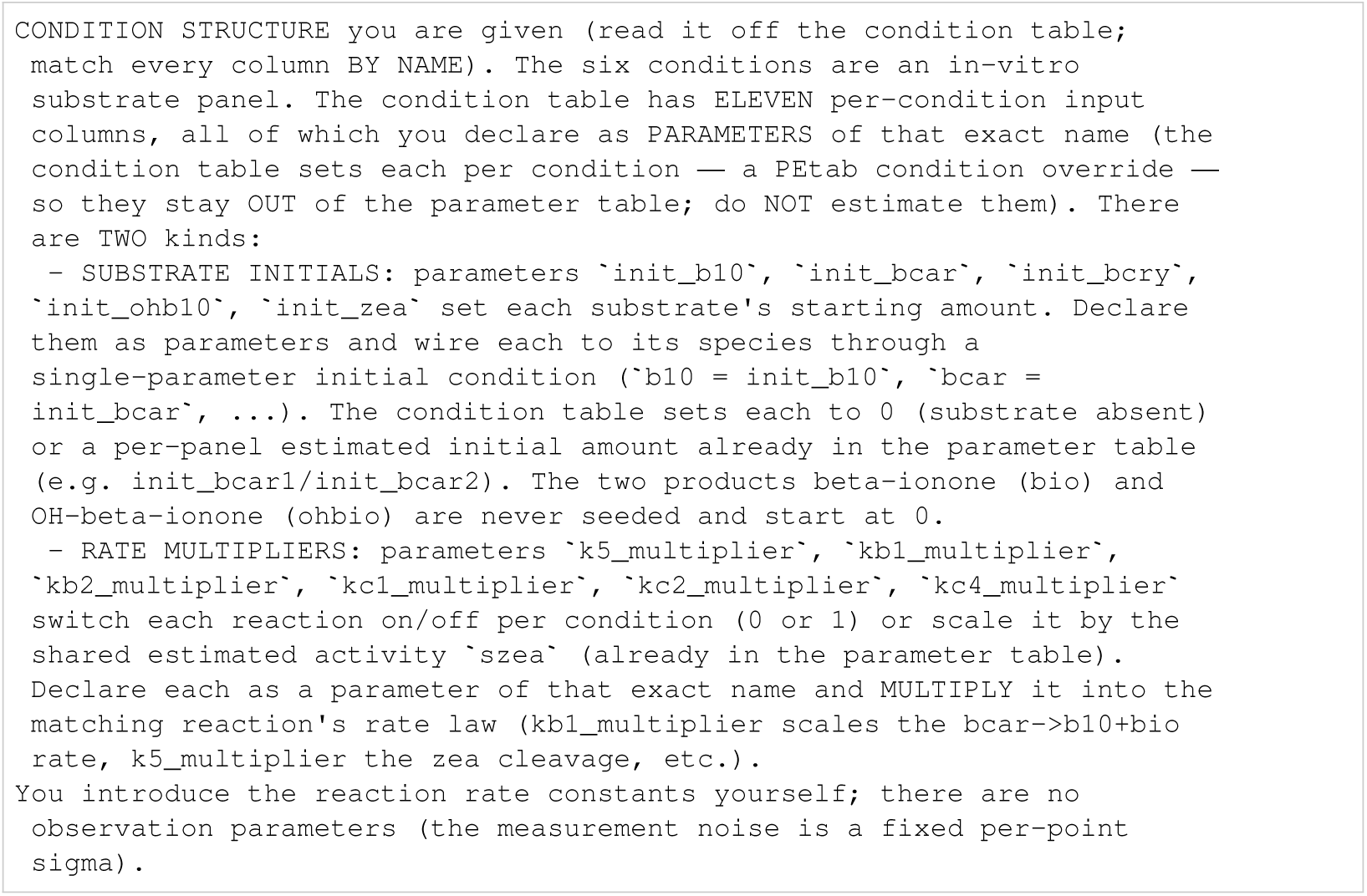
Bruno: known constants.

**Listing 146:**
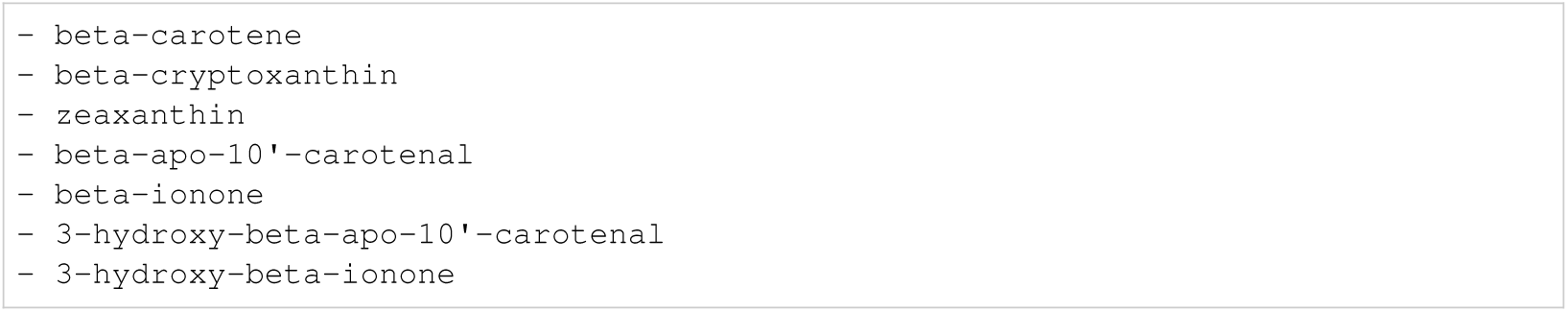
Bruno: biocurator entities.

#### I.5.8 Crauste

**Listing 147:**
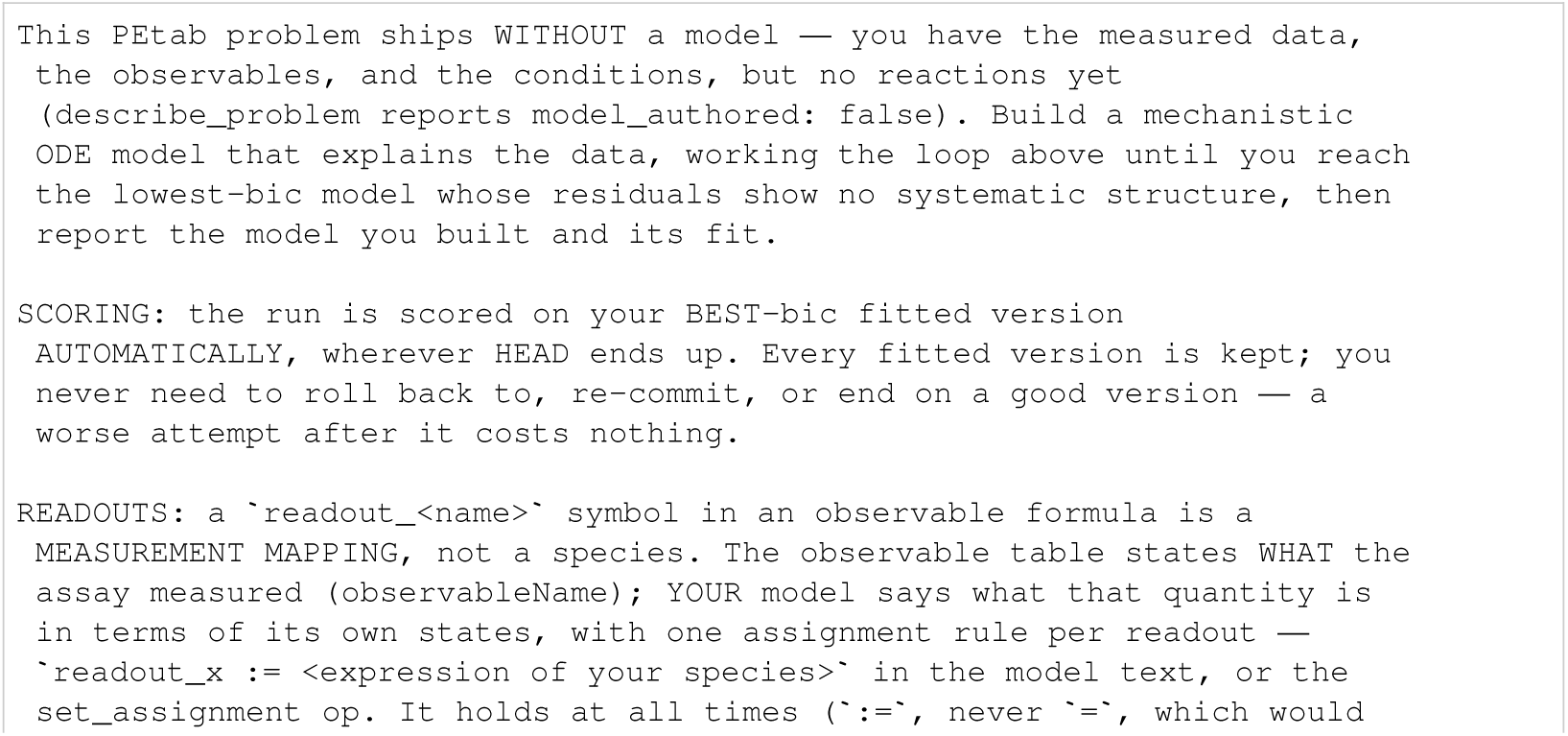
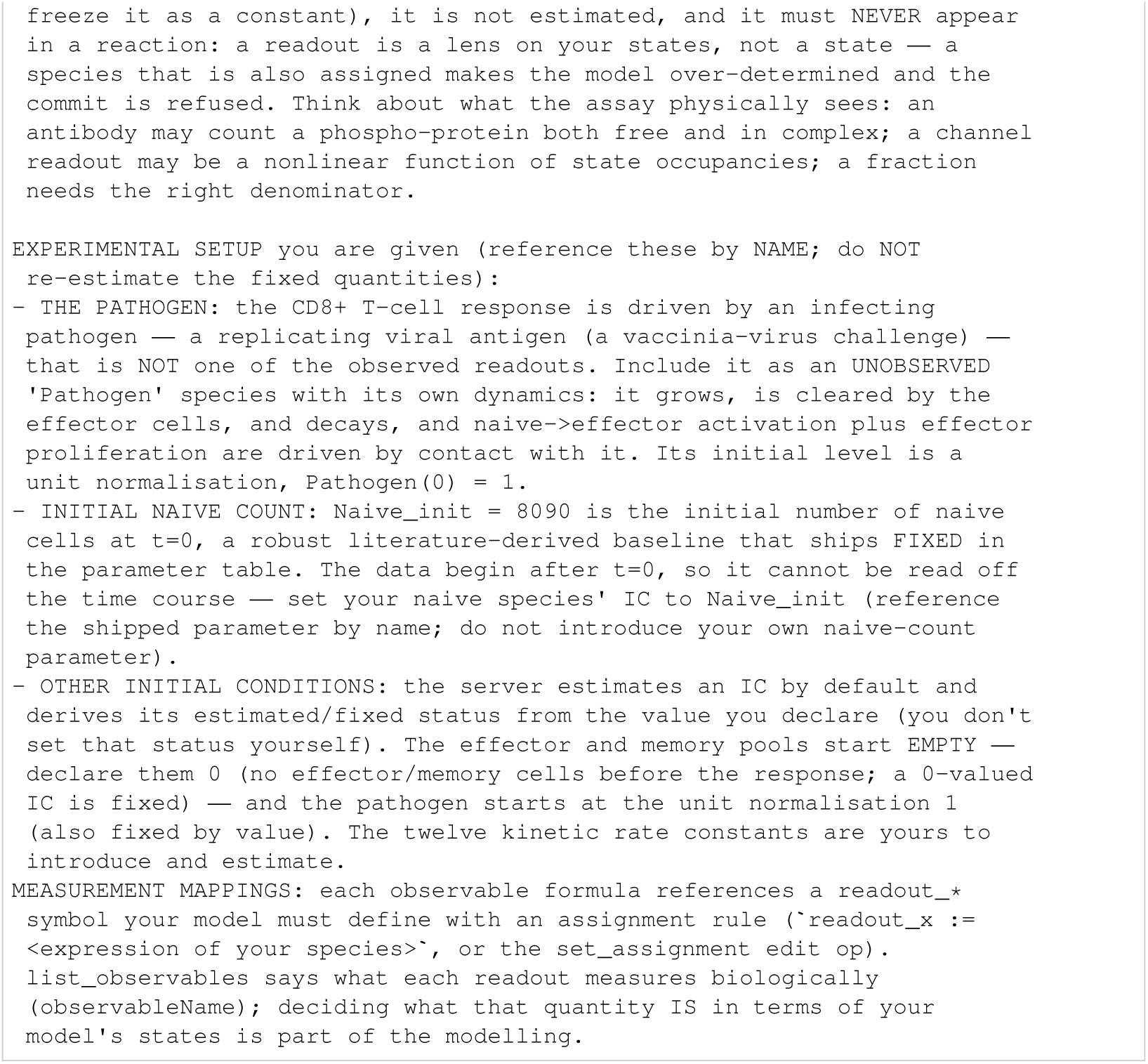
Crauste: initial task.

**Listing 148:**
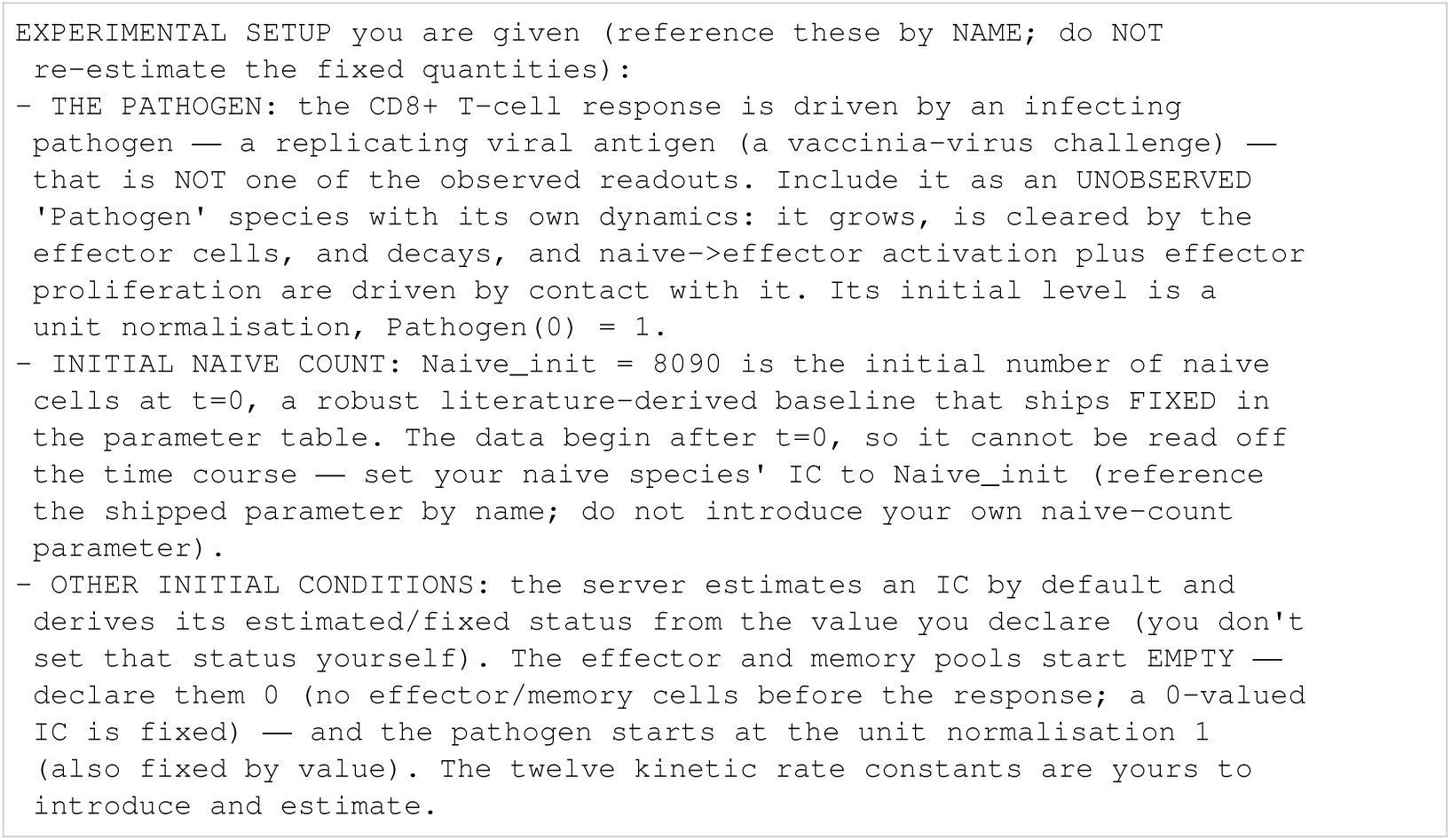
Crauste: known constants.

**Listing 149:**
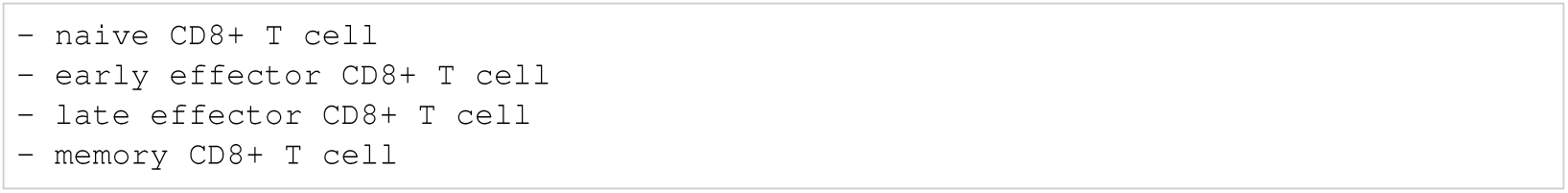
Crauste: biocurator entities.

#### I.5.9 Fiedler

**Listing 150:**
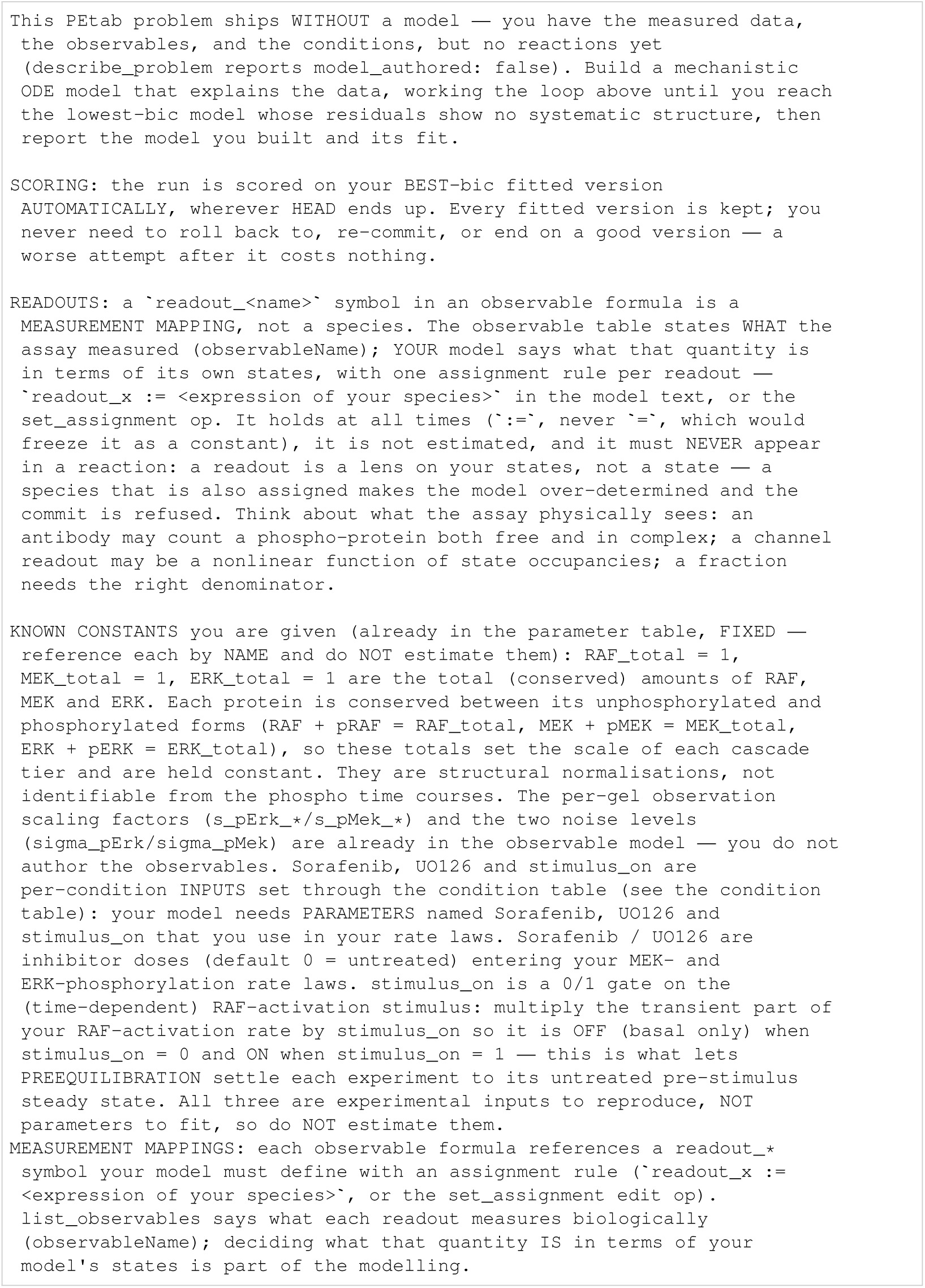
Fiedler: initial task.

**Listing 151:**
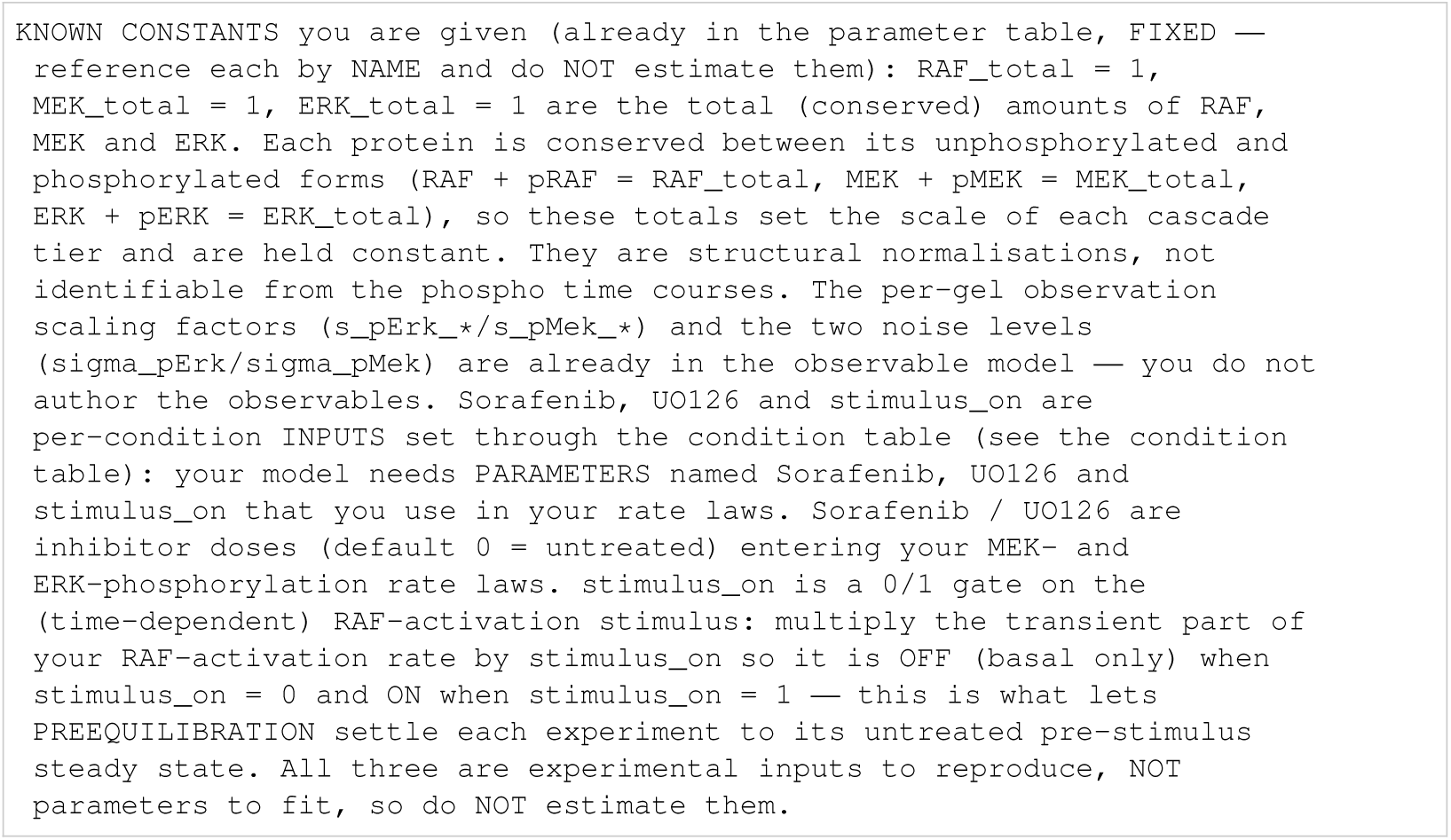
Fiedler: known constants.

**Listing 152:**
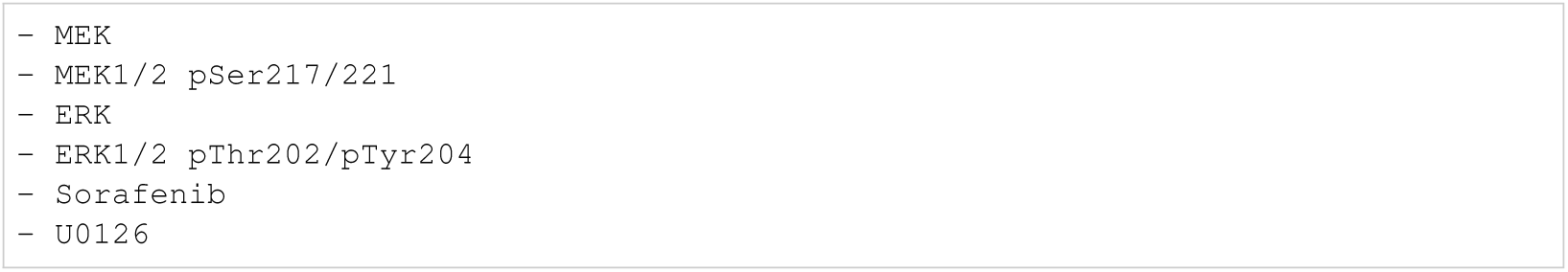
Fiedler: biocurator entities.

#### I.5.10 Fujita

**Listing 153:**
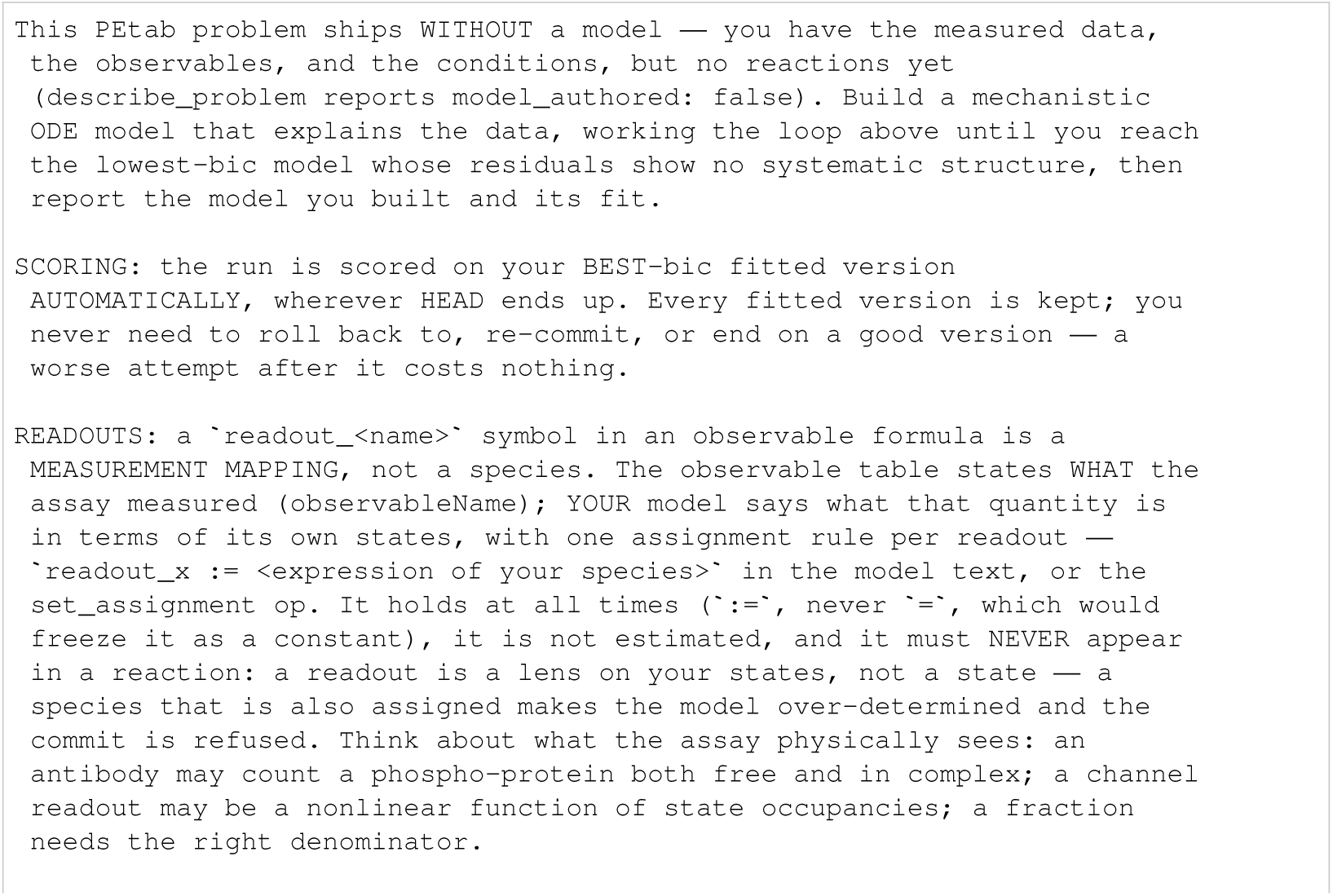
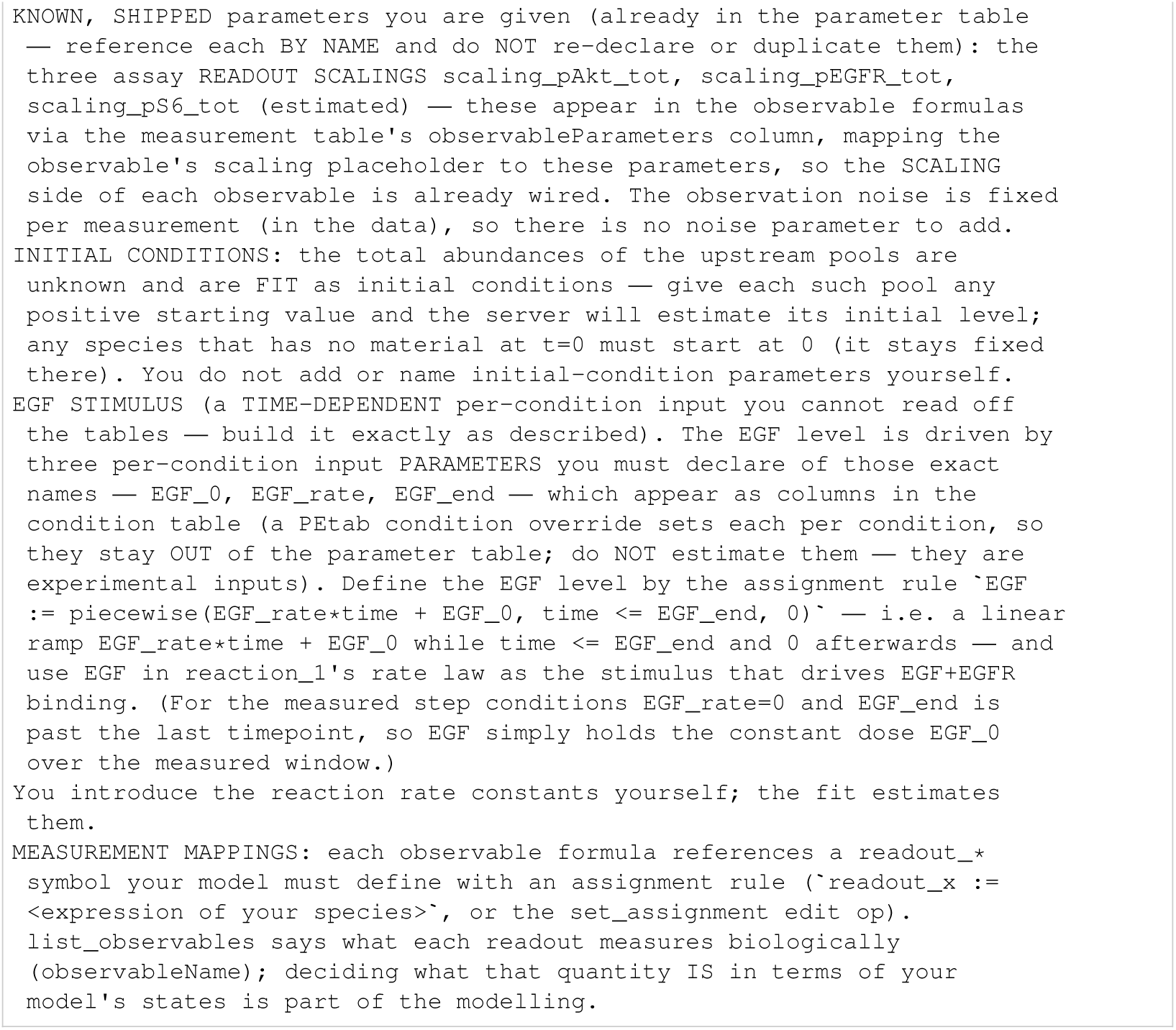
Fujita: initial task.

**Listing 154:**
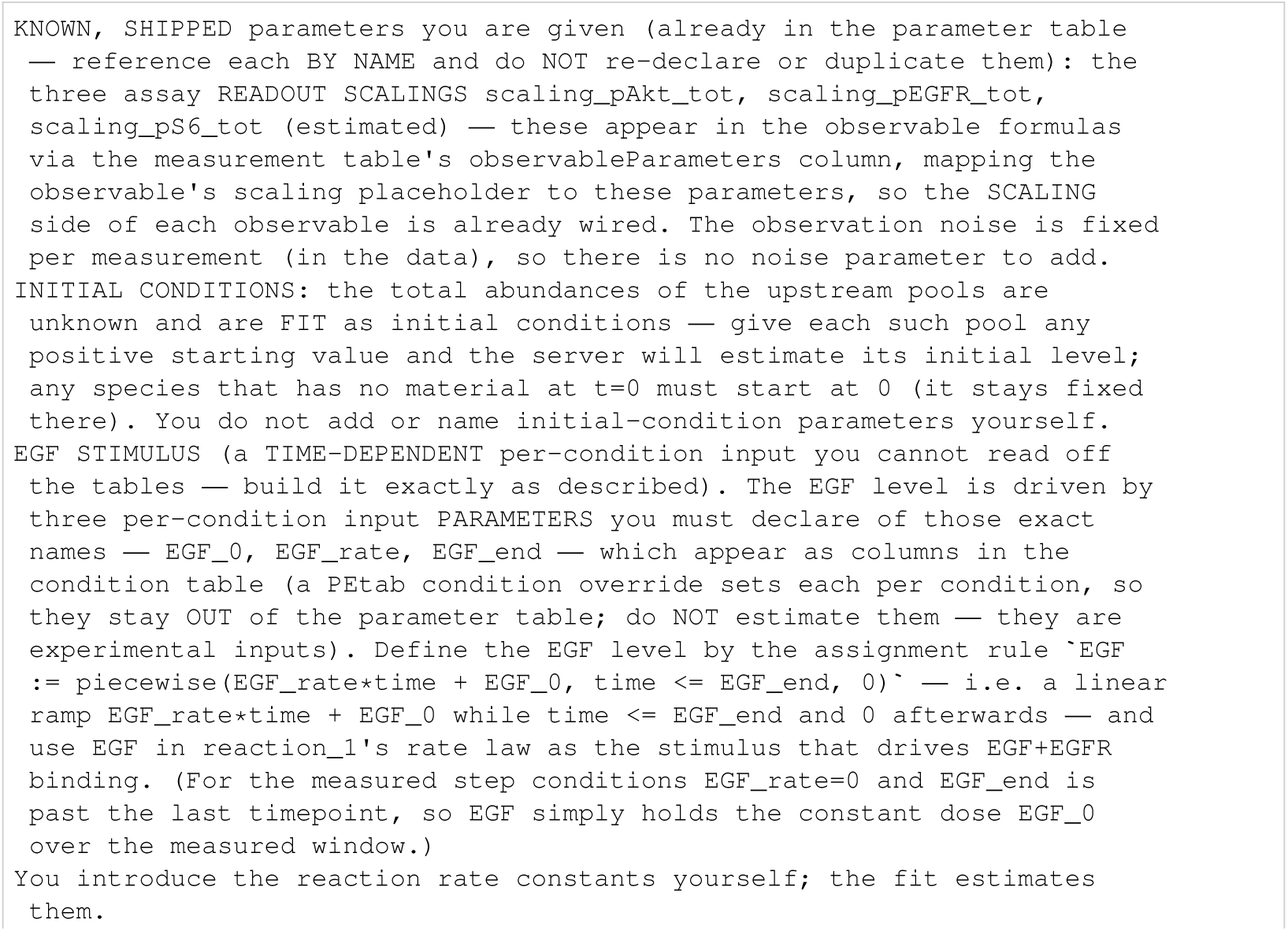
Fujita: known constants.

**Listing 155:**
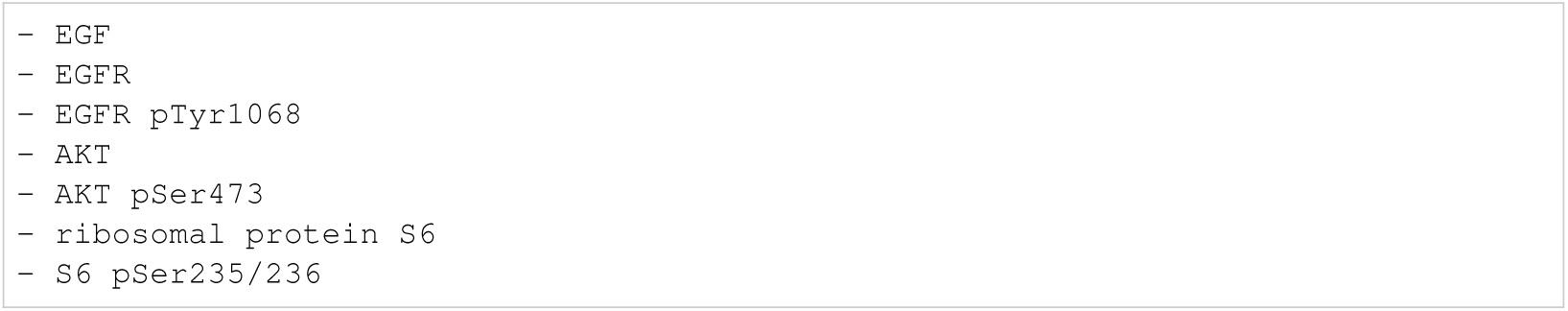
Fujita: biocurator entities.

#### I.5.11 Isensee

**Listing 156:**
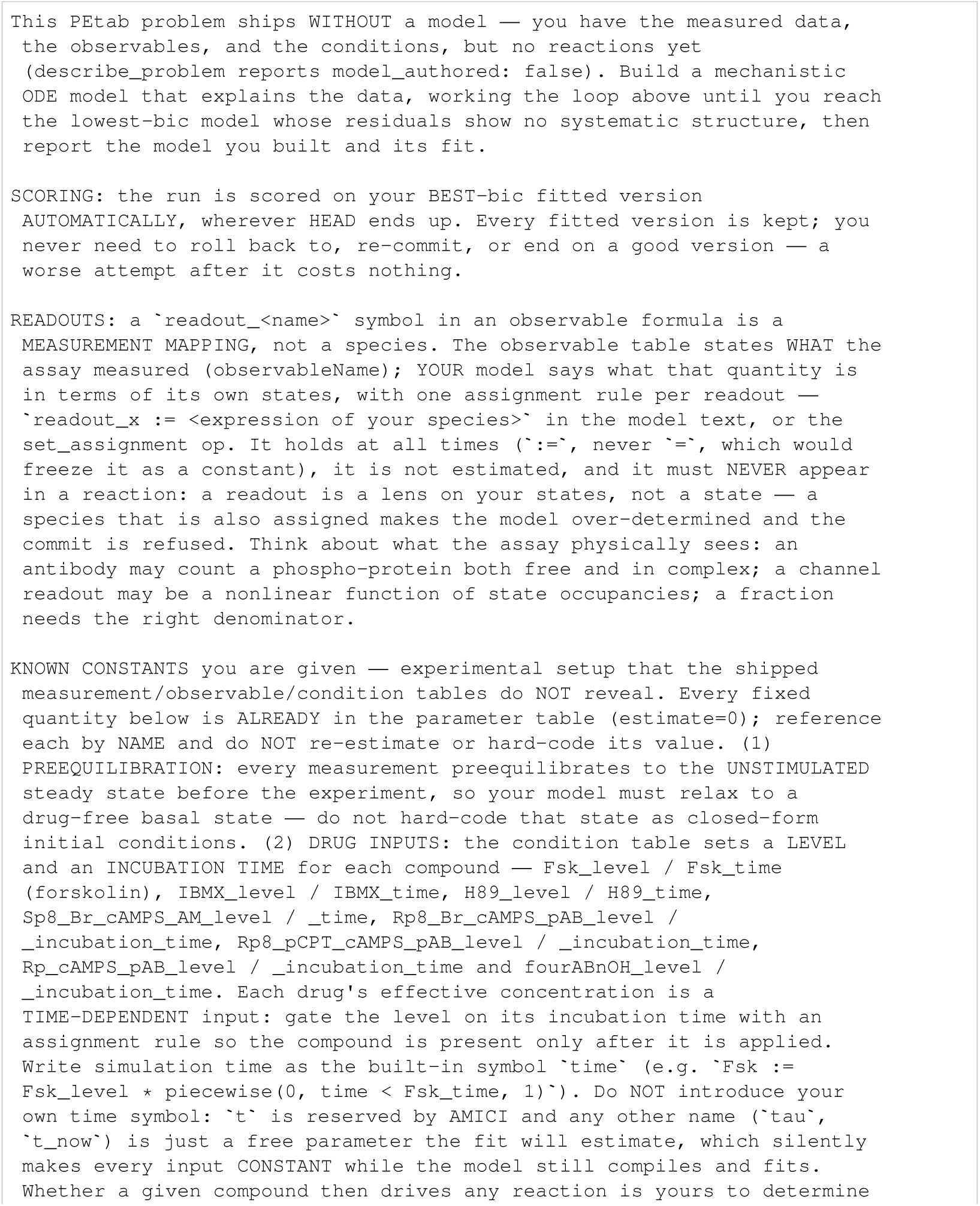
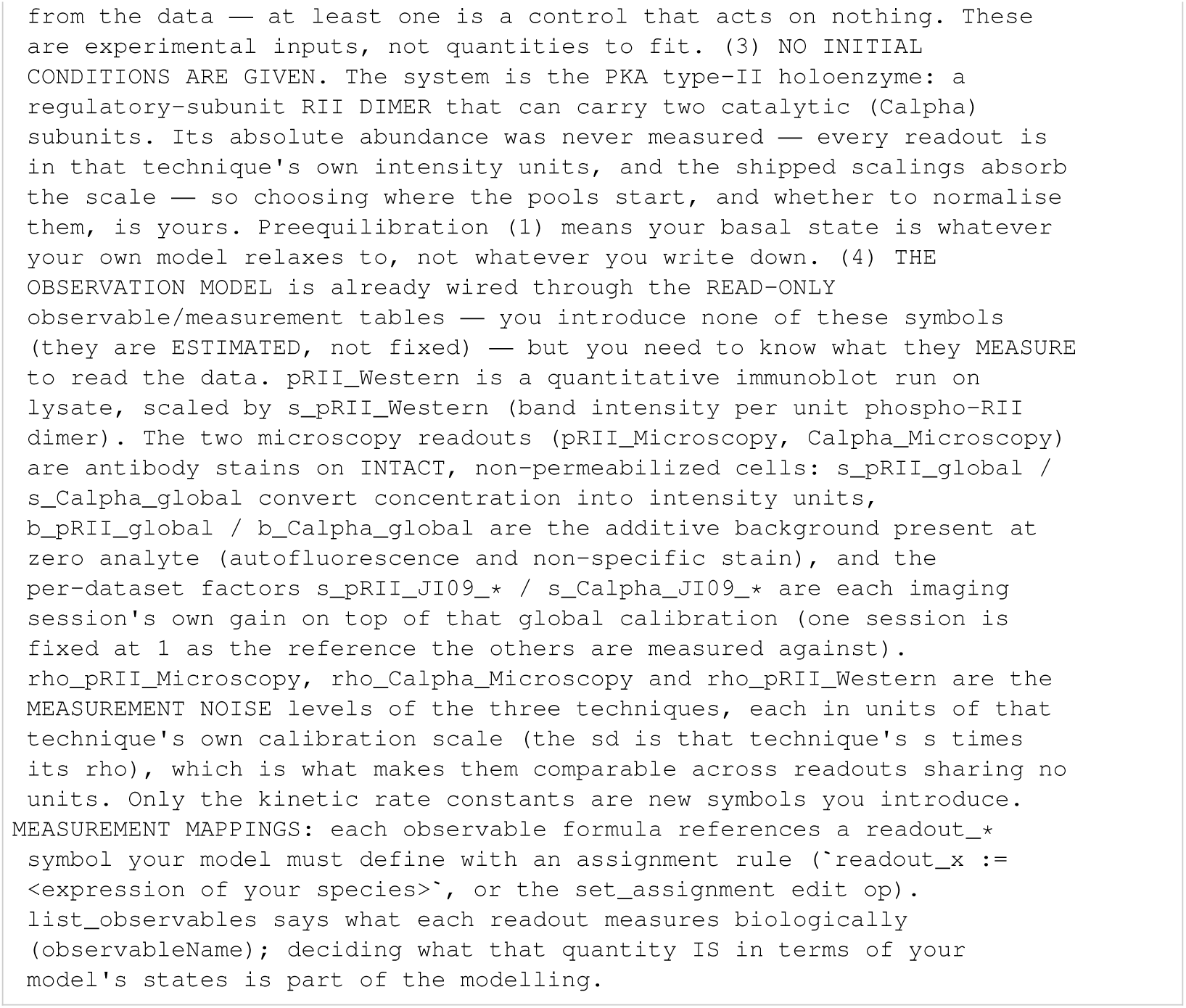
Isensee: initial task.

**Listing 157:**
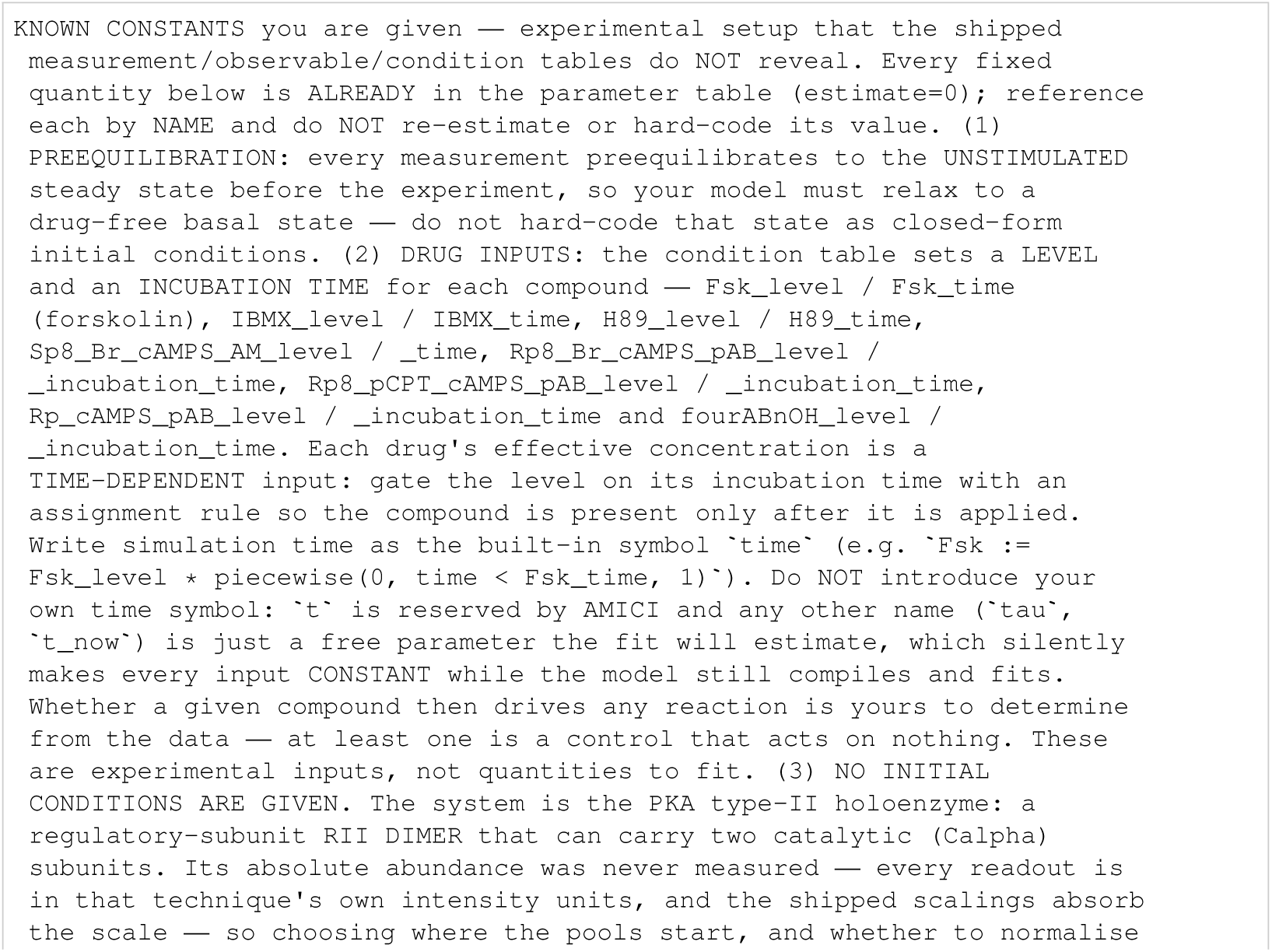
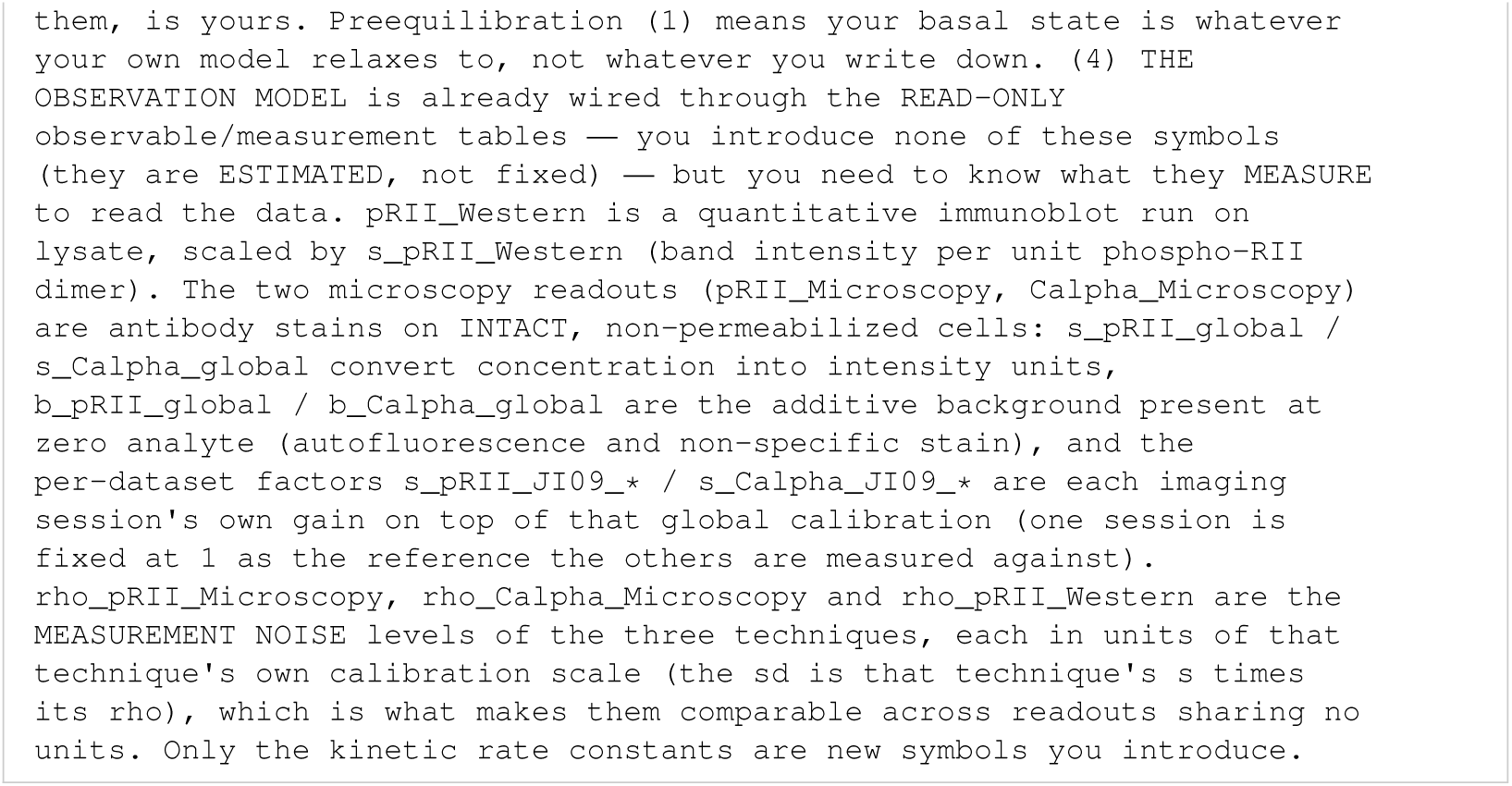
Isensee: known constants.

**Listing 158:**
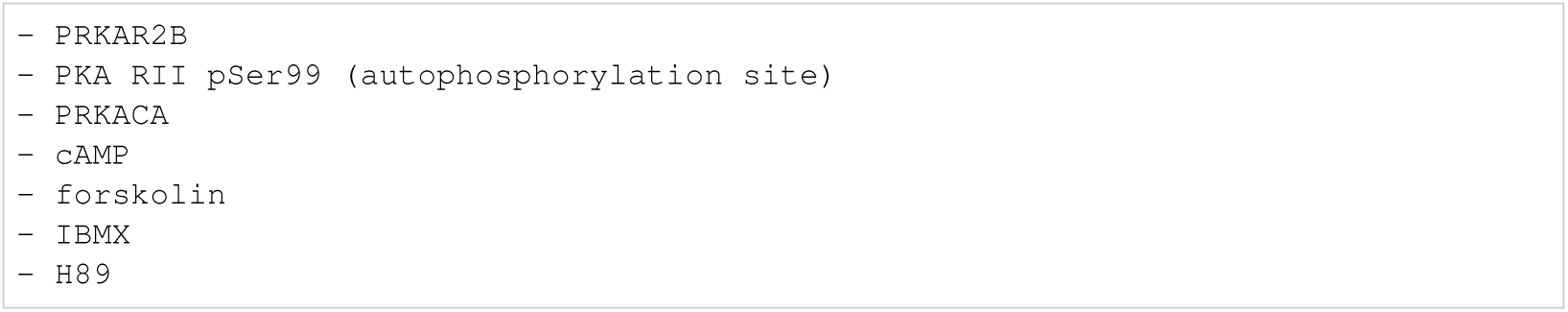
Isensee: biocurator entities.

#### I.5.12 Lucarelli

**Listing 159:**
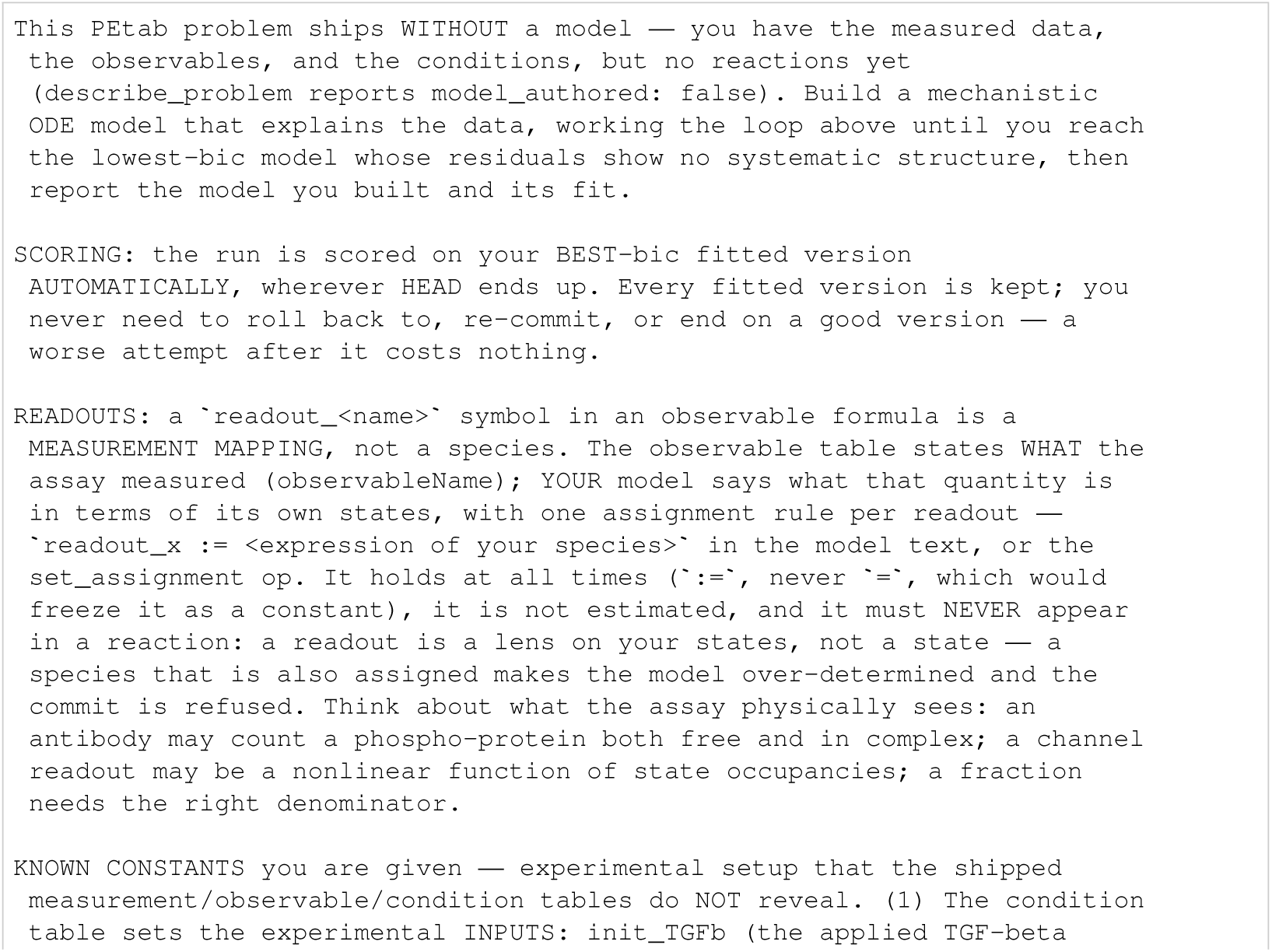
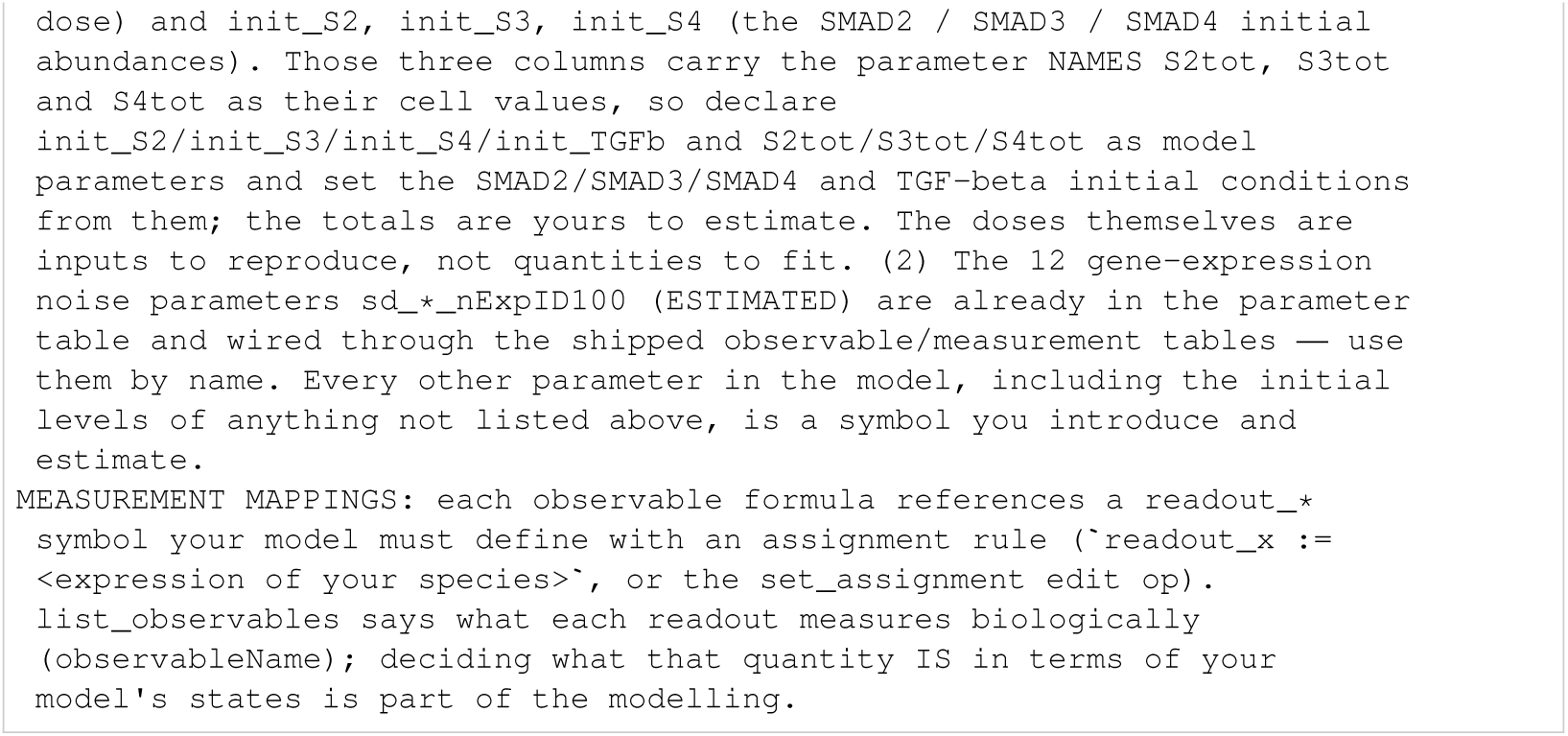
Lucarelli: initial task.

**Listing 160:**
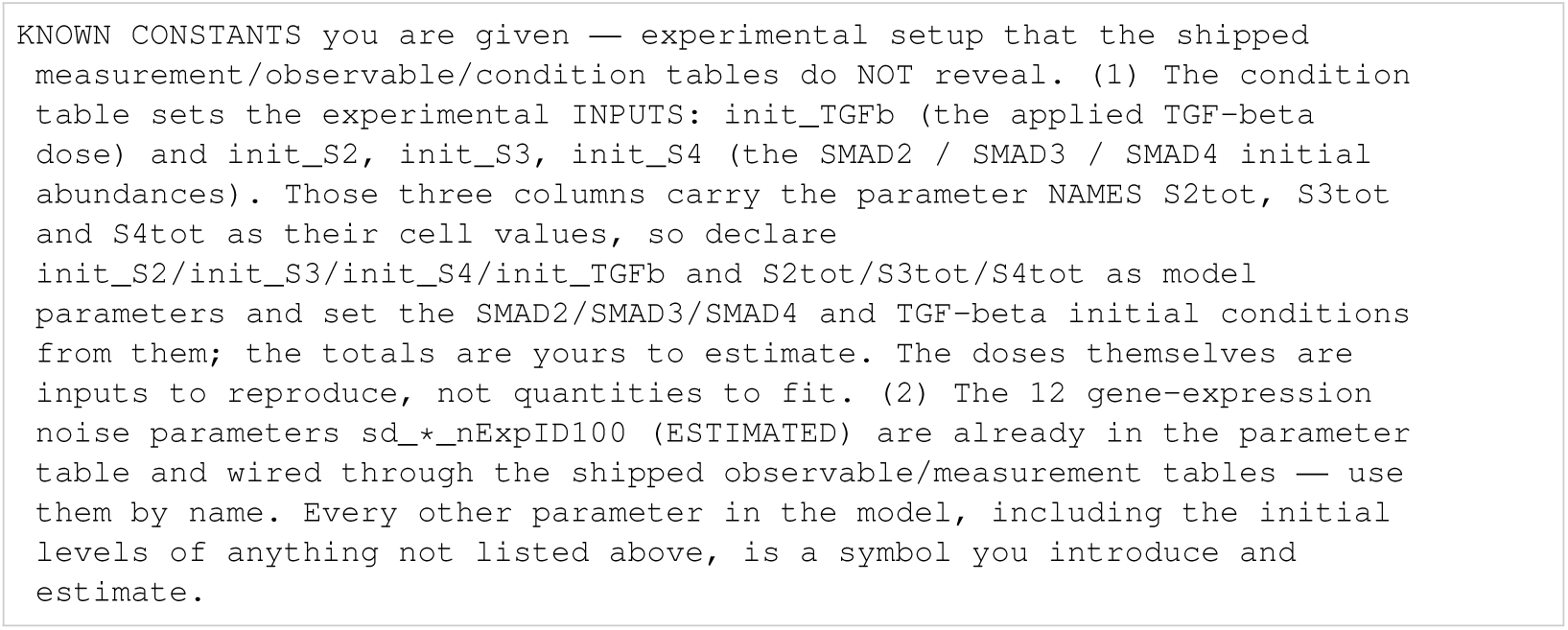
Lucarelli: known constants.

**Listing 161:**
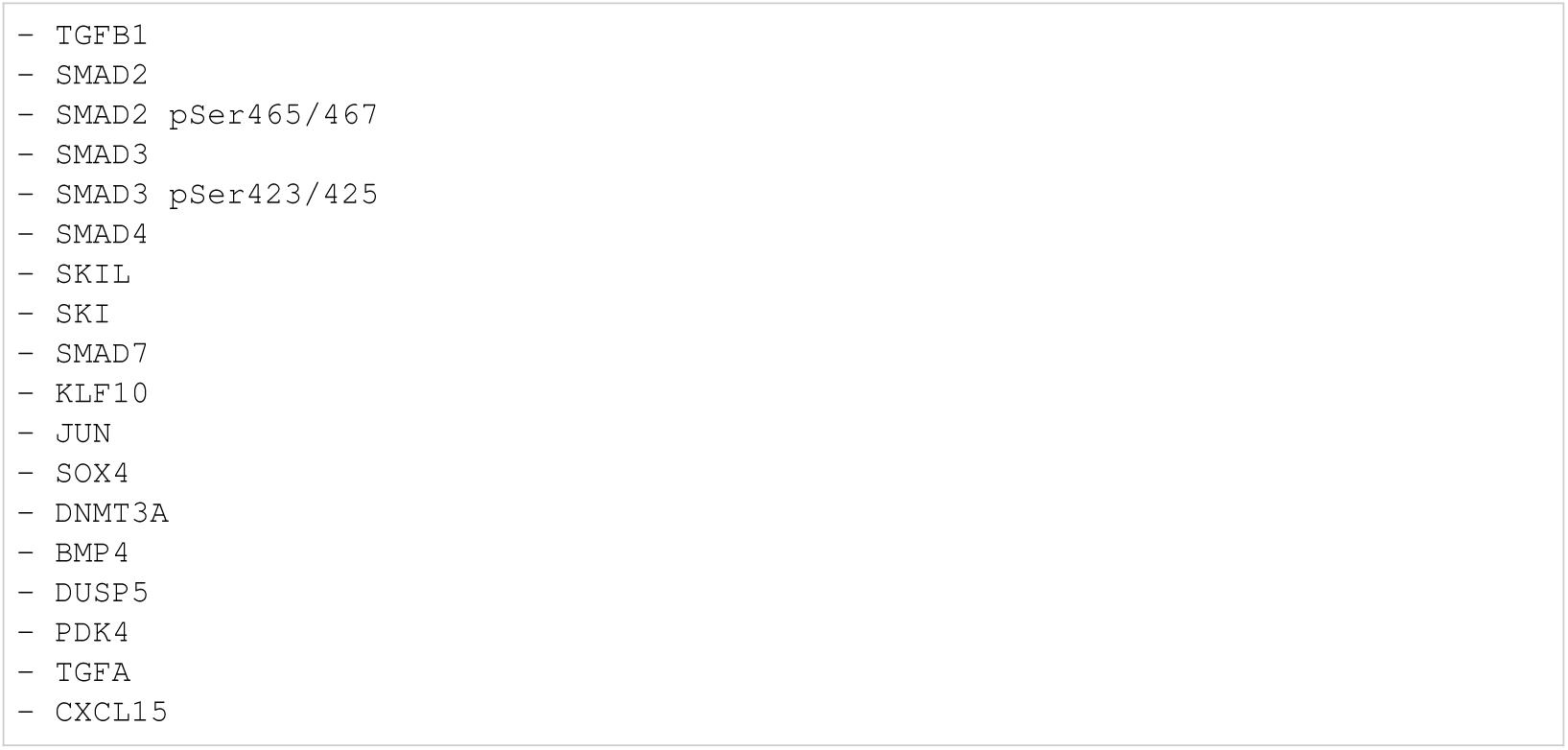
Lucarelli: biocurator entities.

#### I.5.13 Oliveira

**Listing 162:**
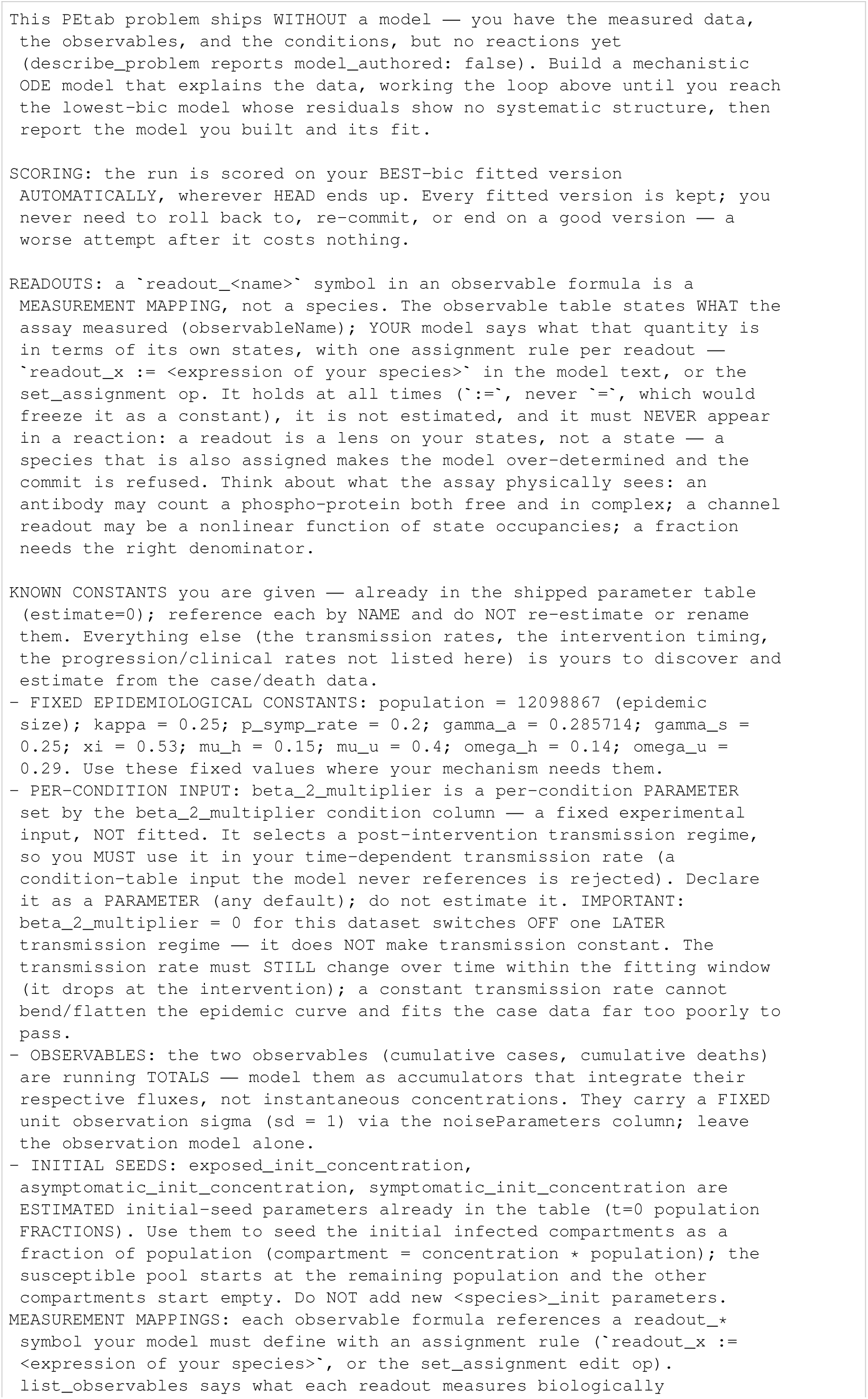
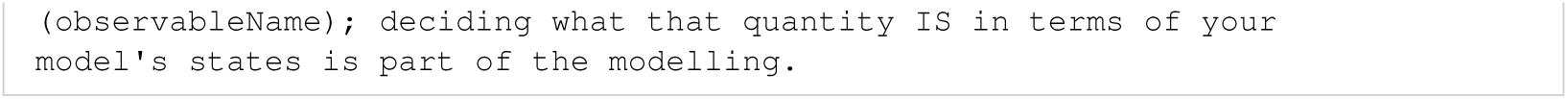
Oliveira: initial task.

**Listing 163:**
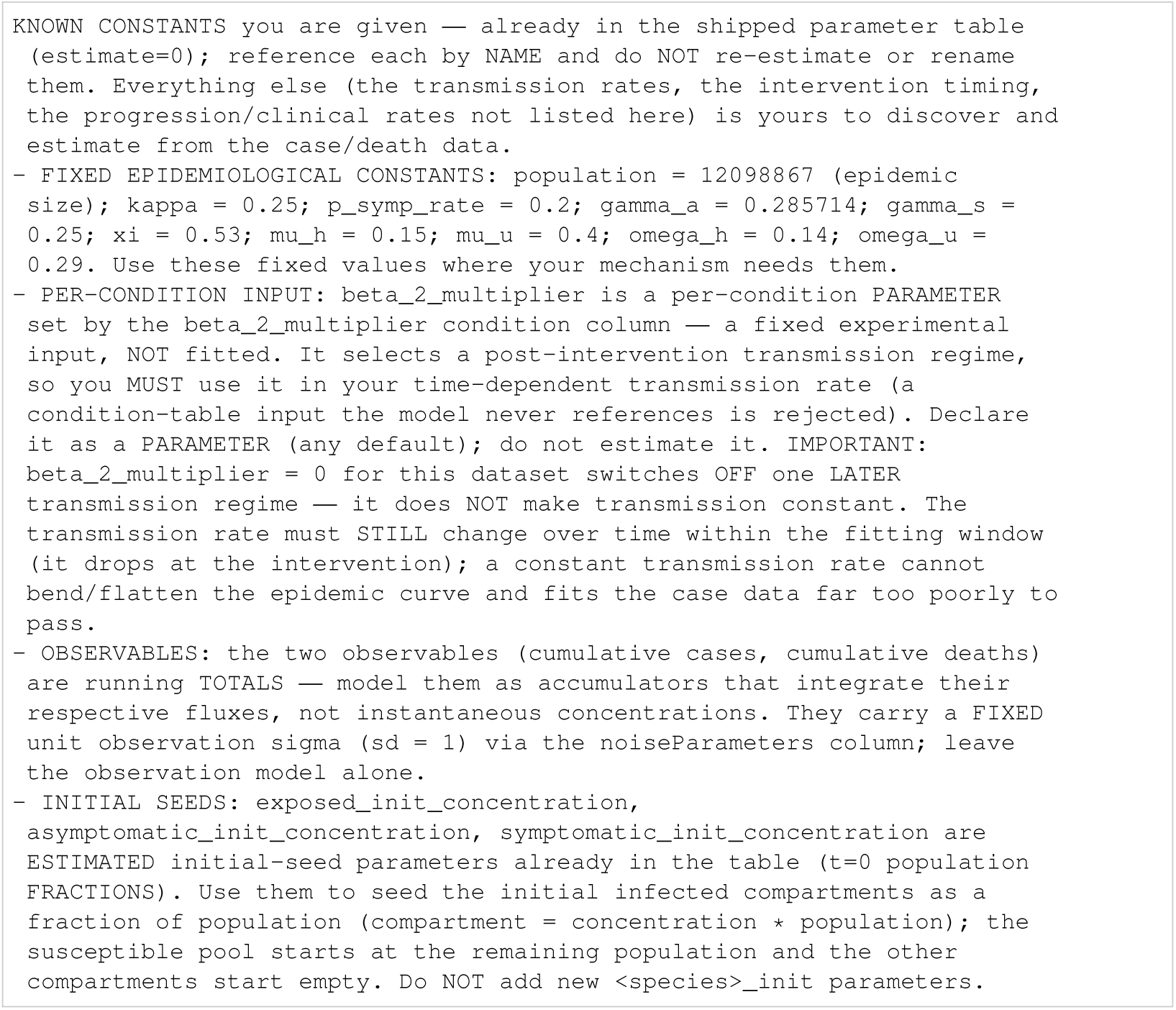
Oliveira: known constants.

**Listing 164:**
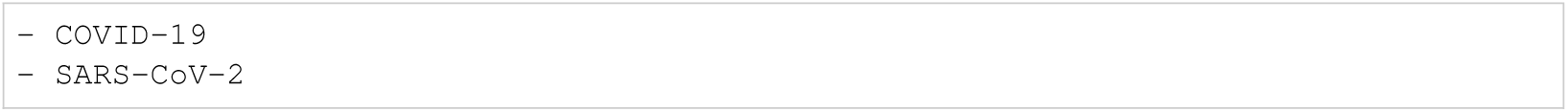
Oliveira: biocurator entities.

#### I.5.14 Perelson

**Listing 165:**
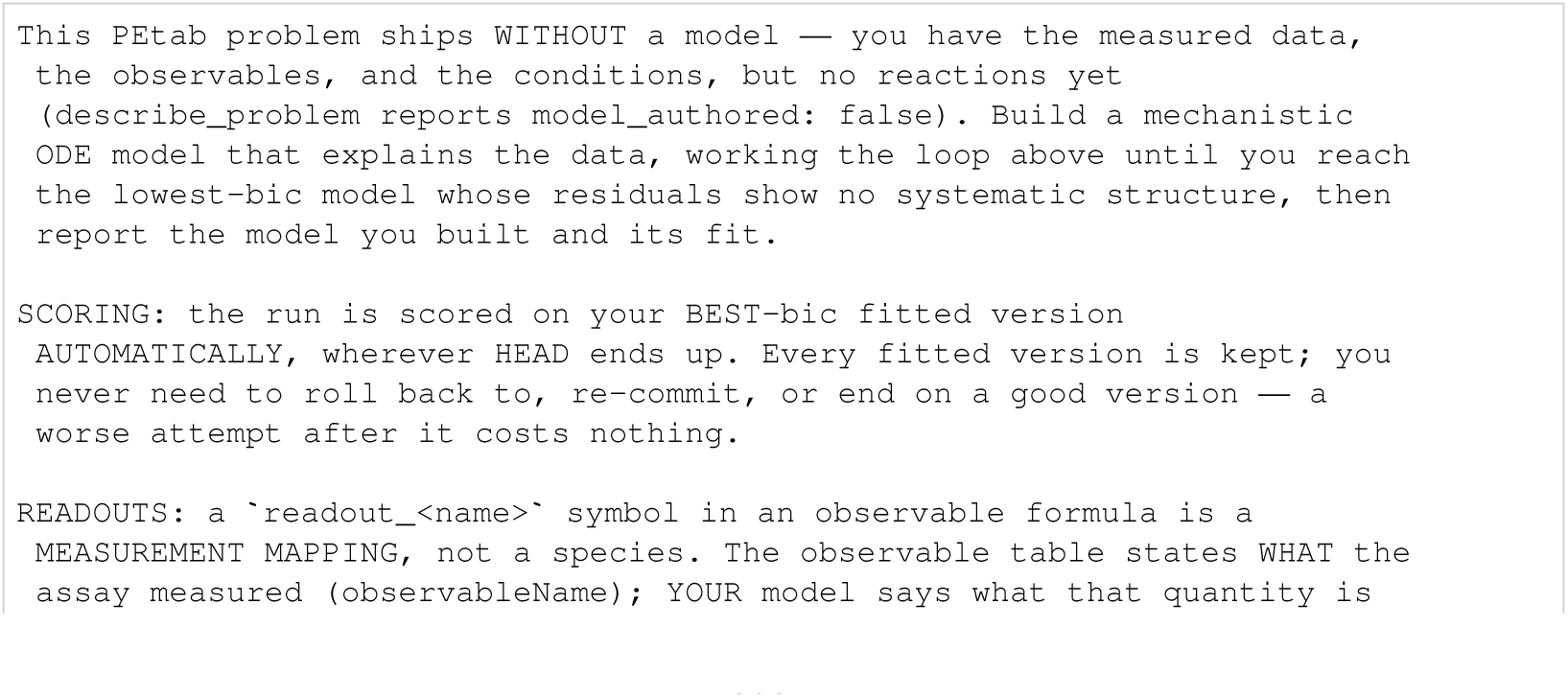
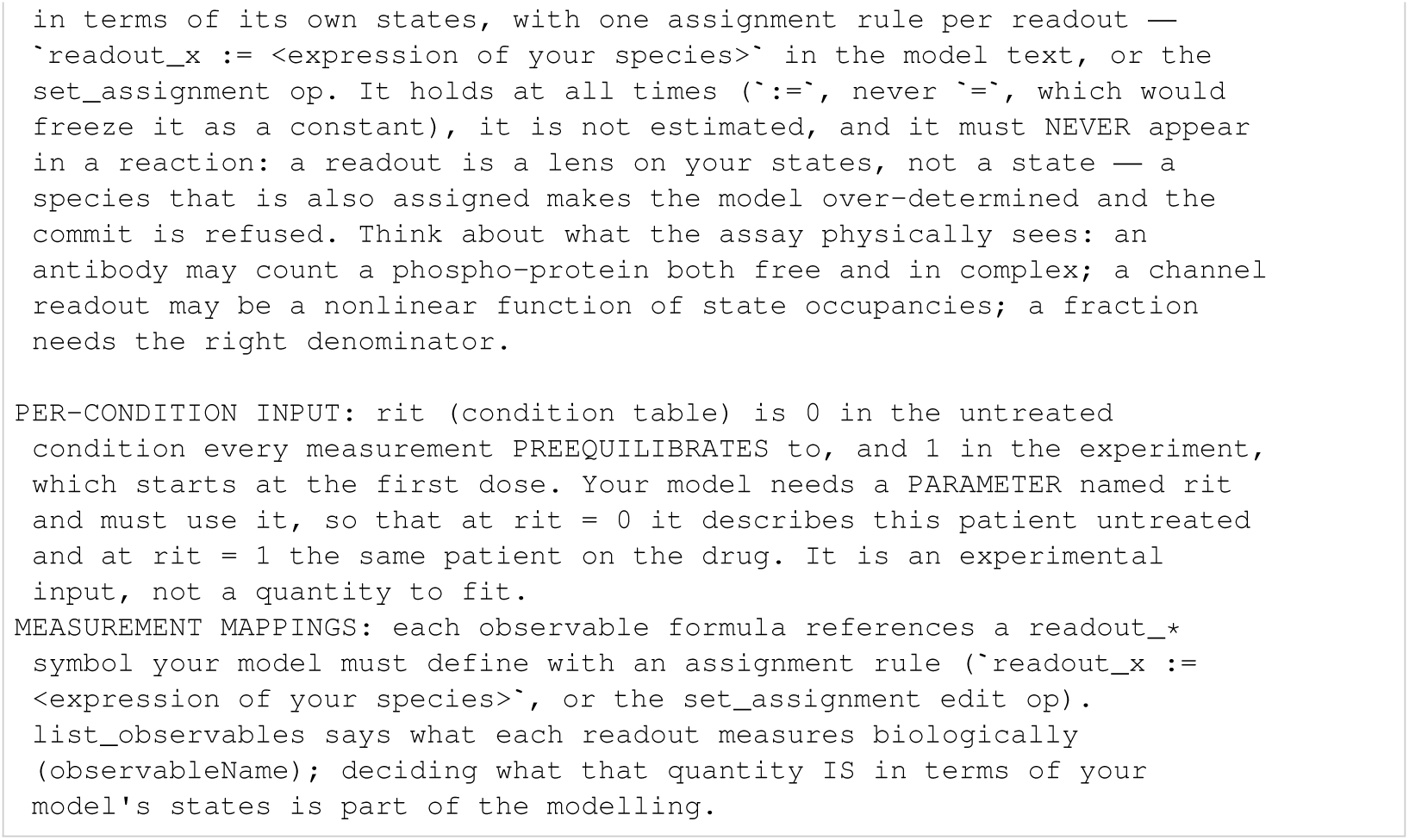
Perelson: initial task.

**Listing 166:**
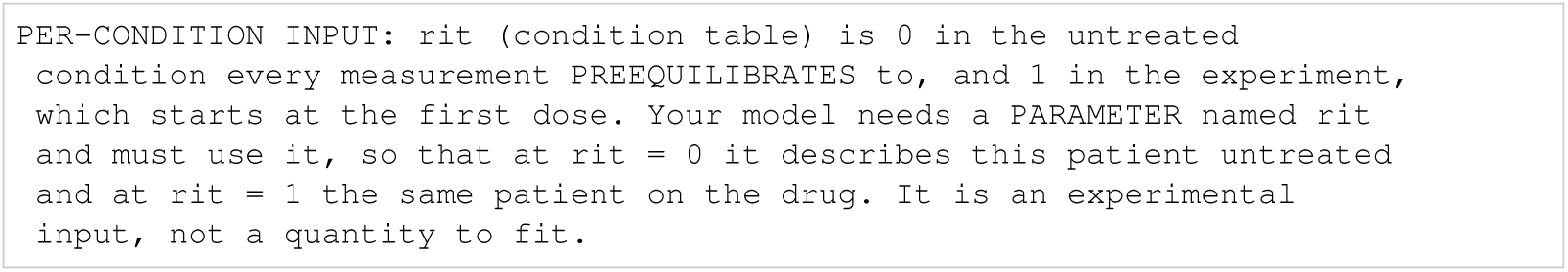
Perelson: known constants.

**Listing 167:**
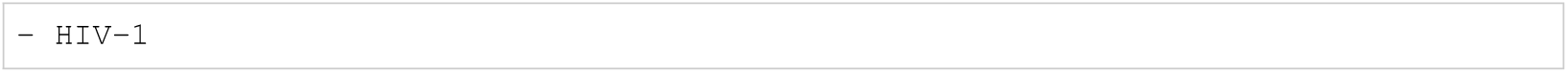
Perelson: biocurator entities.

#### I.5.15 Rahman

**Listing 168:**
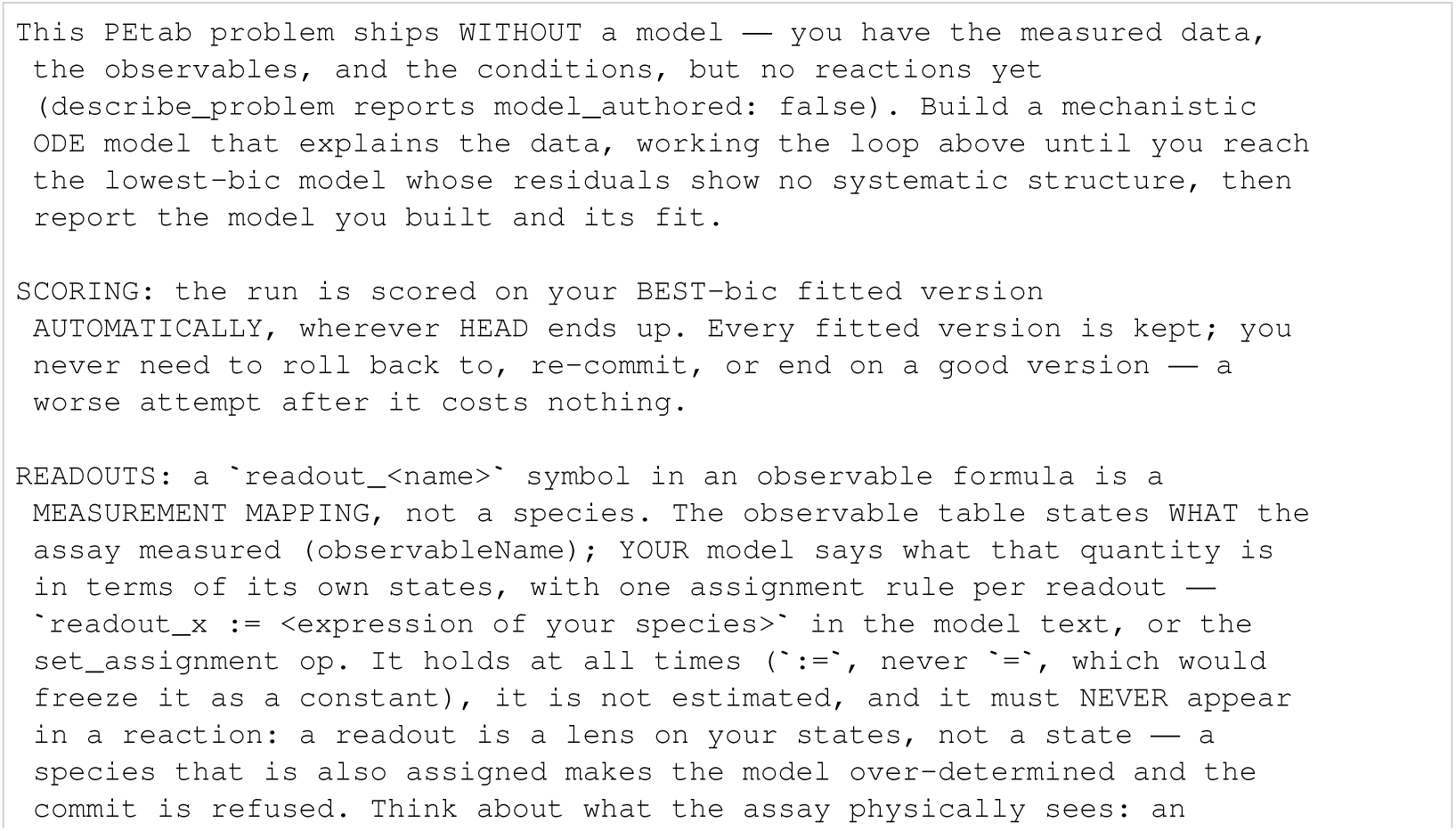
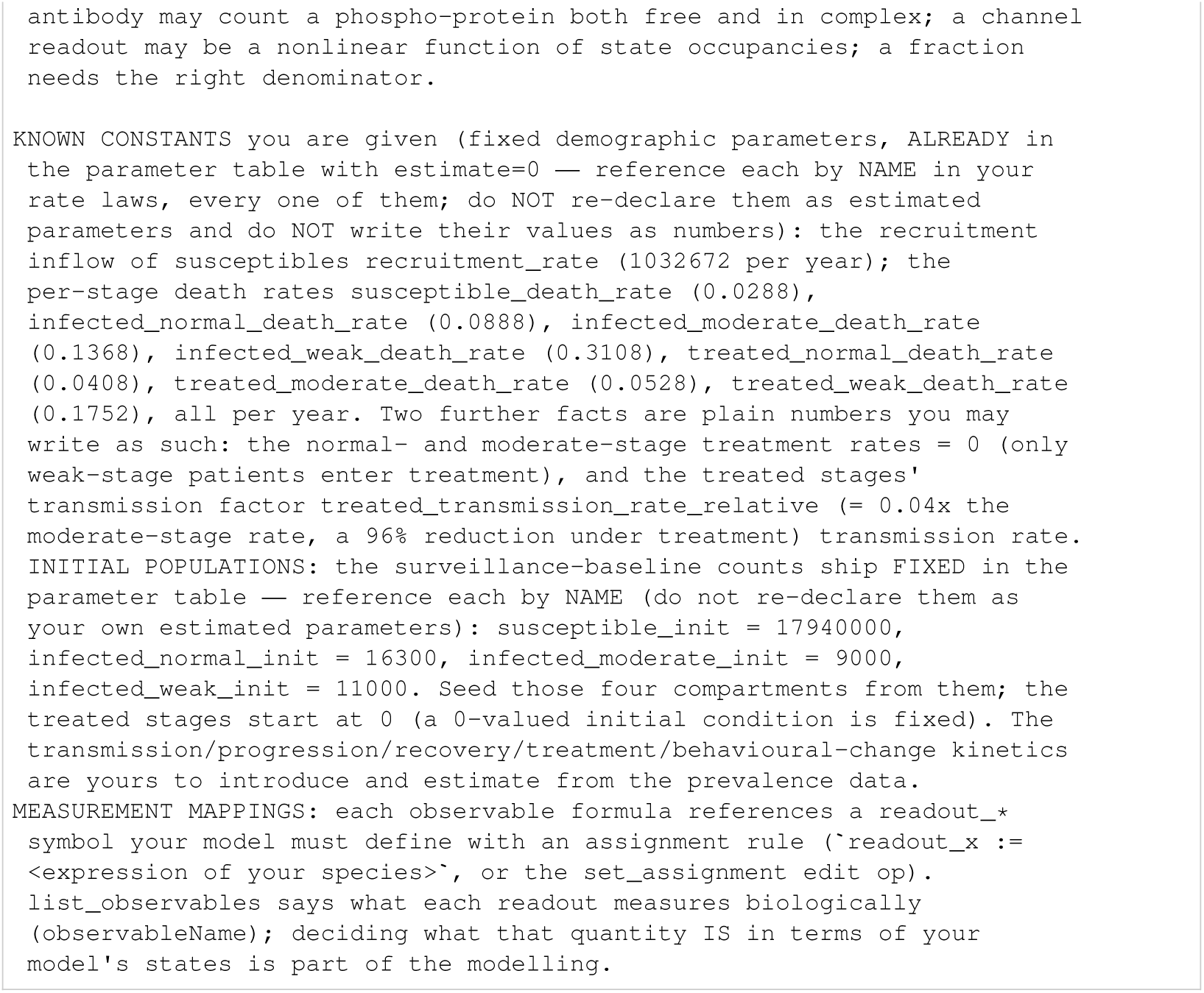
Rahman: initial task.

**Listing 169:**
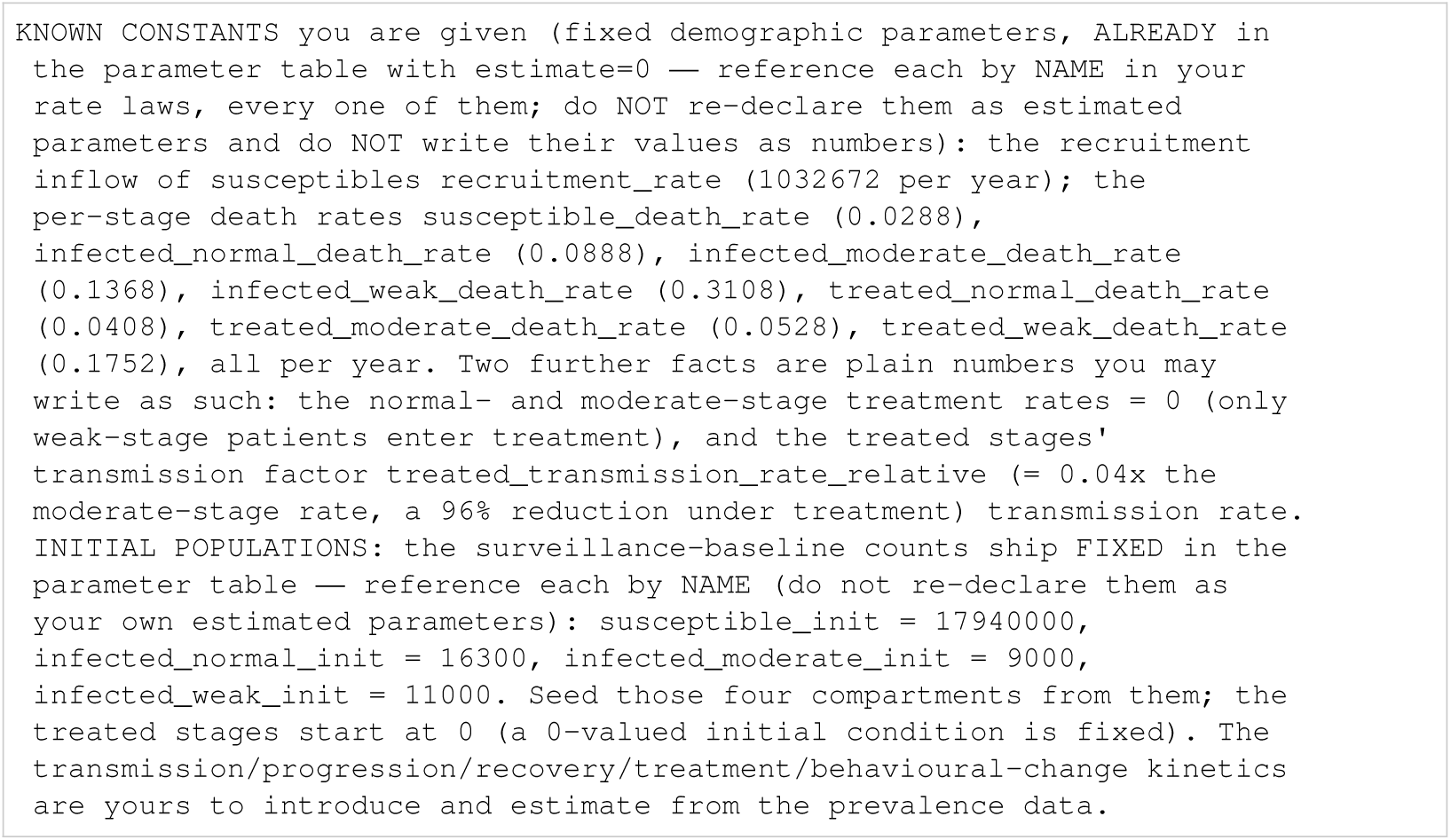
Rahman: known constants.

#### I.5.16 Raimundez

**Listing 170:**
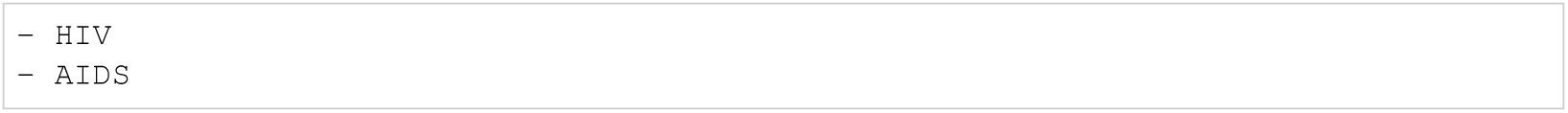
Rahman: biocurator entities.

**Listing 171:**
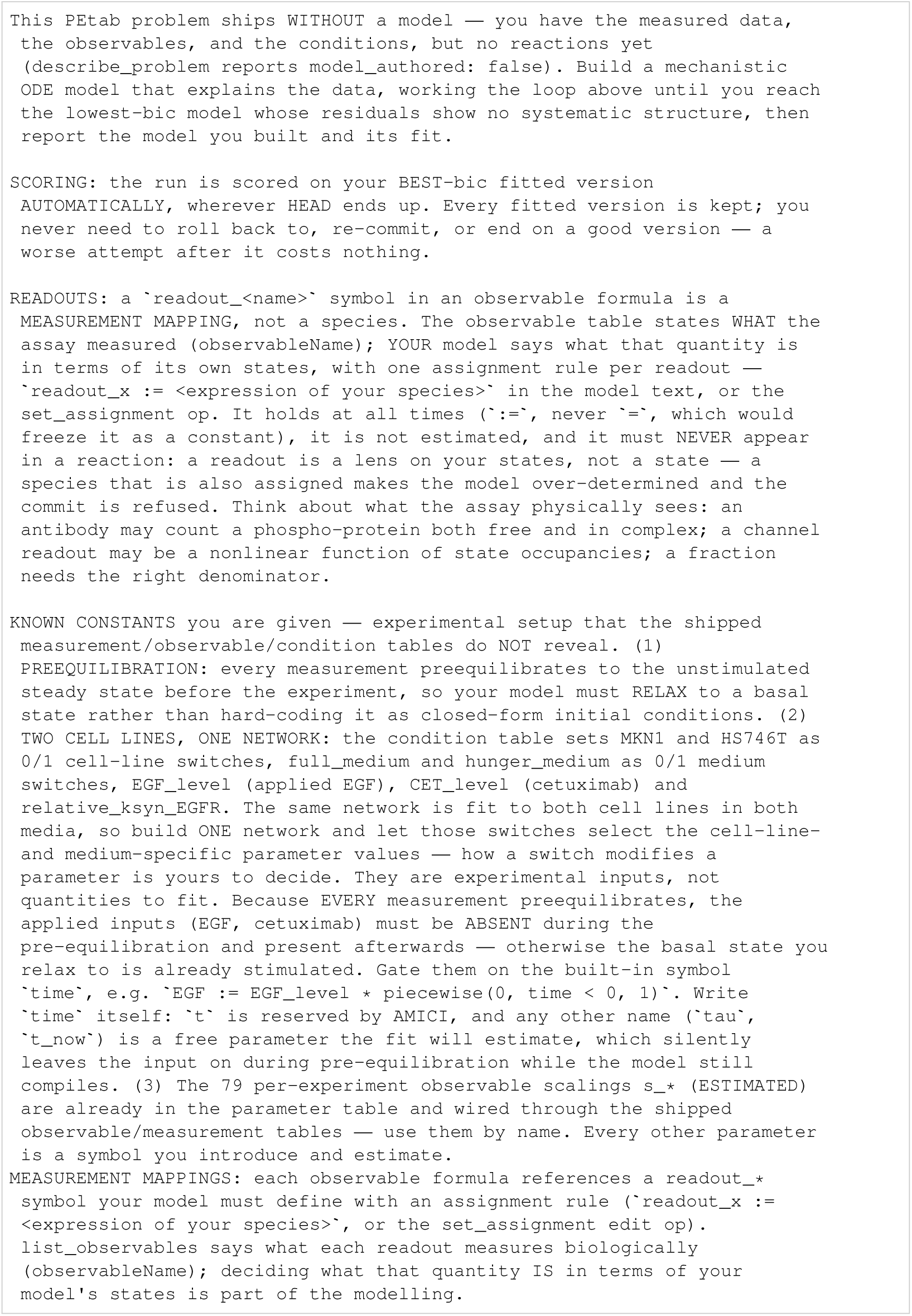
Raimundez: initial task.

**Listing 172:**
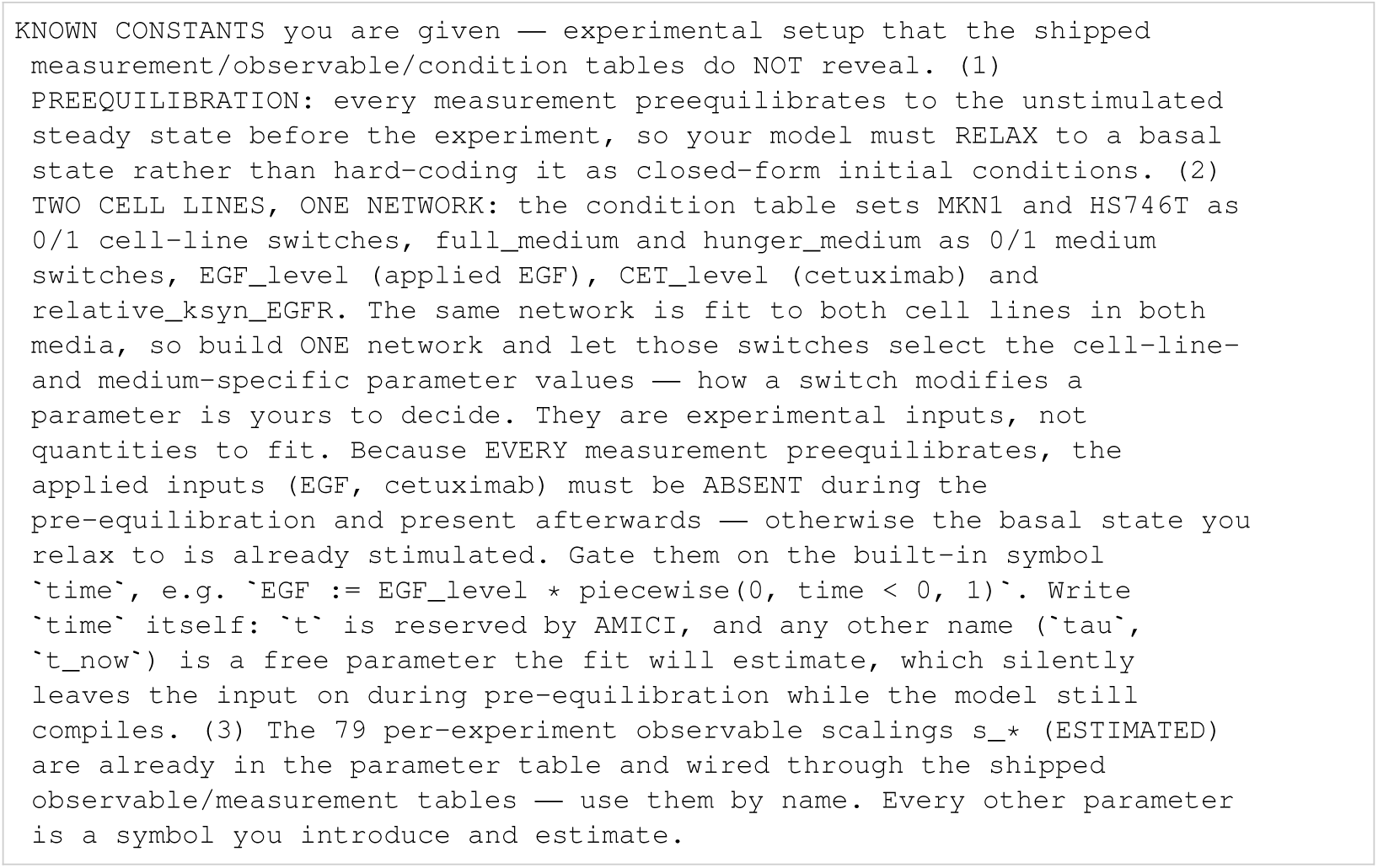
Raimundez: known constants.

**Listing 173:**
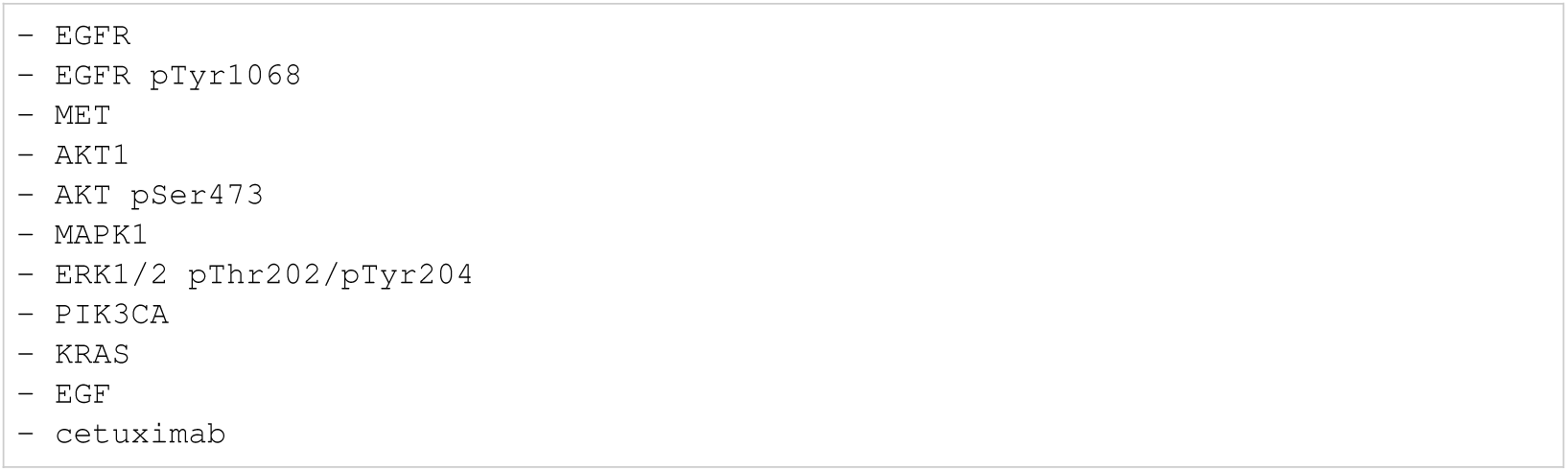
Raimundez: biocurator entities.

#### I.5.17 Schwen

**Listing 174:**
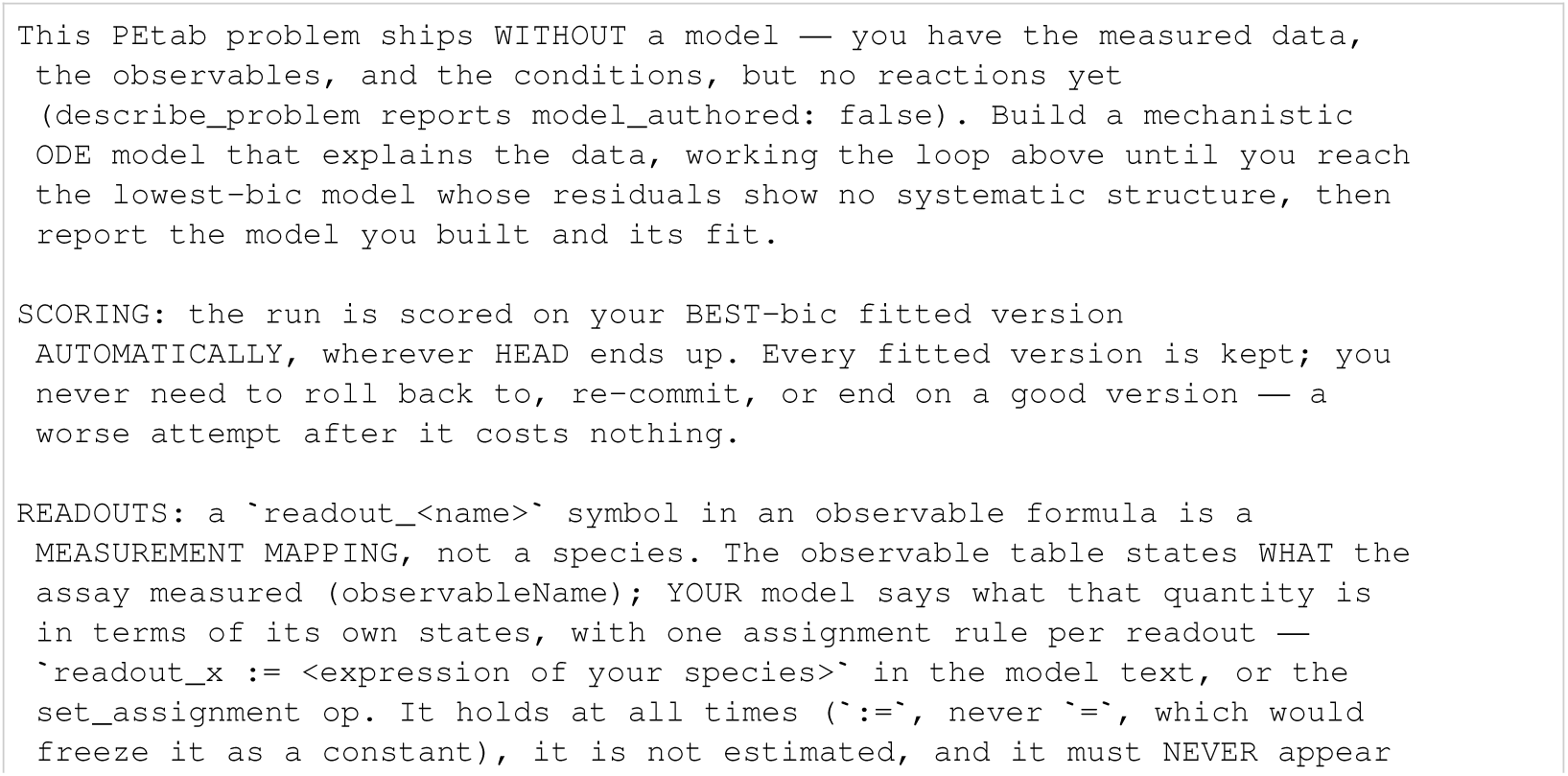
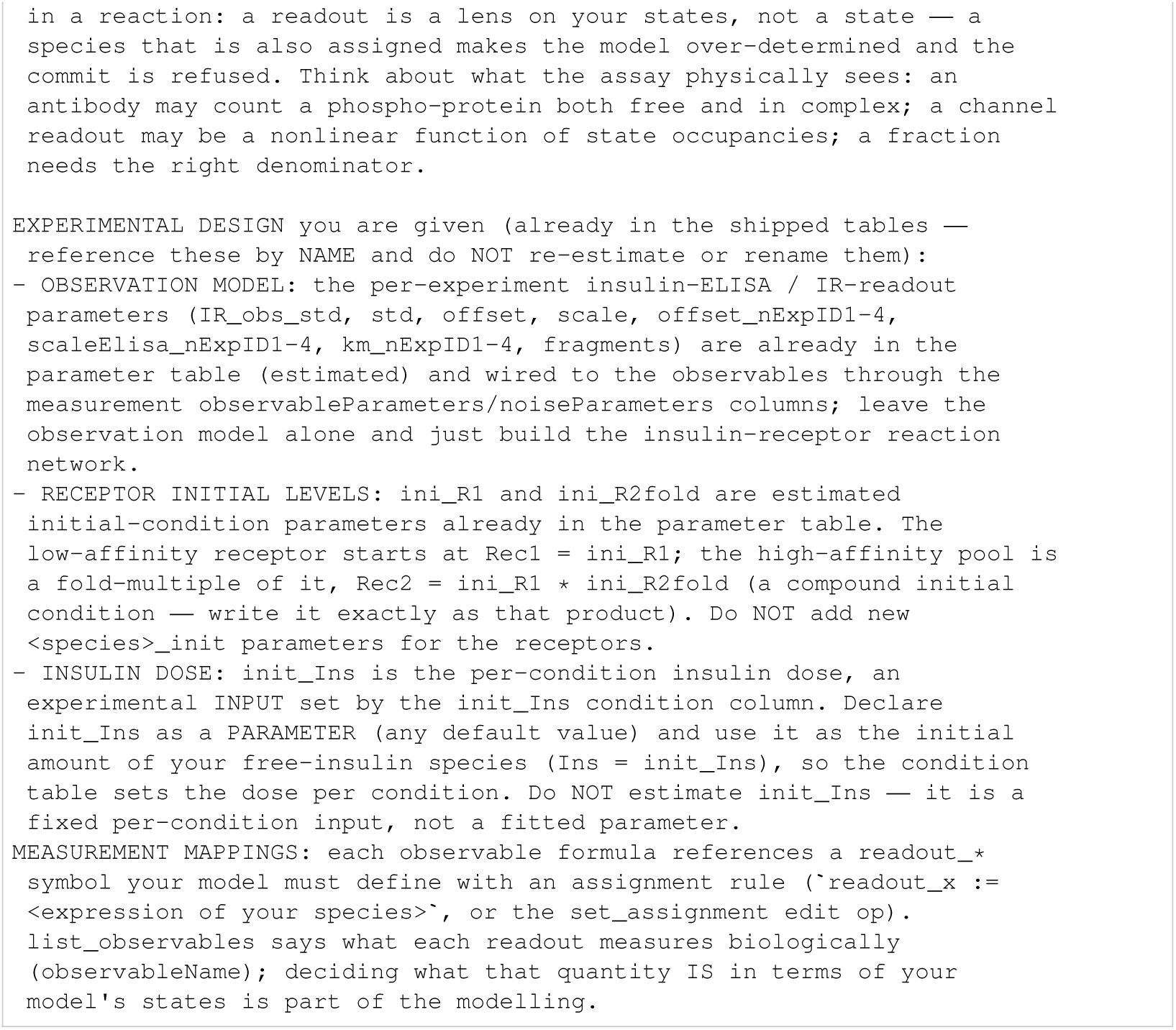
Schwen: initial task.

**Listing 175:**
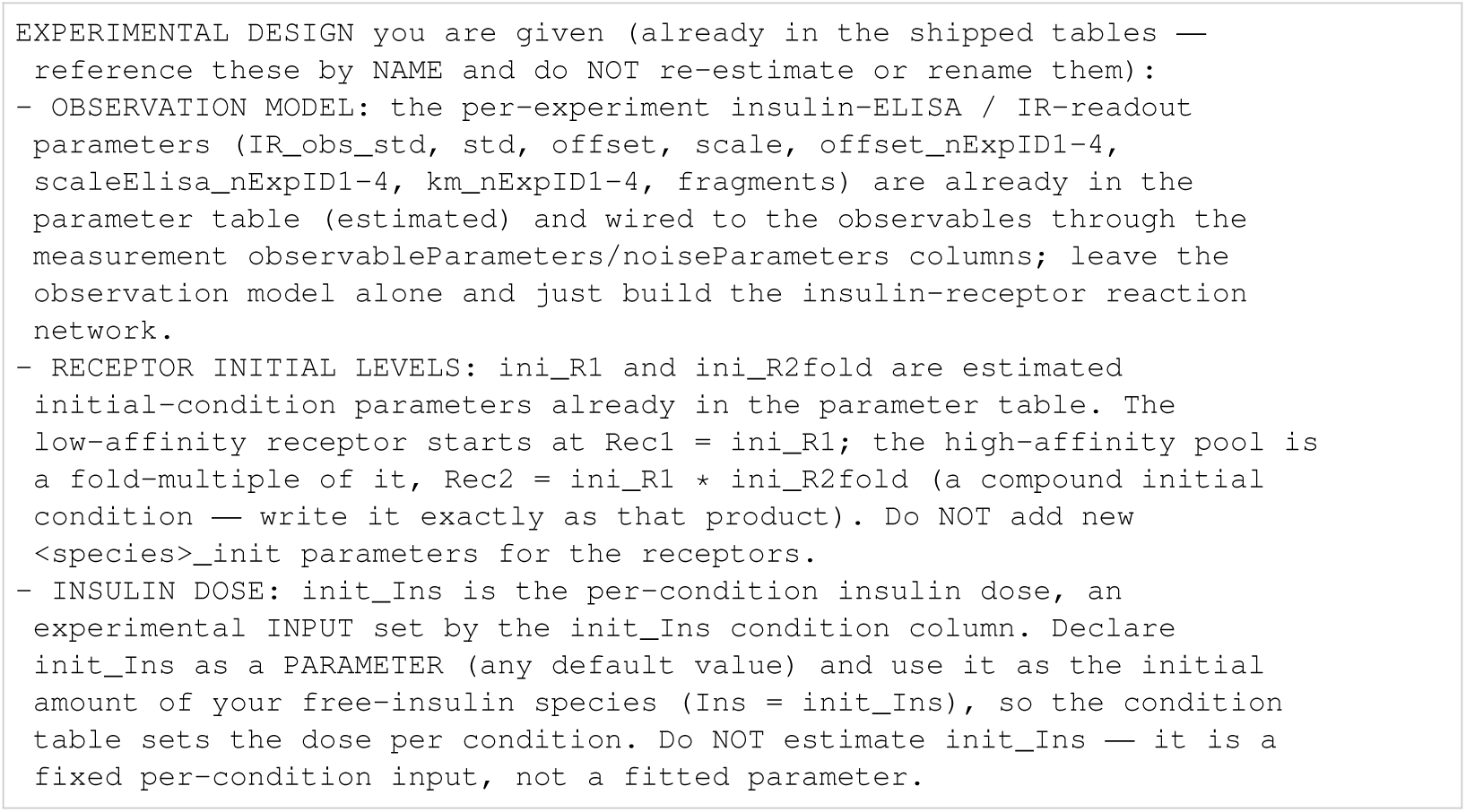
Schwen: known constants.

**Listing 176:**
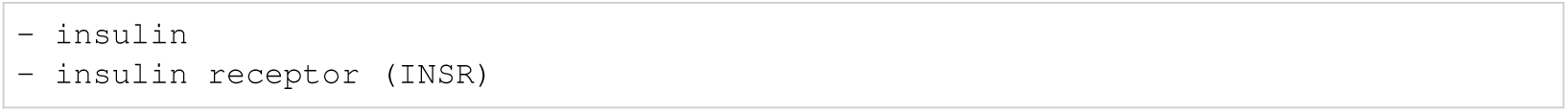
Schwen: biocurator entities.

#### I.5.18 Sneyd

**Listing 177:**
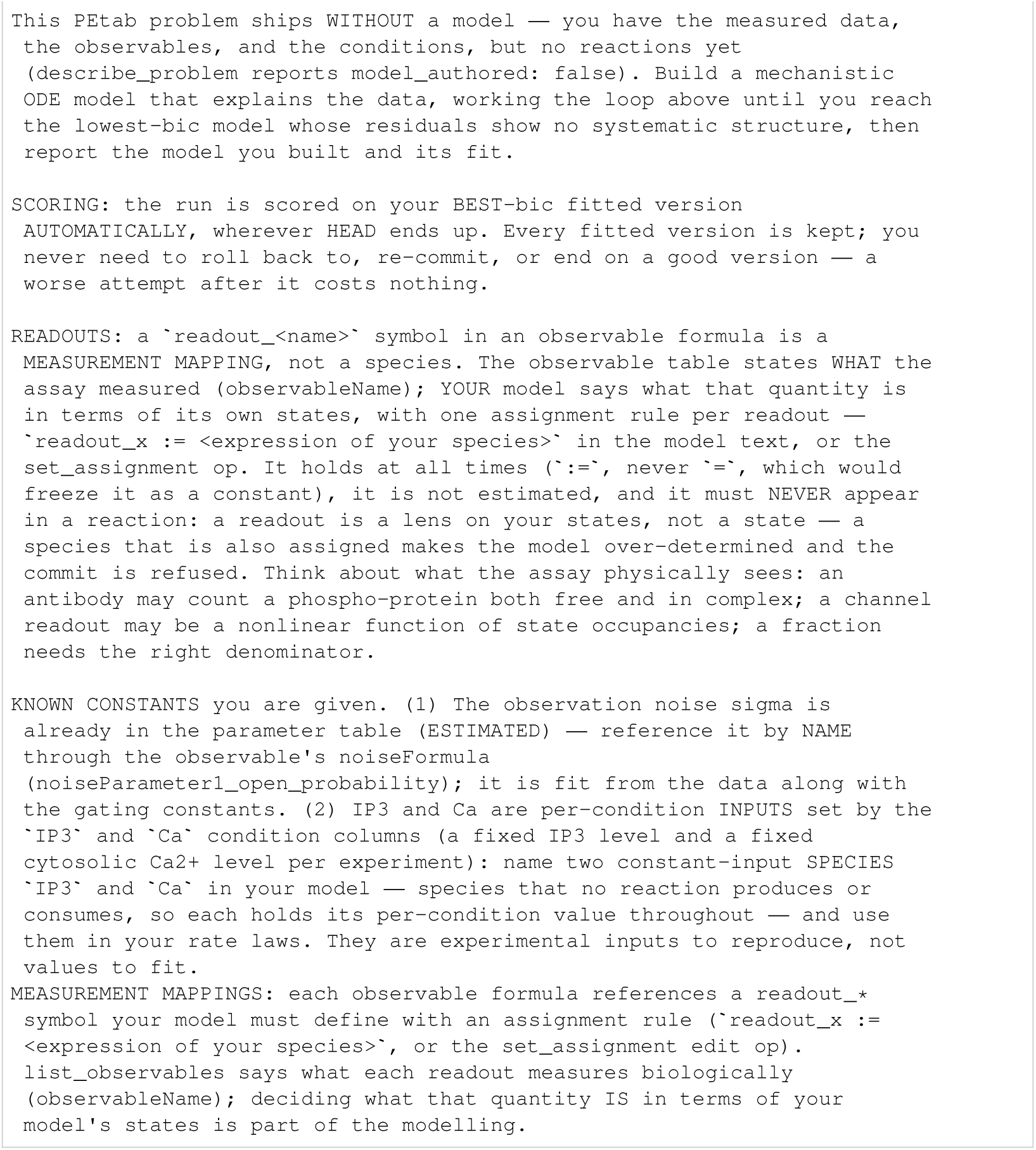
Sneyd: initial task.

**Listing 178:**
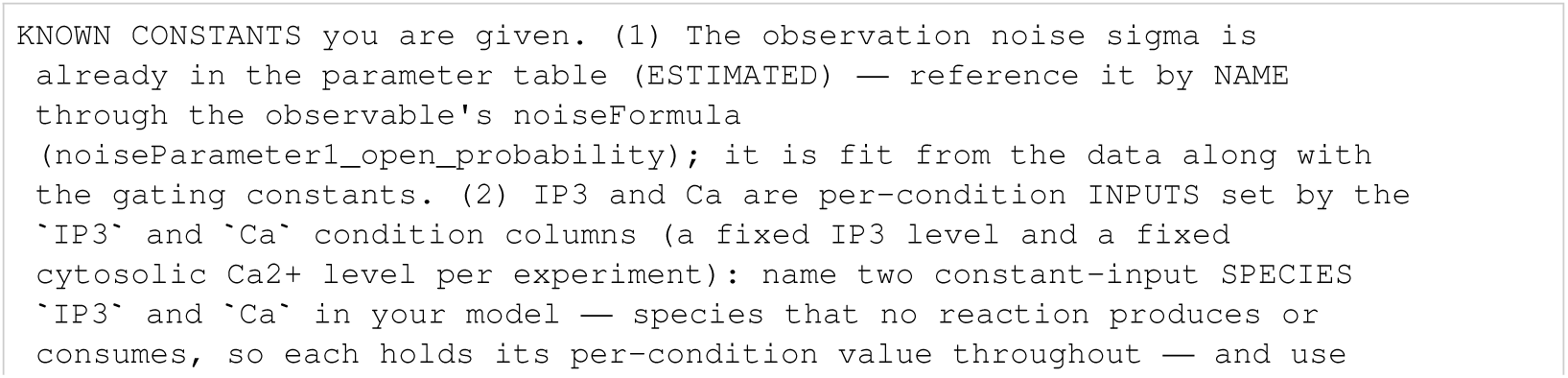
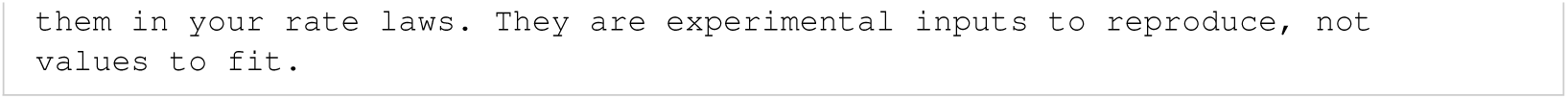
Sneyd: known constants.

**Listing 179:**
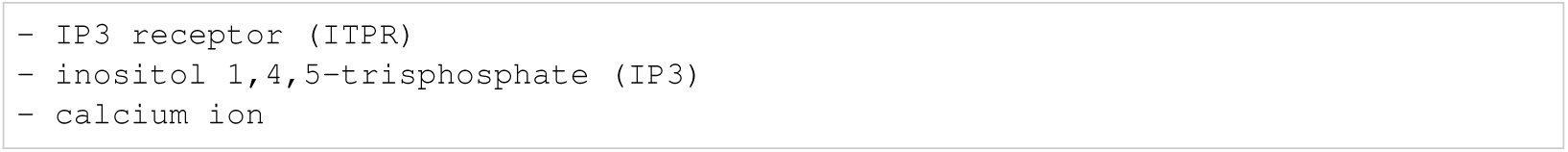
Sneyd: biocurator entities.

## References

Nikhil Abhyankar, Sha Li, Sanchit Kabra, Naren Ramakrishnan, Yulia Gel, and Chandan K. Reddy. LLM-ACES: Closed-loop discovery of dynamical systems with LLM-guided adaptive search. arXiv preprint arXiv:2606.25039, 2026. URL https://arxiv.org/abs/2606.25039.

Hirotugu Akaike. A new look at the statistical model identification. IEEE Transactions on Automatic Control, 19(6):716–723, 1974. doi: 10.1109/TAC.1974.1100705. URL 10.1109/TAC.1974.1100705.

Uri Alon. *An Introduction to Systems Biology: Design Principles of Biological Circuits*. Chapman and Hall/CRC, Boca Raton, 2nd edition, 2019. ISBN 9781439837177. doi: 10.1201/9780429283321. URL 10.1201/9780429283321.

Joy Armistead, Sebastian Höpfl, Pierre Goldhausen, Andrea Müller-Hartmann, Evelin Fahle, Julia Hatzold, Rainer Franzen, Susanne Brodesser, Nicole E. Radde, and Matthias Hammerschmidt. A sphingolipid rheostat controls apoptosis versus apical cell extrusion as alternative tumoursuppressive mechanisms. Cell Death & Disease, 15:746, 2024. doi: 10.1038/s41419-024-07134-2. URL 10.1038/s41419-024-07134-2.

John A. Bachman, Benjamin M. Gyori, and Peter K. Sorger. Automated assembly of molecular mechanisms at scale from text mining and curated databases. Molecular Systems Biology, 19(5): e11325, 2023. doi: 10.15252/msb.202211325. URL https://link.springer.com/article/10.15252/msb.202211325.

Vitaly Balan, Deborah T. Leicht, Jun Zhu, Karina Balan, Alexander Kaplun, Vinita Singh-Gupta, Jun Qin, Hong Ruan, Michael J. Comb, and Guri Tzivion. Identification of novel in vivo Raf-1 phosphorylation sites mediating positive feedback Raf-1 regulation by extracellular signal-regulated kinase. Molecular Biology of the Cell, 17(3):1141–1153, 2006. doi: 10.1091/mbc.E04-12-1123. URL 10.1091/mbc.E04-12-1123.

Hamparsum Bozdogan. Model selection and Akaike’s information criterion (AIC): The general theory and its analytical extensions. Psychometrika, 52(3):345–370, 1987. doi: 10.1007/BF0229 4361. URL 10.1007/BF02294361.

Cecilia Brännmark, Robert Palmér, S. Torkel Glad, Gunnar Cedersund, and Peter Strålfors. Mass and information feedbacks through receptor endocytosis govern insulin signaling as revealed using a parameter-free modeling framework. Journal of Biological Chemistry, 285(26):20171–20179, 2010. doi: 10.1074/jbc.M110.106849. URL 10.1074/jbc.M110.106849.

James Briscoe and Anna Kicheva. The physics of development 100 years after D’Arcy Thompson’s “On Growth and Form”. Mechanisms of Development, 145:26–31, 2017. doi: 10.1016/j.mod.2017.03.005. URL 10.1016/j.mod.2017.03.005.

Eric J. Brown, Peter A. Beal, Curtis T. Keith, Jie Chen, Tae Bum Shin, and Stuart L. Schreiber. Control of p70 S6 kinase by kinase activity of FRAP in vivo. Nature, 377(6548):441–446, 1995. doi: 10.1038/377441a0. URL 10.1038/377441a0.

Mattia Cerrato, Lennart Baur, Jannis Brugger, Sajjad Shumaly, Nicholas Schmitt, Edward Finkelstein, Selina Jukic, Lars Münzel, Felix Peter Paul, Pascal Pfannes, Benedikt Rohr, Julius Schellenberg, Philipp Wolf, and Stefan Kramer. Science-Gym: a simple testbed for AI-driven scientific discovery. Machine Learning, 115:16, 2026. doi: 10.1007/s10994-025-06914-x. URL 10.1007/s10994-025-06914-x.

Ricky T. Q. Chen, Yulia Rubanova, Jesse Bettencourt, and David Duvenaud. Neural ordinary differential equations. In Advances in Neural Information Processing Systems (NeurIPS), 2018. URL https://arxiv.org/abs/1806.07366.

Miles Cranmer. Interpretable machine learning for science with PySR and SymbolicRegression.jl. arXiv preprint arXiv:2305.01582, 2023. URL https://arxiv.org/abs/2305.01582.

Miles Cranmer, Alvaro Sanchez Gonzalez, Peter Battaglia, Rui Xu, Kyle Cranmer, David Spergel, and Shirley Ho. Discovering symbolic models from deep learning with inductive biases. In Advances in Neural Information Processing Systems, volume 33, pp. 17429–17442. Curran Associates, Inc., 2020. URL https://proceedings.neurips.cc/paper_files/paper/2020/file/c9f2f917078bd2db12f23c3b413d9cba-Paper.pdf.

Fabien Crauste, Julien Mafille, Lilia Boucinha, Sophia Djebali, Olivier Gandrillon, Jacqueline Marvel, and Christophe Arpin. Identification of nascent memory CD8 t cells and modeling of their ontogeny. Cell Systems, 4(3):306–317.e4, 2017. doi: 10.1016/j.cels.2017.01.014. URL 10.1016/j.cels.2017.01.014.

Francis Crick. *What Mad Pursuit: A Personal View of Scientific Discovery*. Basic Books, New York, 1988. ISBN 9780465091379. URL https://books.google.com/books?id=NBAhAQAAIAAJ.

Stéphane d’Ascoli, Sören Becker, Philippe Schwaller, Alexander Mathis, and Niki Kilbertus. ODE-Former: Symbolic regression of dynamical systems with transformers. In International Conference on Learning Representations (ICLR), 2024. URL https://proceedings.iclr.cc/paper_files/paper/2024/hash/5ed5c3c846f684a54975ad7a2525199f-Abstract-Conference.html.

Theodore de Pomereu and Fabian Fröhlich. Symbolic regression enables coarse-grained model discovery of intracellular signalling dynamics. bioRxiv, 2026. doi: 10.64898/2026.08.20.745973. URL 10.64898/2026.08.20.745973.

Emek Demir, Michael P. Cary, Suzanne Paley, Ken Fukuda, et al. The BioPAX community standard for pathway data sharing. Nature Biotechnology, 28(9):935–942, 2010. doi: 10.1038/nbt.1666. URL https://www.nature.com/articles/nbt.1666.

Daniel C. Dennett. Real patterns. The Journal of Philosophy, 88(1):27–51, 1991. doi: 10.2307/2027085. URL 10.2307/2027085.

James DiFrisco and Johannes Jaeger. Genetic causation in complex regulatory systems: An integrative dynamic perspective. BioEssays, 42(6):1900226, 2020. doi: 10.1002/bies.201900226. URL 10.1002/bies.201900226.

Michele K. Dougherty, Jürgen Müller, Daniel A. Ritt, Ming Zhou, Xiao Zhen Zhou, Terry D. Copeland, Thomas P. Conrads, Timothy D. Veenstra, Kun Ping Lu, and Deborah K. Morrison. Regulation of Raf-1 by direct feedback phosphorylation. Molecular Cell, 17(2):215–224, 2005. doi: 10.1016/j.molcel.2004.11.055. URL 10.1016/j.molcel.200 4.11.055.

Haonan Duan, Stephen Zhewen Lu, Caitlin Fiona Harrigan, Nishkrit Desai, Jiarui Lu, Michał Koziarski, Leonardo Cotta, and Chris J. Maddison. Measuring scientific capabilities of language models with a systems biology dry lab. In Advances in Neural Information Processing Systems, volume 38, 2025. doi: 10.52202/085713-3733. URL https://proceedings.neurips.cc/paper_files/paper/2025/hash/a23760ba51036d530a2656c3835f826c-Abstract-Datasets_and_Benchmarks_Track.html.

Anna Fiedler, Sebastian Raeth, Fabian J. Theis, Angelika Hausser, and Jan Hasenauer. Tailored parameter optimization methods for ordinary differential equation models with steady-state constraints. BMC Systems Biology, 10:80, 2016. doi: 10.1186/s12918-016-0319-7. URL 10.1186/s12918-016-0319-7.

Malcolm Forster and Elliott Sober. How to tell when simpler, more unified, or less ad hoc theories will provide more accurate predictions. The British Journal for the Philosophy of Science, 45(1): 1–35, 1994. doi: 10.1093/bjps/45.1.1. URL 10.1093/bjps/45.1.1.

Fabian Fröhlich and Peter K. Sorger. Fides: Reliable trust-region optimization for parameter estimation of ordinary differential equation models. PLoS Computational Biology, 18(7):e1010322, 2022. doi: 10.1371/journal.pcbi.1010322.

Fabian Fröhlich, Daniel Weindl, Yannik Schälte, Dilan Pathirana, Łukasz Paszkowski, Glenn Terje Lines, Paul Stapor, and Jan Hasenauer. AMICI: high-performance sensitivity analysis for large ordinary differential equation models. Bioinformatics, 37(20):3676–3677, 2021. doi: 10.1093/bioinformatics/btab227.

Kazuhiro A. Fujita, Yu Toyoshima, Shinsuke Uda, Yu-ichi Ozaki, Hiroyuki Kubota, and Shinya Kuroda. Decoupling of receptor and downstream signals in the Akt pathway by its low-pass filter characteristics. Science Signaling, 3(132):ra56, 2010. doi: 10.1126/scisignal.2000810. URL 10.1126/scisignal.2000810.

Kanishk Gandhi, Michael Y. Li, Lyle Goodyear, Agam Bhatia, Louise Li, Aditi Bhaskar, Mohammed Zaman, and Noah D. Goodman. BoxingGym: Benchmarking progress in automated experimental design and model discovery. arXiv preprint *arXiv:2501.01540*, 2025. doi: 10.48550/arXiv.2501.01540. URL https://arxiv.org/abs/2501.01540.

Stephan Grein, David R. Penas, Daniel Weindl, Polina Lakrisenko, Julio R. Banga, and Jan Hasenauer. Large-scale analysis of optimisation methods for parameter estimation problems in the life sciences. bioRxiv, 2026. doi: 10.64898/2026.07.11.737731.

Benjamin M. Gyori, John A. Bachman, Kartik Subramanian, Jeremy L. Muhlich, Lucian Galescu, and Peter K. Sorger. From word models to executable models of signaling networks using automated assembly. Molecular Systems Biology, 13(11):954, 2017. doi: 10.15252/msb.20177651. URL https://link.springer.com/article/10.15252/msb.20177651.

E. J. Hannan and B. G. Quinn. The determination of the order of an autoregression. Journal of the Royal Statistical Society: Series B (Methodological), 41(2):190–195, 1979. doi: 10.1111/j.2517-6161.1979.tb01072.x. URL 10.1111/j.2517-6161.1979.tb01072.x.

Leonard A. Harris, Justin S. Hogg, José-Juan Tapia, John A. P. Sekar, Sanjana A. Gupta, Ilya Korsunsky, Arshi Arora, Dipak Barua, Robert P. Sheehan, and James R. Faeder. BioNetGen 2.2: advances in rule-based modeling. Bioinformatics, 32(21):3366–3368, 2016. doi: 10.1093/bioinformatics/btw469. URL 10.1093/bioinformatics/btw469.

Helge Hass, Carolin Loos, Elba Raimúndez-Álvarez, Jens Timmer, Jan Hasenauer, and Clemens Kreutz. Benchmark problems for dynamic modeling of intracellular processes. Bioinformatics, 35(17):3073–3082, 2019. doi: 10.1093/bioinformatics/btz020. URL 10.1093/bioinformatics/btz020.

Tom W. Hiscock and Sean G. Megason. Mathematically guided approaches to distinguish models of periodic patterning. Development, 142(3):409–419, 2015. doi: 10.1242/dev.107441. URL 10.1242/dev.107441.

Samuel Holt, Zhaozhi Qian, Tennison Liu, James Weatherall, and Mihaela van der Schaar. Datadriven discovery of dynamical systems in pharmacology using large language models. In Advances in Neural Information Processing Systems, volume 37, 2024. URL https://proceedings.neurips.cc/paper_files/paper/2024/hash/aea8bdc42d8ba3a67a69b3f18be93f69-Abstract-Conference.html.

Michael Hucka, Andrew Finney, Herbert M. Sauro, Hamid Bolouri, John C. Doyle, Hiroaki Kitano, et al. The systems biology markup language (SBML): a medium for representation and exchange of biochemical network models. Bioinformatics, 19(4):524–531, 2003. doi: 10.1093/bioinformatics/btg015. URL 10.1093/bioinformatics/btg015.

Clifford M. Hurvich and Chih-Ling Tsai. Regression and time series model selection in small samples. Biometrika, 76(2):297–307, 1989. doi: 10.1093/biomet/76.2.297. URL 10.1093/biomet/76.2.297.

Ken Inoki, Yong Li, Tianquan Zhu, Jun Wu, and Kun-Liang Guan. TSC2 is phosphorylated and inhibited by Akt and suppresses mTOR signalling. Nature Cell Biology, 4(9):648–657, 2002. doi: 10.1038/ncb839. URL 10.1038/ncb839.

Sanchit Kabra, Nikhil Abhyankar, Saaketh Desai, Prasad Iyer, and Chandan K. Reddy. LLM- AutoSciLab: Closed-loop scientific discovery via active experimentation with LLMs. arXiv preprint *arXiv:2605.24043*, 2026. doi: 10.48550/arXiv.2605.24043. URL https://arxiv.org/abs/2605.24043.

Krzysztof Kacprzyk and Mihaela van der Schaar. No equations needed: Learning system dynamics without relying on closed-form ODEs. In The Thirteenth International Conference on Learning Representations, 2025. URL https://openreview.net/forum?id=kbm6tsICar.

Krzysztof Kacprzyk, Julianna Piskorz, and Mihaela van der Schaar. Skip the equations: Learning behavior of personalized dynamical systems directly from data. In Proceedings of the 42nd International Conference on Machine Learning, volume 267 of Proceedings of Machine Learning Research, pp. 28619–28643. PMLR, 2025. URL https://proceedings.mlr.press/v267/kacprzyk25a.html.

Robert E. Kass and Adrian E. Raftery. Bayes factors. Journal of the American Statistical Association, 90(430):773–795, 1995. doi: 10.1080/01621459.1995.10476572. URL 10.1080/01621459.1995.10476572.

Matt J. Keeling and Pejman Rohani. Modeling Infectious Diseases in Humans and Animals. Princeton University Press, 2008. doi: 10.1515/9781400841035. URL 10.1 515/9781400841035.

Edda Klipp, Wolfram Liebermeister, Christoph Wierling, and Axel Kowald. Systems Biology: A Textbook. Wiley-VCH, Weinheim, 2nd edition, 2016. ISBN 9783527336364. URL https://www.wiley-vch.de/en/areas-interest/natural-sciences/life-sciences-11ls/bioinformatics-computational-biology-11lsg/systems-biology-978-3-527-33636-4.

David Krongauz, Arad Zulti, Eran Segal, and Teddy Lazebnik. Automatic ordinary differential equations discovery for biological systems using large language model powered agentic system. arXiv preprint arXiv:2607.13608, 2026. URL https://arxiv.org/abs/2607.13608.

James Ladyman. Scientific empiricism and modal structure. Synthese, 206:245, 2025. doi: 10.1007/s11229-025-05323-w. URL https://link.springer.com/article/10.1007/s11229-025-05323-w.

James Ladyman. Patterns all the way up: Prolegomena to a future naturalized metaphysics. In Tyler Millhouse, Stephen D. Petersen, and Don Ross (eds.), Dennett’s Real Patterns in Science and Nature, pp. 59–76. MIT Press, 2026. doi: 10.7551/mitpress/15550.003.0006. URL 10.7551/mitpress/15550.003.0006.

LangChain, Inc. Deep Agents. Open-source software, 2025. URL https://github.com/langchain-ai/deepagents. Accessed 22 September 2026.

Carolin Loos, Sabrina Krause, and Jan Hasenauer. Hierarchical optimization for the efficient parametrization of ODE models. Bioinformatics, 34(24):4266–4273, 2018. doi: 10.1093/bioinformatics/bty514. URL 10.1093/bioinformatics/bty514.

Carlos F. Lopez, Jeremy L. Muhlich, John A. Bachman, and Peter K. Sorger. Programming biological models in Python using PySB. Molecular Systems Biology, 9:646, 2013. doi: 10.1038/msb.2013.1. URL 10.1038/msb.2013.1.

Philippe Lucarelli, Marcel Schilling, Clemens Kreutz, Artyom Vlasov, Martin E. Boehm, Nao Iwamoto, Bernhard Steiert, Susen Lattermann, Marvin Wäsch, Markus Stepath, Matthias S. Matter, Mathias Heikenwälder, Katrin Hoffmann, Daniela Deharde, Georg Damm, Daniel Seehofer, Maria Muciek, Norbert Gretz, Wolf D. Lehmann, Jens Timmer, and Ursula Klingmüller. Resolving the combinatorial complexity of Smad protein complex formation and its link to gene expression. Cell Systems, 6(1):75–89.e11, 2018. doi: 10.1016/j.cels.2017.11.010. URL 10.1016/j.cels.2017.11.010.

Don-On Daniel Mak, John E. Pearson, King Pan Campion Loong, Suman Datta, Marisabel Fernández-Mongil, and J. Kevin Foskett. Rapid ligand-regulated gating kinetics of single inositol 1,4,5-trisphosphate receptor Ca2+ release channels. EMBO Reports, 8(11):1044–1051, 2007. doi: 10.1038/sj.embor.7401087. URL 10.1038/sj.embor.7401087.

Niall M. Mangan, Steven L. Brunton, Joshua L. Proctor, and J. Nathan Kutz. Inferring biological networks by sparse identification of nonlinear dynamics. *IEEE Transactions on Molecular*, Biological and Multi-Scale Communications, 2(1):52–63, 2016. doi: 10.1109/TMBMC.2016.2633265. URL 10.1109/TMBMC.2016.2633265.

Huaiyu Mi, Falk Schreiber, Stuart Moodie, Tobias Czauderna, Emek Demir, Robin Haw, Augustin Luna, Nicolas Le Novère, Anatoly Sorokin, and Alice Villéger. Systems biology graphical notation: Activity flow language level 1 version 1.2. Journal of Integrative Bioinformatics, 12(2):265, 2015. doi: 10.2390/biecoll-jib-2015-265. URL 10.2390/biecoll-jib-2015-265.

Marija Milacic, Deidre Beavers, Patrick Conley, et al. The Reactome pathway knowledgebase 2024. Nucleic Acids Research, 52(D1):D672–D678, 2024. doi: 10.1093/nar/gkad1025. URL https://academic.oup.com/nar/article/52/D1/D672/7369850.

Kevin Murphy. Model discovery agent: LLM-assisted Bayesian experiment design for data-efficient discovery of mechanistic world models. arXiv preprint arXiv:2608.09696, 2026. URL https://arxiv.org/abs/2608.09696v4.

Dilan Pathirana, Fabian Fröhlich, Sebastian Persson, Frank T. Bergmann, Matthias König, Paul F. Lang, Severin Bang, Svenja Kemmer, Jan Hasenauer, and Daniel Weindl. The parameter estimation tables (PEtab) format version 2. Journal of Integrative Bioinformatics, 2026. doi: 10.1515/jib-2026-0004. URL 10.1515/jib-2026-0004.

Judea Pearl. Causality: Models, Reasoning, and Inference. Cambridge University Press, 2nd edition, 2009. doi: 10.1017/CBO9780511803161. URL 10.1017/CBO9780511803161.

Alan S. Perelson, Avidan U. Neumann, Martin Markowitz, John M. Leonard, and David D. Ho. HIV-1 dynamics in vivo: virion clearance rate, infected cell life-span, and viral generation time. Science, 271(5255):1582–1586, 1996. doi: 10.1126/science.271.5255.1582. URL 10.1126/science.271.5255.1582.

Sebastian Persson, Branwen Snelling, Maren Philipps, Daniel Weindl, Marija Cvijovic, Jan Hasenauer, Dilan Pathirana, and Fabian Fröhlich. PEtab SciML: an exchange format for specifying and training dynamic scientific machine learning models. arXiv preprint arXiv:2608.20184, 2026. URL https://arxiv.org/abs/2608.20184.

Ingmar Posner, Anson Lei, and Bernhard Schölkopf. From observation to insight: Mechanistic world models and the quest for autonomous discovery. arXiv preprint *arXiv:2607.12474*, 2026. doi: 10.48550/arXiv.2607.12474. URL https://arxiv.org/abs/2607.12474v2.

Mark Raasveldt and Hannes Mühleisen. DuckDB: an embeddable analytical database. In Proceedings of the 2019 International Conference on Management of Data, SIGMOD ’19, pp. 1981– 1984. Association for Computing Machinery, 2019. doi: 10.1145/3299869.3320212.

Elba Raimúndez, Simone Keller, Gwen Zwingenberger, Karolin Ebert, Sabine Hug, Fabian J. Theis, Dieter Maier, Birgit Luber, and Jan Hasenauer. Model-based analysis of response and resistance factors of cetuximab treatment in gastric cancer cell lines. PLoS Computational Biology, 16(3): e1007147, 2020. doi: 10.1371/journal.pcbi.1007147. URL https://journals.plos.org/ploscompbiol/article?id=10.1371/journal.pcbi.1007147.

Ana M. Rossi, Andrew M. Riley, Stephen C. Tovey, Taufiq Rahman, Olivier Dellis, Emily J. A. Taylor, Valery G. Veresov, Barry V. L. Potter, and Colin W. Taylor. Synthetic partial agonists reveal key steps in IP3 receptor activation. Nature Chemical Biology, 5(9):631–639, 2009. doi: 10.1038/nchembio.195. URL 10.1038/nchembio.195.

Surojit Sarkar, Vandana Kalia, W. Nicholas Haining, Bogumila T. Konieczny, Shruti Subramaniam, and Rafi Ahmed. Functional and genomic profiling of effector CD8 T cell subsets with distinct memory fates. Journal of Experimental Medicine, 205(3):625–640, 2008. doi: 10.1084/jem.20 071641. URL 10.1084/jem.20071641.

Yannik Schälte, Fabian Fröhlich, Paul J. Jost, Jakob Vanhoefer, Dilan Pathirana, Paul Stapor, Polina Lakrisenko, Dantong Wang, Elba Raimúndez, Simon Merkt, Leonard Schmiester, Philipp Städter, Stephan Grein, Erika Dudkin, Domagoj Doresic, Daniel Weindl, and Jan Hasenauer. pyPESTO: a modular and scalable tool for parameter estimation for dynamic models. Bioinformatics, 39(11): btad711, 2023. doi: 10.1093/bioinformatics/btad711.

Leonard Schmiester, Yannik Schälte, Fabian Fröhlich, Jan Hasenauer, and Daniel Weindl. Efficient parameterization of large-scale dynamic models based on relative measurements. Bioinformatics, 36(2):594–602, 2020. doi: 10.1093/bioinformatics/btz581. URL 10.1093/bioinformatics/btz581.

Leonard Schmiester, Yannik Schälte, Frank T. Bergmann, Tacio Camba, Erika Dudkin, Janine Egert, Fabian Fröhlich, Lara Fuhrmann, Adrian L. Hauber, Svenja Kemmer, Polina Lakrisenko, Carolin Loos, Simon Merkt, Wolfgang Müller, Dilan Pathirana, Elba Raimúndez, Lukas Refisch, Marcus Rosenblatt, Paul L. Stapor, Philipp Städter, Dantong Wang, Franz-Georg Wieland, Julio R. Banga, Jens Timmer, Alejandro F. Villaverde, Sven Sahle, Clemens Kreutz, Jan Hasenauer, and Daniel Weindl. PEtab—interoperable specification of parameter estimation problems in systems biology. PLoS Computational Biology, 17(1):e1008646, 2021. doi: 10.1371/journal.pcbi.1008646. URL https://journals.plos.org/ploscompbiol/article?id=10.1371/journal.pcbi.1008646.

Gideon Schwarz. Estimating the dimension of a model. The Annals of Statistics, 6(2):461–464, 1978. doi: 10.1214/aos/1176344136. URL 10.1214/aos/1176344136.

Parshin Shojaee, Kazem Meidani, Shashank Gupta, Amir Barati Farimani, and Chandan Reddy. LLM-SR: Scientific equation discovery via programming with large language models. In International Conference on Learning Representations, pp. 16054–16085, 2025a. URL https://proceedings.iclr.cc/paper_files/paper/2025/hash/28df8e730c054c 5331855fd4d5403ba9-Abstract-Conference.html.

Parshin Shojaee, Ngoc-Hieu Nguyen, Kazem Meidani, Amir Barati Farimani, Khoa D Doan, and Chandan K. Reddy. LLM-SRBench: A new benchmark for scientific equation discovery with large language models. In Proceedings of the 42nd International Conference on Machine Learning, volume 267 of Proceedings of Machine Learning Research, pp. 55325–55359. PMLR, 2025b. URL https://proceedings.mlr.press/v267/shojaee25a.html.

Lucian P. Smith, Frank T. Bergmann, Deepak Chandran, and Herbert M. Sauro. Antimony: a modular model definition language. Bioinformatics, 25(18):2452–2454, 2009. doi: 10.1093/bioinformatics/btp401. URL 10.1093/bioinformatics/btp401.

Lucian P. Smith, Michael Hucka, Stefan Hoops, Andrew Finney, Martin Ginkel, Chris J. Myers, Ion Moraru, and Wolfram Liebermeister. SBML level 3 package: Hierarchical model composition, version 1 release 3. Journal of Integrative Bioinformatics, 12(2):268, 2015. doi: 10.2390/biecoll-jib-2015-268. URL 10.2390/biecoll-jib-2015-268.

James Sneyd and Jean-François Dufour. A dynamic model of the type-2 inositol trisphosphate receptor. Proceedings of the National Academy of Sciences, 99(4):2398–2403, 2002. doi: 10.1073/pnas.032281999. URL 10.1073/pnas.032281999.

Sum Kyun Song, Bong Gyun Shin, and Jae Yong Lee. Discovering ordinary differential equations with LLM-based qualitative and quantitative evaluation. In International Conference on Machine Learning (ICML), 2026. URL https://arxiv.org/abs/2605.07323.

Gábor Szederkényi, Julio R. Banga, and Antonio A. Alonso. Inference of complex biological networks: distinguishability issues and optimization-based solutions. BMC Systems Biology, 5:177, 2011. doi: 10.1186/1752-0509-5-177. URL 10.1186/1752-0509-5-177.

Kei Takeuchi. Distribution of information statistics and criteria for adequacy of models. Suri Kagaku, 14(3):12–18, 1976. URL https://ndlsearch.ndl.go.jp/en/books/R000000004-I1701125. In Japanese.

Beth A. Tamburini, Matthew A. Burchill, and Ross M. Kedl. Antigen capture and archiving by lymphatic endothelial cells following vaccination or viral infection. Nature Communications, 5: 3989, 2014. doi: 10.1038/ncomms4989. URL 10.1038/ncomms4989.

The UniProt Consortium. UniProt: the universal protein knowledgebase in 2025. Nucleic Acids Research, 53(D1):D609–D617, 2025. doi: 10.1093/nar/gkae1010. URL https://academic.oup.com/nar/article/53/D1/D609/7902999.

Claire J. Tomlin and Jeffrey D. Axelrod. Biology by numbers: mathematical modelling in developmental biology. Nature Reviews Genetics, 8(5):331–340, 2007. doi: 10.1038/nrg2098. URL 10.1038/nrg2098.

Dénes Türei, Jonathan Schaul, Nicolàs Palacio-Escat, et al. OmniPath: integrated knowledgebase for multi-omics analysis. Nucleic Acids Research, 54(D1):D652–D660, 2026. doi: 10.1093/nar/gkaf 1126. URL https://academic.oup.com/nar/article/54/D1/D652/8326458.

Aki Vehtari, Andrew Gelman, and Jonah Gabry. Practical Bayesian model evaluation using leaveone-out cross-validation and WAIC. Statistics and Computing, 27(5):1413–1432, 2017. doi: 10.1007/s11222-016-9696-4. URL 10.1007/s11222-016-9696-4.

Sumio Watanabe. Asymptotic equivalence of Bayes cross validation and widely applicable information criterion in singular learning theory. Journal of Machine Learning Research, 11:3571–3594, 2010. URL https://www.jmlr.org/papers/v11/watanabe10a.html.

Matt L. Wiemann, Lindsay M. Smith, Peter Melchior, Siddharth Mishra-Sharma, Andrew Gordon Wilson, Pavel Izmailov, and Carolina Cuesta-Lázaro. DiscoverPhysics: Benchmarking LLMs for out-of-the-box scientific thinking. arXiv preprint arXiv*:2605.26087*, 2026. doi: 10.48550/arXiv.2605.26087. URL https://arxiv.org/abs/2605.26087.

James Woodward. Making Things Happen: A Theory of Causal Explanation. Oxford University Press, 2003. doi: 10.1093/0195155270.001.0001. URL https://academic.oup.com/book/4324.

Tianshi Zheng, Kelvin Kiu-Wai Tam, Newt Hue-Nam K. Nguyen, Baixuan Xu, Zhaowei Wang, Jiayang Cheng, Hong Ting Tsang, Weiqi Wang, Jiaxin Bai, Tianqing Fang, Yangqiu Song, Ginny Y. Wong, and Simon See. NewtonBench: Benchmarking generalizable scientific law discovery in LLM agents. In International Conference on Learning Representations (ICLR), 2026. URL https://openreview.net/forum?id=Gk6umqW74m.

Amirmohammad Ziaei Bideh and Jonathan Gryak. LLM-ODE: Data-driven discovery of dynamical systems with large language models. arXiv preprint arXiv:2603.20910, 2026. URL https://arxiv.org/abs/2603.20910.

Amirmohammad Ziaei Bideh, Aleksandra Georgievska, and Jonathan Gryak. MDBench: Benchmarking data-driven methods for model discovery. Proceedings of the AAAI Conference on Artificial Intelligence, 40(24):19746–19754, 2026. doi: 10.1609/aaai.v40i24.39056. URL https://ojs.aaai.org/index.php/AAAI/article/view/39056.

